# On the structure of multiple stable equilibria in competitive ecological systems

**DOI:** 10.1101/2024.11.11.623080

**Authors:** Washington Taylor, James O’Dwyer

## Abstract

For some ecological systems with a large pool of possible species, there can be multiple stable equilibria with different species composition. Natural or anthropogenic disruption can induce a shift between different such equilibria. While some work has been done on ecological systems with multiple equilibria, there is no general theory governing the distribution of equilibria or characterizing the basins of attraction of different equilibria. This article addresses these questions in a simple class of Lotka-Volterra models. We focus on competitive systems of species on a niche axis with multiple equilibria. We find that basins of attraction are generally larger for equilibria with greater biomass; in many cases, the basin of attraction size scales roughly exponentially with the net biomass of equilibria. This is illustrated in two ecologically relevant limits. In a continuous limit with species spaced arbitrarily closely on the niche axis, equilibria with different numbers of species provide new perspective on the notion of limiting similarity. In another limit, akin to a statistical mechanical model, the niche axis becomes infinite while the range of interactions remains fixed; in this limit we prove the exponential relation between basin size and biomass using the Markov chain central limit theorem.

## Introduction

While theoretical models for some ecological systems have a unique globally stable equilibrium, it is well known that there exist many competitive systems and models with multiple stable equilibria (a.k.a. “multiple stable states”), in which different subsets of species from an initial pool persist (Lotka, 1956; Petraitis, 2013). In such situations, the equilibrium reached by a system can depend upon details of the initial conditions and/or the sequence of steps in community assembly. This can be relevant in natural systems, for example, when an ecological system is substantially perturbed by addition of multiple invasive species and extinction of endemic species, or when a new environment is populated by a mixture of existing species from other environments. The possibility of multiple equilibria has recently been explored both in controlled laboratory settings (Estrela et al., 2022; Lopes et al., 2024), and in theoretical models of large numbers of species controlled by random matrix Lotka-Volterra dynamics (Ros et al., 2023). Related situations in which multiple species are present but different species dominate (“alternative stable states”; Beisner et al. (2003)) have been encountered in contexts such as forest/savanna landscapes at intermediate rainfall (Staver et al., 2011) and coral/microalgae dominance on coral reefs (Bruno et al., 2009). Multiple stable states have also been found in other theoretical frameworks besides Lotka-Volterra systems (Goyal et al., 2018; Marsland III et al., 2020). In this work we consider classes of competitive systems that naturally have multiple distinct equilibria with different species composition, and focus in particular on the multiplicity of equilibria and the relative sizes of the basins of attraction of different equilibria.

In this work we restrict attention to competitive models with symmetric inter-species interactions. For such models the existence of a Lyapunov function *H* simplifies theoretical analysis. Equilibria that are locally stable and stable to invasion correspond to local minima of the Lyapunov function. While the analysis in this work applies to symmetric interactions, where the effect of one species on another is equal to that effect it experiences from that other species, it is natural to expect (by continuity) that similar results may hold at least when interactions are only weakly non-symmetric.

Ecological systems where species can be thought of as living on a niche axis (Whittaker, 1972; Whittaker and Levin, 1975) provide a natural context in which to explore the possibility of multiple equilibria, and such systems are the primary focus of this work. These models can be thought of as arising from competition for resources, where the resource type spans a continuum, like seed size for granivores. The notion of “limiting similarity” (MacArthur and Levins, 1967; May and MacArthur, 1972) suggests that an equilibrium will be reached when species are separated by some characteristic minimal distance *d* on the niche axis. Thus, for example, on a niche axis of finite size it has been expected from prior work (Case, 1980; Gatto, 1982) that the number of species *s* reached in equilibrium will be uniquely fixed by the maximum number of species that can be packed at distances greater than *d*, i.e. *s* = l*L*/*d*J, where *L* is the size of the niche axis. In a well-defined set of discrete niche models, however, we find that the number of species in a stable equilibrium can vary across a finite range, with inter-species spacing ranging roughly from *d* to 2*d* where *d* is the minimum possible spacing; for species with discrete niches, local variations in inter-species spacing also contribute to the multiplicity of possible equilibria, which may not have uniform spacing along the niche axis, even when competition is symmetric.

In addition to the multiplicity and distribution of equilibria, we explore the relative sizes of the basins of attraction (also known as domains of attraction; we use the terms interchangeably here) of the different equilibria in models with multiple equilibria. While such questions have been considered in the dynamical systems literature (Genesio et al., 1985), there has been limited attention to this question in the context of ecology (see, however, e.g. Gilpin and Case (1976)). The basin/domain of attraction size can be understood geometrically in some simple models, or, for more general models, in terms of the probability with which different equilibria are reached in numerical simulations with an appropriate distribution on initial conditions. In the niche models that we consider, the size of the domain of attraction of a given equilibrium or class of equilibria is generally larger for equilibria with greater net biomass (summed across all species present in the equilibrium solution), and in many cases the domain size is roughly proportional to the exponential of the biomass in the given equilibrium. We examine this relationship both for finite size systems and in two ecologically natural limits that admit some analytic treatment. Note that for more general systems, the role of biomass here may be played by some more general measure of weighted population size.

The structure of this paper is as follows: We begin by describing several classes of models with multiple equilibria: (1) a simple class of toy “monodominant” models, where the different equilibria each contain a single species, (2) the niche axis models that are the central focus of the paper, and (3) random matrix models that have been studied recently in the literature. We then summarize the various analysis techniques used in the paper, and then present results for each of the model types on the set of equilibria and relationship between domain sizes and biomass. We conclude with some comments on how these results might be extended to other types of systems, and other types of interactions. An Appendix contains some more technical points and more detailed analysis of the various models.

## Methods

### Models

#### Competitive Lotka-Volterra dynamics

We consider ecological systems described by the Lotka-Volterra (LV) model, where populations *n*_*i*_ of *S* species (*i* = 1, …, *S*) obey the dynamical equations

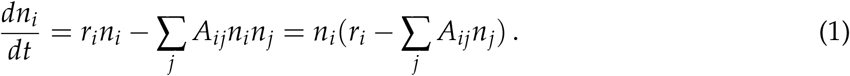

This simple class of equations is well-studied in theoretical ecology (Volterra, 1931; MacArthur and Levins, 1967; May and MacArthur, 1972; Allesina and Tang, 2012). While higher-order terms may be crucial in understanding detailed features of many ecosystems (see, e.g., Levine et al. (2017) for recent discussions), the LV approach may be sufficient for understanding many qualitative or statistical aspects of ecosystem structure and composition (see, e.g., Hu et al. (2022)).

For simplicity, in this paper we focus on symmetric interaction matrices *A* = *A*^*T*^, with purely competitive interactions *A*_*ij*_ ≥ 0, and set all growth rates to *r*_*i*_ = 1. This assumption of symmetry for competitive interactions follows in some cases from the assumptions in models of resource competition (MacArthur, 1970; Chesson, 1990), though it is fair to say that empirical support is at best mixed (Freckleton and Watkinson, 2001). We discuss at various points in the paper and in the Conclusions how the results may be generalized beyond these assumptions.

With these assumptions, as is well known, an equilibrium solution of Eq. 1 is given^1^ by (May and MacArthur, 1972)

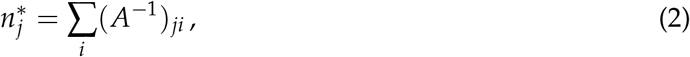

and the system has a Lyapunov function (MacArthur, 1970; Case, 1980; Gatto, 1990)

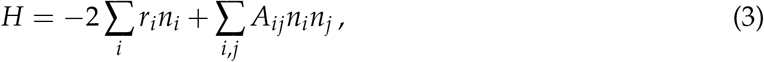

which satisfies *dH*(*t*)/*dt* ≤ 0 and has a value of the negative of the total biomass *B*(*n*^*^) ^2^ in any equilibrium solution (2):

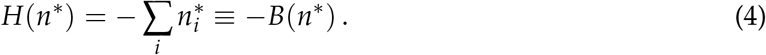

When the interactions between species are weak, there is a stable global equilibrium with 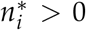 for all species *i* ∈ 𝒮 = {1, …, *S*}. For example, when interspecific interactions vanish, this reduces to each species sitting at its carrying capacity. We focus here, however, on systems with an unstable equilibrium at 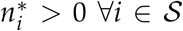 Such instability is generally associated with a negative eigenvalue of the matrix *A* (May, 2019). ^3^

Such a system may have one or more subsets of species 𝒮t ⊂ 𝒮 with stable equilibria 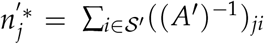, where *A*^t^ is the restriction of the interaction matrix to the subsystem 𝒮 ^′^ and has a strictly positive spectrum of eigenvalues. Note that we consider one of these ‘boundary’ equilibria stable only if it is both stable within the subsystem S^′^and also stable to invasion from other species in 𝒮.

#### Monodominant models

As a particularly simple class of models with multiple stable equilibria, we consider LV models with strong competitive interactions *A*_*ij*_ = 2, *i* ≠ *j*, and *A*_*ii*_ = *a*_*i*_ *<* 2 (Lotka, 1956). When all *a*_*i*_ = 1, this model has an unstable equilibrium with equal populations 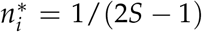, and *S* stable equilibria where a single 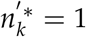 and the remaining 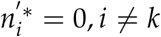. The behavior is similar when all *a*_*i*_ ∼ 1. This represents a class of models with strong competitive interactions between a similar set of species, where each species competes more strongly with the others than with itself (for example in congeneric territorial competition (Graves and Gotelli, 1993)), and provides a simple case study for analyzing basins of attraction

#### Niche axis models

A central class of models that we analyze here is based on the idea of a set of species that are specialized to different locations on a single niche axis, with competitive interactions that are stronger for species more closely positioned on the niche axis. Many ecological communities consist of multiple species competing for resources, and in some cases these resources fall on a continuous spectrum corresponding to resource type. One such example is seeds characterized by their size. Animals may have different efficiencies for feeding on seeds within specific size ranges, and the species niche reflects its resource preferences within this overall spectrum. Similarity in niche values therefore captures the amount of overlap between feeding strategies in this example (MacArthur and Levins, 1967; Leimar et al., 2013; D’Andrea et al., 2018).

For these models, we take *i* as an integer index in the niche axis, and assume that interactions have the form *A*_*ij*_ = *f* (*i* − *j*), where *f* (*x*) is a function that is peaked at *x* = 0 and monotonically decreases for increasing |*x*|, with a limit lim_*x*→±∞_ *f* (*x*) = 0 (Pigolotti et al., 2010; Leimar et al., 2013). We assume a symmetric interaction *f* (*x*) = *f* (−*x*), reflecting our earlier assumption of symmetric competition coefficients.

For our explicit analyses we use periodic boundary conditions on the niche axis^4^; similar results hold for a non-periodic finite axis, but periodicity simplifies the analysis. In the concrete analyses carried out here, we use interactions of the form

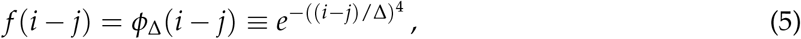

where Δ is a constant that fixes the scale on the lattice. This functional form for the interactions has some precedent in the literature (Leimar et al., 2013), and avoids some degeneracies present for a Gaussian interaction kernel. This is a highly-localized, almost box-shaped kind of interaction kernel; we expect similar results for other similarly localized interactions although the features may be more subtle. Specifically, the Gaussian case separates two different kinds of behavior (Pigolotti et al., 2010; Leimar et al., 2013; Barabas et al., 2013). For more sharply-peaked, less localized kernels (for example, an exponential kernel) there can be solutions in which species with arbitrarily close spacing coexist. ^5^ For interaction kernels more localized than a Gaussian, like ours, however, late time behavior tends to lead to sharp peaks, and solutions generally have a minimum distance between species along the niche axis, known as limiting similarity (see below).

On a periodic lattice of size *S*, the precise interaction matrix is then

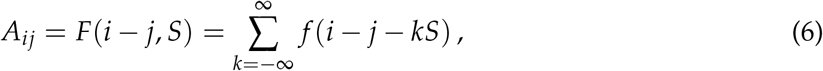

where the summation over *k* implements the effects on species *i* of all copies of species *j* on the periodic lattice. Because of the ultra-local nature of the interactions (5), for *S* » Δ, the only significant contributions to *A*_*ij*_ come from local lattice sites, with negligible effects from the periodicity. The circulant form of the interaction matrix (6) provides some simplifications in analysis, but similar features are expected for local competitive interactions without these symmetry properties (see further discussion in Appendix). The periodic niche models of interest here can thus be characterized by an interaction function 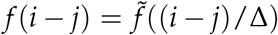, a distance scale (in lattice units) Δ, and a lattice size *S*.

For any form of the interaction function *f* (*i* − *j*), due to the circulant, non-negative form of *A*_*ij*_, there is a global equilibrium with all 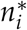 equal (and positive). Similar to the monodominant models described above, this equilibrium is generally unstable but there can be multiple stable subsystem equilibria, at the boundaries of the positive orthant (Case, 1980). In general, for an interaction that decreases sufficiently quickly with increased separation on the niche axis such as (5), the idea of *limiting similarity*, (MacArthur and Levins, 1967; May and MacArthur, 1972) suggests that there should be a characteristic minimum distance scale (in lattice units) *d* ∝ Δ associated with the minimum separation between viable species in stable equilibria, and that species will pack as closely as possible to this minimal separation. As described in detail in the Results section, we find a somewhat richer structure for the set of possible stable and non-invasible equilibria, with the separation between species in equilibria ranging from roughly *d* to 2*d*. This can be understood qualitatively by noting that a new species can only successfully invade when the separation between a pair of species approaches 2*d*.

There are two interesting limits of these models in which the structure simplifies. First is the limit where Δ, *S* → ∞ with fixed *S*/Δ. This corresponds to a continuum limit of a system of fixed size, where the interaction range between species stays constant as a fraction of the system size; in this limit the number of species in stable equilibria takes one of a fixed, finite set of values. This is a limit of high diversity, but specifically where each species interacts with a fixed fraction of the (increasingly large) number of other species. One could think of this as identifying finer-and finer-grained distinctions between individuals, and using these fine-grained distinctions to classify species. This issue arises in classifying bacterial species, where a range of different cutoffs in sequence similarity have been used to define species. The second limit is where we fix Δ and take *S* → ∞; this limit corresponds to a kind of statistical mechanical model where the system size becomes large compared to the range of interactions. This also corresponds to a limit of high diversity, but where each species only ever interacts strongly with a fixed number of other species, even as the number of species becomes large. Ecologically, this limit corresponds to increasing the trait diversity of the entire pool of species, extending in effect the range of the niche axis. In this limit the number of equilibria grows exponentially in *S*, while the number of species in stable equilibria grows linearly with *S*. These limits are discussed in more detail in the Results section. Any model with finite, fixed values for Δ, *S*, such as might describe a natural system, will share features with both the continuum and statistical mechanical limits.

Equilibria for two explicit examples of discrete niche systems are illustrated in Figure 1.

**Figure 1:**
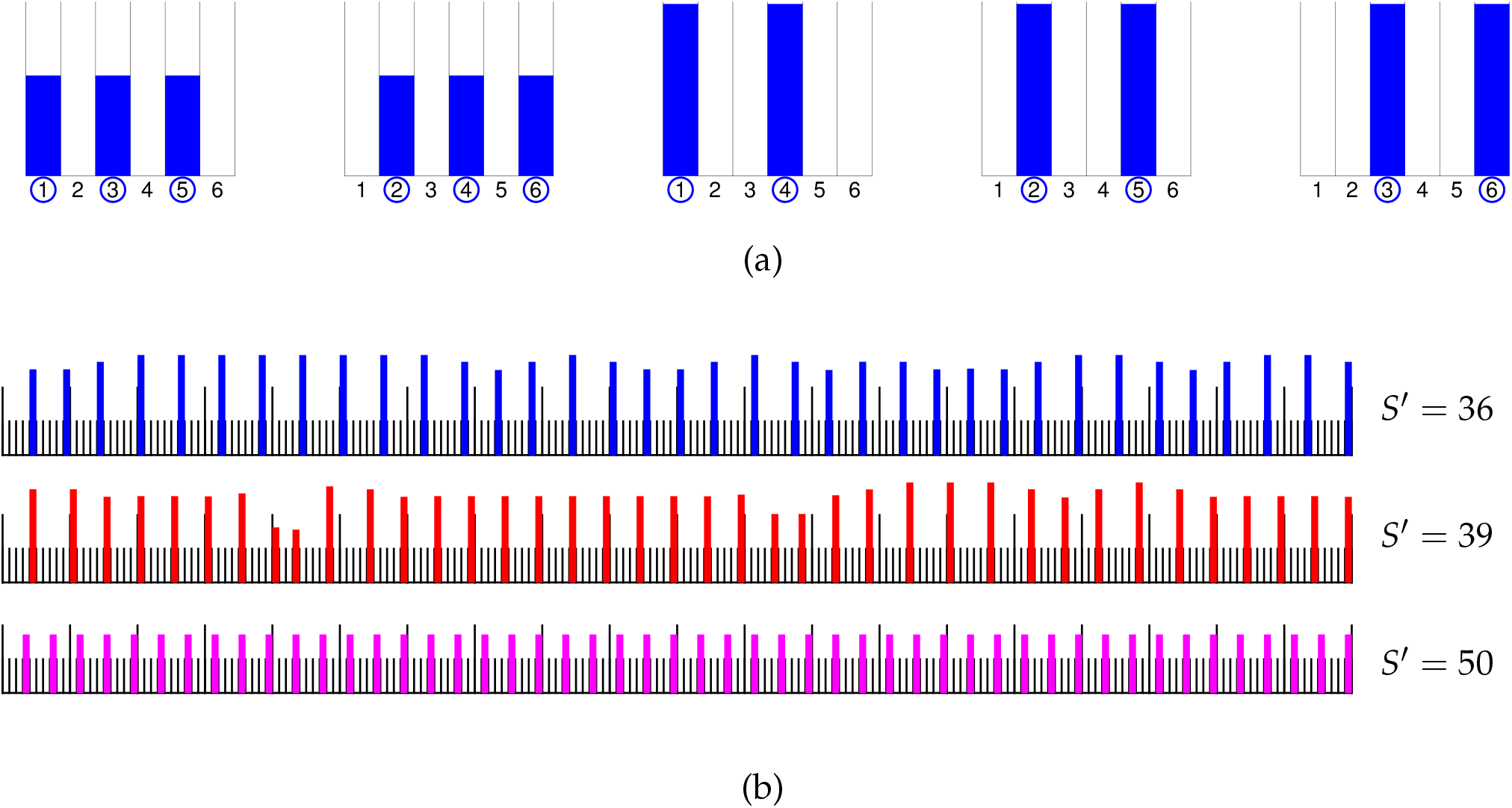
Stable equilibrium solutions for two example discrete periodic niche systems with competitive interaction (5): (a) Complete set of 5 stable equilibria for model with *S* = 6 species and Δ = 2. (b) Three distinct equilibrium solutions of system with *S* = 200 species and Δ = 4, with *S*^′^ = 36, 39, 50 species having nonzero population respectively. This system has thousands of distinct equilibria (number of equilibria grows exponentially in *S* for a given Δ). Equilibria with 36, 39 species are typical results of simulation with random initial conditions. Equilibrium with 50 species is extreme case that realizes “limiting similarity” but is statistically unlikely.

#### Random matrices

Modeling *A*_*ij*_ as a random matrix has a long and rich history in theoretical ecology, following the pioneering work of May (1972). While that original and much subsequent work (Allesina and Tang, 2012) focused on the question of stability of the global equilibrium, more recent work (Bunin, 2017) has identified a regime in which the global equilibrium is unstable but there are many stable subsystem equilibria—similar to the outcome of our niche axis models, above. We thus consider a four-parameter family of random matrix systems, where

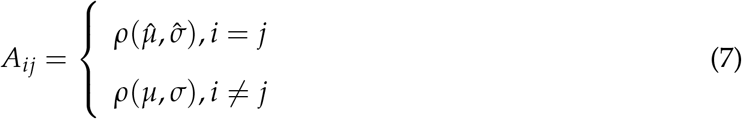

Here, *ρ*(*µ, σ*) is a random distribution with mean *µ* and standard deviation *σ*.

More generally, we can consider a combination of our niche-axis and random matrix models— a semi-local random matrix system where the off-diagonal interactions are suppressed by a factor *f* (|*i* − *j*|) (or the periodic analogue (6)) for some positive localization function *f* with *f* (0) = 1 that decreases towards 0 as |*x*| increases. Thus, in the general form of the model we take *A*_*ij*_ = *ρ*(*µ, σ*) *f* (*i* − *j*), *i* ≠ *j*. This gives a broader class of systems that continuously interpolates between the random matrix systems generally studied in the literature and the niche axis models described above (where in this formulation we have *µ* = 1, *σ* = 0).

### Analysis techniques

#### Numerical analyses

For some smaller systems with *S* ≤ 30, we have carried out explicit analyses of the full set of stable equilibria by a brute force search in which the solution (2) is computed and analyzed for stability for all 2^*S*^ − 1 possible (nonempty) subsets. Such analysis allows definitive identification of the complete set of possible stable equilibria but does not provide information about the sizes of their domains of attraction.

To address the latter question, we have carried out numerical simulations for a number of models. These numerical simulations start with a random initial condition according to a specific distribution, and perform many runs to estimate the distribution of stable equilibria reached, thus estimating the sizes of the basins of attraction. The relative domain sizes of the basins of attraction depend to some extent on the distribution of initial conditions; in most of the work here we consider initial conditions distributed according to a Gaussian with mean 0 and standard deviation equal to the value of each population at carrying capacity, restricted to the positive orthant. From the geometric decomposition described below, and some numerical tests (see Table 1 and associated discussion), we expect that the relative domain sizes are in general relatively insensitive to the specific choice of initial conditions.

**Table 1:**
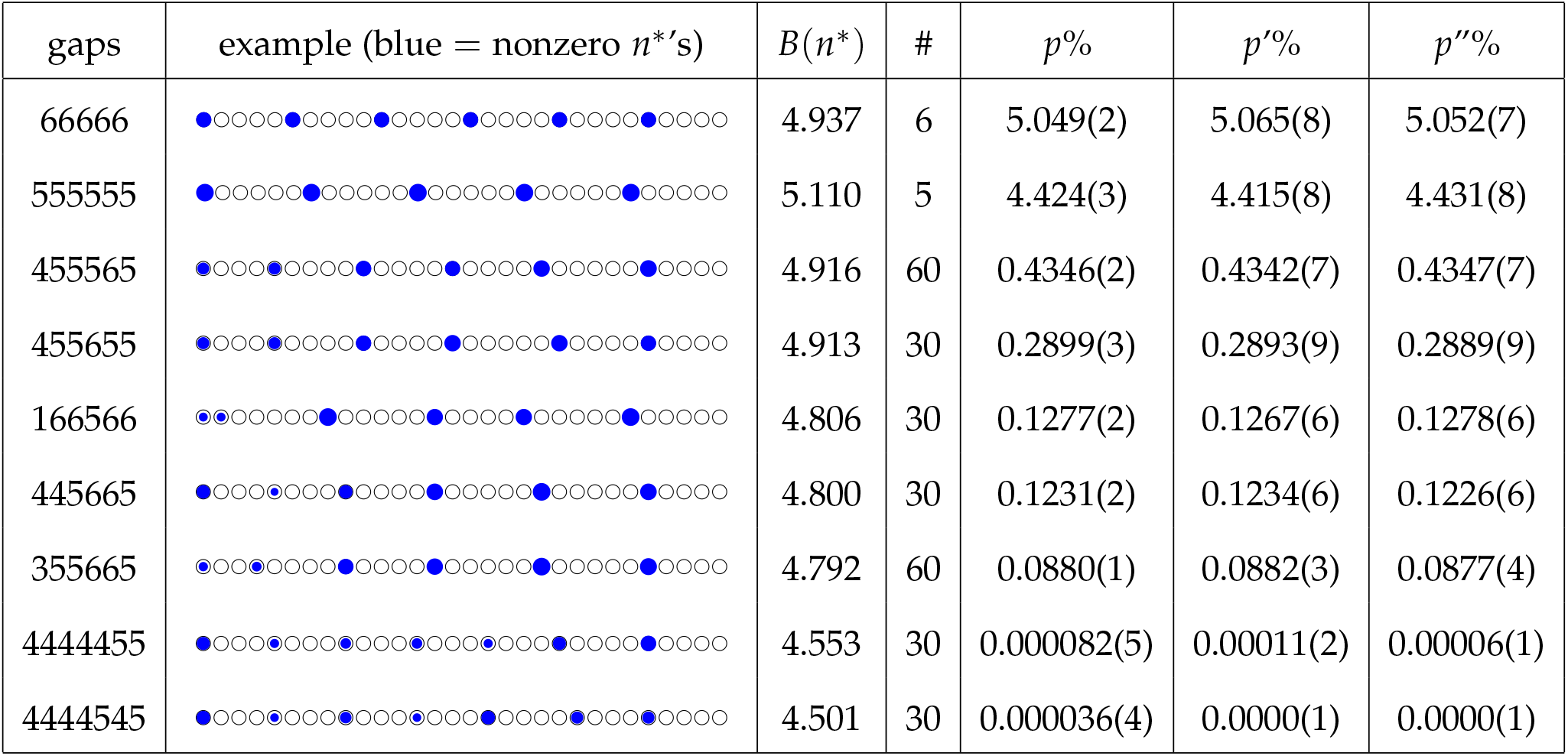
Probabilities and biomass *B*(*n*^*^) of the 281 distinct equilibria for a periodic niche axis model of size *S* = 30 and interaction (5), with Δ = 4. Equilibria are labeled by sequence of distances (“gaps”) between nonzero populations; # (multiplicity) refers to number of equilibria with same gap sequence up to rotation and reflection symmetry. *p*% is percentage of 10^7^ simulations giving any specific equilibrium in class for standard initial conditions (so Σ(*p*×#) = 100); *p*^′^%, *p*^′^% are percentages with two alternative initial distributions, essentially identical statistically to *p*%.

The code to perform the exact analyses was written in Mathematica, and the simulations were performed using Mathematica and Julia.

#### Eigenvalue analysis

It can be useful in models with an unstable global equilibrium to analyze the spectrum of eigenvalues of the *A* matrix, which control the stable and unstable directions around the unstable fixed point. The eigenstates associated with negative eigenvalues correspond to fluctuations around the unstable fixed point that lead to a decrease in the Lyapunov function *H*, generating flow towards a stable equilibrium. In general, more negative eigenvalues lead to equilibria with larger basins of attraction.

In general, as occurs for the niche systems studied here, the number of stable equilibria can be much larger than the number of negative eigenvalues. The eigenvalue/vector structure nonetheless gives insight into the structure of the equilibria. Three different classes of systems studied here exhibit rather different behavior in terms of eigenvalue analysis. For the monodominant systems with *S* species, there is roughly a one-to-one correspondence between negative eigenvalues and equilibria, with *S* stable equilibria, and *S* − 1 negative eigenvalues of *A*. In the continuum limit of niche models, on the other hand, the eigenvectors of *A* with negative eigenvalues correspond to periodic fluctuations where the wave number corresponds to the number of species in the associated stable equilibrium, and pairs of negative eigenvalues are related to O(*S*) distinct solutions with different offsets on the niche axis. And in the statistical mechanical model limit of niche systems, the number of solutions grows exponentially with the number of negative eigenvalues. These relationships are all described in further detail in the Results section.

#### Statistical Mechanical model of niche equilibria

For a periodic niche system as described in the Niche axis models section, the equilibria can be characterized by the positions of the nonzero populations (“spikes”) (i.e., 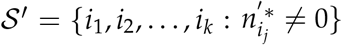. Alternately, we can characterize such an equilibrium by the “gaps” between the spikes, *g*_*j*_ = *i*_*j*+1_ − *i*_*j*_, *j < k, g*_*k*_ = *i*_1_ + *S* − *i*_*k*_. As *S* → ∞, for any fixed local interaction function *f* (*x*) (e.g., (5) with fixed Δ) we expect that the probability (i.e., domain size) of an equilibrium characterized by a sequence of gaps *g*_*j*_ can be described by a local statistical mechanical model

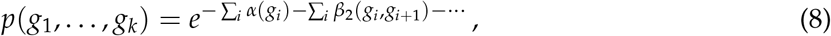

where *α*(= *β*_1_) is a weighting associated with a single gap, *β*_2_ is a weighting for an adjacent pair of gaps, etc.. The expectation is that weightings for the higher-order *n*-gap correlations will become relatively weak as *n* becomes large, since the effect of species at a fixed lattice site *i* on a lattice site *i* + *c* becomes arbitrarily small as *c* → ∞ when lim_*x*→±∞_ *f* (*x*) = 0. We investigate and validate this hypothesis for the niche models in the Results section.

#### Geometry of multiple equilibria

For a system of species 𝒮 with interaction matrix *A* and an equilibrium 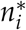, where (𝒮, *A*) may either be the complete system, or a subsystem, and where the equilibrium can be stable or unstable, we note that the assumption that all *r*_*i*_ = 1 implies that the ray (half-line) from the origin (*n*_*i*_ = 0) through the equilibrium 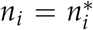 is a fixed locus of the dynamics (proof in Appendix). This observation can be used to give an approximate geometric picture for the basins of attraction of the stable equilibria of simple systems by dividing the positive orthant into cones separated by these rays; the details of this geometric approximation are spelled out in the Appendix. In the Results section, we comment on the accuracy of this geometric approximation. Note that for a system with *S* = 2, this approximation is exact, as described in the first example below.

## Results

### Simple example: Monodominant systems

To illustrate the general approach, we consider first a simple set of monodominant models, beginning with a two-species model with *a*_1_ = *a, a*_2_ = 1, so that the interaction matrix is

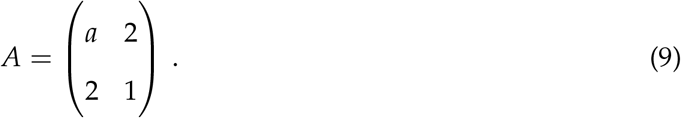

This system has an unstable equilibrium at 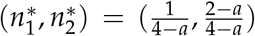, and stable subset equilibria at 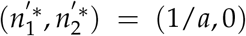 and (0, 1). The basins of attraction are precisely divided by the line connecting the origin to the global unstable equilibrium *n*^*^ (Figure 2(a)), so the geometric decomposition described in the Methods section is exact in this example. Some details of this model are described in the Appendix, including an analytic formula for the relative domain sizes *D*_1_, *D*_2_ (normalized so that *D*_1_ + *D*_2_ = 1) for the two equilibrium for any initial distribution that is independent of angle. The relative domain size *D*_1_ is a monotonically increasing function of the equilibrium population 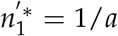, which in this case is the total biomass 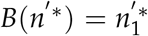 of the single-species stable equilibrium (Figure 2(b)).

**Figure 2:**
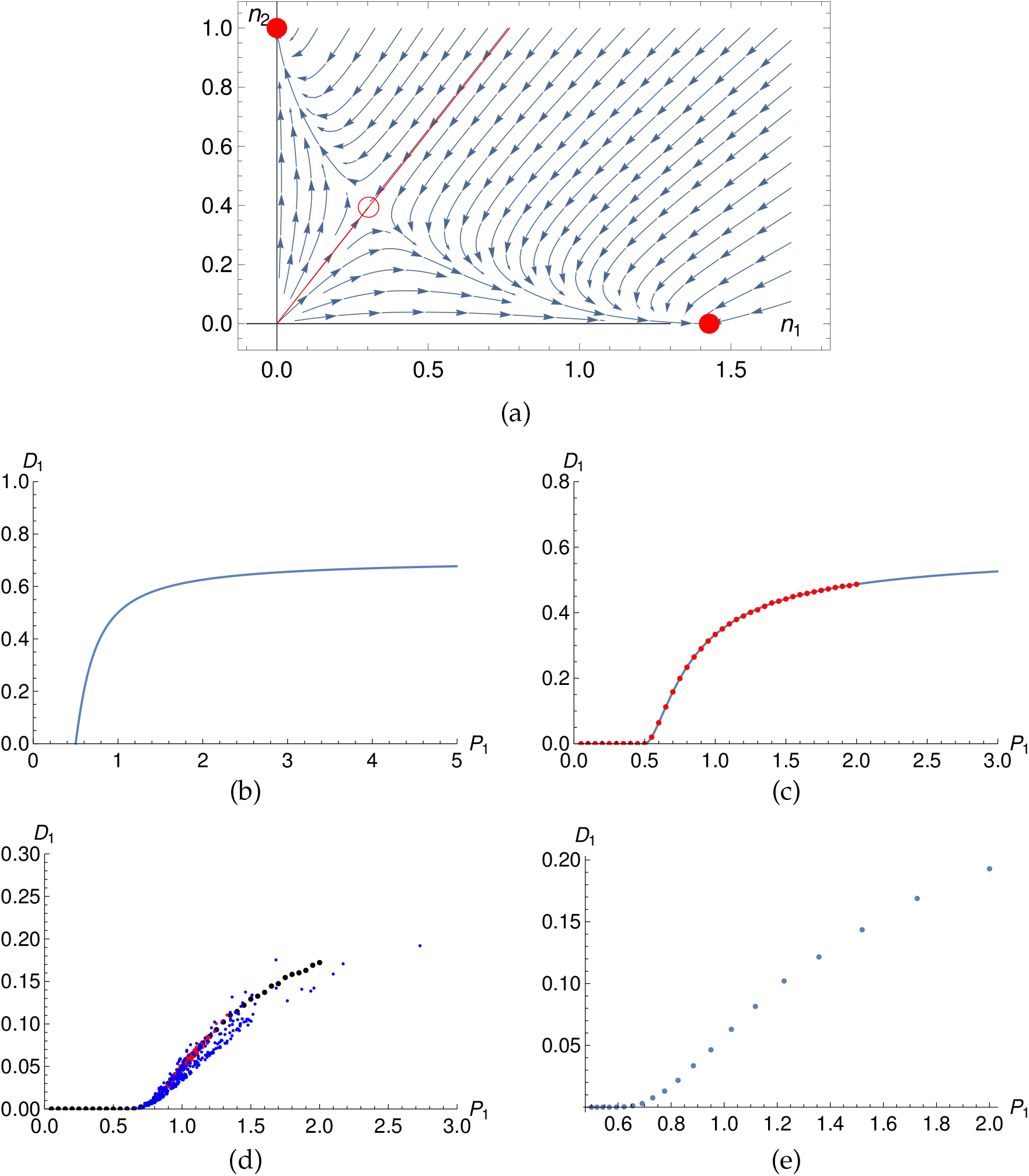
Domain size is monotonically increasing in population for monodominant models. (a) 2-species model, *a* = 0.7: domains of attraction of stable equilibria (disks) are divided by (red) line through unstable equilibrium (circle). (b, c) Domain fraction *D*_1_ as a function of population 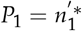 in 2- and 3-species systems; (b) is analytic, (c) solid (blue) curve is geometric approximation, dotted (red) curve is numerical estimate. (d, e) 20-species monodominant system, domain fraction as a function of population; (d) *a*_*i*_ ‘s from Gaussian distribution centered at 1 with *σ* = 0.2(08.0)[0.0] in blue (red) [black]; (e) *a*_*i*_ linearly distributed from 0.5 through 1.9. Further details in Appendix.

For a three-species monodominant model, the results are similar. Considering again the one-parameter model with *a*_1_ = *a, a*_2_ = *a*_3_ = 1, we compare the approximate domain size from the geometric decomposition analysis with numerical computation, with extremely close agreement between the methods (Figure 2(c)). The domain size 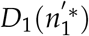 is again a monotonically increasing function of the equilibrium population, and has similar qualitative features.

The results for the monodominant model with large numbers of species *S* are again similar. Numerical computations show that the domain size is a monotonically increasing function of equilibrium population. In general the domain size grows as the *S* − 1 power of the difference of the population just above the threshold of vanishing domain (i.e., linear for *S* = 2 above *n ′* ^*^ = 1/2, quadratic for *S* = 3, as described in Appendix, etc.). We have analyzed such systems in several ways: first allowing a variable single parameter *a*_1_ = *a*, with the other *a*_*j*_ = 1, second taking all *a*_*i*_ from a Gaussian distribution, and third distributing all the *a*_*i*_’s uniformly over a fixed range. We find in the first case that the domain size dependence 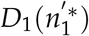 is again a monotonically increasing function of 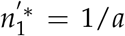. In the second case, we get a distribution of 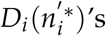 that is centered around the function 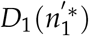 seen in the single-variable case but with linear and higher order fluctuations from the other *a*’s, and in the third case we see a clear monotonic dependence of 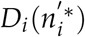 (Figure 2(d, e)).

In the monodominant models the relationship between the eigenvectors of *A* and the *S* stable equilibria of the system is fairly transparent. In the case where all *a*_*i*_ = 1, there are *S* − 1 degenerate negative eigenvalues of *A*, associated with the (*S* − 1)-dimensional space of eigenvectors (*v*_1_, …, *v*_*S*_) with Σ_*i*_ *v*_*i*_ = 0. Geometrically, a fluctuation around the unstable equilibrium in such a direction will put the system in the domain of attraction of the equilibrium associated with the species *j* with the largest *v*_*j*_. For example, in a 3-species system with all *a*_*i*_ = 1, a fluctuation in the direction (−1, −1, +2) will push the system into the domain of attraction of the stable equilibrium with only species 3 (since this fluctuation increases the population of species 3 and decreases the populations of species 1, 2), while a fluctuation in the direction (0.9, 1.1, −2) would push the system into the stable equilibrium with only species 2, by symmetry (*v*_2_ *> v*_1_). As discussed further in the Appendix, with varying values of the *a*_*i*_, the eigenvalues become non-degenerate, and the smallest *a*_*j*_, corresponding to the solution with largest population 1/*a*_*j*_, is generally associated to the (negative) eigenvalue with the largest absolute value, although determining precisely the results of a fluctuation corresponding to a general linear combination of eigenvectors with negative eigenvalues is not straightforward.

To summarize, the monodominant models provide a simple class of model systems where a collection of *S* species has an unstable global equilibrium with all species present, and *S* distinct equilibria with a single species persisting. The relative size of the domain of attraction of each of the distinct equilibria 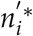 can be described as a monotonically increasing function of the equilibrium population 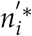, with variability depending on all parameters other than the one (*a*_*i*_) controlling the value of 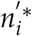.

#### Niche systems

We describe here various results regarding the multiple equilibria of the niche axis systems described in the Methods/Models section. As described there, we consider periodic lattice models with *S* distinct species. We focus particularly on the case of interactions of the form (5), parameterized by a characteristic length Δ, as an illustrative example; qualitative features are expected to be similar for other highly localized interactions. The two distinct limits where analytic treatment is possible provide insight into the general discrete non-limiting systems.

#### Classification of equilibria

For relatively small discrete systems we can simply enumerate all possible subsystems and check for a stable^6^ equilibrium. For example, for a periodic system of size *S* = 30 with Δ = 4, there are 2^30^ ∼ 10^9^ possible equilibria and an explicit check shows that there are 281 of these that are feasible and stable (See Table 1). These equilibria include configurations with 5, 6, and 7 distinct species with nonzero populations.

We have used numerical simulations to estimate the domain sizes of the distinct equilibria in the *S* = 30, Δ = 4 niche model with several different choices of initial distribution, described in the last three columns of Table 1. In the first case (*p*%), we ran 10^7^ simulations with the “standard” initial condition of a Gaussian (restricted to the positive orthant) centered at 0 with standard deviation of 1 (the carrying capacity of each species). In the second case (*p*^′^ %), we did 10^6^ simulations with a Gaussian with standard deviation given by the population value at the unstable equilibrium (∼ 0.138). In the third case (*p*^″^%), we did 10^6^ simulations starting with a uniform distribution in the box with all initial populations in the range 0 ≤ *n*_*i*_ ≤ 1. ^7^ these three initial distributions all give probabilities that agree to better than one part in 1000 for the more common equilibria, and are essentially statistically equivalent, confirming that the domain sizes do not depend strongly on initial conditions.

For larger systems, the number of possible subsystems grows exponentially in *S*, and complete enumeration of all equilibria becomes intractable, but numerical simulations give a sampling of the set of equilibria and a measure of the corresponding domain sizes. For the continuous limit (*S* → ∞ with fixed *S*/Δ) and statistical mechanical limit (*S* → ∞ with fixed Δ), we can systematically analyze many aspects of the equilibria. For finite systems with specific values of *S*, Δ, the solutions combine features that can be seen in these two limits.

#### Continuum limit

In the continuum limit, *S*, Δ → ∞ with fixed *S*/Δ, the dynamical equation (1) approaches the integrodifferential equation

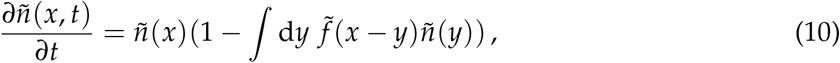

where *x* = *i*/Δ ∈ (0, *S*/Δ] is a continuous parameter replacing the discrete parameter *i* and 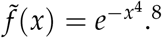

The equilibrium solutions in this limit have equally spaced spikes separated by a distance *d*. In general there is a range of possible values for the separation *d*, with 0.91, :. ≲, *d*: ≲ 1.82. This can be understood in more detail as follows: Working in lattice units *i*, there is an equilibrium solution of (1) with spikes at equal distances *D* = *d*Δ when *n*_*kD*_ = 1/*F*(0, *D*), ∀*k*, with *F* defined in Eq. (6), corresponding to a solution of (10) with spike separation *d*. For the interaction (5), this solution is stable (including against invasion) in the approximate range 0.91 Δ, :≲ *D*, :≲ 1.82 Δ (details in Appendix). This roughly corresponds to the idea that there is a limiting distance within which species cannot coexist (which for this particular interaction is around 0.91Δ); we see, however, that stable equilibria can persist with spacing up to about twice this distance, at which point new species can enter between a pair.

In the continuum limit, there is a clear correspondence between negative eigenvalues of *A* and stable solutions. A fluctuation of the form *δ* ñ = ϵ cos(2*πx*/*d* + *θ*) around the constant unstable equilibrium 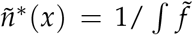 corresponds to a stable solution with spacing *d* in the continuum limit for *d* approximately in the range from 0.9 to 1.8. Furthermore, there is a clear correlation between the size of the eigenvalue and the biomass density for these solutions (see Appendix for more details, particularly Figure A1).

At finite but large *S*, the solutions essentially match the expectation from the continuum limit. We describe in some further detail how this correspondence plays out in practice. For any number of spikes *s* with *D* = *S*/*s* roughly in the allowed range, there is a solution to the discrete system. These solutions have multiplicity of at least *S*/*s*, associated with different locations of the first spike on the lattice. There is also some redundancy in these solutions due to finite size effects. For example, when *S*/*s* is not an integer, the spacings will be slightly uneven. More generally, slight variations in distances give a variety of viable solutions at finite *S* with a fixed number of spikes. Furthermore, in some cases, a spike can be evenly split between a pair of adjacent lattice sites. These considerations explain, for example, why the gap sequences such as 455565, 166566 described in Table 1 are essentially discrete versions of the 6 and 5 spike solutions of the *S*/Δ = 7.5 system at the finite value *S* = 30. Similarly, 4444545, 4444455 are the only 7-spike solutions. Solutions of this nature persist at larger *S* but the spacings approach the expected (approximate) values 1.5Δ, 1.25Δ, 1.07Δ in the continuum limit. Note that in this limit there are also solutions of the *S*/Δ = 7.5 system with 8 spikes (*D* = 0.9375 Δ; for example 44444444 at *S* = 32) but no solutions with only 4 spikes (*D* = 1.875 Δ).

#### Domain sizes in the continuum limit

By performing numerical experiments on finite *S* systems with a fixed value of *L* = *S*/Δ, we can confirm that the (relative) sizes of the domains associated with a fixed number *s* of spikes converge as expected in the continuum limit (see Figure 3(a); for example, probabilities *p*(*s*) of *s* = 5 spikes are 34.13%, 36.18%, 36.21% at *S* = 30, 60, 90 based on 10^7^, 10^5^, 10^5^ simulations respectively.). Furthermore, the numbers of spikes associated with more negative eigenvalues correlate well with solutions having larger domain sizes, as well as with larger biomass as discussed above.

**Figure 3:**
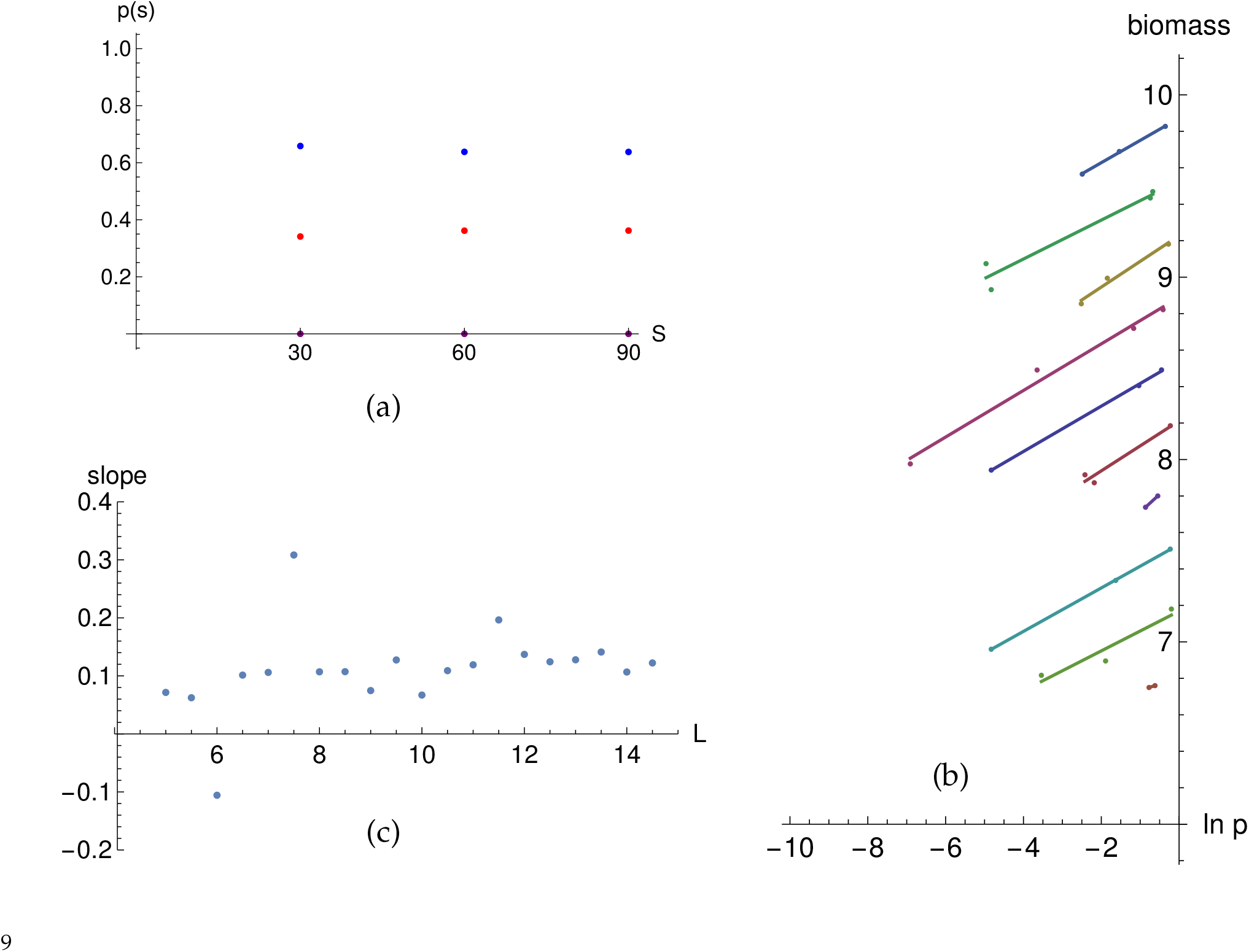
Continuum limit of niche models: (a) Probabilities of *s* = 5 (red), 6 (blue), 7 (purple) spikes (persistent species) in model with *L* = *S*/Δ = 7.5 and *S* = 30, 60, 90 converge rapidly as *S* increases. (b) Relationship between the log of the probability of finding an *s*-spike solution ln *p*(*s*) and average biomass for solutions with *s* spikes, *L* = 10 (bottom plot), 10.5, …, 14.5 (top plot), each in a different color with linear fit, illustrates linear relationship at each *L*. (c) Slopes given by a linear fit to numerical data for the ln *p*/biomass relationship for various values of *L*.

We can directly compare the average biomass *B*(*s*) realized for solutions with *s* spikes with the logarithm of the probability of the solution having *s* spikes. For the range of values of *L* we explored (from *L* = 5 to *L* = 14.5), the relationship between these quantities for each *L* is strikingly close to linear. (See Figure 3(b,c); For each value of *L*, 1000 simulations were done, with *S* = 101 (prime). For example, at *L* = 13, 662 simulations gave 10 spikes, while the number of simulations giving 9, 11, and 8 spikes respectively was 311, 26, and 1.). We do not have a simple and complete theoretical argument for this linear relationship in this limit. In the following discussion of the statistical mechanical limit, on the other hand, we provide a theoretical proof that this kind of linear relationship between biomass and log probability is natural and expected due to a version of the central limit theorem.

The fact that there are distinct solutions with different numbers of spikes *s* in the continuum limit, each of which has a nonzero probability of arising from random initial conditions, provides an interesting perspective on the standard notion of “limiting similarity”. In particular, these models make it clear that while one particular spacing between occupied niches may be preferable, multiple possibilities may be feasible in nature.

#### Statistical mechanical limit

As mentioned in the Methods section, there is an interesting limit of the niche axis models where the interaction function and scale Δ are fixed, but the number of species is taken to be large, *S* → ∞. In this limit the equilibria can be described by a local statistical mechanical (SM) model; we thus refer to this as the SM limit. Basically, the idea is that if the interaction function is local, meaning that *f* (*i* − *j*) vanishes rapidly as |*i* − *j*| grows, a species located at position *i* will only affect those species with nearby values of *j*. If we characterize equilibria by the sequence of gaps *i* − *j* between successive species with non-vanishing populations, then whether a given gap sequence corresponds to an allowed equilibrium can be determined by the local structure of the gap sequence in the SM limit.

As an example, for the models with Δ = 4 and general *S*, all local gap sequences containing an arbitrary sequence of 5’s and 6’s are possible (at least up to 6 gaps, and presumably for any finite gap sequence) except those containing the subsequence 5656 (or 6565); this set of equilibria alone grows exponentially with *S*. On the other hand, gaps of size 2 or greater than 7 do not arise. ^10^ This corresponds well with the expectation from the continuum analysis above that the spacing between species should be in the range 0.9Δ ≅ 3.6 ≤ *D* ≤ 1.8Δ ≅ 7.2.

Using local conditions on which gap sequences are possible gives a simple model (described in more detail in the last two sections of the Appendix) that describes the exponentially large number of equilibria for these niche systems with fixed Δ and large *S*. While the particular local rules depend upon the choice of Δ and the interaction function *f*, we expect a similar structure for any sufficiently local interaction and finite Δ. This kind of local model for the set of possible equilibria naturally generalizes to a statistical model incorporating probabilities representing domain fractions, to which we now turn.

#### Domain sizes in the statistical mechanical limit

Given the local structure of equilibria in the niche axis models, the statistical mechanical model (8) gives a useful characterization of the set of equilibria in the large *S*, fixed Δ (statistical mechanical) limit. In this model the probability of a given equilibrium represents the normalized domain fraction associated with that equilibrium. As described previously, the specific probabilities and domain fractions depends upon the precise choice of distribution over possible initial conditions. For the analysis here we continue to use the standard distribution described in the Methods section, but results should be very similar for other initial distributions with the same angular dependence.

By performing a large number of simulations with random initial conditions, we can numerically estimate the parameters in the statistical model (8) for a given lattice size *S*, parameter Δ, and initial distribution. We briefly summarize some salient aspects in an example here, further details and analysis appear in the last three sections of the Appendix. Based on (10^6^) random simulations with *S* = 90, Δ = 4, the single gap weightings *α*(*g*) = *β*_1_(*g*) can be determined from the observed single-gap probabilities *p*(*g*) = *e*^−*α*(*g*) 11^

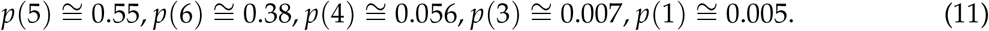

(No gaps of size 2 or greater than 6 are observed^12^.) The probability of a given adjacent pair of gaps is not simply the product of the single-gap probabilities; the most common adjacent gap pairs have probabilities 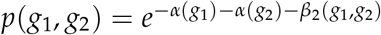 observed as

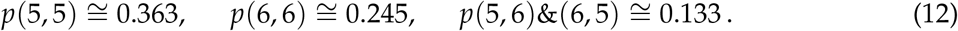

These probabilities satisfy *p*(5, 5) *> p*(5)^2^, *p*(6, 6) *> p*(6)^2^, *p*(5, 6) *< p*(5)*p*(6), i.e. *β*_2_(5, 5), *β*_2_(6, 6) *<* 0, *β*_2_(5, 6) *>* 0. This can be interpreted as local efforts to space evenly. Looking at longer sequences, the additional effects of weightings *β*_3_, *β*_4_ from sequences of length 3, 4 is relatively small and is negligible beyond length 4. This can be quantified by considering the information (Shannon) entropy^13^ (in bits) *σ*_*k*_ per symbol of sequences of *k* gaps,

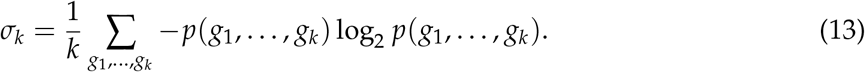

Using data from the sampling described above gives

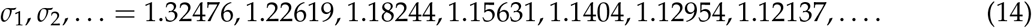

This shows that most of the structure is in the one and two point functions; with only a 10% correction from two-point correlations, 4% from three-point correlations, etc., verifying the local nature of the structure of the set of equilibria. In this way, the local model including only the parameters *α*(*g*), *β*_2_(*g*_1_, *g*_2_) already gives a fairly accurate picture of the set of allowed equilibria and their relative domain sizes.

Note that the number of negative eigenvalues of the interaction matrix is limited by *S*, so that when *S* is large there are exponentially more stable equilibria than there are independent unstable directions around the unstable equilibrium described by distinct negative eigenvalues. Nonetheless, the spectrum of eigenvalues gives an accurate picture of the general structure of the equilibria. For *S* = 200, Δ = 4 the eigenvalues of *A* are graphed in Figure 4. Because of the circulant nature of the interaction matrix, these eigenvalues are associated with periodic eigenvectors (1, *e*^2*πik*/*S*^, *e*^2*πi*2*k*/*S*^, …), *k* = 0, …, *S* − 1. The most negative eigenvalues are associated with eigenvectors with a wavelength of between 5 and 6 lattice sites (wave number *k* ∼ 37), in accord with the observation that 5 and 6 are the most frequently observed gap sizes.

**Figure 4:**
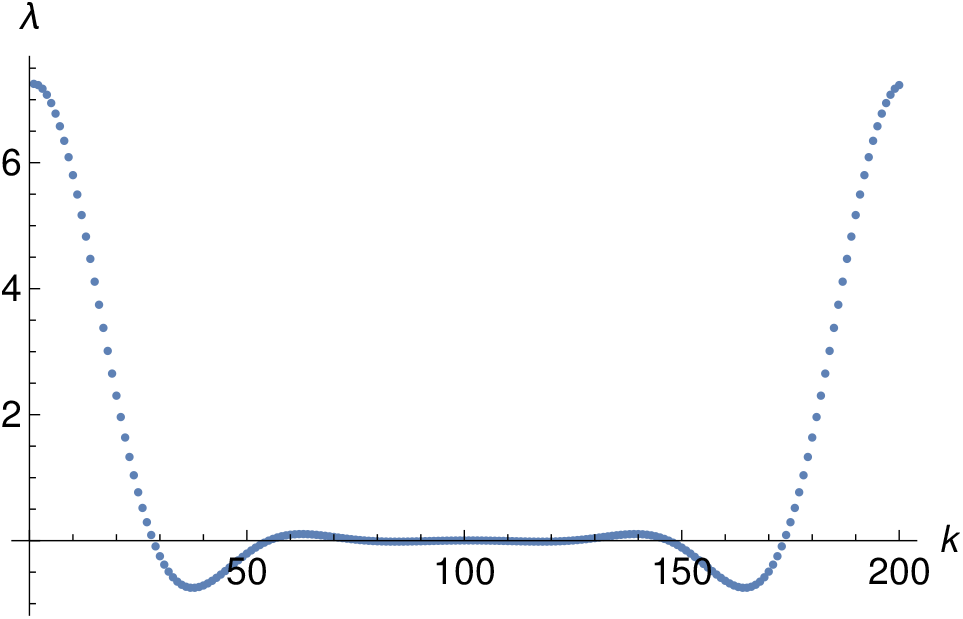
Eigenvalues of *A* for the niche axis model with *S* = 200, Δ = 4, indexed by wave number *k*. The most negative eigenvalue corresponds to a periodicity of ∼ 5.4 (*k* = 37), correlating with the observation that 5 is the most frequently observed gap size, followed by 6.

#### Domain size and populations

In the simple monodominant models studied above, we identified a general monotonic dependence of domain size on equilibrium biomass. For the niche axis models we also find a monotonic dependence, and in the continuum limit we described an apparent exponential dependence of domain size on biomass. In the statistical mechanical limit we observe a similar exponential dependence, which in this limit can be explained analytically. In particular, for the niche axis models in the SM limit we find the approximate relationship

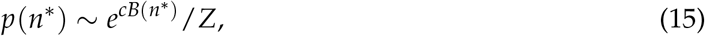

where *p*(*n*^*^) is the relative probability of a given stable equilibrium^14^, *B*(*n*^*^) = −*H*(*n*^*^) is the total biomass 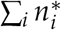 of that equilibrium (i.e., the negative of the Lyapunov function), and *c, Z* are constants. More precisely, the statement is that as *S* → ∞ with fixed Δ, the distribution in (−*H*, ln *p*) space of the set of equilibria found over a large number of simulations with random initial conditions approaches a multivariate normal distribution, so that for equilibria with biomass near a specific value −*H*, the mean value ⟨ln *p*⟩ grows linearly in −*H*.

Numerical evidence for this relation for systems of size *S* = 30, 50 is given in Figure 5(a, b) (the case *S* = 60 is shown in Figure A5). This kind of relation can be shown to arise in any situation where the domain size and biomass are both determined by local features. In particular, in any system described by a statistical model as above with negligible weightings for *n*-gap correlators above some fixed value *n > k*, this result follows from the Markov chain central limit theorem and the fact that the domain size, controlled by the local statistical model (8) is a product of local factors, while the biomass is a sum of local factors, so there is a linear correlation between the log of the domain size and the biomass. Details of the proof of this assertion are given in the Appendix. Note that when there is a positive correlation between domain size and biomass, the constant *c* is positive; this occurs in all situations we have considered but there may be anomalous circumstances where the correlation is negative and *c <* 0. Note also that the statement that can be proven most rigorously involves the distribution where equilibria are sampled with probability *p*(*n*^*^); see the Appendix for further discussion of related subtleties.

**Figure 5:**
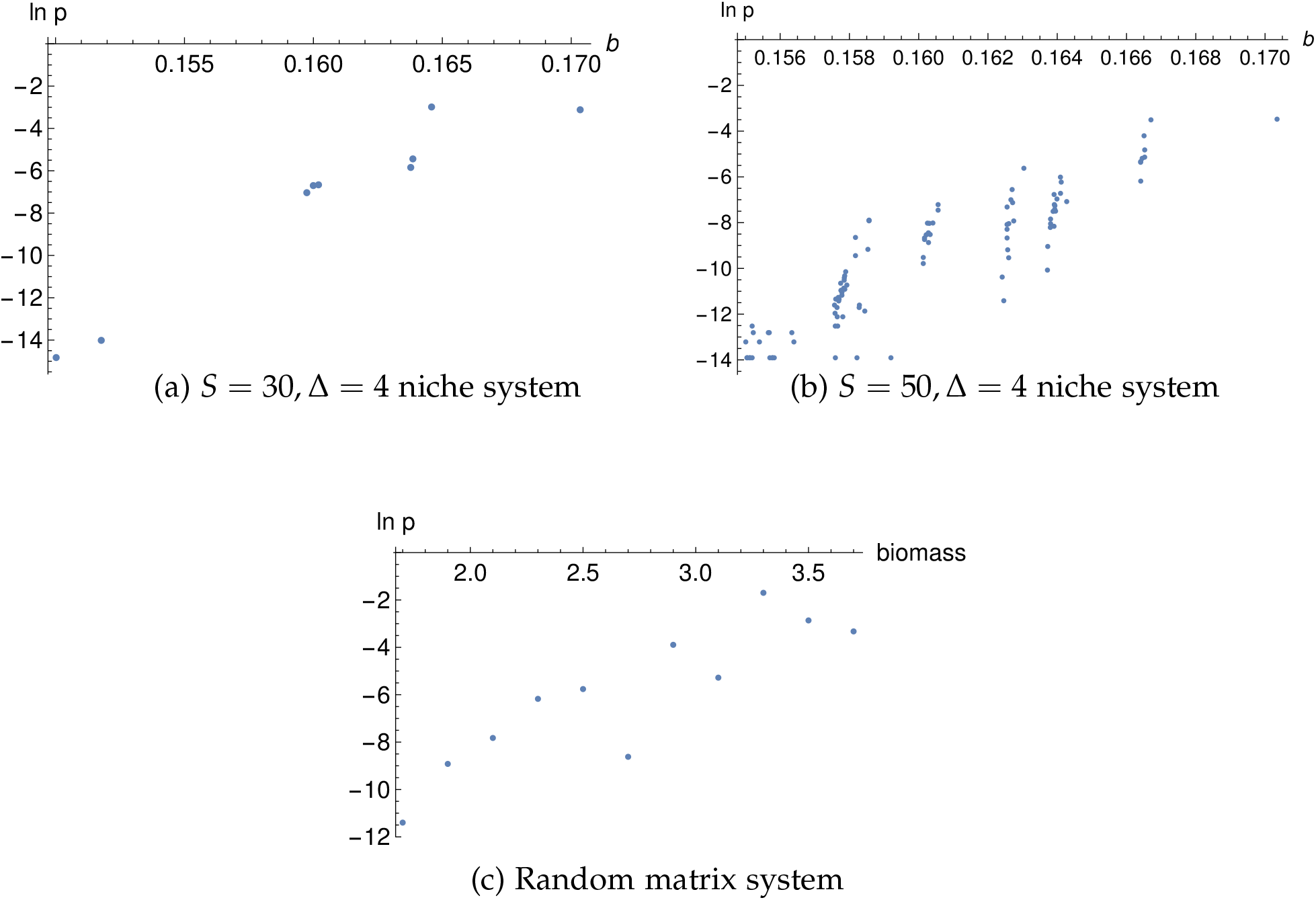
Relationship between biomass density *b* = *B*(*n*^*^)/*S* and ln *p* (logarithm of domain size) is approximately linear for (a, b) niche axis systems of size *S* = 30, 50. A similar approximate log-linear relationship between biomass and domain size appears in a random matrix model of size *S* = 100 with 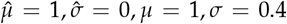 (parameters from Eq. 7). System has ∼ 90 distinct equilibria; results are binned in biomass and averaged.

Note that even when the dependence of the domain size on the net biomass is approximately exponential, or more generally monotonic, this does not mean that typical random initial conditions will give rise to the equilibrium with greatest biomass. In particular, entropic factors may play an important role in determining the features of a typical equilibrium. For example, consider the periodic niche system of size *S* = 150 with Δ = 4 and interaction (5). Considering only the equilibria with gap sizes 5, 6, there is a single equilibrium with 30 species and all gaps 5, and a single equilibrium with 25 species and all gaps 6, but there are of order 10^6^ equilibria with 20 6’s and 6 5’s (roughly 6 * (26 choose 6)), and of order 10^8^ equilibria with 15 6’s and 12 5’s. While of these equilibria with only 5’s and 6’s, the equilibrium with largest biomass is the one with all 5’s, a typical run gives one of the equilibria with a mixture of 5’s and 6’s and 27 or so species since there are so many such equilibria. This helps explain how the biomass/probability relationship on individual equilibria can be compatible with the eigenvalue discussion around Figure 4, which indicates that a “typical” equilibrium should have an intermediate number of species.

In summary, discrete niche axis models provide a class of systems with a large number of stable equilibria. The continuum and statistical mechanical limits describe regimes that can be clearly treated analytically, demonstrating a strong relationship between equilibrium biomass and domain of attraction size. Finite size systems have features that approximate the expectations from both the continuum and statistical mechanical limits.

#### Random matrix systems

While a general random matrix system does not have the locality structure of the niche axis models, there is still some relationship between biomass and domain size. An example with the approximate log-linear relationship is shown in Figure 5(c). This example illustrates that certain random matrix systems may have analogous structure to the statistical mechanical limit of the niche axis models, even though the argument for the linear relationship between biomass −*H* and log domain size ln *p* given in the Appendix does not hold in any obvious way. Further work on these systems is ongoing and will be reported elsewhere.

## Conclusions

### Summary of results and discussion

In this work we have explored the behavior of ecological systems with multiple stable equilibria, focusing in particular on systems of many species with symmetric competitive interactions distributed on a single niche axis. We have demonstrated that different equilibria may contain different numbers of species, with separation ranging from a minimal “limiting similarity” distance to twice that. For systems with finite numbers of species and discrete niches, variation in local separation can enhance the variety of available stable solutions, and these solutions are often unevenly spaced along the niche axis. In the systems we have studied, the likelihood (relative domain size) of different equilibria is generally greater for equilibria with greater biomass, and in many cases the domain size scales roughly exponentially with biomass. In the limit where the niche systems analyzed here can be described by a statistical mechanical model, we have used a version of the central limit theorem to explicitly prove this exponential relation between biomass and domain of attraction size. The same kind of exponential scaling seems to hold in random matrix systems; further results on these systems will appear elsewhere. The strong, monotonically increasing dependence of domain size on biomass suggests more generally that in competitive systems with multiple possible equilibria, the probability of a given equilibrium may be strongly affected by the biomass of the solution, or more generally by an appropriately weighted sum of populations. ^15^

The exponential dependence of the relative probability/domain of attraction size on biomass found here is superficially similar to a related observation that has been made in the literature. The biomass in the equilibrium *n*^*^ is given by *B* = −*H*(*n*^*^) in the models we have considered here; when stochastic noise is added to the system there is a distribution of states of the form *p*(*n*) ∼ *e*^−*cH*(*n*)^ (Biroli et al., 2018) Thus, in a system with noise the probability of being near an equilibrium *n*^*^ is approximated by 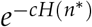. This observation, however, while similar, is logically independent of the conclusion reached here regarding domain sizes. In particular, in the stochastic version the probability distribution refers only to the likelihood of being in the immediate vicinity of a given configuration, and holds across the configuration landscape, including points well away from local equilibria. On the other hand, the domain size that is relevant in the analysis of this paper has to do with the global structure of the space of equilibria. One could certainly artificially construct a Lyapunov function where the size of the domain of attraction is smaller for wells associated with deeper equilibria, so these results cannot be related directly simply from the existence of a Lyapunov function. We are not aware of any observation in the literature of the kind of explicit relationship we have found here between the probability of a given equilibrium being reached after starting from random initial conditions and the biomass or the Lyapunov function of that equilibrium.

The relative sizes of domains of attraction we have estimated through numerical simulations are dependent to some extent on the distribution of initial conditions, so some readers may be concerned about the extent to which our conclusions depend upon the choice of initial conditions.

From the approximate geometric analysis using fixed lines of the dynamics described in the Methods section and Appendix, however, we expect that for distributions that are independent of angle within the positive orthant the results should be fairly insensitive to radial dependence. The explicit numerical test of three different initial condition distributions shown in Table 1 supports the more general conclusion that the domain sizes depend very weakly on the details of the initial distribution. Thus, we expect the conclusions found here to be relatively robust across initial conditions, although the precise details, e.g. the parameters of the statistical model in the SM limit of our niche models, will have some dependence on the specific choice of initial distribution.

### Further directions and ecological applications

There are a number of specific assumptions we have made in the analysis of this paper, and it is natural to investigate the extent to which the conclusions found here persist when these assumptions are relaxed. In particular, we have assumed that the matrix *A* is symmetric with positive off-diagonal terms, corresponding to symmetric competitive interactions. The assumption of symmetric interactions is crucial in the presence of the Lyapunov function *H*. Considering more general matrices is an important direction for future extensions of this work; we may expect, from the continuity of the equations as the interaction parameters are varied, that for matrices that are only weakly non-symmetric, the approximate monotonic relationship between biomass (or a suitable generalization thereof) and domain size may persist, but as antisymmetric components of the interactions become stronger, and systems may become chaotic, the problem becomes more complex and may involve limit cycles, etc.

There are a number of contexts in which it may be interesting to try to find empirical ecological confirmation of the results of this work. There has been some work already (Lopes et al., 2024) in which artificial laboratory ecologies are constructed using different configurations of simple organisms such as yeast types, in which multiple stable equilibria appear to arise. It would be very interesting to characterize such systems in more detail and make a connection between biomass and domain size in such systems. It would also be interesting to identify real ecological systems, such as for example tree species that differentiate niches through height or bird species that differentiate niches through target seed sizes, where different numbers of occupied niches may arise in parallel ecosystems (such as islands with similar environments).

One important additional process that could be considered here is immigration from outside a well-mixed system. It has been argued in prior work that any kind of limiting similarity solution (in the sense of isolated, well-separated species along an niche axis) is likely to be disrupted by immigration, leading to clusters of highly-similar species roughly centered at the locations where the limiting similarity species would have been, both in theory (D’Andrea and Ostling, 2016; D’Andrea et al., 2018) and in empirical data (D’andrea et al., 2020). While we have not derived solutions for models with immigration, our work implies that there may generically be multiple cluster-like solutions with different numbers of clusters. In such systems, we might expect to see qualitatively different solutions dependent on initial conditions. It would be interesting to look for real-world realizations of different niche cluster structures in similar environmental circumstances with a common species pool, such as in similar island ecosystems, although disentangling the effects from local environmental variations would be challenging. Along related lines, it would be interesting to connect the analysis of this paper with work on spatial patterns that can arise for systems where local dynamics has multiple distinct equilibria (Whittaker and Levin, 1977; Levin, 1979).

## Supporting information

Mathematica+Julia code and examples

## Acknowledgments

The authors would like to thank Peter Abrams, Rafael D’Andrea, Jeff Gore, Akshit Goyal, Christopher Klausmeier, Simon Levin, Pankaj Mehta, Nitesh Patro, Serguei Saavedra, and Yuguang Yang for helpful discussions and comments.

## Statement of Authorship

Both authors conceived the ideas and analyses of the paper. WT performed most of the computations and analyses and wrote the original draft. Both authors reviewed and edited the writing at all stages of revision.

## Data and Code Availability

Mathematica code to analyze equilibria of niche axis models and Julia code to run simulations are available as supplementary files.

## Appendix

### Derivation of geometric decomposition into cones

In this appendix we give the details of the approximate geometric decomposition of the positive orthant into basins of attraction outlined in the Methods section.

We consider symmetric systems where all *A*_*ij*_ = *A*_*ji*_, with an equilibrium configuration (stable or unstable) 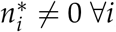, where *A* could either describe a full system or a subset of a larger system. (We are thus assuming that the system is somewhat generic and does not have a degeneracy, i.e. det *A* ≠ 0.) We first prove that the ray from the origin 0 through 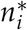 is a fixed locus of the dynamics. We assume that the dynamics is given by (1), with *r*_*i*_ = 1, so 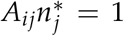. If the populations are given by 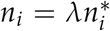, then *dn*_*i*_/*dt* = *n*_*i*_(1 − *λ*), from which it follows that the line containing all points 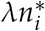 is fixed by the dynamics.

Note that this result on fixed lines of the dynamics holds formally whether or not *n*^*^ is in the positive orthant. For simplicity of exposition, in the following analysis we assume that the equilibrium (stable or unstable) for every subset 𝒮^′^ ⊆ 𝒮 of our full system (including that of the system itself) lies in the positive orthant 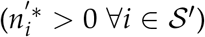 These conditions hold only for rather special matrices, including for example the monodominant models with *a*_*i*_ ∼ 1. ^16^ A similar analysis is possible when this condition does not hold; we comment briefly on how the more general analysis differs in a footnote below.

Given the above result, we can divide the positive orthant into *S*! cones spanned by the fixed lines associated with the vectors *n*^’*^ for each of the *S*! sequences of subsystems 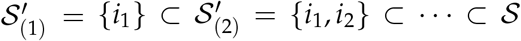. ^17^ It is straightforward to prove that for a system with all *A*_*ij*_ = *A*_*ji*_, each of the *S*! cones spanned by the fixed lines associated with the vectors 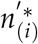 for each of the *S*! sequences of subsystems 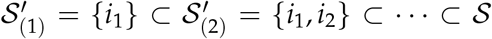 contains exactly one equilibrium, associated with some subset 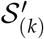, that minimizes the Lyapunov function *H* from (3) across the cone. This can be seen as follows: Consider the values *v*_(*k*)_ of the Lyapunov function for the equilibria associated with subsystems 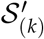. Among these values, there must be a unique minimum *v*_(*m*)_, since the Lyapunov function is quadratic and thus monotonic between any pair of 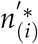 (we use here again the assumption that there are no degeneracies). For each cone, we can then make a simple approximation that the Lyapunov function flows toward the associated minimum *v*_(*k*)_. For many simple systems 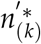 for each cone will be a locally stable equilibrium in the positive orthant, associated with a local minimum *v*_(*k*)_ of the Lyapunov function, and thus a fixed point of the system. This holds, for example, in the monodominant models, since the minimum in each cone occurs at *v*_(1)_ for a single species subset, and this will be the minimum value in every pair of adjacent cones (i.e., cones intersecting on an (*S* − 1)-dimensional hyperplane that contains 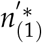. More generally, the dynamics may flow from one cone to an adjacent cone; if cones *C, D* are adjacent and both contain 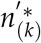, which has the Lyapunov minimum for cone *C* but not for cone *D*, then cone *C* can be associated with the equilibrium of *D* extremizing the Lyapunov function in that cone, etc. ^18^ A full analysis of this geometry becomes exponentially complicated as the number of species increases, since the number of cones goes as *S*! (see Deng et al. (2022)(S4) for a geometric analysis of cones in high-dimensional LV systems in a complementary context).

For many simple systems, including the monodominant models considered here, we can analyze the system in this way and use the *geometric approximation* that all points in each specific cone are in the basin of attraction of a particular associated stable equilibrium, so that the total positive orthant is approximately divided into *S*! cones, each of which contributes to a single basin. Because the higher-dimensional hyperplane boundaries between the cones are not invariant under the dynamics, unlike the lines through the equilibria, however, these hyperplanes do not lie precisely on the domain boundaries, so this approximation is inexact.

### Detailed analysis of monodominant systems

We begin with the two-species monodominant system with diagonal interaction coefficients *a*_1_ = *a, a*_2_ = 1. Several features of this system are worth noting: First, as *a* → 2, the unstable equilibrium approaches (1/2, 0), and the full positive population space lies in the basin of attraction of the stable equilibrium with populations (0, 1). Second, as *a* → 0, the unstable equilibrium approaches (1/4, 1/2) and 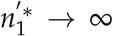. Third when 0 ≤ *a <* 4, the matrix (9) has one positive eigenvalue and one negative eigenvalue; the negative eigenvalue is associated with the unstable direction. Fourth, we can explicitly compute the relative sizes of the basins of attraction of the stable solutions. Given any distribution of initial states that is independent of angle (such as a Gaussian 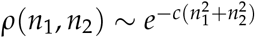, restricted to the positive quadrant), these are given by

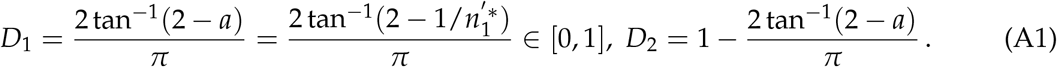

Graphing *D*_1_ as a function of 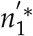 (Figure 2(b)), we see that the domain size is monotonic in the equilibrium population. The domain size vanishes below the threshold 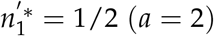, above which it grows linearly for 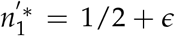, approaching a finite limit of 2 tan^−1^(2)/*π* ≅ 0.705. Allowing the other diagonal element *b* = *a*_22_ of (9) to also vary gives qualitatively similar results, with the unstable equilibrium at (2 − *b*, 2 − *a*)/(4 − *ab*).

For a three-species monodominant model, with the interaction matrix *a*_*ij*_ = 2, *i* ≠ *j* and *a*_*i*_ ≡ *a*_*ii*_, there are generally three stable equilibria with 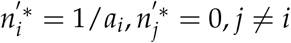. Restricting again to the case where *a*_1_ = *a, a*_2_ = *a*_3_ = 1, the domain size *D*_1_ as a function of population 1/*a* has similar properties to the two-species system: *D*_1_(1/2) = 0, a finite limit 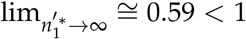, and quadratic growth near 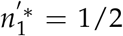 (see Figure 2(c)). To illustrate the relationship between stable equilibria and eigenvectors of the *A* matrix with negative eigenvalues, we consider the case *a*_1_ = 0.75, *a*_2_ = 1, *a*_3_ = 1.25. In this case the most negative eigenvalue of the *A* matrix is *λ* ≅ −1.15, associated with the eigenvector (3.89, −2.69, −1), corresponding to positive fluctuations in the direction of the first species, which has the largest biomass 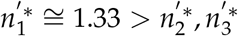.

For larger systems, the relationship between domain size and population continues to be monotonic. Figure 2(d) shows the distribution of relations between domain sizes *D*_*i*_ and population 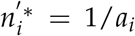 for a set of matrices with random *a*_*i*_ chosen from a Gaussian distribution with several variances (in the case of *σ* = 0, *a*_1_ is fixed and separate simulations are run for each value), and Figure 2(e) shows the distribution of this relation for a fixed matrix with *a*_*i*_ taking values in a uniform discrete distribution in the range [0.5, 1.9]. In both cases the number of species is *S* = 20, and statistics are taken over 10^5^ simulations. It is worth noting that for the fixed matrix with uniformly distributed *a*_*i*_, there is a close match between the eigenvalues and eigenvectors of *A* and the equilibria with largest populations, generalizing the discussion of monodominant models in the main text and the preceding *S* = 3 example; taking *a*_1_ to be the smallest value 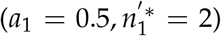, the eigenvector associated with the most negative eigenvalue is of the form (+, −, −, … −), the eigenvector with the next most negative eigenvalue is of the form (−, +, −, −, …, −), etc.

### Derivation of various aspects of continuum limit

We consider the equally spaced equilibrium solution of the periodic niche system discussed in the results section, with spikes at equal distances *D* and 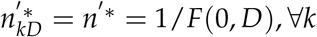, with all other 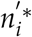 vanishing. This solution is unstable to invasion by a species at *D*/2 when *F*(*D*/2, *D*)*n* ^′ *^ *<* 1. For the interaction function (6), which is highly local, we can approximate this by only considering the neighboring spikes (at *i* = 0, *D*), and this condition is satisfied approximately when

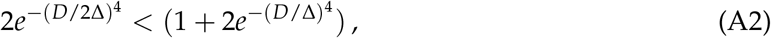

or *D* ≳: 1.8249 Δ. Corrections from the additional spikes only affect this negligibly since the interaction drops off rapidly 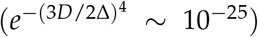 (and indeed at the boundary value of *D, n* ^′ *^ ≅ 1.0 to high precision, so the second term in parentheses in (A2) can be dropped). On the other hand, the periodic solution is unstable against fluctuations of alternating signs when

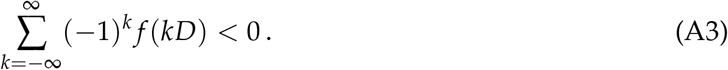

Considering again only nearby spikes (*k* = 0, ±1), for the interaction (6) this occurs when 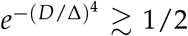, or *D*, :≲ 0.9124Δ, with negligible corrections from including further spikes. We thus expect stable solutions in the range 0.91, :≲ *D*/Δ, :≲ 1.82.

The biomass density for a periodic solution with spikes at distance *D* is given by

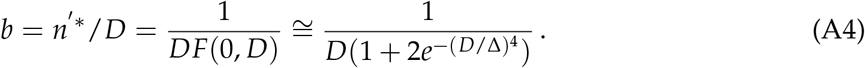

The associated eigenvalue of *A* for a fluctuation with period *D* is given by

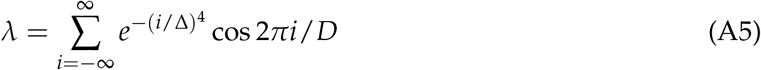

In the continuum limit this becomes

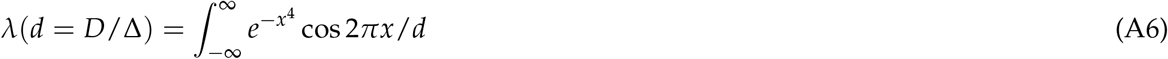

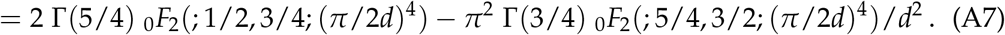

The biomass density of stable periodic solutions and eigenvalue for fluctuations with the associated periodicity around the unstable equilibrium are graphed in Figure A1, illustrating a close (but not precise) correspondence between more negative eigenvalues and greater biomass for corresponding equilibria.

**Figure A1:**
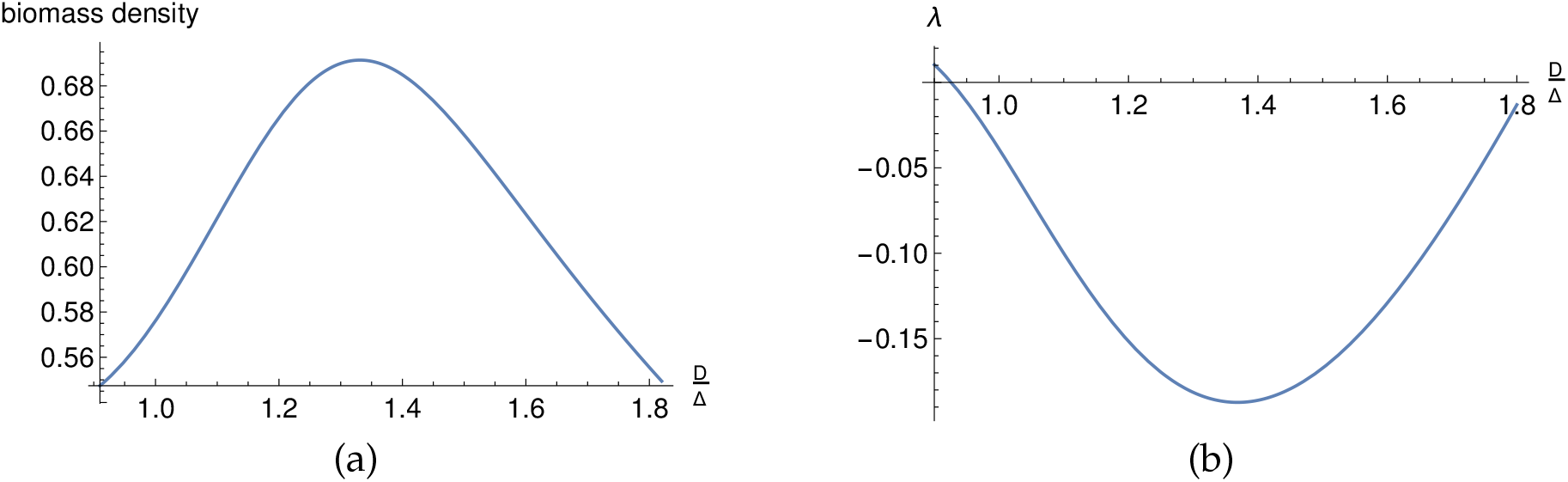
(a) Biomass density for periodic solutions and (b) eigenvalue for fluctuations with corresponding periodicity around stable equilibrium graphed as functions of *D*/Δ. Functions have similar form (with opposite sign) but not exactly identical.

### Markov chain model in the statistical mechanical limit

We describe here in more detail the derivation of the distribution of equilibria and the associated relationship between biomass and domain size using the Markov chain central limit theorem.

It is important to emphasize that there are two different distributions that may be considered relevant when considering the equilibria of the niche models in the statistical mechanical limit. The first distribution is one in which each equilibrium *n*^*^ is sampled with a probability *p*(*n*^*^) given by the likelihood of reaching that equilibrium when initial conditions are chosen from a fixed distribution. The second is one in which we simply look at the ensemble of possible equilibria, taking each distinct equilibrium with unit weight, and look at the distribution of properties such as biomass *B*(*n*^*^) and log probability − ln *p*(*n*^*^) over the set of equilibria.

In this section we focus on the first of these distributions, in which equilibria are sampled with a probability proportional to the (suitably defined) domain size. This is the context in which the relationship between biomass and domain size can be characterized most cleanly. The following section describes the second distribution, in which all equilibria are considered with equal weights. In the final section of the Appendix, we go through an explicit example of these analyses and show a close match with numerical simulations.

The distribution with sampling probability *p*(*n*^*^) is accurately described by a Markov model associated with the statistical model (8). The Markov chain central limit theorem then implies that the distribution on biomass and log probability is a multivariate Gaussian, so that there is statistically a linear relationship between these two quantities. The parameters of the statistical model (8) can be determined “experimentally” by doing a large number of numerical simulations on a system of fixed size *S*. The predictions of this statistical model then can be shown to match very closely with the observed distribution of equilibria. We now describe this analysis more thoroughly; the final section of the Appendix contains further details and numerical checks in some example cases.

We can derive the relation (15) using the Markov chain central limit theorem, given certain natural assumptions about the locality of the system. We assume that at large *S* and fixed Δ, the domain sizes of the equilibria are described by the local model (8)^19^, and furthermore that the weighting factors associated with *n*-gap correlations for *n > k* with *k* a fixed integer are negligible. We also similarly assume that the biomass of the *i*th spike is approximated well as a function that depends only on the finite set of gap values *g*_*i*−*k*_, …, *g*_*i*+*k*−1_ (i.e., the positions of the nearest *k* species in each direction). We have not proven these assumptions rigorously but they are physically plausible given the local nature of the interactions, and appear to be satisfied to a high degree of precision in all systems we have considered.

Given these assumptions, we proceed by first considering sequences of gaps described by the distribution (8), without taking periodicity into account. ^20^ As described below, the periodicity condition only modifies things by a finite shift that becomes irrelevant in the large *S* limit. In the *k*-gap statistical mechanical model, in which we drop weighting factors for more than *k* consecutive gaps, the probability of a given finite gap sequence *p*(*g*_1_, …, *g*_*n*_) with *n* ≫ *k* is approximated by a Markov-type model where *p*(*g*_1_, …, *g*_*j*_) is computed iteratively through an equation of the form

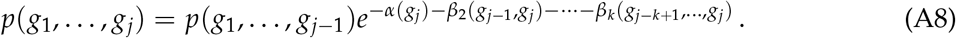

(The initial conditions on the iteration are that 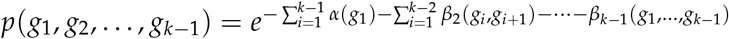

To describe this as a Markov model where the probability of the next state depends only on that of the single previous state, we can consider the sequences of symbols (*g*_1_, … *g*_*k*−1_) as a single index *I*, so that the probability of following a state *I* = (*g*_1_, …, *g*_*k*−1_) with a state *J* = (*g*_2_, …, *g*_*k*_) is given by

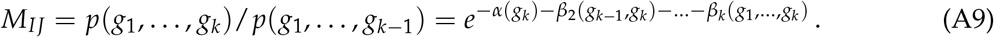

More generally, we write *I*_*i*_ = (*g*_*i*_, …, *g*_*i*+*k*−1_) and the probability distribution on a linear sequence of gaps can be described by the Markov chain

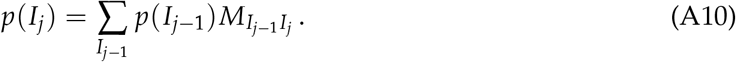

The Markov chain central limit theorem then states that the distribution on any linear function 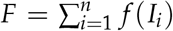 approaches a normal distribution as *n* → ∞. This result naturally generalizes to any local function *f* (*I*_*i*_, …, *I*_*i*+*m*_) for fixed finite *m*. We have assumed that the biomass *B* = −*H*(*n*^*^) is such a local function, as is ln *p* from (8). Indeed, in the Markov model the probability of a sequence *I*_1_, …, *I*_*G*_ is given by 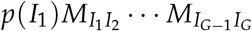, so

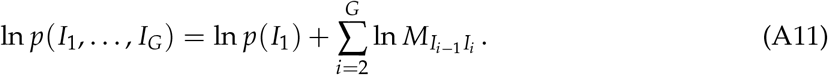

The multivariate version of the Markov chain central limit theorem thus asserts that the set of equilibria with a fixed number of gaps *G* in the large *G*, fixed Δ limit gives a bivariate normal distribution in the variables (−*H*, ln *p*). Note that this normal distribution is reached when the equilibria are sampled with the appropriate probability distribution *p*(*g*_1_, …, *g*_*k*_). As *G* → ∞, the condition of periodicity, i.e. that *I*_*G*+1_ = *I*_1_, gives a finite effect to equations such as (A11), which is negligible in the large *G* limit. Note that we distinguish here between the number of gaps *G* in a periodic sequence and the associated lattice size *S* = *g*_1_ + … + *g*_*G*_; we return to this below.

Let us now assume that we have a statistical *k*-gap model with a fixed value of *k*, and an associated matrix *M*_*IJ*_. The stationary (*k* − 1)-gap probabilities *p*(*I*) = *p*_∞_(*I*) (i.e., the local (*k* − 1)-gap probabilities in the limit of large system size) are a (left) eigenvector of this matrix with eigenvalue 1,

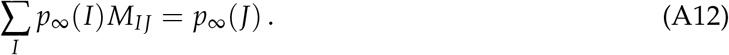

From the matrix *M* we can thus determine the mean of ln *p* per gap

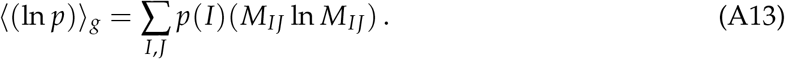

We can similarly determine the mean size of a gap

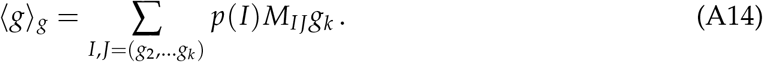

This gives an estimate ^21^ of the mean of the log probability for a periodic configuration on a lattice of size *S* = *g*_1_ + … + *g*_*G*_,

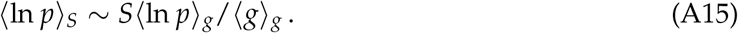

This local *k*-gap model can be extended to describe the mean of the total biomass from an estimate of the biomass at a given lattice site given the adjacent gap sizes on both sides, i.e. *b*_*i*_ ∼ *b*(*I*_*i*−*k*_, …, *I*_*i*+1_) would give the average biomass of a species at the lattice point in the middle of this gap sequence; analytically we expect

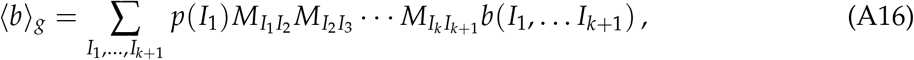

and, similar to Eq.(A15) (and with similar caveats), ⟨ *B*⟩ = ⟨ *b*⟩ _*S*_ ∼ *S*⟨ *b*⟩ _*g*_/⟨ *g*⟩ _*g*_.

Using the local data of the *k*-gap model *M*_*IJ*_, *b*(*I*_1_, …, *I*_*k*+1_), a similar analysis can also be used in principle to predict the higher moments of the log probability and biomass. Because these quantities are correlated between nearby gap positions, however, one would need to compute the two-point correlation between ln *p* and/or biomass contributions between all pairs of nearby lattice sites, and the analysis would become rather involved. Because of the local nature of the model, these correlations will only extend over a finite distance, however for higher moments this distance may be significantly larger than the *k* adequate for analyzing means. For practical purposes, it may be easiest to compute these higher moments by performing a linear fit to data from a construction of all equilibria at relatively large *S*, as described below.

Given a choice of Δ and an integer *k* defining the *k*-gap approximation to the statistical mechanical model, we have thus formulated a hypothesis for a Markov chain model that characterizes the distribution of equilibria of the system in the statistical mechanical limit. We have argued that with some fairly minimal assumptions regarding the locality of the interactions, the distribution on equilibria is a multivariate Gaussian in (*B*, − ln *p*) space. We have tested these results in a number of different circumstances, and find that this model accurately describes the empirically determined distribution of equilibria in those cases studied. Further details are described in the final Appendix.

### Statistical model for all equilibria

In this section we consider the second distribution of interest on the set of equilibria, which contains the set of all equilibria, considered with equal weights. This may seem in some ways to be a natural distribution to consider, however there are some subtle issues regarding this distribution. In particular, very low-probability equilibria may substantially affect the mean and variance of various quantities such as biomass and log probability in this distribution. ^22^ This distribution is also essentially difficult or impossible to experimentally determine precisely and with certainty for large systems with a finite number of samples, as it is hard to to be sure whether one has all the equilibria or some low-probability equilibria are missing. Nonetheless, for this distribution we can still make some useful observations. Given a local statistical model associated with the first distribution above, a modification of this model can give an associated model of the second distribution with uniform weighting factors. This allows us to exactly construct and analyze the second distribution in the approximation of the given statistical model, with the caveat that for computing the mean and variance of quantities of interest, predictions may be off due to low-probability equilibria that are not captured by the given statistical model.

Associated with the *k*-gap Markov chain model described in the preceding section, which generates equilibria with a probability distribution *p*(*g*_1_, … *g*_*p*_), we can use a related model to classify all allowed equilibria *g*_1_, … *g*_*p*_. Associated with the matrix *M*_*IJ*_ characterizing the *k*-gap Markov model, we can define a matrix with 0/1 entries through

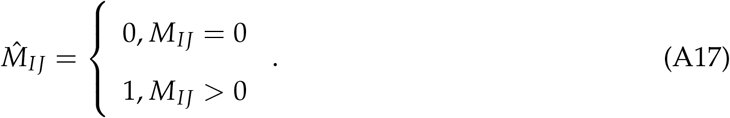

The set of all allowed sequences *g*_1_, …, *g*_*p*_ is then the set of sequences *I*_*k*−1_, …, *I*_*p*_ such that

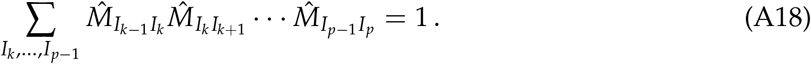

This construction can be used in several ways. First, it can be used to construct an approximation of the complete set of equilibria for the lattice niche models with fixed Δ at large *S*. This extends the construction of equilibria beyond that which is practical from sampling all possible configurations. Note, however, that while we can explicitly check to confirm that any equilibrium suggested by this construction is indeed a stable equilibrium of the lattice model, it is difficult or impossible to be sure that this approximation does not miss very low-likelihood equilibria involving correlations of more than *k* gaps. The number of equilibria given by this model will scale as 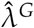, where 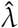 is the largest (real) eigenvalue of 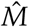. More precisely, the number of equilibria given by this model is just 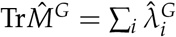, where 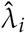 are all the eigenvalues of 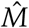.

In the approximation that the *k*-gap model is accurate, this can also give us a distribution on observables such as log probability and biomass sampled over all equilibria counted with equal weights (i.e. the second distribution of interest). We briefly summarize this analysis in the case of log probability.

To determine the distribution over all equally weighted equilibria, we must identify the probability of a given subsequence 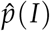 or 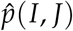 when this subsequence is considered in the context of a larger sequence extending in both directions. The number of sequences in which *I* is preceded by *n* arbitrary gaps and followed by *m* arbitrary gaps satisfying the constraint (A18) is

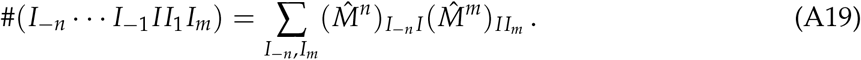

If we denote the left- and right-eigenvectors of 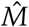 associated with the maximum real eigenvalue 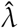 by 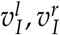, then the desired stationary subsequence probabilities are given by

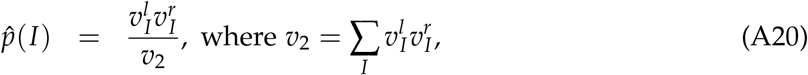

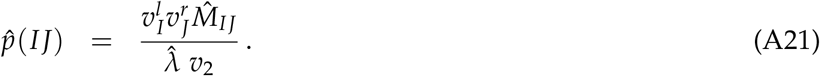

Just as knowledge of the probabilities *p*(*I*), *p*(*IJ*) allowed us to define a Markov chain model with transition matrix *M*_*IJ*_ = *p*(*IJ*)/*p*(*I*) in the previous section, the probabilities (A20, A21) allow us to define a separate Markov chain model generating sequences with the probability distribution associated with the full set of equilibria, through the transition matrix

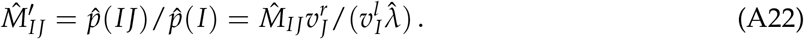

This allows us to derive quantities such as the mean log probability and mean gap size analogous to (A13, A14), but now with unit weighting for all solutions of the *k*-gap model:

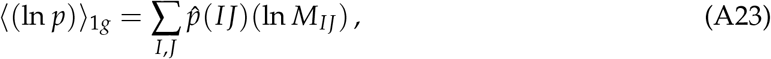

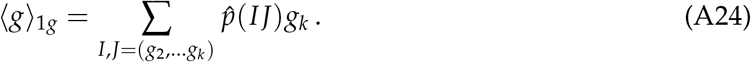

Note that the exponential dependence of the number of sequences on the number of gaps *G* may lead to effects such as an enhanced correlation between log probability and gap size; we give an example of this in the following section.

In the approximation that the *k*-gap model is a correct description of the full set of equilibria, it follows from this analysis that we expect a multivariate normal distribution for quantities such as log probability and biomass for the full set of equilibria of the niche lattice models with fixed Δ as *S* → ∞. As mentioned earlier, however, this model is somewhat less robust as it may be insensitive to very small-probability equilibria that are missed in making the *k*-gap model approximation, and the large tail on the distribution may make predictions from this model less precise than those from the probability-based distribution.

### Details of statistical mechanical model and matching with numerical results

In this final appendix we describe how numerical data from simulation can be used to fix a *k*-gap local statistical mechanical model, and confirm first that this local statistical mechanical model gives a distribution of equilibria with the expected multivariate Gaussian form in (*B*, − ln *p*) space, and second that the distribution from the statistical mechanical model matches the distribution of equilibria found in the simulations.

By performing numerical experiments in which the dynamical equations (1) are solved with random initial conditions, we can numerically determine the finite set of weighting factors in the *k*-gap local statistical mechanical model (8), which allows us to determine the matrix *M*_*IJ*_ for any specific parameter set. In particular, based on empirically measured *k*-gap statistics *p*_*s*_(*g*_1_, …, *g*_*j*_), *j* ≤ *k* computed by compiling all *k*-gap sequences encountered in the equilibria endpoints of a large number of simulations, we can estimate the parameters *α*(*g*), *β*_2_(*g, g*^′^), … of an associated *k*-gap statistical mechanical model, using the relations

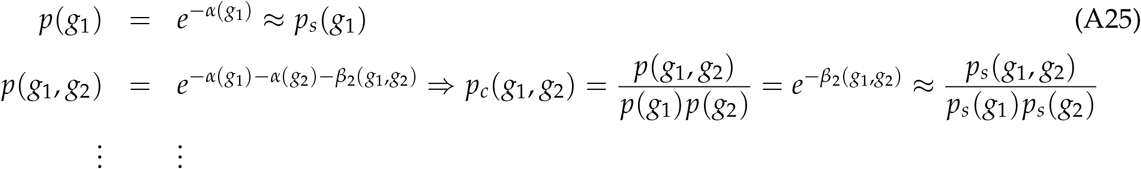

We consider the case Δ = 4 as an example for more detailed analysis; the statistical mechanical models based on other values of Δ should be qualitatively similar. Numerical analysis for Δ = 4 at small *k*, shows rapid convergence both as *S* → ∞ and *k* increases (see Figure A2(a, b)); *k* = 4 at *S* = 90 is fairly accurate. To build a *k* = 4 statistical mechanical model we have used the distribution based on 10^6^ runs at size *S* = 90. This gives, for example, *p*(5, 5, 5) ≈ 0.221 as the most likely 3-gap sequence, as expected from (12).

Using this model of probabilities we can construct the associated Markov model (A8). In particular, we can compute the matrices 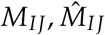 given by (A9, A17) for the 4-gap model. For example, the transition probabilities from the state *I* = (555) are approximately

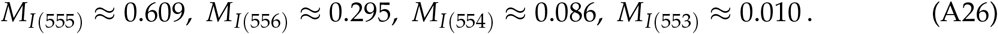

(Note that, for example, the 5553 sequence does not appear in Table 1 of the 281 equilibria for the *S* = 30, Δ = 4 model; this sequence appears for other values of *S*.)

Using the matrix *M*, we can generate random sequences of gaps with a probability distribution given by the associated Markov model, and we can compute the mean of ln *p* and gap size (per gap)

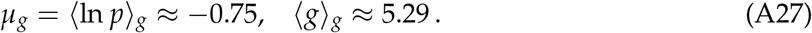

Using the 0/1 transition matrix 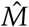, we can construct all allowed sequences of gaps of a given length *G*, and compute the expected means

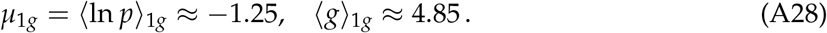

While the results of the Markov chain central limit theorem only apply directly to non-periodic sequences of a fixed number of gaps *G* (or fixed size *S*, in the bitwise version), as discussed earlier we expect that the condition of periodicity will simply cut out a fraction of possible sequences that approaches a constant fraction as *S* → ∞, and we expect that in the large *S* limit the standard deviation in *G* will go as 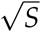 so variations in *G* for a given *S* will be negligible, at least for the *M* Markov model associated with the distribution of equilibria with probability *p*.

As an initial check on this *k* = 4 model, we have used 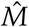 to produce all possible periodic sequences up to length *S* = 90. The number of such sequences grows exponentially, as expected (Figure A2(c)), roughly as 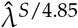, where 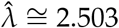. Furthermore, this model precisely reproduces the 281 possible periodic sequences at size *S* = 30. At *S* = 60 the model generates 73,525 distinct sequences associated with stable equilibria, and at *S* = 90 it gives 19.5 million such distinct sequences.

We have next checked to confirm that the equilibria produced by the local statistical mechanical model give a distribution on probabilities that matches the expectation from the Markov chain central limit theorem. For each value of *S*, we can compute the mean of the log probability for all of the equilibrium sequences generated from 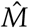, both with the probability-based measure and the unit measure. The results of this computation are shown in Figure A3, and compared to the predictions of (A27, A28). Note that the slight deviation between the red points associated with the mean of the log probability for the equilibria with unit measure and the theoretical prediction from (A28) is likely due to a correlation between the number of gaps and ln *p*; unlike the probabilistic measure, the number of configurations grows exponentially as the number of gaps increases, so the mean gap length for fixed *S* will be slightly smaller than predicted from (A28), pushing the red data points down slightly. We have not tried to analytically compute this effect, which is small. This figure also shows that the variance of log probability (using the probability-based measure) grows linearly at large *S*, and a linear fit gives *σ* ≈ 0.048*S*. A similar analysis can give a theoretical prediction of the multivariate normal distribution when biomass is included purely from the statistical mechanical model, although we have not done that explicitly here.

Finally, we have compared the predictions of the statistical mechanical model to the observed distribution of equilibria in numerical simulations. For those equilibria observed in 10^6^ simulations of the *S* = 60 model, the observed probability is compared to the probability predicted from the *k* = 4 statistical mechanical model in Figure A4. The distribution in (*B*, ln *p*) space of stable solutions observed in numerical simulations is compared with the complete set of equilibria for the *k* = 4 statistical mechanical model in Figure A5. These comparisons show that the statistical mechanical model accurately reproduces the distribution of equilibria and that both the statistical mechanical model and the observed distribution have the expected linear correlation between ln *p* and biomass, which follows from the multivariate Gaussian form of the equilibrium distribution.

**Figure A2:**
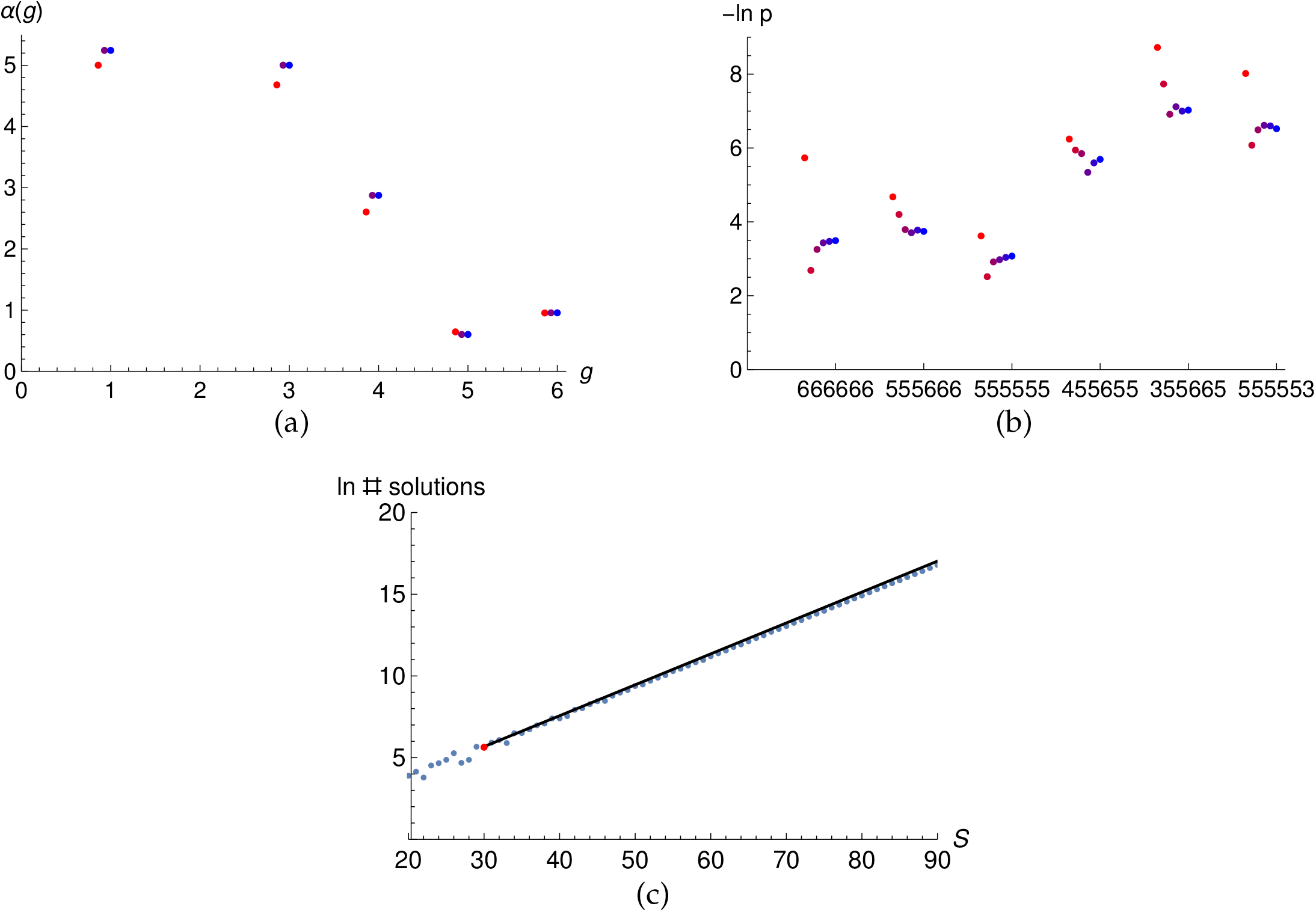
(a) Convergence of *α*(*g*) in statistical mechanical model as *S* → ∞ based on simulations with *S* = 30 (red), *S* = 60 (purple), *S* = 90 (blue). (red and purple points slightly offset horizontally for readability.) (b) Convergence of predicted values for − ln *p*(*g*_1_, …, *g*_6_) for selected sequences in models from *k* = 1 (red) to *k* = 6 (blue). (c) Log of the number of solutions generated by the *k* = 4 statistical mechanical model at lattice size *S*; black line is theoretical prediction 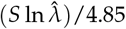, red dot is *S* = 30, with 281 solutions.

**Figure A3:**
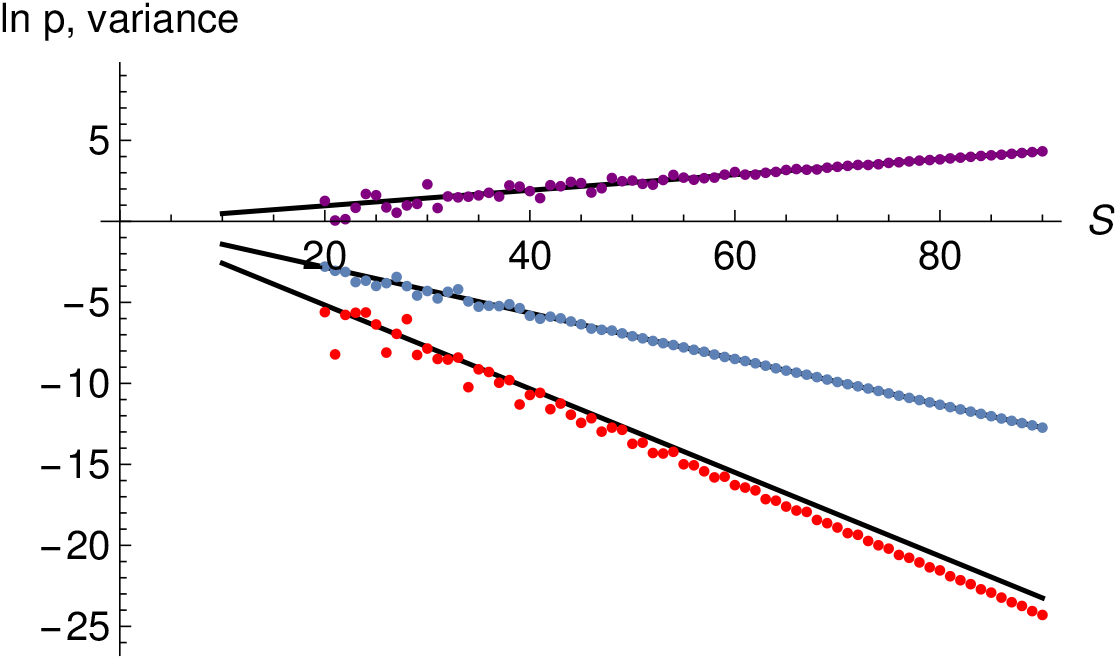
Linear growth of mean and variance of log probability for all periodic sequences of size *S* ≤ 90 in *k* = 4 statistical mechanical model illustrates validity of Markov chain central limit theorem for periodic sequences in this model. Blue points gives means using probability measure, red points use unit measure. Black curves along blue and red are predictions from (A27) and (A28) respectively. Purple points are variances of log probability using probability measure, black line is a linear fit to the values at large *S*.

**Figure A4:**
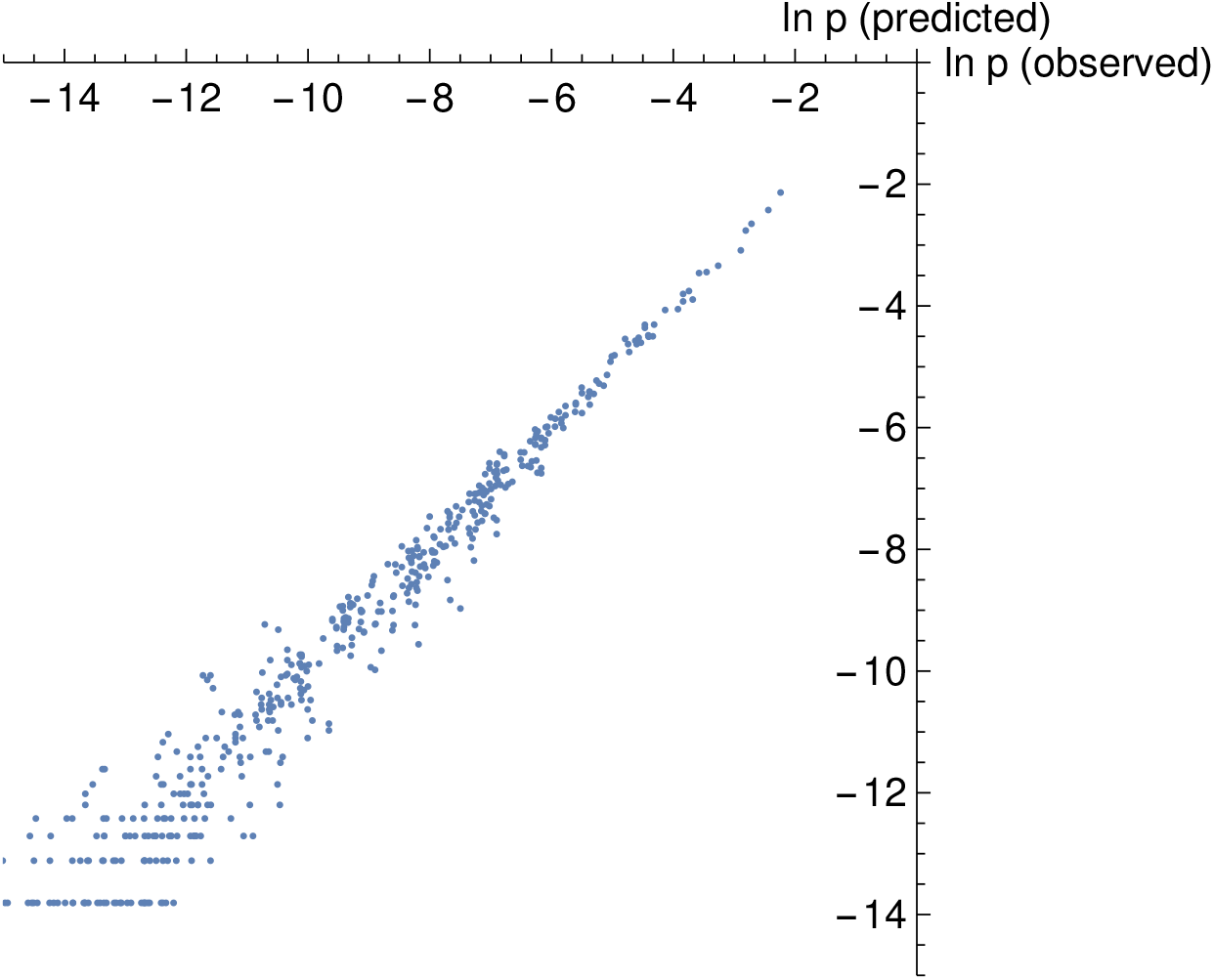
Probabilities of configurations predicted by statistical mechanical model at *k* = 4 vs. observed probabilities in numerical niche model solutions, at size *S* = 60.

**Figure A5:**
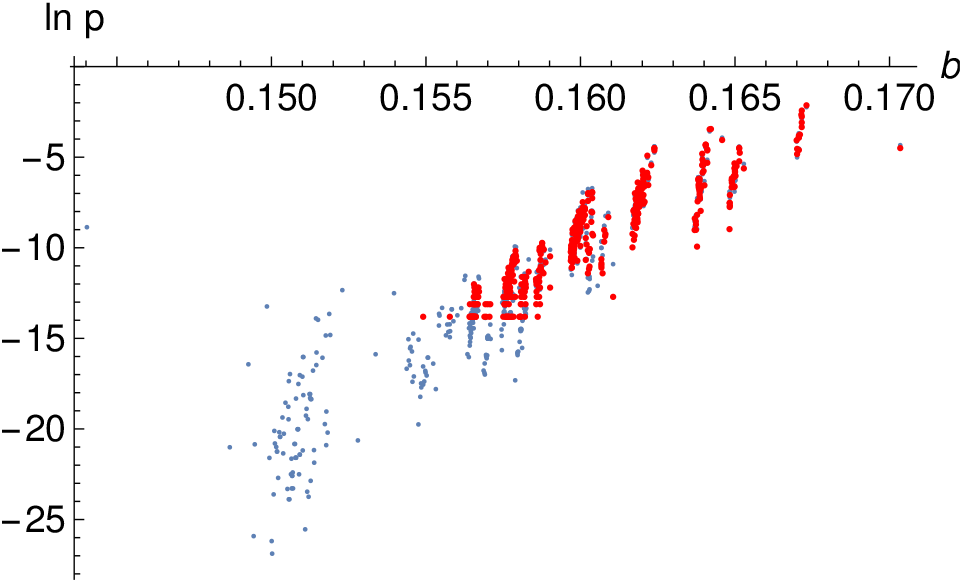
Distribution of equilibria for the *k* = 4, *S* = 60 model in (biomass, ln *p*) space. Red points are those found in 10^6^ numerical simulations; blue points are those realized using the local *k* = 4 statistical mechanical model and the associated 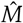 matrix to produce equilibria, for which biomass is computed exactly and ln *p* is estimated using statistical mechanical model.

## Footnotes

1 Note: for most of the systems studied in this paper 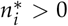 for all *i*; (sub)systems with equilibria that violate this condition and lie outside the positive orthant are discussed in the first section of the Appendix.

2 Note that in these simple models the individuals of each species are assumed to have the same biomass, so biomass is simply the sum of all populations. In more general systems, for example with distinct *r*_*i*_, we expect that the role of biomass will be played by a more general measure associated with some weighted sum of populations.

3 Physically, this is clear from the form of the Lyapunov function since the equilibria lie at local minima. Mathe matically, an instability is generally associated with a positive eigenvalue of the Jacobian matrix, which in this case at the equilibrium is 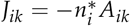 as long as all 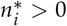 and *A* is symmetric and nondegenerate, the signature (numbers of positive and negative eigenvalues) of *J* and −*A* must be the same — this simply follows from continuity since det 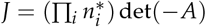 Note that this conclusion can change when *A* is not symmetric, see, e.g., Gibbs et al. (2018), although there are also classes of non-symmetric matrices that may be of ecological interest where the eigenvalues are real and this conclusion still holds (O’Dwyer, 2020). Note also that in much of the ecological literature for interactions that are not purely competitive, the opposite sign convention is often chosen for *A*.

4 By periodic boundary conditions we mean that we consider the species at positions 1 and *S* to be adjacent; this can be thought of on the one hand in terms of species on an infinite lattice where species *k* and *k* + *S* are identified for each *k*, or equivalently as placing the species around the perimeter of a circle.

5 For our regular discrete models, this occurs for any function where 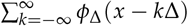 has a local minimum at *x* = Δ/2 and a local maximum at *x* = Δ for any Δ (in which case a set of species regularly spaced at intervals Δ can be invaded by intermediate species); one can check, e.g., numerically that this is true for 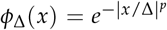 for *p* ≤ 2 but not for *p >* 2.

6 As stated in the Methods section, by a stable equilibrium here we mean one that is both locally stable within the subsystem defined by the set of nonzero species and also stable to invasion from other species in the broader pool.

7 This initial distribution has the feature that the dynamics of every simulation must stay within the initial box for systems with purely competitive interactions; thanks to Jeff Gore for pointing this out.

8 Note that we have also rescaled *ñ*(*x*) ∼ Δ*n*(*i*).

10 These assertions are easy to verify empirically by explicitly checking for systems with relatively small *S* (e.g. *S* ≤ 36). In particular, in general the sequence 5656 is unstable to the sequence 5566, and gaps of size greater than 7 are unstable to invasion; gaps of size 7 primarily arise in uniform sequences of the form 77… 7 when *S* = 7*k*, although for example the sequence 161676 is a similar stable solution for a system of size 27. These constraints can also be proven analytically with some assumptions on the boundary conditions, though the details are not particularly illuminating and are left as an exercise for the reader.

11 Some subtleties in this correspondence related to boundary conditions and finite size effects are discussed in the Appendix.

12 Note in particular that no gaps of size 7 have been found in systems of size *S* = 90, so this model does not include, e.g., gap sequences with all 7’s that are possible when *S* is a multiple of 7.

13 The Shannon entropy of a probability distribution on sequences of symbols from a fixed alphabet characterizes the (logarithm of the) randomness of (i.e. number of bits needed on average to encode) a typical sample from the given distribution.

14 In this and related discussions we drop the prime in *n*^′ *^ for stable equilibria to simplify notation.

15 Note that, for all the models we consider here, we are considering species with identical biomass and growth rate per individual, so biomass here just means total population; more generally the Lyapunov function in a given equilibrium gives population weighted by growth rate, 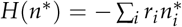, so in more general systems with symmetric interactions we may expect this quantity to govern the domain sizes.

16 Note for example that this does not hold even for the circulant niche interaction matrices; for example, for the *S* = 10, Δ = 4 system the subset 𝒮^′^ = {1, 2, 6, 9} has an equilibrium outside the positive orthant.

17 When some subset equilibria lie outside the positive orthant, there is an additional constraint that we only consider the intersection of each cone with the positive orthant; this intersection will be empty for some cones. Otherwise, the analysis in the general case is identical to the story in the text.

18 To illustrate the flow between adjacent cones, consider the simple example of the 3-species interaction matrix 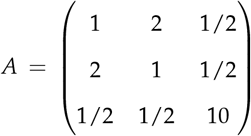. In this case the stable subsystems are {1, 3} and {2, 3}. The cone {*i*_1_, *i*_2_, *i*_3_} = {1, 2, 3} does not contain any of the stable fixed points, and the minimum in this cone is associated with the subset {1}, so this cone is associated with the stable subsystem {1, 3}, while the {2, 1, 3} cone flows to the stable subsystem {2, 3}. Thus, in this example the six cones in the positive orthant divide into two sets of three associated with the domains of attraction of the two stable fixed points. In this case the geometric approximation is exact; since there is essentially a neutral model between species 1 and species 2, the final equilibrium will be determined simply by which species has the greater population in the initial condition, with species 3 coming along for the ride.

19 As described elsewhere, the precise definition of the domain sizes depends upon the distribution of initial conditions.

20 Note that the analysis here could also be carried out directly on the sequence of bits associated with the presence or absence of a population at each niche site. This would give a similar Markov model and corresponding statistical mechanical model on bit strings. In that framework, the number of bits included in the Markov window would be on the order of *gk*, where a typical gap is of size *g*. This form of the analysis similarly leads directly to the conclusion that the set of equilibria gives a bivariate normal distribution in the variables (*B* = −*H*, ln *p*), in the limit of large systems. This is in some sense a more direct approach, and avoids any concerns about higher-order correlations between gap size and probability. Practically, however, the gap-based analysis is simpler in some ways to implement, and ties in more naturally to the other analyses in the paper, so we have focused on that here. The bit-sequence based analysis would proceed similarly, with parallel notation, replacing gaps *g*_*i*_ with bits *b*_*a*_ indicating the presence or absence of a species with nonzero population at lattice site *a*.

21 Note that this estimate may be subject to corrections in situations where there are strong correlations between *g* and ln *p*. As mentioned in the footnote above, no such corrections are needed in the alternative bitwise description of the statistical model; in the cases we have considered explicitly, such corrections are relatively small (see further discussion below).

22 For example, consider a distribution in which the number *k* is chosen with probability *p*_*k*_ = 10^−*k*^, *k* ≥ 1, and in the remainder of cases the number *k* = 0 is chosen. The entropy of this distribution −(ln *p*) = − Σ_*k*_ *p*_*k*_ ln *p*_*k*_ is a finite number, dominated by the most likely small values of *k*. On the other hand, if we sample the *k*’s with equal weights up to some cutoff *K*, then the expectation value of ln *p* diverges, −⟨ ln *p*⟩ ∼ *K*/2 → ∞, dominated by increasingly large values of *k* associated with outcomes that were exponentially suppressed in the original distribution.

