## Supplementary material for "On the structure of multiple stable equilibria in competitive ecological systems": Mathematica+Julia code and examples: 90-sorted.m

```
distribution[90] =
{{9529, {5, 5, 5, 5, 5, 5, 6, 6, 6, 6, 6, 6, 6, 6, 6, 6}}, 
 {9003, {5, 5, 5, 5, 5, 5, 5, 6, 6, 5, 5, 5, 5, 5, 6, 6, 6}}, 
 {8857, {5, 5, 5, 5, 6, 6, 6, 6, 5, 5, 6, 6, 6, 6, 6, 6}}, 
 {8612, {5, 5, 5, 5, 5, 5, 5, 5, 6, 6, 5, 5, 5, 5, 6, 6, 6}}, 
 {8571, {5, 5, 5, 5, 6, 6, 6, 5, 5, 6, 6, 6, 6, 6, 6, 6}}, 
 {8289, {5, 5, 5, 5, 5, 5, 5, 5, 5, 5, 5, 5, 6, 6, 6, 6, 6}}, 
 {7607, {5, 5, 5, 5, 5, 5, 5, 5, 5, 6, 6, 5, 5, 5, 6, 6, 6}}, 
 {7551, {5, 5, 5, 5, 6, 6, 5, 5, 6, 6, 6, 6, 6, 6, 6, 6}}, 
 {7144, {5, 5, 5, 5, 5, 6, 6, 6, 6, 5, 6, 6, 6, 6, 6, 6}}, 
 {6832, {5, 5, 5, 5, 5, 6, 6, 6, 5, 6, 6, 6, 6, 6, 6, 6}}, 
 {6523, {5, 5, 5, 5, 5, 5, 5, 5, 6, 5, 5, 5, 5, 6, 6, 6, 6}}, 
 {6493, {5, 5, 5, 5, 5, 5, 5, 6, 5, 5, 5, 5, 5, 6, 6, 6, 6}}, 
 {6325, {5, 5, 5, 5, 5, 5, 5, 5, 5, 5, 6, 6, 5, 5, 6, 6, 6}}, 
 {5814, {5, 5, 5, 5, 5, 6, 6, 5, 6, 6, 6, 6, 6, 6, 6, 6}}, 
 {5721, {5, 5, 5, 5, 5, 5, 5, 5, 5, 6, 5, 5, 5, 6, 6, 6, 6}}, 
 {5458, {5, 5, 5, 6, 6, 6, 6, 5, 5, 5, 6, 6, 6, 6, 6, 6}}, 
 {5402, {5, 5, 5, 5, 6, 5, 5, 6, 6, 6, 6, 6, 6, 6, 6, 6}}, 
 {5079, {5, 5, 5, 6, 6, 6, 5, 5, 5, 6, 6, 6, 6, 6, 6, 6}}, 
 {4700, {5, 5, 5, 5, 5, 5, 5, 5, 5, 5, 5, 6, 6, 5, 6, 6, 6}}, 
 {4487, {5, 5, 5, 5, 5, 5, 5, 5, 5, 5, 6, 5, 5, 6, 6, 6, 6}}, 
 {4405, {5, 5, 5, 5, 6, 6, 6, 6, 6, 5, 5, 6, 6, 6, 6, 6}}, 
 {4374, {5, 5, 5, 5, 5, 5, 6, 6, 5, 5, 5, 5, 5, 5, 6, 6, 6}}, 
 {4222, {5, 5, 5, 6, 6, 5, 5, 5, 6, 6, 6, 6, 6, 6, 6, 6}}, 
 {3873, {4, 5, 5, 5, 5, 5, 5, 5, 5, 6, 6, 6, 6, 6, 6, 5, 5}}, 
 {3629, {5, 5, 5, 5, 5, 6, 6, 6, 6, 6, 5, 6, 6, 6, 6, 6}}, 
 {3298, {5, 5, 5, 5, 5, 5, 6, 5, 5, 5, 5, 5, 5, 6, 6, 6, 6}}, 
 {3222, {4, 5, 5, 5, 5, 5, 5, 5, 6, 6, 6, 6, 6, 6, 5, 5, 5}}, 
 {3085, {4, 5, 5, 5, 5, 5, 5, 6, 6, 6, 6, 6, 6, 5, 5, 5, 5}}, 
 {2986, {5, 5, 5, 6, 5, 5, 5, 6, 6, 6, 6, 6, 6, 6, 6, 6}}, 
 {2702, {5, 5, 5, 6, 6, 6, 6, 6, 5, 5, 5, 6, 6, 6, 6, 6}}, 
 {2608, {4, 5, 5, 5, 6, 6, 6, 6, 6, 6, 6, 6, 6, 6, 6, 5}}, 
 {2591, {5, 5, 5, 6, 6, 6, 6, 5, 5, 6, 6, 5, 6, 6, 6, 6}}, 
 {2591, {5, 5, 5, 6, 6, 6, 5, 6, 6, 6, 5, 5, 6, 6, 6, 6}}, 
 {2585, {5, 5, 5, 6, 6, 6, 5, 6, 6, 5, 5, 6, 6, 6, 6, 6}}, 
 {2577, {4, 5, 5, 6, 6, 5, 5, 5, 5, 5, 5, 6, 6, 6, 6, 5, 5}}, 
 {2558, {4, 5, 5, 5, 6, 6, 6, 5, 5, 5, 5, 5, 6, 6, 6, 5, 5}}, 
 {2536, {4, 5, 5, 6, 6, 6, 6, 6, 6, 6, 6, 6, 6, 6, 5, 5}}, 
 {2501, {5, 5, 5, 6, 6, 6, 5, 5, 6, 6, 6, 5, 6, 6, 6, 6}}, 
 {2488, {5, 5, 5, 5, 5, 6, 5, 5, 5, 5, 6, 6, 5, 5, 5, 6, 6}}, 
 {2487, {4, 5, 5, 5, 5, 6, 6, 6, 5, 5, 5, 5, 6, 6, 6, 5, 5}}, 
 {2466, {5, 5, 5, 6, 6, 6, 5, 5, 6, 6, 6, 6, 5, 6, 6, 6}}, 
 {2460, {5, 5, 5, 5, 5, 6, 5, 5, 5, 6, 6, 5, 5, 5, 5, 6, 6}}, 
 {2452, {5, 5, 5, 6, 6, 6, 5, 5, 6, 6, 5, 6, 6, 6, 6, 6}}, 
 {2431, {5, 5, 5, 5, 6, 6, 6, 5, 6, 6, 6, 5, 6, 6, 6, 6}}, 
 {2379, {5, 5, 5, 5, 5, 6, 6, 5, 5, 5, 5, 6, 5, 5, 5, 6, 6}}, 
 {2363, {5, 5, 5, 6, 6, 5, 5, 6, 6, 6, 5, 6, 6, 6, 6, 6}}, 
 {2347, {5, 5, 5, 5, 5, 6, 5, 5, 6, 6, 5, 5, 5, 5, 5, 6, 6}}, 
 {2334, {5, 5, 5, 5, 6, 6, 6, 5, 6, 6, 5, 6, 6, 6, 6, 6}}, 
 {2323, {5, 5, 6, 6, 5, 5, 6, 6, 6, 5, 5, 6, 6, 6, 6, 6}}, 
 {2315, {5, 5, 5, 6, 6, 5, 6, 6, 6, 5, 5, 6, 6, 6, 6, 6}}, 
 {2310, {4, 5, 5, 5, 5, 5, 5, 5, 5, 5, 6, 6, 6, 6, 6, 6, 5}}, 
 {2307, {5, 5, 5, 5, 5, 6, 5, 5, 5, 5, 6, 6, 6, 5, 5, 5, 6}}, 
 {2294, {4, 5, 5, 5, 5, 5, 6, 6, 6, 5, 5, 5, 6, 6, 6, 5, 5}}, 
 {2287, {5, 5, 5, 5, 5, 6, 5, 5, 5, 5, 6, 5, 5, 5, 6, 6, 6}}, 
 {2277, {5, 5, 5, 6, 6, 5, 5, 6, 6, 6, 6, 5, 6, 6, 6, 6}}, 
 {2272, {5, 5, 5, 6, 6, 5, 6, 6, 6, 6, 5, 5, 6, 6, 6, 6}}, 
 {2253, {5, 5, 5, 6, 6, 5, 5, 6, 6, 6, 6, 6, 5, 6, 6, 6}}, 
 {2233, {5, 5, 5, 6, 6, 5, 6, 6, 5, 5, 6, 6, 6, 6, 6, 6}}, 
 {2222, {5, 5, 5, 5, 5, 6, 5, 5, 5, 6, 5, 5, 5, 5, 6, 6, 6}}, 
 {2214, {5, 5, 5, 6, 6, 5, 5, 6, 6, 5, 6, 6, 6, 6, 6, 6}}, 
 {2190, {5, 5, 5, 5, 5, 5, 6, 6, 5, 5, 5, 5, 6, 5, 5, 6, 6}}, 
 {2190, {4, 5, 5, 5, 6, 6, 5, 5, 5, 5, 5, 6, 6, 6, 6, 5, 5}}, 
 {2174, {5, 5, 5, 5, 5, 5, 6, 5, 5, 6, 6, 5, 5, 5, 5, 6, 6}}, 
 {2161, {4, 5, 5, 5, 6, 6, 6, 6, 5, 5, 5, 5, 5, 6, 6, 5, 5}}, 
 {2143, {5, 5, 5, 5, 5, 5, 6, 5, 5, 6, 5, 5, 5, 5, 6, 6, 6}}, 
 {2138, {5, 5, 5, 6, 6, 5, 6, 6, 6, 6, 6, 5, 5, 6, 6, 6}}, 
 {2119, {5, 5, 5, 5, 5, 5, 6, 5, 5, 5, 5, 6, 6, 5, 5, 6, 6}}, 
 {2088, {4, 5, 5, 5, 5, 6, 6, 6, 6, 5, 5, 5, 5, 6, 6, 5, 5}}, 
 {2079, {5, 5, 5, 5, 6, 6, 5, 6, 6, 6, 6, 5, 6, 6, 6, 6}}, 
 {2062, {5, 5, 5, 5, 5, 5, 6, 5, 5, 5, 6, 6, 5, 5, 5, 6, 6}}, 
 {2042, {5, 5, 5, 5, 6, 6, 5, 6, 6, 6, 5, 6, 6, 6, 6, 6}}, 
 {2012, {4, 5, 5, 5, 5, 6, 6, 5, 5, 5, 5, 6, 6, 6, 6, 5, 5}}, 
 {2003, {5, 5, 5, 5, 6, 6, 5, 6, 6, 5, 6, 6, 6, 6, 6, 6}}, 
 {1957, {4, 5, 5, 6, 6, 6, 5, 5, 5, 5, 5, 5, 5, 6, 6, 6, 5}}, 
 {1943, {5, 5, 5, 6, 6, 5, 5, 6, 6, 6, 6, 6, 6, 5, 6, 6}}, 
 {1917, {5, 5, 5, 5, 5, 5, 5, 6, 6, 5, 5, 5, 6, 5, 5, 6, 6}}, 
 {1913, {5, 5, 5, 5, 5, 6, 5, 5, 5, 5, 5, 6, 5, 5, 6, 6, 6}}, 
 {1894, {5, 5, 5, 5, 5, 5, 5, 6, 5, 5, 6, 6, 5, 5, 5, 6, 6}}, 
 {1880, {5, 5, 5, 5, 5, 5, 6, 5, 5, 5, 6, 5, 5, 5, 6, 6, 6}}, 
 {1862, {4, 5, 5, 5, 5, 5, 6, 6, 6, 6, 5, 5, 5, 6, 6, 5, 5}}, 
 {1860, {4, 5, 5, 5, 5, 5, 6, 6, 5, 5, 5, 6, 6, 6, 6, 5, 5}}, 
 {1831, {5, 5, 6, 5, 5, 6, 6, 5, 5, 6, 6, 6, 6, 6, 6, 6}}, 
 {1829, {5, 5, 5, 5, 5, 5, 5, 6, 5, 5, 5, 6, 6, 5, 5, 6, 6}}, 
 {1808, {5, 5, 5, 5, 6, 6, 5, 6, 6, 6, 6, 6, 5, 6, 6, 6}}, 
 {1792, {5, 5, 5, 5, 5, 5, 5, 6, 5, 5, 6, 5, 5, 5, 6, 6, 6}}, 
 {1792, {4, 5, 5, 6, 6, 6, 6, 6, 5, 5, 5, 5, 5, 5, 5, 6, 5}}, 
 {1770, {4, 5, 5, 5, 5, 6, 6, 6, 5, 5, 5, 6, 6, 6, 5, 5, 5}}, 
 {1748, {4, 5, 5, 5, 6, 6, 5, 5, 5, 5, 6, 6, 6, 6, 5, 5, 5}}, 
 {1746, {5, 5, 5, 5, 5, 5, 5, 5, 6, 5, 5, 6, 6, 5, 5, 6, 6}}, 
 {1743, {5, 5, 5, 5, 5, 5, 6, 5, 5, 5, 5, 6, 5, 5, 6, 6, 6}}, 
 {1742, {5, 5, 5, 6, 5, 5, 6, 6, 5, 6, 6, 6, 6, 6, 6, 6}}, 
 {1727, {5, 5, 5, 5, 5, 5, 6, 5, 5, 5, 5, 5, 6, 6, 5, 6, 6}}, 
 {1717, {5, 5, 5, 5, 5, 5, 6, 5, 5, 5, 5, 6, 6, 6, 5, 5, 6}}, 
 {1707, {5, 5, 6, 5, 5, 6, 6, 6, 6, 5, 5, 6, 6, 6, 6, 6}}, 
 {1694, {4, 5, 5, 5, 6, 5, 5, 5, 5, 5, 6, 6, 6, 6, 6, 5, 5}}, 
 {1688, {5, 5, 6, 5, 5, 6, 6, 6, 5, 5, 6, 6, 6, 6, 6, 6}}, 
 {1681, {1, 6, 6, 6, 6, 5, 5, 5, 5, 5, 5, 5, 6, 6, 6, 6, 6}}, 
 {1668, {5, 5, 5, 6, 5, 5, 6, 6, 6, 5, 6, 6, 6, 6, 6, 6}}, 
 {1654, {4, 5, 5, 6, 6, 5, 5, 5, 5, 5, 5, 5, 6, 6, 6, 6, 5}}, 
 {1651, {4, 5, 5, 5, 5, 5, 5, 5, 5, 5, 5, 5, 5, 5, 5, 5, 6, 5}}, 
 {1641, {4, 5, 5, 5, 5, 5, 5, 6, 6, 6, 5, 5, 6, 6, 6, 5, 5}}, 
 {1623, {5, 5, 5, 6, 5, 5, 6, 6, 6, 6, 6, 5, 6, 6, 6, 6}}, 
 {1607, {5, 5, 5, 5, 5, 5, 5, 6, 5, 5, 5, 5, 6, 6, 5, 6, 6}}, 
 {1593, {4, 5, 5, 5, 5, 5, 6, 6, 6, 6, 6, 6, 5, 5, 5, 5, 5}}, 
 {1592, {5, 5, 5, 6, 5, 5, 6, 6, 6, 6, 6, 6, 5, 6, 6, 6}}, 
 {1581, {5, 5, 5, 5, 5, 5, 5, 5, 6, 5, 5, 6, 5, 5, 6, 6, 6}}, 
 {1577, {4, 5, 5, 5, 5, 5, 5, 6, 6, 6, 6, 5, 5, 6, 6, 5, 5}}, 
 {1577, {1, 6, 6, 6, 5, 5, 5, 5, 5, 5, 5, 6, 6, 6, 6, 6, 6}}, 
 {1570, {4, 5, 5, 6, 6, 6, 5, 5, 5, 5, 5, 5, 6, 6, 6, 5, 5}}, 
 {1562, {5, 5, 5, 6, 5, 5, 6, 6, 6, 6, 5, 6, 6, 6, 6, 6}}, 
 {1545, {4, 5, 5, 5, 5, 6, 6, 6, 6, 5, 5, 5, 6, 6, 5, 5, 5}}, 
 {1543, {4, 5, 5, 5, 5, 5, 5, 5, 5, 5, 5, 5, 5, 5, 6, 5, 5, 5}}, 
 {1536, {5, 5, 5, 5, 5, 5, 5, 6, 5, 5, 5, 6, 5, 5, 6, 6, 6}}, 
 {1523, {4, 5, 5, 5, 6, 6, 6, 6, 6, 5, 5, 5, 5, 5, 5, 6, 5}}, 
 {1521, {4, 5, 5, 5, 5, 5, 5, 6, 6, 5, 5, 6, 6, 6, 6, 5, 5}}, 
 {1518, {4, 5, 5, 6, 6, 6, 6, 5, 5, 5, 5, 5, 5, 5, 6, 6, 5}}, 
 {1514, {4, 5, 5, 5, 5, 5, 5, 5, 5, 5, 6, 5, 5, 5, 5, 5, 5, 5}}, 
 {1505, {4, 5, 5, 5, 6, 6, 6, 5, 5, 5, 5, 5, 5, 6, 6, 6, 5}}, 
 {1496, {4, 5, 5, 5, 5, 6, 6, 6, 5, 5, 5, 5, 5, 6, 6, 6, 5}}, 
 {1478, {5, 5, 5, 5, 5, 5, 5, 6, 5, 5, 5, 6, 6, 6, 5, 5, 6}}, 
 {1471, {4, 5, 5, 5, 5, 5, 6, 6, 6, 5, 5, 5, 5, 6, 6, 6, 5}}, 
 {1457, {4, 5, 5, 5, 5, 5, 5, 5, 5, 5, 5, 5, 6, 5, 5, 5, 5, 5}}, 
 {1452, {4, 5, 5, 5, 5, 5, 5, 5, 5, 5, 5, 6, 5, 5, 5, 5, 5, 5}}, 
 {1432, {4, 5, 5, 5, 5, 6, 5, 5, 5, 5, 6, 6, 6, 6, 6, 5, 5}}, 
 {1420, {4, 5, 5, 5, 5, 5, 5, 6, 6, 6, 5, 5, 5, 6, 6, 6, 5}}, 
 {1398, {4, 5, 5, 5, 5, 5, 5, 5, 5, 5, 5, 5, 5, 5, 5, 6, 5, 5}}, 
 {1397, {4, 5, 5, 5, 5, 5, 6, 6, 6, 6, 6, 5, 5, 5, 5, 6, 5}}, 
 {1394, {4, 5, 5, 5, 5, 5, 5, 5, 6, 6, 6, 5, 6, 6, 6, 5, 5}}, 
 {1393, {4, 5, 5, 5, 5, 6, 6, 5, 5, 5, 6, 6, 6, 6, 5, 5, 5}}, 
 {1385, {5, 5, 5, 5, 5, 5, 5, 5, 6, 5, 5, 5, 6, 6, 5, 6, 6}}, 
 {1384, {4, 5, 5, 6, 5, 5, 5, 5, 5, 5, 6, 6, 6, 6, 6, 5, 5}}, 
 {1376, {4, 5, 5, 5, 5, 6, 6, 6, 6, 6, 5, 5, 5, 5, 5, 6, 5}}, 
 {1374, {4, 5, 5, 5, 5, 5, 5, 5, 5, 5, 5, 5, 5, 6, 5, 5, 5, 5}}, 
 {1365, {5, 5, 5, 5, 5, 5, 5, 5, 5, 6, 5, 5, 6, 6, 5, 6, 6}}, 
 {1353, {5, 5, 5, 6, 5, 5, 6, 6, 6, 6, 6, 6, 6, 5, 6, 6}}, 
 {1350, {4, 5, 5, 5, 6, 5, 5, 5, 5, 6, 6, 6, 6, 6, 5, 5, 5}}, 
 {1344, {4, 5, 5, 5, 5, 5, 6, 6, 6, 6, 5, 5, 6, 6, 5, 5, 5}}, 
 {1300, {4, 5, 5, 5, 5, 5, 6, 6, 6, 5, 5, 6, 6, 6, 5, 5, 5}}, 
 {1298, {5, 5, 5, 5, 6, 5, 5, 5, 5, 6, 6, 5, 5, 5, 5, 6, 6}}, 
 {1298, {5, 5, 5, 5, 6, 5, 5, 5, 5, 6, 5, 5, 5, 5, 6, 6, 6}}, 
 {1294, {4, 5, 5, 5, 5, 5, 6, 5, 5, 5, 6, 6, 6, 6, 6, 5, 5}}, 
 {1285, {4, 5, 5, 5, 6, 6, 6, 6, 5, 5, 5, 5, 5, 5, 6, 6, 5}}, 
 {1281, {4, 4, 5, 5, 5, 5, 5, 5, 6, 6, 6, 6, 6, 6, 6, 5, 5}}, 
 {1263, {4, 5, 5, 5, 6, 6, 5, 5, 5, 5, 5, 5, 6, 6, 6, 6, 5}}, 
 {1259, {4, 5, 5, 5, 6, 6, 6, 6, 6, 5, 5, 5, 5, 5, 6, 5, 5}}, 
 {1241, {4, 5, 5, 5, 5, 5, 6, 6, 6, 6, 5, 5, 5, 5, 6, 6, 5}}, 
 {1241, {4, 5, 5, 5, 5, 5, 6, 6, 5, 5, 5, 5, 6, 6, 6, 6, 5}}, 
 {1236, {4, 5, 5, 6, 6, 6, 6, 5, 6, 6, 6, 6, 6, 6, 6, 5}}, 
 {1233, {4, 5, 5, 5, 5, 6, 6, 6, 6, 6, 5, 5, 5, 5, 6, 5, 5}}, 
 {1233, {4, 5, 5, 5, 5, 5, 5, 6, 6, 6, 6, 6, 5, 5, 5, 6, 5}}, 
 {1227, {4, 5, 5, 5, 5, 6, 6, 6, 6, 5, 5, 5, 5, 5, 6, 6, 5}}, 
 {1211, {4, 5, 5, 5, 5, 5, 5, 5, 6, 6, 6, 6, 5, 6, 6, 5, 5}}, 
 {1200, {5, 5, 6, 6, 5, 5, 6, 6, 5, 5, 6, 6, 6, 6, 6, 6}}, 
 {1199, {4, 5, 5, 6, 6, 6, 5, 6, 6, 6, 6, 6, 6, 6, 6, 5}}, 
 {1196, {4, 5, 5, 6, 6, 6, 6, 6, 6, 6, 5, 6, 6, 6, 6, 5}}, 
 {1191, {5, 5, 6, 6, 6, 5, 5, 6, 6, 6, 5, 5, 6, 6, 6, 6}}, 
 {1181, {4, 4, 5, 5, 5, 5, 5, 6, 6, 6, 6, 6, 6, 6, 5, 5, 5}}, 
 {1179, {4, 5, 5, 6, 6, 6, 6, 6, 5, 6, 6, 6, 6, 6, 6, 5}}, 
 {1176, {5, 5, 5, 5, 5, 6, 5, 5, 5, 5, 5, 6, 6, 5, 5, 6, 6}}, 
 {1172, {4, 5, 6, 5, 5, 5, 5, 5, 5, 5, 5, 6, 6, 6, 6, 6, 5}}, 
 {1171, {4, 5, 5, 5, 5, 6, 6, 6, 6, 6, 5, 5, 5, 6, 5, 5, 5}}, 
 {1170, {5, 5, 5, 5, 6, 6, 6, 6, 5, 6, 6, 5, 6, 6, 6, 6}}, 
 {1169, {5, 5, 5, 5, 5, 6, 5, 5, 6, 5, 5, 5, 5, 5, 6, 6, 6}}, 
 {1161, {5, 5, 5, 5, 6, 6, 6, 5, 6, 6, 6, 6, 5, 6, 6, 6}}, 
 {1155, {4, 5, 5, 5, 5, 6, 6, 5, 5, 5, 5, 5, 6, 6, 6, 6, 5}}, 
 {1155, {5, 5, 6, 6, 5, 5, 6, 6, 6, 6, 5, 5, 6, 6, 6, 6}}, 
 {1146, {4, 5, 5, 5, 5, 5, 6, 6, 5, 5, 6, 6, 6, 6, 5, 5, 5}}, 
 {1131, {4, 5, 5, 6, 6, 6, 6, 6, 6, 6, 6, 6, 5, 6, 6, 5}}, 
 {1129, {4, 5, 5, 6, 6, 6, 6, 6, 6, 6, 6, 5, 6, 6, 6, 5}}, 
 {1116, {4, 5, 5, 5, 5, 5, 5, 5, 6, 6, 5, 6, 6, 6, 6, 5, 5}}, 
 {1115, {4, 5, 5, 5, 5, 5, 6, 6, 6, 6, 6, 5, 5, 5, 6, 5, 5}}, 
 {1115, {4, 5, 5, 5, 5, 5, 5, 6, 6, 5, 5, 5, 6, 6, 6, 6, 5}}, 
 {1110, {4, 5, 5, 6, 6, 6, 6, 6, 6, 5, 6, 6, 6, 6, 6, 5}}, 
 {1093, {4, 5, 5, 5, 5, 5, 5, 6, 5, 5, 6, 6, 6, 6, 6, 5, 5}}, 
 {1088, {4, 5, 6, 6, 6, 6, 6, 5, 5, 6, 6, 6, 6, 6, 6, 5}}, 
 {1080, {4, 5, 5, 5, 5, 5, 5, 6, 6, 6, 5, 6, 6, 6, 5, 5, 5}}, 
 {1066, {4, 5, 5, 5, 5, 6, 6, 5, 5, 6, 6, 6, 6, 5, 5, 5, 5}}, 
 {1055, {1, 6, 6, 5, 5, 5, 5, 5, 5, 5, 6, 6, 6, 6, 6, 6, 6}}, 
 {1053, {4, 5, 5, 5, 5, 6, 5, 5, 5, 6, 6, 6, 6, 6, 5, 5, 5}}, 
 {1050, {4, 5, 5, 5, 5, 5, 5, 6, 6, 6, 6, 5, 5, 5, 6, 6, 5}}, 
 {1049, {4, 5, 5, 6, 6, 5, 6, 6, 6, 6, 6, 6, 6, 6, 6, 5}}, 
 {1040, {4, 5, 5, 5, 5, 5, 5, 6, 6, 6, 6, 5, 6, 6, 5, 5, 5}}, 
 {1030, {4, 5, 5, 5, 5, 5, 6, 6, 6, 5, 6, 6, 6, 5, 5, 5, 5}}, 
 {1010, {4, 5, 5, 5, 6, 6, 6, 5, 5, 5, 5, 6, 6, 6, 5, 5, 5}}, 
 {997, {4, 5, 6, 6, 6, 6, 5, 5, 6, 6, 6, 6, 6, 6, 6, 5}}, 
 {995, {4, 5, 5, 6, 5, 5, 5, 5, 5, 5, 5, 6, 6, 6, 6, 6, 5}}, 
 {989, {5, 5, 5, 5, 5, 5, 6, 6, 5, 5, 5, 6, 5, 5, 5, 6, 6}}, 
 {984, {4, 5, 5, 5, 6, 5, 5, 5, 5, 5, 5, 6, 6, 6, 6, 6, 5}}, 
 {981, {4, 5, 5, 5, 5, 5, 5, 5, 6, 6, 6, 6, 6, 5, 5, 6, 5}}, 
 {972, {5, 5, 5, 5, 5, 5, 5, 5, 6, 6, 5, 5, 6, 5, 5, 6, 6}}, 
 {958, {4, 5, 6, 6, 5, 5, 5, 5, 5, 5, 5, 5, 6, 6, 6, 6, 5}}, 
 {949, {1, 6, 6, 6, 5, 5, 5, 5, 6, 6, 6, 5, 5, 5, 6, 6, 6}}, 
 {938, {4, 5, 5, 5, 5, 5, 5, 5, 6, 6, 6, 5, 5, 6, 6, 6, 5}}, 
 {927, {4, 5, 6, 6, 6, 5, 5, 6, 6, 6, 6, 6, 6, 6, 6, 5}}, 
 {925, {1, 6, 6, 6, 6, 6, 6, 5, 6, 6, 6, 6, 6, 6, 6, 6}}, 
 {917, {6, 6, 6, 6, 6, 6, 6, 6, 6, 6, 6, 6, 6, 6, 6}}, 
 {913, {4, 5, 5, 5, 5, 5, 5, 6, 6, 6, 6, 6, 5, 5, 6, 5, 5}}, 
 {910, {4, 5, 5, 5, 5, 5, 5, 6, 6, 5, 6, 6, 6, 6, 5, 5, 5}}, 
 {897, {5, 5, 5, 5, 5, 5, 6, 5, 5, 5, 6, 6, 6, 5, 5, 5, 6}}, 
 {891, {4, 5, 5, 6, 6, 5, 5, 5, 5, 6, 6, 5, 5, 6, 6, 5, 5}}, 
 {890, {1, 6, 6, 6, 5, 6, 6, 6, 6, 6, 6, 6, 6, 6, 6, 6}}, 
 {886, {4, 5, 5, 5, 5, 5, 6, 5, 5, 5, 5, 6, 6, 6, 6, 6, 5}}, 
 {885, {1, 6, 6, 6, 6, 6, 5, 6, 6, 6, 6, 6, 6, 6, 6, 6}}, 
 {883, {4, 5, 5, 5, 5, 5, 5, 5, 6, 6, 5, 5, 6, 6, 6, 6, 5}}, 
 {868, {4, 5, 5, 6, 6, 5, 5, 5, 5, 6, 6, 6, 5, 5, 5, 6, 5}}, 
 {867, {4, 5, 5, 6, 6, 5, 5, 5, 6, 6, 6, 5, 5, 5, 5, 6, 5}}, 
 {867, {4, 5, 5, 5, 5, 5, 5, 6, 5, 5, 5, 6, 6, 6, 6, 6, 5}}, 
 {860, {4, 5, 5, 6, 6, 6, 5, 5, 5, 6, 6, 5, 5, 5, 5, 6, 5}}, 
 {860, {4, 5, 6, 5, 5, 6, 6, 6, 6, 6, 6, 6, 6, 6, 6, 5}}, 
 {857, {4, 5, 5, 5, 5, 5, 6, 6, 6, 6, 6, 5, 5, 6, 5, 5, 5}}, 
 {857, {4, 4, 5, 5, 6, 6, 6, 5, 5, 5, 5, 6, 6, 6, 6, 5, 5}}, 
 {856, {4, 5, 5, 5, 5, 6, 5, 5, 5, 5, 5, 6, 6, 6, 6, 6, 5}}, 
 {855, {4, 5, 5, 5, 5, 5, 6, 6, 6, 6, 5, 6, 6, 5, 5, 5, 5}}, 
 {852, {1, 6, 6, 6, 6, 5, 6, 6, 6, 6, 6, 6, 6, 6, 6, 6}}, 
 {849, {5, 5, 5, 5, 6, 6, 5, 6, 6, 6, 6, 6, 6, 5, 6, 6}}, 
 {847, {4, 4, 5, 5, 5, 5, 5, 5, 5, 6, 6, 6, 6, 6, 6, 6, 5}}, 
 {847, {4, 5, 6, 6, 5, 5, 6, 6, 6, 6, 6, 6, 6, 6, 6, 5}}, 
 {846, {4, 5, 5, 5, 5, 5, 5, 5, 6, 6, 6, 6, 5, 5, 6, 6, 5}}, 
 {844, {4, 5, 5, 5, 5, 5, 6, 6, 5, 6, 6, 6, 6, 5, 5, 5, 5}}, 
 {838, {4, 5, 5, 5, 5, 5, 6, 5, 5, 6, 6, 6, 6, 6, 5, 5, 5}}, 
 {838, {4, 5, 5, 5, 4, 5, 5, 5, 6, 6, 6, 6, 6, 6, 6, 5, 5}}, 
 {834, {4, 4, 5, 5, 5, 5, 5, 5, 5, 5, 5, 5, 5, 6, 6, 5, 5, 5}}, 
 {816, {4, 5, 5, 6, 6, 6, 5, 5, 5, 5, 5, 6, 6, 5, 5, 6, 5}}, 
 {815, {1, 6, 6, 6, 5, 5, 5, 5, 6, 6, 5, 5, 5, 6, 6, 6, 6}}, 
 {814, {4, 5, 5, 6, 6, 6, 5, 5, 5, 5, 6, 6, 5, 5, 5, 6, 5}}, 
 {799, {1, 6, 6, 6, 5, 5, 5, 6, 6, 5, 5, 5, 5, 6, 6, 6, 6}}, 
 {796, {4, 5, 5, 6, 6, 5, 5, 6, 5, 5, 5, 5, 6, 6, 6, 5, 5}}, 
 {795, {4, 5, 5, 5, 5, 6, 5, 5, 6, 6, 6, 6, 6, 5, 5, 5, 5}}, 
 {771, {4, 4, 5, 5, 5, 5, 5, 5, 5, 5, 5, 5, 5, 5, 6, 6, 5, 5}}, 
 {764, {4, 4, 5, 5, 5, 5, 5, 5, 5, 5, 5, 5, 6, 6, 5, 5, 5, 5}}, 
 {761, {4, 5, 5, 6, 6, 5, 5, 6, 6, 6, 5, 5, 5, 5, 5, 6, 5}}, 
 {761, {4, 5, 5, 6, 6, 5, 5, 5, 6, 5, 5, 5, 6, 6, 6, 5, 5}}, 
 {760, {4, 5, 5, 5, 5, 5, 5, 5, 5, 6, 6, 6, 5, 6, 6, 6, 5}}, 
 {753, {4, 5, 5, 6, 6, 6, 5, 5, 6, 6, 5, 5, 5, 5, 5, 6, 5}}, 
 {751, {1, 6, 6, 5, 6, 6, 6, 6, 6, 6, 6, 6, 6, 6, 6, 6}}, 
 {750, {4, 5, 5, 5, 5, 5, 5, 5, 5, 6, 5, 5, 5, 5, 5, 5, 5, 5}}, 
 {743, {4, 5, 5, 6, 6, 5, 5, 5, 5, 5, 6, 6, 6, 5, 5, 6, 5}}, 
 {742, {4, 4, 5, 5, 5, 5, 5, 5, 5, 5, 6, 6, 5, 5, 5, 5, 5, 5}}, 
 {742, {4, 4, 5, 5, 5, 6, 6, 6, 5, 5, 5, 6, 6, 6, 6, 5, 5}}, 
 {735, {4, 4, 5, 5, 5, 6, 6, 6, 6, 5, 5, 5, 6, 6, 6, 5, 5}}, 
 {728, {4, 4, 5, 5, 5, 5, 5, 5, 5, 5, 5, 6, 6, 5, 5, 5, 5, 5}}, 
 {713, {4, 5, 5, 6, 6, 5, 5, 5, 5, 6, 5, 5, 6, 6, 6, 5, 5}}, 
 {709, {1, 6, 6, 6, 5, 5, 5, 5, 5, 6, 6, 6, 5, 5, 6, 6, 6}}, 
 {707, {4, 5, 5, 5, 5, 5, 5, 5, 5, 6, 6, 6, 6, 5, 6, 6, 5}}, 
 {696, {1, 6, 6, 5, 5, 5, 5, 6, 6, 6, 6, 5, 5, 5, 6, 6, 6}}, 
 {693, {5, 5, 6, 5, 5, 6, 5, 5, 6, 6, 6, 6, 6, 6, 6, 6}}, 
 {690, {1, 6, 6, 5, 5, 5, 5, 6, 6, 6, 5, 5, 5, 6, 6, 6, 6}}, 
 {687, {5, 5, 6, 6, 5, 5, 6, 6, 5, 6, 6, 5, 6, 6, 6, 6}}, 
 {677, {4, 5, 5, 6, 6, 5, 5, 5, 5, 5, 6, 6, 5, 6, 6, 5, 5}}, 
 {670, {4, 5, 5, 5, 5, 5, 5, 5, 5, 6, 6, 5, 6, 6, 6, 6, 5}}, 
 {659, {4, 5, 5, 6, 6, 5, 5, 5, 6, 6, 5, 5, 5, 5, 6, 6, 5}}, 
 {659, {4, 5, 5, 5, 6, 6, 5, 5, 6, 6, 5, 5, 5, 6, 6, 5, 5}}, 
 {657, {4, 4, 5, 5, 6, 6, 5, 5, 5, 5, 6, 6, 6, 6, 6, 5, 5}}, 
 {645, {5, 5, 6, 6, 5, 5, 6, 6, 5, 6, 6, 6, 5, 6, 6, 6}}, 
 {638, {1, 6, 6, 6, 6, 5, 5, 5, 5, 6, 5, 5, 5, 6, 6, 6, 6}}, 
 {638, {1, 6, 6, 6, 5, 5, 6, 6, 5, 5, 5, 5, 5, 6, 6, 6, 6}}, 
 {637, {4, 4, 5, 5, 5, 5, 5, 5, 5, 5, 5, 5, 5, 5, 5, 6, 6, 5}}, 
 {636, {4, 5, 5, 4, 5, 5, 5, 5, 6, 6, 6, 6, 6, 6, 6, 5, 5}}, 
 {635, {1, 6, 6, 5, 5, 5, 6, 6, 6, 5, 5, 5, 5, 6, 6, 6, 6}}, 
 {633, {5, 5, 6, 6, 5, 6, 6, 5, 6, 6, 6, 5, 5, 6, 6, 6}}, 
 {630, {5, 5, 6, 6, 5, 6, 6, 5, 5, 6, 6, 5, 6, 6, 6, 6}}, 
 {628, {4, 5, 5, 5, 5, 5, 5, 5, 6, 5, 5, 6, 6, 6, 6, 6, 5}}, 
 {626, {4, 4, 5, 5, 6, 6, 6, 5, 5, 5, 5, 5, 6, 6, 6, 6, 5}}, 
 {625, {4, 5, 5, 6, 6, 6, 6, 5, 5, 5, 6, 5, 5, 5, 5, 6, 5}}, 
 {625, {4, 5, 5, 6, 6, 6, 6, 5, 5, 5, 5, 6, 5, 5, 5, 6, 5}}, 
 {625, {4, 5, 5, 6, 6, 5, 6, 6, 6, 5, 5, 5, 5, 5, 5, 6, 5}}, 
 {624, {4, 5, 5, 6, 5, 5, 5, 5, 6, 6, 6, 5, 5, 6, 6, 5, 5}}, 
 {621, {1, 6, 6, 6, 5, 5, 5, 5, 5, 6, 6, 5, 5, 6, 6, 6, 6}}, 
 {616, {5, 5, 5, 6, 6, 5, 6, 6, 5, 6, 6, 5, 6, 6, 6, 6}}, 
 {614, {4, 5, 6, 6, 6, 5, 5, 5, 5, 5, 5, 5, 5, 6, 6, 6, 5}}, 
 {614, {1, 6, 6, 5, 5, 5, 6, 6, 6, 6, 5, 5, 5, 5, 6, 6, 6}}, 
 {611, {5, 5, 5, 5, 5, 5, 5, 5, 6, 5, 5, 6, 6, 6, 5, 5, 6}}, 
 {611, {4, 4, 5, 5, 5, 5, 6, 6, 6, 6, 6, 6, 6, 5, 5, 5, 5}}, 
 {610, {4, 5, 5, 6, 6, 6, 6, 5, 5, 5, 5, 5, 6, 5, 5, 6, 5}}, 
 {610, {4, 4, 5, 5, 6, 6, 6, 6, 5, 5, 5, 5, 5, 6, 6, 6, 5}}, 
 {597, {4, 5, 5, 6, 6, 5, 5, 5, 5, 5, 6, 6, 5, 5, 6, 6, 5}}, 
 {595, {4, 5, 5, 5, 6, 6, 6, 5, 5, 5, 5, 6, 6, 5, 5, 6, 5}}, 
 {595, {4, 5, 5, 5, 6, 6, 5, 5, 6, 5, 5, 5, 6, 6, 6, 5, 5}}, 
 {595, {5, 5, 5, 6, 6, 5, 6, 6, 6, 5, 6, 6, 5, 6, 6, 6}}, 
 {594, {4, 5, 5, 5, 5, 6, 6, 6, 5, 5, 6, 6, 6, 5, 5, 5, 5}}, 
 {592, {4, 5, 5, 6, 5, 5, 6, 6, 5, 5, 5, 5, 6, 6, 6, 5, 5}}, 
 {590, {4, 4, 5, 5, 6, 6, 6, 6, 6, 6, 5, 5, 5, 5, 5, 6, 5}}, 
 {589, {4, 5, 5, 5, 6, 6, 6, 5, 5, 5, 6, 6, 5, 5, 5, 6, 5}}, 
 {588, {3, 5, 5, 6, 6, 6, 6, 6, 6, 6, 6, 6, 6, 6, 6, 5}}, 
 {585, {4, 4, 5, 5, 5, 6, 6, 6, 6, 6, 5, 5, 5, 6, 6, 5, 5}}, 
 {583, {4, 5, 5, 6, 6, 6, 5, 5, 5, 6, 5, 5, 5, 5, 6, 6, 5}}, 
 {583, {4, 5, 5, 6, 6, 5, 5, 5, 5, 6, 6, 5, 5, 5, 6, 6, 5}}, 
 {582, {4, 5, 5, 5, 6, 6, 5, 5, 5, 5, 6, 6, 6, 5, 5, 6, 5}}, 
 {581, {4, 5, 5, 5, 4, 5, 5, 5, 5, 5, 5, 5, 5, 5, 6, 6, 5, 5}}, 
 {581, {4, 5, 5, 6, 5, 5, 5, 6, 6, 5, 5, 5, 6, 6, 6, 5, 5}}, 
 {579, {4, 4, 5, 5, 6, 6, 6, 6, 6, 5, 5, 5, 5, 5, 6, 6, 5}}, 
 {578, {4, 5, 5, 6, 6, 5, 5, 6, 6, 5, 5, 5, 5, 5, 6, 6, 5}}, 
 {577, {4, 5, 5, 6, 5, 5, 5, 6, 6, 6, 5, 5, 5, 6, 6, 5, 5}}, 
 {576, {4, 5, 5, 5, 6, 6, 5, 5, 5, 6, 6, 5, 5, 6, 6, 5, 5}}, 
 {573, {4, 5, 5, 5, 6, 6, 5, 5, 6, 6, 6, 5, 5, 5, 5, 6, 5}}, 
 {573, {1, 6, 6, 6, 5, 5, 5, 5, 6, 5, 5, 5, 6, 6, 6, 6, 6}}, 
 {569, {4, 4, 5, 5, 5, 6, 6, 6, 6, 5, 5, 5, 5, 6, 6, 6, 5}}, 
 {566, {4, 4, 5, 5, 5, 6, 6, 6, 5, 5, 5, 5, 6, 6, 6, 6, 5}}, 
 {564, {1, 6, 6, 5, 5, 5, 5, 6, 6, 5, 5, 5, 6, 6, 6, 6, 6}}, 
 {563, {4, 5, 5, 5, 6, 5, 5, 6, 6, 5, 5, 5, 6, 6, 6, 5, 5}}, 
 {562, {4, 5, 5, 6, 5, 5, 6, 6, 6, 5, 5, 5, 5, 6, 6, 5, 5}}, 
 {558, {4, 5, 6, 5, 5, 5, 5, 6, 6, 6, 5, 5, 5, 5, 6, 6, 5}}, 
 {558, {4, 5, 5, 5, 6, 6, 6, 5, 5, 5, 6, 5, 5, 6, 6, 5, 5}}, 
 {557, {5, 5, 5, 6, 6, 5, 6, 6, 5, 6, 6, 6, 5, 6, 6, 6}}, 
 {556, {5, 5, 5, 5, 6, 5, 5, 6, 5, 5, 5, 5, 6, 5, 5, 6, 6}}, 
 {556, {4, 5, 5, 6, 6, 6, 5, 6, 6, 5, 5, 5, 5, 5, 5, 6, 5}}, 
 {556, {4, 4, 5, 5, 5, 6, 6, 5, 5, 5, 6, 6, 6, 6, 6, 5, 5}}, 
 {550, {4, 4, 5, 5, 5, 5, 6, 6, 6, 6, 5, 5, 6, 6, 6, 5, 5}}, 
 {549, {4, 5, 5, 5, 6, 6, 5, 5, 5, 6, 6, 6, 5, 5, 5, 6, 5}}, 
 {548, {4, 5, 5, 6, 6, 6, 5, 5, 5, 5, 6, 5, 5, 5, 6, 6, 5}}, 
 {548, {4, 5, 5, 5, 5, 6, 6, 5, 5, 6, 6, 5, 5, 6, 6, 5, 5}}, 
 {547, {4, 5, 5, 6, 6, 6, 6, 5, 5, 6, 5, 5, 5, 5, 5, 6, 5}}, 
 {547, {4, 5, 5, 6, 6, 5, 5, 5, 5, 6, 5, 5, 5, 6, 6, 6, 5}}, 
 {546, {4, 5, 5, 5, 5, 5, 4, 5, 5, 6, 6, 6, 6, 6, 6, 6, 5}}, 
 {542, {4, 5, 5, 5, 4, 5, 5, 5, 5, 6, 6, 6, 6, 6, 6, 6, 5}}, 
 {540, {4, 5, 5, 6, 6, 5, 5, 5, 6, 5, 5, 5, 5, 6, 6, 6, 5}}, 
 {540, {4, 5, 5, 5, 6, 5, 5, 6, 6, 6, 5, 5, 5, 6, 6, 5, 5}}, 
 {540, {1, 6, 6, 6, 5, 5, 5, 6, 5, 5, 5, 5, 6, 6, 6, 6, 6}}, 
 {539, {4, 4, 5, 5, 5, 6, 6, 6, 6, 6, 6, 5, 5, 5, 5, 6, 5}}, 
 {538, {4, 5, 5, 6, 6, 6, 5, 5, 5, 5, 5, 6, 5, 5, 6, 6, 5}}, 
 {538, {4, 4, 5, 5, 5, 6, 6, 6, 5, 5, 6, 6, 6, 6, 5, 5, 5}}, 
 {535, {5, 5, 5, 5, 6, 5, 5, 5, 5, 6, 5, 5, 6, 5, 5, 6, 6}}, 
 {534, {5, 5, 5, 5, 5, 6, 5, 5, 6, 5, 5, 6, 5, 5, 5, 6, 6}}, 
 {534, {4, 5, 6, 5, 5, 5, 5, 5, 6, 6, 6, 6, 5, 5, 5, 6, 5}}, 
 {534, {4, 4, 5, 5, 5, 5, 6, 6, 6, 5, 5, 6, 6, 6, 6, 5, 5}}, 
 {533, {1, 6, 6, 5, 5, 6, 6, 6, 5, 5, 5, 5, 5, 6, 6, 6, 6}}, 
 {532, {4, 5, 5, 6, 5, 5, 5, 6, 6, 6, 6, 5, 5, 5, 5, 6, 5}}, 
 {531, {1, 6, 6, 6, 5, 5, 5, 5, 5, 5, 6, 6, 6, 5, 6, 6, 6}}, 
 {530, {4, 5, 5, 5, 6, 6, 6, 5, 5, 6, 6, 5, 5, 5, 5, 6, 5}}, 
 {530, {5, 5, 5, 6, 6, 5, 6, 6, 5, 6, 6, 6, 6, 5, 6, 6}}, 
 {526, {5, 5, 5, 5, 5, 6, 5, 5, 5, 6, 6, 5, 5, 6, 5, 5, 6}}, 
 {525, {4, 5, 6, 5, 5, 5, 5, 5, 6, 6, 5, 5, 5, 6, 6, 6, 5}}, 
 {523, {4, 5, 5, 5, 5, 5, 5, 4, 5, 5, 5, 5, 5, 5, 6, 6, 5, 5}}, 
 {521, {5, 5, 5, 5, 5, 6, 5, 5, 5, 6, 5, 5, 6, 6, 5, 5, 6}}, 
 {520, {4, 5, 5, 5, 6, 5, 5, 5, 6, 6, 6, 5, 5, 6, 6, 5, 5}}, 
 {519, {4, 4, 5, 5, 5, 6, 6, 6, 6, 6, 5, 5, 5, 5, 6, 6, 5}}, 
 {518, {4, 5, 6, 5, 5, 5, 5, 6, 6, 5, 5, 5, 5, 6, 6, 6, 5}}, 
 {516, {4, 5, 5, 5, 5, 5, 5, 5, 4, 5, 5, 5, 5, 5, 6, 6, 5, 5}}, 
 {515, {4, 5, 5, 5, 6, 6, 5, 5, 5, 5, 6, 6, 5, 6, 6, 5, 5}}, 
 {514, {5, 5, 5, 5, 6, 5, 5, 6, 5, 5, 5, 6, 5, 5, 5, 6, 6}}, 
 {514, {4, 5, 5, 6, 5, 5, 5, 5, 6, 6, 5, 5, 6, 6, 6, 5, 5}}, 
 {514, {4, 5, 5, 5, 6, 5, 5, 5, 5, 6, 6, 5, 6, 6, 6, 5, 5}}, 
 {509, {5, 5, 5, 5, 5, 5, 5, 5, 5, 5, 5, 5, 5, 5, 5, 5, 5, 5}}, 
 {509, {4, 5, 5, 5, 5, 4, 5, 5, 5, 5, 5, 5, 5, 5, 6, 6, 5, 5}}, 
 {508, {4, 5, 5, 6, 6, 5, 5, 6, 5, 5, 5, 5, 5, 6, 6, 6, 5}}, 
 {505, {5, 5, 5, 5, 5, 6, 5, 5, 5, 6, 5, 5, 6, 5, 5, 6, 6}}, 
 {505, {4, 5, 5, 5, 6, 6, 5, 5, 5, 6, 5, 5, 6, 6, 6, 5, 5}}, 
 {504, {4, 5, 5, 6, 5, 5, 5, 5, 6, 6, 6, 6, 5, 5, 5, 6, 5}}, 
 {502, {4, 5, 5, 5, 5, 5, 5, 5, 5, 5, 4, 5, 5, 5, 6, 6, 5, 5}}, 
 {502, {4, 5, 5, 5, 6, 6, 5, 6, 6, 5, 5, 5, 5, 6, 6, 5, 5}}, 
 {501, {5, 5, 5, 5, 6, 5, 5, 5, 6, 5, 5, 6, 5, 5, 5, 6, 6}}, 
 {500, {1, 6, 6, 6, 6, 5, 5, 5, 5, 5, 6, 5, 5, 6, 6, 6, 6}}, 
 {498, {4, 5, 5, 5, 6, 6, 6, 5, 5, 6, 5, 5, 5, 6, 6, 5, 5}}, 
 {496, {5, 5, 5, 5, 6, 5, 5, 5, 6, 5, 5, 6, 6, 5, 5, 5, 6}}, 
 {493, {4, 5, 6, 5, 5, 5, 5, 5, 6, 6, 6, 5, 5, 5, 6, 6, 5}}, 
 {492, {5, 5, 6, 5, 5, 6, 6, 5, 6, 6, 6, 5, 6, 6, 6, 6}}, 
 {490, {4, 5, 5, 6, 6, 6, 5, 5, 6, 5, 5, 5, 5, 5, 6, 6, 5}}, 
 {490, {4, 4, 5, 5, 5, 5, 6, 6, 6, 6, 5, 5, 5, 6, 6, 6, 5}}, 
 {489, {5, 5, 5, 5, 6, 5, 5, 5, 6, 5, 5, 5, 6, 5, 5, 6, 6}}, 
 {489, {4, 5, 5, 6, 5, 5, 5, 5, 5, 6, 6, 6, 5, 6, 6, 5, 5}}, 
 {487, {3, 5, 5, 5, 5, 5, 5, 5, 6, 6, 6, 6, 6, 6, 6, 5, 5}}, 
 {487, {1, 6, 6, 6, 5, 6, 6, 5, 5, 5, 5, 5, 5, 6, 6, 6, 6}}, 
 {487, {1, 6, 6, 5, 6, 6, 6, 6, 5, 5, 5, 5, 5, 5, 6, 6, 6}}, 
 {487, {1, 6, 6, 5, 5, 6, 6, 5, 5, 5, 5, 5, 6, 6, 6, 6, 6}}, 
 {486, {4, 4, 5, 5, 5, 5, 6, 6, 6, 6, 6, 5, 5, 6, 6, 5, 5}}, 
 {484, {4, 5, 5, 5, 5, 5, 5, 5, 5, 4, 5, 5, 5, 5, 6, 6, 5, 5}}, 
 {484, {4, 5, 5, 5, 6, 5, 5, 5, 6, 6, 5, 5, 6, 6, 6, 5, 5}}, 
 {483, {5, 5, 5, 5, 5, 6, 5, 5, 6, 5, 5, 5, 6, 5, 5, 6, 6}}, 
 {481, {5, 5, 5, 5, 5, 5, 6, 5, 5, 6, 5, 5, 6, 5, 5, 6, 6}}, 
 {481, {5, 5, 6, 5, 5, 6, 6, 5, 6, 6, 5, 6, 6, 6, 6, 6}}, 
 {480, {4, 5, 6, 5, 5, 5, 6, 6, 5, 5, 5, 5, 5, 6, 6, 6, 5}}, 
 {478, {4, 4, 5, 5, 6, 6, 5, 5, 5, 5, 5, 6, 6, 6, 6, 6, 5}}, 
 {475, {4, 5, 5, 5, 4, 5, 5, 5, 5, 5, 5, 6, 6, 5, 5, 5, 5, 5}}, 
 {475, {4, 5, 5, 5, 6, 5, 5, 5, 5, 6, 6, 6, 6, 5, 5, 6, 5}}, 
 {475, {4, 5, 5, 5, 5, 4, 5, 5, 5, 6, 6, 6, 6, 6, 6, 6, 5}}, 
 {475, {1, 6, 6, 5, 6, 6, 6, 5, 5, 5, 5, 5, 5, 6, 6, 6, 6}}, 
 {472, {4, 5, 5, 5, 4, 5, 5, 5, 5, 5, 5, 5, 5, 6, 6, 5, 5, 5}}, 
 {472, {4, 5, 5, 5, 6, 6, 6, 5, 5, 4, 5, 6, 6, 6, 6, 5, 5}}, 
 {472, {4, 4, 5, 5, 5, 5, 6, 6, 6, 6, 6, 6, 5, 5, 5, 6, 5}}, 
 {472, {4, 4, 5, 5, 5, 5, 6, 6, 6, 5, 5, 5, 6, 6, 6, 6, 5}}, 
 {470, {5, 5, 5, 5, 6, 5, 5, 5, 6, 6, 5, 5, 5, 6, 5, 5, 6}}, 
 {470, {5, 5, 5, 5, 5, 6, 5, 5, 6, 5, 5, 5, 6, 6, 5, 5, 6}}, 
 {470, {4, 5, 5, 6, 5, 5, 6, 5, 5, 5, 5, 6, 6, 6, 6, 5, 5}}, 
 {470, {1, 6, 6, 5, 5, 5, 5, 5, 6, 6, 6, 6, 5, 5, 6, 6, 6}}, 
 {468, {4, 4, 5, 5, 5, 5, 6, 6, 5, 5, 6, 6, 6, 6, 6, 5, 5}}, 
 {468, {1, 6, 6, 5, 5, 5, 6, 6, 5, 5, 5, 5, 6, 6, 6, 6, 6}}, 
 {466, {4, 5, 5, 5, 6, 6, 6, 6, 5, 5, 4, 5, 6, 6, 6, 5, 5}}, 
 {466, {4, 4, 5, 5, 5, 5, 6, 6, 6, 6, 6, 5, 5, 5, 6, 6, 5}}, 
 {465, {1, 6, 6, 6, 5, 5, 5, 5, 5, 6, 5, 5, 6, 6, 6, 6, 6}}, 
 {465, {1, 6, 6, 5, 5, 6, 6, 6, 6, 5, 5, 5, 5, 5, 6, 6, 6}}, 
 {464, {4, 5, 5, 5, 5, 5, 6, 6, 5, 6, 6, 5, 5, 6, 6, 5, 5}}, 
 {463, {1, 6, 6, 6, 5, 5, 6, 5, 5, 5, 5, 5, 6, 6, 6, 6, 6}}, 
 {463, {1, 6, 6, 5, 5, 5, 5, 5, 6, 6, 6, 5, 5, 6, 6, 6, 6}}, 
 {463, {1, 6, 6, 6, 6, 6, 6, 6, 5, 6, 6, 6, 6, 6, 6, 6}}, 
 {462, {4, 5, 5, 5, 6, 5, 4, 5, 5, 6, 6, 6, 6, 6, 6, 5, 5}}, 
 {460, {4, 5, 5, 5, 5, 6, 6, 6, 5, 5, 5, 6, 6, 5, 5, 6, 5}}, 
 {460, {4, 5, 5, 5, 5, 4, 5, 5, 6, 6, 6, 6, 6, 6, 6, 5, 5}}, 
 {459, {4, 4, 5, 6, 6, 6, 5, 5, 5, 5, 5, 5, 6, 6, 6, 6, 5}}, 
 {459, {5, 5, 6, 5, 5, 6, 6, 5, 6, 6, 6, 6, 5, 6, 6, 6}}, 
 {458, {5, 5, 5, 5, 6, 5, 5, 5, 6, 5, 5, 5, 6, 6, 5, 5, 6}}, 
 {457, {4, 5, 5, 6, 5, 5, 5, 6, 6, 6, 5, 5, 5, 5, 6, 6, 5}}, 
 {457, {4, 5, 5, 5, 5, 6, 6, 6, 5, 5, 6, 5, 5, 6, 6, 5, 5}}, 
 {457, {4, 4, 5, 6, 5, 5, 5, 5, 5, 5, 6, 6, 6, 6, 6, 6, 5}}, 
 {456, {1, 6, 6, 6, 5, 5, 5, 5, 5, 5, 6, 6, 5, 6, 6, 6, 6}}, 
 {453, {4, 5, 5, 5, 5, 6, 6, 5, 5, 5, 6, 6, 6, 5, 5, 6, 5}}, 
 {452, {4, 5, 5, 5, 5, 5, 4, 5, 5, 5, 5, 5, 5, 5, 6, 6, 5, 5}}, 
 {452, {4, 5, 5, 6, 5, 5, 5, 5, 6, 6, 6, 5, 5, 5, 6, 6, 5}}, 
 {452, {4, 5, 5, 5, 6, 6, 6, 5, 5, 4, 5, 5, 6, 6, 6, 6, 5}}, 
 {452, {4, 4, 5, 5, 5, 6, 6, 5, 5, 5, 5, 6, 6, 6, 6, 6, 5}}, 
 {451, {4, 4, 5, 5, 5, 5, 5, 6, 6, 6, 6, 5, 6, 6, 6, 5, 5}}, 
 {450, {4, 5, 5, 5, 6, 6, 6, 6, 5, 5, 5, 5, 6, 5, 5, 6, 5}}, 
 {447, {4, 5, 5, 6, 5, 5, 6, 6, 6, 6, 5, 5, 5, 5, 5, 6, 5}}, 
 {447, {4, 5, 5, 5, 6, 6, 6, 6, 5, 4, 5, 5, 6, 6, 6, 5, 5}}, 
 {446, {4, 5, 5, 5, 6, 5, 5, 6, 5, 5, 5, 6, 6, 6, 6, 5, 5}}, 
 {446, {4, 5, 5, 5, 5, 6, 5, 5, 6, 6, 5, 5, 6, 6, 6, 5, 5}}, 
 {444, {4, 5, 6, 5, 5, 5, 6, 6, 6, 5, 5, 5, 5, 5, 6, 6, 5}}, 
 {444, {4, 5, 5, 5, 6, 6, 5, 5, 6, 6, 5, 5, 5, 5, 6, 6, 5}}, 
 {444, {5, 5, 6, 5, 5, 6, 6, 6, 5, 6, 6, 5, 6, 6, 6, 6}}, 
 {443, {4, 5, 5, 5, 6, 6, 6, 5, 6, 6, 5, 5, 5, 5, 5, 6, 5}}, 
 {442, {4, 5, 5, 5, 5, 5, 4, 5, 5, 5, 5, 5, 5, 6, 6, 5, 5, 5}}, 
 {442, {4, 5, 6, 5, 5, 5, 5, 6, 5, 5, 5, 5, 6, 6, 6, 6, 5}}, 
 {441, {4, 5, 5, 6, 6, 5, 5, 5, 5, 5, 6, 5, 5, 6, 6, 6, 5}}, 
 {441, {4, 4, 5, 5, 5, 6, 5, 5, 5, 6, 6, 6, 6, 6, 6, 5, 5}}, 
 {440, {3, 5, 5, 6, 6, 6, 5, 5, 5, 5, 5, 6, 6, 6, 6, 5, 5}}, 
 {439, {4, 5, 5, 6, 6, 6, 5, 5, 4, 5, 5, 6, 6, 6, 6, 5, 5}}, 
 {439, {4, 5, 5, 5, 6, 5, 5, 5, 6, 6, 6, 6, 5, 5, 5, 6, 5}}, 
 {439, {4, 4, 5, 5, 5, 6, 6, 5, 5, 6, 6, 6, 6, 6, 5, 5, 5}}, 
 {438, {4, 5, 5, 5, 6, 6, 5, 5, 5, 5, 6, 6, 5, 5, 6, 6, 5}}, 
 {438, {4, 5, 5, 5, 6, 5, 5, 5, 5, 6, 6, 6, 5, 6, 6, 5, 5}}, 
 {435, {4, 5, 5, 6, 5, 5, 5, 5, 6, 6, 5, 5, 5, 6, 6, 6, 5}}, 
 {435, {4, 5, 5, 5, 6, 6, 5, 5, 5, 6, 6, 5, 6, 6, 5, 5, 5}}, 
 {434, {4, 5, 5, 4, 5, 5, 5, 5, 5, 5, 5, 5, 5, 5, 6, 6, 5, 5}}, 
 {433, {4, 5, 5, 5, 6, 5, 5, 6, 6, 6, 6, 5, 5, 5, 5, 6, 5}}, 
 {432, {4, 5, 5, 6, 6, 5, 6, 6, 5, 5, 5, 5, 5, 5, 6, 6, 5}}, 
 {432, {1, 6, 6, 5, 5, 5, 5, 6, 6, 6, 6, 6, 5, 5, 5, 6, 6}}, 
 {431, {5, 5, 5, 5, 5, 5, 6, 5, 5, 6, 5, 5, 6, 6, 5, 5, 6}}, 
 {431, {4, 5, 5, 5, 5, 6, 6, 5, 5, 6, 5, 5, 6, 6, 6, 5, 5}}, 
 {431, {4, 5, 5, 5, 5, 6, 6, 5, 5, 5, 6, 6, 5, 6, 6, 5, 5}}, 
 {428, {4, 5, 5, 6, 6, 5, 5, 5, 6, 6, 5, 5, 5, 6, 6, 5, 5}}, 
 {427, {4, 5, 5, 6, 6, 5, 5, 5, 5, 5, 5, 6, 6, 5, 6, 6, 5}}, 
 {427, {4, 5, 5, 5, 6, 6, 6, 5, 4, 5, 5, 6, 6, 6, 6, 5, 5}}, 
 {425, {4, 5, 5, 5, 5, 5, 6, 6, 5, 5, 6, 6, 5, 6, 6, 5, 5}}, 
 {425, {4, 4, 5, 5, 6, 5, 5, 5, 5, 6, 6, 6, 6, 6, 6, 5, 5}}, 
 {424, {4, 5, 5, 6, 5, 5, 5, 5, 5, 6, 6, 6, 5, 5, 6, 6, 5}}, 
 {423, {4, 5, 5, 5, 4, 5, 5, 5, 5, 5, 5, 5, 6, 6, 5, 5, 5, 5}}, 
 {422, {4, 5, 5, 5, 5, 4, 5, 5, 5, 5, 5, 5, 6, 6, 5, 5, 5, 5}}, 
 {420, {4, 5, 5, 6, 5, 5, 5, 5, 6, 5, 5, 6, 6, 6, 6, 5, 5}}, 
 {419, {4, 5, 6, 5, 5, 6, 6, 5, 5, 5, 5, 5, 5, 6, 6, 6, 5}}, 
 {417, {4, 5, 5, 6, 5, 5, 5, 5, 5, 6, 6, 5, 6, 6, 6, 5, 5}}, 
 {417, {4, 5, 5, 5, 6, 6, 6, 5, 5, 5, 6, 6, 5, 5, 6, 5, 5}}, 
 {417, {4, 5, 5, 5, 5, 6, 6, 5, 5, 6, 6, 6, 5, 5, 5, 6, 5}}, 
 {416, {4, 5, 5, 5, 6, 6, 5, 6, 6, 6, 5, 5, 5, 5, 5, 6, 5}}, 
 {414, {4, 5, 5, 6, 5, 5, 5, 6, 5, 5, 5, 6, 6, 6, 6, 5, 5}}, 
 {414, {4, 5, 5, 5, 6, 5, 5, 6, 6, 5, 5, 5, 5, 6, 6, 6, 5}}, 
 {414, {1, 6, 6, 5, 6, 6, 5, 5, 5, 5, 5, 5, 6, 6, 6, 6, 6}}, 
 {413, {4, 5, 5, 5, 5, 5, 5, 5, 4, 5, 5, 5, 5, 6, 6, 5, 5, 5}}, 
 {412, {4, 5, 5, 5, 5, 5, 5, 4, 5, 5, 5, 5, 5, 6, 6, 5, 5, 5}}, 
 {412, {4, 5, 5, 6, 5, 5, 5, 6, 6, 5, 5, 5, 5, 6, 6, 6, 5}}, 
 {412, {4, 5, 5, 5, 5, 6, 6, 6, 5, 5, 6, 6, 5, 5, 5, 6, 5}}, 
 {412, {4, 5, 5, 5, 5, 6, 6, 5, 6, 6, 6, 5, 5, 5, 5, 6, 5}}, 
 {411, {4, 5, 5, 5, 6, 6, 5, 5, 5, 6, 6, 6, 5, 5, 6, 5, 5}}, 
 {411, {4, 5, 5, 5, 6, 5, 5, 6, 6, 5, 5, 6, 6, 6, 5, 5, 5}}, 
 {410, {4, 5, 5, 5, 6, 6, 6, 5, 5, 6, 6, 5, 5, 5, 6, 5, 5}}, 
 {410, {4, 4, 5, 5, 5, 5, 5, 6, 6, 6, 5, 6, 6, 6, 6, 5, 5}}, 
 {409, {4, 5, 5, 5, 6, 6, 5, 5, 6, 6, 6, 5, 5, 5, 6, 5, 5}}, 
 {407, {1, 6, 6, 5, 5, 5, 5, 5, 6, 6, 5, 5, 6, 6, 6, 6, 6}}, 
 {404, {1, 6, 6, 5, 5, 5, 5, 6, 5, 5, 5, 6, 6, 6, 6, 6, 6}}, 
 {403, {4, 5, 5, 5, 5, 4, 5, 5, 5, 5, 5, 5, 5, 6, 6, 5, 5, 5}}, 
 {403, {4, 5, 5, 5, 5, 6, 6, 5, 6, 6, 5, 5, 5, 6, 6, 5, 5}}, 
 {401, {4, 5, 6, 5, 5, 5, 5, 5, 5, 6, 6, 6, 6, 5, 5, 6, 5}}, 
 {401, {4, 5, 5, 5, 6, 6, 5, 5, 6, 5, 5, 6, 6, 6, 5, 5, 5}}, 
 {400, {4, 5, 5, 5, 6, 6, 6, 6, 5, 5, 4, 5, 5, 6, 6, 6, 5}}, 
 {400, {4, 5, 5, 5, 6, 5, 5, 5, 6, 6, 5, 5, 5, 6, 6, 6, 5}}, 
 {400, {3, 5, 5, 5, 5, 5, 6, 6, 6, 6, 6, 6, 6, 5, 5, 5, 5}}, 
 {399, {4, 5, 6, 5, 5, 5, 5, 5, 6, 5, 5, 5, 6, 6, 6, 6, 5}}, 
 {397, {4, 5, 5, 5, 5, 5, 6, 6, 6, 5, 5, 6, 6, 5, 5, 6, 5}}, 
 {397, {3, 5, 5, 6, 6, 6, 6, 6, 6, 5, 5, 5, 5, 5, 5, 6, 5}}, 
 {396, {4, 5, 6, 6, 5, 5, 5, 5, 6, 5, 5, 5, 5, 6, 6, 6, 5}}, 
 {396, {4, 5, 5, 6, 5, 5, 5, 5, 5, 6, 6, 6, 6, 5, 5, 6, 5}}, 
 {396, {4, 5, 5, 5, 6, 6, 6, 6, 5, 5, 6, 5, 5, 5, 5, 6, 5}}, 
 {396, {4, 5, 5, 5, 6, 5, 5, 6, 6, 6, 5, 5, 5, 5, 6, 6, 5}}, 
 {396, {3, 5, 5, 5, 5, 5, 5, 5, 5, 6, 6, 6, 6, 6, 6, 6, 5}}, 
 {394, {4, 4, 5, 5, 5, 5, 6, 6, 5, 5, 5, 6, 6, 6, 6, 6, 5}}, 
 {393, {4, 5, 5, 5, 6, 5, 5, 6, 6, 6, 5, 5, 6, 6, 5, 5, 5}}, 
 {393, {4, 4, 5, 5, 5, 5, 5, 6, 6, 5, 6, 6, 6, 6, 6, 5, 5}}, 
 {392, {4, 5, 5, 5, 6, 6, 6, 6, 6, 5, 5, 4, 5, 5, 6, 6, 5}}, 
 {392, {4, 5, 5, 5, 6, 6, 6, 6, 5, 5, 5, 6, 5, 5, 5, 6, 5}}, 
 {388, {4, 5, 5, 5, 6, 6, 5, 5, 6, 5, 5, 5, 5, 6, 6, 6, 5}}, 
 {386, {4, 4, 5, 5, 5, 5, 5, 5, 5, 6, 6, 5, 5, 5, 5, 5, 5, 5}}, 
 {386, {4, 5, 6, 5, 5, 5, 5, 5, 5, 6, 6, 6, 5, 5, 6, 6, 5}}, 
 {386, {4, 5, 5, 6, 5, 5, 6, 6, 6, 5, 5, 5, 5, 5, 6, 6, 5}}, 
 {386, {1, 6, 6, 5, 5, 6, 5, 5, 5, 5, 5, 6, 6, 6, 6, 6, 6}}, 
 {385, {4, 5, 5, 5, 5, 6, 5, 5, 5, 6, 6, 5, 6, 6, 6, 5, 5}}, 
 {384, {4, 5, 5, 6, 6, 5, 5, 4, 5, 5, 6, 6, 6, 6, 6, 5, 5}}, 
 {383, {4, 5, 5, 5, 6, 6, 5, 5, 5, 6, 6, 5, 5, 5, 6, 6, 5}}, 
 {382, {4, 4, 5, 6, 6, 5, 5, 5, 5, 5, 5, 6, 6, 6, 6, 6, 5}}, 
 {381, {4, 4, 5, 5, 5, 6, 6, 6, 6, 6, 6, 5, 5, 5, 6, 5, 5}}, 
 {380, {4, 5, 5, 5, 5, 6, 5, 5, 6, 6, 6, 5, 5, 6, 6, 5, 5}}, 
 {379, {4, 5, 5, 5, 6, 6, 6, 6, 6, 5, 5, 4, 5, 6, 6, 5, 5}}, 
 {378, {4, 4, 5, 5, 5, 5, 6, 6, 6, 6, 5, 6, 6, 6, 5, 5, 5}}, 
 {377, {4, 5, 5, 5, 6, 6, 6, 5, 5, 5, 5, 6, 5, 5, 6, 6, 5}}, 
 {377, {4, 5, 5, 5, 6, 6, 5, 5, 4, 5, 5, 6, 6, 6, 6, 6, 5}}, 
 {377, {3, 5, 5, 6, 6, 6, 6, 5, 5, 5, 5, 5, 5, 6, 6, 6, 5}}, 
 {376, {4, 4, 5, 5, 5, 5, 6, 6, 6, 5, 6, 6, 6, 6, 5, 5, 5}}, 
 {375, {3, 5, 6, 5, 5, 5, 5, 5, 5, 5, 6, 6, 6, 6, 6, 6, 5}}, 
 {374, {4, 5, 5, 6, 5, 5, 5, 5, 5, 6, 6, 5, 5, 6, 6, 6, 5}}, 
 {374, {4, 5, 5, 5, 6, 6, 5, 5, 4, 5, 6, 6, 6, 6, 6, 5, 5}}, 
 {374, {4, 4, 5, 5, 5, 5, 5, 6, 6, 6, 6, 6, 6, 5, 5, 6, 5}}, 
 {373, {3, 5, 6, 6, 6, 5, 5, 5, 5, 5, 5, 5, 6, 6, 6, 6, 5}}, 
 {372, {4, 5, 6, 5, 5, 5, 6, 5, 5, 5, 5, 5, 6, 6, 6, 6, 5}}, 
 {372, {4, 5, 5, 5, 6, 6, 5, 5, 5, 5, 5, 6, 6, 5, 6, 6, 5}}, 
 {371, {4, 5, 5, 5, 6, 5, 5, 5, 6, 6, 6, 5, 6, 6, 5, 5, 5}}, 
 {370, {4, 5, 6, 5, 5, 5, 5, 5, 5, 6, 6, 5, 5, 6, 6, 6, 5}}, 
 {369, {4, 5, 5, 5, 6, 5, 5, 5, 6, 6, 6, 5, 5, 5, 6, 6, 5}}, 
 {369, {4, 5, 5, 5, 5, 6, 6, 6, 5, 6, 6, 5, 5, 5, 5, 6, 5}}, 
 {369, {1, 6, 6, 6, 6, 6, 5, 5, 4, 5, 5, 5, 6, 6, 6, 6, 6}}, 
 {367, {4, 5, 6, 6, 5, 5, 5, 5, 5, 6, 6, 5, 5, 5, 6, 6, 5}}, 
 {367, {4, 4, 5, 5, 5, 5, 6, 6, 6, 6, 6, 5, 6, 6, 5, 5, 5}}, 
 {367, {1, 6, 6, 6, 6, 5, 5, 5, 4, 5, 5, 6, 6, 6, 6, 6, 6}}, 
 {365, {4, 5, 5, 5, 6, 6, 6, 6, 6, 5, 4, 5, 5, 6, 6, 5, 5}}, 
 {364, {4, 5, 5, 5, 6, 5, 5, 5, 5, 6, 6, 6, 5, 5, 6, 6, 5}}, 
 {363, {4, 5, 5, 4, 5, 5, 6, 6, 5, 5, 6, 6, 6, 6, 6, 5, 5}}, 
 {363, {3, 5, 5, 6, 6, 6, 5, 5, 5, 5, 5, 5, 6, 6, 6, 6, 5}}, 
 {363, {1, 6, 6, 5, 5, 5, 5, 5, 5, 6, 6, 6, 6, 5, 6, 6, 6}}, 
 {362, {4, 5, 5, 5, 6, 6, 6, 5, 6, 6, 5, 5, 5, 5, 6, 5, 5}}, 
 {359, {4, 5, 6, 5, 5, 6, 6, 6, 5, 5, 5, 5, 5, 5, 6, 6, 5}}, 
 {359, {4, 5, 5, 5, 6, 6, 6, 5, 5, 5, 6, 5, 5, 5, 6, 6, 5}}, 
 {359, {4, 5, 5, 5, 6, 5, 5, 5, 6, 5, 5, 6, 6, 6, 6, 5, 5}}, 
 {359, {4, 5, 5, 5, 5, 6, 5, 5, 5, 6, 6, 6, 5, 6, 6, 5, 5}}, 
 {359, {4, 5, 5, 4, 5, 5, 6, 6, 6, 5, 5, 6, 6, 6, 6, 5, 5}}, 
 {358, {1, 6, 6, 5, 5, 5, 6, 5, 5, 5, 5, 6, 6, 6, 6, 6, 6}}, 
 {357, {4, 5, 6, 6, 5, 5, 5, 5, 5, 6, 5, 5, 5, 6, 6, 6, 5}}, 
 {357, {4, 4, 5, 5, 5, 5, 6, 5, 5, 6, 6, 6, 6, 6, 6, 5, 5}}, 
 {357, {1, 6, 6, 6, 5, 5, 4, 5, 5, 5, 6, 6, 6, 6, 6, 6, 6}}, 
 {355, {4, 5, 5, 4, 5, 5, 5, 5, 5, 6, 6, 6, 6, 6, 6, 6, 5}}, 
 {354, {4, 5, 5, 6, 5, 5, 6, 6, 5, 5, 5, 5, 5, 6, 6, 6, 5}}, 
 {354, {4, 5, 5, 5, 6, 5, 5, 5, 6, 6, 5, 6, 6, 6, 5, 5, 5}}, 
 {353, {4, 5, 5, 5, 5, 5, 5, 5, 5, 5, 5, 5, 4, 5, 5, 6, 6, 5}}, 
 {353, {4, 5, 5, 4, 5, 5, 5, 5, 5, 5, 5, 6, 6, 5, 5, 5, 5, 5}}, 
 {353, {4, 5, 5, 5, 5, 6, 6, 6, 6, 5, 5, 5, 6, 5, 5, 6, 5}}, 
 {352, {3, 5, 5, 6, 6, 5, 5, 5, 5, 5, 6, 6, 6, 6, 6, 5, 5}}, 
 {351, {4, 5, 5, 4, 5, 5, 5, 5, 5, 5, 5, 5, 5, 6, 6, 5, 5, 5}}, 
 {351, {4, 5, 6, 5, 5, 5, 5, 5, 5, 5, 6, 6, 6, 5, 6, 6, 5}}, 
 {351, {4, 5, 5, 5, 6, 6, 5, 6, 6, 6, 5, 5, 5, 5, 6, 5, 5}}, 
 {351, {4, 5, 5, 5, 5, 5, 5, 6, 6, 5, 6, 6, 5, 6, 6, 5, 5}}, 
 {351, {1, 6, 6, 5, 5, 5, 5, 5, 5, 6, 6, 6, 5, 6, 6, 6, 6}}, 
 {350, {4, 5, 5, 5, 5, 6, 6, 5, 6, 6, 5, 5, 6, 6, 5, 5, 5}}, 
 {350, {3, 5, 5, 5, 5, 5, 5, 6, 6, 6, 6, 6, 6, 6, 5, 5, 5}}, 
 {349, {4, 5, 5, 5, 6, 6, 6, 5, 5, 6, 5, 5, 5, 5, 6, 6, 5}}, 
 {349, {5, 5, 6, 6, 5, 6, 6, 6, 5, 6, 6, 5, 5, 6, 6, 6}}, 
 {347, {4, 5, 5, 5, 6, 6, 6, 6, 5, 5, 5, 6, 5, 5, 6, 5, 5}}, 
 {347, {4, 5, 5, 5, 5, 5, 6, 6, 5, 6, 6, 6, 5, 5, 5, 6, 5}}, 
 {347, {4, 4, 5, 5, 5, 5, 5, 6, 6, 6, 5, 5, 6, 6, 6, 6, 5}}, 
 {347, {1, 6, 6, 6, 6, 5, 5, 4, 5, 5, 5, 6, 6, 6, 6, 6, 6}}, 
 {346, {4, 5, 5, 6, 5, 5, 5, 5, 5, 5, 6, 6, 6, 5, 6, 6, 5}}, 
 {346, {4, 5, 5, 5, 6, 6, 5, 5, 5, 6, 5, 5, 5, 6, 6, 6, 5}}, 
 {346, {4, 5, 5, 5, 5, 5, 6, 6, 5, 5, 6, 6, 5, 5, 6, 6, 5}}, 
 {345, {4, 5, 5, 5, 5, 6, 6, 6, 5, 5, 6, 6, 5, 5, 6, 5, 5}}, 
 {343, {4, 4, 5, 5, 5, 5, 5, 6, 6, 6, 6, 6, 5, 5, 6, 6, 5}}, 
 {342, {4, 5, 5, 5, 6, 6, 5, 6, 6, 5, 5, 5, 5, 5, 6, 6, 5}}, 
 {342, {4, 5, 5, 5, 5, 6, 6, 5, 5, 6, 6, 5, 6, 6, 5, 5, 5}}, 
 {341, {4, 4, 5, 5, 5, 5, 5, 6, 6, 5, 5, 6, 6, 6, 6, 6, 5}}, 
 {340, {5, 5, 5, 6, 6, 6, 5, 6, 6, 5, 6, 6, 5, 6, 6, 6}}, 
 {339, {4, 5, 5, 5, 4, 5, 5, 6, 6, 6, 5, 6, 6, 6, 6, 5, 5}}, 
 {337, {4, 5, 5, 5, 5, 6, 6, 5, 5, 5, 5, 6, 6, 5, 6, 6, 5}}, 
 {337, {4, 5, 5, 5, 5, 5, 6, 6, 6, 5, 6, 6, 5, 5, 5, 6, 5}}, 
 {336, {3, 5, 5, 5, 5, 5, 5, 5, 5, 5, 5, 5, 5, 5, 6, 6, 5, 5}}, 
 {334, {5, 5, 6, 6, 5, 5, 6, 6, 6, 5, 6, 6, 5, 6, 6, 6}}, 
 {334, {4, 4, 5, 6, 6, 6, 6, 6, 6, 6, 6, 6, 6, 6, 6, 5}}, 
 {333, {4, 5, 5, 5, 4, 5, 5, 5, 5, 5, 5, 5, 5, 5, 5, 6, 6, 5}}, 
 {333, {4, 5, 5, 5, 6, 6, 6, 6, 6, 6, 5, 5, 4, 5, 6, 5, 5}}, 
 {333, {4, 5, 5, 5, 6, 6, 5, 4, 5, 5, 6, 6, 6, 6, 6, 5, 5}}, 
 {333, {4, 5, 5, 5, 5, 5, 6, 5, 5, 6, 6, 5, 6, 6, 6, 5, 5}}, 
 {332, {4, 5, 5, 5, 5, 6, 6, 6, 5, 6, 6, 5, 5, 6, 5, 5, 5}}, 
 {331, {4, 5, 5, 5, 5, 5, 4, 5, 5, 5, 5, 5, 6, 6, 5, 5, 5, 5}}, 
 {331, {5, 5, 6, 6, 5, 6, 6, 5, 5, 6, 6, 6, 5, 6, 6, 6}}, 
 {330, {4, 5, 5, 5, 5, 5, 5, 6, 6, 6, 5, 6, 6, 5, 5, 6, 5}}, 
 {330, {4, 4, 5, 5, 5, 5, 6, 5, 5, 5, 6, 6, 6, 6, 6, 6, 5}}, 
 {330, {4, 4, 5, 5, 5, 5, 5, 6, 6, 6, 6, 6, 5, 6, 6, 5, 5}}, 
 {330, {4, 4, 5, 5, 5, 5, 5, 6, 6, 6, 6, 5, 5, 6, 6, 6, 5}}, 
 {329, {4, 5, 5, 5, 6, 5, 5, 5, 5, 5, 6, 6, 6, 5, 6, 6, 5}}, 
 {328, {4, 5, 5, 6, 5, 5, 5, 5, 6, 6, 6, 6, 5, 5, 6, 5, 5}}, 
 {327, {1, 6, 6, 6, 5, 5, 5, 4, 5, 5, 6, 6, 6, 6, 6, 6, 6}}, 
 {326, {4, 4, 5, 5, 5, 5, 5, 5, 6, 6, 6, 6, 6, 5, 6, 6, 5}}, 
 {325, {4, 5, 5, 5, 6, 5, 5, 5, 5, 6, 6, 5, 5, 6, 6, 6, 5}}, 
 {324, {4, 5, 6, 5, 5, 6, 5, 5, 5, 5, 5, 5, 6, 6, 6, 6, 5}}, 
 {324, {4, 5, 5, 6, 5, 5, 5, 6, 5, 5, 5, 5, 6, 6, 6, 6, 5}}, 
 {323, {5, 5, 5, 5, 6, 5, 5, 6, 5, 5, 6, 5, 5, 5, 5, 6, 6}}, 
 {322, {4, 5, 5, 5, 5, 5, 6, 6, 5, 5, 6, 6, 6, 5, 5, 6, 5}}, 
 {321, {4, 5, 5, 5, 6, 6, 5, 5, 5, 5, 6, 5, 5, 6, 6, 6, 5}}, 
 {320, {4, 5, 5, 5, 5, 6, 6, 6, 5, 5, 5, 6, 5, 5, 6, 6, 5}}, 
 {320, {4, 5, 5, 5, 5, 6, 6, 5, 5, 6, 6, 5, 5, 5, 6, 6, 5}}, 
 {320, {3, 5, 6, 6, 5, 5, 5, 5, 5, 5, 5, 6, 6, 6, 6, 6, 5}}, 
 {320, {5, 5, 6, 6, 5, 6, 6, 6, 5, 5, 6, 6, 5, 6, 6, 6}}, 
 {317, {4, 5, 6, 5, 5, 5, 5, 5, 5, 6, 5, 5, 6, 6, 6, 6, 5}}, 
 {317, {4, 5, 5, 6, 5, 5, 6, 5, 5, 5, 5, 5, 6, 6, 6, 6, 5}}, 
 {317, {4, 5, 5, 5, 5, 5, 6, 5, 5, 6, 6, 6, 5, 6, 6, 5, 5}}, 
 {317, {3, 5, 5, 5, 5, 6, 6, 6, 6, 5, 5, 5, 6, 6, 6, 5, 5}}, 
 {316, {5, 5, 6, 6, 5, 6, 6, 6, 5, 5, 6, 6, 6, 5, 6, 6}}, 
 {315, {4, 5, 6, 6, 5, 5, 6, 5, 5, 5, 5, 5, 5, 6, 6, 6, 5}}, 
 {314, {4, 4, 5, 5, 5, 6, 5, 5, 6, 6, 6, 6, 6, 6, 5, 5, 5}}, 
 {313, {4, 5, 5, 5, 5, 6, 6, 5, 5, 5, 6, 6, 5, 5, 6, 6, 5}}, 
 {313, {4, 5, 5, 5, 5, 5, 6, 6, 6, 5, 6, 6, 5, 5, 6, 5, 5}}, 
 {313, {1, 6, 6, 5, 5, 5, 5, 5, 6, 6, 6, 6, 6, 5, 5, 6, 6}}, 
 {311, {4, 5, 5, 6, 5, 5, 5, 5, 6, 5, 5, 5, 6, 6, 6, 6, 5}}, 
 {310, {4, 5, 6, 5, 5, 5, 5, 5, 5, 5, 6, 6, 5, 6, 6, 6, 5}}, 
 {310, {4, 5, 5, 5, 5, 6, 5, 5, 6, 6, 5, 5, 5, 6, 6, 6, 5}}, 
 {309, {1, 6, 6, 5, 5, 5, 5, 5, 5, 6, 6, 6, 6, 6, 5, 6, 6}}, 
 {308, {4, 5, 6, 6, 5, 5, 5, 5, 5, 5, 6, 6, 5, 5, 6, 6, 5}}, 
 {308, {4, 5, 5, 5, 5, 6, 6, 5, 5, 6, 5, 5, 5, 6, 6, 6, 5}}, 
 {308, {1, 6, 6, 5, 5, 5, 5, 5, 5, 6, 6, 5, 6, 6, 6, 6, 6}}, 
 {307, {4, 5, 5, 5, 6, 6, 6, 6, 6, 6, 5, 5, 4, 5, 5, 6, 5}}, 
 {307, {4, 5, 5, 5, 5, 5, 6, 6, 5, 5, 5, 6, 6, 5, 6, 6, 5}}, 
 {306, {4, 4, 5, 5, 5, 6, 5, 5, 5, 5, 6, 6, 6, 6, 6, 6, 5}}, 
 {305, {4, 5, 5, 5, 5, 5, 5, 5, 5, 5, 5, 4, 5, 5, 6, 6, 5, 5}}, 
 {305, {4, 5, 6, 5, 5, 5, 5, 6, 6, 6, 6, 5, 5, 5, 5, 6, 5}}, 
 {305, {4, 5, 5, 5, 5, 6, 6, 6, 6, 5, 5, 6, 5, 5, 5, 6, 5}}, 
 {304, {4, 5, 5, 5, 5, 4, 5, 5, 5, 5, 5, 5, 5, 5, 5, 6, 6, 5}}, 
 {304, {4, 5, 5, 5, 5, 6, 5, 5, 6, 5, 5, 6, 6, 6, 6, 5, 5}}, 
 {303, {4, 5, 5, 5, 5, 6, 6, 5, 6, 6, 5, 5, 5, 5, 6, 6, 5}}, 
 {303, {4, 5, 5, 4, 5, 5, 6, 6, 6, 5, 5, 5, 6, 6, 6, 6, 5}}, 
 {302, {4, 5, 6, 6, 5, 5, 5, 6, 5, 5, 5, 5, 5, 6, 6, 6, 5}}, 
 {302, {4, 5, 5, 5, 5, 6, 5, 5, 5, 6, 6, 6, 6, 5, 5, 6, 5}}, 
 {302, {4, 4, 5, 5, 5, 5, 6, 6, 5, 6, 6, 6, 6, 6, 5, 5, 5}}, 
 {299, {4, 5, 5, 5, 6, 5, 5, 6, 6, 6, 6, 5, 5, 5, 6, 5, 5}}, 
 {299, {3, 5, 5, 6, 6, 5, 5, 5, 5, 5, 5, 6, 6, 6, 6, 6, 5}}, 
 {298, {4, 5, 5, 5, 5, 6, 6, 6, 5, 6, 6, 5, 5, 5, 6, 5, 5}}, 
 {297, {4, 5, 5, 5, 5, 6, 6, 6, 6, 5, 4, 5, 6, 6, 6, 5, 5}}, 
 {297, {3, 5, 5, 6, 6, 6, 6, 6, 5, 5, 5, 5, 5, 5, 6, 6, 5}}, 
 {297, {5, 5, 6, 6, 5, 6, 6, 5, 6, 6, 5, 5, 6, 6, 6, 6}}, 
 {296, {4, 5, 5, 5, 5, 6, 5, 5, 6, 6, 5, 6, 6, 6, 5, 5, 5}}, 
 {296, {4, 5, 5, 5, 5, 5, 6, 6, 6, 6, 5, 5, 6, 5, 5, 6, 5}}, 
 {294, {4, 5, 5, 5, 5, 5, 5, 5, 5, 4, 5, 5, 5, 5, 5, 6, 6, 5}}, 
 {294, {4, 5, 5, 5, 5, 6, 6, 6, 6, 5, 4, 5, 5, 6, 6, 6, 5}}, 
 {293, {3, 5, 5, 5, 5, 5, 5, 5, 5, 5, 5, 6, 6, 5, 5, 5, 5, 5}}, 
 {293, {4, 5, 5, 5, 4, 5, 5, 6, 6, 6, 5, 5, 6, 6, 6, 6, 5}}, 
 {293, {4, 5, 6, 6, 6, 6, 5, 6, 6, 5, 6, 6, 6, 6, 6, 5}}, 
 {292, {4, 5, 5, 5, 6, 5, 5, 6, 5, 5, 5, 5, 6, 6, 6, 6, 5}}, 
 {291, {4, 5, 5, 5, 5, 5, 5, 5, 5, 5, 5, 4, 5, 5, 5, 6, 6, 5}}, 
 {291, {3, 5, 5, 5, 5, 5, 5, 5, 5, 5, 6, 6, 5, 5, 5, 5, 5, 5}}, 
 {291, {4, 5, 5, 5, 5, 5, 5, 6, 6, 5, 5, 6, 6, 5, 6, 6, 5}}, 
 {291, {3, 5, 5, 5, 5, 5, 6, 6, 6, 6, 5, 5, 5, 6, 6, 6, 5}}, 
 {290, {4, 5, 5, 4, 5, 5, 5, 5, 5, 5, 5, 5, 6, 6, 5, 5, 5, 5}}, 
 {290, {4, 4, 5, 5, 5, 5, 5, 5, 6, 6, 6, 5, 6, 6, 6, 6, 5}}, 
 {287, {5, 5, 6, 6, 5, 5, 6, 6, 5, 6, 6, 6, 6, 5, 6, 6}}, 
 {286, {4, 5, 5, 5, 5, 5, 5, 4, 5, 5, 5, 5, 5, 5, 5, 6, 6, 5}}, 
 {286, {4, 5, 5, 6, 5, 5, 5, 5, 5, 5, 6, 6, 5, 6, 6, 6, 5}}, 
 {286, {4, 5, 5, 5, 6, 5, 5, 6, 5, 5, 6, 6, 6, 6, 5, 5, 5}}, 
 {286, {4, 5, 5, 5, 6, 5, 5, 5, 6, 6, 6, 6, 5, 5, 6, 5, 5}}, 
 {286, {4, 5, 5, 5, 5, 6, 6, 5, 6, 6, 6, 5, 5, 5, 6, 5, 5}}, 
 {286, {4, 5, 5, 5, 5, 6, 6, 5, 5, 6, 6, 6, 5, 5, 6, 5, 5}}, 
 {286, {4, 5, 5, 5, 5, 6, 5, 5, 6, 6, 6, 6, 5, 5, 5, 6, 5}}, 
 {285, {4, 5, 5, 5, 5, 5, 6, 6, 5, 6, 6, 5, 6, 6, 5, 5, 5}}, 
 {285, {3, 5, 5, 5, 6, 6, 6, 6, 6, 6, 5, 5, 5, 5, 5, 6, 5}}, 
 {282, {3, 5, 5, 5, 5, 6, 6, 6, 6, 5, 5, 5, 5, 6, 6, 6, 5}}, 
 {282, {1, 6, 6, 5, 5, 5, 5, 5, 6, 5, 5, 6, 6, 6, 6, 6, 6}}, 
 {282, {4, 5, 6, 6, 5, 6, 6, 6, 5, 6, 6, 6, 6, 6, 6, 5}}, 
 {281, {4, 5, 5, 5, 5, 5, 5, 5, 4, 5, 5, 5, 5, 5, 5, 6, 6, 5}}, 
 {281, {4, 5, 5, 5, 4, 5, 5, 6, 6, 6, 6, 6, 5, 5, 6, 6, 5}}, 
 {281, {4, 4, 5, 5, 4, 5, 5, 6, 6, 6, 6, 6, 6, 6, 6, 5, 5}}, 
 {281, {3, 5, 5, 5, 5, 5, 6, 6, 6, 6, 6, 6, 5, 5, 5, 6, 5}}, 
 {280, {4, 5, 5, 5, 5, 6, 5, 5, 5, 6, 6, 5, 5, 6, 6, 6, 5}}, 
 {280, {4, 5, 5, 5, 5, 5, 5, 6, 6, 5, 6, 6, 6, 5, 5, 6, 5}}, 
 {280, {4, 4, 5, 5, 5, 5, 6, 6, 6, 6, 6, 6, 5, 5, 6, 5, 5}}, 
 {279, {4, 5, 5, 5, 5, 6, 6, 5, 5, 5, 6, 5, 5, 6, 6, 6, 5}}, 
 {278, {4, 5, 5, 4, 5, 5, 6, 6, 6, 6, 5, 5, 5, 6, 6, 6, 5}}, 
 {277, {4, 5, 6, 6, 5, 5, 5, 5, 5, 5, 5, 6, 6, 5, 6, 6, 5}}, 
 {277, {3, 5, 5, 5, 5, 6, 6, 6, 6, 6, 6, 5, 5, 5, 5, 6, 5}}, 
 {276, {4, 5, 5, 4, 5, 5, 6, 6, 5, 5, 5, 6, 6, 6, 6, 6, 5}}, 
 {275, {4, 5, 5, 5, 5, 5, 4, 5, 5, 5, 5, 5, 5, 5, 5, 6, 6, 5}}, 
 {275, {4, 5, 5, 5, 5, 6, 6, 6, 5, 5, 6, 5, 5, 5, 6, 6, 5}}, 
 {274, {4, 5, 6, 6, 6, 5, 6, 6, 6, 5, 6, 6, 6, 6, 6, 5}}, 
 {273, {4, 5, 5, 5, 5, 6, 6, 6, 6, 5, 5, 6, 5, 5, 6, 5, 5}}, 
 {273, {4, 5, 5, 5, 5, 6, 5, 5, 5, 6, 6, 6, 5, 5, 6, 6, 5}}, 
 {273, {1, 6, 6, 5, 5, 4, 5, 5, 5, 6, 6, 6, 6, 6, 6, 6, 6}}, 
 {271, {4, 4, 5, 5, 5, 5, 5, 5, 6, 6, 6, 6, 5, 6, 6, 6, 5}}, 
 {271, {3, 5, 5, 5, 6, 6, 6, 5, 5, 5, 5, 6, 6, 6, 6, 5, 5}}, 
 {271, {5, 5, 6, 5, 5, 6, 6, 5, 6, 6, 6, 6, 6, 5, 6, 6}}, 
 {269, {4, 5, 5, 6, 5, 5, 5, 5, 5, 6, 5, 5, 6, 6, 6, 6, 5}}, 
 {269, {4, 5, 5, 5, 5, 6, 6, 6, 6, 5, 5, 4, 5, 6, 6, 6, 5}}, 
 {269, {4, 5, 5, 5, 5, 6, 6, 6, 5, 4, 5, 6, 6, 6, 6, 5, 5}}, 
 {269, {4, 5, 5, 5, 5, 5, 6, 6, 6, 5, 5, 6, 5, 5, 6, 6, 5}}, 
 {269, {4, 5, 5, 5, 4, 5, 5, 6, 6, 5, 6, 6, 6, 6, 6, 5, 5}}, 
 {269, {5, 5, 5, 6, 6, 5, 6, 6, 6, 5, 6, 6, 6, 5, 6, 6}}, 
 {268, {4, 5, 5, 5, 6, 5, 5, 5, 6, 5, 5, 5, 6, 6, 6, 6, 5}}, 
 {268, {4, 5, 5, 5, 5, 5, 5, 6, 6, 5, 6, 6, 5, 5, 6, 6, 5}}, 
 {268, {4, 5, 6, 6, 6, 5, 6, 6, 5, 6, 6, 6, 6, 6, 6, 5}}, 
 {267, {4, 5, 5, 5, 5, 6, 6, 6, 6, 6, 5, 4, 5, 6, 6, 5, 5}}, 
 {267, {4, 4, 5, 5, 6, 5, 5, 5, 5, 5, 6, 6, 6, 6, 6, 6, 5}}, 
 {267, {3, 5, 5, 5, 6, 6, 6, 6, 5, 5, 5, 5, 6, 6, 6, 5, 5}}, 
 {267, {3, 5, 5, 5, 5, 5, 6, 6, 6, 5, 5, 5, 6, 6, 6, 6, 5}}, 
 {266, {3, 5, 5, 5, 5, 6, 6, 6, 5, 5, 5, 6, 6, 6, 6, 5, 5}}, 
 {265, {4, 5, 5, 5, 6, 5, 5, 5, 5, 5, 6, 6, 5, 6, 6, 6, 5}}, 
 {265, {4, 5, 5, 5, 5, 6, 5, 5, 5, 5, 6, 6, 6, 5, 6, 6, 5}}, 
 {265, {4, 5, 5, 5, 4, 5, 5, 6, 6, 6, 6, 6, 6, 5, 5, 6, 5}}, 
 {264, {4, 4, 5, 5, 5, 5, 5, 6, 5, 5, 5, 5, 5, 5, 5, 5, 6, 5}}, 
 {263, {4, 4, 5, 5, 5, 5, 5, 5, 5, 5, 5, 6, 5, 5, 5, 5, 6, 5}}, 
 {263, {3, 5, 5, 5, 5, 6, 6, 6, 5, 5, 5, 5, 6, 6, 6, 6, 5}}, 
 {262, {4, 5, 5, 5, 5, 6, 6, 5, 6, 6, 6, 5, 5, 6, 5, 5, 5}}, 
 {261, {4, 5, 5, 5, 5, 5, 6, 5, 5, 5, 6, 6, 6, 5, 6, 6, 5}}, 
 {260, {4, 5, 5, 4, 5, 5, 5, 6, 6, 6, 6, 6, 6, 6, 5, 5, 5}}, 
 {259, {3, 5, 5, 5, 5, 5, 5, 5, 5, 6, 6, 5, 5, 5, 5, 5, 5, 5}}, 
 {259, {4, 5, 5, 5, 5, 6, 6, 6, 5, 5, 4, 5, 6, 6, 6, 6, 5}}, 
 {258, {4, 5, 5, 5, 5, 5, 6, 6, 5, 6, 6, 5, 5, 5, 6, 6, 5}}, 
 {257, {4, 5, 6, 6, 6, 5, 6, 6, 6, 6, 5, 6, 6, 6, 6, 5}}, 
 {256, {3, 5, 5, 5, 5, 5, 5, 5, 5, 5, 5, 5, 5, 5, 5, 6, 6, 5}}, 
 {256, {4, 5, 6, 6, 5, 5, 5, 5, 5, 5, 6, 5, 5, 6, 6, 6, 5}}, 
 {256, {4, 5, 5, 5, 6, 5, 5, 5, 5, 6, 5, 5, 6, 6, 6, 6, 5}}, 
 {256, {1, 6, 6, 6, 6, 6, 5, 4, 5, 5, 5, 5, 6, 6, 6, 6, 6}}, 
 {256, {4, 5, 6, 6, 5, 6, 6, 6, 6, 5, 6, 6, 6, 6, 6, 5}}, 
 {255, {4, 5, 5, 5, 5, 6, 6, 6, 5, 4, 5, 5, 6, 6, 6, 6, 5}}, 
 {255, {4, 5, 5, 5, 5, 5, 6, 6, 5, 5, 6, 5, 5, 6, 6, 6, 5}}, 
 {254, {4, 4, 5, 5, 5, 5, 5, 5, 5, 5, 6, 5, 5, 5, 5, 5, 6, 5}}, 
 {254, {4, 5, 5, 5, 5, 5, 6, 5, 5, 6, 6, 5, 5, 6, 6, 6, 5}}, 
 {253, {4, 4, 5, 5, 5, 5, 5, 5, 6, 5, 5, 5, 5, 5, 5, 5, 6, 5}}, 
 {253, {4, 4, 5, 5, 5, 5, 5, 5, 5, 6, 5, 5, 5, 5, 5, 5, 6, 5}}, 
 {253, {5, 5, 5, 5, 6, 5, 5, 5, 5, 6, 5, 5, 6, 6, 5, 5, 6}}, 
 {253, {4, 5, 5, 5, 5, 6, 5, 5, 6, 6, 6, 5, 6, 6, 5, 5, 5}}, 
 {253, {4, 5, 5, 5, 4, 5, 5, 6, 6, 5, 5, 6, 6, 6, 6, 6, 5}}, 
 {252, {4, 5, 5, 5, 5, 6, 5, 5, 6, 6, 6, 5, 5, 5, 6, 6, 5}}, 
 {251, {4, 5, 5, 5, 5, 5, 6, 6, 5, 6, 6, 6, 5, 5, 6, 5, 5}}, 
 {251, {4, 4, 5, 5, 5, 5, 5, 6, 5, 5, 6, 6, 6, 6, 6, 6, 5}}, 
 {250, {4, 5, 5, 5, 6, 5, 4, 5, 6, 6, 6, 6, 6, 6, 5, 5, 5}}, 
 {250, {3, 5, 5, 5, 5, 6, 6, 6, 6, 6, 5, 5, 5, 5, 6, 6, 5}}, 
 {249, {4, 4, 5, 5, 5, 5, 6, 5, 5, 5, 5, 5, 5, 5, 5, 5, 6, 5}}, 
 {249, {4, 5, 5, 5, 5, 6, 5, 4, 5, 6, 6, 6, 6, 6, 6, 5, 5}}, 
 {249, {4, 5, 5, 4, 5, 5, 5, 6, 6, 6, 6, 5, 6, 6, 6, 5, 5}}, 
 {248, {4, 4, 5, 5, 5, 5, 5, 6, 5, 5, 5, 5, 5, 6, 5, 5, 5, 5}}, 
 {248, {3, 5, 5, 5, 5, 5, 5, 6, 6, 6, 6, 6, 6, 5, 5, 6, 5}}, 
 {247, {4, 4, 5, 5, 5, 5, 5, 5, 6, 5, 5, 5, 5, 5, 6, 5, 5, 5}}, 
 {247, {3, 5, 5, 5, 5, 5, 5, 5, 5, 5, 5, 5, 5, 6, 6, 5, 5, 5}}, 
 {247, {4, 5, 5, 5, 6, 6, 5, 5, 6, 6, 5, 5, 6, 6, 5, 5, 5}}, 
 {246, {4, 4, 5, 5, 5, 4, 5, 5, 5, 6, 6, 6, 5, 5, 5, 5, 5, 5}}, 
 {246, {4, 5, 5, 5, 6, 6, 6, 5, 4, 5, 6, 6, 6, 6, 5, 5, 5}}, 
 {246, {4, 5, 5, 5, 4, 5, 5, 6, 6, 6, 6, 5, 5, 6, 6, 6, 5}}, 
 {246, {4, 5, 5, 4, 5, 5, 6, 6, 6, 6, 6, 6, 5, 5, 5, 6, 5}}, 
 {246, {4, 4, 5, 5, 5, 5, 5, 5, 6, 6, 5, 6, 6, 6, 6, 6, 5}}, 
 {245, {4, 4, 5, 5, 5, 5, 5, 5, 5, 5, 4, 5, 5, 6, 6, 6, 5, 5}}, 
 {245, {4, 5, 5, 5, 6, 6, 6, 6, 5, 5, 6, 5, 5, 5, 6, 5, 5}}, 
 {244, {1, 6, 6, 5, 5, 5, 5, 5, 5, 5, 5, 5, 5, 5, 5, 5, 6, 6}}, 
 {243, {3, 5, 5, 5, 5, 5, 6, 6, 6, 6, 5, 5, 6, 6, 6, 5, 5}}, 
 {242, {4, 4, 5, 5, 5, 5, 5, 5, 6, 5, 5, 5, 5, 6, 5, 5, 5, 5}}, 
 {242, {3, 5, 5, 5, 5, 5, 5, 5, 5, 5, 5, 5, 6, 6, 5, 5, 5, 5}}, 
 {242, {4, 5, 6, 6, 5, 6, 6, 6, 6, 6, 5, 6, 6, 6, 6, 5}}, 
 {241, {4, 4, 5, 5, 5, 5, 5, 4, 5, 5, 6, 6, 6, 5, 5, 5, 5, 5}}, 
 {241, {4, 5, 5, 5, 5, 6, 6, 6, 6, 6, 5, 4, 5, 5, 6, 6, 5}}, 
 {240, {3, 5, 5, 5, 6, 6, 6, 6, 5, 5, 5, 5, 5, 6, 6, 6, 5}}, 
 {239, {4, 5, 5, 5, 6, 5, 5, 4, 5, 5, 6, 6, 6, 6, 6, 6, 5}}, 
 {237, {4, 5, 5, 5, 5, 6, 5, 5, 6, 5, 5, 5, 6, 6, 6, 6, 5}}, 
 {237, {3, 5, 6, 5, 5, 5, 5, 6, 6, 6, 6, 6, 5, 5, 5, 6, 5}}, 
 {236, {4, 5, 5, 4, 5, 5, 6, 6, 6, 6, 6, 5, 5, 5, 6, 6, 5}}, 
 {235, {4, 5, 5, 5, 6, 5, 5, 4, 5, 6, 6, 6, 6, 6, 6, 5, 5}}, 
 {234, {4, 4, 5, 5, 5, 6, 5, 5, 5, 5, 5, 5, 5, 5, 5, 6, 5, 5}}, 
 {234, {5, 5, 5, 6, 5, 5, 5, 6, 5, 5, 5, 6, 5, 5, 5, 6, 6}}, 
 {234, {3, 5, 5, 5, 6, 6, 5, 5, 5, 5, 6, 6, 6, 6, 6, 5, 5}}, 
 {232, {4, 5, 5, 4, 5, 5, 5, 6, 6, 6, 5, 6, 6, 6, 6, 5, 5}}, 
 {232, {4, 4, 5, 5, 6, 6, 6, 5, 5, 5, 6, 6, 6, 5, 5, 6, 5}}, 
 {232, {3, 5, 5, 5, 5, 5, 6, 6, 6, 5, 5, 6, 6, 6, 6, 5, 5}}, 
 {230, {4, 4, 5, 5, 5, 4, 5, 5, 5, 5, 5, 5, 5, 6, 6, 6, 5, 5}}, 
 {230, {4, 5, 5, 5, 5, 5, 5, 5, 6, 6, 5, 6, 6, 5, 6, 6, 5}}, 
 {230, {4, 4, 5, 6, 5, 5, 5, 6, 6, 6, 6, 5, 5, 5, 6, 6, 5}}, 
 {228, {4, 5, 5, 5, 5, 6, 6, 6, 6, 6, 6, 5, 4, 5, 6, 5, 5}}, 
 {228, {4, 5, 5, 5, 5, 5, 6, 5, 5, 6, 6, 6, 6, 5, 5, 6, 5}}, 
 {228, {3, 5, 5, 5, 5, 6, 6, 6, 6, 6, 5, 5, 5, 6, 6, 5, 5}}, 
 {228, {3, 5, 5, 5, 5, 6, 6, 5, 5, 5, 6, 6, 6, 6, 6, 5, 5}}, 
 {227, {4, 5, 5, 5, 5, 5, 5, 5, 5, 5, 4, 5, 5, 5, 5, 6, 6, 5}}, 
 {227, {4, 4, 5, 5, 5, 4, 5, 5, 6, 6, 6, 5, 5, 5, 5, 5, 5, 5}}, 
 {227, {3, 5, 6, 5, 5, 5, 6, 6, 6, 5, 5, 5, 5, 6, 6, 6, 5}}, 
 {227, {3, 5, 5, 6, 6, 6, 5, 5, 5, 6, 6, 6, 5, 5, 5, 6, 5}}, 
 {225, {4, 4, 5, 5, 6, 6, 6, 5, 5, 6, 6, 6, 5, 5, 5, 6, 5}}, 
 {224, {4, 5, 5, 5, 5, 4, 5, 5, 5, 5, 5, 6, 6, 5, 5, 5, 5, 5}}, 
 {224, {4, 5, 5, 5, 5, 5, 6, 5, 5, 5, 6, 6, 5, 6, 6, 6, 5}}, 
 {224, {4, 4, 5, 5, 6, 6, 5, 5, 6, 6, 5, 5, 6, 6, 6, 5, 5}}, 
 {223, {4, 4, 5, 5, 5, 5, 5, 5, 5, 6, 5, 5, 5, 5, 6, 5, 5, 5}}, 
 {222, {4, 4, 5, 5, 5, 6, 5, 5, 5, 5, 5, 5, 5, 5, 5, 5, 6, 5}}, 
 {222, {4, 4, 5, 5, 5, 5, 4, 5, 5, 6, 6, 6, 5, 5, 5, 5, 5, 5}}, 
 {221, {4, 4, 5, 6, 5, 5, 6, 6, 6, 5, 5, 5, 5, 6, 6, 6, 5}}, 
 {221, {3, 5, 5, 5, 5, 5, 5, 6, 6, 6, 5, 5, 6, 6, 6, 6, 5}}, 
 {220, {3, 5, 5, 5, 5, 5, 6, 6, 6, 6, 6, 5, 5, 5, 6, 6, 5}}, 
 {220, {1, 6, 6, 5, 5, 5, 4, 5, 5, 6, 6, 6, 6, 6, 6, 6, 6}}, 
 {219, {4, 4, 5, 5, 5, 5, 5, 5, 5, 4, 5, 5, 6, 6, 6, 5, 5, 5}}, 
 {219, {4, 5, 5, 5, 5, 6, 6, 5, 5, 4, 5, 6, 6, 6, 6, 6, 5}}, 
 {219, {4, 5, 5, 5, 5, 5, 6, 5, 5, 6, 6, 6, 5, 5, 6, 6, 5}}, 
 {219, {3, 5, 6, 5, 5, 6, 6, 6, 5, 5, 5, 5, 5, 6, 6, 6, 5}}, 
 {219, {3, 5, 5, 5, 6, 6, 6, 6, 6, 5, 5, 5, 5, 6, 6, 5, 5}}, 
 {218, {1, 6, 6, 5, 5, 4, 5, 5, 5, 5, 5, 5, 5, 5, 5, 6, 6, 6}}, 
 {218, {4, 5, 5, 5, 6, 6, 5, 4, 5, 6, 6, 6, 6, 6, 5, 5, 5}}, 
 {218, {4, 5, 6, 6, 5, 6, 6, 5, 6, 6, 6, 6, 6, 6, 6, 5}}, 
 {217, {4, 5, 5, 5, 5, 5, 5, 5, 5, 4, 5, 5, 5, 6, 6, 5, 5, 5}}, 
 {217, {4, 4, 5, 5, 5, 5, 5, 5, 4, 5, 5, 6, 6, 6, 5, 5, 5, 5}}, 
 {217, {4, 5, 5, 5, 5, 6, 6, 6, 6, 6, 6, 5, 4, 5, 5, 6, 5}}, 
 {217, {1, 6, 6, 6, 5, 5, 5, 5, 4, 5, 6, 6, 6, 6, 6, 6, 6}}, 
 {216, {4, 4, 5, 5, 4, 5, 5, 6, 6, 6, 5, 5, 5, 5, 5, 5, 5, 5}}, 
 {216, {4, 4, 5, 6, 5, 5, 5, 5, 6, 6, 6, 6, 6, 5, 5, 6, 5}}, 
 {215, {4, 5, 5, 5, 5, 6, 5, 4, 5, 5, 6, 6, 6, 6, 6, 6, 5}}, 
 {215, {4, 5, 5, 4, 5, 5, 5, 6, 6, 5, 6, 6, 6, 6, 6, 5, 5}}, 
 {214, {4, 4, 5, 5, 5, 5, 5, 6, 5, 5, 5, 5, 5, 5, 6, 5, 5, 5}}, 
 {214, {4, 4, 5, 5, 5, 4, 5, 5, 5, 5, 5, 5, 6, 6, 6, 5, 5, 5}}, 
 {213, {4, 4, 5, 5, 5, 5, 5, 5, 5, 5, 6, 5, 5, 5, 5, 6, 5, 5}}, 
 {213, {4, 5, 5, 5, 5, 6, 5, 5, 6, 6, 6, 6, 5, 5, 6, 5, 5}}, 
 {213, {1, 6, 6, 6, 6, 5, 4, 5, 5, 5, 5, 6, 6, 6, 6, 6, 6}}, 
 {212, {4, 5, 5, 5, 5, 5, 5, 6, 5, 5, 6, 6, 5, 6, 6, 6, 5}}, 
 {212, {4, 4, 5, 5, 5, 4, 5, 5, 6, 6, 6, 6, 6, 6, 6, 6, 5}}, 
 {211, {4, 5, 5, 4, 5, 5, 5, 5, 5, 5, 5, 5, 5, 5, 5, 6, 6, 5}}, 
 {211, {4, 4, 5, 5, 5, 5, 5, 6, 5, 5, 5, 5, 5, 5, 5, 6, 5, 5}}, 
 {211, {4, 4, 5, 5, 5, 5, 4, 5, 5, 5, 5, 5, 5, 6, 6, 6, 5, 5}}, 
 {211, {4, 5, 5, 5, 6, 5, 4, 5, 5, 5, 6, 6, 6, 6, 6, 6, 5}}, 
 {211, {4, 5, 5, 5, 5, 6, 6, 6, 6, 6, 6, 5, 5, 4, 5, 6, 5}}, 
 {211, {3, 5, 5, 5, 6, 6, 6, 5, 5, 5, 5, 5, 6, 6, 6, 6, 5}}, 
 {211, {3, 5, 5, 5, 5, 6, 6, 5, 5, 5, 5, 6, 6, 6, 6, 6, 5}}, 
 {210, {4, 4, 5, 5, 5, 5, 5, 5, 6, 5, 5, 5, 6, 5, 5, 5, 5, 5}}, 
 {210, {4, 4, 5, 5, 5, 5, 5, 5, 5, 6, 5, 5, 5, 5, 5, 6, 5, 5}}, 
 {210, {1, 6, 6, 5, 5, 5, 5, 5, 5, 5, 5, 5, 4, 5, 5, 6, 6, 6}}, 
 {210, {3, 5, 6, 5, 5, 5, 5, 6, 6, 6, 5, 5, 5, 6, 6, 6, 5}}, 
 {210, {3, 5, 5, 5, 5, 5, 6, 6, 6, 6, 6, 5, 5, 6, 6, 5, 5}}, 
 {210, {3, 5, 5, 5, 5, 5, 6, 6, 5, 5, 5, 6, 6, 6, 6, 6, 5}}, 
 {209, {4, 4, 5, 5, 5, 5, 5, 5, 6, 5, 5, 5, 5, 5, 5, 6, 5, 5}}, 
 {209, {4, 5, 5, 5, 5, 6, 5, 5, 5, 5, 6, 6, 5, 6, 6, 6, 5}}, 
 {209, {4, 5, 6, 6, 5, 6, 6, 6, 6, 6, 6, 5, 6, 6, 6, 5}}, 
 {208, {4, 4, 5, 5, 5, 5, 6, 5, 5, 5, 5, 5, 5, 5, 6, 5, 5, 5}}, 
 {208, {4, 4, 5, 5, 5, 5, 4, 5, 5, 5, 6, 6, 6, 5, 5, 5, 5, 5}}, 
 {208, {3, 5, 5, 5, 5, 5, 5, 6, 6, 6, 6, 5, 6, 6, 6, 5, 5}}, 
 {207, {3, 5, 5, 5, 5, 5, 5, 6, 6, 6, 6, 5, 5, 6, 6, 6, 5}}, 
 {206, {4, 4, 5, 5, 5, 5, 5, 4, 5, 5, 5, 5, 5, 6, 6, 6, 5, 5}}, 
 {206, {4, 5, 5, 5, 5, 6, 6, 6, 6, 6, 5, 5, 4, 5, 6, 6, 5}}, 
 {206, {3, 5, 5, 5, 6, 6, 5, 5, 5, 5, 5, 6, 6, 6, 6, 6, 5}}, 
 {205, {4, 4, 5, 5, 5, 5, 6, 5, 5, 5, 5, 5, 5, 5, 5, 6, 5, 5}}, 
 {205, {4, 5, 5, 5, 4, 5, 6, 6, 6, 5, 5, 5, 6, 6, 6, 6, 5}}, 
 {205, {3, 5, 6, 6, 6, 6, 6, 5, 6, 6, 6, 6, 6, 6, 6, 5}}, 
 {204, {4, 4, 5, 5, 6, 5, 5, 5, 5, 5, 5, 5, 5, 5, 5, 5, 6, 5}}, 
 {203, {4, 4, 5, 5, 5, 5, 4, 5, 5, 5, 5, 6, 6, 6, 5, 5, 5, 5}}, 
 {203, {4, 4, 5, 5, 4, 5, 5, 5, 6, 6, 6, 5, 5, 5, 5, 5, 5, 5}}, 
 {203, {3, 5, 5, 5, 6, 6, 6, 6, 6, 5, 5, 5, 5, 5, 6, 6, 5}}, 
 {203, {1, 6, 6, 6, 5, 4, 5, 5, 5, 5, 6, 6, 6, 6, 6, 6, 6}}, 
 {203, {5, 5, 6, 5, 5, 6, 6, 6, 5, 6, 6, 6, 5, 6, 6, 6}}, 
 {202, {4, 4, 5, 5, 5, 5, 5, 5, 5, 5, 5, 5, 6, 5, 5, 5, 6, 5}}, 
 {202, {4, 5, 5, 4, 5, 5, 5, 6, 6, 6, 6, 6, 5, 6, 6, 5, 5}}, 
 {201, {4, 4, 5, 5, 6, 6, 6, 6, 5, 5, 5, 6, 6, 5, 5, 6, 5}}, 
 {200, {4, 5, 5, 4, 5, 5, 5, 5, 5, 5, 6, 6, 5, 5, 5, 5, 5, 5}}, 
 {200, {4, 4, 5, 5, 4, 5, 5, 5, 5, 5, 5, 6, 6, 6, 5, 5, 5, 5}}, 
 {199, {4, 5, 5, 5, 5, 5, 5, 6, 5, 5, 6, 6, 6, 5, 6, 6, 5}}, 
 {199, {3, 5, 6, 5, 5, 5, 5, 6, 6, 5, 5, 5, 6, 6, 6, 6, 5}}, 
 {199, {3, 5, 5, 6, 6, 6, 5, 5, 5, 5, 6, 6, 6, 5, 5, 6, 5}}, 
 {199, {3, 5, 6, 6, 6, 6, 5, 6, 6, 6, 6, 6, 6, 6, 6, 5}}, 
 {198, {4, 4, 5, 5, 5, 5, 5, 5, 5, 5, 6, 5, 5, 5, 6, 5, 5, 5}}, 
 {197, {4, 5, 5, 5, 5, 5, 6, 5, 5, 6, 5, 5, 6, 6, 6, 6, 5}}, 
 {197, {1, 6, 6, 6, 6, 5, 5, 5, 5, 4, 5, 6, 6, 6, 6, 6, 6}}, 
 {196, {4, 5, 5, 5, 5, 6, 6, 5, 4, 5, 5, 6, 6, 6, 6, 6, 5}}, 
 {196, {4, 4, 5, 6, 5, 5, 6, 6, 5, 5, 5, 5, 6, 6, 6, 6, 5}}, 
 {196, {4, 4, 5, 5, 6, 6, 5, 5, 6, 6, 6, 6, 5, 5, 5, 6, 5}}, 
 {195, {3, 5, 5, 5, 6, 5, 5, 5, 5, 6, 6, 6, 6, 6, 6, 5, 5}}, 
 {194, {4, 5, 5, 5, 6, 6, 6, 5, 4, 5, 5, 5, 6, 6, 6, 6, 5}}, 
 {194, {4, 5, 5, 5, 5, 6, 5, 5, 5, 6, 5, 5, 6, 6, 6, 6, 5}}, 
 {194, {4, 5, 5, 5, 5, 4, 5, 5, 6, 6, 6, 6, 6, 5, 6, 6, 5}}, 
 {194, {4, 5, 5, 5, 4, 5, 6, 5, 5, 5, 6, 6, 6, 6, 6, 6, 5}}, 
 {194, {4, 4, 5, 6, 6, 5, 5, 5, 5, 6, 6, 6, 5, 5, 6, 6, 5}}, 
 {194, {4, 4, 5, 6, 5, 5, 5, 6, 6, 6, 5, 5, 5, 6, 6, 6, 5}}, 
 {193, {4, 5, 5, 5, 5, 6, 6, 5, 4, 5, 6, 6, 6, 6, 6, 5, 5}}, 
 {193, {3, 5, 6, 5, 5, 5, 5, 6, 6, 6, 6, 5, 5, 5, 6, 6, 5}}, 
 {192, {4, 5, 5, 5, 5, 5, 6, 6, 6, 5, 4, 5, 6, 6, 6, 6, 5}}, 
 {192, {4, 5, 5, 5, 5, 4, 5, 5, 6, 6, 6, 5, 6, 6, 6, 6, 5}}, 
 {192, {3, 5, 6, 5, 5, 5, 5, 5, 6, 6, 6, 6, 6, 5, 5, 6, 5}}, 
 {191, {4, 5, 6, 6, 5, 5, 5, 5, 6, 6, 5, 5, 5, 5, 6, 6, 5}}, 
 {191, {4, 5, 5, 5, 4, 5, 5, 5, 6, 6, 6, 6, 5, 6, 6, 6, 5}}, 
 {191, {4, 4, 5, 5, 6, 6, 6, 5, 5, 5, 6, 6, 5, 5, 6, 6, 5}}, 
 {191, {4, 4, 5, 5, 6, 6, 5, 5, 5, 6, 6, 6, 6, 5, 5, 6, 5}}, 
 {191, {3, 5, 6, 6, 5, 6, 6, 6, 6, 6, 6, 6, 6, 6, 6, 5}}, 
 {190, {4, 4, 5, 5, 5, 4, 5, 5, 5, 5, 5, 6, 6, 6, 5, 5, 5, 5}}, 
 {190, {4, 5, 5, 6, 6, 5, 5, 5, 6, 5, 5, 6, 6, 5, 5, 6, 5}}, 
 {190, {4, 5, 5, 5, 6, 6, 5, 4, 5, 5, 5, 6, 6, 6, 6, 6, 5}}, 
 {190, {4, 4, 5, 6, 5, 5, 6, 6, 6, 6, 5, 5, 5, 5, 6, 6, 5}}, 
 {189, {4, 4, 5, 5, 5, 5, 5, 5, 5, 5, 5, 5, 5, 6, 5, 5, 6, 5}}, 
 {189, {4, 5, 5, 6, 6, 5, 5, 6, 6, 5, 5, 6, 5, 5, 5, 6, 5}}, 
 {189, {4, 5, 5, 5, 4, 5, 6, 6, 5, 5, 5, 6, 6, 6, 6, 6, 5}}, 
 {188, {4, 4, 5, 5, 6, 6, 6, 6, 5, 5, 6, 6, 5, 5, 5, 6, 5}}, 
 {188, {4, 4, 5, 5, 6, 6, 5, 5, 5, 6, 6, 6, 5, 6, 6, 5, 5}}, 
 {188, {3, 5, 5, 5, 5, 5, 5, 6, 6, 6, 6, 6, 5, 5, 6, 6, 5}}, 
 {187, {4, 4, 5, 5, 5, 5, 5, 5, 5, 5, 5, 6, 5, 5, 6, 5, 5, 5}}, 
 {187, {4, 5, 5, 6, 6, 5, 5, 6, 5, 5, 5, 6, 6, 5, 5, 6, 5}}, 
 {187, {4, 4, 5, 6, 6, 5, 5, 6, 6, 5, 5, 5, 5, 6, 6, 6, 5}}, 
 {187, {4, 4, 5, 6, 5, 5, 5, 5, 6, 6, 6, 6, 5, 5, 6, 6, 5}}, 
 {187, {4, 4, 5, 5, 6, 6, 6, 5, 5, 5, 5, 6, 6, 5, 6, 6, 5}}, 
 {186, {4, 4, 5, 5, 5, 5, 5, 5, 5, 4, 5, 5, 5, 6, 6, 6, 5, 5}}, 
 {186, {4, 4, 5, 5, 5, 5, 5, 4, 5, 5, 5, 5, 6, 6, 6, 5, 5, 5}}, 
 {186, {4, 4, 5, 5, 5, 4, 5, 5, 5, 5, 6, 6, 6, 5, 5, 5, 5, 5}}, 
 {186, {3, 5, 5, 5, 5, 5, 5, 5, 5, 5, 6, 5, 5, 5, 5, 5, 6, 5}}, 
 {186, {3, 5, 5, 6, 6, 6, 5, 5, 6, 6, 6, 5, 5, 5, 5, 6, 5}}, 
 {185, {4, 4, 5, 5, 5, 5, 5, 5, 4, 5, 5, 5, 6, 6, 6, 5, 5, 5}}, 
 {185, {4, 4, 5, 5, 4, 5, 5, 5, 5, 5, 5, 5, 5, 6, 6, 6, 5, 5}}, 
 {185, {4, 4, 5, 6, 5, 5, 5, 5, 6, 6, 6, 5, 5, 6, 6, 6, 5}}, 
 {185, {4, 4, 5, 5, 5, 4, 5, 6, 6, 6, 6, 6, 6, 6, 6, 5, 5}}, 
 {184, {4, 4, 5, 5, 5, 5, 4, 5, 5, 5, 5, 5, 6, 6, 6, 5, 5, 5}}, 
 {184, {1, 6, 6, 5, 5, 5, 5, 5, 4, 5, 5, 5, 5, 5, 5, 6, 6, 6}}, 
 {184, {4, 4, 5, 6, 5, 5, 5, 5, 6, 6, 5, 5, 6, 6, 6, 6, 5}}, 
 {184, {4, 4, 5, 5, 6, 6, 5, 6, 6, 5, 5, 5, 6, 6, 6, 5, 5}}, 
 {184, {3, 5, 6, 5, 5, 5, 5, 5, 6, 6, 5, 5, 6, 6, 6, 6, 5}}, 
 {183, {4, 4, 5, 5, 4, 5, 5, 5, 5, 5, 6, 6, 6, 5, 5, 5, 5, 5}}, 
 {183, {3, 5, 5, 5, 5, 5, 6, 6, 5, 5, 6, 6, 6, 6, 6, 5, 5}}, 
 {182, {3, 5, 6, 5, 5, 6, 6, 5, 5, 5, 5, 5, 6, 6, 6, 6, 5}}, 
 {182, {3, 5, 5, 6, 6, 6, 6, 5, 5, 5, 5, 6, 6, 5, 5, 6, 5}}, 
 {181, {4, 4, 5, 5, 5, 5, 5, 5, 5, 6, 5, 5, 6, 5, 5, 5, 5, 5}}, 
 {181, {4, 4, 5, 5, 6, 6, 6, 5, 5, 6, 6, 5, 5, 5, 6, 6, 5}}, 
 {180, {4, 5, 5, 5, 5, 5, 5, 4, 5, 5, 5, 5, 6, 6, 5, 5, 5, 5}}, 
 {180, {3, 5, 5, 5, 5, 5, 6, 5, 5, 5, 5, 5, 5, 5, 5, 5, 6, 5}}, 
 {180, {1, 6, 6, 5, 5, 5, 5, 5, 5, 4, 5, 5, 5, 5, 5, 6, 6, 6}}, 
 {180, {4, 4, 5, 5, 6, 6, 5, 5, 6, 6, 5, 5, 5, 6, 6, 6, 5}}, 
 {179, {4, 5, 5, 5, 5, 5, 5, 5, 5, 5, 5, 6, 5, 4, 5, 5, 6, 5}}, 
 {179, {4, 5, 5, 6, 6, 5, 5, 6, 5, 5, 6, 6, 5, 5, 5, 6, 5}}, 
 {179, {4, 5, 5, 6, 6, 5, 4, 5, 5, 6, 6, 5, 6, 6, 6, 5, 5}}, 
 {179, {4, 5, 5, 5, 4, 5, 5, 5, 6, 6, 6, 5, 6, 6, 6, 6, 5}}, 
 {178, {4, 4, 5, 5, 4, 5, 5, 5, 5, 6, 6, 6, 5, 5, 5, 5, 5, 5}}, 
 {178, {4, 5, 5, 6, 6, 5, 6, 6, 5, 4, 5, 5, 6, 6, 6, 5, 5}}, 
 {178, {1, 6, 6, 6, 5, 5, 5, 6, 6, 5, 5, 6, 5, 5, 6, 6, 6}}, 
 {177, {4, 5, 5, 5, 5, 5, 5, 5, 5, 5, 6, 5, 4, 5, 6, 5, 5, 5}}, 
 {177, {4, 5, 5, 5, 4, 5, 5, 5, 5, 4, 5, 5, 6, 6, 6, 5, 5, 5}}, 
 {177, {4, 5, 5, 6, 5, 5, 5, 6, 6, 6, 6, 5, 5, 5, 6, 5, 5}}, 
 {177, {4, 4, 5, 5, 6, 6, 5, 5, 5, 6, 6, 6, 5, 5, 6, 6, 5}}, 
 {177, {4, 4, 5, 5, 5, 6, 6, 6, 5, 5, 5, 6, 6, 5, 6, 6, 5}}, 
 {177, {3, 5, 5, 5, 5, 5, 5, 6, 6, 5, 5, 6, 6, 6, 6, 6, 5}}, 
 {176, {4, 4, 5, 6, 5, 5, 5, 6, 6, 5, 5, 5, 6, 6, 6, 6, 5}}, 
 {176, {3, 5, 6, 5, 5, 6, 6, 6, 6, 5, 5, 5, 5, 5, 6, 6, 5}}, 
 {176, {1, 6, 6, 6, 5, 6, 6, 6, 5, 5, 4, 5, 5, 6, 6, 6, 6}}, 
 {175, {4, 4, 5, 6, 5, 5, 5, 5, 5, 5, 5, 5, 5, 5, 5, 5, 6, 5}}, 
 {175, {4, 4, 5, 5, 5, 5, 5, 5, 4, 5, 5, 5, 5, 6, 6, 6, 5, 5}}, 
 {174, {4, 5, 5, 5, 5, 4, 5, 5, 6, 6, 5, 6, 6, 6, 6, 6, 5}}, 
 {174, {4, 4, 5, 5, 6, 6, 6, 5, 6, 6, 6, 5, 5, 5, 5, 6, 5}}, 
 {174, {1, 6, 6, 5, 6, 6, 5, 5, 6, 6, 5, 5, 5, 5, 6, 6, 6}}, 
 {173, {4, 4, 5, 5, 5, 6, 6, 5, 5, 6, 6, 5, 6, 6, 6, 5, 5}}, 
 {173, {3, 5, 6, 5, 5, 5, 6, 6, 6, 6, 5, 5, 5, 5, 6, 6, 5}}, 
 {173, {1, 6, 6, 5, 6, 6, 6, 5, 5, 5, 6, 5, 5, 5, 6, 6, 6}}, 
 {173, {1, 6, 6, 5, 6, 6, 5, 5, 5, 5, 6, 6, 5, 5, 6, 6, 6}}, 
 {173, {1, 6, 6, 5, 5, 6, 6, 5, 5, 6, 6, 5, 5, 5, 6, 6, 6}}, 
 {172, {4, 4, 5, 5, 5, 5, 5, 5, 5, 5, 6, 5, 5, 6, 5, 5, 5, 5}}, 
 {172, {4, 5, 5, 6, 6, 5, 5, 6, 6, 5, 5, 5, 6, 5, 5, 6, 5}}, 
 {172, {4, 5, 5, 5, 5, 5, 6, 6, 6, 6, 5, 4, 5, 6, 6, 6, 5}}, 
 {172, {4, 5, 5, 5, 4, 5, 5, 5, 6, 6, 6, 6, 6, 5, 6, 6, 5}}, 
 {172, {4, 5, 5, 4, 5, 6, 6, 6, 5, 5, 5, 5, 6, 6, 6, 6, 5}}, 
 {171, {4, 5, 5, 5, 4, 5, 5, 5, 4, 5, 5, 5, 5, 6, 6, 6, 5, 5}}, 
 {171, {4, 5, 5, 5, 6, 6, 6, 6, 6, 6, 5, 4, 5, 5, 6, 5, 5}}, 
 {171, {3, 5, 6, 5, 5, 5, 6, 6, 5, 5, 5, 5, 6, 6, 6, 6, 5}}, 
 {170, {4, 4, 5, 5, 4, 5, 5, 5, 5, 5, 5, 5, 6, 6, 6, 5, 5, 5}}, 
 {170, {3, 5, 6, 5, 5, 5, 5, 5, 6, 6, 6, 5, 5, 6, 6, 6, 5}}, 
 {170, {3, 5, 5, 5, 5, 6, 6, 6, 6, 6, 5, 5, 6, 6, 5, 5, 5}}, 
 {169, {1, 6, 6, 5, 4, 5, 5, 5, 5, 5, 5, 5, 5, 5, 5, 6, 6, 6}}, 
 {169, {3, 5, 5, 5, 5, 5, 5, 5, 6, 6, 6, 6, 5, 6, 6, 6, 5}}, 
 {168, {4, 5, 5, 5, 5, 5, 5, 5, 5, 6, 5, 5, 4, 5, 6, 5, 5, 5}}, 
 {168, {4, 4, 5, 5, 5, 5, 5, 5, 5, 5, 5, 5, 6, 5, 5, 6, 5, 5}}, 
 {168, {4, 4, 5, 5, 5, 5, 5, 5, 5, 5, 5, 4, 5, 5, 6, 6, 6, 5}}, 
 {168, {4, 5, 5, 6, 6, 5, 5, 5, 6, 6, 5, 5, 6, 5, 5, 6, 5}}, 
 {168, {4, 5, 5, 4, 5, 5, 6, 5, 5, 6, 6, 6, 6, 6, 6, 5, 5}}, 
 {168, {4, 4, 5, 5, 6, 6, 5, 5, 6, 6, 6, 5, 5, 5, 6, 6, 5}}, 
 {168, {4, 4, 5, 5, 6, 5, 5, 6, 6, 5, 5, 6, 6, 6, 6, 5, 5}}, 
 {168, {4, 4, 5, 5, 4, 5, 5, 5, 6, 6, 6, 6, 6, 6, 6, 6, 5}}, 
 {168, {3, 5, 5, 6, 6, 5, 5, 5, 6, 6, 6, 6, 5, 5, 5, 6, 5}}, 
 {168, {3, 5, 5, 5, 5, 5, 5, 6, 6, 6, 5, 6, 6, 6, 6, 5, 5}}, 
 {168, {3, 5, 5, 5, 5, 5, 5, 6, 6, 5, 6, 6, 6, 6, 6, 5, 5}}, 
 {168, {1, 6, 6, 5, 5, 5, 6, 6, 5, 6, 6, 5, 5, 5, 6, 6, 6}}, 
 {167, {3, 5, 5, 6, 6, 6, 6, 5, 5, 5, 6, 6, 5, 5, 5, 6, 5}}, 
 {167, {1, 6, 6, 5, 4, 5, 5, 5, 5, 6, 6, 6, 6, 6, 6, 6, 6}}, 
 {167, {3, 5, 6, 6, 6, 5, 6, 6, 6, 6, 6, 6, 6, 6, 6, 5}}, 
 {166, {4, 5, 5, 5, 4, 5, 5, 5, 5, 5, 4, 5, 5, 6, 6, 6, 5, 5}}, 
 {166, {4, 4, 5, 5, 5, 5, 5, 4, 5, 5, 5, 6, 6, 6, 5, 5, 5, 5}}, 
 {166, {4, 5, 5, 6, 6, 6, 5, 4, 5, 5, 6, 6, 6, 5, 5, 6, 5}}, 
 {166, {4, 4, 5, 5, 6, 6, 6, 6, 6, 5, 5, 5, 6, 5, 5, 6, 5}}, 
 {165, {4, 5, 5, 5, 5, 5, 5, 5, 5, 5, 6, 5, 4, 5, 5, 5, 6, 5}}, 
 {165, {4, 5, 5, 5, 4, 5, 5, 5, 5, 5, 5, 6, 5, 5, 5, 5, 6, 5}}, 
 {164, {4, 5, 5, 5, 5, 5, 5, 5, 6, 5, 5, 5, 4, 5, 6, 5, 5, 5}}, 
 {164, {4, 4, 5, 5, 5, 5, 4, 5, 5, 5, 5, 5, 5, 5, 6, 6, 6, 5}}, 
 {164, {4, 5, 5, 6, 6, 6, 5, 5, 6, 5, 5, 6, 5, 5, 5, 6, 5}}, 
 {164, {4, 5, 5, 6, 6, 5, 6, 6, 5, 5, 5, 6, 5, 5, 5, 6, 5}}, 
 {164, {4, 5, 5, 6, 5, 5, 4, 5, 5, 6, 6, 6, 6, 6, 6, 5, 5}}, 
 {164, {3, 5, 5, 5, 6, 5, 5, 5, 5, 5, 6, 6, 6, 6, 6, 6, 5}}, 
 {163, {4, 4, 5, 5, 5, 5, 5, 5, 5, 5, 5, 6, 5, 5, 5, 6, 5, 5}}, 
 {163, {1, 6, 6, 5, 5, 5, 4, 5, 5, 5, 5, 5, 5, 5, 5, 6, 6, 6}}, 
 {163, {4, 4, 5, 5, 6, 6, 6, 6, 5, 6, 6, 5, 5, 5, 5, 6, 5}}, 
 {163, {3, 5, 5, 5, 5, 5, 6, 5, 5, 5, 6, 6, 6, 6, 6, 6, 5}}, 
 {163, {1, 6, 6, 6, 5, 5, 5, 6, 5, 5, 6, 6, 5, 5, 6, 6, 6}}, 
 {163, {1, 6, 6, 6, 5, 5, 4, 5, 5, 6, 6, 6, 5, 6, 6, 6, 6}}, 
 {162, {4, 5, 5, 6, 6, 5, 5, 6, 5, 4, 5, 5, 6, 6, 6, 6, 5}}, 
 {162, {4, 5, 5, 4, 5, 5, 5, 6, 6, 6, 6, 6, 6, 5, 5, 6, 5}}, 
 {162, {4, 4, 5, 5, 6, 6, 5, 5, 5, 6, 6, 5, 6, 6, 6, 5, 5}}, 
 {161, {4, 5, 5, 5, 5, 5, 5, 5, 5, 5, 5, 6, 5, 4, 5, 6, 5, 5}}, 
 {161, {1, 6, 6, 5, 5, 5, 5, 5, 5, 5, 4, 5, 5, 5, 5, 6, 6, 6}}, 
 {160, {1, 6, 6, 6, 5, 5, 5, 5, 6, 5, 5, 6, 6, 5, 6, 6, 6}}, 
 {159, {3, 5, 6, 5, 5, 5, 5, 5, 5, 5, 5, 5, 5, 5, 5, 5, 6, 5}}, 
 {159, {4, 4, 5, 5, 6, 6, 5, 6, 6, 6, 6, 5, 5, 5, 5, 6, 5}}, 
 {159, {3, 5, 6, 6, 5, 5, 6, 6, 5, 5, 5, 5, 5, 6, 6, 6, 5}}, 
 {159, {1, 6, 6, 5, 6, 6, 5, 5, 5, 6, 6, 5, 5, 5, 6, 6, 6}}, 
 {158, {4, 4, 5, 5, 5, 5, 5, 5, 5, 6, 5, 5, 5, 6, 5, 5, 5, 5}}, 
 {158, {4, 5, 5, 6, 5, 5, 6, 5, 5, 6, 6, 5, 5, 6, 6, 5, 5}}, 
 {158, {4, 4, 5, 5, 6, 6, 6, 5, 5, 6, 5, 5, 5, 6, 6, 6, 5}}, 
 {158, {4, 4, 5, 5, 5, 6, 6, 5, 6, 6, 5, 5, 6, 6, 6, 5, 5}}, 
 {158, {3, 5, 6, 6, 5, 5, 5, 5, 6, 6, 6, 5, 5, 5, 6, 6, 5}}, 
 {157, {4, 4, 5, 5, 6, 6, 6, 5, 6, 6, 5, 5, 5, 5, 6, 6, 5}}, 
 {157, {4, 4, 5, 5, 5, 6, 6, 6, 5, 6, 6, 5, 5, 6, 6, 5, 5}}, 
 {156, {4, 5, 5, 4, 5, 5, 6, 5, 5, 5, 6, 6, 6, 6, 6, 6, 5}}, 
 {156, {4, 4, 5, 6, 5, 5, 5, 5, 5, 6, 6, 6, 6, 5, 6, 6, 5}}, 
 {156, {4, 4, 5, 5, 5, 6, 6, 6, 5, 6, 6, 6, 5, 5, 5, 6, 5}}, 
 {156, {3, 5, 5, 6, 6, 5, 5, 5, 5, 6, 6, 6, 6, 5, 5, 6, 5}}, 
 {156, {3, 5, 5, 5, 6, 6, 5, 5, 5, 6, 6, 6, 6, 6, 5, 5, 5}}, 
 {155, {4, 5, 5, 6, 6, 6, 5, 4, 5, 6, 5, 5, 6, 6, 6, 5, 5}}, 
 {155, {4, 5, 5, 5, 4, 5, 5, 5, 6, 6, 5, 6, 6, 6, 6, 6, 5}}, 
 {155, {4, 4, 5, 5, 5, 6, 6, 6, 6, 5, 5, 6, 6, 5, 5, 6, 5}}, 
 {154, {4, 5, 6, 5, 5, 5, 5, 6, 6, 5, 5, 6, 6, 5, 5, 6, 5}}, 
 {154, {4, 4, 5, 5, 6, 6, 5, 6, 6, 5, 5, 5, 5, 6, 6, 6, 5}}, 
 {154, {3, 5, 6, 5, 5, 5, 5, 6, 5, 5, 5, 6, 6, 6, 6, 6, 5}}, 
 {154, {3, 5, 5, 5, 5, 5, 5, 5, 6, 6, 6, 5, 6, 6, 6, 6, 5}}, 
 {153, {4, 5, 5, 5, 5, 5, 5, 6, 5, 5, 5, 5, 4, 5, 5, 5, 6, 5}}, 
 {153, {4, 5, 5, 5, 4, 5, 5, 5, 5, 5, 6, 5, 5, 5, 5, 5, 6, 5}}, 
 {153, {3, 5, 5, 5, 5, 6, 5, 5, 5, 5, 5, 5, 5, 5, 5, 5, 6, 5}}, 
 {153, {3, 5, 5, 5, 5, 5, 5, 5, 5, 6, 5, 5, 5, 5, 5, 5, 6, 5}}, 
 {153, {4, 5, 5, 6, 5, 5, 6, 6, 5, 5, 6, 5, 5, 6, 6, 5, 5}}, 
 {153, {3, 5, 6, 6, 5, 5, 5, 5, 6, 6, 5, 5, 5, 6, 6, 6, 5}}, 
 {152, {4, 4, 5, 5, 5, 4, 5, 6, 6, 6, 5, 5, 5, 5, 5, 5, 5, 5}}, 
 {152, {4, 5, 6, 5, 5, 5, 6, 6, 5, 5, 5, 6, 6, 5, 5, 6, 5}}, 
 {152, {4, 5, 5, 6, 5, 5, 6, 6, 5, 5, 6, 6, 5, 5, 5, 6, 5}}, 
 {152, {4, 5, 5, 4, 5, 5, 5, 6, 6, 5, 5, 6, 6, 6, 6, 6, 5}}, 
 {152, {4, 4, 5, 5, 6, 6, 5, 6, 6, 6, 5, 5, 5, 5, 6, 6, 5}}, 
 {152, {4, 4, 5, 5, 6, 6, 5, 5, 5, 5, 6, 6, 6, 5, 6, 6, 5}}, 
 {152, {3, 5, 5, 5, 6, 6, 6, 5, 5, 5, 6, 6, 6, 6, 5, 5, 5}}, 
 {151, {4, 5, 5, 5, 5, 5, 6, 6, 5, 4, 5, 6, 6, 6, 6, 6, 5}}, 
 {151, {4, 5, 5, 5, 5, 4, 5, 6, 5, 5, 6, 6, 6, 6, 6, 6, 5}}, 
 {151, {4, 4, 5, 5, 5, 6, 6, 6, 5, 5, 6, 6, 5, 5, 6, 6, 5}}, 
 {150, {4, 5, 5, 5, 5, 5, 5, 5, 5, 5, 6, 5, 5, 4, 5, 6, 5, 5}}, 
 {150, {3, 5, 5, 5, 5, 5, 5, 6, 5, 5, 5, 5, 5, 5, 5, 5, 6, 5}}, 
 {150, {4, 5, 5, 6, 6, 5, 6, 6, 5, 5, 6, 5, 5, 5, 5, 6, 5}}, 
 {150, {4, 5, 5, 5, 5, 5, 6, 6, 6, 6, 6, 6, 5, 4, 5, 6, 5}}, 
 {150, {4, 4, 5, 5, 6, 6, 5, 5, 6, 5, 5, 6, 6, 6, 6, 5, 5}}, 
 {150, {4, 4, 5, 5, 4, 5, 6, 6, 6, 6, 6, 6, 6, 6, 5, 5, 5}}, 
 {150, {1, 6, 6, 5, 6, 6, 5, 5, 6, 5, 5, 5, 5, 6, 6, 6, 6}}, 
 {149, {4, 4, 5, 5, 5, 5, 4, 5, 6, 6, 6, 5, 5, 5, 5, 5, 5, 5}}, 
 {149, {4, 4, 5, 5, 5, 6, 6, 6, 5, 5, 6, 6, 6, 5, 5, 6, 5}}, 
 {148, {4, 4, 5, 5, 5, 5, 5, 5, 5, 5, 5, 4, 5, 6, 6, 6, 5, 5}}, 
 {148, {4, 5, 5, 4, 5, 6, 6, 5, 5, 5, 5, 6, 6, 6, 6, 6, 5}}, 
 {148, {4, 5, 5, 4, 5, 6, 5, 5, 5, 5, 6, 6, 6, 6, 6, 6, 5}}, 
 {148, {4, 4, 5, 6, 6, 5, 6, 6, 5, 5, 5, 5, 5, 6, 6, 6, 5}}, 
 {148, {4, 4, 5, 6, 6, 5, 5, 6, 5, 5, 5, 5, 6, 6, 6, 6, 5}}, 
 {148, {3, 5, 5, 5, 5, 6, 5, 5, 5, 5, 6, 6, 6, 6, 6, 6, 5}}, 
 {148, {1, 6, 6, 5, 6, 6, 5, 6, 6, 5, 5, 5, 5, 5, 6, 6, 6}}, 
 {147, {4, 5, 5, 5, 4, 5, 5, 5, 5, 6, 5, 5, 5, 5, 5, 5, 6, 5}}, 
 {147, {4, 5, 5, 5, 5, 5, 6, 5, 4, 5, 6, 6, 6, 6, 6, 6, 5}}, 
 {147, {4, 5, 5, 4, 5, 5, 5, 6, 6, 6, 6, 5, 5, 6, 6, 6, 5}}, 
 {147, {4, 5, 5, 4, 5, 5, 5, 6, 6, 6, 5, 5, 6, 6, 6, 6, 5}}, 
 {147, {3, 5, 5, 6, 6, 6, 5, 6, 6, 6, 5, 5, 5, 5, 5, 6, 5}}, 
 {147, {1, 6, 6, 5, 4, 5, 5, 6, 6, 6, 5, 5, 6, 6, 6, 6, 6}}, 
 {146, {4, 5, 5, 5, 5, 4, 5, 5, 5, 6, 5, 5, 5, 5, 5, 5, 6, 5}}, 
 {146, {4, 4, 5, 5, 5, 4, 5, 5, 5, 5, 5, 5, 5, 5, 6, 6, 6, 5}}, 
 {146, {3, 5, 6, 5, 5, 5, 5, 5, 6, 6, 6, 6, 5, 5, 6, 6, 5}}, 
 {146, {3, 5, 5, 6, 6, 6, 6, 5, 5, 6, 6, 5, 5, 5, 5, 6, 5}}, 
 {146, {3, 5, 5, 5, 5, 5, 5, 6, 6, 6, 6, 6, 5, 6, 6, 5, 5}}, 
 {146, {1, 6, 6, 6, 5, 5, 4, 5, 5, 6, 6, 5, 6, 6, 6, 6, 6}}, 
 {145, {1, 6, 6, 5, 5, 5, 5, 5, 5, 5, 5, 4, 5, 5, 5, 6, 6, 6}}, 
 {145, {4, 5, 5, 6, 6, 5, 6, 6, 5, 5, 5, 5, 6, 5, 5, 6, 5}}, 
 {145, {4, 5, 5, 5, 5, 4, 5, 6, 6, 5, 5, 6, 6, 6, 6, 6, 5}}, 
 {145, {4, 4, 5, 5, 5, 6, 6, 5, 6, 6, 6, 6, 5, 5, 5, 6, 5}}, 
 {145, {4, 4, 5, 5, 5, 6, 6, 5, 6, 6, 6, 5, 5, 6, 6, 5, 5}}, 
 {145, {4, 4, 5, 5, 5, 6, 6, 5, 5, 5, 6, 6, 6, 5, 6, 6, 5}}, 
 {145, {3, 5, 5, 5, 5, 6, 6, 5, 5, 6, 6, 6, 6, 6, 5, 5, 5}}, 
 {144, {4, 5, 5, 5, 5, 5, 5, 6, 5, 4, 5, 5, 6, 5, 5, 5, 5, 5}}, 
 {144, {4, 5, 5, 5, 5, 5, 5, 6, 5, 4, 5, 5, 5, 5, 6, 5, 5, 5}}, 
 {144, {4, 5, 5, 5, 4, 5, 5, 5, 5, 6, 5, 5, 5, 5, 5, 6, 5, 5}}, 
 {144, {3, 5, 5, 5, 5, 5, 5, 5, 5, 5, 5, 5, 6, 5, 5, 5, 6, 5}}, 
 {144, {4, 5, 5, 6, 6, 5, 5, 6, 5, 4, 5, 6, 6, 6, 6, 5, 5}}, 
 {144, {4, 5, 5, 6, 6, 5, 5, 4, 5, 5, 6, 6, 6, 5, 6, 6, 5}}, 
 {144, {4, 5, 5, 5, 6, 5, 5, 6, 6, 5, 5, 6, 6, 5, 5, 6, 5}}, 
 {144, {4, 5, 5, 5, 5, 4, 5, 5, 6, 6, 6, 6, 5, 6, 6, 6, 5}}, 
 {144, {4, 4, 5, 6, 6, 5, 5, 5, 6, 6, 5, 5, 5, 6, 6, 6, 5}}, 
 {144, {4, 4, 5, 6, 5, 5, 5, 5, 5, 6, 6, 6, 5, 6, 6, 6, 5}}, 
 {144, {3, 5, 6, 5, 5, 5, 6, 5, 5, 5, 5, 6, 6, 6, 6, 6, 5}}, 
 {144, {3, 5, 5, 5, 5, 6, 5, 5, 5, 6, 6, 6, 6, 6, 6, 5, 5}}, 
 {143, {4, 5, 5, 5, 5, 5, 4, 5, 5, 5, 6, 5, 5, 5, 5, 5, 6, 5}}, 
 {143, {4, 5, 5, 5, 4, 5, 5, 6, 5, 5, 5, 5, 5, 5, 5, 5, 6, 5}}, 
 {143, {4, 5, 5, 5, 4, 5, 5, 5, 6, 5, 5, 5, 5, 5, 5, 5, 6, 5}}, 
 {143, {4, 5, 5, 6, 5, 4, 5, 5, 6, 6, 6, 6, 5, 6, 6, 5, 5}}, 
 {143, {4, 5, 5, 4, 5, 5, 5, 6, 6, 6, 6, 6, 5, 5, 6, 6, 5}}, 
 {143, {3, 5, 5, 6, 6, 6, 5, 5, 5, 5, 6, 6, 5, 5, 6, 6, 5}}, 
 {143, {3, 5, 5, 5, 5, 5, 5, 5, 6, 6, 5, 6, 6, 6, 6, 6, 5}}, 
 {143, {1, 6, 6, 6, 6, 5, 4, 5, 5, 6, 6, 5, 5, 6, 6, 6, 6}}, 
 {143, {1, 6, 6, 5, 5, 5, 6, 6, 6, 5, 5, 5, 6, 6, 5, 6, 6}}, 
 {143, {1, 6, 6, 5, 5, 5, 6, 6, 5, 5, 5, 6, 6, 6, 5, 6, 6}}, 
 {142, {4, 5, 5, 6, 6, 5, 6, 6, 5, 5, 4, 5, 5, 6, 6, 6, 5}}, 
 {142, {4, 5, 5, 6, 6, 5, 4, 5, 5, 6, 6, 6, 5, 6, 6, 5, 5}}, 
 {142, {4, 4, 5, 6, 6, 5, 5, 5, 5, 6, 6, 5, 5, 6, 6, 6, 5}}, 
 {141, {4, 5, 5, 5, 5, 5, 5, 5, 5, 6, 5, 4, 5, 5, 5, 6, 5, 5}}, 
 {141, {4, 5, 6, 5, 5, 5, 5, 5, 6, 6, 5, 6, 6, 5, 5, 6, 5}}, 
 {141, {4, 5, 5, 6, 6, 6, 5, 5, 5, 6, 5, 5, 6, 5, 5, 6, 5}}, 
 {141, {4, 5, 5, 6, 6, 6, 5, 5, 4, 5, 6, 5, 5, 6, 6, 6, 5}}, 
 {141, {4, 5, 5, 6, 6, 5, 4, 5, 5, 6, 6, 6, 6, 5, 5, 6, 5}}, 
 {141, {4, 5, 5, 5, 5, 5, 6, 6, 6, 6, 6, 5, 4, 5, 6, 6, 5}}, 
 {141, {4, 5, 5, 5, 5, 5, 5, 4, 5, 6, 6, 6, 6, 6, 6, 6, 5}}, 
 {141, {4, 5, 5, 5, 5, 4, 5, 6, 6, 6, 5, 5, 6, 6, 6, 6, 5}}, 
 {141, {3, 5, 6, 6, 5, 5, 5, 5, 5, 6, 6, 6, 5, 5, 6, 6, 5}}, 
 {141, {3, 5, 5, 6, 6, 6, 6, 5, 6, 6, 5, 5, 5, 5, 5, 6, 5}}, 
 {141, {3, 5, 5, 6, 6, 5, 5, 6, 6, 6, 6, 5, 5, 5, 5, 6, 5}}, 
 {141, {1, 6, 6, 6, 6, 5, 5, 4, 5, 5, 6, 6, 5, 6, 6, 6, 6}}, 
 {141, {1, 6, 6, 6, 5, 5, 6, 5, 5, 5, 6, 6, 5, 5, 6, 6, 6}}, 
 {141, {1, 6, 6, 6, 5, 5, 5, 6, 5, 5, 6, 5, 5, 6, 6, 6, 6}}, 
 {141, {1, 6, 6, 6, 5, 5, 4, 5, 5, 6, 6, 6, 6, 5, 6, 6, 6}}, 
 {140, {4, 4, 5, 5, 5, 5, 5, 4, 5, 5, 5, 5, 5, 5, 6, 6, 6, 5}}, 
 {140, {1, 6, 6, 5, 5, 5, 5, 4, 5, 5, 5, 5, 5, 5, 5, 6, 6, 6}}, 
 {140, {4, 5, 5, 6, 5, 5, 5, 6, 6, 5, 5, 6, 6, 5, 5, 6, 5}}, 
 {140, {4, 4, 5, 5, 5, 6, 6, 5, 5, 6, 6, 6, 6, 5, 5, 6, 5}}, 
 {140, {4, 4, 5, 5, 5, 5, 4, 5, 6, 6, 6, 6, 6, 6, 6, 6, 5}}, 
 {140, {3, 5, 5, 5, 4, 5, 5, 6, 6, 6, 6, 6, 6, 6, 6, 5, 5}}, 
 {140, {1, 6, 6, 5, 6, 6, 6, 6, 5, 5, 4, 5, 5, 6, 6, 6, 6}}, 
 {139, {4, 5, 6, 5, 5, 5, 6, 6, 6, 5, 5, 5, 6, 5, 5, 6, 5}}, 
 {139, {1, 6, 6, 5, 5, 6, 6, 5, 5, 5, 6, 6, 5, 5, 6, 6, 6}}, 
 {138, {4, 5, 5, 5, 5, 5, 5, 5, 6, 5, 4, 5, 5, 5, 6, 5, 5, 5}}, 
 {138, {4, 5, 5, 6, 5, 4, 5, 5, 6, 6, 5, 6, 6, 6, 6, 5, 5}}, 
 {138, {4, 4, 5, 5, 6, 6, 6, 6, 5, 5, 5, 6, 5, 5, 6, 6, 5}}, 
 {138, {1, 6, 6, 5, 5, 5, 6, 6, 5, 5, 6, 6, 5, 5, 6, 6, 6}}, 
 {138, {1, 6, 6, 5, 5, 5, 5, 6, 6, 6, 5, 5, 6, 6, 5, 6, 6}}, 
 {138, {4, 5, 6, 6, 6, 6, 5, 6, 6, 6, 5, 6, 6, 6, 6, 5}}, 
 {137, {4, 5, 5, 5, 5, 5, 5, 6, 5, 5, 4, 5, 5, 5, 5, 5, 6, 5}}, 
 {137, {4, 5, 5, 5, 5, 5, 4, 5, 5, 6, 5, 5, 5, 5, 5, 5, 6, 5}}, 
 {137, {4, 5, 5, 6, 5, 5, 5, 5, 6, 6, 5, 6, 6, 5, 5, 6, 5}}, 
 {137, {4, 5, 5, 5, 6, 6, 5, 5, 6, 6, 5, 5, 6, 5, 5, 6, 5}}, 
 {137, {4, 4, 5, 6, 6, 6, 5, 5, 5, 5, 6, 5, 5, 6, 6, 6, 5}}, 
 {137, {4, 4, 5, 5, 5, 6, 6, 5, 5, 6, 6, 5, 5, 6, 6, 6, 5}}, 
 {137, {1, 6, 6, 5, 6, 6, 6, 5, 5, 4, 5, 5, 6, 6, 6, 6, 6}}, 
 {137, {1, 6, 6, 5, 6, 6, 5, 5, 5, 6, 5, 5, 5, 6, 6, 6, 6}}, 
 {136, {4, 5, 5, 5, 5, 5, 6, 5, 5, 5, 4, 5, 5, 5, 6, 5, 5, 5}}, 
 {136, {4, 5, 5, 5, 5, 5, 5, 6, 5, 5, 5, 4, 5, 6, 5, 5, 5, 5}}, 
 {136, {4, 5, 5, 5, 5, 5, 5, 6, 5, 5, 5, 4, 5, 5, 6, 5, 5, 5}}, 
 {136, {4, 4, 5, 5, 5, 5, 5, 5, 5, 4, 5, 6, 6, 6, 5, 5, 5, 5}}, 
 {136, {4, 5, 5, 6, 6, 6, 5, 5, 6, 5, 5, 5, 6, 5, 5, 6, 5}}, 
 {136, {4, 5, 5, 6, 6, 5, 5, 6, 6, 5, 4, 5, 5, 6, 6, 6, 5}}, 
 {136, {4, 5, 5, 6, 5, 4, 5, 5, 6, 6, 5, 5, 6, 6, 6, 6, 5}}, 
 {136, {3, 5, 5, 5, 5, 6, 6, 6, 5, 5, 6, 6, 6, 6, 5, 5, 5}}, 
 {136, {1, 6, 6, 6, 5, 6, 6, 5, 5, 4, 5, 5, 6, 6, 6, 6, 6}}, 
 {136, {1, 6, 6, 5, 5, 5, 5, 6, 6, 5, 5, 6, 6, 6, 5, 6, 6}}, 
 {135, {4, 5, 5, 5, 5, 5, 5, 6, 5, 4, 5, 5, 5, 6, 5, 5, 5, 5}}, 
 {135, {4, 5, 5, 5, 5, 5, 5, 5, 6, 5, 5, 5, 4, 5, 5, 5, 6, 5}}, 
 {135, {4, 5, 5, 5, 5, 5, 5, 5, 5, 6, 5, 5, 5, 4, 5, 5, 6, 5}}, 
 {135, {4, 5, 5, 5, 6, 6, 5, 5, 6, 5, 5, 6, 6, 5, 5, 6, 5}}, 
 {135, {4, 5, 5, 5, 4, 5, 5, 6, 5, 5, 6, 6, 6, 6, 6, 6, 5}}, 
 {135, {4, 4, 5, 6, 5, 5, 6, 5, 5, 5, 5, 6, 6, 6, 6, 6, 5}}, 
 {135, {4, 4, 5, 6, 5, 5, 5, 6, 5, 5, 5, 6, 6, 6, 6, 6, 5}}, 
 {135, {4, 4, 5, 5, 5, 6, 6, 6, 5, 5, 6, 6, 5, 6, 6, 5, 5}}, 
 {135, {4, 4, 5, 5, 5, 6, 6, 5, 5, 6, 6, 6, 5, 6, 6, 5, 5}}, 
 {135, {4, 4, 5, 5, 5, 6, 5, 5, 6, 6, 6, 5, 6, 6, 6, 5, 5}}, 
 {135, {1, 6, 6, 6, 6, 5, 5, 6, 5, 4, 5, 5, 6, 6, 6, 6, 6}}, 
 {134, {4, 5, 5, 5, 5, 5, 5, 5, 6, 5, 5, 4, 5, 6, 5, 5, 5, 5}}, 
 {134, {4, 5, 5, 5, 5, 4, 5, 5, 5, 5, 5, 6, 5, 5, 5, 5, 6, 5}}, 
 {134, {4, 5, 5, 5, 4, 5, 5, 5, 5, 4, 5, 5, 5, 6, 6, 6, 5, 5}}, 
 {134, {4, 4, 5, 5, 5, 5, 5, 5, 5, 4, 5, 5, 5, 5, 6, 6, 6, 5}}, 
 {134, {3, 5, 5, 5, 5, 5, 5, 5, 6, 5, 5, 5, 5, 5, 5, 5, 6, 5}}, 
 {134, {4, 5, 5, 6, 6, 5, 5, 6, 5, 5, 6, 5, 5, 5, 6, 6, 5}}, 
 {134, {4, 4, 5, 6, 6, 5, 5, 5, 5, 5, 6, 6, 6, 5, 6, 6, 5}}, 
 {134, {4, 4, 5, 5, 5, 6, 6, 6, 6, 6, 5, 5, 6, 5, 5, 6, 5}}, 
 {134, {4, 4, 5, 5, 5, 6, 6, 5, 6, 6, 6, 5, 5, 5, 6, 6, 5}}, 
 {134, {1, 6, 6, 6, 5, 5, 6, 6, 6, 5, 5, 4, 5, 6, 6, 6, 6}}, 
 {134, {4, 5, 6, 6, 6, 5, 6, 6, 6, 6, 6, 5, 6, 6, 6, 5}}, 
 {133, {4, 5, 5, 5, 5, 5, 5, 6, 5, 5, 5, 4, 5, 5, 5, 5, 6, 5}}, 
 {133, {4, 5, 5, 5, 5, 5, 5, 5, 6, 5, 4, 5, 5, 5, 5, 5, 6, 5}}, 
 {133, {4, 5, 5, 4, 5, 5, 5, 4, 5, 5, 5, 5, 5, 6, 6, 6, 5, 5}}, 
 {133, {4, 4, 5, 5, 5, 5, 5, 5, 5, 5, 4, 5, 6, 6, 6, 5, 5, 5}}, 
 {133, {4, 5, 5, 6, 6, 6, 5, 4, 5, 6, 6, 6, 5, 5, 5, 6, 5}}, 
 {133, {4, 5, 5, 6, 6, 5, 5, 4, 5, 5, 6, 6, 5, 6, 6, 6, 5}}, 
 {133, {4, 5, 5, 6, 5, 5, 5, 6, 6, 5, 6, 6, 5, 5, 5, 6, 5}}, 
 {133, {4, 4, 5, 5, 5, 5, 6, 6, 6, 5, 5, 6, 6, 5, 6, 6, 5}}, 
 {133, {3, 5, 5, 5, 5, 5, 6, 5, 5, 6, 6, 6, 6, 6, 6, 5, 5}}, 
 {133, {1, 6, 6, 5, 6, 6, 6, 6, 6, 5, 5, 4, 5, 5, 6, 6, 6}}, 
 {133, {1, 6, 6, 5, 5, 6, 6, 5, 6, 6, 5, 5, 5, 5, 6, 6, 6}}, 
 {132, {4, 5, 5, 5, 5, 5, 5, 6, 5, 5, 5, 5, 5, 4, 5, 5, 6, 5}}, 
 {132, {4, 5, 5, 5, 5, 5, 5, 5, 5, 5, 6, 5, 4, 5, 5, 6, 5, 5}}, 
 {132, {4, 5, 5, 5, 5, 5, 5, 4, 5, 5, 5, 6, 5, 5, 5, 5, 6, 5}}, 
 {132, {4, 5, 5, 5, 4, 5, 5, 5, 5, 5, 6, 5, 5, 5, 5, 6, 5, 5}}, 
 {132, {4, 4, 5, 5, 4, 5, 6, 6, 6, 5, 5, 5, 5, 5, 5, 5, 5, 5}}, 
 {132, {4, 5, 6, 5, 5, 5, 5, 6, 6, 6, 5, 5, 6, 5, 5, 6, 5}}, 
 {132, {4, 5, 5, 5, 5, 6, 5, 5, 4, 5, 6, 6, 6, 6, 6, 6, 5}}, 
 {132, {4, 4, 5, 6, 6, 5, 5, 5, 6, 5, 5, 5, 6, 6, 6, 6, 5}}, 
 {132, {4, 4, 5, 5, 6, 6, 6, 5, 5, 5, 6, 5, 5, 6, 6, 6, 5}}, 
 {132, {4, 4, 5, 5, 6, 6, 5, 5, 6, 5, 5, 5, 6, 6, 6, 6, 5}}, 
 {132, {4, 4, 5, 5, 6, 5, 5, 5, 6, 6, 6, 5, 6, 6, 6, 5, 5}}, 
 {132, {1, 6, 6, 5, 5, 5, 5, 4, 5, 6, 6, 6, 6, 6, 6, 6, 6}}, 
 {132, {1, 6, 6, 5, 4, 5, 6, 6, 6, 5, 5, 5, 6, 6, 6, 6, 6}}, 
 {132, {1, 6, 6, 1, 6, 6, 6, 6, 6, 5, 5, 6, 6, 6, 6, 6, 6}}, 
 {131, {4, 5, 5, 5, 5, 5, 5, 5, 5, 6, 5, 4, 5, 5, 6, 5, 5, 5}}, 
 {131, {4, 5, 5, 5, 4, 5, 5, 5, 5, 5, 6, 5, 5, 5, 6, 5, 5, 5}}, 
 {131, {4, 5, 5, 4, 5, 5, 5, 5, 5, 5, 4, 5, 5, 6, 6, 6, 5, 5}}, 
 {131, {4, 5, 5, 6, 6, 5, 5, 5, 6, 5, 5, 6, 5, 5, 6, 6, 5}}, 
 {131, {4, 4, 5, 5, 6, 6, 6, 6, 6, 6, 5, 4, 5, 5, 6, 6, 5}}, 
 {131, {3, 5, 5, 5, 5, 6, 6, 6, 6, 5, 5, 6, 6, 6, 5, 5, 5}}, 
 {131, {3, 5, 5, 5, 5, 5, 6, 6, 6, 6, 5, 6, 6, 6, 5, 5, 5}}, 
 {131, {1, 6, 6, 6, 5, 5, 6, 6, 6, 5, 4, 5, 5, 6, 6, 6, 6}}, 
 {131, {1, 6, 6, 6, 5, 5, 5, 6, 5, 5, 5, 6, 6, 5, 6, 6, 6}}, 
 {131, {1, 6, 6, 5, 6, 6, 6, 5, 5, 6, 5, 5, 5, 5, 6, 6, 6}}, 
 {131, {1, 6, 6, 5, 6, 6, 6, 5, 5, 5, 5, 6, 5, 5, 6, 6, 6}}, 
 {131, {1, 6, 6, 5, 6, 6, 5, 5, 5, 5, 5, 6, 6, 5, 6, 6, 6}}, 
 {131, {1, 6, 6, 5, 5, 6, 6, 5, 5, 5, 5, 6, 6, 5, 6, 6, 6}}, 
 {131, {1, 6, 6, 5, 5, 5, 5, 5, 6, 6, 6, 5, 6, 6, 5, 6, 6}}, 
 {130, {4, 5, 5, 5, 5, 5, 5, 5, 5, 6, 5, 4, 5, 6, 5, 5, 5, 5}}, 
 {130, {4, 4, 5, 5, 4, 5, 5, 5, 5, 5, 5, 5, 5, 5, 6, 6, 6, 5}}, 
 {130, {4, 5, 5, 6, 6, 5, 5, 4, 5, 6, 5, 5, 6, 6, 6, 6, 5}}, 
 {130, {4, 5, 5, 6, 5, 5, 6, 6, 5, 6, 6, 5, 5, 5, 5, 6, 5}}, 
 {130, {4, 4, 5, 5, 5, 6, 5, 5, 6, 6, 5, 6, 6, 6, 6, 5, 5}}, 
 {130, {3, 5, 5, 6, 6, 5, 5, 6, 6, 5, 5, 5, 6, 6, 6, 5, 5}}, 
 {130, {1, 6, 6, 6, 5, 5, 6, 6, 5, 5, 4, 5, 6, 6, 6, 6, 6}}, 
 {130, {1, 6, 6, 6, 5, 5, 4, 5, 6, 6, 6, 6, 5, 5, 6, 6, 6}}, 
 {130, {1, 6, 6, 5, 5, 5, 5, 6, 6, 5, 6, 6, 5, 5, 6, 6, 6}}, 
 {130, {1, 6, 6, 5, 4, 5, 6, 6, 6, 6, 6, 5, 5, 5, 6, 6, 6}}, 
 {129, {4, 5, 5, 5, 5, 5, 5, 5, 6, 5, 4, 5, 5, 6, 5, 5, 5, 5}}, 
 {129, {4, 5, 5, 5, 5, 4, 5, 5, 6, 5, 5, 5, 5, 5, 5, 5, 6, 5}}, 
 {129, {4, 5, 5, 5, 4, 5, 5, 5, 5, 5, 5, 5, 6, 5, 5, 5, 6, 5}}, 
 {129, {4, 4, 5, 5, 6, 6, 5, 5, 5, 5, 6, 6, 5, 6, 6, 6, 5}}, 
 {129, {4, 4, 5, 5, 6, 5, 5, 5, 6, 6, 6, 6, 6, 5, 5, 6, 5}}, 
 {129, {1, 6, 6, 6, 5, 5, 6, 5, 5, 6, 5, 5, 5, 6, 6, 6, 6}}, 
 {129, {1, 6, 6, 5, 6, 6, 5, 5, 5, 5, 5, 6, 6, 6, 5, 6, 6}}, 
 {129, {1, 6, 6, 5, 5, 6, 5, 5, 5, 6, 6, 5, 5, 6, 6, 6, 6}}, 
 {128, {4, 5, 5, 5, 5, 5, 5, 6, 5, 5, 5, 5, 4, 5, 6, 5, 5, 5}}, 
 {128, {4, 5, 5, 5, 5, 5, 5, 6, 5, 5, 4, 5, 5, 5, 6, 5, 5, 5}}, 
 {128, {4, 5, 5, 5, 5, 4, 5, 5, 5, 5, 6, 5, 5, 5, 5, 6, 5, 5}}, 
 {128, {4, 5, 5, 5, 5, 4, 5, 5, 5, 5, 5, 5, 6, 5, 5, 5, 6, 5}}, 
 {128, {4, 5, 6, 5, 5, 5, 5, 6, 6, 5, 6, 6, 5, 5, 5, 6, 5}}, 
 {128, {4, 5, 5, 6, 6, 6, 6, 5, 5, 4, 5, 6, 5, 5, 6, 6, 5}}, 
 {128, {4, 5, 5, 6, 5, 4, 5, 5, 6, 6, 6, 5, 5, 6, 6, 6, 5}}, 
 {128, {4, 5, 5, 5, 6, 5, 5, 6, 5, 5, 6, 6, 5, 6, 6, 5, 5}}, 
 {128, {3, 5, 6, 6, 6, 5, 5, 5, 5, 6, 5, 5, 5, 6, 6, 6, 5}}, 
 {128, {3, 5, 5, 5, 5, 6, 6, 6, 6, 6, 6, 5, 5, 6, 5, 5, 5}}, 
 {128, {1, 6, 6, 6, 6, 5, 5, 4, 5, 6, 6, 5, 5, 6, 6, 6, 6}}, 
 {128, {1, 6, 6, 5, 4, 5, 6, 6, 6, 6, 5, 5, 5, 6, 6, 6, 6}}, 
 {128, {1, 6, 6, 1, 6, 6, 6, 6, 5, 5, 6, 6, 6, 6, 6, 6, 6}}, 
 {127, {4, 5, 5, 5, 5, 5, 5, 6, 5, 5, 5, 5, 5, 4, 5, 6, 5, 5}}, 
 {127, {4, 5, 5, 6, 6, 5, 5, 6, 6, 6, 6, 5, 5, 4, 5, 6, 5}}, 
 {127, {4, 5, 5, 5, 6, 5, 5, 6, 6, 6, 6, 5, 5, 6, 5, 5, 5}}, 
 {127, {4, 4, 5, 5, 5, 6, 6, 6, 5, 6, 6, 5, 5, 5, 6, 6, 5}}, 
 {127, {4, 4, 5, 5, 5, 6, 6, 5, 6, 6, 5, 6, 6, 6, 5, 5, 5}}, 
 {127, {4, 4, 5, 5, 5, 5, 6, 6, 6, 6, 5, 6, 6, 5, 5, 6, 5}}, 
 {127, {4, 4, 5, 5, 5, 5, 6, 6, 5, 5, 6, 6, 6, 5, 6, 6, 5}}, 
 {127, {4, 4, 5, 4, 5, 5, 5, 6, 6, 6, 6, 6, 6, 6, 6, 5, 5}}, 
 {127, {3, 5, 6, 6, 5, 5, 5, 6, 6, 5, 5, 5, 5, 6, 6, 6, 5}}, 
 {127, {3, 5, 6, 5, 5, 6, 5, 5, 5, 5, 5, 6, 6, 6, 6, 6, 5}}, 
 {127, {3, 5, 5, 6, 6, 6, 6, 6, 5, 5, 5, 5, 6, 5, 5, 6, 5}}, 
 {127, {1, 6, 6, 6, 6, 1, 6, 6, 6, 6, 5, 5, 6, 6, 6, 6, 6}}, 
 {127, {1, 6, 6, 6, 5, 4, 5, 6, 6, 6, 6, 5, 5, 5, 6, 6, 6}}, 
 {127, {1, 6, 6, 5, 5, 6, 5, 5, 6, 6, 6, 5, 5, 5, 6, 6, 6}}, 
 {127, {1, 6, 6, 5, 5, 6, 5, 5, 6, 6, 5, 5, 5, 6, 6, 6, 6}}, 
 {126, {4, 5, 5, 5, 5, 4, 5, 5, 5, 5, 6, 5, 5, 5, 5, 5, 6, 5}}, 
 {126, {4, 4, 5, 5, 5, 5, 5, 5, 4, 5, 6, 6, 6, 5, 5, 5, 5, 5}}, 
 {126, {4, 4, 5, 5, 5, 5, 5, 5, 4, 5, 5, 5, 5, 5, 6, 6, 6, 5}}, 
 {126, {4, 5, 6, 5, 5, 5, 5, 5, 6, 5, 5, 6, 6, 5, 6, 6, 5}}, 
 {126, {4, 5, 5, 6, 6, 5, 4, 5, 5, 6, 6, 6, 5, 5, 6, 6, 5}}, 
 {126, {4, 5, 5, 6, 5, 4, 5, 5, 6, 6, 6, 6, 6, 5, 5, 6, 5}}, 
 {126, {4, 5, 5, 4, 5, 5, 5, 5, 6, 6, 6, 6, 5, 6, 6, 6, 5}}, 
 {126, {1, 6, 6, 6, 1, 6, 6, 6, 6, 5, 5, 6, 6, 6, 6, 6, 6}}, 
 {126, {1, 6, 6, 5, 5, 6, 5, 5, 6, 5, 5, 5, 6, 6, 6, 6, 6}}, 
 {126, {1, 6, 6, 5, 5, 5, 5, 6, 6, 5, 5, 6, 6, 5, 6, 6, 6}}, 
 {125, {4, 5, 5, 5, 5, 5, 6, 5, 4, 5, 5, 5, 6, 5, 5, 5, 5, 5}}, 
 {125, {4, 5, 5, 6, 5, 4, 5, 5, 6, 6, 6, 5, 6, 6, 6, 5, 5}}, 
 {125, {4, 5, 5, 5, 6, 5, 5, 5, 6, 6, 5, 6, 6, 5, 5, 6, 5}}, 
 {125, {4, 4, 5, 5, 6, 6, 6, 6, 6, 5, 5, 6, 5, 5, 5, 6, 5}}, 
 {125, {4, 4, 5, 5, 5, 5, 6, 6, 5, 6, 6, 6, 6, 5, 5, 6, 5}}, 
 {125, {3, 5, 6, 5, 5, 5, 5, 5, 5, 6, 6, 6, 5, 6, 6, 6, 5}}, 
 {125, {3, 5, 5, 6, 6, 6, 5, 5, 6, 6, 5, 5, 5, 5, 6, 6, 5}}, 
 {125, {3, 5, 5, 6, 6, 6, 5, 5, 5, 6, 6, 5, 5, 5, 6, 6, 5}}, 
 {125, {3, 5, 5, 5, 5, 5, 5, 5, 6, 6, 6, 6, 6, 5, 6, 6, 5}}, 
 {125, {1, 6, 6, 6, 6, 5, 5, 4, 5, 6, 5, 5, 6, 6, 6, 6, 6}}, 
 {125, {1, 6, 6, 6, 5, 5, 6, 5, 5, 5, 5, 6, 6, 5, 6, 6, 6}}, 
 {124, {4, 5, 5, 5, 4, 5, 5, 5, 6, 5, 5, 5, 5, 5, 5, 6, 5, 5}}, 
 {124, {3, 5, 5, 5, 5, 5, 5, 5, 5, 5, 5, 5, 5, 6, 5, 5, 6, 5}}, 
 {124, {4, 5, 5, 6, 6, 5, 5, 6, 6, 6, 5, 5, 4, 5, 6, 6, 5}}, 
 {124, {4, 4, 5, 6, 6, 5, 5, 5, 5, 5, 6, 6, 5, 6, 6, 6, 5}}, 
 {124, {4, 4, 5, 6, 5, 5, 5, 5, 5, 6, 6, 5, 6, 6, 6, 6, 5}}, 
 {124, {4, 4, 5, 5, 6, 6, 6, 6, 6, 5, 5, 4, 5, 6, 6, 6, 5}}, 
 {124, {4, 4, 5, 5, 6, 6, 5, 5, 5, 6, 5, 5, 6, 6, 6, 6, 5}}, 
 {124, {3, 5, 6, 5, 5, 5, 5, 5, 5, 6, 6, 6, 6, 5, 6, 6, 5}}, 
 {123, {4, 5, 5, 5, 5, 5, 5, 5, 6, 5, 4, 5, 6, 5, 5, 5, 5, 5}}, 
 {123, {4, 5, 5, 5, 5, 5, 5, 5, 6, 5, 4, 5, 5, 5, 5, 6, 5, 5}}, 
 {123, {1, 6, 6, 5, 5, 5, 5, 5, 5, 5, 5, 5, 5, 4, 5, 6, 6, 6}}, 
 {123, {4, 5, 6, 5, 5, 6, 5, 5, 5, 5, 6, 6, 6, 5, 5, 6, 5}}, 
 {123, {4, 5, 6, 5, 5, 5, 6, 5, 5, 6, 6, 6, 5, 5, 5, 6, 5}}, 
 {123, {4, 5, 5, 6, 5, 5, 6, 6, 5, 5, 5, 6, 6, 5, 5, 6, 5}}, 
 {123, {4, 5, 5, 6, 5, 5, 6, 5, 5, 6, 5, 5, 6, 6, 6, 5, 5}}, 
 {123, {3, 5, 6, 6, 5, 5, 5, 5, 5, 6, 6, 5, 5, 6, 6, 6, 5}}, 
 {123, {3, 5, 6, 5, 5, 5, 5, 5, 5, 6, 6, 5, 6, 6, 6, 6, 5}}, 
 {123, {1, 6, 6, 5, 5, 6, 6, 5, 5, 6, 5, 5, 5, 6, 6, 6, 6}}, 
 {122, {4, 5, 5, 5, 5, 5, 5, 5, 5, 6, 5, 4, 5, 5, 5, 5, 6, 5}}, 
 {122, {3, 5, 5, 5, 5, 5, 5, 5, 5, 5, 5, 6, 5, 5, 5, 5, 6, 5}}, 
 {122, {3, 5, 5, 6, 6, 5, 6, 6, 6, 6, 5, 5, 5, 5, 5, 6, 5}}, 
 {122, {3, 5, 5, 6, 6, 5, 5, 6, 6, 6, 5, 5, 5, 5, 6, 6, 5}}, 
 {121, {4, 5, 5, 5, 5, 5, 5, 5, 4, 5, 5, 5, 6, 5, 5, 5, 6, 5}}, 
 {121, {4, 5, 5, 5, 5, 5, 4, 5, 5, 5, 5, 6, 5, 5, 6, 5, 5, 5}}, 
 {121, {4, 5, 5, 5, 5, 4, 5, 5, 5, 6, 5, 5, 5, 5, 5, 6, 5, 5}}, 
 {121, {4, 5, 5, 5, 5, 4, 5, 5, 5, 5, 5, 6, 5, 5, 6, 5, 5, 5}}, 
 {121, {4, 4, 5, 5, 5, 5, 6, 6, 5, 5, 6, 6, 5, 6, 6, 6, 5}}, 
 {121, {3, 5, 5, 6, 6, 5, 5, 6, 6, 5, 5, 5, 5, 6, 6, 6, 5}}, 
 {121, {1, 6, 6, 6, 5, 4, 5, 5, 6, 6, 6, 6, 5, 5, 6, 6, 6}}, 
 {120, {4, 5, 5, 5, 5, 5, 6, 5, 5, 5, 5, 4, 5, 5, 5, 5, 6, 5}}, 
 {120, {4, 5, 5, 5, 5, 5, 5, 5, 5, 6, 5, 5, 5, 4, 5, 6, 5, 5}}, 
 {120, {3, 5, 5, 5, 6, 5, 5, 5, 5, 5, 5, 5, 5, 5, 5, 5, 6, 5}}, 
 {120, {4, 5, 5, 4, 5, 5, 5, 6, 5, 5, 6, 6, 6, 6, 6, 6, 5}}, 
 {120, {4, 4, 5, 5, 6, 6, 5, 5, 5, 6, 6, 5, 5, 6, 6, 6, 5}}, 
 {120, {4, 4, 5, 5, 6, 5, 5, 6, 6, 5, 5, 5, 6, 6, 6, 6, 5}}, 
 {120, {4, 4, 5, 5, 5, 5, 6, 6, 6, 5, 6, 6, 6, 5, 5, 6, 5}}, 
 {120, {3, 5, 5, 5, 5, 5, 6, 6, 5, 6, 6, 6, 6, 6, 5, 5, 5}}, 
 {120, {1, 6, 6, 6, 5, 5, 4, 5, 6, 6, 6, 5, 5, 6, 6, 6, 6}}, 
 {119, {4, 5, 5, 5, 5, 5, 5, 6, 5, 5, 5, 4, 5, 5, 5, 6, 5, 5}}, 
 {119, {4, 5, 5, 5, 5, 5, 4, 5, 5, 5, 5, 6, 5, 5, 5, 5, 6, 5}}, 
 {119, {4, 4, 5, 4, 5, 5, 6, 6, 6, 5, 5, 5, 5, 5, 5, 5, 5, 5}}, 
 {119, {4, 4, 5, 5, 6, 5, 5, 6, 6, 6, 6, 6, 5, 5, 5, 6, 5}}, 
 {119, {4, 4, 5, 5, 5, 5, 6, 6, 6, 5, 6, 6, 5, 6, 6, 5, 5}}, 
 {119, {3, 5, 5, 6, 6, 6, 5, 5, 5, 5, 5, 6, 6, 5, 6, 6, 5}}, 
 {119, {3, 5, 5, 6, 6, 5, 5, 5, 6, 6, 6, 5, 5, 5, 6, 6, 5}}, 
 {119, {1, 6, 6, 5, 5, 6, 5, 5, 5, 5, 6, 6, 6, 5, 6, 6, 6}}, 
 {118, {4, 5, 5, 5, 5, 5, 6, 5, 5, 5, 4, 5, 5, 5, 5, 6, 5, 5}}, 
 {118, {4, 5, 5, 5, 5, 5, 5, 5, 5, 4, 5, 5, 5, 6, 5, 5, 6, 5}}, 
 {118, {4, 5, 5, 5, 4, 5, 5, 5, 5, 6, 5, 5, 5, 5, 6, 5, 5, 5}}, 
 {118, {4, 4, 5, 5, 5, 5, 5, 5, 5, 5, 4, 5, 5, 5, 6, 6, 6, 5}}, 
 {118, {4, 5, 5, 6, 5, 5, 6, 6, 6, 5, 5, 5, 6, 5, 5, 6, 5}}, 
 {118, {4, 4, 5, 5, 5, 6, 6, 5, 5, 6, 5, 5, 6, 6, 6, 6, 5}}, 
 {118, {4, 4, 5, 5, 5, 5, 6, 6, 5, 6, 6, 6, 5, 5, 6, 6, 5}}, 
 {118, {3, 5, 6, 6, 5, 5, 5, 5, 6, 5, 5, 5, 6, 6, 6, 6, 5}}, 
 {118, {3, 5, 6, 6, 5, 5, 5, 5, 5, 5, 6, 6, 6, 5, 6, 6, 5}}, 
 {118, {3, 5, 5, 6, 6, 5, 5, 5, 6, 6, 5, 5, 6, 6, 6, 5, 5}}, 
 {118, {3, 5, 5, 6, 6, 5, 5, 5, 5, 6, 6, 6, 5, 5, 6, 6, 5}}, 
 {118, {3, 5, 5, 5, 5, 5, 6, 6, 6, 6, 6, 5, 6, 6, 5, 5, 5}}, 
 {118, {1, 6, 6, 5, 5, 6, 6, 6, 5, 5, 5, 5, 6, 6, 5, 6, 6}}, 
 {118, {1, 6, 6, 5, 5, 6, 6, 5, 5, 5, 6, 5, 5, 6, 6, 6, 6}}, 
 {118, {1, 6, 6, 5, 5, 6, 5, 5, 5, 6, 6, 6, 5, 5, 6, 6, 6}}, 
 {118, {1, 6, 6, 5, 5, 5, 6, 6, 6, 5, 5, 6, 6, 5, 5, 6, 6}}, 
 {117, {4, 5, 5, 5, 5, 5, 5, 5, 5, 6, 5, 5, 4, 5, 5, 5, 6, 5}}, 
 {117, {4, 5, 5, 5, 5, 5, 5, 4, 5, 5, 6, 5, 5, 5, 5, 5, 6, 5}}, 
 {117, {4, 5, 5, 4, 5, 5, 5, 5, 5, 4, 5, 5, 5, 6, 6, 6, 5, 5}}, 
 {117, {4, 5, 5, 4, 5, 5, 5, 5, 4, 5, 5, 5, 5, 6, 6, 6, 5, 5}}, 
 {117, {4, 5, 5, 6, 6, 5, 5, 6, 5, 5, 6, 5, 5, 6, 6, 5, 5}}, 
 {117, {4, 5, 5, 4, 5, 5, 5, 5, 6, 6, 6, 5, 6, 6, 6, 6, 5}}, 
 {117, {3, 5, 5, 5, 5, 6, 6, 6, 5, 6, 6, 6, 6, 5, 5, 5, 5}}, 
 {116, {4, 5, 5, 5, 5, 5, 6, 5, 5, 4, 5, 5, 5, 6, 5, 5, 5, 5}}, 
 {116, {4, 5, 5, 6, 6, 5, 4, 5, 5, 6, 6, 5, 5, 6, 6, 6, 5}}, 
 {116, {4, 4, 5, 5, 6, 6, 6, 6, 5, 5, 6, 5, 5, 5, 6, 6, 5}}, 
 {116, {1, 6, 6, 6, 5, 5, 6, 6, 5, 4, 5, 5, 6, 6, 6, 6, 6}}, 
 {116, {1, 6, 6, 5, 5, 6, 6, 6, 5, 5, 6, 5, 5, 5, 6, 6, 6}}, 
 {116, {1, 6, 6, 5, 5, 6, 6, 5, 5, 5, 6, 6, 6, 5, 5, 6, 6}}, 
 {116, {1, 6, 6, 5, 5, 5, 5, 6, 6, 5, 6, 6, 6, 5, 5, 6, 6}}, 
 {115, {4, 5, 5, 5, 5, 5, 5, 5, 6, 5, 5, 4, 5, 5, 5, 5, 6, 5}}, 
 {115, {4, 5, 5, 5, 5, 5, 5, 5, 4, 5, 5, 6, 5, 5, 5, 5, 6, 5}}, 
 {115, {1, 6, 6, 1, 6, 6, 5, 5, 5, 5, 5, 5, 5, 5, 6, 6, 6, 6}}, 
 {115, {4, 5, 6, 5, 5, 5, 5, 6, 5, 5, 6, 6, 5, 5, 6, 6, 5}}, 
 {115, {4, 5, 5, 5, 6, 6, 6, 5, 5, 6, 5, 5, 6, 5, 5, 6, 5}}, 
 {115, {4, 4, 5, 6, 5, 5, 5, 6, 6, 6, 6, 6, 5, 5, 5, 6, 5}}, 
 {115, {4, 4, 5, 6, 5, 5, 5, 5, 6, 5, 5, 6, 6, 6, 6, 6, 5}}, 
 {115, {3, 5, 6, 5, 5, 5, 5, 5, 6, 5, 5, 6, 6, 6, 6, 6, 5}}, 
 {115, {1, 6, 6, 6, 5, 5, 4, 5, 6, 5, 5, 6, 6, 6, 6, 6, 6}}, 
 {115, {1, 6, 6, 5, 4, 5, 5, 6, 6, 6, 6, 5, 5, 6, 6, 6, 6}}, 
 {114, {4, 5, 5, 5, 5, 5, 5, 5, 5, 5, 6, 5, 5, 4, 5, 5, 6, 5}}, 
 {114, {4, 5, 6, 5, 5, 6, 5, 5, 5, 5, 6, 6, 5, 5, 6, 6, 5}}, 
 {114, {4, 5, 6, 5, 5, 5, 6, 5, 5, 5, 6, 6, 6, 5, 5, 6, 5}}, 
 {114, {4, 5, 5, 6, 5, 4, 5, 6, 6, 6, 6, 5, 5, 6, 6, 5, 5}}, 
 {114, {4, 5, 5, 5, 6, 5, 5, 6, 6, 5, 6, 6, 5, 5, 6, 5, 5}}, 
 {114, {4, 4, 5, 5, 6, 5, 5, 5, 6, 6, 5, 6, 6, 6, 6, 5, 5}}, 
 {114, {4, 4, 5, 5, 5, 6, 6, 5, 6, 6, 5, 5, 5, 6, 6, 6, 5}}, 
 {114, {3, 5, 5, 6, 6, 5, 5, 5, 6, 6, 5, 5, 5, 6, 6, 6, 5}}, 
 {113, {4, 5, 5, 5, 4, 5, 5, 5, 5, 5, 5, 6, 5, 5, 5, 6, 5, 5}}, 
 {113, {4, 4, 5, 5, 5, 6, 5, 5, 5, 5, 5, 5, 5, 5, 6, 5, 5, 5}}, 
 {113, {4, 5, 5, 6, 6, 5, 5, 6, 5, 5, 5, 6, 5, 5, 6, 6, 5}}, 
 {113, {4, 5, 5, 6, 6, 5, 5, 4, 5, 6, 6, 5, 5, 6, 6, 6, 5}}, 
 {113, {4, 5, 4, 5, 5, 6, 6, 6, 5, 5, 5, 6, 6, 6, 6, 5, 5}}, 
 {113, {4, 4, 5, 5, 6, 6, 6, 6, 6, 5, 4, 5, 5, 6, 6, 6, 5}}, 
 {113, {4, 4, 5, 5, 6, 5, 5, 5, 6, 6, 6, 6, 5, 6, 6, 5, 5}}, 
 {113, {3, 5, 6, 6, 5, 5, 5, 6, 5, 5, 5, 5, 6, 6, 6, 6, 5}}, 
 {113, {3, 5, 5, 5, 6, 6, 6, 5, 5, 5, 6, 6, 6, 5, 5, 6, 5}}, 
 {113, {1, 6, 6, 5, 5, 6, 6, 6, 6, 6, 6, 6, 6, 1, 6, 6, 6}}, 
 {113, {1, 6, 6, 5, 5, 5, 5, 5, 6, 6, 5, 6, 6, 6, 5, 6, 6}}, 
 {113, {1, 6, 6, 5, 4, 5, 5, 6, 6, 5, 5, 6, 6, 6, 6, 6, 6}}, 
 {112, {4, 5, 5, 5, 5, 5, 5, 5, 6, 5, 5, 5, 4, 5, 5, 6, 5, 5}}, 
 {112, {4, 4, 5, 5, 5, 5, 5, 4, 5, 6, 6, 6, 5, 5, 5, 5, 5, 5}}, 
 {112, {1, 6, 6, 5, 5, 5, 5, 5, 5, 5, 5, 6, 6, 6, 1, 6, 6, 6}}, 
 {112, {4, 5, 5, 6, 5, 5, 5, 6, 5, 5, 6, 6, 5, 6, 6, 5, 5}}, 
 {112, {4, 5, 5, 5, 6, 6, 6, 6, 6, 6, 5, 5, 5, 4, 5, 6, 5}}, 
 {112, {4, 4, 5, 6, 6, 5, 5, 4, 5, 5, 6, 6, 6, 6, 6, 6, 5}}, 
 {112, {4, 4, 5, 5, 6, 6, 6, 6, 5, 4, 5, 5, 6, 6, 6, 6, 5}}, 
 {112, {4, 4, 5, 5, 5, 6, 6, 6, 5, 6, 6, 6, 5, 5, 6, 5, 5}}, 
 {112, {4, 4, 5, 5, 5, 6, 6, 5, 5, 6, 6, 6, 5, 5, 6, 6, 5}}, 
 {112, {3, 5, 6, 6, 6, 5, 5, 5, 5, 5, 6, 5, 5, 6, 6, 6, 5}}, 
 {112, {1, 6, 6, 6, 6, 5, 5, 5, 6, 5, 4, 5, 6, 6, 6, 6, 6}}, 
 {112, {1, 6, 6, 6, 5, 4, 5, 6, 6, 6, 5, 5, 5, 6, 6, 6, 6}}, 
 {112, {1, 6, 6, 6, 5, 4, 5, 5, 6, 6, 6, 5, 5, 6, 6, 6, 6}}, 
 {112, {1, 6, 6, 6, 5, 4, 5, 5, 6, 6, 5, 5, 6, 6, 6, 6, 6}}, 
 {112, {1, 6, 6, 5, 5, 6, 6, 5, 5, 5, 5, 6, 6, 6, 5, 6, 6}}, 
 {111, {4, 5, 5, 5, 5, 5, 5, 5, 6, 5, 5, 5, 5, 4, 5, 5, 6, 5}}, 
 {111, {4, 4, 5, 5, 5, 5, 5, 5, 6, 6, 5, 5, 4, 5, 6, 5, 5, 5}}, 
 {111, {4, 5, 5, 6, 6, 6, 5, 5, 5, 6, 5, 4, 5, 6, 6, 6, 5}}, 
 {111, {4, 5, 5, 5, 6, 5, 5, 6, 5, 5, 6, 6, 6, 5, 5, 6, 5}}, 
 {111, {4, 4, 5, 6, 5, 5, 4, 5, 5, 6, 6, 6, 6, 6, 6, 6, 5}}, 
 {111, {3, 5, 5, 6, 6, 6, 5, 6, 6, 5, 5, 5, 5, 5, 6, 6, 5}}, 
 {111, {3, 5, 5, 6, 6, 5, 5, 5, 6, 6, 6, 5, 5, 6, 6, 5, 5}}, 
 {111, {3, 5, 5, 5, 6, 5, 5, 5, 6, 6, 6, 6, 6, 6, 5, 5, 5}}, 
 {111, {1, 6, 6, 6, 1, 6, 6, 6, 5, 5, 6, 6, 6, 6, 6, 6, 6}}, 
 {111, {1, 6, 6, 5, 5, 5, 5, 6, 5, 5, 6, 6, 5, 6, 6, 6, 6}}, 
 {111, {4, 5, 6, 6, 5, 6, 6, 6, 6, 6, 6, 6, 5, 6, 6, 5}}, 
 {110, {4, 5, 5, 5, 5, 5, 5, 6, 5, 5, 4, 5, 6, 5, 5, 5, 5, 5}}, 
 {110, {4, 5, 5, 5, 5, 5, 5, 6, 5, 4, 5, 5, 5, 5, 5, 6, 5, 5}}, 
 {110, {4, 5, 5, 5, 5, 5, 5, 5, 6, 5, 5, 4, 5, 5, 5, 6, 5, 5}}, 
 {110, {4, 5, 5, 4, 5, 5, 5, 5, 5, 6, 5, 5, 5, 5, 5, 5, 6, 5}}, 
 {110, {4, 5, 5, 4, 5, 5, 5, 5, 5, 5, 6, 5, 5, 5, 6, 5, 5, 5}}, 
 {110, {4, 5, 5, 4, 5, 5, 5, 5, 4, 5, 5, 6, 6, 6, 5, 5, 5, 5}}, 
 {110, {4, 4, 5, 5, 5, 5, 5, 5, 5, 5, 5, 5, 4, 5, 6, 6, 6, 5}}, 
 {110, {1, 6, 6, 6, 5, 5, 5, 5, 4, 4, 5, 5, 5, 5, 5, 6, 6, 6}}, 
 {110, {4, 5, 6, 5, 5, 6, 5, 5, 6, 6, 5, 5, 5, 5, 6, 6, 5}}, 
 {110, {4, 5, 5, 6, 6, 5, 5, 6, 6, 5, 4, 5, 6, 6, 6, 5, 5}}, 
 {110, {4, 5, 5, 6, 5, 5, 5, 6, 6, 6, 5, 5, 6, 5, 5, 6, 5}}, 
 {110, {4, 5, 5, 4, 5, 5, 5, 5, 6, 6, 6, 6, 6, 5, 6, 6, 5}}, 
 {110, {4, 4, 5, 5, 6, 5, 5, 6, 6, 6, 5, 5, 5, 6, 6, 6, 5}}, 
 {110, {3, 5, 5, 6, 6, 5, 5, 5, 5, 6, 6, 5, 5, 6, 6, 6, 5}}, 
 {110, {3, 5, 5, 6, 5, 5, 5, 5, 5, 6, 6, 6, 6, 6, 6, 5, 5}}, 
 {110, {3, 5, 5, 6, 5, 5, 5, 5, 5, 5, 6, 6, 6, 6, 6, 6, 5}}, 
 {110, {1, 6, 6, 6, 5, 5, 4, 5, 6, 6, 5, 5, 6, 6, 6, 6, 6}}, 
 {110, {1, 6, 6, 5, 6, 6, 5, 5, 5, 5, 6, 5, 5, 6, 6, 6, 6}}, 
 {110, {1, 6, 6, 5, 5, 5, 6, 6, 6, 5, 5, 6, 5, 5, 6, 6, 6}}, 
 {110, {3, 5, 6, 6, 6, 6, 6, 6, 5, 6, 6, 6, 6, 6, 6, 5}}, 
 {109, {4, 5, 5, 5, 5, 5, 6, 5, 5, 5, 5, 5, 4, 5, 5, 5, 6, 5}}, 
 {109, {4, 5, 5, 5, 5, 5, 6, 5, 4, 5, 5, 5, 5, 5, 6, 5, 5, 5}}, 
 {109, {4, 5, 6, 5, 5, 6, 6, 5, 5, 5, 5, 6, 5, 5, 6, 6, 5}}, 
 {109, {4, 5, 6, 5, 5, 5, 5, 6, 5, 5, 5, 6, 6, 5, 6, 6, 5}}, 
 {109, {4, 5, 5, 5, 6, 6, 6, 6, 5, 5, 5, 4, 5, 6, 6, 6, 5}}, 
 {109, {4, 4, 5, 5, 6, 5, 5, 6, 6, 6, 6, 5, 5, 5, 6, 6, 5}}, 
 {109, {4, 4, 5, 5, 5, 6, 6, 6, 6, 5, 6, 6, 5, 5, 6, 5, 5}}, 
 {109, {4, 4, 5, 5, 5, 5, 6, 6, 6, 5, 6, 6, 5, 5, 6, 6, 5}}, 
 {109, {3, 5, 5, 6, 6, 6, 6, 6, 5, 5, 4, 5, 5, 6, 6, 6, 5}}, 
 {109, {3, 5, 5, 5, 6, 6, 6, 5, 6, 6, 6, 5, 5, 5, 5, 6, 5}}, 
 {109, {1, 6, 6, 1, 6, 6, 6, 5, 5, 6, 6, 6, 6, 6, 6, 6, 6}}, 
 {108, {4, 5, 5, 5, 5, 5, 6, 5, 5, 5, 5, 4, 5, 6, 5, 5, 5, 5}}, 
 {108, {4, 5, 6, 5, 5, 5, 6, 5, 5, 5, 5, 6, 6, 5, 6, 6, 5}}, 
 {108, {4, 5, 5, 6, 6, 5, 4, 5, 6, 6, 6, 5, 5, 6, 6, 5, 5}}, 
 {107, {4, 5, 5, 5, 5, 5, 5, 5, 6, 5, 5, 4, 5, 5, 6, 5, 5, 5}}, 
 {107, {4, 5, 5, 5, 5, 5, 5, 5, 4, 5, 5, 5, 6, 5, 5, 6, 5, 5}}, 
 {107, {4, 5, 5, 4, 5, 5, 5, 5, 5, 6, 5, 5, 5, 5, 5, 6, 5, 5}}, 
 {107, {4, 5, 5, 6, 5, 5, 6, 5, 5, 5, 6, 6, 5, 5, 6, 6, 5}}, 
 {107, {4, 4, 5, 5, 6, 6, 6, 5, 5, 4, 5, 6, 6, 6, 6, 6, 5}}, 
 {107, {4, 4, 5, 5, 5, 6, 5, 5, 5, 6, 6, 6, 6, 5, 6, 6, 5}}, 
 {107, {4, 4, 5, 5, 5, 6, 5, 5, 5, 6, 6, 6, 5, 6, 6, 6, 5}}, 
 {107, {4, 4, 5, 5, 5, 5, 6, 6, 5, 6, 6, 5, 5, 6, 6, 6, 5}}, 
 {107, {3, 5, 5, 6, 6, 6, 6, 6, 5, 5, 5, 6, 5, 5, 5, 6, 5}}, 
 {107, {3, 5, 5, 6, 6, 6, 5, 5, 6, 5, 5, 5, 5, 6, 6, 6, 5}}, 
 {107, {3, 5, 5, 5, 5, 5, 6, 6, 6, 5, 6, 6, 6, 6, 5, 5, 5}}, 
 {107, {1, 6, 6, 5, 5, 5, 6, 6, 6, 6, 6, 5, 4, 5, 6, 6, 6}}, 
 {106, {4, 5, 5, 5, 5, 5, 6, 5, 5, 5, 5, 4, 5, 5, 5, 6, 5, 5}}, 
 {106, {4, 5, 5, 5, 5, 5, 5, 5, 5, 5, 6, 5, 5, 5, 4, 5, 6, 5}}, 
 {106, {4, 5, 5, 5, 5, 4, 5, 5, 5, 5, 5, 5, 6, 5, 5, 6, 5, 5}}, 
 {106, {4, 5, 5, 5, 4, 5, 5, 5, 5, 5, 5, 6, 5, 5, 6, 5, 5, 5}}, 
 {106, {4, 5, 5, 4, 5, 5, 5, 5, 5, 6, 5, 5, 5, 5, 6, 5, 5, 5}}, 
 {106, {4, 5, 5, 4, 5, 5, 5, 5, 5, 4, 5, 5, 6, 6, 6, 5, 5, 5}}, 
 {106, {4, 5, 5, 4, 5, 5, 5, 4, 5, 5, 6, 6, 6, 5, 5, 5, 5, 5}}, 
 {106, {4, 4, 5, 5, 5, 5, 5, 5, 6, 5, 5, 6, 5, 5, 5, 5, 5, 5}}, 
 {106, {4, 5, 5, 6, 6, 6, 5, 4, 5, 6, 5, 5, 5, 6, 6, 6, 5}}, 
 {106, {4, 5, 5, 6, 5, 5, 6, 5, 5, 6, 6, 6, 5, 5, 5, 6, 5}}, 
 {106, {4, 5, 5, 6, 5, 5, 6, 5, 5, 6, 6, 5, 5, 5, 6, 6, 5}}, 
 {106, {4, 5, 5, 6, 5, 5, 5, 6, 5, 5, 6, 6, 5, 5, 6, 6, 5}}, 
 {106, {4, 5, 5, 6, 5, 4, 5, 5, 6, 6, 6, 6, 5, 5, 6, 6, 5}}, 
 {106, {4, 5, 5, 5, 6, 6, 6, 5, 5, 5, 4, 5, 6, 6, 6, 6, 5}}, 
 {106, {4, 5, 5, 5, 6, 6, 5, 5, 5, 4, 5, 6, 6, 6, 6, 6, 5}}, 
 {106, {4, 4, 5, 5, 5, 6, 6, 5, 5, 5, 6, 6, 5, 6, 6, 6, 5}}, 
 {106, {4, 4, 5, 5, 5, 5, 6, 6, 5, 6, 6, 5, 6, 6, 6, 5, 5}}, 
 {106, {3, 5, 5, 6, 6, 6, 6, 6, 6, 6, 5, 5, 4, 5, 5, 6, 5}}, 
 {106, {3, 5, 5, 5, 5, 4, 5, 5, 6, 6, 6, 6, 6, 6, 6, 6, 5}}, 
 {106, {1, 6, 6, 5, 5, 4, 5, 5, 6, 6, 6, 6, 5, 6, 6, 6, 6}}, 
 {105, {4, 5, 5, 5, 5, 5, 6, 5, 5, 4, 5, 5, 5, 5, 6, 5, 5, 5}}, 
 {105, {4, 5, 5, 5, 5, 5, 5, 4, 5, 5, 5, 5, 6, 5, 5, 5, 6, 5}}, 
 {105, {4, 5, 5, 5, 5, 5, 4, 5, 5, 5, 5, 5, 5, 6, 5, 5, 6, 5}}, 
 {105, {4, 5, 5, 4, 5, 5, 5, 6, 5, 5, 5, 5, 5, 5, 5, 6, 5, 5}}, 
 {105, {4, 5, 5, 6, 6, 5, 4, 5, 6, 6, 5, 5, 6, 6, 6, 5, 5}}, 
 {105, {4, 5, 5, 6, 6, 5, 4, 5, 6, 5, 5, 6, 6, 6, 6, 5, 5}}, 
 {105, {4, 5, 5, 5, 6, 5, 5, 6, 6, 5, 6, 6, 5, 5, 5, 6, 5}}, 
 {105, {1, 6, 6, 5, 5, 4, 5, 5, 6, 6, 6, 5, 6, 6, 6, 6, 6}}, 
 {104, {4, 5, 5, 5, 5, 5, 6, 5, 5, 5, 4, 5, 5, 6, 5, 5, 5, 5}}, 
 {104, {4, 5, 5, 5, 5, 5, 4, 5, 5, 5, 6, 5, 5, 5, 5, 6, 5, 5}}, 
 {104, {4, 5, 5, 5, 4, 5, 5, 5, 5, 5, 5, 5, 5, 6, 5, 5, 6, 5}}, 
 {104, {4, 5, 5, 6, 6, 6, 5, 5, 5, 6, 6, 5, 4, 5, 6, 6, 5}}, 
 {104, {4, 5, 5, 6, 6, 5, 4, 5, 6, 6, 6, 6, 5, 5, 5, 6, 5}}, 
 {104, {4, 4, 5, 5, 6, 5, 5, 6, 6, 6, 6, 5, 5, 6, 6, 5, 5}}, 
 {104, {4, 4, 5, 5, 5, 6, 6, 6, 6, 5, 6, 6, 5, 5, 5, 6, 5}}, 
 {104, {4, 4, 5, 4, 5, 5, 6, 6, 6, 6, 6, 6, 6, 6, 5, 5, 5}}, 
 {104, {1, 6, 6, 6, 6, 5, 4, 5, 6, 6, 5, 5, 5, 6, 6, 6, 6}}, 
 {104, {1, 6, 6, 5, 5, 5, 6, 6, 5, 5, 5, 6, 6, 5, 6, 6, 6}}, 
 {104, {1, 6, 6, 5, 5, 5, 5, 6, 5, 5, 6, 6, 6, 5, 6, 6, 6}}, 
 {103, {4, 5, 5, 5, 5, 5, 6, 5, 5, 5, 5, 4, 5, 5, 6, 5, 5, 5}}, 
 {103, {3, 5, 5, 5, 4, 5, 5, 5, 5, 5, 5, 5, 5, 6, 6, 6, 5, 5}}, 
 {103, {4, 5, 5, 6, 6, 5, 5, 5, 6, 6, 6, 6, 5, 4, 5, 6, 5}}, 
 {103, {4, 5, 5, 6, 6, 5, 5, 5, 6, 6, 6, 5, 4, 5, 6, 6, 5}}, 
 {103, {4, 5, 5, 5, 6, 6, 6, 5, 6, 6, 6, 5, 5, 4, 5, 6, 5}}, 
 {103, {4, 5, 4, 5, 5, 5, 5, 6, 6, 6, 6, 6, 6, 6, 5, 5, 5}}, 
 {103, {4, 4, 5, 6, 6, 5, 5, 5, 5, 6, 5, 5, 6, 6, 6, 6, 5}}, 
 {103, {4, 4, 5, 5, 6, 5, 5, 5, 6, 6, 6, 6, 5, 5, 6, 6, 5}}, 
 {103, {4, 4, 5, 5, 5, 5, 6, 6, 5, 6, 6, 6, 5, 6, 6, 5, 5}}, 
 {103, {3, 5, 5, 6, 6, 6, 6, 5, 5, 4, 5, 5, 6, 6, 6, 6, 5}}, 
 {103, {3, 5, 5, 6, 6, 6, 5, 5, 5, 6, 5, 5, 6, 6, 6, 5, 5}}, 
 {103, {3, 5, 5, 6, 6, 6, 5, 5, 4, 5, 5, 6, 6, 6, 6, 6, 5}}, 
 {103, {1, 6, 6, 6, 6, 5, 4, 5, 6, 5, 5, 5, 6, 6, 6, 6, 6}}, 
 {103, {1, 6, 6, 6, 5, 5, 6, 6, 6, 6, 1, 6, 6, 6, 6, 6, 6}}, 
 {103, {1, 6, 6, 5, 4, 5, 5, 6, 6, 6, 6, 6, 5, 5, 6, 6, 6}}, 
 {102, {4, 5, 5, 5, 4, 5, 5, 5, 5, 5, 5, 5, 6, 5, 5, 6, 5, 5}}, 
 {102, {4, 5, 5, 4, 5, 5, 6, 5, 5, 5, 5, 5, 5, 5, 5, 5, 6, 5}}, 
 {102, {4, 5, 5, 4, 5, 5, 5, 6, 5, 5, 5, 5, 5, 5, 5, 5, 6, 5}}, 
 {102, {4, 5, 5, 4, 5, 5, 5, 5, 5, 5, 6, 5, 5, 5, 5, 5, 6, 5}}, 
 {102, {4, 4, 5, 5, 5, 5, 5, 4, 5, 5, 6, 6, 5, 5, 5, 5, 6, 5}}, 
 {102, {3, 5, 5, 5, 5, 5, 5, 5, 5, 5, 4, 5, 5, 6, 6, 6, 5, 5}}, 
 {102, {1, 6, 6, 6, 5, 5, 4, 4, 5, 5, 5, 5, 5, 5, 5, 6, 6, 6}}, 
 {102, {4, 5, 5, 6, 5, 4, 5, 6, 6, 6, 5, 5, 6, 6, 6, 5, 5}}, 
 {102, {4, 5, 5, 5, 6, 6, 5, 6, 6, 5, 5, 5, 6, 5, 5, 6, 5}}, 
 {102, {4, 5, 5, 5, 6, 5, 5, 6, 5, 5, 6, 6, 5, 5, 6, 6, 5}}, 
 {102, {4, 5, 5, 4, 5, 5, 5, 5, 6, 6, 5, 6, 6, 6, 6, 6, 5}}, 
 {102, {4, 5, 4, 5, 5, 5, 5, 5, 6, 6, 6, 6, 6, 6, 6, 5, 5}}, 
 {102, {4, 4, 5, 6, 6, 6, 5, 5, 4, 5, 5, 6, 6, 6, 6, 6, 5}}, 
 {102, {4, 4, 5, 5, 6, 5, 5, 6, 5, 5, 6, 6, 6, 6, 6, 5, 5}}, 
 {102, {4, 4, 5, 5, 6, 5, 5, 5, 5, 6, 6, 6, 5, 6, 6, 6, 5}}, 
 {102, {4, 4, 5, 5, 5, 6, 5, 5, 6, 6, 6, 6, 5, 6, 6, 5, 5}}, 
 {102, {4, 4, 5, 5, 5, 6, 5, 5, 6, 6, 5, 5, 6, 6, 6, 6, 5}}, 
 {101, {4, 5, 5, 5, 5, 5, 5, 5, 5, 5, 5, 5, 6, 5, 4, 5, 6, 5}}, 
 {101, {4, 5, 5, 4, 5, 5, 5, 5, 5, 5, 5, 5, 6, 5, 5, 5, 6, 5}}, 
 {101, {4, 4, 5, 5, 5, 5, 6, 5, 5, 5, 5, 5, 5, 6, 5, 5, 5, 5}}, 
 {101, {4, 4, 5, 5, 5, 5, 5, 5, 6, 6, 5, 5, 4, 5, 5, 5, 6, 5}}, 
 {101, {4, 4, 5, 4, 5, 5, 5, 5, 5, 5, 5, 5, 5, 6, 6, 6, 5, 5}}, 
 {101, {3, 5, 5, 5, 4, 5, 5, 5, 5, 6, 6, 6, 5, 5, 5, 5, 5, 5}}, 
 {101, {4, 5, 6, 5, 5, 5, 6, 5, 5, 6, 5, 5, 5, 6, 6, 6, 5}}, 
 {101, {4, 5, 6, 5, 5, 5, 5, 6, 5, 5, 6, 6, 6, 5, 5, 6, 5}}, 
 {101, {4, 5, 5, 6, 6, 6, 5, 5, 5, 6, 6, 6, 5, 4, 5, 6, 5}}, 
 {101, {4, 5, 5, 6, 5, 5, 6, 5, 5, 5, 6, 6, 5, 6, 6, 5, 5}}, 
 {101, {4, 5, 5, 5, 6, 6, 5, 6, 6, 5, 5, 6, 5, 5, 5, 6, 5}}, 
 {101, {4, 5, 5, 5, 6, 6, 5, 6, 6, 5, 4, 5, 6, 6, 6, 5, 5}}, 
 {101, {4, 4, 5, 5, 5, 6, 5, 5, 6, 5, 5, 6, 6, 6, 6, 6, 5}}, 
 {101, {3, 5, 5, 6, 6, 6, 6, 6, 5, 5, 6, 5, 5, 5, 5, 6, 5}}, 
 {101, {3, 5, 5, 6, 6, 6, 6, 5, 5, 5, 6, 5, 5, 5, 6, 6, 5}}, 
 {101, {3, 5, 5, 5, 5, 5, 5, 6, 5, 5, 6, 6, 6, 6, 6, 6, 5}}, 
 {101, {1, 6, 6, 6, 5, 5, 5, 6, 6, 6, 5, 4, 5, 6, 6, 6, 6}}, 
 {101, {1, 6, 6, 5, 5, 6, 6, 6, 5, 5, 5, 6, 5, 5, 6, 6, 6}}, 
 {101, {1, 6, 6, 5, 5, 5, 6, 5, 5, 6, 6, 5, 5, 6, 6, 6, 6}}, 
 {101, {1, 6, 6, 5, 5, 4, 5, 5, 6, 6, 6, 6, 6, 5, 6, 6, 6}}, 
 {100, {4, 5, 5, 5, 6, 5, 4, 5, 6, 6, 6, 5, 6, 6, 6, 5, 5}}, 
 {100, {4, 4, 5, 5, 6, 6, 6, 6, 5, 5, 4, 5, 6, 6, 6, 6, 5}}, 
 {100, {4, 4, 5, 5, 6, 6, 6, 5, 4, 5, 6, 6, 6, 6, 6, 5, 5}}, 
 {100, {3, 5, 6, 6, 5, 6, 6, 5, 5, 5, 5, 5, 5, 6, 6, 6, 5}}, 
 {100, {3, 5, 5, 6, 6, 6, 6, 5, 5, 5, 5, 6, 5, 5, 6, 6, 5}}, 
 {100, {3, 5, 5, 6, 6, 5, 6, 6, 5, 5, 5, 5, 6, 6, 6, 5, 5}}, 
 {100, {3, 5, 5, 6, 6, 5, 5, 6, 5, 5, 5, 6, 6, 6, 6, 5, 5}}, 
 {100, {1, 6, 6, 6, 5, 5, 6, 6, 6, 6, 6, 6, 1, 6, 6, 6, 6}}, 
 {100, {1, 6, 6, 5, 5, 5, 6, 6, 5, 5, 6, 5, 5, 6, 6, 6, 6}}, 
 {99, {4, 5, 5, 5, 5, 4, 5, 5, 5, 5, 5, 6, 5, 5, 5, 6, 5, 5}}, 
 {99, {4, 5, 6, 5, 5, 5, 5, 6, 6, 5, 5, 6, 5, 5, 6, 6, 5}}, 
 {99, {4, 5, 5, 6, 5, 5, 5, 6, 6, 5, 5, 6, 5, 5, 6, 6, 5}}, 
 {99, {4, 5, 5, 5, 5, 5, 4, 5, 6, 6, 5, 6, 6, 6, 6, 6, 5}}, 
 {99, {4, 4, 5, 5, 5, 6, 6, 6, 6, 5, 5, 6, 5, 5, 6, 6, 5}}, 
 {99, {4, 4, 5, 5, 5, 5, 6, 5, 5, 6, 6, 5, 6, 6, 6, 6, 5}}, 
 {99, {3, 5, 6, 6, 5, 5, 6, 5, 5, 5, 5, 5, 6, 6, 6, 6, 5}}, 
 {99, {3, 5, 5, 6, 6, 6, 6, 6, 6, 5, 5, 4, 5, 5, 6, 6, 5}}, 
 {99, {3, 5, 5, 6, 6, 5, 5, 5, 5, 6, 6, 5, 6, 6, 6, 5, 5}}, 
 {99, {3, 5, 5, 5, 6, 6, 6, 6, 5, 5, 5, 6, 6, 5, 5, 6, 5}}, 
 {99, {1, 6, 6, 6, 5, 5, 6, 5, 5, 5, 6, 5, 5, 6, 6, 6, 6}}, 
 {99, {1, 6, 6, 5, 6, 6, 5, 5, 4, 5, 5, 6, 6, 6, 6, 6, 6}}, 
 {98, {4, 5, 5, 5, 5, 5, 5, 5, 5, 5, 5, 5, 5, 4, 5, 6, 6, 5}}, 
 {98, {4, 5, 6, 5, 5, 6, 5, 5, 6, 5, 5, 5, 5, 6, 6, 6, 5}}, 
 {98, {4, 5, 5, 6, 5, 4, 5, 6, 6, 5, 5, 6, 6, 6, 6, 5, 5}}, 
 {98, {4, 5, 5, 5, 6, 5, 4, 5, 6, 6, 6, 6, 5, 6, 6, 5, 5}}, 
 {98, {4, 4, 5, 5, 6, 5, 5, 6, 6, 6, 5, 5, 6, 6, 6, 5, 5}}, 
 {98, {4, 4, 5, 5, 5, 6, 6, 6, 5, 5, 6, 5, 5, 6, 6, 6, 5}}, 
 {98, {3, 5, 5, 6, 6, 5, 5, 5, 5, 5, 6, 6, 6, 5, 6, 6, 5}}, 
 {98, {1, 6, 6, 5, 5, 5, 5, 6, 5, 5, 6, 6, 6, 6, 5, 6, 6}}, 
 {98, {1, 6, 6, 5, 5, 5, 5, 5, 6, 6, 5, 6, 6, 5, 6, 6, 6}}, 
 {98, {1, 6, 6, 5, 5, 4, 5, 6, 6, 5, 5, 6, 6, 6, 6, 6, 6}}, 
 {97, {4, 5, 5, 5, 5, 5, 5, 6, 5, 5, 4, 5, 5, 5, 5, 6, 5, 5}}, 
 {97, {4, 5, 5, 5, 5, 5, 5, 5, 6, 5, 5, 5, 5, 4, 5, 6, 5, 5}}, 
 {97, {4, 5, 5, 5, 5, 5, 5, 5, 5, 4, 5, 5, 6, 5, 5, 5, 6, 5}}, 
 {97, {4, 5, 5, 5, 5, 5, 5, 4, 5, 5, 5, 5, 6, 5, 5, 6, 5, 5}}, 
 {97, {4, 5, 5, 5, 5, 5, 4, 5, 5, 5, 5, 5, 6, 5, 5, 5, 6, 5}}, 
 {97, {4, 5, 5, 5, 5, 4, 5, 5, 5, 5, 5, 5, 5, 6, 5, 5, 6, 5}}, 
 {97, {4, 5, 5, 5, 4, 5, 5, 5, 5, 5, 6, 5, 5, 6, 5, 5, 5, 5}}, 
 {97, {4, 5, 5, 5, 4, 5, 5, 5, 5, 5, 5, 4, 5, 5, 6, 6, 6, 5}}, 
 {97, {4, 5, 5, 5, 4, 5, 5, 5, 5, 4, 5, 5, 5, 5, 6, 6, 6, 5}}, 
 {97, {4, 5, 5, 4, 5, 5, 5, 5, 5, 5, 6, 5, 5, 5, 5, 6, 5, 5}}, 
 {97, {4, 4, 5, 5, 5, 5, 5, 6, 5, 5, 5, 5, 6, 5, 5, 5, 5, 5}}, 
 {97, {4, 4, 5, 4, 5, 5, 5, 5, 5, 6, 6, 6, 5, 5, 5, 5, 5, 5}}, 
 {97, {3, 5, 5, 5, 5, 4, 5, 5, 5, 5, 5, 5, 5, 6, 6, 6, 5, 5}}, 
 {97, {4, 5, 6, 5, 5, 6, 6, 5, 5, 6, 5, 5, 5, 5, 6, 6, 5}}, 
 {97, {4, 5, 5, 6, 6, 6, 6, 6, 5, 4, 5, 6, 5, 5, 5, 6, 5}}, 
 {97, {4, 5, 5, 6, 5, 5, 5, 5, 6, 5, 5, 6, 6, 5, 6, 6, 5}}, 
 {97, {4, 5, 5, 6, 5, 4, 5, 6, 6, 6, 6, 6, 5, 5, 5, 6, 5}}, 
 {97, {1, 6, 6, 6, 5, 5, 6, 5, 4, 5, 5, 6, 6, 6, 6, 6, 6}}, 
 {96, {4, 5, 5, 5, 5, 5, 5, 6, 5, 5, 4, 5, 5, 6, 5, 5, 5, 5}}, 
 {96, {4, 5, 5, 5, 5, 5, 4, 5, 5, 5, 5, 5, 6, 5, 5, 6, 5, 5}}, 
 {96, {4, 4, 5, 4, 5, 5, 5, 5, 5, 5, 6, 6, 6, 5, 5, 5, 5, 5}}, 
 {96, {4, 5, 5, 6, 6, 5, 5, 5, 6, 5, 4, 5, 6, 6, 6, 6, 5}}, 
 {96, {4, 4, 5, 5, 5, 5, 5, 6, 6, 6, 5, 6, 6, 5, 6, 6, 5}}, 
 {96, {3, 5, 5, 5, 5, 6, 6, 5, 6, 6, 6, 6, 6, 5, 5, 5, 5}}, 
 {96, {1, 6, 6, 5, 5, 6, 5, 5, 5, 5, 6, 6, 6, 6, 5, 6, 6}}, 
 {96, {1, 6, 6, 5, 5, 5, 6, 5, 5, 6, 6, 6, 5, 5, 6, 6, 6}}, 
 {96, {1, 6, 6, 5, 5, 5, 6, 5, 5, 5, 6, 6, 6, 6, 5, 6, 6}}, 
 {96, {1, 6, 6, 5, 5, 5, 6, 5, 5, 5, 6, 6, 5, 6, 6, 6, 6}}, 
 {95, {4, 5, 5, 5, 5, 4, 5, 5, 5, 5, 6, 5, 5, 5, 6, 5, 5, 5}}, 
 {95, {4, 4, 5, 4, 5, 5, 5, 6, 6, 6, 5, 5, 5, 5, 5, 5, 5, 5}}, 
 {95, {4, 5, 6, 5, 5, 6, 5, 5, 5, 5, 5, 6, 6, 5, 6, 6, 5}}, 
 {95, {4, 4, 5, 5, 6, 6, 6, 6, 6, 6, 6, 5, 5, 4, 5, 6, 5}}, 
 {95, {1, 6, 6, 5, 5, 6, 6, 6, 6, 6, 6, 1, 6, 6, 6, 6, 6}}, 
 {95, {1, 6, 6, 5, 5, 4, 5, 6, 5, 5, 6, 6, 6, 6, 6, 6, 6}}, 
 {95, {1, 6, 6, 5, 5, 4, 5, 5, 6, 6, 6, 6, 6, 6, 5, 6, 6}}, 
 {94, {4, 5, 5, 5, 5, 5, 5, 6, 5, 5, 5, 5, 4, 5, 5, 6, 5, 5}}, 
 {94, {4, 5, 5, 5, 5, 5, 5, 5, 5, 5, 4, 5, 5, 6, 5, 5, 6, 5}}, 
 {94, {4, 5, 5, 4, 5, 5, 5, 5, 5, 5, 5, 6, 5, 5, 5, 6, 5, 5}}, 
 {94, {4, 4, 5, 5, 5, 4, 4, 5, 5, 5, 5, 6, 6, 6, 6, 5, 5, 5}}, 
 {94, {4, 4, 5, 4, 5, 5, 5, 5, 5, 5, 5, 5, 6, 6, 6, 5, 5, 5}}, 
 {94, {3, 5, 5, 5, 4, 5, 5, 6, 6, 6, 5, 5, 5, 5, 5, 5, 5, 5}}, 
 {94, {4, 5, 5, 6, 6, 6, 6, 5, 4, 5, 6, 5, 5, 5, 6, 6, 5}}, 
 {94, {4, 5, 5, 5, 6, 5, 5, 6, 6, 5, 5, 6, 5, 5, 6, 6, 5}}, 
 {94, {4, 5, 5, 5, 6, 5, 5, 6, 5, 5, 5, 6, 6, 5, 6, 6, 5}}, 
 {94, {4, 5, 4, 5, 5, 6, 6, 5, 5, 5, 6, 6, 6, 6, 6, 5, 5}}, 
 {94, {4, 4, 5, 5, 6, 5, 5, 5, 6, 6, 6, 5, 5, 6, 6, 6, 5}}, 
 {94, {4, 4, 5, 5, 5, 6, 5, 5, 6, 6, 6, 6, 5, 5, 6, 6, 5}}, 
 {94, {3, 5, 5, 5, 6, 6, 5, 5, 6, 6, 5, 5, 6, 6, 6, 5, 5}}, 
 {94, {1, 6, 6, 6, 5, 5, 4, 4, 5, 6, 6, 6, 6, 6, 6, 6, 6}}, 
 {94, {1, 6, 6, 5, 5, 6, 6, 6, 6, 5, 5, 4, 5, 6, 6, 6, 6}}, 
 {94, {1, 6, 6, 5, 5, 5, 5, 6, 6, 6, 5, 6, 6, 5, 5, 6, 6}}, 
 {93, {4, 5, 5, 5, 5, 5, 6, 5, 4, 5, 5, 5, 5, 6, 5, 5, 5, 5}}, 
 {93, {4, 5, 5, 5, 5, 5, 5, 4, 5, 5, 5, 5, 5, 6, 5, 5, 6, 5}}, 
 {93, {4, 5, 5, 4, 5, 5, 5, 5, 6, 5, 5, 5, 5, 5, 6, 5, 5, 5}}, 
 {93, {4, 5, 5, 4, 5, 5, 5, 5, 5, 5, 6, 5, 5, 6, 5, 5, 5, 5}}, 
 {93, {4, 5, 5, 4, 5, 5, 5, 5, 5, 5, 5, 6, 5, 5, 5, 5, 6, 5}}, 
 {93, {4, 4, 5, 5, 5, 5, 5, 6, 6, 5, 5, 4, 5, 6, 5, 5, 5, 5}}, 
 {93, {4, 4, 5, 5, 5, 5, 5, 5, 6, 5, 4, 5, 5, 6, 6, 5, 5, 5}}, 
 {93, {4, 4, 5, 4, 5, 5, 5, 5, 5, 5, 5, 6, 6, 6, 5, 5, 5, 5}}, 
 {93, {4, 4, 5, 4, 5, 5, 5, 5, 5, 5, 5, 5, 5, 5, 6, 6, 6, 5}}, 
 {93, {4, 5, 5, 5, 5, 6, 6, 5, 6, 6, 5, 6, 6, 5, 5, 5, 5}}, 
 {93, {4, 4, 5, 5, 5, 6, 6, 5, 6, 6, 6, 6, 5, 5, 6, 5, 5}}, 
 {93, {4, 4, 5, 5, 5, 6, 5, 5, 6, 6, 6, 5, 5, 6, 6, 6, 5}}, 
 {93, {3, 5, 5, 5, 6, 6, 5, 5, 5, 6, 6, 6, 6, 5, 5, 6, 5}}, 
 {93, {3, 5, 5, 4, 5, 5, 5, 6, 6, 6, 6, 6, 6, 6, 6, 5, 5}}, 
 {93, {1, 6, 6, 6, 6, 6, 5, 4, 4, 5, 5, 6, 6, 6, 6, 6, 6}}, 
 {93, {1, 6, 6, 5, 5, 6, 6, 6, 6, 5, 4, 5, 5, 6, 6, 6, 6}}, 
 {93, {1, 6, 6, 5, 5, 6, 5, 4, 5, 5, 6, 6, 6, 6, 6, 6, 6}}, 
 {92, {4, 5, 5, 5, 5, 5, 5, 5, 5, 5, 5, 6, 5, 5, 4, 5, 6, 5}}, 
 {92, {4, 5, 5, 5, 5, 4, 5, 5, 5, 5, 5, 4, 5, 5, 6, 6, 6, 5}}, 
 {92, {4, 4, 5, 5, 6, 5, 5, 5, 5, 5, 5, 5, 5, 5, 5, 6, 5, 5}}, 
 {92, {4, 4, 5, 5, 5, 5, 5, 6, 5, 4, 5, 5, 6, 6, 5, 5, 5, 5}}, 
 {92, {3, 5, 5, 5, 5, 5, 5, 5, 5, 5, 5, 4, 5, 5, 6, 6, 6, 5}}, 
 {92, {4, 5, 5, 6, 6, 5, 5, 5, 6, 6, 5, 4, 5, 6, 6, 6, 5}}, 
 {92, {4, 5, 5, 6, 5, 5, 6, 6, 6, 5, 4, 5, 5, 6, 6, 6, 5}}, 
 {92, {4, 4, 5, 6, 6, 5, 5, 5, 6, 6, 6, 5, 5, 5, 6, 6, 5}}, 
 {92, {3, 5, 5, 5, 5, 6, 6, 6, 5, 5, 6, 6, 6, 5, 5, 6, 5}}, 
 {92, {1, 6, 6, 6, 5, 5, 6, 6, 6, 6, 6, 1, 6, 6, 6, 6, 6}}, 
 {92, {1, 6, 6, 6, 5, 4, 5, 6, 5, 5, 5, 6, 6, 6, 6, 6, 6}}, 
 {92, {1, 6, 6, 5, 5, 6, 6, 6, 5, 5, 4, 5, 6, 6, 6, 6, 6}}, 
 {92, {1, 6, 6, 5, 5, 6, 6, 6, 5, 4, 5, 5, 6, 6, 6, 6, 6}}, 
 {92, {1, 6, 6, 5, 5, 5, 6, 6, 6, 6, 5, 5, 6, 5, 5, 6, 6}}, 
 {92, {1, 6, 6, 5, 5, 4, 5, 5, 6, 6, 5, 6, 6, 6, 6, 6, 6}}, 
 {91, {4, 5, 5, 5, 4, 5, 5, 5, 4, 5, 5, 5, 5, 5, 6, 6, 6, 5}}, 
 {91, {4, 4, 5, 5, 6, 6, 5, 5, 4, 5, 5, 5, 5, 5, 5, 5, 6, 5}}, 
 {91, {4, 5, 6, 5, 5, 5, 6, 5, 5, 5, 6, 6, 5, 5, 6, 6, 5}}, 
 {91, {4, 5, 5, 6, 5, 5, 5, 6, 5, 5, 6, 6, 6, 5, 5, 6, 5}}, 
 {91, {4, 5, 5, 6, 5, 4, 5, 6, 5, 5, 6, 6, 6, 6, 6, 5, 5}}, 
 {91, {4, 5, 5, 5, 6, 6, 6, 5, 6, 6, 5, 4, 5, 6, 6, 5, 5}}, 
 {91, {4, 5, 5, 5, 6, 6, 6, 5, 5, 4, 5, 6, 6, 5, 6, 6, 5}}, 
 {91, {4, 5, 5, 5, 6, 5, 4, 5, 6, 6, 5, 6, 6, 6, 6, 5, 5}}, 
 {91, {4, 5, 5, 4, 5, 5, 6, 6, 5, 6, 6, 5, 6, 6, 6, 5, 5}}, 
 {91, {1, 6, 6, 5, 5, 6, 6, 5, 5, 4, 5, 6, 6, 6, 6, 6, 6}}, 
 {91, {1, 6, 6, 5, 5, 5, 6, 6, 6, 6, 5, 4, 5, 6, 6, 6, 6}}, 
 {91, {1, 6, 6, 5, 5, 4, 5, 6, 6, 6, 6, 6, 5, 5, 6, 6, 6}}, 
 {90, {4, 5, 5, 5, 5, 5, 5, 5, 5, 6, 5, 5, 4, 5, 5, 6, 5, 5}}, 
 {90, {4, 5, 5, 5, 4, 5, 5, 5, 5, 5, 4, 5, 5, 5, 6, 6, 6, 5}}, 
 {90, {4, 4, 5, 5, 5, 5, 5, 6, 6, 5, 5, 4, 5, 5, 5, 5, 6, 5}}, 
 {90, {4, 4, 5, 5, 5, 5, 4, 5, 5, 6, 6, 5, 5, 5, 5, 5, 6, 5}}, 
 {90, {3, 5, 5, 6, 5, 5, 5, 5, 5, 5, 5, 5, 5, 5, 5, 5, 6, 5}}, 
 {90, {4, 5, 5, 5, 6, 5, 5, 6, 6, 6, 5, 5, 6, 5, 5, 6, 5}}, 
 {90, {4, 4, 5, 5, 6, 5, 5, 5, 5, 6, 6, 6, 6, 5, 6, 6, 5}}, 
 {90, {3, 5, 5, 6, 6, 5, 6, 6, 5, 5, 5, 5, 5, 6, 6, 6, 5}}, 
 {90, {3, 5, 5, 5, 5, 6, 6, 6, 6, 6, 6, 5, 5, 5, 6, 5, 5}}, 
 {90, {3, 5, 5, 4, 5, 5, 5, 5, 6, 6, 6, 6, 6, 6, 6, 6, 5}}, 
 {90, {1, 6, 6, 6, 6, 5, 4, 4, 5, 5, 6, 6, 6, 6, 6, 6, 6}}, 
 {90, {1, 6, 6, 6, 5, 5, 5, 6, 5, 4, 5, 6, 6, 6, 6, 6, 6}}, 
 {90, {1, 6, 6, 5, 5, 5, 6, 6, 5, 6, 6, 6, 5, 5, 5, 6, 6}}, 
 {89, {4, 5, 5, 4, 5, 5, 5, 5, 5, 5, 5, 6, 5, 5, 6, 5, 5, 5}}, 
 {89, {4, 5, 5, 4, 5, 5, 5, 5, 4, 5, 5, 5, 6, 6, 6, 5, 5, 5}}, 
 {89, {4, 4, 5, 5, 5, 5, 6, 5, 5, 5, 5, 4, 5, 5, 6, 6, 5, 5}}, 
 {89, {4, 4, 5, 5, 5, 5, 5, 5, 5, 6, 5, 4, 5, 5, 6, 6, 5, 5}}, 
 {89, {4, 4, 5, 5, 5, 4, 5, 5, 5, 6, 6, 5, 5, 5, 5, 5, 6, 5}}, 
 {89, {3, 5, 5, 5, 4, 5, 5, 5, 5, 5, 5, 6, 6, 6, 5, 5, 5, 5}}, 
 {89, {4, 5, 5, 6, 5, 5, 6, 6, 5, 5, 5, 6, 5, 5, 6, 6, 5}}, 
 {89, {4, 5, 5, 6, 5, 5, 6, 5, 5, 5, 6, 6, 6, 5, 5, 6, 5}}, 
 {89, {4, 5, 5, 6, 5, 4, 5, 6, 6, 6, 5, 5, 5, 6, 6, 6, 5}}, 
 {89, {4, 4, 5, 5, 6, 6, 5, 5, 6, 6, 6, 5, 5, 6, 6, 5, 5}}, 
 {89, {4, 4, 5, 5, 5, 5, 5, 6, 6, 5, 6, 6, 5, 6, 6, 6, 5}}, 
 {89, {3, 5, 6, 5, 5, 5, 4, 5, 5, 6, 6, 6, 6, 6, 6, 6, 5}}, 
 {89, {3, 5, 5, 6, 6, 6, 5, 5, 5, 6, 5, 5, 5, 6, 6, 6, 5}}, 
 {89, {3, 5, 5, 6, 6, 5, 5, 5, 5, 6, 6, 6, 5, 6, 6, 5, 5}}, 
 {89, {1, 6, 6, 6, 5, 4, 5, 5, 5, 6, 6, 6, 5, 6, 6, 6, 6}}, 
 {89, {1, 6, 6, 5, 5, 5, 6, 5, 5, 5, 6, 6, 6, 5, 6, 6, 6}}, 
 {89, {1, 6, 6, 5, 5, 4, 5, 6, 6, 6, 6, 5, 5, 6, 6, 6, 6}}, 
 {89, {1, 6, 6, 5, 4, 5, 6, 5, 5, 5, 6, 6, 6, 6, 6, 6, 6}}, 
 {89, {5, 5, 6, 6, 5, 6, 6, 5, 6, 6, 5, 6, 6, 5, 6, 6}}, 
 {88, {4, 5, 5, 5, 4, 5, 5, 5, 5, 5, 4, 5, 6, 6, 6, 5, 5, 5}}, 
 {88, {4, 4, 5, 5, 5, 4, 5, 6, 5, 5, 5, 5, 5, 5, 6, 6, 5, 5}}, 
 {88, {4, 4, 5, 5, 4, 4, 5, 5, 5, 5, 5, 5, 6, 6, 6, 6, 5, 5}}, 
 {88, {3, 5, 5, 4, 5, 5, 6, 6, 6, 5, 5, 5, 5, 5, 5, 5, 5, 5}}, 
 {88, {4, 5, 6, 5, 5, 6, 5, 5, 5, 6, 6, 5, 5, 5, 6, 6, 5}}, 
 {88, {4, 5, 6, 5, 5, 5, 6, 6, 5, 5, 5, 6, 5, 5, 6, 6, 5}}, 
 {88, {4, 5, 5, 6, 6, 6, 5, 5, 6, 6, 6, 5, 5, 4, 5, 6, 5}}, 
 {88, {4, 5, 5, 5, 5, 5, 4, 5, 6, 6, 6, 5, 6, 6, 6, 6, 5}}, 
 {88, {4, 4, 5, 5, 5, 5, 6, 5, 5, 6, 6, 6, 5, 6, 6, 6, 5}}, 
 {88, {4, 4, 5, 4, 5, 5, 5, 5, 6, 6, 6, 6, 6, 6, 6, 6, 5}}, 
 {88, {3, 5, 5, 6, 6, 5, 6, 6, 6, 5, 5, 5, 5, 5, 6, 6, 5}}, 
 {88, {3, 5, 5, 5, 6, 6, 6, 6, 6, 6, 5, 5, 5, 5, 6, 5, 5}}, 
 {88, {1, 6, 6, 5, 4, 5, 6, 6, 5, 5, 5, 6, 6, 6, 6, 6, 6}}, 
 {87, {4, 5, 5, 5, 5, 6, 5, 5, 4, 5, 5, 5, 5, 6, 5, 5, 5, 5}}, 
 {87, {4, 5, 5, 5, 5, 4, 5, 5, 5, 5, 4, 5, 5, 6, 6, 6, 5, 5}}, 
 {87, {4, 5, 5, 4, 5, 5, 5, 5, 6, 5, 5, 5, 5, 5, 5, 5, 6, 5}}, 
 {87, {3, 5, 5, 5, 5, 5, 5, 5, 6, 5, 5, 5, 5, 6, 5, 5, 5, 5}}, 
 {87, {4, 5, 5, 6, 6, 5, 4, 5, 6, 6, 5, 5, 5, 6, 6, 6, 5}}, 
 {87, {4, 4, 5, 5, 6, 6, 6, 6, 6, 6, 6, 5, 4, 5, 5, 6, 5}}, 
 {87, {4, 4, 5, 5, 5, 6, 5, 5, 6, 6, 6, 6, 6, 5, 5, 6, 5}}, 
 {87, {3, 5, 5, 6, 6, 5, 5, 4, 5, 5, 6, 6, 6, 6, 6, 6, 5}}, 
 {87, {3, 5, 5, 5, 6, 6, 5, 6, 6, 6, 6, 5, 5, 5, 5, 6, 5}}, 
 {87, {3, 5, 5, 5, 6, 6, 5, 5, 6, 6, 6, 6, 5, 5, 5, 6, 5}}, 
 {87, {1, 6, 6, 5, 5, 5, 6, 5, 5, 6, 5, 5, 6, 6, 6, 6, 6}}, 
 {86, {4, 4, 5, 5, 4, 5, 5, 6, 6, 5, 5, 5, 5, 5, 5, 5, 6, 5}}, 
 {86, {4, 5, 6, 6, 5, 5, 5, 6, 5, 5, 5, 6, 5, 5, 6, 6, 5}}, 
 {86, {4, 5, 6, 5, 5, 6, 5, 5, 5, 6, 5, 5, 5, 6, 6, 6, 5}}, 
 {86, {4, 5, 6, 5, 5, 5, 6, 5, 5, 6, 6, 5, 5, 5, 6, 6, 5}}, 
 {86, {4, 5, 5, 6, 6, 6, 6, 5, 5, 5, 6, 5, 4, 5, 6, 6, 5}}, 
 {86, {4, 5, 5, 5, 6, 6, 6, 5, 6, 6, 5, 4, 5, 5, 6, 6, 5}}, 
 {86, {4, 5, 5, 5, 6, 6, 5, 6, 6, 5, 4, 5, 5, 6, 6, 6, 5}}, 
 {86, {3, 5, 6, 6, 5, 5, 5, 5, 5, 6, 5, 5, 6, 6, 6, 6, 5}}, 
 {86, {1, 6, 6, 6, 6, 5, 4, 5, 5, 6, 5, 5, 6, 6, 6, 6, 6}}, 
 {85, {4, 5, 5, 4, 5, 5, 5, 4, 5, 5, 5, 6, 6, 6, 5, 5, 5, 5}}, 
 {85, {3, 5, 5, 5, 5, 5, 5, 6, 5, 5, 5, 5, 5, 5, 6, 5, 5, 5}}, 
 {85, {3, 5, 5, 5, 5, 5, 5, 5, 4, 5, 5, 5, 5, 6, 6, 6, 5, 5}}, 
 {85, {3, 5, 5, 5, 5, 4, 5, 5, 5, 5, 6, 6, 6, 5, 5, 5, 5, 5}}, 
 {85, {4, 5, 6, 6, 5, 5, 5, 5, 6, 5, 5, 6, 5, 5, 6, 6, 5}}, 
 {85, {4, 5, 6, 5, 5, 6, 6, 5, 5, 5, 6, 5, 5, 5, 6, 6, 5}}, 
 {85, {4, 5, 5, 6, 6, 6, 5, 4, 5, 6, 6, 5, 5, 5, 6, 6, 5}}, 
 {85, {4, 5, 5, 6, 5, 5, 5, 6, 6, 5, 6, 6, 5, 5, 6, 5, 5}}, 
 {85, {4, 5, 5, 6, 5, 4, 5, 6, 6, 6, 6, 5, 5, 5, 6, 6, 5}}, 
 {85, {4, 5, 5, 5, 6, 5, 4, 5, 5, 6, 6, 6, 5, 6, 6, 6, 5}}, 
 {85, {4, 4, 5, 5, 6, 5, 5, 6, 5, 5, 5, 6, 6, 6, 6, 6, 5}}, 
 {85, {4, 4, 5, 5, 6, 5, 5, 5, 6, 6, 5, 5, 6, 6, 6, 6, 5}}, 
 {85, {4, 4, 5, 5, 5, 6, 5, 5, 5, 6, 6, 5, 6, 6, 6, 6, 5}}, 
 {85, {3, 5, 5, 6, 6, 6, 5, 5, 5, 5, 6, 5, 5, 6, 6, 6, 5}}, 
 {85, {3, 5, 5, 6, 6, 5, 5, 5, 6, 5, 5, 5, 6, 6, 6, 6, 5}}, 
 {85, {3, 5, 5, 5, 5, 5, 6, 6, 6, 5, 6, 6, 6, 5, 5, 6, 5}}, 
 {85, {1, 6, 6, 5, 5, 5, 6, 6, 6, 5, 4, 5, 6, 6, 6, 6, 6}}, 
 {84, {4, 5, 5, 5, 4, 5, 6, 5, 5, 5, 5, 5, 5, 5, 5, 5, 6, 5}}, 
 {84, {4, 4, 5, 5, 6, 5, 5, 5, 5, 5, 5, 4, 5, 5, 6, 6, 5, 5}}, 
 {84, {4, 5, 6, 5, 5, 5, 6, 6, 5, 5, 6, 5, 5, 5, 6, 6, 5}}, 
 {84, {4, 5, 5, 6, 6, 5, 4, 5, 6, 6, 6, 5, 5, 5, 6, 6, 5}}, 
 {84, {4, 5, 5, 6, 5, 5, 6, 6, 5, 5, 4, 5, 6, 6, 6, 6, 5}}, 
 {84, {4, 4, 5, 5, 6, 6, 6, 6, 6, 6, 5, 5, 4, 5, 6, 6, 5}}, 
 {84, {4, 4, 5, 5, 5, 6, 6, 6, 5, 4, 5, 6, 6, 6, 6, 6, 5}}, 
 {84, {4, 4, 5, 5, 5, 5, 6, 5, 5, 6, 6, 6, 6, 5, 6, 6, 5}}, 
 {84, {3, 5, 6, 6, 6, 5, 5, 5, 4, 5, 5, 6, 6, 6, 6, 6, 5}}, 
 {84, {3, 5, 5, 5, 5, 6, 5, 5, 6, 6, 6, 6, 6, 6, 5, 5, 5}}, 
 {84, {3, 5, 5, 5, 4, 5, 5, 5, 6, 6, 6, 6, 6, 6, 6, 6, 5}}, 
 {84, {1, 6, 6, 6, 6, 6, 5, 5, 4, 4, 5, 6, 6, 6, 6, 6, 6}}, 
 {84, {1, 6, 6, 6, 5, 6, 6, 5, 5, 5, 4, 5, 6, 6, 6, 6, 6}}, 
 {84, {1, 6, 6, 5, 5, 5, 6, 6, 5, 5, 6, 6, 6, 5, 5, 6, 6}}, 
 {83, {4, 5, 5, 5, 5, 5, 5, 5, 5, 4, 5, 6, 5, 5, 5, 5, 6, 5}}, 
 {83, {4, 5, 5, 4, 5, 5, 5, 4, 5, 5, 5, 5, 6, 6, 6, 5, 5, 5}}, 
 {83, {4, 5, 4, 5, 5, 5, 5, 5, 5, 5, 5, 5, 5, 5, 6, 6, 5, 5}}, 
 {83, {4, 4, 5, 5, 5, 5, 4, 5, 5, 5, 6, 6, 5, 5, 5, 5, 6, 5}}, 
 {83, {3, 5, 5, 4, 5, 5, 5, 5, 5, 5, 5, 5, 5, 6, 6, 6, 5, 5}}, 
 {83, {1, 6, 6, 6, 5, 5, 5, 4, 4, 5, 5, 5, 5, 5, 5, 6, 6, 6}}, 
 {83, {4, 5, 5, 6, 6, 6, 6, 6, 5, 5, 5, 6, 5, 4, 5, 6, 5}}, 
 {83, {4, 5, 5, 6, 5, 5, 6, 6, 6, 5, 5, 6, 5, 5, 5, 6, 5}}, 
 {83, {4, 5, 5, 6, 5, 5, 6, 5, 5, 6, 5, 5, 5, 6, 6, 6, 5}}, 
 {83, {4, 5, 5, 5, 5, 6, 5, 5, 6, 6, 5, 6, 6, 5, 5, 6, 5}}, 
 {83, {3, 5, 6, 5, 5, 4, 5, 5, 5, 6, 6, 6, 6, 6, 6, 6, 5}}, 
 {83, {3, 5, 5, 5, 5, 4, 5, 6, 6, 6, 6, 6, 6, 6, 6, 5, 5}}, 
 {83, {1, 6, 6, 5, 5, 6, 6, 5, 4, 5, 5, 6, 6, 6, 6, 6, 6}}, 
 {82, {4, 5, 5, 5, 4, 5, 5, 5, 5, 5, 5, 4, 5, 6, 6, 6, 5, 5}}, 
 {82, {4, 5, 5, 5, 4, 5, 5, 5, 5, 4, 5, 6, 6, 6, 5, 5, 5, 5}}, 
 {82, {4, 4, 5, 5, 5, 5, 5, 5, 6, 5, 4, 5, 5, 5, 6, 6, 5, 5}}, 
 {82, {4, 4, 5, 5, 5, 4, 5, 6, 5, 5, 5, 5, 5, 6, 6, 5, 5, 5}}, 
 {82, {4, 4, 5, 5, 4, 4, 5, 5, 5, 5, 5, 6, 6, 6, 6, 5, 5, 5}}, 
 {82, {3, 5, 5, 5, 5, 5, 6, 5, 5, 5, 5, 5, 5, 5, 6, 5, 5, 5}}, 
 {82, {3, 5, 5, 5, 4, 5, 5, 5, 5, 5, 6, 6, 6, 5, 5, 5, 5, 5}}, 
 {82, {3, 5, 5, 5, 4, 5, 5, 5, 5, 5, 5, 5, 6, 6, 6, 5, 5, 5}}, 
 {82, {4, 5, 6, 5, 5, 6, 5, 5, 5, 5, 6, 5, 5, 6, 6, 6, 5}}, 
 {82, {4, 5, 6, 5, 5, 5, 5, 6, 6, 6, 5, 4, 5, 6, 6, 6, 5}}, 
 {82, {4, 5, 5, 6, 5, 5, 6, 5, 5, 5, 5, 6, 6, 5, 6, 6, 5}}, 
 {82, {4, 5, 5, 5, 6, 6, 6, 6, 6, 5, 5, 5, 4, 5, 6, 6, 5}}, 
 {82, {4, 5, 5, 5, 6, 6, 5, 6, 6, 5, 5, 6, 5, 5, 6, 5, 5}}, 
 {82, {4, 5, 5, 5, 6, 6, 5, 5, 6, 5, 5, 6, 5, 5, 6, 6, 5}}, 
 {82, {4, 5, 5, 4, 5, 5, 6, 6, 6, 5, 6, 6, 6, 5, 5, 6, 5}}, 
 {82, {4, 4, 5, 5, 6, 6, 6, 5, 5, 6, 5, 5, 6, 6, 6, 5, 5}}, 
 {82, {4, 4, 5, 5, 6, 6, 6, 5, 4, 5, 5, 6, 6, 6, 6, 6, 5}}, 
 {82, {3, 5, 6, 6, 5, 5, 5, 5, 5, 5, 6, 6, 5, 6, 6, 6, 5}}, 
 {82, {3, 5, 5, 5, 6, 6, 5, 5, 6, 6, 5, 5, 5, 6, 6, 6, 5}}, 
 {82, {1, 6, 6, 6, 5, 4, 5, 5, 5, 6, 6, 6, 6, 5, 6, 6, 6}}, 
 {82, {1, 6, 6, 5, 6, 6, 5, 4, 5, 5, 5, 6, 6, 6, 6, 6, 6}}, 
 {81, {4, 5, 5, 5, 5, 5, 5, 5, 4, 5, 5, 5, 5, 6, 5, 5, 6, 5}}, 
 {81, {4, 5, 5, 5, 5, 5, 5, 4, 5, 5, 5, 6, 5, 5, 5, 6, 5, 5}}, 
 {81, {4, 5, 5, 5, 5, 4, 5, 5, 5, 5, 5, 4, 5, 6, 6, 6, 5, 5}}, 
 {81, {4, 5, 5, 5, 5, 4, 5, 5, 5, 5, 4, 5, 5, 5, 6, 6, 6, 5}}, 
 {81, {4, 4, 5, 5, 5, 5, 6, 6, 5, 5, 4, 5, 5, 5, 5, 5, 6, 5}}, 
 {81, {4, 4, 5, 5, 5, 5, 6, 5, 5, 5, 4, 5, 5, 6, 6, 5, 5, 5}}, 
 {81, {4, 4, 5, 5, 5, 5, 5, 5, 4, 5, 5, 6, 6, 5, 5, 5, 6, 5}}, 
 {81, {3, 5, 5, 5, 5, 5, 5, 5, 5, 5, 5, 4, 5, 6, 6, 6, 5, 5}}, 
 {81, {3, 5, 5, 5, 4, 5, 5, 5, 5, 5, 5, 5, 5, 5, 6, 6, 6, 5}}, 
 {81, {3, 5, 5, 4, 5, 5, 5, 5, 5, 5, 5, 5, 5, 5, 6, 6, 6, 5}}, 
 {81, {4, 5, 6, 5, 5, 6, 6, 5, 5, 5, 5, 6, 6, 5, 5, 6, 5}}, 
 {81, {4, 5, 6, 5, 5, 5, 5, 6, 6, 5, 4, 5, 6, 6, 6, 6, 5}}, 
 {81, {4, 5, 5, 6, 6, 6, 6, 5, 5, 5, 6, 6, 5, 4, 5, 6, 5}}, 
 {81, {4, 5, 5, 6, 5, 5, 6, 6, 5, 4, 5, 5, 6, 6, 6, 6, 5}}, 
 {81, {4, 5, 5, 6, 5, 4, 5, 6, 6, 5, 5, 5, 6, 6, 6, 6, 5}}, 
 {81, {4, 5, 5, 5, 6, 6, 6, 5, 4, 5, 6, 6, 5, 6, 6, 5, 5}}, 
 {81, {4, 4, 5, 5, 6, 6, 5, 4, 5, 5, 6, 6, 6, 6, 6, 6, 5}}, 
 {81, {3, 5, 5, 5, 6, 6, 6, 5, 5, 6, 6, 6, 5, 5, 5, 6, 5}}, 
 {81, {3, 5, 5, 5, 5, 6, 6, 6, 6, 5, 6, 6, 5, 5, 5, 6, 5}}, 
 {81, {1, 6, 6, 6, 5, 4, 5, 6, 6, 5, 5, 5, 6, 6, 6, 6, 6}}, 
 {81, {1, 6, 6, 5, 5, 6, 5, 5, 5, 5, 6, 6, 5, 6, 6, 6, 6}}, 
 {80, {4, 5, 5, 5, 5, 5, 4, 5, 5, 5, 5, 6, 5, 5, 5, 6, 5, 5}}, 
 {80, {4, 4, 5, 5, 5, 6, 6, 5, 5, 4, 5, 5, 5, 5, 5, 5, 6, 5}}, 
 {80, {4, 4, 5, 5, 5, 5, 6, 5, 5, 4, 5, 5, 5, 6, 6, 5, 5, 5}}, 
 {80, {4, 4, 5, 5, 5, 5, 5, 4, 5, 6, 5, 5, 5, 5, 6, 6, 5, 5}}, 
 {80, {4, 4, 5, 5, 5, 4, 4, 5, 5, 5, 5, 5, 6, 6, 6, 6, 5, 5}}, 
 {80, {3, 5, 5, 5, 5, 5, 5, 6, 5, 5, 5, 5, 5, 6, 5, 5, 5, 5}}, 
 {80, {3, 5, 5, 5, 5, 5, 5, 4, 5, 5, 5, 5, 5, 6, 6, 6, 5, 5}}, 
 {80, {4, 5, 5, 6, 5, 4, 5, 6, 5, 5, 5, 6, 6, 6, 6, 6, 5}}, 
 {80, {4, 5, 5, 5, 6, 6, 5, 5, 4, 5, 6, 6, 6, 5, 6, 6, 5}}, 
 {80, {4, 5, 5, 5, 6, 6, 5, 5, 4, 5, 6, 6, 5, 6, 6, 6, 5}}, 
 {80, {4, 5, 5, 5, 6, 5, 5, 5, 6, 5, 5, 6, 6, 5, 6, 6, 5}}, 
 {80, {4, 5, 5, 5, 6, 5, 4, 5, 6, 6, 6, 6, 6, 5, 5, 6, 5}}, 
 {80, {4, 5, 5, 5, 6, 5, 4, 5, 5, 6, 6, 6, 6, 5, 6, 6, 5}}, 
 {80, {3, 5, 5, 5, 6, 6, 6, 6, 5, 5, 6, 6, 5, 5, 5, 6, 5}}, 
 {80, {3, 5, 5, 5, 6, 6, 5, 6, 6, 6, 5, 5, 5, 6, 6, 5, 5}}, 
 {80, {3, 5, 5, 5, 6, 5, 5, 6, 6, 6, 6, 6, 5, 5, 5, 6, 5}}, 
 {80, {3, 5, 5, 5, 5, 5, 6, 6, 6, 6, 5, 6, 6, 5, 5, 6, 5}}, 
 {80, {1, 6, 6, 6, 5, 5, 5, 4, 5, 6, 6, 6, 6, 5, 6, 6, 6}}, 
 {80, {1, 6, 6, 6, 5, 4, 4, 5, 5, 6, 6, 6, 6, 6, 6, 6, 6}}, 
 {79, {4, 5, 5, 5, 5, 4, 5, 6, 5, 5, 5, 5, 5, 5, 5, 5, 6, 5}}, 
 {79, {4, 5, 5, 4, 5, 5, 5, 5, 5, 5, 5, 5, 5, 6, 5, 5, 6, 5}}, 
 {79, {4, 4, 5, 6, 5, 5, 5, 5, 5, 5, 5, 5, 4, 5, 5, 6, 6, 5}}, 
 {79, {4, 4, 5, 5, 5, 5, 5, 5, 5, 6, 6, 5, 5, 4, 5, 6, 5, 5}}, 
 {79, {3, 5, 5, 5, 5, 6, 5, 5, 5, 5, 5, 5, 5, 5, 6, 5, 5, 5}}, 
 {79, {4, 5, 6, 5, 5, 5, 5, 6, 5, 5, 6, 5, 5, 6, 6, 6, 5}}, 
 {79, {4, 5, 5, 6, 5, 5, 6, 6, 6, 5, 4, 5, 6, 6, 6, 5, 5}}, 
 {79, {4, 5, 5, 5, 6, 6, 6, 5, 6, 6, 5, 5, 4, 5, 6, 6, 5}}, 
 {79, {4, 5, 5, 5, 6, 5, 4, 5, 5, 6, 6, 5, 6, 6, 6, 6, 5}}, 
 {79, {3, 5, 5, 6, 6, 5, 5, 6, 5, 5, 5, 5, 6, 6, 6, 6, 5}}, 
 {79, {3, 5, 5, 5, 6, 5, 5, 5, 6, 6, 6, 6, 6, 5, 5, 6, 5}}, 
 {79, {3, 5, 5, 5, 5, 6, 6, 6, 6, 5, 5, 6, 6, 5, 5, 6, 5}}, 
 {79, {1, 6, 6, 5, 6, 6, 6, 5, 5, 5, 4, 5, 6, 6, 6, 6, 6}}, 
 {79, {1, 6, 6, 5, 4, 5, 5, 5, 6, 6, 6, 6, 6, 5, 6, 6, 6}}, 
 {78, {4, 5, 5, 5, 5, 5, 5, 5, 5, 6, 5, 5, 5, 5, 4, 5, 6, 5}}, 
 {78, {4, 4, 5, 5, 5, 5, 6, 5, 5, 5, 4, 5, 5, 5, 6, 6, 5, 5}}, 
 {78, {4, 4, 5, 5, 5, 5, 5, 6, 6, 5, 5, 4, 5, 5, 6, 5, 5, 5}}, 
 {78, {3, 5, 5, 5, 5, 4, 5, 5, 6, 6, 6, 5, 5, 5, 5, 5, 5, 5}}, 
 {78, {1, 6, 6, 5, 5, 5, 5, 5, 5, 5, 4, 4, 5, 5, 6, 6, 6, 6}}, 
 {78, {4, 5, 5, 6, 6, 5, 4, 5, 6, 5, 5, 5, 6, 6, 6, 6, 5}}, 
 {78, {4, 5, 5, 6, 5, 5, 6, 6, 5, 5, 6, 5, 5, 5, 6, 6, 5}}, 
 {78, {4, 5, 5, 6, 5, 4, 5, 5, 6, 5, 5, 6, 6, 6, 6, 6, 5}}, 
 {78, {4, 5, 5, 5, 6, 6, 5, 6, 6, 6, 5, 5, 4, 5, 6, 6, 5}}, 
 {78, {4, 4, 5, 5, 5, 5, 5, 6, 6, 5, 6, 6, 6, 5, 6, 6, 5}}, 
 {78, {3, 5, 5, 6, 6, 5, 5, 5, 5, 5, 6, 6, 5, 6, 6, 6, 5}}, 
 {78, {3, 5, 5, 5, 6, 6, 6, 6, 5, 6, 6, 5, 5, 5, 5, 6, 5}}, 
 {78, {3, 5, 5, 5, 5, 6, 6, 6, 5, 6, 6, 6, 5, 5, 5, 6, 5}}, 
 {78, {1, 6, 6, 5, 5, 4, 5, 6, 6, 6, 5, 5, 6, 6, 6, 6, 6}}, 
 {77, {4, 5, 5, 4, 5, 5, 5, 5, 6, 5, 5, 5, 5, 5, 5, 6, 5, 5}}, 
 {77, {4, 5, 5, 4, 5, 5, 5, 5, 5, 5, 5, 5, 6, 5, 5, 6, 5, 5}}, 
 {77, {4, 4, 5, 5, 5, 5, 6, 6, 5, 5, 5, 5, 5, 4, 5, 5, 6, 5}}, 
 {77, {4, 4, 5, 5, 5, 5, 5, 5, 5, 6, 6, 5, 5, 4, 5, 5, 6, 5}}, 
 {77, {4, 4, 5, 5, 5, 5, 5, 4, 4, 5, 5, 5, 6, 6, 6, 6, 5, 5}}, 
 {77, {3, 5, 5, 5, 5, 5, 5, 6, 5, 5, 5, 5, 6, 5, 5, 5, 5, 5}}, 
 {77, {3, 5, 5, 5, 5, 5, 5, 5, 6, 5, 5, 5, 6, 5, 5, 5, 5, 5}}, 
 {77, {3, 5, 5, 5, 5, 5, 5, 5, 6, 5, 5, 5, 5, 5, 6, 5, 5, 5}}, 
 {77, {3, 5, 5, 5, 5, 5, 5, 5, 5, 6, 5, 5, 5, 5, 6, 5, 5, 5}}, 
 {77, {1, 6, 6, 5, 5, 5, 5, 4, 4, 5, 5, 5, 5, 5, 6, 6, 6, 6}}, 
 {77, {1, 6, 6, 1, 6, 6, 6, 5, 5, 5, 5, 5, 5, 5, 5, 6, 6, 6}}, 
 {77, {4, 5, 5, 6, 5, 5, 5, 6, 5, 5, 5, 6, 6, 5, 6, 6, 5}}, 
 {77, {4, 5, 5, 5, 6, 5, 5, 6, 5, 5, 6, 5, 5, 6, 6, 6, 5}}, 
 {77, {3, 5, 5, 6, 6, 5, 5, 5, 6, 5, 5, 6, 6, 6, 6, 5, 5}}, 
 {77, {1, 6, 6, 5, 6, 6, 6, 6, 5, 4, 5, 5, 5, 6, 6, 6, 6}}, 
 {77, {1, 6, 6, 5, 5, 6, 5, 5, 5, 6, 6, 6, 6, 5, 5, 6, 6}}, 
 {76, {4, 4, 5, 5, 5, 5, 5, 5, 6, 6, 5, 5, 4, 5, 5, 6, 5, 5}}, 
 {76, {4, 4, 5, 5, 5, 5, 5, 5, 5, 5, 6, 6, 5, 5, 4, 5, 6, 5}}, 
 {76, {4, 4, 5, 4, 5, 5, 5, 5, 6, 6, 6, 5, 5, 5, 5, 5, 5, 5}}, 
 {76, {3, 5, 5, 5, 5, 5, 5, 5, 5, 4, 5, 5, 5, 5, 6, 6, 6, 5}}, 
 {76, {3, 5, 5, 5, 5, 5, 5, 5, 4, 5, 5, 6, 6, 6, 5, 5, 5, 5}}, 
 {76, {1, 6, 6, 5, 4, 4, 5, 5, 5, 5, 5, 5, 5, 5, 6, 6, 6, 6}}, 
 {76, {4, 5, 5, 6, 5, 5, 6, 6, 6, 6, 5, 4, 5, 5, 6, 6, 5}}, 
 {76, {4, 4, 5, 6, 6, 5, 5, 5, 4, 5, 6, 6, 6, 6, 6, 6, 5}}, 
 {76, {3, 5, 5, 6, 6, 6, 5, 4, 5, 5, 6, 6, 6, 6, 6, 5, 5}}, 
 {76, {3, 5, 5, 5, 6, 6, 6, 5, 5, 6, 6, 5, 5, 5, 6, 6, 5}}, 
 {76, {3, 5, 5, 5, 6, 6, 5, 6, 6, 5, 5, 5, 5, 6, 6, 6, 5}}, 
 {76, {3, 5, 5, 5, 5, 5, 6, 6, 5, 6, 6, 6, 6, 5, 5, 6, 5}}, 
 {76, {1, 6, 6, 5, 5, 6, 5, 5, 5, 6, 5, 5, 6, 6, 6, 6, 6}}, 
 {75, {4, 5, 5, 5, 5, 5, 5, 6, 5, 4, 5, 5, 5, 5, 5, 5, 6, 5}}, 
 {75, {4, 5, 5, 5, 5, 5, 5, 5, 6, 5, 5, 5, 5, 5, 4, 5, 6, 5}}, 
 {75, {4, 4, 5, 5, 6, 6, 5, 5, 5, 5, 5, 5, 4, 5, 5, 5, 6, 5}}, 
 {75, {4, 4, 5, 5, 5, 5, 5, 6, 6, 5, 5, 4, 5, 5, 5, 6, 5, 5}}, 
 {75, {3, 5, 5, 5, 5, 4, 5, 5, 5, 5, 5, 5, 5, 5, 6, 6, 6, 5}}, 
 {75, {4, 5, 5, 6, 6, 6, 6, 5, 4, 5, 6, 6, 5, 5, 5, 6, 5}}, 
 {75, {4, 5, 5, 5, 6, 6, 6, 5, 5, 6, 5, 4, 5, 6, 6, 6, 5}}, 
 {75, {4, 5, 5, 5, 4, 5, 5, 6, 6, 5, 6, 6, 6, 5, 6, 6, 5}}, 
 {75, {4, 4, 5, 5, 6, 6, 5, 4, 5, 6, 6, 6, 6, 6, 6, 5, 5}}, 
 {75, {4, 4, 5, 5, 6, 5, 4, 5, 6, 6, 6, 6, 6, 6, 6, 5, 5}}, 
 {75, {3, 5, 5, 6, 6, 5, 5, 5, 5, 6, 5, 5, 6, 6, 6, 6, 5}}, 
 {75, {3, 5, 5, 5, 6, 6, 6, 5, 5, 5, 5, 6, 6, 5, 6, 6, 5}}, 
 {75, {1, 6, 6, 6, 5, 5, 5, 4, 5, 6, 6, 6, 5, 6, 6, 6, 6}}, 
 {75, {1, 6, 6, 5, 4, 5, 5, 6, 6, 6, 6, 6, 6, 5, 5, 6, 6}}, 
 {74, {4, 5, 5, 5, 5, 4, 5, 5, 5, 6, 5, 4, 5, 5, 6, 6, 5, 5}}, 
 {74, {4, 4, 5, 5, 6, 6, 5, 5, 5, 4, 5, 5, 5, 5, 5, 5, 6, 5}}, 
 {74, {4, 4, 5, 5, 5, 5, 5, 5, 5, 5, 6, 5, 4, 5, 5, 6, 6, 5}}, 
 {74, {4, 4, 5, 5, 5, 5, 4, 5, 5, 6, 6, 5, 5, 5, 5, 6, 5, 5}}, 
 {74, {4, 4, 5, 5, 5, 4, 5, 5, 6, 6, 5, 5, 5, 5, 6, 5, 5, 5}}, 
 {74, {3, 5, 5, 5, 5, 5, 5, 5, 5, 4, 5, 5, 5, 6, 6, 6, 5, 5}}, 
 {74, {3, 5, 5, 5, 5, 5, 4, 5, 5, 6, 6, 6, 5, 5, 5, 5, 5, 5}}, 
 {74, {4, 5, 5, 6, 5, 5, 6, 6, 6, 6, 5, 4, 5, 6, 6, 5, 5}}, 
 {74, {4, 5, 5, 5, 6, 6, 6, 5, 4, 5, 5, 6, 6, 5, 6, 6, 5}}, 
 {74, {3, 5, 5, 6, 6, 6, 5, 5, 4, 5, 6, 6, 6, 6, 6, 5, 5}}, 
 {74, {3, 5, 5, 5, 5, 6, 6, 6, 5, 5, 5, 6, 6, 5, 6, 6, 5}}, 
 {74, {3, 5, 5, 5, 5, 5, 6, 6, 6, 6, 6, 6, 5, 5, 6, 5, 5}}, 
 {74, {3, 5, 5, 4, 5, 5, 6, 6, 6, 6, 6, 6, 6, 6, 5, 5, 5}}, 
 {74, {1, 6, 6, 5, 4, 5, 6, 6, 6, 6, 6, 6, 5, 5, 5, 6, 6}}, 
 {73, {4, 4, 5, 5, 5, 5, 5, 5, 6, 6, 5, 5, 5, 4, 5, 6, 5, 5}}, 
 {73, {4, 4, 5, 5, 5, 5, 5, 5, 5, 4, 5, 5, 6, 6, 5, 5, 6, 5}}, 
 {73, {4, 4, 5, 5, 5, 5, 5, 5, 4, 5, 6, 5, 5, 5, 6, 6, 5, 5}}, 
 {73, {4, 4, 5, 5, 4, 5, 5, 5, 6, 5, 5, 5, 5, 6, 6, 5, 5, 5}}, 
 {73, {3, 5, 5, 4, 5, 5, 5, 5, 5, 6, 6, 6, 5, 5, 5, 5, 5, 5}}, 
 {73, {1, 6, 6, 5, 5, 5, 5, 5, 4, 4, 5, 5, 5, 5, 6, 6, 6, 6}}, 
 {73, {4, 5, 5, 5, 4, 5, 5, 6, 6, 6, 5, 6, 6, 5, 6, 6, 5}}, 
 {73, {4, 4, 5, 6, 5, 5, 5, 6, 6, 6, 5, 6, 6, 5, 5, 6, 5}}, 
 {73, {4, 4, 5, 5, 6, 5, 5, 5, 6, 5, 5, 6, 6, 6, 6, 6, 5}}, 
 {73, {4, 4, 5, 5, 4, 5, 5, 6, 6, 6, 6, 6, 6, 5, 6, 6, 5}}, 
 {73, {3, 5, 6, 5, 5, 5, 6, 6, 6, 5, 5, 6, 6, 5, 5, 6, 5}}, 
 {73, {1, 6, 6, 6, 6, 5, 5, 3, 5, 5, 6, 6, 6, 6, 6, 6, 6}}, 
 {73, {1, 6, 6, 6, 5, 6, 6, 5, 4, 5, 5, 5, 6, 6, 6, 6, 6}}, 
 {73, {1, 6, 6, 6, 1, 6, 6, 6, 6, 6, 5, 5, 6, 6, 6, 6, 6}}, 
 {72, {4, 5, 5, 5, 5, 5, 5, 5, 4, 5, 6, 5, 5, 5, 5, 5, 6, 5}}, 
 {72, {4, 4, 5, 5, 6, 6, 5, 5, 5, 5, 4, 5, 5, 5, 5, 5, 6, 5}}, 
 {72, {4, 4, 5, 5, 5, 5, 6, 5, 5, 4, 5, 5, 5, 5, 6, 6, 5, 5}}, 
 {72, {4, 4, 5, 5, 5, 5, 5, 6, 5, 4, 5, 5, 5, 5, 6, 6, 5, 5}}, 
 {72, {4, 4, 5, 5, 5, 5, 4, 4, 5, 5, 5, 5, 6, 6, 6, 6, 5, 5}}, 
 {72, {3, 5, 5, 4, 5, 5, 5, 6, 6, 6, 5, 5, 5, 5, 5, 5, 5, 5}}, 
 {72, {3, 5, 5, 4, 5, 5, 5, 5, 5, 5, 6, 6, 6, 5, 5, 5, 5, 5}}, 
 {72, {1, 6, 6, 6, 5, 4, 4, 5, 5, 5, 5, 5, 5, 5, 5, 6, 6, 6}}, 
 {72, {4, 5, 5, 6, 5, 5, 6, 6, 5, 4, 5, 6, 6, 6, 6, 5, 5}}, 
 {72, {4, 5, 5, 6, 5, 5, 6, 5, 4, 5, 6, 6, 6, 6, 6, 5, 5}}, 
 {72, {4, 5, 5, 5, 6, 6, 5, 6, 6, 6, 6, 5, 5, 4, 5, 6, 5}}, 
 {72, {4, 5, 5, 5, 6, 6, 5, 4, 5, 6, 6, 5, 6, 6, 6, 5, 5}}, 
 {72, {4, 5, 5, 5, 6, 6, 5, 4, 5, 5, 6, 6, 5, 6, 6, 6, 5}}, 
 {72, {4, 4, 5, 5, 6, 6, 5, 5, 4, 5, 6, 6, 6, 6, 6, 6, 5}}, 
 {72, {4, 4, 5, 5, 4, 5, 5, 6, 6, 6, 5, 6, 6, 6, 6, 6, 5}}, 
 {72, {3, 5, 6, 6, 5, 5, 4, 5, 5, 5, 6, 6, 6, 6, 6, 6, 5}}, 
 {72, {3, 5, 6, 5, 5, 5, 5, 6, 6, 6, 5, 6, 6, 5, 5, 6, 5}}, 
 {72, {3, 5, 5, 5, 5, 6, 6, 5, 6, 6, 6, 6, 5, 5, 5, 6, 5}}, 
 {72, {1, 6, 6, 6, 6, 5, 5, 6, 5, 5, 4, 5, 6, 6, 6, 6, 6}}, 
 {72, {1, 6, 6, 5, 5, 5, 6, 5, 5, 6, 6, 6, 6, 5, 5, 6, 6}}, 
 {72, {1, 6, 6, 5, 4, 4, 5, 5, 6, 6, 6, 6, 6, 6, 6, 6, 6}}, 
 {71, {4, 5, 4, 5, 5, 5, 5, 5, 5, 5, 5, 5, 5, 6, 6, 5, 5, 5}}, 
 {71, {4, 4, 5, 5, 5, 6, 5, 5, 5, 5, 5, 4, 5, 5, 6, 6, 5, 5}}, 
 {71, {4, 4, 5, 5, 5, 5, 6, 6, 5, 5, 5, 4, 5, 5, 5, 6, 5, 5}}, 
 {71, {4, 4, 5, 5, 5, 5, 5, 6, 5, 5, 4, 5, 5, 5, 6, 6, 5, 5}}, 
 {71, {4, 4, 5, 5, 5, 5, 5, 4, 4, 5, 5, 5, 5, 6, 6, 6, 6, 5}}, 
 {71, {4, 4, 5, 5, 5, 5, 4, 5, 6, 5, 5, 5, 5, 5, 6, 6, 5, 5}}, 
 {71, {4, 4, 5, 5, 5, 5, 4, 5, 5, 6, 6, 5, 5, 5, 6, 5, 5, 5}}, 
 {71, {4, 4, 5, 5, 5, 4, 5, 5, 6, 6, 5, 5, 5, 5, 5, 5, 6, 5}}, 
 {71, {4, 4, 5, 5, 4, 5, 5, 6, 6, 5, 5, 5, 5, 6, 5, 5, 5, 5}}, 
 {71, {4, 4, 5, 5, 4, 5, 5, 6, 6, 5, 5, 5, 5, 5, 6, 5, 5, 5}}, 
 {71, {3, 5, 5, 5, 5, 4, 5, 5, 5, 6, 6, 6, 5, 5, 5, 5, 5, 5}}, 
 {71, {3, 5, 5, 5, 4, 5, 5, 6, 6, 5, 5, 5, 5, 5, 5, 5, 6, 5}}, 
 {71, {4, 5, 6, 5, 5, 5, 6, 5, 5, 5, 6, 5, 5, 6, 6, 6, 5}}, 
 {71, {4, 5, 5, 5, 5, 6, 6, 5, 6, 6, 5, 5, 6, 5, 5, 6, 5}}, 
 {71, {4, 4, 5, 5, 6, 5, 5, 5, 5, 6, 6, 5, 6, 6, 6, 6, 5}}, 
 {71, {3, 5, 6, 6, 6, 5, 5, 4, 5, 5, 5, 6, 6, 6, 6, 6, 5}}, 
 {71, {3, 5, 5, 5, 6, 6, 6, 6, 6, 5, 5, 5, 6, 5, 5, 6, 5}}, 
 {71, {3, 5, 5, 5, 6, 6, 6, 5, 5, 5, 6, 6, 5, 6, 6, 5, 5}}, 
 {71, {3, 5, 5, 5, 5, 6, 6, 5, 5, 6, 6, 6, 6, 5, 5, 6, 5}}, 
 {71, {1, 6, 6, 6, 6, 5, 4, 5, 5, 5, 6, 6, 5, 6, 6, 6, 6}}, 
 {71, {1, 6, 6, 6, 5, 6, 6, 6, 5, 5, 5, 4, 5, 6, 6, 6, 6}}, 
 {71, {1, 6, 6, 5, 5, 6, 6, 6, 6, 6, 5, 5, 4, 5, 6, 6, 6}}, 
 {71, {1, 6, 6, 5, 5, 6, 6, 6, 6, 6, 5, 4, 5, 5, 6, 6, 6}}, 
 {71, {1, 6, 6, 5, 5, 6, 6, 6, 6, 6, 1, 6, 6, 6, 6, 6, 6}}, 
 {70, {4, 5, 5, 4, 5, 5, 5, 5, 5, 5, 5, 4, 5, 5, 6, 6, 6, 5}}, 
 {70, {4, 5, 5, 4, 5, 5, 4, 5, 5, 5, 5, 5, 6, 6, 6, 5, 5, 5}}, 
 {70, {4, 5, 5, 4, 5, 5, 4, 5, 5, 5, 5, 5, 5, 6, 6, 6, 5, 5}}, 
 {70, {4, 4, 5, 6, 5, 5, 4, 5, 5, 5, 5, 5, 5, 5, 5, 6, 6, 5}}, 
 {70, {4, 4, 5, 5, 5, 6, 5, 5, 5, 5, 4, 5, 5, 6, 6, 5, 5, 5}}, 
 {70, {4, 4, 5, 5, 5, 5, 6, 5, 5, 5, 5, 5, 4, 5, 5, 6, 6, 5}}, 
 {70, {4, 4, 5, 5, 5, 5, 5, 5, 6, 5, 5, 4, 5, 5, 6, 6, 5, 5}}, 
 {70, {4, 4, 5, 5, 5, 4, 5, 6, 5, 5, 5, 6, 6, 5, 5, 5, 5, 5}}, 
 {70, {3, 5, 6, 5, 5, 5, 5, 5, 5, 5, 5, 5, 4, 5, 5, 6, 6, 5}}, 
 {70, {3, 5, 5, 5, 5, 5, 5, 4, 5, 5, 6, 6, 6, 5, 5, 5, 5, 5}}, 
 {70, {3, 5, 5, 4, 5, 5, 5, 5, 5, 5, 5, 6, 6, 6, 5, 5, 5, 5}}, 
 {70, {1, 6, 6, 6, 5, 5, 4, 5, 5, 5, 5, 4, 5, 5, 5, 6, 6, 6}}, 
 {70, {1, 6, 6, 5, 5, 4, 5, 5, 5, 5, 5, 4, 5, 5, 6, 6, 6, 6}}, 
 {70, {4, 5, 5, 6, 6, 6, 6, 6, 5, 5, 4, 5, 6, 5, 5, 6, 5}}, 
 {70, {4, 5, 5, 6, 5, 5, 6, 6, 6, 6, 6, 5, 5, 4, 5, 6, 5}}, 
 {70, {4, 5, 5, 5, 6, 6, 6, 5, 6, 6, 6, 5, 4, 5, 5, 6, 5}}, 
 {70, {4, 5, 5, 5, 5, 6, 5, 5, 6, 5, 5, 6, 6, 5, 6, 6, 5}}, 
 {70, {4, 4, 5, 5, 5, 6, 6, 6, 6, 5, 4, 5, 6, 6, 6, 6, 5}}, 
 {70, {3, 5, 5, 5, 6, 6, 5, 6, 6, 6, 5, 5, 5, 5, 6, 6, 5}}, 
 {70, {3, 5, 5, 5, 5, 6, 6, 5, 5, 6, 6, 5, 5, 6, 6, 6, 5}}, 
 {70, {3, 5, 5, 5, 5, 5, 6, 6, 6, 5, 5, 6, 6, 5, 6, 6, 5}}, 
 {70, {1, 6, 6, 6, 5, 6, 6, 6, 5, 4, 5, 5, 5, 6, 6, 6, 6}}, 
 {70, {1, 6, 6, 6, 5, 5, 5, 4, 5, 6, 6, 5, 6, 6, 6, 6, 6}}, 
 {70, {1, 6, 6, 5, 6, 6, 6, 6, 5, 5, 5, 4, 5, 6, 6, 6, 6}}, 
 {70, {1, 6, 6, 5, 5, 6, 6, 6, 6, 1, 6, 6, 6, 6, 6, 6, 6}}, 
 {69, {4, 5, 5, 5, 5, 5, 5, 6, 5, 4, 5, 6, 5, 5, 5, 5, 5, 5}}, 
 {69, {4, 4, 5, 6, 5, 5, 5, 5, 5, 4, 5, 5, 5, 5, 5, 6, 6, 5}}, 
 {69, {4, 4, 5, 5, 6, 6, 5, 5, 5, 5, 5, 5, 5, 4, 5, 5, 6, 5}}, 
 {69, {4, 4, 5, 5, 5, 4, 5, 5, 6, 6, 5, 5, 5, 6, 5, 5, 5, 5}}, 
 {69, {4, 4, 5, 5, 5, 4, 5, 5, 5, 5, 6, 6, 5, 5, 5, 5, 6, 5}}, 
 {69, {3, 5, 5, 5, 5, 5, 5, 5, 4, 5, 5, 5, 5, 5, 6, 6, 6, 5}}, 
 {69, {3, 5, 5, 5, 4, 5, 6, 6, 6, 5, 5, 5, 5, 5, 5, 5, 5, 5}}, 
 {69, {1, 6, 6, 5, 5, 5, 4, 4, 5, 5, 5, 5, 5, 5, 6, 6, 6, 6}}, 
 {69, {4, 5, 6, 5, 5, 5, 5, 6, 6, 6, 6, 5, 4, 5, 6, 6, 5}}, 
 {69, {4, 5, 5, 5, 6, 6, 6, 6, 5, 6, 6, 5, 5, 4, 5, 6, 5}}, 
 {69, {4, 5, 5, 5, 6, 6, 5, 6, 6, 6, 5, 4, 5, 5, 6, 6, 5}}, 
 {69, {4, 4, 5, 5, 4, 5, 5, 6, 6, 6, 6, 5, 6, 6, 6, 6, 5}}, 
 {69, {3, 5, 6, 5, 5, 5, 6, 6, 5, 5, 6, 6, 6, 5, 5, 6, 5}}, 
 {69, {3, 5, 5, 6, 6, 6, 6, 5, 5, 6, 5, 5, 5, 5, 6, 6, 5}}, 
 {69, {3, 5, 5, 6, 6, 6, 6, 5, 4, 5, 5, 6, 6, 6, 6, 5, 5}}, 
 {69, {3, 5, 5, 5, 6, 6, 5, 5, 6, 6, 6, 5, 5, 5, 6, 6, 5}}, 
 {69, {3, 5, 5, 5, 6, 5, 5, 5, 6, 6, 6, 5, 6, 6, 6, 5, 5}}, 
 {69, {1, 6, 6, 6, 6, 6, 5, 5, 3, 5, 5, 6, 6, 6, 6, 6, 6}}, 
 {69, {1, 6, 6, 5, 5, 6, 6, 6, 6, 6, 6, 6, 1, 6, 6, 6, 6}}, 
 {68, {4, 5, 5, 5, 4, 5, 5, 5, 5, 6, 5, 4, 5, 5, 6, 6, 5, 5}}, 
 {68, {4, 4, 5, 5, 5, 5, 6, 6, 5, 5, 5, 5, 4, 5, 5, 5, 6, 5}}, 
 {68, {4, 4, 5, 5, 5, 5, 6, 6, 5, 5, 4, 5, 5, 5, 6, 5, 5, 5}}, 
 {68, {4, 4, 5, 5, 5, 4, 5, 5, 6, 6, 5, 5, 5, 5, 5, 6, 5, 5}}, 
 {68, {4, 4, 5, 5, 5, 4, 5, 5, 5, 6, 5, 5, 5, 5, 6, 6, 5, 5}}, 
 {68, {4, 4, 5, 5, 4, 5, 5, 5, 6, 5, 5, 5, 5, 5, 6, 6, 5, 5}}, 
 {68, {4, 4, 5, 5, 4, 5, 5, 5, 5, 6, 6, 5, 5, 5, 5, 5, 6, 5}}, 
 {68, {3, 5, 5, 6, 6, 5, 5, 4, 5, 5, 5, 5, 5, 5, 5, 5, 6, 5}}, 
 {68, {3, 5, 5, 5, 5, 5, 5, 5, 5, 5, 4, 5, 5, 5, 6, 6, 6, 5}}, 
 {68, {3, 5, 5, 5, 5, 5, 5, 5, 5, 4, 5, 5, 6, 6, 6, 5, 5, 5}}, 
 {68, {3, 5, 5, 5, 5, 5, 5, 4, 5, 6, 6, 6, 5, 5, 5, 5, 5, 5}}, 
 {68, {5, 5, 5, 6, 5, 5, 6, 5, 5, 5, 6, 5, 5, 6, 5, 5, 6}}, 
 {68, {4, 5, 5, 6, 5, 5, 6, 5, 5, 6, 6, 6, 5, 5, 6, 5, 5}}, 
 {68, {4, 5, 5, 6, 5, 5, 5, 6, 5, 5, 6, 5, 5, 6, 6, 6, 5}}, 
 {68, {4, 5, 5, 5, 6, 6, 5, 5, 6, 5, 4, 5, 6, 6, 6, 6, 5}}, 
 {68, {4, 5, 5, 5, 6, 6, 5, 4, 5, 6, 6, 6, 5, 6, 6, 5, 5}}, 
 {68, {3, 5, 5, 5, 6, 6, 5, 6, 6, 5, 5, 5, 6, 6, 6, 5, 5}}, 
 {68, {3, 5, 5, 5, 5, 6, 6, 6, 5, 5, 6, 6, 5, 6, 6, 5, 5}}, 
 {68, {3, 5, 5, 5, 4, 5, 6, 6, 6, 6, 6, 6, 6, 6, 5, 5, 5}}, 
 {68, {3, 5, 5, 4, 5, 6, 6, 6, 6, 6, 6, 6, 6, 5, 5, 5, 5}}, 
 {68, {1, 6, 6, 5, 4, 5, 5, 5, 6, 6, 6, 6, 5, 6, 6, 6, 6}}, 
 {68, {1, 6, 6, 1, 6, 6, 5, 5, 6, 6, 6, 6, 6, 6, 6, 6, 6}}, 
 {67, {4, 5, 5, 4, 5, 5, 5, 5, 5, 5, 5, 4, 5, 6, 6, 6, 5, 5}}, 
 {67, {4, 4, 5, 5, 5, 6, 6, 5, 5, 5, 5, 5, 5, 4, 5, 6, 5, 5}}, 
 {67, {4, 4, 5, 5, 5, 6, 5, 5, 5, 4, 5, 5, 5, 5, 6, 6, 5, 5}}, 
 {67, {4, 4, 5, 5, 5, 5, 6, 6, 5, 5, 5, 4, 5, 5, 5, 5, 6, 5}}, 
 {67, {4, 4, 5, 5, 5, 5, 5, 6, 6, 5, 5, 5, 4, 5, 5, 5, 6, 5}}, 
 {67, {4, 4, 5, 5, 5, 5, 5, 6, 5, 5, 5, 5, 4, 5, 5, 6, 6, 5}}, 
 {67, {4, 4, 5, 5, 5, 5, 5, 6, 5, 5, 5, 4, 5, 5, 6, 6, 5, 5}}, 
 {67, {4, 4, 5, 5, 5, 5, 5, 5, 6, 6, 5, 5, 5, 4, 5, 5, 6, 5}}, 
 {67, {4, 4, 5, 5, 5, 5, 5, 5, 5, 6, 6, 5, 5, 5, 4, 5, 6, 5}}, 
 {67, {4, 4, 5, 5, 5, 5, 4, 5, 5, 5, 6, 5, 5, 5, 6, 6, 5, 5}}, 
 {67, {3, 5, 5, 5, 5, 5, 5, 5, 6, 5, 5, 6, 5, 5, 5, 5, 5, 5}}, 
 {67, {3, 5, 5, 5, 5, 5, 5, 5, 5, 5, 5, 6, 5, 5, 6, 5, 5, 5}}, 
 {67, {3, 5, 5, 4, 5, 5, 5, 5, 6, 6, 6, 5, 5, 5, 5, 5, 5, 5}}, 
 {67, {4, 5, 5, 6, 5, 5, 6, 5, 4, 5, 5, 6, 6, 6, 6, 6, 5}}, 
 {67, {4, 5, 5, 5, 6, 6, 6, 6, 5, 6, 6, 5, 4, 5, 5, 6, 5}}, 
 {67, {4, 5, 5, 5, 6, 6, 6, 5, 4, 5, 6, 6, 6, 5, 5, 6, 5}}, 
 {67, {4, 5, 5, 5, 6, 6, 5, 5, 6, 6, 5, 4, 5, 6, 6, 6, 5}}, 
 {67, {4, 5, 5, 4, 5, 5, 6, 6, 6, 5, 5, 6, 6, 5, 6, 6, 5}}, 
 {67, {3, 5, 5, 5, 5, 6, 6, 6, 5, 6, 6, 5, 5, 5, 6, 6, 5}}, 
 {67, {1, 6, 6, 6, 6, 5, 5, 4, 4, 5, 6, 6, 6, 6, 6, 6, 6}}, 
 {67, {1, 6, 6, 6, 5, 5, 5, 6, 6, 5, 4, 5, 6, 6, 6, 6, 6}}, 
 {66, {4, 5, 5, 5, 5, 4, 5, 5, 5, 6, 5, 5, 5, 5, 6, 5, 5, 5}}, 
 {66, {4, 5, 5, 4, 5, 5, 5, 5, 5, 6, 5, 5, 5, 6, 5, 5, 5, 5}}, 
 {66, {4, 5, 5, 4, 5, 5, 5, 5, 4, 5, 6, 6, 6, 5, 5, 5, 5, 5}}, 
 {66, {4, 4, 5, 6, 5, 4, 5, 5, 5, 5, 5, 5, 5, 5, 5, 6, 6, 5}}, 
 {66, {4, 4, 5, 5, 5, 5, 5, 6, 5, 5, 4, 5, 5, 6, 6, 5, 5, 5}}, 
 {66, {4, 4, 5, 5, 5, 4, 5, 5, 5, 6, 6, 5, 5, 5, 6, 5, 5, 5}}, 
 {66, {4, 4, 5, 5, 4, 5, 6, 5, 5, 5, 5, 5, 6, 6, 5, 5, 5, 5}}, 
 {66, {3, 5, 5, 5, 5, 5, 5, 5, 5, 6, 5, 5, 6, 5, 5, 5, 5, 5}}, 
 {66, {1, 6, 6, 5, 5, 4, 5, 6, 5, 5, 5, 5, 5, 5, 5, 5, 6, 6}}, 
 {66, {4, 5, 5, 5, 6, 6, 6, 5, 6, 6, 6, 5, 4, 5, 6, 5, 5}}, 
 {66, {4, 5, 5, 5, 6, 6, 5, 4, 5, 5, 6, 6, 6, 5, 6, 6, 5}}, 
 {66, {4, 5, 5, 5, 4, 5, 6, 5, 5, 6, 6, 6, 5, 6, 6, 6, 5}}, 
 {66, {4, 4, 5, 5, 6, 5, 4, 5, 5, 6, 6, 6, 6, 6, 6, 6, 5}}, 
 {66, {4, 4, 5, 5, 4, 5, 5, 6, 6, 5, 6, 6, 6, 6, 6, 6, 5}}, 
 {66, {4, 4, 5, 4, 5, 5, 6, 6, 6, 5, 5, 6, 6, 6, 6, 6, 5}}, 
 {66, {3, 5, 6, 6, 5, 5, 5, 4, 5, 5, 6, 6, 6, 6, 6, 6, 5}}, 
 {66, {3, 5, 5, 6, 6, 6, 6, 6, 6, 6, 5, 4, 5, 5, 5, 6, 5}}, 
 {66, {3, 5, 5, 5, 6, 6, 5, 5, 5, 6, 6, 6, 5, 5, 6, 6, 5}}, 
 {66, {3, 5, 5, 5, 5, 6, 6, 6, 6, 6, 5, 5, 6, 5, 5, 6, 5}}, 
 {66, {3, 5, 5, 5, 5, 6, 6, 5, 6, 6, 5, 5, 6, 6, 6, 5, 5}}, 
 {66, {3, 5, 5, 4, 5, 5, 6, 6, 5, 5, 6, 6, 6, 6, 6, 6, 5}}, 
 {66, {1, 6, 6, 6, 6, 5, 5, 5, 4, 5, 6, 6, 5, 6, 6, 6, 6}}, 
 {66, {1, 6, 6, 6, 5, 4, 5, 5, 5, 6, 6, 5, 6, 6, 6, 6, 6}}, 
 {65, {4, 5, 5, 5, 4, 5, 5, 5, 6, 5, 5, 5, 5, 5, 6, 5, 5, 5}}, 
 {65, {4, 4, 5, 6, 5, 5, 5, 5, 4, 5, 5, 5, 5, 5, 5, 6, 6, 5}}, 
 {65, {4, 4, 5, 5, 5, 6, 5, 5, 5, 5, 5, 5, 4, 5, 5, 6, 6, 5}}, 
 {65, {4, 4, 5, 5, 5, 5, 6, 6, 5, 5, 5, 5, 4, 5, 6, 5, 5, 5}}, 
 {65, {4, 4, 5, 5, 5, 5, 5, 6, 6, 5, 5, 5, 5, 4, 5, 5, 6, 5}}, 
 {65, {4, 4, 5, 5, 5, 5, 5, 6, 5, 4, 5, 5, 5, 6, 6, 5, 5, 5}}, 
 {65, {4, 4, 5, 5, 5, 5, 5, 5, 5, 6, 6, 5, 4, 5, 5, 5, 6, 5}}, 
 {65, {4, 4, 5, 5, 5, 4, 5, 6, 5, 5, 5, 5, 6, 6, 5, 5, 5, 5}}, 
 {65, {4, 4, 5, 5, 5, 4, 5, 5, 6, 5, 5, 5, 5, 6, 6, 5, 5, 5}}, 
 {65, {4, 4, 5, 5, 5, 4, 5, 5, 5, 5, 5, 6, 6, 5, 5, 5, 6, 5}}, 
 {65, {3, 5, 5, 6, 6, 5, 5, 5, 5, 5, 5, 5, 4, 5, 5, 5, 6, 5}}, 
 {65, {3, 5, 5, 4, 5, 5, 6, 6, 5, 5, 5, 5, 5, 5, 5, 5, 6, 5}}, 
 {65, {1, 6, 6, 6, 5, 5, 4, 5, 5, 4, 5, 5, 5, 5, 5, 6, 6, 6}}, 
 {65, {4, 5, 6, 6, 5, 4, 5, 6, 6, 5, 5, 5, 5, 6, 6, 6, 5}}, 
 {65, {4, 5, 6, 5, 5, 5, 6, 6, 5, 5, 6, 6, 5, 5, 5, 6, 5}}, 
 {65, {4, 5, 5, 6, 5, 5, 4, 5, 5, 6, 6, 6, 6, 5, 6, 6, 5}}, 
 {65, {4, 5, 5, 6, 5, 5, 4, 5, 5, 6, 6, 6, 5, 6, 6, 6, 5}}, 
 {65, {4, 5, 5, 5, 6, 6, 6, 6, 6, 5, 4, 5, 6, 5, 5, 6, 5}}, 
 {65, {4, 5, 5, 5, 6, 6, 5, 6, 6, 5, 5, 4, 5, 6, 6, 6, 5}}, 
 {65, {4, 5, 5, 5, 6, 5, 4, 5, 6, 5, 5, 6, 6, 6, 6, 6, 5}}, 
 {65, {4, 5, 5, 5, 4, 5, 6, 5, 5, 6, 6, 5, 6, 6, 6, 6, 5}}, 
 {65, {4, 5, 5, 4, 5, 5, 6, 6, 6, 6, 5, 6, 6, 5, 5, 6, 5}}, 
 {65, {4, 5, 5, 4, 5, 5, 6, 6, 6, 5, 6, 6, 5, 5, 6, 6, 5}}, 
 {65, {4, 4, 5, 4, 5, 6, 6, 6, 6, 6, 6, 6, 6, 5, 5, 5, 5}}, 
 {65, {3, 5, 5, 5, 6, 6, 5, 5, 6, 5, 5, 6, 6, 6, 6, 5, 5}}, 
 {65, {1, 6, 6, 6, 5, 5, 3, 5, 5, 6, 6, 6, 6, 6, 6, 6, 6}}, 
 {65, {1, 6, 6, 5, 4, 5, 5, 5, 6, 6, 6, 5, 6, 6, 6, 6, 6}}, 
 {64, {4, 5, 5, 5, 5, 5, 4, 5, 6, 5, 5, 5, 5, 5, 5, 5, 6, 5}}, 
 {64, {4, 5, 5, 5, 5, 5, 4, 5, 5, 6, 5, 4, 5, 5, 6, 6, 5, 5}}, 
 {64, {4, 4, 5, 6, 5, 5, 5, 5, 5, 5, 4, 5, 5, 5, 5, 6, 6, 5}}, 
 {64, {4, 4, 5, 6, 5, 5, 5, 4, 5, 5, 5, 5, 5, 5, 5, 6, 6, 5}}, 
 {64, {4, 4, 5, 5, 5, 5, 5, 6, 6, 5, 5, 5, 5, 5, 4, 5, 6, 5}}, 
 {64, {4, 4, 5, 5, 5, 5, 5, 5, 6, 5, 5, 5, 4, 5, 5, 6, 6, 5}}, 
 {64, {4, 4, 5, 5, 5, 5, 5, 5, 5, 6, 5, 4, 5, 5, 5, 6, 6, 5}}, 
 {64, {3, 5, 5, 6, 6, 5, 5, 5, 4, 5, 5, 5, 5, 5, 5, 5, 6, 5}}, 
 {64, {3, 5, 5, 5, 5, 5, 4, 5, 5, 5, 5, 5, 5, 5, 6, 6, 6, 5}}, 
 {64, {1, 6, 6, 6, 5, 5, 4, 5, 5, 5, 4, 5, 5, 5, 5, 6, 6, 6}}, 
 {64, {4, 5, 5, 5, 6, 6, 5, 6, 6, 6, 6, 5, 4, 5, 5, 6, 5}}, 
 {64, {4, 5, 5, 5, 6, 6, 5, 6, 6, 6, 5, 4, 5, 6, 6, 5, 5}}, 
 {64, {4, 5, 5, 5, 4, 5, 5, 6, 6, 5, 6, 6, 5, 6, 6, 6, 5}}, 
 {64, {3, 5, 5, 5, 6, 6, 5, 5, 6, 6, 6, 5, 5, 6, 6, 5, 5}}, 
 {64, {3, 5, 5, 5, 6, 5, 5, 6, 6, 6, 5, 5, 5, 6, 6, 6, 5}}, 
 {64, {3, 5, 5, 5, 6, 5, 5, 6, 6, 5, 5, 5, 6, 6, 6, 6, 5}}, 
 {64, {3, 5, 5, 5, 6, 5, 5, 5, 6, 6, 6, 6, 5, 6, 6, 5, 5}}, 
 {64, {3, 5, 5, 5, 5, 6, 6, 6, 5, 5, 6, 6, 5, 5, 6, 6, 5}}, 
 {64, {1, 6, 6, 6, 5, 5, 6, 5, 5, 4, 5, 6, 6, 6, 6, 6, 6}}, 
 {64, {1, 6, 6, 5, 4, 5, 5, 6, 5, 5, 6, 6, 6, 6, 6, 6, 6}}, 
 {63, {4, 4, 5, 5, 5, 6, 6, 5, 5, 5, 5, 5, 5, 4, 5, 5, 6, 5}}, 
 {63, {4, 4, 5, 5, 5, 5, 6, 5, 4, 5, 5, 5, 6, 6, 5, 5, 5, 5}}, 
 {63, {4, 4, 5, 5, 5, 4, 5, 5, 6, 5, 5, 5, 5, 5, 6, 6, 5, 5}}, 
 {63, {4, 4, 5, 5, 4, 5, 5, 5, 5, 5, 6, 6, 5, 5, 5, 5, 6, 5}}, 
 {63, {3, 5, 5, 5, 5, 5, 4, 5, 5, 5, 6, 6, 6, 5, 5, 5, 5, 5}}, 
 {63, {1, 6, 6, 5, 5, 4, 5, 5, 5, 5, 6, 5, 5, 5, 5, 5, 6, 6}}, 
 {63, {1, 6, 6, 5, 4, 5, 6, 5, 5, 5, 5, 5, 5, 5, 5, 5, 6, 6}}, 
 {63, {5, 5, 5, 5, 6, 5, 5, 6, 5, 5, 6, 5, 5, 6, 5, 5, 6}}, 
 {63, {4, 5, 5, 5, 6, 5, 5, 5, 4, 5, 6, 6, 6, 6, 6, 6, 5}}, 
 {63, {4, 4, 5, 5, 4, 5, 5, 6, 6, 6, 6, 6, 5, 6, 6, 6, 5}}, 
 {63, {3, 5, 5, 6, 6, 5, 5, 4, 5, 6, 6, 6, 6, 6, 6, 5, 5}}, 
 {63, {3, 5, 5, 6, 5, 5, 6, 6, 6, 6, 6, 5, 5, 5, 5, 6, 5}}, 
 {63, {3, 5, 5, 5, 6, 6, 6, 5, 5, 6, 5, 5, 5, 6, 6, 6, 5}}, 
 {63, {3, 5, 5, 5, 6, 6, 6, 5, 5, 5, 6, 6, 5, 5, 6, 6, 5}}, 
 {63, {3, 5, 5, 5, 6, 6, 5, 5, 5, 5, 6, 6, 6, 5, 6, 6, 5}}, 
 {63, {3, 5, 5, 5, 6, 5, 5, 5, 6, 6, 6, 5, 5, 6, 6, 6, 5}}, 
 {63, {3, 5, 5, 5, 5, 6, 6, 5, 5, 5, 6, 6, 6, 5, 6, 6, 5}}, 
 {62, {4, 5, 5, 5, 5, 6, 5, 5, 5, 4, 5, 5, 5, 6, 5, 5, 5, 5}}, 
 {62, {4, 5, 5, 5, 5, 5, 5, 4, 5, 6, 5, 5, 5, 5, 5, 5, 6, 5}}, 
 {62, {4, 5, 5, 5, 5, 4, 5, 5, 5, 5, 5, 5, 4, 5, 6, 6, 6, 5}}, 
 {62, {4, 5, 5, 5, 4, 5, 5, 6, 5, 5, 5, 4, 5, 5, 6, 6, 5, 5}}, 
 {62, {4, 4, 5, 5, 5, 6, 6, 5, 5, 5, 5, 4, 5, 5, 5, 6, 5, 5}}, 
 {62, {4, 4, 5, 5, 5, 5, 6, 6, 5, 5, 5, 5, 5, 4, 5, 6, 5, 5}}, 
 {62, {4, 4, 5, 5, 5, 5, 6, 6, 5, 5, 5, 4, 5, 5, 6, 5, 5, 5}}, 
 {62, {4, 4, 5, 5, 5, 5, 5, 5, 6, 6, 5, 5, 5, 5, 4, 5, 6, 5}}, 
 {62, {4, 4, 5, 5, 5, 4, 5, 6, 6, 5, 5, 5, 5, 6, 5, 5, 5, 5}}, 
 {62, {4, 4, 5, 5, 5, 4, 5, 5, 6, 6, 5, 5, 6, 5, 5, 5, 5, 5}}, 
 {62, {4, 4, 5, 5, 4, 5, 5, 5, 6, 6, 5, 5, 5, 5, 6, 5, 5, 5}}, 
 {62, {3, 5, 5, 5, 5, 5, 5, 5, 5, 5, 6, 5, 5, 6, 5, 5, 5, 5}}, 
 {62, {3, 5, 5, 5, 5, 5, 4, 5, 5, 5, 5, 6, 6, 6, 5, 5, 5, 5}}, 
 {62, {3, 5, 5, 5, 4, 5, 5, 5, 6, 6, 6, 5, 5, 5, 5, 5, 5, 5}}, 
 {62, {4, 5, 5, 6, 5, 5, 6, 5, 5, 5, 6, 5, 5, 6, 6, 6, 5}}, 
 {62, {4, 5, 5, 6, 5, 5, 5, 6, 6, 6, 5, 4, 5, 6, 6, 6, 5}}, 
 {62, {4, 5, 5, 5, 6, 6, 6, 5, 4, 5, 6, 5, 5, 6, 6, 6, 5}}, 
 {62, {4, 5, 5, 5, 6, 6, 5, 6, 6, 6, 6, 5, 4, 5, 6, 5, 5}}, 
 {62, {4, 5, 5, 5, 6, 6, 5, 4, 5, 6, 5, 5, 6, 6, 6, 6, 5}}, 
 {62, {4, 4, 5, 6, 6, 6, 5, 5, 5, 6, 5, 5, 5, 6, 6, 6, 5}}, 
 {62, {4, 4, 5, 6, 6, 6, 5, 4, 5, 5, 5, 6, 6, 6, 6, 6, 5}}, 
 {62, {3, 5, 5, 5, 6, 6, 6, 5, 5, 6, 6, 5, 5, 6, 6, 5, 5}}, 
 {62, {3, 5, 5, 5, 6, 6, 5, 5, 5, 6, 6, 5, 5, 6, 6, 6, 5}}, 
 {62, {1, 6, 6, 6, 5, 4, 5, 5, 6, 5, 5, 6, 6, 6, 6, 6, 6}}, 
 {62, {1, 6, 6, 5, 5, 6, 6, 6, 1, 6, 6, 6, 6, 6, 6, 6, 6}}, 
 {61, {4, 5, 5, 5, 5, 6, 5, 5, 5, 5, 4, 5, 5, 5, 6, 5, 5, 5}}, 
 {61, {4, 5, 5, 5, 4, 5, 5, 6, 6, 5, 5, 4, 5, 5, 5, 5, 6, 5}}, 
 {61, {4, 5, 5, 5, 4, 5, 5, 5, 6, 6, 5, 5, 4, 5, 5, 5, 6, 5}}, 
 {61, {4, 5, 5, 5, 4, 5, 5, 5, 6, 5, 4, 5, 5, 6, 6, 5, 5, 5}}, 
 {61, {4, 5, 5, 5, 4, 5, 5, 5, 6, 5, 4, 5, 5, 5, 6, 6, 5, 5}}, 
 {61, {4, 4, 5, 5, 5, 6, 6, 5, 5, 5, 5, 5, 4, 5, 5, 5, 6, 5}}, 
 {61, {4, 4, 5, 5, 5, 6, 5, 4, 5, 5, 5, 5, 5, 5, 5, 6, 6, 5}}, 
 {61, {4, 4, 5, 5, 5, 5, 4, 5, 6, 5, 5, 5, 6, 6, 5, 5, 5, 5}}, 
 {61, {4, 4, 5, 5, 5, 4, 5, 6, 6, 5, 5, 5, 5, 5, 5, 5, 6, 5}}, 
 {61, {4, 4, 5, 5, 5, 4, 5, 6, 5, 5, 6, 6, 5, 5, 5, 5, 5, 5}}, 
 {61, {4, 4, 5, 5, 4, 5, 6, 5, 5, 5, 5, 5, 5, 5, 5, 6, 6, 5}}, 
 {61, {4, 4, 5, 5, 4, 5, 5, 6, 6, 5, 5, 5, 6, 5, 5, 5, 5, 5}}, 
 {61, {4, 4, 5, 5, 4, 5, 5, 5, 6, 6, 5, 5, 5, 5, 5, 5, 6, 5}}, 
 {61, {3, 5, 5, 5, 5, 5, 5, 5, 5, 6, 5, 5, 5, 6, 5, 5, 5, 5}}, 
 {61, {4, 5, 5, 5, 6, 6, 6, 6, 5, 4, 5, 6, 6, 5, 5, 6, 5}}, 
 {61, {4, 5, 5, 5, 6, 6, 5, 5, 6, 6, 6, 5, 4, 5, 6, 6, 5}}, 
 {61, {4, 5, 5, 5, 6, 5, 4, 5, 6, 6, 6, 6, 5, 5, 6, 6, 5}}, 
 {61, {4, 5, 5, 5, 6, 5, 4, 5, 6, 6, 6, 5, 5, 6, 6, 6, 5}}, 
 {61, {4, 5, 5, 5, 6, 5, 4, 5, 6, 6, 5, 5, 6, 6, 6, 6, 5}}, 
 {61, {4, 5, 5, 4, 5, 5, 6, 6, 5, 5, 6, 6, 6, 5, 6, 6, 5}}, 
 {61, {4, 5, 4, 5, 5, 5, 5, 5, 5, 6, 6, 6, 6, 6, 6, 6, 5}}, 
 {61, {4, 4, 5, 5, 5, 6, 6, 6, 6, 6, 6, 5, 4, 5, 6, 6, 5}}, 
 {61, {4, 4, 5, 5, 5, 6, 5, 4, 5, 6, 6, 6, 6, 6, 6, 6, 5}}, 
 {61, {4, 4, 5, 4, 5, 5, 6, 6, 6, 6, 6, 5, 5, 6, 6, 6, 5}}, 
 {61, {3, 5, 5, 6, 6, 6, 6, 5, 5, 5, 4, 5, 6, 6, 6, 6, 5}}, 
 {61, {1, 6, 6, 5, 5, 5, 6, 5, 4, 5, 6, 6, 6, 6, 6, 6, 6}}, 
 {61, {1, 6, 6, 5, 4, 5, 5, 5, 6, 6, 6, 6, 6, 6, 5, 6, 6}}, 
 {60, {4, 4, 5, 5, 5, 6, 6, 5, 5, 4, 5, 5, 5, 5, 5, 6, 5, 5}}, 
 {60, {4, 4, 5, 5, 5, 5, 6, 6, 5, 5, 4, 5, 5, 5, 5, 6, 5, 5}}, 
 {60, {4, 4, 5, 5, 5, 5, 5, 6, 6, 5, 5, 5, 4, 5, 6, 5, 5, 5}}, 
 {60, {4, 4, 5, 5, 5, 5, 5, 4, 5, 6, 5, 5, 5, 5, 5, 6, 6, 5}}, 
 {60, {4, 4, 5, 5, 5, 4, 5, 5, 5, 4, 5, 5, 6, 6, 6, 6, 5, 5}}, 
 {60, {4, 4, 5, 5, 4, 5, 5, 6, 6, 5, 5, 5, 5, 5, 5, 6, 5, 5}}, 
 {60, {3, 5, 5, 5, 5, 5, 5, 5, 4, 5, 5, 5, 6, 6, 6, 5, 5, 5}}, 
 {60, {3, 5, 5, 5, 5, 5, 5, 4, 5, 5, 5, 5, 5, 5, 6, 6, 6, 5}}, 
 {60, {3, 5, 5, 5, 5, 5, 4, 5, 5, 5, 5, 5, 5, 6, 6, 6, 5, 5}}, 
 {60, {1, 6, 6, 5, 5, 5, 5, 5, 5, 4, 5, 6, 5, 5, 5, 5, 6, 6}}, 
 {60, {4, 5, 5, 6, 5, 5, 6, 6, 6, 5, 5, 4, 5, 6, 6, 6, 5}}, 
 {60, {4, 4, 5, 6, 5, 5, 6, 5, 5, 5, 6, 6, 6, 5, 6, 6, 5}}, 
 {60, {4, 4, 5, 6, 5, 5, 5, 6, 6, 5, 6, 6, 5, 5, 6, 6, 5}}, 
 {60, {4, 4, 5, 5, 5, 6, 6, 6, 6, 6, 6, 6, 5, 4, 5, 6, 5}}, 
 {60, {4, 4, 5, 5, 4, 5, 6, 6, 6, 6, 6, 6, 6, 5, 5, 6, 5}}, 
 {60, {4, 4, 5, 4, 5, 5, 6, 6, 6, 6, 5, 6, 6, 6, 6, 5, 5}}, 
 {60, {3, 5, 5, 6, 6, 6, 5, 5, 5, 4, 5, 6, 6, 6, 6, 6, 5}}, 
 {60, {3, 5, 5, 6, 6, 5, 4, 5, 5, 6, 6, 6, 6, 6, 6, 5, 5}}, 
 {60, {3, 5, 5, 5, 6, 6, 6, 6, 6, 5, 5, 6, 5, 5, 5, 6, 5}}, 
 {60, {3, 5, 5, 5, 6, 6, 5, 5, 5, 5, 6, 6, 5, 6, 6, 6, 5}}, 
 {60, {3, 5, 5, 5, 6, 5, 5, 6, 6, 5, 5, 6, 6, 6, 6, 5, 5}}, 
 {60, {1, 6, 6, 5, 6, 6, 6, 6, 6, 5, 5, 5, 4, 5, 6, 6, 6}}, 
 {59, {4, 5, 5, 5, 5, 4, 5, 5, 6, 6, 5, 5, 4, 5, 5, 5, 6, 5}}, 
 {59, {4, 5, 5, 5, 4, 5, 5, 6, 6, 5, 5, 5, 4, 5, 5, 5, 6, 5}}, 
 {59, {4, 4, 5, 5, 6, 5, 4, 5, 5, 5, 5, 5, 5, 5, 6, 6, 5, 5}}, 
 {59, {4, 4, 5, 5, 5, 6, 5, 5, 5, 4, 5, 5, 5, 6, 6, 5, 5, 5}}, 
 {59, {4, 4, 5, 5, 5, 5, 6, 5, 4, 5, 5, 5, 5, 5, 6, 6, 5, 5}}, 
 {59, {4, 4, 5, 5, 5, 5, 5, 4, 5, 5, 6, 6, 5, 5, 5, 6, 5, 5}}, 
 {59, {4, 4, 5, 5, 5, 5, 4, 5, 5, 5, 5, 6, 6, 5, 5, 5, 6, 5}}, 
 {59, {4, 4, 5, 5, 5, 5, 4, 4, 5, 5, 5, 5, 5, 6, 6, 6, 6, 5}}, 
 {59, {4, 4, 5, 5, 5, 4, 5, 6, 5, 5, 5, 5, 5, 5, 5, 6, 6, 5}}, 
 {59, {4, 4, 5, 5, 5, 4, 5, 5, 5, 6, 5, 5, 5, 5, 5, 6, 6, 5}}, 
 {59, {4, 4, 5, 5, 4, 5, 6, 5, 5, 5, 5, 5, 5, 6, 6, 5, 5, 5}}, 
 {59, {4, 4, 5, 5, 4, 5, 5, 6, 6, 6, 6, 5, 5, 4, 5, 5, 5, 5}}, 
 {59, {4, 4, 5, 5, 4, 5, 5, 6, 5, 5, 5, 5, 6, 6, 5, 5, 5, 5}}, 
 {59, {4, 4, 5, 5, 4, 5, 5, 5, 5, 6, 5, 5, 5, 5, 6, 6, 5, 5}}, 
 {59, {3, 5, 6, 5, 5, 5, 5, 5, 4, 5, 5, 5, 5, 5, 5, 6, 6, 5}}, 
 {59, {3, 5, 5, 5, 4, 5, 5, 5, 5, 6, 6, 5, 5, 5, 5, 5, 6, 5}}, 
 {59, {1, 6, 6, 5, 5, 5, 5, 5, 4, 5, 6, 5, 5, 5, 5, 5, 6, 6}}, 
 {59, {4, 5, 5, 6, 6, 6, 5, 5, 4, 5, 6, 6, 5, 5, 6, 6, 5}}, 
 {59, {4, 5, 5, 5, 6, 6, 6, 6, 5, 6, 6, 5, 4, 5, 6, 5, 5}}, 
 {59, {4, 5, 5, 5, 6, 6, 5, 4, 5, 6, 6, 6, 6, 5, 5, 6, 5}}, 
 {59, {4, 5, 5, 5, 6, 5, 5, 6, 6, 6, 6, 5, 4, 5, 6, 6, 5}}, 
 {59, {4, 5, 5, 5, 4, 5, 6, 6, 5, 5, 6, 6, 6, 5, 6, 6, 5}}, 
 {59, {4, 4, 5, 6, 6, 6, 6, 5, 4, 5, 5, 5, 6, 6, 6, 6, 5}}, 
 {59, {4, 4, 5, 6, 5, 5, 6, 6, 5, 5, 5, 6, 6, 5, 6, 6, 5}}, 
 {59, {4, 4, 5, 6, 5, 4, 5, 5, 5, 6, 6, 6, 6, 6, 6, 6, 5}}, 
 {59, {4, 4, 5, 5, 4, 5, 6, 6, 5, 5, 6, 6, 6, 6, 6, 6, 5}}, 
 {59, {4, 4, 5, 4, 5, 5, 6, 6, 6, 6, 5, 5, 6, 6, 6, 6, 5}}, 
 {59, {3, 5, 6, 6, 6, 6, 5, 5, 4, 5, 5, 5, 6, 6, 6, 6, 5}}, 
 {59, {3, 5, 5, 5, 6, 6, 6, 5, 6, 6, 5, 5, 5, 5, 6, 6, 5}}, 
 {59, {3, 5, 5, 5, 6, 6, 5, 5, 5, 6, 6, 5, 6, 6, 6, 5, 5}}, 
 {59, {3, 5, 5, 5, 5, 6, 6, 5, 6, 6, 6, 5, 5, 6, 6, 5, 5}}, 
 {59, {3, 5, 5, 5, 5, 6, 6, 5, 6, 6, 6, 5, 5, 5, 6, 6, 5}}, 
 {58, {4, 5, 5, 4, 5, 5, 6, 5, 5, 5, 5, 5, 5, 5, 5, 6, 5, 5}}, 
 {58, {4, 4, 5, 6, 5, 5, 5, 5, 5, 5, 5, 4, 5, 5, 5, 6, 6, 5}}, 
 {58, {4, 4, 5, 5, 5, 6, 6, 5, 5, 5, 5, 5, 4, 5, 5, 6, 5, 5}}, 
 {58, {4, 4, 5, 5, 5, 5, 5, 5, 6, 5, 4, 5, 5, 5, 5, 6, 6, 5}}, 
 {58, {4, 4, 5, 5, 5, 5, 5, 5, 5, 4, 5, 6, 5, 5, 6, 6, 5, 5}}, 
 {58, {4, 4, 5, 5, 5, 5, 4, 5, 5, 6, 5, 5, 6, 6, 5, 5, 5, 5}}, 
 {58, {4, 4, 5, 5, 4, 5, 5, 6, 5, 5, 5, 5, 5, 6, 6, 5, 5, 5}}, 
 {58, {4, 4, 5, 5, 4, 5, 5, 5, 5, 5, 5, 5, 6, 6, 5, 5, 6, 5}}, 
 {58, {4, 4, 5, 5, 4, 4, 5, 5, 5, 5, 5, 5, 5, 6, 6, 6, 6, 5}}, 
 {58, {3, 5, 6, 5, 5, 5, 5, 5, 5, 4, 5, 5, 5, 5, 5, 6, 6, 5}}, 
 {58, {3, 5, 5, 4, 5, 5, 5, 5, 5, 6, 6, 5, 5, 5, 5, 5, 6, 5}}, 
 {58, {1, 6, 6, 5, 5, 5, 5, 5, 5, 4, 4, 5, 5, 5, 6, 6, 6, 6}}, 
 {58, {4, 5, 6, 5, 4, 5, 6, 6, 5, 5, 5, 5, 6, 6, 6, 6, 5}}, 
 {58, {4, 5, 5, 6, 5, 5, 5, 6, 6, 6, 6, 5, 4, 5, 6, 6, 5}}, 
 {58, {4, 5, 5, 6, 5, 5, 4, 5, 5, 6, 6, 5, 6, 6, 6, 6, 5}}, 
 {58, {4, 5, 5, 5, 6, 6, 5, 4, 5, 6, 6, 6, 5, 5, 6, 6, 5}}, 
 {58, {4, 5, 5, 5, 6, 5, 5, 6, 6, 5, 4, 5, 6, 6, 6, 6, 5}}, 
 {58, {4, 5, 5, 5, 6, 5, 5, 4, 5, 6, 6, 5, 6, 6, 6, 6, 5}}, 
 {58, {4, 5, 5, 4, 5, 5, 6, 6, 5, 6, 6, 6, 6, 5, 5, 6, 5}}, 
 {58, {4, 5, 5, 4, 5, 5, 6, 6, 5, 6, 6, 6, 5, 5, 6, 6, 5}}, 
 {58, {4, 5, 4, 5, 5, 5, 6, 6, 6, 6, 6, 5, 5, 6, 6, 5, 5}}, 
 {58, {4, 4, 5, 6, 5, 5, 6, 6, 5, 5, 6, 6, 6, 5, 5, 6, 5}}, 
 {58, {4, 4, 5, 5, 6, 6, 6, 6, 5, 4, 5, 6, 6, 6, 6, 5, 5}}, 
 {58, {4, 4, 5, 5, 4, 5, 6, 6, 6, 5, 6, 6, 6, 6, 6, 5, 5}}, 
 {58, {4, 4, 5, 4, 5, 5, 6, 6, 6, 5, 6, 6, 6, 6, 6, 5, 5}}, 
 {58, {3, 5, 6, 5, 5, 6, 6, 5, 5, 5, 6, 6, 6, 5, 5, 6, 5}}, 
 {58, {3, 5, 5, 5, 5, 6, 6, 6, 5, 6, 6, 5, 5, 6, 6, 5, 5}}, 
 {58, {3, 5, 5, 5, 5, 5, 6, 6, 5, 5, 6, 6, 6, 5, 6, 6, 5}}, 
 {58, {3, 5, 5, 5, 5, 5, 5, 6, 6, 5, 6, 6, 5, 6, 6, 6, 5}}, 
 {58, {3, 5, 5, 4, 5, 6, 6, 6, 6, 6, 6, 6, 5, 5, 5, 6, 5}}, 
 {58, {1, 6, 6, 5, 6, 6, 6, 5, 4, 5, 5, 5, 6, 6, 6, 6, 6}}, 
 {58, {1, 6, 6, 5, 5, 4, 5, 6, 6, 6, 6, 6, 6, 5, 5, 6, 6}}, 
 {57, {4, 5, 5, 5, 5, 5, 5, 5, 5, 5, 4, 5, 6, 5, 5, 5, 6, 5}}, 
 {57, {4, 5, 5, 5, 4, 5, 5, 6, 5, 5, 5, 5, 5, 5, 5, 6, 5, 5}}, 
 {57, {4, 5, 5, 4, 5, 5, 5, 4, 5, 5, 5, 5, 5, 5, 6, 6, 6, 5}}, 
 {57, {4, 4, 5, 5, 6, 5, 5, 5, 5, 5, 5, 5, 4, 5, 5, 6, 6, 5}}, 
 {57, {4, 4, 5, 5, 5, 6, 5, 4, 5, 5, 5, 5, 5, 6, 6, 5, 5, 5}}, 
 {57, {4, 4, 5, 5, 5, 5, 6, 5, 4, 5, 5, 5, 5, 6, 6, 5, 5, 5}}, 
 {57, {4, 4, 5, 5, 5, 5, 4, 5, 5, 6, 5, 5, 5, 5, 5, 6, 6, 5}}, 
 {57, {4, 4, 5, 5, 4, 5, 5, 5, 6, 5, 5, 5, 6, 6, 5, 5, 5, 5}}, 
 {57, {4, 4, 5, 5, 4, 5, 5, 5, 5, 5, 6, 5, 5, 6, 6, 5, 5, 5}}, 
 {57, {3, 5, 5, 5, 5, 5, 5, 5, 5, 6, 6, 5, 5, 4, 5, 5, 6, 5}}, 
 {57, {3, 5, 5, 5, 5, 5, 5, 5, 5, 5, 6, 5, 5, 5, 6, 5, 5, 5}}, 
 {57, {3, 5, 5, 5, 5, 5, 5, 5, 5, 5, 4, 5, 6, 6, 6, 5, 5, 5}}, 
 {57, {3, 5, 5, 5, 5, 5, 5, 5, 4, 5, 6, 6, 6, 5, 5, 5, 5, 5}}, 
 {57, {3, 5, 5, 5, 5, 4, 5, 5, 5, 5, 5, 6, 6, 6, 5, 5, 5, 5}}, 
 {57, {3, 5, 5, 4, 5, 5, 5, 5, 5, 5, 5, 5, 6, 6, 6, 5, 5, 5}}, 
 {57, {1, 6, 6, 5, 5, 4, 4, 5, 5, 5, 5, 5, 5, 5, 6, 6, 6, 6}}, 
 {57, {4, 5, 5, 5, 6, 6, 6, 5, 5, 6, 6, 6, 5, 4, 5, 6, 5}}, 
 {57, {4, 5, 5, 4, 5, 5, 6, 6, 5, 6, 6, 5, 5, 6, 6, 6, 5}}, 
 {57, {4, 4, 5, 6, 5, 5, 5, 6, 6, 5, 6, 6, 6, 5, 5, 6, 5}}, 
 {57, {4, 4, 5, 5, 5, 6, 6, 6, 6, 6, 5, 4, 5, 6, 6, 6, 5}}, 
 {57, {4, 4, 5, 4, 5, 5, 6, 6, 6, 6, 6, 6, 5, 5, 6, 6, 5}}, 
 {57, {3, 5, 6, 5, 5, 6, 6, 5, 5, 6, 6, 5, 5, 5, 6, 6, 5}}, 
 {57, {3, 5, 5, 5, 6, 5, 5, 6, 6, 6, 6, 5, 5, 6, 6, 5, 5}}, 
 {57, {3, 5, 5, 5, 5, 5, 6, 6, 5, 5, 6, 6, 5, 6, 6, 6, 5}}, 
 {57, {3, 5, 5, 4, 5, 5, 6, 6, 6, 6, 6, 5, 5, 6, 6, 6, 5}}, 
 {57, {1, 6, 6, 5, 6, 6, 6, 6, 6, 5, 4, 5, 5, 5, 6, 6, 6}}, 
 {56, {4, 5, 5, 5, 5, 5, 6, 5, 5, 4, 5, 5, 5, 5, 5, 6, 5, 5}}, 
 {56, {4, 5, 5, 5, 5, 4, 5, 5, 5, 6, 6, 5, 5, 4, 5, 5, 6, 5}}, 
 {56, {4, 5, 5, 4, 5, 5, 5, 5, 5, 6, 5, 4, 5, 5, 6, 6, 5, 5}}, 
 {56, {4, 5, 5, 4, 5, 5, 5, 5, 4, 5, 5, 5, 5, 5, 6, 6, 6, 5}}, 
 {56, {4, 5, 4, 5, 5, 5, 5, 5, 5, 5, 5, 5, 6, 6, 5, 5, 5, 5}}, 
 {56, {4, 4, 5, 5, 6, 6, 5, 5, 5, 5, 5, 4, 5, 5, 5, 5, 6, 5}}, 
 {56, {4, 4, 5, 5, 5, 6, 6, 5, 5, 5, 5, 5, 5, 5, 4, 5, 6, 5}}, 
 {56, {4, 4, 5, 5, 5, 5, 4, 5, 5, 6, 5, 5, 5, 5, 6, 6, 5, 5}}, 
 {56, {4, 4, 5, 5, 5, 5, 4, 5, 5, 5, 5, 5, 6, 6, 5, 5, 6, 5}}, 
 {56, {4, 4, 5, 5, 4, 5, 6, 5, 5, 5, 5, 5, 5, 5, 6, 6, 5, 5}}, 
 {56, {4, 4, 5, 5, 4, 5, 5, 6, 6, 5, 5, 6, 5, 5, 5, 5, 5, 5}}, 
 {56, {4, 4, 5, 5, 4, 5, 5, 6, 5, 5, 5, 5, 5, 5, 5, 6, 6, 5}}, 
 {56, {1, 6, 6, 5, 5, 4, 5, 5, 5, 5, 5, 6, 5, 5, 5, 5, 6, 6}}, 
 {56, {4, 5, 5, 6, 5, 5, 5, 6, 6, 6, 6, 6, 5, 4, 5, 6, 5}}, 
 {56, {4, 5, 5, 5, 6, 6, 6, 6, 6, 5, 5, 6, 5, 4, 5, 6, 5}}, 
 {56, {4, 5, 5, 5, 6, 6, 6, 6, 5, 5, 6, 5, 4, 5, 6, 6, 5}}, 
 {56, {4, 4, 5, 6, 5, 5, 6, 5, 5, 6, 6, 6, 6, 5, 5, 6, 5}}, 
 {56, {4, 4, 5, 5, 5, 4, 5, 6, 6, 6, 6, 5, 6, 6, 6, 6, 5}}, 
 {56, {4, 4, 5, 5, 4, 5, 6, 6, 6, 6, 6, 5, 5, 6, 6, 6, 5}}, 
 {56, {4, 4, 5, 5, 4, 5, 6, 6, 6, 6, 5, 5, 6, 6, 6, 6, 5}}, 
 {56, {3, 5, 5, 6, 5, 5, 6, 6, 6, 5, 5, 5, 5, 6, 6, 6, 5}}, 
 {56, {3, 5, 5, 4, 5, 5, 6, 6, 6, 6, 6, 6, 6, 5, 5, 6, 5}}, 
 {55, {4, 4, 5, 5, 6, 6, 5, 4, 5, 5, 5, 5, 5, 5, 5, 5, 6, 5}}, 
 {55, {4, 4, 5, 5, 5, 6, 5, 5, 4, 5, 5, 5, 5, 6, 6, 5, 5, 5}}, 
 {55, {4, 4, 5, 5, 5, 6, 5, 5, 4, 5, 5, 5, 5, 5, 5, 6, 6, 5}}, 
 {55, {4, 4, 5, 5, 5, 6, 5, 4, 5, 5, 5, 5, 5, 5, 6, 6, 5, 5}}, 
 {55, {4, 4, 5, 5, 5, 5, 6, 6, 5, 5, 5, 5, 5, 5, 4, 5, 6, 5}}, 
 {55, {4, 4, 5, 5, 5, 5, 5, 6, 5, 4, 5, 5, 5, 5, 5, 6, 6, 5}}, 
 {55, {4, 4, 5, 5, 5, 5, 5, 5, 6, 5, 4, 5, 6, 6, 5, 5, 5, 5}}, 
 {55, {4, 4, 5, 5, 5, 5, 5, 4, 5, 6, 5, 5, 6, 6, 5, 5, 5, 5}}, 
 {55, {4, 4, 5, 5, 5, 5, 4, 5, 6, 5, 5, 5, 5, 6, 6, 5, 5, 5}}, 
 {55, {4, 4, 5, 5, 5, 5, 4, 5, 5, 5, 6, 5, 5, 5, 5, 6, 6, 5}}, 
 {55, {4, 4, 5, 5, 5, 4, 5, 5, 5, 6, 5, 5, 5, 6, 6, 5, 5, 5}}, 
 {55, {4, 4, 5, 5, 4, 5, 6, 5, 5, 5, 5, 6, 6, 5, 5, 5, 5, 5}}, 
 {55, {4, 4, 5, 5, 4, 5, 5, 6, 5, 5, 5, 5, 5, 5, 6, 6, 5, 5}}, 
 {55, {4, 4, 5, 5, 4, 5, 5, 5, 5, 5, 5, 6, 6, 5, 5, 5, 6, 5}}, 
 {55, {4, 4, 5, 5, 4, 5, 5, 5, 5, 4, 5, 5, 6, 6, 6, 6, 5, 5}}, 
 {55, {3, 5, 5, 5, 5, 5, 5, 5, 6, 6, 5, 5, 4, 5, 5, 5, 6, 5}}, 
 {55, {1, 6, 6, 5, 4, 5, 5, 6, 5, 5, 5, 5, 5, 5, 5, 5, 6, 6}}, 
 {55, {1, 6, 6, 5, 4, 5, 5, 5, 5, 4, 5, 5, 5, 5, 6, 6, 6, 6}}, 
 {55, {4, 5, 6, 5, 4, 5, 6, 5, 5, 5, 5, 6, 6, 6, 6, 6, 5}}, 
 {55, {4, 4, 5, 5, 4, 5, 6, 6, 6, 6, 5, 6, 6, 6, 6, 5, 5}}, 
 {55, {4, 4, 5, 4, 5, 6, 6, 6, 6, 5, 5, 5, 6, 6, 6, 6, 5}}, 
 {55, {3, 5, 6, 5, 5, 4, 5, 5, 6, 6, 6, 6, 6, 5, 6, 6, 5}}, 
 {55, {3, 5, 5, 5, 6, 6, 5, 5, 5, 6, 6, 6, 5, 6, 6, 5, 5}}, 
 {55, {3, 5, 5, 5, 6, 5, 5, 6, 6, 6, 6, 5, 5, 5, 6, 6, 5}}, 
 {55, {3, 5, 5, 5, 5, 6, 6, 5, 5, 6, 6, 6, 5, 5, 6, 6, 5}}, 
 {55, {3, 5, 5, 5, 5, 5, 6, 6, 5, 6, 6, 6, 5, 5, 6, 6, 5}}, 
 {54, {4, 5, 5, 5, 5, 5, 5, 4, 5, 5, 5, 6, 5, 5, 6, 5, 5, 5}}, 
 {54, {4, 5, 5, 5, 5, 5, 4, 5, 5, 5, 6, 5, 5, 5, 6, 5, 5, 5}}, 
 {54, {4, 5, 5, 5, 4, 5, 5, 5, 6, 5, 5, 4, 5, 5, 6, 6, 5, 5}}, 
 {54, {4, 5, 5, 5, 4, 5, 5, 5, 4, 5, 5, 5, 6, 6, 6, 5, 5, 5}}, 
 {54, {4, 5, 5, 4, 5, 5, 5, 5, 5, 4, 5, 6, 6, 6, 5, 5, 5, 5}}, 
 {54, {4, 5, 5, 4, 5, 5, 5, 4, 5, 6, 5, 5, 5, 5, 6, 6, 5, 5}}, 
 {54, {4, 4, 5, 5, 5, 6, 6, 5, 5, 5, 4, 5, 5, 5, 5, 5, 6, 5}}, 
 {54, {4, 4, 5, 5, 5, 6, 5, 5, 4, 5, 5, 5, 5, 5, 6, 6, 5, 5}}, 
 {54, {4, 4, 5, 5, 5, 5, 6, 6, 5, 4, 5, 5, 5, 5, 5, 5, 6, 5}}, 
 {54, {4, 4, 5, 5, 5, 5, 6, 5, 5, 4, 5, 5, 6, 6, 5, 5, 5, 5}}, 
 {54, {4, 4, 5, 5, 5, 5, 5, 5, 6, 6, 5, 4, 5, 6, 5, 5, 5, 5}}, 
 {54, {4, 4, 5, 5, 5, 4, 5, 5, 5, 6, 6, 5, 5, 5, 5, 6, 5, 5}}, 
 {54, {4, 4, 5, 5, 5, 4, 5, 5, 5, 6, 5, 5, 6, 6, 5, 5, 5, 5}}, 
 {54, {4, 4, 5, 5, 5, 4, 5, 5, 5, 5, 6, 5, 5, 5, 5, 6, 6, 5}}, 
 {54, {4, 4, 5, 5, 5, 4, 5, 5, 5, 5, 5, 5, 6, 6, 5, 5, 6, 5}}, 
 {54, {4, 4, 5, 5, 4, 5, 6, 5, 5, 6, 6, 5, 5, 5, 5, 5, 5, 5}}, 
 {54, {4, 4, 5, 5, 4, 5, 5, 5, 5, 5, 6, 6, 5, 5, 6, 5, 5, 5}}, 
 {54, {4, 4, 5, 5, 4, 5, 5, 5, 5, 5, 6, 5, 5, 5, 5, 6, 6, 5}}, 
 {54, {4, 4, 5, 4, 5, 5, 5, 5, 5, 5, 6, 6, 5, 5, 5, 5, 6, 5}}, 
 {54, {3, 5, 5, 6, 6, 5, 5, 5, 5, 5, 5, 5, 5, 4, 5, 5, 6, 5}}, 
 {54, {3, 5, 5, 5, 5, 6, 6, 5, 5, 4, 5, 5, 5, 5, 5, 5, 6, 5}}, 
 {54, {3, 5, 5, 5, 5, 5, 6, 6, 5, 5, 5, 5, 4, 5, 5, 5, 6, 5}}, 
 {54, {3, 5, 5, 5, 5, 5, 4, 5, 5, 5, 5, 5, 6, 6, 6, 5, 5, 5}}, 
 {54, {3, 5, 5, 5, 5, 4, 5, 5, 6, 6, 5, 5, 5, 5, 5, 5, 6, 5}}, 
 {54, {3, 5, 5, 5, 4, 5, 5, 5, 5, 5, 6, 6, 5, 5, 5, 5, 6, 5}}, 
 {54, {4, 5, 5, 5, 6, 6, 6, 5, 4, 5, 6, 6, 5, 5, 6, 6, 5}}, 
 {54, {4, 5, 5, 5, 5, 6, 5, 4, 5, 6, 6, 6, 6, 5, 6, 6, 5}}, 
 {54, {4, 5, 5, 5, 4, 5, 6, 6, 5, 6, 6, 5, 5, 6, 6, 6, 5}}, 
 {54, {4, 5, 4, 5, 5, 5, 6, 6, 6, 6, 5, 5, 6, 6, 6, 5, 5}}, 
 {54, {4, 5, 4, 5, 5, 5, 6, 6, 6, 5, 5, 6, 6, 6, 6, 5, 5}}, 
 {54, {4, 5, 4, 5, 5, 5, 6, 6, 5, 5, 6, 6, 6, 6, 6, 5, 5}}, 
 {54, {4, 4, 5, 6, 6, 6, 5, 5, 5, 4, 5, 6, 6, 6, 6, 6, 5}}, 
 {54, {4, 4, 5, 5, 6, 6, 5, 6, 6, 5, 5, 6, 6, 5, 5, 6, 5}}, 
 {54, {3, 5, 6, 5, 5, 6, 6, 5, 5, 6, 5, 5, 5, 6, 6, 6, 5}}, 
 {54, {3, 5, 5, 6, 5, 5, 6, 6, 5, 5, 5, 6, 6, 6, 6, 5, 5}}, 
 {54, {3, 5, 5, 6, 5, 5, 5, 6, 6, 6, 5, 5, 5, 6, 6, 6, 5}}, 
 {54, {3, 5, 5, 5, 6, 6, 6, 5, 5, 5, 6, 5, 5, 6, 6, 6, 5}}, 
 {54, {3, 5, 5, 5, 5, 6, 6, 6, 5, 5, 6, 5, 5, 6, 6, 6, 5}}, 
 {54, {3, 5, 5, 5, 5, 6, 6, 5, 5, 6, 6, 6, 5, 6, 6, 5, 5}}, 
 {54, {3, 5, 5, 5, 5, 5, 6, 6, 6, 5, 6, 6, 5, 5, 6, 6, 5}}, 
 {54, {3, 5, 5, 5, 5, 5, 5, 6, 6, 6, 5, 6, 6, 5, 6, 6, 5}}, 
 {54, {3, 5, 5, 4, 5, 5, 6, 6, 6, 6, 6, 6, 5, 5, 6, 6, 5}}, 
 {53, {4, 5, 5, 5, 5, 6, 5, 5, 5, 4, 5, 5, 5, 5, 6, 5, 5, 5}}, 
 {53, {4, 5, 5, 5, 5, 5, 4, 5, 5, 6, 5, 5, 5, 5, 5, 6, 5, 5}}, 
 {53, {4, 5, 5, 4, 5, 5, 5, 5, 5, 5, 4, 5, 6, 6, 6, 5, 5, 5}}, 
 {53, {4, 5, 5, 4, 5, 5, 5, 5, 5, 5, 4, 5, 5, 5, 6, 6, 6, 5}}, 
 {53, {4, 4, 5, 5, 6, 5, 5, 4, 5, 5, 5, 5, 5, 5, 6, 6, 5, 5}}, 
 {53, {4, 4, 5, 5, 5, 5, 6, 5, 5, 5, 4, 5, 5, 5, 5, 6, 6, 5}}, 
 {53, {4, 4, 5, 5, 5, 5, 5, 6, 6, 5, 4, 5, 5, 5, 5, 5, 6, 5}}, 
 {53, {4, 4, 5, 5, 5, 5, 5, 6, 5, 4, 5, 6, 6, 5, 5, 5, 5, 5}}, 
 {53, {4, 4, 5, 5, 5, 5, 5, 4, 5, 6, 6, 5, 5, 5, 5, 5, 6, 5}}, 
 {53, {4, 4, 5, 5, 5, 5, 5, 4, 5, 5, 6, 5, 5, 5, 6, 6, 5, 5}}, 
 {53, {4, 4, 5, 5, 5, 5, 4, 5, 5, 6, 6, 5, 5, 6, 5, 5, 5, 5}}, 
 {53, {4, 4, 5, 5, 5, 5, 4, 5, 5, 5, 6, 5, 5, 6, 6, 5, 5, 5}}, 
 {53, {3, 5, 5, 5, 5, 5, 5, 4, 5, 5, 6, 6, 5, 5, 5, 5, 6, 5}}, 
 {53, {3, 5, 5, 5, 5, 5, 5, 4, 5, 5, 5, 6, 6, 6, 5, 5, 5, 5}}, 
 {53, {3, 5, 5, 5, 5, 5, 5, 4, 5, 5, 5, 5, 6, 6, 6, 5, 5, 5}}, 
 {53, {3, 5, 5, 5, 5, 5, 4, 5, 5, 6, 6, 5, 5, 5, 5, 5, 6, 5}}, 
 {53, {4, 5, 5, 6, 6, 5, 5, 6, 6, 5, 5, 4, 5, 6, 6, 6, 5}}, 
 {53, {4, 5, 5, 5, 6, 6, 6, 5, 5, 6, 6, 5, 4, 5, 6, 6, 5}}, 
 {53, {4, 5, 5, 4, 5, 5, 6, 6, 5, 5, 6, 6, 5, 6, 6, 6, 5}}, 
 {53, {4, 4, 5, 6, 5, 5, 5, 6, 6, 5, 5, 6, 6, 5, 6, 6, 5}}, 
 {53, {4, 4, 5, 6, 5, 5, 5, 6, 5, 5, 6, 6, 6, 5, 6, 6, 5}}, 
 {53, {4, 4, 5, 6, 5, 5, 5, 4, 5, 6, 6, 6, 6, 6, 6, 6, 5}}, 
 {53, {4, 4, 5, 5, 4, 5, 6, 6, 6, 6, 6, 5, 6, 6, 6, 5, 5}}, 
 {53, {3, 5, 5, 6, 5, 5, 5, 5, 6, 6, 6, 6, 6, 5, 5, 6, 5}}, 
 {53, {3, 5, 5, 5, 6, 6, 6, 5, 5, 6, 5, 5, 6, 6, 6, 5, 5}}, 
 {53, {3, 5, 5, 5, 6, 5, 5, 6, 6, 6, 5, 5, 6, 6, 6, 5, 5}}, 
 {53, {3, 5, 5, 5, 6, 5, 5, 5, 6, 6, 5, 6, 6, 6, 6, 5, 5}}, 
 {53, {1, 6, 6, 5, 5, 6, 5, 5, 4, 5, 6, 6, 6, 6, 6, 6, 6}}, 
 {53, {1, 6, 6, 5, 5, 5, 4, 5, 6, 6, 6, 6, 5, 6, 6, 6, 6}}, 
 {52, {4, 5, 5, 5, 5, 5, 6, 5, 5, 5, 5, 5, 4, 5, 5, 6, 5, 5}}, 
 {52, {4, 5, 5, 5, 5, 5, 5, 4, 5, 5, 6, 5, 5, 5, 5, 6, 5, 5}}, 
 {52, {4, 5, 5, 5, 5, 5, 4, 5, 5, 6, 6, 5, 4, 5, 5, 5, 6, 5}}, 
 {52, {4, 5, 5, 4, 5, 5, 5, 4, 5, 6, 6, 6, 5, 5, 5, 5, 5, 5}}, 
 {52, {4, 5, 4, 5, 5, 5, 5, 5, 5, 5, 5, 6, 6, 5, 5, 5, 5, 5}}, 
 {52, {4, 4, 5, 5, 6, 6, 5, 5, 5, 5, 5, 5, 5, 5, 4, 5, 6, 5}}, 
 {52, {4, 4, 5, 5, 5, 5, 5, 5, 5, 4, 5, 6, 5, 5, 5, 6, 6, 5}}, 
 {52, {4, 4, 5, 5, 5, 5, 5, 5, 5, 4, 4, 5, 5, 6, 6, 6, 6, 5}}, 
 {52, {4, 4, 5, 5, 5, 5, 5, 5, 4, 5, 5, 6, 6, 5, 5, 6, 5, 5}}, 
 {52, {4, 4, 5, 5, 5, 5, 5, 5, 4, 4, 5, 5, 5, 6, 6, 6, 6, 5}}, 
 {52, {4, 4, 5, 5, 5, 5, 5, 4, 5, 5, 6, 6, 5, 5, 6, 5, 5, 5}}, 
 {52, {4, 4, 5, 5, 5, 4, 5, 5, 5, 5, 6, 6, 5, 5, 6, 5, 5, 5}}, 
 {52, {4, 4, 5, 5, 5, 4, 5, 5, 5, 5, 5, 6, 6, 5, 5, 6, 5, 5}}, 
 {52, {4, 4, 5, 5, 5, 4, 4, 5, 5, 5, 5, 5, 5, 6, 6, 6, 6, 5}}, 
 {52, {4, 4, 5, 5, 4, 5, 5, 5, 6, 6, 6, 6, 5, 5, 4, 5, 5, 5}}, 
 {52, {4, 4, 5, 5, 4, 5, 5, 5, 6, 5, 5, 5, 5, 5, 5, 6, 6, 5}}, 
 {52, {4, 4, 5, 5, 4, 5, 5, 5, 5, 5, 5, 6, 5, 5, 6, 6, 5, 5}}, 
 {52, {4, 4, 5, 4, 5, 5, 6, 6, 5, 5, 5, 5, 5, 5, 5, 5, 6, 5}}, 
 {52, {4, 4, 5, 4, 5, 5, 5, 5, 5, 5, 5, 6, 6, 5, 5, 5, 6, 5}}, 
 {52, {3, 5, 6, 5, 5, 5, 5, 4, 5, 5, 5, 5, 5, 5, 5, 6, 6, 5}}, 
 {52, {3, 5, 5, 5, 5, 5, 6, 6, 5, 5, 4, 5, 5, 5, 5, 5, 6, 5}}, 
 {52, {3, 5, 5, 5, 5, 5, 6, 5, 5, 5, 5, 5, 5, 6, 5, 5, 5, 5}}, 
 {52, {4, 5, 5, 5, 6, 6, 6, 6, 5, 5, 6, 6, 5, 4, 5, 6, 5}}, 
 {52, {4, 5, 5, 5, 6, 6, 5, 5, 6, 6, 6, 6, 5, 4, 5, 6, 5}}, 
 {52, {4, 5, 5, 5, 4, 5, 6, 5, 5, 6, 6, 6, 6, 5, 6, 6, 5}}, 
 {52, {4, 4, 5, 6, 6, 5, 4, 5, 5, 5, 6, 6, 6, 6, 6, 6, 5}}, 
 {52, {4, 4, 5, 5, 4, 5, 6, 6, 6, 6, 6, 6, 5, 5, 6, 6, 5}}, 
 {52, {4, 4, 5, 5, 4, 5, 6, 6, 6, 5, 5, 6, 6, 6, 6, 6, 5}}, 
 {52, {4, 4, 5, 4, 5, 5, 6, 6, 6, 6, 6, 6, 6, 5, 5, 6, 5}}, 
 {52, {3, 5, 6, 5, 5, 6, 5, 5, 6, 6, 5, 5, 5, 6, 6, 6, 5}}, 
 {52, {3, 5, 5, 6, 5, 5, 6, 6, 5, 5, 5, 5, 6, 6, 6, 6, 5}}, 
 {52, {3, 5, 5, 6, 5, 5, 5, 6, 6, 6, 6, 6, 5, 5, 5, 6, 5}}, 
 {52, {3, 5, 5, 5, 5, 6, 6, 5, 5, 6, 6, 5, 6, 6, 6, 5, 5}}, 
 {52, {1, 6, 6, 6, 5, 5, 6, 6, 6, 1, 6, 6, 6, 6, 6, 6, 6}}, 
 {51, {4, 5, 5, 5, 5, 4, 5, 5, 6, 5, 5, 5, 5, 5, 5, 6, 5, 5}}, 
 {51, {4, 5, 5, 5, 5, 4, 5, 5, 5, 5, 6, 5, 5, 6, 5, 5, 5, 5}}, 
 {51, {4, 5, 5, 5, 4, 5, 5, 5, 5, 6, 6, 5, 5, 4, 5, 6, 5, 5}}, 
 {51, {4, 5, 5, 4, 5, 6, 5, 5, 5, 5, 5, 5, 5, 5, 5, 5, 6, 5}}, 
 {51, {4, 4, 5, 5, 5, 5, 5, 5, 4, 5, 6, 5, 5, 6, 6, 5, 5, 5}}, 
 {51, {4, 4, 5, 5, 5, 5, 4, 5, 6, 6, 5, 5, 5, 5, 5, 5, 6, 5}}, 
 {51, {4, 4, 5, 5, 5, 4, 5, 5, 6, 5, 5, 5, 6, 6, 5, 5, 5, 5}}, 
 {51, {4, 4, 5, 5, 5, 4, 5, 5, 5, 5, 6, 5, 5, 6, 6, 5, 5, 5}}, 
 {51, {4, 4, 5, 5, 5, 4, 5, 5, 5, 5, 6, 5, 5, 5, 6, 6, 5, 5}}, 
 {51, {4, 4, 5, 5, 5, 4, 5, 5, 5, 5, 5, 6, 5, 5, 6, 6, 5, 5}}, 
 {51, {4, 4, 5, 5, 4, 5, 6, 5, 5, 5, 6, 6, 5, 5, 5, 5, 5, 5}}, 
 {51, {4, 4, 5, 5, 4, 5, 5, 6, 5, 5, 6, 6, 5, 5, 5, 5, 5, 5}}, 
 {51, {4, 4, 5, 5, 4, 5, 5, 6, 5, 5, 5, 6, 6, 5, 5, 5, 5, 5}}, 
 {51, {4, 4, 5, 5, 4, 5, 5, 5, 4, 5, 5, 6, 6, 6, 6, 5, 5, 5}}, 
 {51, {4, 4, 5, 4, 4, 5, 5, 5, 5, 5, 5, 6, 6, 6, 6, 5, 5, 5}}, 
 {51, {3, 5, 5, 5, 5, 6, 6, 5, 5, 5, 5, 5, 4, 5, 5, 5, 6, 5}}, 
 {51, {3, 5, 5, 5, 5, 5, 5, 5, 5, 5, 5, 5, 4, 5, 6, 6, 6, 5}}, 
 {51, {3, 5, 5, 4, 5, 6, 6, 6, 5, 5, 5, 5, 5, 5, 5, 5, 5, 5}}, 
 {51, {1, 6, 6, 1, 6, 6, 5, 5, 5, 6, 5, 5, 5, 5, 5, 6, 6, 6}}, 
 {51, {5, 5, 5, 6, 5, 5, 5, 6, 5, 5, 6, 5, 5, 6, 5, 5, 6}}, 
 {51, {4, 5, 5, 6, 5, 5, 6, 6, 6, 6, 5, 5, 4, 5, 6, 6, 5}}, 
 {51, {4, 5, 5, 6, 5, 5, 6, 6, 5, 5, 6, 6, 5, 5, 6, 5, 5}}, 
 {51, {4, 5, 5, 6, 5, 5, 4, 5, 6, 5, 5, 6, 6, 6, 6, 6, 5}}, 
 {51, {4, 4, 5, 6, 5, 5, 6, 6, 5, 6, 6, 5, 5, 5, 6, 6, 5}}, 
 {51, {4, 4, 5, 5, 6, 6, 5, 5, 6, 6, 5, 6, 6, 5, 5, 6, 5}}, 
 {51, {4, 4, 5, 4, 5, 5, 6, 6, 6, 6, 6, 5, 6, 6, 6, 5, 5}}, 
 {51, {3, 5, 5, 6, 6, 6, 6, 6, 5, 4, 5, 5, 5, 6, 6, 6, 5}}, 
 {51, {3, 5, 5, 5, 6, 6, 6, 5, 6, 6, 5, 5, 5, 6, 6, 5, 5}}, 
 {51, {3, 5, 5, 5, 6, 6, 5, 4, 5, 5, 6, 6, 6, 6, 6, 6, 5}}, 
 {51, {3, 5, 5, 5, 5, 6, 5, 5, 6, 6, 5, 6, 6, 6, 6, 5, 5}}, 
 {51, {3, 5, 5, 5, 4, 5, 5, 6, 6, 6, 5, 6, 6, 6, 6, 6, 5}}, 
 {51, {3, 5, 5, 4, 5, 5, 6, 6, 6, 6, 6, 5, 6, 6, 6, 5, 5}}, 
 {50, {4, 5, 5, 5, 5, 4, 5, 5, 6, 6, 5, 5, 5, 4, 5, 5, 6, 5}}, 
 {50, {4, 5, 5, 5, 4, 5, 5, 6, 6, 5, 5, 5, 5, 4, 5, 5, 6, 5}}, 
 {50, {4, 5, 5, 4, 5, 5, 5, 5, 5, 4, 5, 5, 5, 5, 6, 6, 6, 5}}, 
 {50, {4, 5, 4, 5, 5, 5, 5, 5, 5, 5, 6, 6, 5, 5, 5, 5, 5, 5}}, 
 {50, {4, 4, 5, 5, 5, 6, 6, 5, 5, 5, 4, 5, 5, 5, 5, 6, 5, 5}}, 
 {50, {4, 4, 5, 5, 5, 6, 5, 5, 5, 5, 5, 5, 4, 5, 6, 6, 5, 5}}, 
 {50, {4, 4, 5, 5, 5, 5, 5, 6, 6, 5, 4, 5, 5, 6, 5, 5, 5, 5}}, 
 {50, {4, 4, 5, 5, 5, 5, 5, 6, 5, 5, 5, 4, 5, 5, 5, 6, 6, 5}}, 
 {50, {4, 4, 5, 5, 5, 5, 5, 5, 5, 5, 6, 6, 5, 4, 5, 6, 5, 5}}, 
 {50, {4, 4, 5, 5, 5, 5, 5, 5, 4, 5, 6, 5, 5, 5, 5, 6, 6, 5}}, 
 {50, {3, 5, 6, 5, 5, 4, 5, 5, 5, 5, 5, 5, 5, 5, 5, 6, 6, 5}}, 
 {50, {3, 5, 5, 6, 6, 5, 5, 5, 5, 4, 5, 5, 5, 5, 5, 5, 6, 5}}, 
 {50, {3, 5, 5, 5, 5, 5, 5, 6, 5, 5, 5, 5, 5, 5, 5, 6, 5, 5}}, 
 {50, {3, 5, 5, 5, 5, 5, 5, 5, 5, 5, 5, 5, 6, 5, 5, 6, 5, 5}}, 
 {50, {3, 5, 5, 5, 5, 4, 5, 6, 6, 6, 5, 5, 5, 5, 5, 5, 5, 5}}, 
 {50, {1, 6, 6, 5, 5, 4, 5, 5, 5, 6, 5, 5, 5, 5, 5, 5, 6, 6}}, 
 {50, {1, 6, 6, 5, 4, 5, 5, 5, 5, 6, 5, 5, 5, 5, 5, 5, 6, 6}}, 
 {50, {4, 5, 5, 6, 5, 5, 5, 6, 6, 5, 4, 5, 6, 6, 6, 6, 5}}, 
 {50, {4, 5, 5, 4, 5, 6, 5, 5, 5, 6, 6, 6, 6, 5, 6, 6, 5}}, 
 {50, {4, 4, 5, 6, 5, 5, 6, 6, 5, 5, 6, 6, 5, 5, 6, 6, 5}}, 
 {50, {4, 4, 5, 4, 5, 6, 6, 6, 6, 6, 5, 5, 5, 6, 6, 6, 5}}, 
 {50, {3, 5, 6, 5, 5, 6, 5, 5, 6, 6, 6, 5, 5, 5, 6, 6, 5}}, 
 {50, {3, 5, 5, 6, 6, 6, 6, 5, 4, 5, 5, 5, 6, 6, 6, 6, 5}}, 
 {50, {3, 5, 5, 6, 5, 5, 5, 6, 6, 5, 5, 5, 6, 6, 6, 6, 5}}, 
 {50, {3, 5, 5, 5, 5, 6, 6, 5, 6, 6, 5, 5, 5, 6, 6, 6, 5}}, 
 {50, {3, 5, 5, 5, 5, 6, 5, 5, 6, 6, 6, 6, 6, 5, 5, 6, 5}}, 
 {50, {3, 5, 5, 5, 5, 6, 5, 5, 5, 6, 6, 5, 6, 6, 6, 6, 5}}, 
 {50, {3, 5, 5, 5, 5, 5, 6, 6, 5, 6, 6, 5, 6, 6, 6, 5, 5}}, 
 {50, {1, 6, 6, 5, 5, 6, 6, 5, 5, 6, 5, 5, 6, 6, 5, 6, 6}}, 
 {49, {4, 5, 5, 5, 5, 5, 4, 5, 5, 6, 6, 5, 5, 4, 5, 5, 6, 5}}, 
 {49, {4, 5, 5, 4, 5, 5, 6, 5, 5, 5, 5, 4, 5, 5, 6, 6, 5, 5}}, 
 {49, {4, 4, 5, 5, 6, 5, 5, 5, 4, 5, 5, 5, 5, 5, 6, 6, 5, 5}}, 
 {49, {4, 4, 5, 5, 5, 6, 6, 5, 5, 5, 5, 4, 5, 5, 5, 5, 6, 5}}, 
 {49, {4, 4, 5, 5, 5, 6, 5, 5, 5, 4, 5, 5, 5, 5, 5, 6, 6, 5}}, 
 {49, {4, 4, 5, 5, 5, 5, 5, 5, 6, 5, 5, 4, 5, 5, 5, 6, 6, 5}}, 
 {49, {4, 4, 5, 5, 5, 5, 5, 5, 5, 6, 5, 5, 4, 5, 5, 6, 6, 5}}, 
 {49, {4, 4, 5, 5, 5, 5, 5, 5, 5, 5, 6, 6, 5, 4, 5, 5, 6, 5}}, 
 {49, {4, 4, 5, 5, 5, 5, 5, 4, 5, 5, 5, 6, 6, 5, 5, 5, 6, 5}}, 
 {49, {4, 4, 5, 5, 5, 5, 5, 4, 5, 5, 5, 6, 5, 5, 6, 6, 5, 5}}, 
 {49, {4, 4, 5, 5, 5, 4, 5, 6, 6, 5, 5, 5, 5, 5, 6, 5, 5, 5}}, 
 {49, {4, 4, 5, 5, 4, 5, 5, 5, 6, 6, 5, 5, 6, 5, 5, 5, 5, 5}}, 
 {49, {1, 6, 6, 5, 5, 4, 5, 5, 6, 5, 5, 5, 5, 5, 5, 5, 6, 6}}, 
 {49, {1, 6, 6, 5, 5, 4, 5, 5, 5, 5, 5, 5, 6, 5, 5, 5, 6, 6}}, 
 {49, {1, 6, 6, 1, 6, 6, 5, 5, 5, 5, 6, 5, 5, 5, 5, 6, 6, 6}}, 
 {49, {4, 5, 5, 6, 5, 4, 5, 6, 6, 6, 6, 6, 5, 5, 6, 5, 5}}, 
 {49, {4, 5, 5, 5, 6, 5, 5, 6, 6, 6, 5, 4, 5, 6, 6, 6, 5}}, 
 {49, {4, 5, 5, 4, 5, 6, 6, 5, 6, 6, 5, 5, 5, 6, 6, 6, 5}}, 
 {49, {4, 4, 5, 6, 6, 6, 6, 5, 5, 4, 5, 5, 6, 6, 6, 6, 5}}, 
 {49, {3, 5, 6, 6, 6, 5, 4, 5, 5, 5, 5, 6, 6, 6, 6, 6, 5}}, 
 {49, {3, 5, 6, 5, 5, 6, 5, 5, 5, 6, 6, 6, 5, 5, 6, 6, 5}}, 
 {49, {3, 5, 6, 5, 5, 5, 6, 6, 6, 6, 5, 5, 6, 5, 5, 6, 5}}, 
 {49, {3, 5, 6, 5, 5, 5, 6, 6, 5, 6, 6, 6, 5, 5, 5, 6, 5}}, 
 {49, {3, 5, 6, 5, 5, 5, 6, 5, 5, 5, 6, 6, 6, 5, 6, 6, 5}}, 
 {49, {3, 5, 5, 6, 6, 6, 5, 5, 6, 5, 5, 6, 6, 5, 5, 6, 5}}, 
 {49, {3, 5, 5, 6, 6, 6, 5, 4, 5, 5, 5, 6, 6, 6, 6, 6, 5}}, 
 {49, {3, 5, 5, 5, 5, 6, 6, 6, 6, 5, 5, 6, 5, 5, 6, 6, 5}}, 
 {49, {3, 5, 5, 5, 5, 5, 6, 6, 5, 6, 6, 5, 5, 6, 6, 6, 5}}, 
 {48, {4, 5, 5, 5, 5, 5, 6, 5, 5, 4, 5, 5, 6, 5, 5, 5, 5, 5}}, 
 {48, {4, 5, 5, 5, 4, 5, 5, 5, 5, 5, 5, 5, 4, 5, 6, 6, 6, 5}}, 
 {48, {4, 5, 5, 4, 5, 5, 5, 6, 5, 4, 5, 5, 5, 5, 6, 6, 5, 5}}, 
 {48, {4, 4, 5, 5, 6, 5, 5, 5, 5, 5, 5, 4, 5, 5, 5, 6, 6, 5}}, 
 {48, {4, 4, 5, 5, 5, 6, 6, 5, 4, 5, 5, 5, 5, 5, 5, 5, 6, 5}}, 
 {48, {4, 4, 5, 5, 5, 5, 6, 5, 5, 4, 5, 5, 5, 5, 5, 6, 6, 5}}, 
 {48, {4, 4, 5, 5, 5, 5, 5, 6, 6, 5, 5, 5, 4, 5, 5, 6, 5, 5}}, 
 {48, {4, 4, 5, 5, 5, 5, 5, 6, 5, 5, 4, 5, 5, 5, 5, 6, 6, 5}}, 
 {48, {4, 4, 5, 5, 5, 5, 5, 5, 6, 5, 5, 4, 5, 6, 6, 5, 5, 5}}, 
 {48, {4, 4, 5, 5, 5, 5, 5, 5, 5, 6, 6, 5, 4, 5, 5, 6, 5, 5}}, 
 {48, {4, 4, 5, 5, 5, 5, 5, 4, 5, 5, 6, 5, 5, 5, 5, 6, 6, 5}}, 
 {48, {4, 4, 5, 5, 5, 5, 5, 4, 5, 5, 5, 5, 6, 6, 5, 5, 6, 5}}, 
 {48, {4, 4, 5, 5, 5, 4, 5, 5, 5, 6, 6, 5, 5, 6, 5, 5, 5, 5}}, 
 {48, {4, 4, 5, 5, 4, 5, 5, 5, 6, 6, 5, 5, 5, 6, 5, 5, 5, 5}}, 
 {48, {4, 4, 5, 5, 4, 5, 5, 5, 5, 6, 6, 5, 5, 5, 6, 5, 5, 5}}, 
 {48, {4, 4, 5, 4, 5, 6, 6, 6, 5, 5, 5, 5, 5, 5, 5, 5, 5, 5}}, 
 {48, {4, 4, 5, 4, 4, 5, 5, 5, 5, 5, 6, 6, 6, 6, 5, 5, 5, 5}}, 
 {48, {3, 5, 6, 5, 5, 5, 5, 5, 5, 5, 5, 4, 5, 5, 5, 6, 6, 5}}, 
 {48, {3, 5, 6, 5, 5, 5, 5, 5, 5, 5, 4, 5, 5, 5, 5, 6, 6, 5}}, 
 {48, {3, 5, 6, 5, 5, 5, 4, 5, 5, 5, 5, 5, 5, 5, 5, 6, 6, 5}}, 
 {48, {3, 5, 5, 6, 6, 5, 5, 5, 5, 5, 5, 4, 5, 5, 5, 5, 6, 5}}, 
 {48, {3, 5, 5, 5, 5, 5, 5, 6, 5, 5, 5, 6, 5, 5, 5, 5, 5, 5}}, 
 {48, {3, 5, 5, 5, 5, 4, 5, 5, 5, 5, 5, 5, 6, 6, 6, 5, 5, 5}}, 
 {48, {1, 6, 6, 6, 5, 4, 5, 5, 5, 4, 5, 5, 5, 5, 5, 6, 6, 6}}, 
 {48, {1, 6, 6, 6, 5, 3, 5, 5, 5, 5, 5, 5, 5, 5, 5, 6, 6, 6}}, 
 {48, {1, 6, 6, 5, 5, 5, 5, 4, 5, 6, 5, 5, 5, 5, 5, 5, 6, 6}}, 
 {48, {1, 6, 6, 5, 5, 5, 5, 4, 5, 5, 6, 5, 5, 5, 5, 5, 6, 6}}, 
 {48, {1, 6, 6, 5, 5, 4, 5, 5, 5, 5, 5, 5, 5, 6, 5, 5, 6, 6}}, 
 {48, {1, 6, 6, 5, 4, 5, 5, 5, 6, 5, 5, 5, 5, 5, 5, 5, 6, 6}}, 
 {48, {1, 6, 6, 5, 3, 5, 5, 5, 5, 5, 5, 5, 5, 5, 6, 6, 6, 6}}, 
 {48, {4, 5, 5, 5, 5, 6, 6, 5, 6, 6, 6, 5, 4, 5, 6, 6, 5}}, 
 {48, {4, 5, 5, 5, 5, 6, 6, 5, 6, 6, 5, 4, 5, 6, 6, 6, 5}}, 
 {48, {4, 5, 5, 5, 5, 6, 5, 4, 5, 6, 6, 5, 6, 6, 6, 6, 5}}, 
 {48, {4, 5, 4, 5, 5, 6, 6, 6, 5, 5, 5, 5, 6, 6, 6, 6, 5}}, 
 {48, {4, 4, 5, 6, 5, 5, 6, 5, 5, 6, 6, 6, 5, 5, 6, 6, 5}}, 
 {48, {4, 4, 5, 5, 5, 6, 6, 5, 6, 6, 6, 5, 6, 6, 5, 5, 5}}, 
 {48, {4, 4, 5, 4, 5, 5, 6, 6, 5, 5, 6, 6, 6, 6, 6, 6, 5}}, 
 {48, {3, 5, 6, 5, 5, 5, 6, 5, 5, 6, 6, 6, 5, 5, 6, 6, 5}}, 
 {48, {3, 5, 5, 6, 6, 5, 5, 6, 6, 5, 5, 6, 6, 5, 5, 6, 5}}, 
 {48, {3, 5, 5, 5, 6, 6, 5, 6, 6, 5, 5, 6, 6, 6, 5, 5, 5}}, 
 {48, {3, 5, 5, 5, 6, 6, 5, 5, 6, 5, 5, 5, 6, 6, 6, 6, 5}}, 
 {48, {3, 5, 5, 5, 5, 6, 6, 5, 5, 6, 5, 5, 6, 6, 6, 6, 5}}, 
 {48, {3, 5, 5, 5, 4, 5, 5, 6, 6, 6, 6, 6, 6, 5, 6, 6, 5}}, 
 {48, {3, 5, 5, 4, 5, 5, 6, 6, 6, 5, 5, 6, 6, 6, 6, 6, 5}}, 
 {48, {1, 6, 6, 5, 5, 5, 6, 6, 5, 4, 5, 6, 6, 6, 6, 6, 6}}, 
 {48, {1, 6, 6, 5, 4, 5, 5, 5, 6, 6, 5, 6, 6, 6, 6, 6, 6}}, 
 {47, {4, 5, 5, 5, 5, 5, 5, 5, 4, 5, 5, 6, 5, 5, 5, 6, 5, 5}}, 
 {47, {4, 5, 5, 5, 5, 4, 5, 5, 5, 6, 5, 5, 4, 5, 5, 6, 6, 5}}, 
 {47, {4, 4, 5, 5, 6, 5, 5, 5, 5, 5, 4, 5, 5, 5, 6, 6, 5, 5}}, 
 {47, {4, 4, 5, 5, 5, 5, 6, 5, 5, 5, 5, 5, 5, 4, 5, 6, 6, 5}}, 
 {47, {4, 4, 5, 5, 5, 5, 5, 5, 6, 5, 5, 5, 4, 5, 6, 6, 5, 5}}, 
 {47, {4, 4, 5, 5, 5, 5, 5, 5, 5, 5, 6, 5, 5, 4, 5, 6, 6, 5}}, 
 {47, {4, 4, 5, 5, 5, 5, 5, 4, 5, 6, 5, 5, 5, 6, 6, 5, 5, 5}}, 
 {47, {4, 4, 5, 5, 5, 4, 5, 5, 5, 5, 5, 6, 5, 5, 5, 6, 6, 5}}, 
 {47, {3, 5, 5, 6, 6, 5, 5, 5, 5, 5, 4, 5, 5, 5, 5, 5, 6, 5}}, 
 {47, {3, 5, 5, 5, 4, 5, 5, 5, 5, 5, 5, 5, 6, 6, 5, 5, 6, 5}}, 
 {47, {1, 6, 6, 5, 5, 4, 5, 5, 5, 5, 4, 5, 5, 5, 6, 6, 6, 6}}, 
 {47, {4, 5, 5, 5, 6, 5, 5, 4, 5, 6, 6, 6, 6, 5, 6, 6, 5}}, 
 {47, {4, 5, 5, 5, 5, 6, 6, 6, 5, 4, 5, 6, 6, 5, 6, 6, 5}}, 
 {47, {4, 5, 5, 4, 5, 5, 6, 6, 5, 6, 6, 6, 5, 6, 6, 5, 5}}, 
 {47, {4, 5, 4, 5, 5, 5, 6, 5, 5, 6, 6, 6, 6, 6, 6, 5, 5}}, 
 {47, {4, 4, 5, 5, 6, 6, 6, 5, 6, 6, 5, 5, 6, 5, 5, 6, 5}}, 
 {47, {4, 4, 5, 5, 5, 6, 6, 5, 4, 5, 6, 6, 6, 6, 6, 6, 5}}, 
 {47, {3, 5, 6, 5, 5, 6, 6, 5, 5, 5, 6, 6, 5, 5, 6, 6, 5}}, 
 {47, {3, 5, 6, 5, 5, 6, 6, 5, 5, 5, 5, 6, 6, 5, 6, 6, 5}}, 
 {47, {3, 5, 5, 5, 6, 5, 5, 5, 5, 6, 6, 6, 6, 5, 6, 6, 5}}, 
 {47, {3, 5, 5, 5, 5, 6, 5, 5, 6, 6, 5, 5, 6, 6, 6, 6, 5}}, 
 {47, {3, 5, 5, 5, 5, 5, 5, 6, 6, 5, 6, 6, 6, 5, 6, 6, 5}}, 
 {47, {3, 5, 5, 5, 4, 5, 5, 6, 6, 5, 6, 6, 6, 6, 6, 6, 5}}, 
 {47, {3, 5, 5, 4, 5, 6, 6, 6, 6, 6, 5, 5, 5, 6, 6, 6, 5}}, 
 {46, {4, 5, 5, 5, 4, 5, 5, 6, 5, 4, 5, 5, 5, 5, 6, 6, 5, 5}}, 
 {46, {4, 5, 5, 5, 4, 5, 5, 5, 5, 6, 5, 5, 5, 6, 5, 5, 5, 5}}, 
 {46, {4, 5, 5, 5, 4, 5, 5, 5, 5, 4, 5, 6, 5, 5, 6, 6, 5, 5}}, 
 {46, {4, 5, 5, 4, 5, 5, 5, 6, 5, 5, 5, 5, 5, 5, 6, 5, 5, 5}}, 
 {46, {4, 4, 5, 5, 6, 5, 4, 5, 5, 5, 5, 5, 5, 5, 5, 6, 6, 5}}, 
 {46, {4, 4, 5, 5, 5, 5, 5, 6, 6, 5, 5, 5, 5, 4, 5, 6, 5, 5}}, 
 {46, {4, 4, 5, 5, 5, 5, 5, 5, 6, 6, 5, 4, 5, 5, 6, 5, 5, 5}}, 
 {46, {4, 4, 5, 5, 5, 5, 5, 4, 5, 6, 6, 5, 5, 5, 5, 6, 5, 5}}, 
 {46, {4, 4, 5, 5, 5, 5, 5, 4, 5, 5, 5, 6, 6, 5, 5, 6, 5, 5}}, 
 {46, {4, 4, 5, 5, 5, 5, 4, 5, 5, 5, 6, 6, 5, 5, 5, 6, 5, 5}}, 
 {46, {4, 4, 5, 5, 5, 4, 5, 5, 4, 5, 5, 6, 6, 6, 6, 5, 5, 5}}, 
 {46, {4, 4, 5, 5, 4, 5, 6, 6, 5, 5, 5, 6, 5, 5, 5, 5, 5, 5}}, 
 {46, {4, 4, 5, 5, 4, 5, 6, 6, 5, 5, 5, 5, 5, 5, 5, 5, 6, 5}}, 
 {46, {4, 4, 5, 5, 4, 5, 5, 5, 5, 6, 6, 6, 6, 5, 5, 4, 5, 5}}, 
 {46, {3, 5, 5, 5, 5, 6, 6, 5, 5, 5, 5, 5, 5, 4, 5, 5, 6, 5}}, 
 {46, {3, 5, 5, 5, 5, 5, 5, 6, 6, 5, 5, 4, 5, 5, 5, 5, 6, 5}}, 
 {46, {1, 6, 6, 5, 4, 5, 5, 5, 5, 5, 6, 5, 5, 5, 5, 5, 6, 6}}, 
 {46, {1, 6, 6, 5, 4, 5, 5, 5, 4, 5, 5, 5, 5, 5, 6, 6, 6, 6}}, 
 {46, {4, 5, 5, 5, 6, 5, 5, 4, 5, 6, 6, 6, 5, 6, 6, 6, 5}}, 
 {46, {4, 5, 5, 4, 5, 6, 5, 5, 6, 6, 5, 5, 6, 6, 6, 6, 5}}, 
 {46, {4, 4, 5, 5, 6, 6, 5, 6, 6, 6, 5, 5, 6, 5, 5, 6, 5}}, 
 {46, {4, 4, 5, 5, 4, 5, 6, 6, 5, 6, 6, 6, 6, 6, 6, 5, 5}}, 
 {46, {3, 5, 5, 6, 6, 6, 6, 6, 5, 5, 5, 4, 5, 6, 6, 6, 5}}, 
 {46, {3, 5, 5, 6, 6, 5, 5, 6, 6, 5, 6, 6, 5, 5, 5, 6, 5}}, 
 {46, {3, 5, 5, 6, 5, 5, 6, 6, 6, 5, 5, 5, 6, 6, 6, 5, 5}}, 
 {46, {3, 5, 5, 6, 5, 5, 5, 5, 6, 6, 6, 5, 6, 6, 6, 5, 5}}, 
 {46, {3, 5, 5, 6, 5, 4, 5, 5, 6, 6, 6, 6, 6, 6, 6, 5, 5}}, 
 {46, {3, 5, 5, 5, 6, 6, 6, 6, 5, 5, 5, 6, 5, 5, 6, 6, 5}}, 
 {46, {3, 5, 5, 5, 6, 6, 5, 5, 6, 6, 5, 6, 6, 6, 5, 5, 5}}, 
 {46, {3, 5, 5, 5, 6, 5, 5, 6, 5, 5, 6, 6, 6, 6, 6, 5, 5}}, 
 {46, {3, 5, 5, 4, 5, 6, 6, 6, 6, 5, 5, 5, 6, 6, 6, 6, 5}}, 
 {46, {1, 6, 6, 5, 6, 6, 5, 5, 5, 4, 5, 6, 6, 6, 6, 6, 6}}, 
 {46, {1, 6, 6, 5, 5, 4, 4, 5, 6, 6, 6, 6, 6, 6, 6, 6, 6}}, 
 {45, {4, 5, 5, 5, 5, 5, 5, 6, 5, 5, 5, 5, 5, 5, 4, 5, 6, 5}}, 
 {45, {4, 5, 5, 5, 4, 5, 5, 5, 5, 5, 6, 5, 4, 5, 6, 6, 5, 5}}, 
 {45, {4, 5, 5, 4, 5, 5, 6, 6, 5, 5, 5, 5, 5, 4, 5, 5, 6, 5}}, 
 {45, {4, 4, 5, 6, 5, 5, 5, 5, 5, 5, 5, 5, 5, 4, 5, 6, 6, 5}}, 
 {45, {4, 4, 5, 5, 5, 5, 6, 5, 4, 5, 5, 5, 5, 5, 5, 6, 6, 5}}, 
 {45, {4, 4, 5, 5, 5, 5, 5, 5, 5, 6, 5, 4, 5, 6, 6, 5, 5, 5}}, 
 {45, {4, 4, 5, 5, 5, 5, 5, 5, 5, 4, 5, 6, 6, 5, 5, 5, 6, 5}}, 
 {45, {4, 4, 5, 5, 5, 5, 4, 5, 6, 6, 5, 5, 5, 5, 6, 5, 5, 5}}, 
 {45, {4, 4, 5, 5, 5, 5, 4, 5, 5, 5, 5, 6, 5, 5, 6, 6, 5, 5}}, 
 {45, {4, 4, 5, 5, 5, 4, 5, 5, 5, 5, 6, 6, 5, 5, 5, 6, 5, 5}}, 
 {45, {4, 4, 5, 5, 4, 5, 5, 6, 6, 6, 6, 5, 5, 5, 4, 5, 5, 5}}, 
 {45, {4, 4, 5, 5, 4, 5, 5, 5, 6, 6, 5, 5, 5, 5, 5, 6, 5, 5}}, 
 {45, {4, 4, 5, 5, 4, 5, 5, 4, 5, 5, 6, 6, 6, 6, 5, 5, 5, 5}}, 
 {45, {4, 4, 5, 4, 4, 5, 5, 5, 5, 5, 5, 5, 5, 6, 6, 6, 6, 5}}, 
 {45, {3, 5, 5, 5, 5, 5, 6, 6, 5, 5, 5, 5, 5, 4, 5, 5, 6, 5}}, 
 {45, {3, 5, 5, 5, 5, 5, 6, 5, 5, 5, 5, 5, 5, 5, 5, 6, 5, 5}}, 
 {45, {3, 5, 5, 5, 5, 5, 5, 5, 6, 6, 5, 5, 5, 4, 5, 5, 6, 5}}, 
 {45, {3, 5, 5, 5, 5, 5, 5, 5, 5, 5, 6, 6, 5, 4, 5, 5, 6, 5}}, 
 {45, {1, 6, 6, 6, 5, 4, 5, 5, 5, 5, 5, 5, 4, 5, 5, 6, 6, 6}}, 
 {45, {4, 5, 5, 6, 5, 5, 5, 6, 5, 4, 5, 6, 6, 6, 6, 6, 5}}, 
 {45, {4, 5, 4, 5, 5, 6, 6, 6, 6, 5, 5, 5, 5, 6, 6, 6, 5}}, 
 {45, {4, 4, 5, 6, 5, 5, 6, 6, 6, 5, 5, 6, 5, 5, 6, 6, 5}}, 
 {45, {4, 4, 5, 4, 5, 6, 6, 6, 6, 6, 6, 5, 5, 5, 6, 6, 5}}, 
 {45, {4, 4, 5, 4, 5, 6, 6, 6, 5, 5, 5, 6, 6, 6, 6, 6, 5}}, 
 {45, {4, 4, 5, 4, 5, 5, 6, 6, 6, 6, 6, 6, 5, 6, 6, 5, 5}}, 
 {45, {3, 5, 6, 5, 5, 6, 6, 5, 5, 5, 6, 5, 5, 6, 6, 6, 5}}, 
 {45, {3, 5, 6, 5, 5, 5, 5, 6, 6, 5, 6, 6, 6, 5, 5, 6, 5}}, 
 {45, {3, 5, 5, 5, 6, 6, 6, 6, 5, 5, 6, 5, 5, 6, 6, 5, 5}}, 
 {45, {3, 5, 5, 5, 6, 5, 5, 5, 6, 6, 5, 5, 6, 6, 6, 6, 5}}, 
 {45, {3, 5, 5, 5, 5, 6, 6, 5, 5, 5, 6, 6, 5, 6, 6, 6, 5}}, 
 {45, {3, 5, 5, 5, 5, 5, 6, 6, 6, 5, 6, 6, 5, 6, 6, 5, 5}}, 
 {45, {3, 5, 5, 5, 5, 5, 6, 6, 5, 6, 6, 6, 5, 6, 6, 5, 5}}, 
 {44, {4, 5, 5, 5, 5, 5, 5, 5, 5, 5, 5, 4, 5, 6, 5, 5, 6, 5}}, 
 {44, {4, 5, 5, 5, 4, 5, 5, 5, 6, 6, 5, 5, 5, 4, 5, 5, 6, 5}}, 
 {44, {4, 5, 5, 5, 4, 5, 5, 5, 5, 6, 6, 5, 5, 4, 5, 5, 6, 5}}, 
 {44, {4, 5, 5, 4, 5, 5, 6, 6, 5, 5, 4, 5, 5, 5, 5, 5, 6, 5}}, 
 {44, {4, 5, 5, 4, 5, 5, 5, 6, 6, 5, 5, 4, 5, 5, 5, 6, 5, 5}}, 
 {44, {4, 5, 5, 4, 5, 5, 5, 6, 5, 5, 5, 4, 5, 5, 6, 6, 5, 5}}, 
 {44, {4, 5, 5, 4, 5, 5, 5, 6, 5, 4, 5, 5, 5, 6, 6, 5, 5, 5}}, 
 {44, {4, 4, 5, 5, 5, 6, 5, 5, 5, 5, 5, 4, 5, 5, 5, 6, 6, 5}}, 
 {44, {4, 4, 5, 5, 5, 5, 6, 6, 5, 5, 5, 5, 4, 5, 5, 6, 5, 5}}, 
 {44, {4, 4, 5, 5, 5, 5, 6, 6, 5, 4, 5, 5, 5, 5, 5, 6, 5, 5}}, 
 {44, {4, 4, 5, 5, 5, 5, 6, 5, 5, 5, 4, 5, 6, 6, 5, 5, 5, 5}}, 
 {44, {4, 4, 5, 5, 5, 5, 5, 5, 6, 6, 5, 4, 5, 5, 5, 6, 5, 5}}, 
 {44, {4, 4, 5, 5, 5, 4, 5, 6, 6, 5, 5, 5, 6, 5, 5, 5, 5, 5}}, 
 {44, {4, 4, 5, 5, 5, 4, 5, 5, 6, 5, 5, 6, 6, 5, 5, 5, 5, 5}}, 
 {44, {4, 4, 5, 4, 4, 5, 5, 5, 5, 5, 5, 5, 6, 6, 6, 6, 5, 5}}, 
 {44, {3, 5, 5, 5, 5, 4, 5, 5, 5, 5, 5, 6, 6, 5, 5, 5, 6, 5}}, 
 {44, {3, 5, 5, 4, 5, 5, 5, 5, 6, 6, 5, 5, 5, 5, 5, 5, 6, 5}}, 
 {44, {4, 5, 5, 5, 6, 6, 6, 6, 5, 4, 5, 6, 5, 5, 6, 6, 5}}, 
 {44, {4, 5, 5, 5, 6, 5, 5, 6, 5, 4, 5, 6, 6, 6, 6, 6, 5}}, 
 {44, {3, 5, 6, 5, 5, 6, 5, 5, 5, 6, 6, 6, 6, 5, 5, 6, 5}}, 
 {44, {3, 5, 6, 5, 5, 5, 5, 6, 6, 5, 6, 6, 5, 5, 6, 6, 5}}, 
 {44, {3, 5, 5, 6, 5, 5, 5, 6, 6, 6, 6, 5, 5, 6, 6, 5, 5}}, 
 {44, {3, 5, 5, 5, 6, 6, 6, 6, 5, 5, 6, 5, 5, 5, 6, 6, 5}}, 
 {44, {3, 5, 5, 5, 5, 6, 6, 6, 6, 5, 4, 5, 6, 6, 6, 6, 5}}, 
 {44, {3, 5, 5, 5, 5, 6, 5, 5, 6, 6, 6, 5, 6, 6, 6, 5, 5}}, 
 {44, {3, 5, 5, 5, 5, 6, 5, 5, 6, 6, 6, 5, 5, 6, 6, 6, 5}}, 
 {44, {3, 5, 5, 5, 5, 5, 4, 5, 6, 6, 6, 6, 6, 6, 6, 6, 5}}, 
 {44, {3, 5, 5, 4, 5, 5, 6, 6, 6, 6, 6, 6, 5, 6, 6, 5, 5}}, 
 {44, {3, 5, 5, 4, 5, 5, 6, 6, 5, 6, 6, 6, 6, 6, 6, 5, 5}}, 
 {44, {1, 6, 6, 5, 5, 5, 4, 5, 6, 6, 5, 6, 6, 6, 6, 6, 6}}, 
 {43, {4, 5, 5, 5, 4, 5, 5, 5, 4, 5, 5, 6, 6, 5, 5, 5, 6, 5}}, 
 {43, {4, 5, 5, 4, 5, 5, 6, 5, 4, 5, 5, 5, 5, 5, 6, 6, 5, 5}}, 
 {43, {4, 5, 5, 4, 5, 5, 5, 5, 6, 6, 5, 5, 4, 5, 6, 5, 5, 5}}, 
 {43, {4, 4, 5, 5, 6, 5, 5, 5, 5, 4, 5, 5, 5, 5, 5, 6, 6, 5}}, 
 {43, {4, 4, 5, 5, 5, 5, 5, 6, 6, 5, 4, 5, 5, 5, 6, 5, 5, 5}}, 
 {43, {4, 4, 5, 5, 5, 5, 5, 5, 5, 5, 4, 5, 6, 6, 5, 5, 6, 5}}, 
 {43, {4, 4, 5, 5, 5, 5, 5, 5, 4, 5, 5, 6, 5, 5, 6, 6, 5, 5}}, 
 {43, {4, 4, 5, 5, 5, 5, 5, 5, 4, 5, 5, 6, 5, 5, 5, 6, 6, 5}}, 
 {43, {4, 4, 5, 5, 5, 4, 5, 5, 6, 6, 6, 6, 5, 4, 5, 5, 5, 5}}, 
 {43, {4, 4, 5, 5, 4, 5, 5, 5, 5, 5, 6, 5, 5, 5, 6, 6, 5, 5}}, 
 {43, {4, 4, 5, 5, 4, 5, 5, 5, 5, 5, 5, 6, 5, 5, 5, 6, 6, 5}}, 
 {43, {3, 5, 6, 5, 5, 4, 5, 6, 5, 5, 5, 5, 5, 5, 5, 5, 6, 5}}, 
 {43, {3, 5, 5, 6, 6, 5, 5, 5, 5, 5, 5, 5, 5, 5, 4, 5, 6, 5}}, 
 {43, {3, 5, 5, 5, 5, 6, 6, 5, 5, 5, 4, 5, 5, 5, 5, 5, 6, 5}}, 
 {43, {3, 5, 5, 5, 5, 5, 5, 6, 6, 5, 5, 5, 5, 4, 5, 5, 6, 5}}, 
 {43, {3, 5, 5, 5, 5, 5, 5, 5, 6, 5, 5, 5, 4, 5, 5, 6, 6, 5}}, 
 {43, {3, 5, 5, 5, 5, 5, 5, 5, 5, 6, 5, 5, 5, 5, 5, 6, 5, 5}}, 
 {43, {3, 5, 5, 5, 5, 5, 5, 5, 5, 5, 6, 5, 4, 5, 5, 6, 6, 5}}, 
 {43, {3, 5, 5, 5, 5, 5, 5, 5, 5, 4, 5, 6, 6, 6, 5, 5, 5, 5}}, 
 {43, {3, 5, 5, 5, 4, 5, 6, 6, 5, 5, 5, 5, 5, 5, 5, 5, 6, 5}}, 
 {43, {3, 5, 5, 5, 4, 5, 5, 5, 5, 5, 5, 6, 6, 5, 5, 5, 6, 5}}, 
 {43, {1, 6, 6, 5, 5, 5, 4, 5, 6, 5, 5, 5, 5, 5, 5, 5, 6, 6}}, 
 {43, {1, 6, 6, 5, 5, 4, 5, 5, 4, 5, 5, 5, 5, 5, 6, 6, 6, 6}}, 
 {43, {4, 5, 5, 5, 4, 5, 6, 6, 5, 5, 6, 6, 5, 6, 6, 6, 5}}, 
 {43, {4, 5, 4, 5, 5, 5, 5, 6, 6, 6, 6, 6, 5, 6, 6, 5, 5}}, 
 {43, {4, 4, 5, 5, 6, 6, 5, 6, 6, 5, 6, 6, 5, 5, 5, 6, 5}}, 
 {43, {4, 4, 5, 5, 4, 5, 6, 6, 6, 6, 6, 6, 5, 6, 6, 5, 5}}, 
 {43, {4, 4, 5, 5, 4, 5, 6, 5, 5, 6, 6, 6, 6, 6, 6, 6, 5}}, 
 {43, {3, 5, 6, 5, 5, 6, 5, 5, 5, 5, 6, 6, 6, 5, 6, 6, 5}}, 
 {43, {3, 5, 5, 6, 5, 5, 5, 6, 6, 6, 5, 5, 6, 6, 6, 5, 5}}, 
 {43, {3, 5, 5, 5, 6, 5, 5, 5, 5, 6, 6, 5, 6, 6, 6, 6, 5}}, 
 {43, {3, 5, 5, 5, 6, 5, 4, 5, 5, 6, 6, 6, 6, 6, 6, 6, 5}}, 
 {43, {3, 5, 5, 5, 5, 6, 6, 6, 6, 6, 5, 4, 5, 6, 6, 6, 5}}, 
 {43, {3, 5, 5, 5, 5, 6, 5, 5, 5, 6, 6, 6, 6, 5, 6, 6, 5}}, 
 {43, {3, 5, 5, 5, 4, 5, 6, 6, 6, 6, 5, 5, 6, 6, 6, 6, 5}}, 
 {43, {3, 5, 5, 4, 5, 6, 5, 5, 5, 6, 6, 6, 6, 6, 6, 6, 5}}, 
 {43, {1, 6, 6, 5, 5, 5, 4, 5, 6, 6, 6, 6, 6, 5, 6, 6, 6}}, 
 {42, {4, 5, 5, 5, 5, 5, 4, 5, 5, 5, 6, 5, 4, 5, 6, 6, 5, 5}}, 
 {42, {4, 5, 5, 5, 4, 5, 5, 5, 5, 5, 6, 5, 4, 5, 5, 6, 6, 5}}, 
 {42, {4, 5, 5, 4, 5, 5, 6, 6, 5, 5, 5, 5, 4, 5, 5, 5, 6, 5}}, 
 {42, {4, 5, 5, 4, 5, 5, 6, 5, 5, 5, 4, 5, 5, 5, 6, 6, 5, 5}}, 
 {42, {4, 5, 5, 4, 5, 5, 5, 5, 6, 5, 5, 5, 5, 6, 5, 5, 5, 5}}, 
 {42, {4, 5, 5, 4, 5, 5, 5, 5, 5, 5, 5, 5, 4, 5, 6, 6, 6, 5}}, 
 {42, {4, 5, 5, 4, 5, 5, 4, 5, 5, 5, 5, 5, 5, 5, 6, 6, 6, 5}}, 
 {42, {4, 4, 5, 5, 5, 6, 5, 5, 5, 5, 4, 5, 5, 5, 6, 6, 5, 5}}, 
 {42, {4, 4, 5, 5, 5, 5, 5, 6, 5, 5, 5, 5, 4, 5, 6, 6, 5, 5}}, 
 {42, {4, 4, 5, 5, 5, 5, 5, 5, 5, 4, 5, 5, 6, 5, 5, 6, 6, 5}}, 
 {42, {4, 4, 5, 5, 5, 5, 4, 5, 6, 5, 5, 5, 5, 5, 5, 6, 6, 5}}, 
 {42, {4, 4, 5, 5, 5, 5, 4, 5, 5, 6, 5, 5, 5, 6, 6, 5, 5, 5}}, 
 {42, {4, 4, 5, 5, 5, 5, 4, 5, 5, 5, 6, 6, 5, 5, 6, 5, 5, 5}}, 
 {42, {4, 4, 5, 5, 5, 4, 5, 5, 6, 5, 5, 5, 5, 5, 5, 6, 6, 5}}, 
 {42, {4, 4, 5, 5, 4, 5, 6, 6, 6, 6, 5, 5, 5, 5, 4, 5, 5, 5}}, 
 {42, {4, 4, 5, 5, 4, 5, 6, 6, 6, 6, 5, 5, 4, 5, 5, 5, 5, 5}}, 
 {42, {4, 4, 5, 5, 4, 5, 5, 5, 6, 5, 5, 6, 6, 5, 5, 5, 5, 5}}, 
 {42, {4, 4, 5, 5, 4, 5, 5, 5, 5, 6, 5, 5, 6, 6, 5, 5, 5, 5}}, 
 {42, {4, 4, 5, 5, 4, 5, 5, 5, 5, 6, 5, 5, 5, 6, 6, 5, 5, 5}}, 
 {42, {4, 4, 5, 5, 4, 5, 5, 5, 4, 5, 5, 5, 6, 6, 6, 6, 5, 5}}, 
 {42, {3, 5, 5, 5, 5, 5, 5, 6, 6, 5, 5, 5, 4, 5, 5, 5, 6, 5}}, 
 {42, {3, 5, 5, 5, 5, 5, 5, 4, 5, 5, 5, 6, 6, 5, 5, 5, 6, 5}}, 
 {42, {1, 6, 6, 5, 5, 5, 5, 5, 5, 5, 5, 4, 4, 5, 6, 6, 6, 6}}, 
 {42, {1, 6, 6, 5, 5, 5, 5, 4, 5, 5, 5, 6, 5, 5, 5, 5, 6, 6}}, 
 {42, {1, 6, 6, 5, 4, 5, 5, 5, 5, 5, 5, 6, 5, 5, 5, 5, 6, 6}}, 
 {42, {4, 5, 5, 5, 6, 5, 5, 6, 6, 6, 6, 6, 5, 4, 5, 6, 5}}, 
 {42, {3, 5, 6, 6, 6, 6, 5, 4, 5, 5, 5, 5, 6, 6, 6, 6, 5}}, 
 {42, {3, 5, 6, 6, 6, 5, 5, 5, 5, 4, 5, 6, 6, 6, 6, 6, 5}}, 
 {42, {3, 5, 6, 5, 5, 6, 6, 5, 6, 6, 5, 5, 5, 5, 6, 6, 5}}, 
 {42, {3, 5, 6, 5, 5, 5, 6, 5, 5, 6, 6, 6, 6, 5, 5, 6, 5}}, 
 {42, {3, 5, 5, 6, 6, 6, 6, 6, 6, 5, 4, 5, 5, 5, 6, 6, 5}}, 
 {42, {3, 5, 5, 6, 6, 5, 6, 6, 5, 5, 6, 6, 5, 5, 5, 6, 5}}, 
 {42, {3, 5, 5, 6, 5, 5, 5, 5, 6, 6, 5, 6, 6, 6, 6, 5, 5}}, 
 {42, {3, 5, 5, 5, 6, 6, 5, 5, 6, 6, 6, 5, 6, 6, 5, 5, 5}}, 
 {42, {3, 5, 5, 5, 6, 5, 5, 6, 5, 5, 5, 6, 6, 6, 6, 6, 5}}, 
 {42, {3, 5, 5, 5, 4, 5, 6, 6, 6, 6, 6, 6, 5, 5, 6, 6, 5}}, 
 {42, {3, 5, 5, 5, 4, 5, 5, 6, 6, 6, 6, 6, 5, 6, 6, 6, 5}}, 
 {42, {1, 6, 6, 6, 6, 5, 5, 6, 6, 6, 6, 1, 6, 6, 6, 6, 6}}, 
 {42, {1, 6, 6, 6, 5, 6, 6, 5, 4, 5, 5, 6, 6, 5, 6, 6, 6}}, 
 {42, {1, 6, 6, 5, 5, 5, 4, 5, 6, 6, 6, 6, 6, 6, 5, 6, 6}}, 
 {41, {4, 5, 5, 5, 5, 4, 5, 5, 5, 6, 6, 5, 5, 4, 5, 6, 5, 5}}, 
 {41, {4, 5, 5, 5, 4, 5, 6, 5, 5, 5, 5, 5, 4, 5, 5, 6, 6, 5}}, 
 {41, {4, 5, 5, 5, 4, 5, 5, 6, 6, 5, 4, 5, 5, 5, 5, 5, 6, 5}}, 
 {41, {4, 4, 5, 5, 5, 6, 5, 5, 5, 5, 5, 5, 5, 4, 5, 6, 6, 5}}, 
 {41, {4, 4, 5, 5, 5, 5, 6, 6, 6, 5, 5, 4, 4, 5, 6, 5, 5, 5}}, 
 {41, {4, 4, 5, 5, 5, 5, 6, 5, 5, 5, 5, 4, 5, 5, 5, 6, 6, 5}}, 
 {41, {4, 4, 5, 5, 5, 5, 5, 5, 5, 6, 6, 5, 4, 5, 6, 5, 5, 5}}, 
 {41, {4, 4, 5, 5, 5, 5, 5, 5, 5, 6, 5, 5, 5, 4, 5, 6, 6, 5}}, 
 {41, {4, 4, 5, 5, 5, 5, 5, 5, 5, 5, 6, 5, 4, 5, 6, 6, 5, 5}}, 
 {41, {4, 4, 5, 5, 5, 4, 5, 6, 6, 5, 5, 5, 5, 5, 5, 6, 5, 5}}, 
 {41, {4, 4, 5, 5, 4, 5, 5, 5, 5, 6, 6, 5, 5, 6, 5, 5, 5, 5}}, 
 {41, {4, 4, 5, 4, 5, 5, 6, 6, 5, 5, 5, 5, 6, 5, 5, 5, 5, 5}}, 
 {41, {3, 5, 5, 5, 5, 5, 5, 5, 5, 5, 6, 6, 5, 5, 4, 5, 6, 5}}, 
 {41, {1, 6, 6, 6, 5, 5, 5, 4, 5, 5, 4, 5, 5, 5, 5, 6, 6, 6}}, 
 {41, {1, 6, 6, 6, 5, 4, 5, 5, 5, 5, 4, 5, 5, 5, 5, 6, 6, 6}}, 
 {41, {1, 6, 6, 5, 5, 5, 5, 5, 5, 5, 4, 5, 6, 5, 5, 5, 6, 6}}, 
 {41, {1, 6, 6, 5, 5, 5, 5, 5, 4, 5, 5, 4, 5, 5, 6, 6, 6, 6}}, 
 {41, {1, 6, 6, 5, 5, 5, 4, 5, 5, 5, 5, 6, 5, 5, 5, 5, 6, 6}}, 
 {41, {1, 6, 6, 5, 5, 5, 4, 5, 5, 5, 5, 4, 5, 5, 6, 6, 6, 6}}, 
 {41, {1, 6, 6, 1, 6, 6, 6, 5, 5, 4, 5, 5, 5, 5, 6, 6, 6, 6}}, 
 {41, {4, 5, 6, 6, 5, 5, 5, 6, 5, 5, 6, 5, 5, 5, 6, 6, 5}}, 
 {41, {4, 5, 5, 5, 6, 6, 5, 4, 5, 6, 6, 5, 5, 6, 6, 6, 5}}, 
 {41, {4, 5, 5, 4, 5, 5, 6, 5, 5, 6, 6, 5, 6, 6, 6, 6, 5}}, 
 {41, {4, 4, 5, 6, 6, 5, 5, 6, 5, 5, 6, 6, 5, 5, 6, 6, 5}}, 
 {41, {4, 4, 5, 6, 6, 5, 5, 6, 5, 5, 6, 5, 5, 6, 6, 6, 5}}, 
 {41, {4, 4, 5, 5, 6, 5, 5, 6, 6, 5, 6, 6, 6, 5, 5, 6, 5}}, 
 {41, {4, 4, 5, 5, 6, 5, 5, 4, 5, 6, 6, 6, 6, 6, 6, 6, 5}}, 
 {41, {4, 4, 5, 4, 5, 5, 6, 6, 5, 6, 6, 6, 6, 6, 6, 5, 5}}, 
 {41, {3, 5, 6, 6, 5, 4, 5, 5, 5, 5, 6, 6, 6, 6, 6, 6, 5}}, 
 {41, {3, 5, 6, 5, 5, 4, 5, 6, 6, 6, 6, 6, 6, 5, 5, 6, 5}}, 
 {41, {3, 5, 5, 5, 6, 6, 6, 6, 6, 6, 6, 5, 4, 5, 5, 6, 5}}, 
 {41, {3, 5, 5, 5, 6, 6, 6, 6, 5, 4, 5, 5, 6, 6, 6, 6, 5}}, 
 {41, {3, 5, 5, 5, 6, 6, 5, 5, 5, 6, 5, 5, 6, 6, 6, 6, 5}}, 
 {41, {3, 5, 5, 5, 4, 5, 5, 6, 6, 6, 6, 5, 6, 6, 6, 6, 5}}, 
 {41, {1, 6, 6, 1, 6, 6, 6, 6, 5, 6, 6, 5, 6, 6, 6, 6, 6}}, 
 {40, {4, 5, 5, 5, 5, 4, 5, 5, 6, 5, 5, 4, 5, 5, 6, 6, 5, 5}}, 
 {40, {4, 5, 5, 5, 4, 5, 5, 6, 5, 4, 5, 5, 5, 5, 5, 6, 6, 5}}, 
 {40, {4, 4, 5, 5, 5, 5, 5, 5, 5, 6, 5, 5, 4, 5, 6, 6, 5, 5}}, 
 {40, {4, 4, 5, 5, 5, 5, 5, 4, 5, 5, 6, 5, 5, 6, 6, 5, 5, 5}}, 
 {40, {4, 4, 5, 5, 5, 5, 5, 4, 5, 5, 5, 6, 5, 5, 5, 6, 6, 5}}, 
 {40, {4, 4, 5, 5, 5, 5, 4, 5, 6, 6, 5, 5, 5, 6, 5, 5, 5, 5}}, 
 {40, {4, 4, 5, 5, 4, 4, 5, 5, 5, 5, 6, 6, 6, 6, 5, 5, 5, 5}}, 
 {40, {4, 4, 5, 4, 5, 5, 6, 6, 5, 5, 5, 6, 5, 5, 5, 5, 5, 5}}, 
 {40, {3, 5, 5, 5, 6, 6, 5, 5, 5, 5, 5, 5, 4, 5, 5, 5, 6, 5}}, 
 {40, {3, 5, 5, 5, 6, 6, 5, 5, 4, 5, 5, 5, 5, 5, 5, 5, 6, 5}}, 
 {40, {3, 5, 5, 5, 6, 5, 5, 5, 5, 5, 5, 5, 5, 5, 6, 5, 5, 5}}, 
 {40, {3, 5, 5, 5, 6, 5, 5, 5, 5, 5, 5, 5, 5, 5, 5, 6, 5, 5}}, 
 {40, {3, 5, 5, 5, 5, 5, 5, 5, 4, 5, 5, 6, 6, 5, 5, 5, 6, 5}}, 
 {40, {3, 5, 5, 5, 5, 4, 5, 5, 5, 6, 6, 5, 5, 5, 5, 5, 6, 5}}, 
 {40, {1, 6, 6, 6, 5, 4, 5, 5, 5, 5, 5, 4, 5, 5, 5, 6, 6, 6}}, 
 {40, {1, 6, 6, 5, 5, 5, 5, 5, 4, 5, 5, 6, 5, 5, 5, 5, 6, 6}}, 
 {40, {1, 6, 6, 5, 5, 5, 4, 5, 5, 5, 5, 5, 6, 5, 5, 5, 6, 6}}, 
 {40, {1, 6, 1, 6, 6, 6, 5, 5, 5, 5, 5, 5, 5, 5, 6, 6, 6, 6}}, 
 {40, {4, 5, 5, 5, 5, 6, 6, 6, 5, 6, 6, 5, 4, 5, 6, 6, 5}}, 
 {40, {4, 4, 5, 6, 5, 5, 6, 6, 5, 5, 6, 5, 5, 6, 6, 6, 5}}, 
 {40, {4, 4, 5, 6, 5, 5, 6, 5, 5, 6, 6, 5, 5, 6, 6, 6, 5}}, 
 {40, {4, 4, 5, 6, 5, 5, 5, 5, 6, 6, 5, 6, 6, 5, 6, 6, 5}}, 
 {40, {4, 4, 5, 5, 6, 6, 5, 5, 6, 5, 5, 6, 6, 5, 6, 6, 5}}, 
 {40, {4, 4, 5, 5, 6, 5, 5, 6, 6, 5, 6, 6, 5, 5, 6, 6, 5}}, 
 {40, {4, 4, 5, 5, 5, 4, 5, 6, 6, 6, 6, 6, 6, 5, 6, 6, 5}}, 
 {40, {4, 4, 5, 5, 5, 4, 5, 6, 6, 6, 5, 6, 6, 6, 6, 6, 5}}, 
 {40, {3, 5, 6, 6, 5, 6, 6, 5, 5, 4, 5, 5, 6, 6, 6, 6, 5}}, 
 {40, {3, 5, 6, 5, 4, 5, 5, 5, 5, 6, 6, 6, 6, 6, 6, 6, 5}}, 
 {40, {3, 5, 5, 6, 6, 6, 6, 6, 6, 5, 5, 5, 4, 5, 6, 6, 5}}, 
 {40, {3, 5, 5, 6, 6, 5, 6, 6, 5, 6, 6, 5, 5, 5, 5, 6, 5}}, 
 {40, {3, 5, 5, 5, 4, 5, 6, 6, 6, 6, 6, 5, 5, 6, 6, 6, 5}}, 
 {40, {3, 5, 5, 5, 4, 5, 6, 5, 5, 6, 6, 6, 6, 6, 6, 6, 5}}, 
 {40, {3, 5, 5, 4, 5, 5, 6, 6, 6, 6, 5, 6, 6, 6, 6, 5, 5}}, 
 {39, {4, 5, 5, 5, 5, 5, 4, 5, 5, 5, 6, 5, 4, 5, 5, 6, 6, 5}}, 
 {39, {4, 5, 5, 5, 5, 4, 5, 5, 6, 5, 5, 5, 4, 5, 6, 6, 5, 5}}, 
 {39, {4, 5, 5, 5, 4, 5, 5, 5, 4, 5, 6, 5, 5, 5, 5, 6, 6, 5}}, 
 {39, {4, 5, 5, 5, 4, 5, 5, 5, 4, 5, 5, 6, 5, 5, 6, 6, 5, 5}}, 
 {39, {4, 4, 5, 5, 6, 5, 5, 5, 5, 5, 4, 5, 5, 5, 5, 6, 6, 5}}, 
 {39, {4, 4, 5, 5, 5, 5, 6, 5, 5, 5, 5, 4, 5, 6, 6, 5, 5, 5}}, 
 {39, {4, 4, 5, 5, 5, 5, 5, 6, 6, 6, 5, 5, 5, 4, 4, 5, 6, 5}}, 
 {39, {4, 4, 5, 5, 5, 5, 5, 5, 5, 5, 4, 5, 6, 5, 5, 6, 6, 5}}, 
 {39, {4, 4, 5, 5, 5, 5, 4, 4, 5, 6, 5, 5, 5, 5, 6, 6, 6, 5}}, 
 {39, {4, 4, 5, 5, 4, 5, 6, 6, 5, 5, 5, 5, 6, 5, 5, 5, 5, 5}}, 
 {39, {4, 4, 5, 5, 4, 5, 6, 6, 5, 5, 5, 5, 5, 6, 5, 5, 5, 5}}, 
 {39, {4, 4, 5, 5, 4, 5, 6, 6, 5, 5, 5, 5, 5, 5, 6, 5, 5, 5}}, 
 {39, {4, 4, 5, 5, 4, 5, 5, 5, 5, 6, 5, 5, 5, 5, 5, 6, 6, 5}}, 
 {39, {4, 4, 5, 5, 4, 5, 5, 5, 5, 5, 5, 6, 6, 5, 5, 6, 5, 5}}, 
 {39, {4, 4, 5, 5, 4, 5, 5, 5, 5, 5, 5, 5, 6, 5, 5, 6, 6, 5}}, 
 {39, {4, 4, 5, 5, 4, 5, 5, 5, 5, 5, 4, 5, 5, 6, 6, 6, 6, 5}}, 
 {39, {4, 4, 5, 4, 5, 5, 5, 6, 6, 5, 5, 5, 5, 5, 5, 5, 6, 5}}, 
 {39, {3, 5, 6, 5, 4, 5, 5, 5, 5, 5, 5, 5, 5, 5, 5, 6, 6, 5}}, 
 {39, {3, 5, 5, 5, 5, 6, 5, 5, 5, 5, 5, 5, 5, 6, 5, 5, 5, 5}}, 
 {39, {3, 5, 5, 5, 5, 6, 5, 5, 5, 5, 5, 5, 5, 5, 5, 6, 5, 5}}, 
 {39, {3, 5, 5, 5, 5, 5, 4, 5, 5, 5, 6, 6, 5, 5, 5, 5, 6, 5}}, 
 {39, {3, 5, 5, 5, 5, 4, 5, 5, 5, 6, 5, 5, 5, 5, 6, 6, 5, 5}}, 
 {39, {3, 5, 5, 5, 4, 5, 5, 5, 6, 6, 5, 5, 5, 5, 5, 5, 6, 5}}, 
 {39, {3, 5, 5, 4, 5, 5, 5, 6, 6, 5, 5, 5, 5, 5, 5, 5, 6, 5}}, 
 {39, {1, 6, 6, 6, 5, 5, 3, 5, 5, 5, 5, 5, 5, 5, 5, 6, 6, 6}}, 
 {39, {1, 6, 6, 5, 5, 5, 5, 5, 4, 5, 5, 5, 5, 6, 5, 5, 6, 6}}, 
 {39, {1, 6, 6, 5, 5, 5, 5, 4, 5, 5, 5, 4, 5, 5, 6, 6, 6, 6}}, 
 {39, {1, 6, 6, 5, 5, 4, 5, 5, 5, 4, 5, 5, 5, 5, 6, 6, 6, 6}}, 
 {39, {1, 6, 6, 5, 4, 5, 5, 5, 5, 5, 5, 4, 5, 5, 6, 6, 6, 6}}, 
 {39, {4, 5, 5, 6, 5, 5, 4, 5, 6, 6, 5, 5, 6, 6, 6, 6, 5}}, 
 {39, {4, 5, 5, 6, 5, 4, 5, 5, 6, 6, 5, 6, 6, 5, 6, 6, 5}}, 
 {39, {4, 5, 5, 5, 5, 6, 6, 6, 5, 6, 6, 6, 5, 4, 5, 6, 5}}, 
 {39, {4, 5, 4, 5, 5, 5, 5, 6, 6, 6, 6, 5, 6, 6, 6, 5, 5}}, 
 {39, {4, 4, 5, 5, 6, 5, 5, 6, 6, 6, 6, 6, 5, 5, 6, 5, 5}}, 
 {39, {4, 4, 5, 5, 5, 4, 5, 6, 6, 5, 6, 6, 6, 6, 6, 6, 5}}, 
 {39, {4, 4, 5, 4, 4, 5, 5, 6, 6, 6, 6, 6, 6, 6, 6, 6, 5}}, 
 {39, {3, 5, 6, 5, 5, 6, 6, 6, 6, 6, 5, 4, 5, 5, 6, 6, 5}}, 
 {39, {3, 5, 6, 5, 5, 5, 6, 6, 6, 5, 5, 6, 5, 5, 6, 6, 5}}, 
 {39, {3, 5, 6, 5, 5, 5, 6, 6, 5, 5, 6, 6, 5, 5, 6, 6, 5}}, 
 {39, {3, 5, 6, 5, 5, 5, 6, 6, 5, 5, 6, 5, 5, 6, 6, 6, 5}}, 
 {39, {3, 5, 6, 5, 5, 5, 6, 5, 5, 6, 6, 5, 5, 6, 6, 6, 5}}, 
 {39, {3, 5, 6, 5, 5, 5, 5, 5, 6, 6, 5, 6, 6, 5, 6, 6, 5}}, 
 {39, {3, 5, 6, 5, 5, 5, 5, 4, 5, 6, 6, 6, 6, 6, 6, 6, 5}}, 
 {39, {3, 5, 5, 6, 6, 6, 6, 6, 6, 6, 5, 5, 5, 4, 5, 6, 5}}, 
 {39, {3, 5, 5, 6, 6, 6, 5, 6, 6, 5, 5, 6, 5, 5, 5, 6, 5}}, 
 {39, {3, 5, 5, 6, 6, 5, 5, 6, 5, 5, 6, 6, 6, 5, 5, 6, 5}}, 
 {39, {3, 5, 5, 6, 5, 5, 5, 5, 6, 6, 6, 5, 5, 6, 6, 6, 5}}, 
 {39, {3, 5, 5, 5, 6, 6, 6, 5, 4, 5, 5, 6, 6, 6, 6, 6, 5}}, 
 {39, {3, 5, 5, 5, 6, 5, 5, 5, 6, 6, 6, 6, 5, 5, 6, 6, 5}}, 
 {39, {3, 5, 5, 5, 6, 5, 5, 5, 5, 6, 6, 6, 5, 6, 6, 6, 5}}, 
 {39, {3, 5, 5, 5, 5, 6, 6, 6, 5, 6, 6, 5, 6, 6, 5, 5, 5}}, 
 {39, {3, 5, 5, 4, 5, 5, 6, 6, 6, 5, 6, 6, 6, 6, 6, 5, 5}}, 
 {39, {1, 6, 6, 6, 5, 4, 4, 5, 6, 6, 5, 6, 6, 6, 6, 6, 6}}, 
 {39, {1, 6, 6, 1, 6, 6, 6, 5, 6, 6, 6, 5, 6, 6, 6, 6, 6}}, 
 {38, {4, 5, 5, 5, 5, 4, 5, 6, 5, 5, 5, 5, 4, 5, 5, 6, 6, 5}}, 
 {38, {4, 5, 5, 4, 5, 5, 5, 4, 5, 5, 6, 6, 5, 5, 5, 5, 6, 5}}, 
 {38, {4, 4, 5, 5, 5, 5, 6, 5, 5, 5, 5, 5, 4, 5, 6, 6, 5, 5}}, 
 {38, {4, 4, 5, 5, 5, 5, 5, 5, 6, 6, 5, 4, 5, 5, 5, 5, 6, 5}}, 
 {38, {4, 4, 5, 5, 5, 5, 5, 5, 4, 5, 5, 5, 6, 6, 5, 5, 6, 5}}, 
 {38, {4, 4, 5, 5, 5, 5, 5, 5, 4, 5, 5, 5, 6, 5, 5, 6, 6, 5}}, 
 {38, {4, 4, 5, 5, 5, 5, 5, 5, 4, 4, 5, 5, 6, 6, 6, 6, 5, 5}}, 
 {38, {4, 4, 5, 5, 5, 5, 5, 4, 5, 6, 6, 5, 5, 6, 5, 5, 5, 5}}, 
 {38, {4, 4, 5, 5, 5, 5, 4, 5, 6, 5, 5, 6, 6, 5, 5, 5, 5, 5}}, 
 {38, {4, 4, 5, 5, 5, 5, 4, 5, 5, 5, 5, 6, 5, 5, 5, 6, 6, 5}}, 
 {38, {4, 4, 5, 5, 5, 5, 4, 5, 5, 5, 4, 5, 5, 6, 6, 6, 6, 5}}, 
 {38, {4, 4, 5, 5, 5, 4, 5, 6, 6, 5, 5, 6, 5, 5, 5, 5, 5, 5}}, 
 {38, {4, 4, 5, 5, 4, 5, 5, 5, 5, 6, 6, 5, 5, 5, 5, 6, 5, 5}}, 
 {38, {4, 4, 5, 5, 4, 5, 5, 5, 5, 5, 6, 6, 5, 5, 5, 6, 5, 5}}, 
 {38, {3, 5, 5, 5, 6, 6, 5, 5, 5, 5, 5, 5, 5, 4, 5, 5, 6, 5}}, 
 {38, {3, 5, 5, 5, 5, 5, 5, 5, 6, 5, 5, 5, 5, 5, 5, 6, 5, 5}}, 
 {38, {3, 5, 5, 5, 5, 5, 5, 5, 5, 5, 5, 6, 5, 5, 5, 6, 5, 5}}, 
 {38, {1, 6, 6, 5, 4, 5, 5, 5, 5, 5, 5, 5, 6, 5, 5, 5, 6, 6}}, 
 {38, {1, 6, 6, 1, 6, 6, 5, 5, 6, 5, 5, 5, 5, 5, 5, 6, 6, 6}}, 
 {38, {4, 5, 6, 5, 5, 5, 5, 6, 5, 4, 5, 6, 6, 6, 6, 6, 5}}, 
 {38, {4, 5, 5, 4, 5, 6, 5, 5, 6, 6, 6, 5, 5, 6, 6, 6, 5}}, 
 {38, {4, 5, 4, 5, 5, 6, 6, 6, 6, 6, 5, 5, 5, 5, 6, 6, 5}}, 
 {38, {4, 4, 5, 6, 6, 6, 5, 4, 5, 5, 6, 6, 5, 6, 6, 6, 5}}, 
 {38, {4, 4, 5, 6, 6, 5, 5, 6, 5, 5, 5, 6, 6, 5, 6, 6, 5}}, 
 {38, {4, 4, 5, 6, 5, 5, 6, 5, 5, 5, 6, 6, 5, 6, 6, 6, 5}}, 
 {38, {4, 4, 5, 5, 6, 5, 5, 6, 6, 6, 5, 6, 6, 5, 5, 6, 5}}, 
 {38, {3, 5, 6, 5, 5, 6, 6, 5, 4, 5, 5, 6, 6, 6, 6, 6, 5}}, 
 {38, {3, 5, 6, 5, 5, 6, 5, 5, 6, 5, 5, 5, 6, 6, 6, 6, 5}}, 
 {38, {3, 5, 6, 5, 5, 5, 6, 5, 5, 6, 5, 5, 6, 6, 6, 6, 5}}, 
 {38, {3, 5, 5, 6, 5, 5, 6, 5, 5, 5, 5, 6, 6, 6, 6, 6, 5}}, 
 {38, {3, 5, 5, 6, 5, 5, 5, 6, 6, 5, 5, 6, 6, 6, 6, 5, 5}}, 
 {38, {3, 5, 5, 6, 5, 5, 5, 5, 6, 6, 6, 6, 5, 5, 6, 6, 5}}, 
 {38, {3, 5, 5, 5, 6, 6, 6, 6, 6, 5, 5, 4, 5, 6, 6, 6, 5}}, 
 {38, {3, 5, 5, 5, 4, 5, 6, 6, 6, 6, 6, 6, 5, 6, 6, 5, 5}}, 
 {38, {3, 5, 5, 4, 5, 6, 6, 6, 6, 6, 6, 5, 5, 6, 6, 5, 5}}, 
 {38, {3, 5, 5, 4, 5, 6, 6, 6, 5, 5, 5, 6, 6, 6, 6, 6, 5}}, 
 {37, {4, 5, 5, 5, 5, 5, 4, 5, 5, 5, 6, 6, 5, 4, 5, 6, 5, 5}}, 
 {37, {4, 5, 5, 5, 5, 4, 5, 5, 5, 6, 5, 4, 5, 6, 6, 5, 5, 5}}, 
 {37, {4, 5, 5, 5, 5, 4, 5, 5, 5, 5, 6, 5, 4, 5, 5, 6, 6, 5}}, 
 {37, {4, 5, 5, 5, 4, 5, 5, 5, 6, 6, 5, 5, 5, 4, 5, 6, 5, 5}}, 
 {37, {4, 5, 5, 4, 5, 5, 5, 6, 6, 5, 5, 5, 4, 5, 5, 6, 5, 5}}, 
 {37, {4, 4, 5, 5, 6, 5, 5, 5, 5, 4, 5, 5, 5, 5, 6, 6, 5, 5}}, 
 {37, {4, 4, 5, 5, 6, 5, 5, 5, 4, 5, 5, 5, 5, 5, 5, 6, 6, 5}}, 
 {37, {4, 4, 5, 5, 6, 5, 5, 4, 5, 5, 5, 5, 5, 5, 5, 6, 6, 5}}, 
 {37, {4, 4, 5, 5, 5, 6, 6, 5, 4, 5, 5, 5, 5, 5, 5, 6, 5, 5}}, 
 {37, {4, 4, 5, 5, 5, 5, 5, 6, 5, 5, 4, 5, 6, 6, 5, 5, 5, 5}}, 
 {37, {4, 4, 5, 5, 5, 5, 5, 5, 5, 4, 5, 6, 6, 5, 5, 6, 5, 5}}, 
 {37, {4, 4, 5, 5, 5, 5, 5, 5, 4, 5, 6, 6, 5, 5, 5, 5, 6, 5}}, 
 {37, {4, 4, 5, 5, 5, 5, 4, 5, 5, 4, 5, 5, 6, 6, 6, 6, 5, 5}}, 
 {37, {4, 4, 5, 5, 5, 4, 5, 5, 4, 5, 6, 6, 6, 6, 5, 5, 5, 5}}, 
 {37, {4, 4, 5, 5, 5, 4, 5, 5, 4, 5, 5, 5, 6, 6, 6, 6, 5, 5}}, 
 {37, {4, 4, 5, 5, 4, 4, 5, 5, 6, 6, 6, 5, 5, 5, 5, 5, 6, 5}}, 
 {37, {4, 4, 5, 4, 5, 5, 6, 6, 5, 5, 5, 5, 5, 5, 6, 5, 5, 5}}, 
 {37, {4, 4, 5, 4, 5, 5, 5, 5, 5, 6, 6, 5, 5, 5, 5, 5, 6, 5}}, 
 {37, {4, 4, 5, 4, 5, 5, 5, 5, 5, 5, 5, 5, 6, 6, 5, 5, 6, 5}}, 
 {37, {3, 5, 6, 5, 5, 5, 5, 5, 5, 5, 5, 5, 5, 4, 5, 6, 6, 5}}, 
 {37, {3, 5, 6, 5, 5, 5, 5, 4, 5, 5, 6, 5, 5, 5, 5, 5, 6, 5}}, 
 {37, {3, 5, 5, 5, 5, 6, 6, 5, 5, 5, 5, 4, 5, 5, 5, 5, 6, 5}}, 
 {37, {3, 5, 5, 5, 5, 5, 6, 5, 5, 5, 5, 4, 5, 5, 6, 6, 5, 5}}, 
 {37, {3, 5, 5, 5, 5, 5, 5, 6, 5, 5, 5, 4, 5, 5, 6, 6, 5, 5}}, 
 {37, {3, 5, 5, 5, 5, 5, 5, 5, 6, 5, 4, 5, 5, 5, 6, 6, 5, 5}}, 
 {37, {3, 5, 5, 5, 5, 5, 5, 4, 5, 6, 5, 5, 5, 5, 5, 6, 6, 5}}, 
 {37, {3, 5, 5, 5, 5, 4, 5, 5, 5, 5, 6, 6, 5, 5, 5, 5, 6, 5}}, 
 {37, {3, 5, 5, 4, 4, 5, 5, 5, 5, 6, 6, 6, 6, 5, 5, 5, 5, 5}}, 
 {37, {1, 6, 6, 5, 5, 4, 5, 5, 5, 5, 6, 6, 6, 1, 6, 6, 6, 6}}, 
 {37, {4, 5, 5, 4, 5, 5, 5, 6, 6, 5, 6, 6, 6, 5, 6, 6, 5}}, 
 {37, {4, 5, 4, 5, 5, 5, 6, 6, 5, 6, 6, 6, 6, 6, 5, 5, 5}}, 
 {37, {4, 4, 5, 4, 5, 6, 6, 6, 6, 6, 6, 6, 5, 5, 5, 6, 5}}, 
 {37, {3, 5, 6, 6, 5, 5, 6, 5, 5, 5, 6, 6, 5, 5, 6, 6, 5}}, 
 {37, {3, 5, 6, 5, 5, 6, 5, 5, 5, 6, 6, 5, 5, 6, 6, 6, 5}}, 
 {37, {3, 5, 5, 6, 6, 6, 5, 5, 6, 6, 5, 5, 6, 5, 5, 6, 5}}, 
 {37, {3, 5, 5, 6, 6, 5, 5, 6, 6, 5, 5, 6, 5, 5, 6, 6, 5}}, 
 {37, {3, 5, 5, 6, 5, 5, 6, 5, 5, 5, 6, 6, 6, 6, 6, 5, 5}}, 
 {37, {3, 5, 5, 6, 5, 5, 5, 6, 6, 6, 6, 5, 5, 5, 6, 6, 5}}, 
 {37, {3, 5, 5, 5, 6, 6, 6, 6, 6, 6, 5, 4, 5, 5, 6, 6, 5}}, 
 {37, {3, 5, 5, 5, 6, 6, 5, 5, 4, 5, 6, 6, 6, 6, 6, 6, 5}}, 
 {37, {3, 5, 5, 5, 5, 6, 5, 5, 6, 6, 6, 6, 5, 6, 6, 5, 5}}, 
 {37, {3, 5, 5, 4, 5, 6, 6, 6, 6, 5, 5, 6, 6, 6, 6, 5, 5}}, 
 {37, {3, 5, 5, 4, 5, 6, 6, 5, 5, 5, 6, 6, 6, 6, 6, 6, 5}}, 
 {37, {1, 6, 6, 6, 1, 6, 6, 6, 5, 6, 6, 6, 5, 6, 6, 6, 6}}, 
 {37, {1, 6, 6, 5, 6, 6, 5, 6, 6, 6, 5, 4, 5, 5, 6, 6, 6}}, 
 {37, {1, 6, 6, 5, 4, 5, 6, 6, 6, 6, 5, 5, 6, 6, 5, 6, 6}}, 
 {36, {4, 5, 5, 5, 5, 5, 4, 5, 5, 6, 5, 5, 4, 5, 6, 6, 5, 5}}, 
 {36, {4, 5, 5, 5, 4, 5, 5, 6, 5, 5, 5, 5, 4, 5, 5, 6, 6, 5}}, 
 {36, {4, 5, 5, 5, 4, 5, 5, 6, 5, 5, 4, 5, 5, 5, 6, 6, 5, 5}}, 
 {36, {4, 5, 5, 5, 4, 5, 5, 5, 6, 5, 5, 4, 5, 5, 5, 6, 6, 5}}, 
 {36, {4, 5, 5, 4, 5, 5, 6, 5, 5, 5, 5, 5, 4, 5, 6, 6, 5, 5}}, 
 {36, {4, 5, 5, 4, 5, 5, 5, 5, 6, 5, 4, 5, 5, 5, 6, 6, 5, 5}}, 
 {36, {4, 5, 5, 4, 5, 5, 5, 5, 5, 6, 5, 5, 6, 5, 5, 5, 5, 5}}, 
 {36, {4, 5, 5, 4, 5, 5, 5, 5, 4, 5, 6, 5, 5, 5, 6, 6, 5, 5}}, 
 {36, {4, 4, 5, 5, 5, 6, 5, 5, 5, 5, 4, 5, 5, 5, 5, 6, 6, 5}}, 
 {36, {4, 4, 5, 5, 5, 5, 6, 6, 5, 5, 4, 4, 5, 5, 5, 6, 6, 5}}, 
 {36, {4, 4, 5, 5, 5, 5, 6, 6, 5, 4, 5, 5, 5, 5, 6, 5, 5, 5}}, 
 {36, {4, 4, 5, 5, 5, 5, 5, 6, 6, 6, 5, 5, 4, 4, 5, 6, 5, 5}}, 
 {36, {4, 4, 5, 5, 5, 5, 5, 6, 6, 6, 5, 5, 4, 4, 5, 5, 6, 5}}, 
 {36, {4, 4, 5, 5, 5, 5, 5, 6, 6, 5, 4, 5, 5, 5, 5, 6, 5, 5}}, 
 {36, {4, 4, 5, 5, 5, 5, 5, 6, 5, 4, 4, 5, 5, 6, 6, 6, 5, 5}}, 
 {36, {4, 4, 5, 5, 5, 5, 5, 5, 5, 5, 5, 6, 6, 5, 4, 5, 6, 5}}, 
 {36, {4, 4, 5, 4, 5, 5, 5, 6, 5, 5, 5, 6, 6, 5, 5, 5, 5, 5}}, 
 {36, {4, 4, 5, 4, 5, 5, 5, 5, 5, 5, 5, 6, 5, 5, 6, 6, 5, 5}}, 
 {36, {3, 5, 5, 5, 5, 5, 5, 5, 6, 6, 5, 4, 5, 5, 5, 5, 6, 5}}, 
 {36, {3, 5, 5, 5, 5, 5, 5, 5, 4, 5, 5, 5, 6, 6, 5, 5, 6, 5}}, 
 {36, {3, 5, 5, 5, 5, 5, 4, 5, 6, 6, 6, 5, 5, 5, 5, 5, 5, 5}}, 
 {36, {3, 5, 5, 5, 5, 4, 5, 5, 5, 6, 5, 5, 5, 5, 5, 6, 6, 5}}, 
 {36, {3, 5, 5, 5, 4, 5, 5, 6, 6, 5, 5, 5, 5, 6, 5, 5, 5, 5}}, 
 {36, {3, 5, 5, 5, 4, 5, 5, 6, 5, 5, 5, 5, 5, 5, 5, 6, 6, 5}}, 
 {36, {1, 6, 6, 6, 5, 4, 5, 5, 4, 5, 5, 5, 5, 5, 5, 6, 6, 6}}, 
 {36, {1, 6, 6, 5, 5, 5, 5, 5, 5, 5, 5, 6, 6, 1, 6, 6, 6, 6}}, 
 {36, {1, 6, 6, 5, 5, 5, 5, 5, 5, 5, 5, 4, 5, 6, 5, 5, 6, 6}}, 
 {36, {1, 6, 6, 5, 5, 5, 5, 5, 5, 5, 5, 3, 5, 5, 6, 6, 6, 6}}, 
 {36, {1, 6, 6, 5, 5, 4, 5, 5, 5, 5, 5, 5, 4, 5, 6, 6, 6, 6}}, 
 {36, {1, 6, 6, 1, 6, 6, 5, 5, 5, 5, 5, 6, 5, 5, 5, 6, 6, 6}}, 
 {36, {4, 5, 5, 4, 5, 6, 5, 5, 5, 6, 6, 5, 6, 6, 6, 6, 5}}, 
 {36, {4, 5, 5, 4, 5, 5, 5, 6, 6, 5, 6, 6, 5, 6, 6, 6, 5}}, 
 {36, {4, 5, 4, 5, 5, 5, 6, 6, 6, 6, 6, 6, 5, 5, 5, 6, 5}}, 
 {36, {4, 4, 5, 6, 5, 5, 5, 6, 5, 5, 6, 6, 5, 6, 6, 6, 5}}, 
 {36, {4, 4, 5, 5, 5, 6, 6, 5, 6, 6, 5, 6, 6, 5, 5, 6, 5}}, 
 {36, {4, 4, 5, 5, 5, 4, 5, 6, 6, 6, 6, 6, 5, 6, 6, 6, 5}}, 
 {36, {4, 4, 5, 4, 5, 5, 5, 6, 6, 6, 6, 6, 5, 6, 6, 6, 5}}, 
 {36, {3, 5, 6, 5, 5, 6, 6, 6, 5, 5, 5, 6, 5, 5, 6, 6, 5}}, 
 {36, {3, 5, 6, 5, 5, 6, 5, 5, 5, 5, 6, 6, 5, 6, 6, 6, 5}}, 
 {36, {3, 5, 6, 5, 5, 5, 5, 6, 6, 5, 5, 6, 6, 5, 6, 6, 5}}, 
 {36, {3, 5, 6, 5, 5, 4, 5, 5, 6, 6, 5, 6, 6, 6, 6, 6, 5}}, 
 {36, {3, 5, 5, 6, 6, 5, 5, 6, 6, 6, 5, 5, 6, 5, 5, 6, 5}}, 
 {36, {3, 5, 5, 6, 6, 5, 5, 5, 6, 6, 5, 6, 6, 5, 5, 6, 5}}, 
 {36, {3, 5, 5, 6, 6, 5, 4, 5, 5, 5, 6, 6, 6, 6, 6, 6, 5}}, 
 {36, {3, 5, 5, 6, 5, 5, 5, 5, 6, 6, 5, 5, 6, 6, 6, 6, 5}}, 
 {36, {3, 5, 5, 6, 5, 5, 5, 5, 5, 6, 6, 6, 5, 6, 6, 6, 5}}, 
 {36, {3, 5, 5, 5, 5, 6, 6, 6, 6, 6, 6, 5, 4, 5, 6, 6, 5}}, 
 {36, {3, 5, 5, 5, 5, 6, 5, 5, 6, 6, 6, 6, 5, 5, 6, 6, 5}}, 
 {36, {3, 5, 5, 4, 5, 5, 6, 6, 6, 6, 5, 5, 6, 6, 6, 6, 5}}, 
 {36, {3, 5, 5, 4, 5, 5, 5, 6, 6, 6, 6, 5, 6, 6, 6, 6, 5}}, 
 {36, {1, 6, 6, 5, 5, 5, 6, 5, 5, 6, 6, 5, 6, 6, 5, 6, 6}}, 
 {36, {1, 6, 6, 1, 6, 6, 5, 6, 6, 6, 6, 5, 6, 6, 6, 6, 6}}, 
 {35, {4, 5, 5, 5, 5, 5, 5, 4, 5, 5, 6, 5, 4, 5, 5, 6, 6, 5}}, 
 {35, {4, 5, 5, 5, 5, 4, 5, 5, 6, 5, 4, 5, 5, 5, 6, 6, 5, 5}}, 
 {35, {4, 5, 5, 5, 4, 5, 5, 5, 6, 5, 5, 5, 4, 5, 5, 6, 6, 5}}, 
 {35, {4, 5, 5, 5, 4, 5, 5, 5, 5, 4, 5, 5, 6, 6, 5, 5, 6, 5}}, 
 {35, {4, 5, 5, 4, 5, 5, 5, 5, 6, 5, 4, 5, 5, 6, 6, 5, 5, 5}}, 
 {35, {4, 5, 5, 4, 5, 5, 5, 4, 5, 6, 5, 5, 5, 6, 6, 5, 5, 5}}, 
 {35, {4, 5, 5, 4, 5, 5, 5, 4, 5, 5, 6, 6, 5, 5, 6, 5, 5, 5}}, 
 {35, {4, 4, 5, 6, 5, 5, 5, 5, 5, 4, 5, 6, 5, 5, 5, 5, 6, 5}}, 
 {35, {4, 4, 5, 5, 5, 5, 5, 5, 6, 6, 6, 5, 4, 4, 5, 6, 5, 5}}, 
 {35, {4, 4, 5, 5, 5, 5, 5, 5, 4, 5, 6, 6, 5, 5, 5, 6, 5, 5}}, 
 {35, {4, 4, 5, 5, 5, 5, 4, 5, 6, 6, 5, 5, 6, 5, 5, 5, 5, 5}}, 
 {35, {4, 4, 5, 5, 5, 4, 5, 5, 5, 5, 4, 5, 6, 6, 6, 6, 5, 5}}, 
 {35, {4, 4, 5, 5, 5, 4, 5, 5, 5, 5, 4, 5, 5, 6, 6, 6, 6, 5}}, 
 {35, {4, 4, 5, 4, 5, 5, 6, 6, 5, 5, 5, 5, 5, 5, 5, 6, 5, 5}}, 
 {35, {4, 4, 5, 4, 5, 5, 5, 6, 5, 5, 5, 5, 5, 5, 5, 6, 6, 5}}, 
 {35, {4, 4, 5, 4, 5, 5, 5, 5, 6, 6, 5, 5, 5, 5, 5, 5, 6, 5}}, 
 {35, {3, 5, 6, 5, 5, 5, 5, 5, 4, 5, 6, 5, 5, 5, 5, 5, 6, 5}}, 
 {35, {3, 5, 6, 5, 5, 4, 5, 5, 5, 5, 6, 5, 5, 5, 5, 5, 6, 5}}, 
 {35, {3, 5, 5, 5, 6, 6, 5, 5, 5, 5, 4, 5, 5, 5, 5, 5, 6, 5}}, 
 {35, {3, 5, 5, 5, 5, 5, 6, 5, 5, 5, 5, 5, 4, 5, 5, 6, 6, 5}}, 
 {35, {3, 5, 5, 5, 5, 5, 5, 4, 5, 6, 5, 5, 6, 6, 5, 5, 5, 5}}, 
 {35, {3, 5, 5, 5, 4, 5, 6, 5, 5, 5, 5, 5, 5, 5, 5, 6, 6, 5}}, 
 {35, {1, 6, 6, 6, 5, 5, 4, 5, 5, 5, 5, 5, 4, 5, 5, 6, 6, 6}}, 
 {35, {1, 6, 6, 5, 5, 5, 4, 5, 5, 5, 6, 5, 5, 5, 5, 5, 6, 6}}, 
 {35, {4, 5, 5, 6, 6, 5, 6, 6, 5, 4, 5, 6, 6, 5, 5, 6, 5}}, 
 {35, {4, 5, 5, 6, 5, 4, 5, 6, 6, 5, 5, 6, 6, 5, 6, 6, 5}}, 
 {35, {4, 5, 5, 5, 5, 6, 5, 4, 5, 6, 6, 6, 5, 6, 6, 6, 5}}, 
 {35, {4, 4, 5, 4, 5, 5, 6, 5, 5, 6, 6, 6, 6, 6, 6, 6, 5}}, 
 {35, {4, 4, 5, 4, 5, 5, 5, 6, 6, 5, 6, 6, 6, 6, 6, 6, 5}}, 
 {35, {3, 5, 6, 6, 5, 6, 6, 6, 5, 5, 4, 5, 5, 6, 6, 6, 5}}, 
 {35, {3, 5, 6, 6, 5, 5, 4, 5, 5, 6, 6, 6, 5, 6, 6, 6, 5}}, 
 {35, {3, 5, 6, 5, 5, 6, 6, 6, 5, 4, 5, 5, 6, 6, 6, 6, 5}}, 
 {35, {3, 5, 6, 5, 5, 6, 6, 5, 5, 4, 5, 6, 6, 6, 6, 6, 5}}, 
 {35, {3, 5, 6, 5, 5, 4, 5, 6, 6, 6, 6, 6, 5, 5, 6, 6, 5}}, 
 {35, {3, 5, 6, 5, 5, 4, 5, 6, 6, 6, 6, 5, 5, 6, 6, 6, 5}}, 
 {35, {3, 5, 6, 5, 5, 4, 5, 5, 6, 6, 6, 6, 5, 6, 6, 6, 5}}, 
 {35, {3, 5, 5, 6, 5, 5, 5, 5, 5, 6, 6, 6, 6, 5, 6, 6, 5}}, 
 {35, {3, 5, 5, 5, 6, 6, 6, 5, 5, 4, 5, 6, 6, 6, 6, 6, 5}}, 
 {35, {3, 5, 5, 5, 6, 6, 6, 5, 4, 5, 6, 6, 6, 6, 6, 5, 5}}, 
 {35, {3, 5, 5, 5, 5, 6, 6, 5, 6, 6, 6, 5, 6, 6, 5, 5, 5}}, 
 {35, {3, 5, 5, 5, 5, 5, 6, 5, 5, 6, 6, 6, 6, 5, 6, 6, 5}}, 
 {35, {3, 5, 5, 5, 5, 5, 6, 5, 5, 6, 6, 6, 5, 6, 6, 6, 5}}, 
 {35, {3, 5, 5, 5, 4, 5, 6, 6, 6, 5, 5, 6, 6, 6, 6, 6, 5}}, 
 {35, {1, 6, 6, 5, 6, 6, 5, 6, 6, 6, 5, 5, 4, 5, 6, 6, 6}}, 
 {34, {4, 5, 5, 5, 5, 5, 5, 5, 5, 4, 5, 5, 6, 5, 5, 6, 5, 5}}, 
 {34, {4, 5, 5, 5, 5, 5, 5, 4, 5, 5, 6, 5, 4, 5, 6, 6, 5, 5}}, 
 {34, {4, 5, 5, 5, 5, 4, 5, 5, 5, 6, 6, 5, 4, 5, 5, 5, 6, 5}}, 
 {34, {4, 5, 5, 5, 5, 4, 5, 5, 5, 6, 5, 5, 4, 5, 6, 6, 5, 5}}, 
 {34, {4, 5, 5, 5, 4, 5, 5, 5, 6, 5, 5, 5, 4, 5, 6, 6, 5, 5}}, 
 {34, {4, 5, 5, 5, 4, 5, 5, 5, 5, 5, 6, 6, 5, 4, 5, 6, 5, 5}}, 
 {34, {4, 5, 5, 4, 5, 5, 5, 6, 5, 5, 5, 5, 4, 5, 6, 6, 5, 5}}, 
 {34, {4, 5, 5, 4, 5, 5, 5, 5, 5, 6, 6, 5, 5, 4, 5, 6, 5, 5}}, 
 {34, {4, 5, 5, 4, 5, 5, 5, 5, 5, 6, 6, 5, 5, 4, 5, 5, 6, 5}}, 
 {34, {4, 4, 5, 6, 5, 5, 4, 5, 6, 5, 5, 5, 5, 5, 5, 5, 6, 5}}, 
 {34, {4, 4, 5, 5, 5, 5, 6, 6, 6, 5, 5, 4, 4, 5, 5, 5, 6, 5}}, 
 {34, {4, 4, 5, 5, 5, 5, 5, 5, 5, 5, 5, 6, 5, 4, 5, 6, 6, 5}}, 
 {34, {4, 4, 5, 5, 5, 5, 4, 5, 5, 5, 5, 6, 6, 5, 5, 6, 5, 5}}, 
 {34, {4, 4, 5, 5, 4, 5, 6, 6, 5, 5, 6, 5, 5, 5, 5, 5, 5, 5}}, 
 {34, {4, 4, 5, 5, 4, 5, 6, 5, 5, 5, 5, 6, 5, 5, 5, 5, 6, 5}}, 
 {34, {3, 5, 5, 5, 5, 5, 6, 6, 5, 5, 5, 4, 5, 5, 5, 5, 6, 5}}, 
 {34, {3, 5, 5, 5, 5, 5, 5, 5, 4, 4, 5, 5, 5, 6, 6, 6, 6, 5}}, 
 {34, {3, 5, 5, 5, 5, 4, 5, 5, 5, 5, 5, 5, 6, 6, 5, 5, 6, 5}}, 
 {34, {3, 5, 5, 5, 4, 5, 5, 6, 6, 5, 5, 5, 6, 5, 5, 5, 5, 5}}, 
 {34, {3, 5, 5, 5, 4, 5, 5, 5, 4, 5, 5, 5, 6, 6, 6, 6, 5, 5}}, 
 {34, {3, 5, 5, 5, 4, 4, 5, 5, 5, 5, 5, 5, 5, 6, 6, 6, 6, 5}}, 
 {34, {3, 5, 5, 4, 4, 5, 5, 5, 5, 5, 5, 5, 6, 6, 6, 6, 5, 5}}, 
 {34, {1, 6, 6, 6, 5, 5, 5, 4, 5, 5, 5, 4, 5, 5, 5, 6, 6, 6}}, 
 {34, {1, 6, 6, 5, 5, 5, 5, 6, 5, 4, 4, 5, 5, 5, 5, 6, 6, 6}}, 
 {34, {4, 5, 4, 5, 5, 6, 5, 5, 5, 6, 6, 6, 6, 6, 6, 5, 5}}, 
 {34, {4, 5, 4, 5, 5, 5, 6, 6, 5, 5, 5, 6, 6, 6, 6, 6, 5}}, 
 {34, {4, 5, 4, 5, 5, 5, 5, 6, 6, 6, 5, 6, 6, 6, 6, 5, 5}}, 
 {34, {4, 4, 5, 5, 6, 5, 5, 6, 6, 5, 6, 6, 5, 6, 6, 5, 5}}, 
 {34, {4, 4, 5, 4, 5, 6, 6, 6, 6, 6, 6, 5, 5, 6, 6, 5, 5}}, 
 {34, {4, 4, 5, 4, 5, 6, 5, 5, 6, 6, 6, 6, 6, 6, 6, 5, 5}}, 
 {34, {3, 5, 6, 6, 5, 5, 5, 6, 5, 5, 5, 6, 6, 5, 6, 6, 5}}, 
 {34, {3, 5, 6, 5, 5, 6, 6, 6, 6, 5, 5, 4, 5, 6, 6, 6, 5}}, 
 {34, {3, 5, 6, 5, 5, 6, 6, 6, 6, 5, 4, 5, 5, 6, 6, 6, 5}}, 
 {34, {3, 5, 6, 5, 5, 5, 6, 6, 5, 6, 6, 5, 5, 5, 6, 6, 5}}, 
 {34, {3, 5, 6, 5, 5, 5, 6, 6, 5, 5, 5, 6, 6, 5, 6, 6, 5}}, 
 {34, {3, 5, 6, 5, 5, 5, 5, 6, 5, 5, 6, 6, 6, 5, 6, 6, 5}}, 
 {34, {3, 5, 6, 5, 4, 5, 5, 6, 6, 5, 5, 6, 6, 6, 6, 6, 5}}, 
 {34, {3, 5, 5, 6, 5, 5, 6, 6, 6, 6, 5, 5, 5, 6, 6, 5, 5}}, 
 {34, {3, 5, 5, 6, 5, 5, 6, 6, 6, 6, 5, 5, 5, 5, 6, 6, 5}}, 
 {34, {3, 5, 5, 6, 5, 5, 5, 5, 5, 6, 6, 5, 6, 6, 6, 6, 5}}, 
 {34, {3, 5, 5, 5, 6, 6, 6, 6, 6, 6, 6, 5, 5, 4, 5, 6, 5}}, 
 {34, {3, 5, 5, 5, 6, 6, 6, 6, 6, 5, 4, 5, 5, 6, 6, 6, 5}}, 
 {34, {3, 5, 5, 5, 5, 6, 6, 6, 5, 4, 5, 6, 6, 6, 6, 6, 5}}, 
 {34, {3, 5, 5, 5, 5, 6, 6, 5, 6, 6, 5, 6, 6, 6, 5, 5, 5}}, 
 {34, {3, 5, 5, 5, 5, 6, 5, 5, 6, 5, 5, 6, 6, 6, 6, 6, 5}}, 
 {34, {3, 5, 5, 5, 5, 5, 6, 5, 5, 6, 6, 5, 6, 6, 6, 6, 5}}, 
 {34, {3, 5, 5, 4, 5, 6, 6, 6, 6, 6, 6, 5, 5, 5, 6, 6, 5}}, 
 {34, {3, 5, 5, 4, 5, 6, 6, 6, 6, 6, 5, 5, 6, 6, 6, 5, 5}}, 
 {34, {3, 5, 5, 4, 5, 6, 6, 5, 5, 6, 6, 6, 6, 6, 6, 5, 5}}, 
 {34, {3, 5, 5, 4, 5, 5, 5, 6, 6, 6, 6, 6, 5, 6, 6, 6, 5}}, 
 {34, {1, 6, 6, 6, 6, 6, 5, 5, 5, 3, 5, 6, 6, 6, 6, 6, 6}}, 
 {34, {1, 6, 6, 6, 6, 5, 3, 5, 5, 5, 6, 6, 6, 6, 6, 6, 6}}, 
 {34, {1, 6, 6, 6, 5, 5, 5, 3, 5, 6, 6, 6, 6, 6, 6, 6, 6}}, 
 {34, {1, 6, 6, 6, 5, 4, 5, 6, 6, 5, 6, 6, 5, 5, 6, 6, 6}}, 
 {34, {1, 6, 6, 5, 6, 6, 6, 6, 1, 6, 6, 6, 6, 5, 6, 6, 6}}, 
 {34, {1, 6, 6, 5, 5, 5, 4, 5, 6, 6, 6, 5, 6, 6, 6, 6, 6}}, 
 {34, {1, 6, 6, 1, 6, 6, 6, 5, 6, 6, 6, 6, 5, 6, 6, 6, 6}}, 
 {33, {4, 5, 5, 5, 5, 4, 5, 5, 5, 5, 6, 6, 5, 4, 5, 5, 6, 5}}, 
 {33, {4, 5, 5, 5, 4, 5, 5, 5, 5, 6, 5, 5, 4, 5, 5, 6, 6, 5}}, 
 {33, {4, 5, 5, 5, 4, 5, 5, 5, 5, 6, 5, 4, 5, 5, 5, 6, 6, 5}}, 
 {33, {4, 5, 5, 4, 5, 5, 5, 6, 5, 5, 4, 5, 5, 5, 6, 6, 5, 5}}, 
 {33, {4, 5, 5, 4, 5, 5, 5, 5, 6, 6, 5, 5, 5, 4, 5, 6, 5, 5}}, 
 {33, {4, 5, 5, 4, 5, 5, 5, 5, 6, 5, 5, 4, 5, 5, 6, 6, 5, 5}}, 
 {33, {4, 5, 5, 4, 5, 5, 5, 5, 5, 5, 6, 5, 4, 5, 5, 6, 6, 5}}, 
 {33, {4, 5, 5, 4, 5, 5, 5, 5, 5, 4, 5, 6, 5, 5, 6, 6, 5, 5}}, 
 {33, {4, 5, 5, 4, 5, 5, 4, 5, 5, 5, 6, 5, 5, 5, 6, 6, 5, 5}}, 
 {33, {4, 4, 5, 6, 5, 5, 5, 5, 4, 5, 5, 6, 5, 5, 5, 5, 6, 5}}, 
 {33, {4, 4, 5, 6, 5, 5, 4, 5, 5, 5, 6, 5, 5, 5, 5, 5, 6, 5}}, 
 {33, {4, 4, 5, 5, 6, 5, 5, 5, 5, 5, 5, 5, 4, 5, 6, 6, 5, 5}}, 
 {33, {4, 4, 5, 5, 5, 6, 5, 5, 5, 5, 5, 4, 5, 6, 6, 5, 5, 5}}, 
 {33, {4, 4, 5, 5, 5, 5, 6, 5, 5, 5, 5, 6, 5, 4, 5, 5, 6, 5}}, 
 {33, {4, 4, 5, 5, 5, 5, 5, 5, 4, 5, 6, 6, 5, 5, 6, 5, 5, 5}}, 
 {33, {4, 4, 5, 5, 5, 5, 4, 4, 5, 5, 6, 6, 6, 5, 5, 5, 6, 5}}, 
 {33, {4, 4, 5, 5, 5, 5, 4, 4, 5, 5, 5, 6, 6, 6, 6, 5, 5, 5}}, 
 {33, {4, 4, 5, 5, 5, 4, 5, 5, 6, 6, 6, 6, 5, 5, 4, 5, 5, 5}}, 
 {33, {4, 4, 5, 5, 5, 4, 5, 5, 5, 5, 5, 5, 6, 5, 5, 6, 6, 5}}, 
 {33, {4, 4, 5, 4, 5, 5, 6, 5, 5, 5, 5, 6, 6, 5, 5, 5, 5, 5}}, 
 {33, {4, 4, 5, 4, 5, 5, 6, 5, 5, 5, 5, 5, 5, 5, 6, 6, 5, 5}}, 
 {33, {4, 4, 5, 4, 5, 5, 5, 5, 5, 6, 6, 5, 5, 5, 5, 6, 5, 5}}, 
 {33, {3, 5, 5, 6, 6, 5, 4, 5, 5, 5, 5, 5, 5, 5, 5, 5, 6, 5}}, 
 {33, {3, 5, 5, 5, 6, 5, 5, 5, 5, 5, 5, 4, 5, 5, 6, 6, 5, 5}}, 
 {33, {3, 5, 5, 5, 6, 5, 4, 5, 5, 5, 5, 5, 5, 5, 6, 6, 5, 5}}, 
 {33, {3, 5, 5, 5, 5, 6, 5, 4, 5, 5, 5, 5, 5, 5, 6, 6, 5, 5}}, 
 {33, {3, 5, 5, 5, 5, 5, 6, 5, 5, 5, 5, 5, 6, 5, 5, 5, 5, 5}}, 
 {33, {3, 5, 5, 5, 5, 5, 5, 4, 5, 6, 5, 5, 5, 5, 6, 6, 5, 5}}, 
 {33, {3, 5, 5, 5, 5, 5, 4, 5, 5, 5, 5, 5, 6, 6, 5, 5, 6, 5}}, 
 {33, {3, 5, 5, 5, 4, 5, 6, 5, 5, 5, 5, 6, 6, 5, 5, 5, 5, 5}}, 
 {33, {3, 5, 5, 5, 4, 5, 5, 6, 6, 6, 6, 5, 5, 4, 5, 5, 5, 5}}, 
 {33, {3, 5, 5, 5, 4, 5, 5, 5, 6, 5, 5, 5, 5, 5, 6, 6, 5, 5}}, 
 {33, {3, 5, 5, 5, 4, 5, 5, 5, 5, 5, 6, 5, 5, 5, 6, 6, 5, 5}}, 
 {33, {3, 5, 5, 4, 5, 5, 6, 6, 5, 5, 5, 5, 5, 5, 6, 5, 5, 5}}, 
 {33, {3, 5, 5, 4, 5, 5, 6, 5, 5, 5, 5, 5, 5, 5, 5, 6, 6, 5}}, 
 {33, {3, 5, 5, 4, 4, 5, 5, 5, 5, 5, 6, 6, 6, 6, 5, 5, 5, 5}}, 
 {33, {1, 6, 6, 5, 5, 5, 5, 6, 5, 5, 5, 4, 4, 5, 5, 6, 6, 6}}, 
 {33, {1, 6, 6, 1, 6, 6, 5, 5, 4, 5, 5, 5, 5, 6, 6, 6, 6, 6}}, 
 {33, {4, 5, 6, 5, 5, 6, 5, 5, 6, 5, 5, 6, 6, 5, 5, 6, 5}}, 
 {33, {4, 5, 5, 5, 5, 6, 6, 5, 6, 6, 6, 6, 5, 4, 5, 6, 5}}, 
 {33, {4, 5, 5, 4, 5, 6, 5, 5, 5, 6, 6, 6, 5, 6, 6, 6, 5}}, 
 {33, {4, 5, 4, 5, 5, 5, 6, 6, 6, 5, 6, 6, 6, 6, 5, 5, 5}}, 
 {33, {4, 4, 5, 6, 5, 4, 5, 5, 6, 6, 6, 6, 5, 6, 6, 6, 5}}, 
 {33, {4, 4, 5, 5, 6, 5, 5, 6, 6, 5, 5, 6, 6, 5, 6, 6, 5}}, 
 {33, {4, 4, 5, 4, 5, 6, 6, 6, 6, 6, 5, 5, 6, 6, 6, 5, 5}}, 
 {33, {4, 4, 5, 4, 5, 6, 6, 6, 6, 5, 5, 6, 6, 6, 6, 5, 5}}, 
 {33, {3, 5, 6, 6, 6, 5, 5, 4, 5, 5, 6, 6, 5, 6, 6, 6, 5}}, 
 {33, {3, 5, 6, 5, 4, 5, 5, 6, 6, 6, 6, 6, 5, 5, 6, 6, 5}}, 
 {33, {3, 5, 5, 6, 6, 5, 6, 6, 6, 5, 5, 5, 6, 5, 5, 6, 5}}, 
 {33, {3, 5, 5, 6, 5, 5, 5, 6, 5, 5, 6, 6, 6, 6, 6, 5, 5}}, 
 {33, {3, 5, 5, 5, 6, 5, 5, 6, 6, 6, 5, 6, 6, 6, 5, 5, 5}}, 
 {33, {1, 6, 6, 6, 5, 3, 5, 6, 6, 5, 5, 6, 6, 6, 6, 6, 6}}, 
 {33, {1, 6, 6, 5, 6, 6, 6, 5, 6, 6, 5, 5, 4, 5, 6, 6, 6}}, 
 {33, {1, 6, 6, 5, 6, 6, 5, 6, 6, 5, 4, 5, 5, 6, 6, 6, 6}}, 
 {33, {1, 6, 6, 5, 6, 6, 5, 5, 5, 6, 5, 5, 6, 6, 5, 6, 6}}, 
 {33, {1, 6, 6, 5, 6, 6, 5, 5, 4, 5, 6, 6, 5, 6, 6, 6, 6}}, 
 {33, {1, 6, 6, 5, 5, 6, 5, 5, 6, 6, 5, 6, 6, 5, 5, 6, 6}}, 
 {33, {1, 6, 6, 5, 4, 5, 6, 6, 6, 5, 5, 6, 6, 5, 6, 6, 6}}, 
 {33, {1, 6, 6, 5, 4, 5, 5, 6, 6, 6, 5, 6, 6, 6, 5, 6, 6}}, 
 {33, {1, 6, 6, 5, 3, 5, 6, 6, 6, 5, 5, 6, 6, 6, 6, 6, 6}}, 
 {33, {1, 6, 6, 1, 6, 6, 5, 6, 6, 6, 6, 6, 5, 6, 6, 6, 6}}, 
 {32, {4, 5, 5, 5, 5, 4, 5, 6, 5, 5, 5, 4, 5, 5, 5, 6, 6, 5}}, 
 {32, {4, 5, 5, 5, 5, 4, 5, 5, 6, 5, 5, 5, 4, 5, 5, 6, 6, 5}}, 
 {32, {4, 5, 5, 5, 4, 5, 6, 5, 5, 5, 5, 4, 5, 5, 5, 6, 6, 5}}, 
 {32, {4, 5, 5, 5, 4, 5, 6, 5, 5, 5, 4, 5, 5, 5, 5, 6, 6, 5}}, 
 {32, {4, 5, 5, 5, 4, 5, 5, 5, 5, 5, 6, 6, 5, 5, 4, 5, 6, 5}}, 
 {32, {4, 5, 5, 4, 5, 5, 6, 6, 5, 5, 5, 4, 5, 5, 5, 5, 6, 5}}, 
 {32, {4, 5, 5, 4, 5, 5, 5, 6, 6, 5, 5, 4, 5, 5, 5, 5, 6, 5}}, 
 {32, {4, 5, 5, 4, 5, 5, 5, 5, 4, 5, 6, 6, 5, 5, 5, 5, 6, 5}}, 
 {32, {4, 5, 5, 4, 5, 5, 5, 5, 4, 5, 5, 6, 5, 5, 6, 6, 5, 5}}, 
 {32, {4, 4, 5, 6, 5, 5, 5, 5, 5, 5, 4, 5, 6, 5, 5, 5, 6, 5}}, 
 {32, {4, 4, 5, 5, 5, 6, 5, 5, 5, 5, 6, 5, 4, 5, 5, 5, 6, 5}}, 
 {32, {4, 4, 5, 5, 5, 6, 5, 5, 5, 5, 5, 6, 5, 4, 5, 5, 6, 5}}, 
 {32, {4, 4, 5, 5, 5, 5, 6, 6, 6, 5, 5, 5, 4, 4, 5, 5, 6, 5}}, 
 {32, {4, 4, 5, 5, 5, 5, 6, 5, 4, 4, 5, 5, 5, 6, 6, 6, 5, 5}}, 
 {32, {4, 4, 5, 5, 5, 5, 5, 4, 5, 5, 5, 5, 6, 5, 5, 6, 6, 5}}, 
 {32, {4, 4, 5, 5, 5, 5, 4, 5, 6, 6, 5, 5, 5, 5, 5, 6, 5, 5}}, 
 {32, {4, 4, 5, 5, 5, 4, 5, 6, 6, 6, 6, 5, 5, 4, 5, 5, 5, 5}}, 
 {32, {4, 4, 5, 5, 5, 4, 5, 6, 6, 6, 6, 5, 4, 5, 5, 5, 5, 5}}, 
 {32, {4, 4, 5, 4, 5, 5, 6, 6, 5, 5, 6, 5, 5, 5, 5, 5, 5, 5}}, 
 {32, {4, 4, 5, 4, 5, 5, 5, 5, 5, 6, 5, 5, 5, 5, 6, 6, 5, 5}}, 
 {32, {4, 4, 5, 4, 5, 5, 5, 5, 5, 4, 5, 5, 6, 6, 6, 6, 5, 5}}, 
 {32, {3, 5, 6, 5, 5, 5, 4, 5, 6, 5, 5, 5, 5, 5, 5, 5, 6, 5}}, 
 {32, {3, 5, 6, 5, 5, 5, 4, 5, 5, 6, 5, 5, 5, 5, 5, 5, 6, 5}}, 
 {32, {3, 5, 6, 5, 5, 5, 4, 5, 5, 5, 6, 5, 5, 5, 5, 5, 6, 5}}, 
 {32, {3, 5, 5, 5, 5, 5, 5, 6, 6, 5, 5, 4, 5, 5, 6, 5, 5, 5}}, 
 {32, {3, 5, 5, 5, 5, 5, 5, 6, 6, 5, 4, 5, 5, 5, 5, 5, 6, 5}}, 
 {32, {3, 5, 5, 5, 5, 5, 5, 6, 5, 4, 5, 5, 6, 6, 5, 5, 5, 5}}, 
 {32, {3, 5, 5, 5, 5, 5, 5, 5, 5, 5, 6, 5, 5, 5, 5, 6, 5, 5}}, 
 {32, {3, 5, 5, 5, 5, 4, 5, 5, 6, 6, 5, 5, 5, 5, 6, 5, 5, 5}}, 
 {32, {3, 5, 5, 5, 4, 5, 5, 6, 5, 5, 5, 5, 5, 5, 6, 6, 5, 5}}, 
 {32, {3, 5, 5, 4, 5, 6, 6, 5, 5, 5, 5, 5, 5, 5, 5, 5, 6, 5}}, 
 {32, {3, 5, 5, 4, 5, 5, 5, 5, 5, 6, 5, 5, 5, 5, 5, 6, 6, 5}}, 
 {32, {3, 5, 5, 4, 5, 5, 5, 5, 5, 5, 5, 6, 6, 5, 5, 5, 6, 5}}, 
 {32, {3, 5, 5, 4, 5, 5, 5, 5, 5, 5, 5, 5, 6, 6, 5, 5, 6, 5}}, 
 {32, {3, 5, 5, 4, 4, 5, 5, 5, 6, 6, 6, 6, 5, 5, 5, 5, 5, 5}}, 
 {32, {1, 6, 6, 6, 1, 6, 6, 6, 5, 5, 4, 5, 5, 5, 5, 6, 6, 6}}, 
 {32, {1, 6, 6, 5, 5, 5, 5, 5, 4, 5, 5, 5, 4, 5, 6, 6, 6, 6}}, 
 {32, {1, 6, 6, 5, 5, 4, 4, 5, 5, 6, 6, 5, 5, 5, 5, 5, 6, 6}}, 
 {32, {1, 6, 6, 1, 6, 6, 5, 5, 5, 5, 5, 5, 6, 5, 5, 6, 6, 6}}, 
 {32, {4, 5, 5, 5, 5, 6, 6, 5, 4, 5, 6, 6, 5, 6, 6, 6, 5}}, 
 {32, {4, 5, 5, 4, 5, 6, 6, 5, 5, 5, 6, 6, 6, 5, 6, 6, 5}}, 
 {32, {4, 4, 5, 6, 6, 5, 6, 6, 5, 5, 4, 5, 6, 6, 6, 6, 5}}, 
 {32, {3, 5, 6, 5, 5, 6, 6, 6, 5, 5, 4, 5, 6, 6, 6, 6, 5}}, 
 {32, {3, 5, 6, 5, 5, 6, 5, 4, 5, 5, 6, 6, 6, 6, 6, 6, 5}}, 
 {32, {3, 5, 6, 5, 5, 5, 6, 6, 6, 6, 5, 4, 5, 6, 6, 6, 5}}, 
 {32, {3, 5, 6, 5, 5, 5, 6, 5, 5, 5, 6, 6, 5, 6, 6, 6, 5}}, 
 {32, {3, 5, 5, 6, 6, 5, 6, 6, 5, 5, 5, 6, 6, 5, 5, 6, 5}}, 
 {32, {3, 5, 5, 6, 5, 5, 5, 5, 6, 6, 6, 6, 5, 6, 6, 5, 5}}, 
 {32, {3, 5, 5, 5, 6, 6, 6, 6, 5, 4, 5, 6, 6, 6, 6, 5, 5}}, 
 {32, {3, 5, 5, 5, 6, 5, 5, 6, 6, 5, 6, 6, 6, 6, 5, 5, 5}}, 
 {32, {3, 5, 5, 4, 5, 6, 5, 5, 6, 6, 6, 6, 6, 6, 6, 5, 5}}, 
 {32, {1, 6, 6, 5, 5, 6, 5, 5, 6, 5, 5, 6, 6, 5, 6, 6, 6}}, 
 {32, {1, 6, 6, 5, 5, 3, 5, 5, 6, 6, 6, 6, 6, 6, 6, 6, 6}}, 
 {31, {4, 5, 5, 5, 5, 4, 5, 5, 6, 6, 5, 4, 5, 5, 5, 5, 6, 5}}, 
 {31, {4, 5, 5, 5, 5, 4, 5, 5, 5, 6, 5, 4, 5, 5, 5, 6, 6, 5}}, 
 {31, {4, 5, 5, 5, 5, 4, 5, 5, 5, 5, 6, 6, 5, 5, 4, 5, 6, 5}}, 
 {31, {4, 5, 5, 5, 4, 5, 5, 5, 6, 6, 5, 4, 5, 5, 5, 6, 5, 5}}, 
 {31, {4, 5, 5, 4, 5, 5, 5, 4, 5, 6, 6, 5, 5, 5, 5, 6, 5, 5}}, 
 {31, {4, 4, 5, 5, 5, 5, 6, 6, 6, 5, 5, 4, 4, 5, 5, 6, 5, 5}}, 
 {31, {4, 4, 5, 5, 5, 5, 5, 6, 5, 5, 5, 6, 5, 4, 5, 5, 6, 5}}, 
 {31, {4, 4, 5, 5, 5, 5, 5, 6, 5, 5, 5, 5, 5, 4, 5, 6, 6, 5}}, 
 {31, {4, 4, 5, 5, 5, 4, 5, 5, 5, 4, 5, 6, 6, 6, 6, 5, 5, 5}}, 
 {31, {4, 4, 5, 5, 5, 4, 4, 5, 5, 6, 6, 6, 5, 5, 5, 5, 6, 5}}, 
 {31, {4, 4, 5, 5, 4, 5, 5, 5, 5, 4, 5, 5, 5, 6, 6, 6, 6, 5}}, 
 {31, {4, 4, 5, 5, 4, 4, 5, 5, 5, 6, 6, 6, 5, 5, 5, 5, 6, 5}}, 
 {31, {4, 4, 5, 5, 4, 4, 5, 5, 5, 5, 6, 6, 5, 5, 6, 6, 5, 5}}, 
 {31, {4, 4, 5, 4, 5, 6, 5, 5, 5, 6, 6, 5, 5, 5, 5, 5, 5, 5}}, 
 {31, {4, 4, 5, 4, 5, 6, 5, 5, 5, 5, 5, 6, 6, 5, 5, 5, 5, 5}}, 
 {31, {4, 4, 5, 4, 5, 6, 5, 5, 5, 5, 5, 5, 5, 5, 6, 6, 5, 5}}, 
 {31, {4, 4, 5, 4, 5, 5, 6, 6, 5, 5, 5, 5, 5, 6, 5, 5, 5, 5}}, 
 {31, {4, 4, 5, 4, 5, 5, 6, 5, 5, 5, 5, 5, 6, 6, 5, 5, 5, 5}}, 
 {31, {4, 4, 5, 4, 5, 5, 6, 5, 5, 5, 5, 5, 5, 6, 6, 5, 5, 5}}, 
 {31, {4, 4, 5, 4, 5, 5, 5, 6, 5, 5, 5, 5, 6, 6, 5, 5, 5, 5}}, 
 {31, {4, 4, 5, 4, 5, 5, 5, 5, 6, 6, 5, 5, 5, 5, 5, 6, 5, 5}}, 
 {31, {4, 4, 5, 4, 5, 5, 5, 5, 6, 5, 5, 5, 5, 5, 6, 6, 5, 5}}, 
 {31, {4, 4, 5, 4, 5, 5, 5, 5, 5, 5, 6, 5, 5, 6, 6, 5, 5, 5}}, 
 {31, {4, 4, 5, 4, 4, 5, 5, 5, 6, 6, 6, 5, 5, 5, 5, 5, 6, 5}}, 
 {31, {3, 5, 5, 5, 5, 5, 6, 5, 5, 5, 5, 4, 5, 5, 5, 6, 6, 5}}, 
 {31, {3, 5, 5, 5, 5, 5, 5, 6, 6, 5, 5, 5, 5, 5, 4, 5, 6, 5}}, 
 {31, {3, 5, 5, 5, 5, 5, 5, 5, 6, 5, 4, 5, 5, 6, 6, 5, 5, 5}}, 
 {31, {3, 5, 5, 5, 5, 5, 5, 5, 5, 5, 4, 5, 6, 6, 5, 5, 6, 5}}, 
 {31, {3, 5, 5, 5, 4, 5, 6, 5, 5, 5, 5, 5, 5, 5, 6, 6, 5, 5}}, 
 {31, {3, 5, 5, 5, 4, 4, 5, 5, 5, 5, 6, 6, 6, 6, 5, 5, 5, 5}}, 
 {31, {3, 5, 5, 4, 5, 6, 5, 5, 5, 5, 5, 5, 5, 5, 5, 6, 6, 5}}, 
 {31, {3, 5, 5, 4, 5, 5, 6, 6, 5, 5, 5, 5, 6, 5, 5, 5, 5, 5}}, 
 {31, {3, 5, 5, 4, 5, 5, 5, 5, 6, 5, 5, 5, 5, 5, 5, 6, 6, 5}}, 
 {31, {3, 5, 5, 4, 5, 5, 5, 5, 5, 5, 6, 6, 5, 5, 5, 5, 6, 5}}, 
 {31, {1, 6, 6, 5, 5, 5, 5, 5, 5, 4, 5, 5, 5, 6, 5, 5, 6, 6}}, 
 {31, {1, 6, 6, 5, 5, 5, 5, 5, 4, 5, 5, 5, 6, 5, 5, 5, 6, 6}}, 
 {31, {1, 6, 6, 5, 5, 4, 5, 5, 5, 5, 6, 6, 6, 6, 1, 6, 6, 6}}, 
 {31, {1, 6, 6, 5, 4, 5, 5, 5, 5, 5, 4, 5, 5, 5, 6, 6, 6, 6}}, 
 {31, {1, 6, 6, 5, 3, 5, 6, 5, 5, 5, 5, 5, 5, 5, 5, 6, 6, 6}}, 
 {31, {4, 5, 6, 6, 5, 5, 6, 5, 5, 5, 5, 6, 5, 5, 6, 6, 5}}, 
 {31, {4, 5, 5, 6, 6, 5, 5, 6, 6, 5, 6, 6, 5, 4, 5, 6, 5}}, 
 {31, {4, 5, 5, 4, 5, 6, 6, 5, 5, 5, 6, 6, 5, 6, 6, 6, 5}}, 
 {31, {4, 5, 4, 5, 5, 6, 6, 5, 5, 5, 5, 6, 6, 6, 6, 6, 5}}, 
 {31, {4, 5, 4, 5, 5, 5, 6, 6, 6, 5, 5, 5, 6, 6, 6, 6, 5}}, 
 {31, {4, 4, 5, 6, 6, 5, 6, 6, 6, 5, 5, 4, 5, 6, 6, 6, 5}}, 
 {31, {4, 4, 5, 4, 5, 5, 5, 6, 6, 6, 6, 6, 6, 5, 6, 6, 5}}, 
 {31, {4, 4, 5, 4, 5, 5, 5, 6, 6, 6, 6, 5, 6, 6, 6, 6, 5}}, 
 {31, {3, 5, 6, 5, 5, 5, 6, 5, 4, 5, 6, 6, 6, 6, 6, 6, 5}}, 
 {31, {3, 5, 6, 5, 5, 4, 5, 6, 5, 5, 6, 6, 6, 6, 6, 6, 5}}, 
 {31, {3, 5, 5, 5, 6, 5, 4, 5, 6, 6, 6, 6, 6, 6, 6, 5, 5}}, 
 {31, {3, 5, 5, 5, 5, 6, 5, 5, 5, 6, 6, 6, 5, 6, 6, 6, 5}}, 
 {31, {3, 5, 5, 5, 4, 5, 6, 6, 6, 6, 5, 6, 6, 6, 6, 5, 5}}, 
 {31, {3, 5, 5, 5, 4, 5, 6, 6, 5, 5, 6, 6, 6, 6, 6, 6, 5}}, 
 {31, {1, 6, 6, 5, 6, 6, 6, 5, 6, 6, 6, 1, 6, 6, 6, 6, 6}}, 
 {31, {1, 6, 6, 5, 6, 6, 6, 5, 4, 5, 5, 6, 6, 5, 6, 6, 6}}, 
 {31, {1, 6, 6, 5, 6, 6, 5, 6, 6, 6, 1, 6, 6, 6, 6, 6, 6}}, 
 {31, {1, 6, 6, 5, 6, 6, 5, 5, 6, 5, 5, 6, 5, 5, 6, 6, 6}}, 
 {31, {1, 6, 6, 5, 4, 5, 6, 6, 6, 5, 5, 6, 6, 6, 5, 6, 6}}, 
 {31, {1, 6, 6, 5, 3, 5, 6, 6, 6, 6, 5, 5, 6, 6, 6, 6, 6}}, 
 {31, {1, 6, 6, 1, 6, 6, 5, 6, 6, 5, 6, 6, 6, 6, 6, 6, 6}}, 
 {30, {4, 5, 5, 5, 5, 5, 5, 4, 5, 5, 6, 6, 5, 5, 4, 5, 6, 5}}, 
 {30, {4, 5, 5, 5, 4, 5, 6, 5, 5, 4, 5, 5, 5, 5, 5, 6, 6, 5}}, 
 {30, {4, 5, 5, 5, 4, 5, 5, 6, 6, 5, 5, 5, 5, 5, 4, 5, 6, 5}}, 
 {30, {4, 5, 5, 4, 5, 5, 6, 5, 5, 5, 5, 5, 4, 5, 5, 6, 6, 5}}, 
 {30, {4, 5, 5, 4, 5, 5, 5, 6, 6, 5, 5, 5, 5, 4, 5, 5, 6, 5}}, 
 {30, {4, 5, 5, 4, 5, 5, 5, 4, 5, 6, 6, 5, 5, 5, 5, 5, 6, 5}}, 
 {30, {4, 5, 5, 4, 5, 5, 5, 4, 5, 5, 5, 6, 5, 5, 6, 6, 5, 5}}, 
 {30, {4, 4, 5, 6, 5, 5, 4, 5, 5, 6, 5, 5, 5, 5, 5, 5, 6, 5}}, 
 {30, {4, 4, 5, 5, 5, 6, 6, 5, 5, 5, 4, 4, 5, 5, 6, 6, 5, 5}}, 
 {30, {4, 4, 5, 5, 5, 6, 5, 5, 4, 4, 5, 5, 5, 6, 6, 6, 5, 5}}, 
 {30, {4, 4, 5, 5, 5, 5, 6, 6, 5, 5, 4, 4, 5, 5, 6, 6, 5, 5}}, 
 {30, {4, 4, 5, 5, 5, 5, 5, 6, 5, 5, 5, 4, 5, 6, 6, 5, 5, 5}}, 
 {30, {4, 4, 5, 5, 5, 4, 4, 5, 5, 6, 5, 5, 5, 6, 6, 6, 5, 5}}, 
 {30, {4, 4, 5, 5, 5, 4, 4, 5, 5, 5, 6, 6, 5, 5, 6, 6, 5, 5}}, 
 {30, {4, 4, 5, 5, 4, 5, 5, 6, 6, 6, 5, 5, 5, 4, 5, 5, 6, 5}}, 
 {30, {4, 4, 5, 5, 4, 5, 5, 6, 6, 6, 5, 5, 4, 5, 6, 5, 5, 5}}, 
 {30, {4, 4, 5, 4, 5, 5, 5, 6, 6, 6, 6, 5, 5, 5, 4, 5, 5, 5}}, 
 {30, {4, 4, 5, 4, 5, 5, 5, 6, 6, 5, 5, 5, 5, 6, 5, 5, 5, 5}}, 
 {30, {4, 4, 5, 4, 5, 5, 5, 6, 5, 5, 5, 5, 5, 6, 6, 5, 5, 5}}, 
 {30, {4, 4, 5, 4, 5, 5, 5, 5, 5, 6, 5, 5, 5, 5, 5, 6, 6, 5}}, 
 {30, {4, 4, 5, 4, 5, 5, 5, 5, 5, 5, 5, 5, 6, 5, 5, 6, 6, 5}}, 
 {30, {3, 5, 6, 5, 5, 5, 5, 5, 4, 5, 5, 5, 6, 5, 5, 5, 6, 5}}, 
 {30, {3, 5, 6, 5, 5, 5, 4, 5, 5, 5, 5, 5, 6, 5, 5, 5, 6, 5}}, 
 {30, {3, 5, 5, 5, 5, 6, 5, 5, 5, 5, 5, 5, 4, 5, 5, 6, 6, 5}}, 
 {30, {3, 5, 5, 5, 5, 5, 6, 5, 4, 5, 5, 5, 5, 5, 5, 6, 6, 5}}, 
 {30, {3, 5, 5, 5, 5, 5, 5, 5, 5, 5, 4, 4, 5, 6, 6, 6, 6, 5}}, 
 {30, {3, 5, 5, 5, 5, 5, 4, 5, 5, 5, 5, 6, 6, 5, 5, 5, 6, 5}}, 
 {30, {3, 5, 5, 5, 5, 4, 4, 5, 5, 5, 6, 6, 6, 6, 5, 5, 5, 5}}, 
 {30, {3, 5, 5, 5, 5, 4, 4, 5, 5, 5, 5, 5, 6, 6, 6, 6, 5, 5}}, 
 {30, {3, 5, 5, 5, 4, 5, 5, 6, 6, 5, 5, 5, 5, 5, 6, 5, 5, 5}}, 
 {30, {3, 5, 5, 5, 4, 5, 5, 6, 5, 5, 5, 6, 6, 5, 5, 5, 5, 5}}, 
 {30, {3, 5, 5, 4, 5, 5, 6, 6, 5, 5, 6, 5, 5, 5, 5, 5, 5, 5}}, 
 {30, {1, 6, 6, 5, 5, 5, 5, 4, 5, 5, 6, 6, 6, 1, 6, 6, 6, 6}}, 
 {30, {1, 6, 6, 5, 5, 5, 4, 5, 5, 5, 4, 5, 5, 5, 6, 6, 6, 6}}, 
 {30, {4, 5, 5, 5, 5, 6, 6, 6, 6, 5, 6, 6, 5, 4, 5, 6, 5}}, 
 {30, {4, 5, 5, 5, 5, 4, 5, 6, 6, 5, 6, 6, 5, 6, 6, 6, 5}}, 
 {30, {4, 5, 5, 4, 5, 6, 6, 5, 5, 6, 6, 5, 5, 6, 6, 6, 5}}, 
 {30, {4, 5, 4, 5, 5, 6, 6, 6, 6, 6, 6, 5, 5, 5, 5, 6, 5}}, 
 {30, {4, 5, 4, 5, 5, 5, 6, 6, 6, 6, 6, 6, 5, 5, 6, 5, 5}}, 
 {30, {4, 4, 5, 6, 6, 5, 4, 5, 5, 6, 6, 6, 6, 5, 6, 6, 5}}, 
 {30, {4, 4, 5, 5, 6, 6, 5, 6, 6, 5, 5, 6, 5, 5, 6, 6, 5}}, 
 {30, {4, 4, 5, 5, 6, 5, 5, 6, 5, 5, 6, 6, 6, 5, 6, 6, 5}}, 
 {30, {3, 5, 6, 6, 6, 5, 4, 5, 6, 5, 5, 5, 6, 6, 6, 6, 5}}, 
 {30, {3, 5, 6, 6, 5, 5, 5, 5, 4, 5, 6, 6, 6, 6, 6, 6, 5}}, 
 {30, {3, 5, 6, 5, 5, 4, 5, 5, 6, 6, 6, 5, 6, 6, 6, 6, 5}}, 
 {30, {3, 5, 6, 5, 4, 5, 6, 6, 6, 5, 5, 5, 6, 6, 6, 6, 5}}, 
 {30, {3, 5, 5, 6, 6, 6, 5, 6, 6, 5, 5, 5, 6, 5, 5, 6, 5}}, 
 {30, {3, 5, 5, 6, 5, 5, 5, 6, 5, 5, 5, 6, 6, 6, 6, 6, 5}}, 
 {30, {3, 5, 5, 5, 6, 6, 6, 6, 5, 5, 4, 5, 6, 6, 6, 6, 5}}, 
 {30, {3, 5, 5, 5, 6, 5, 5, 4, 5, 6, 6, 6, 6, 6, 6, 6, 5}}, 
 {30, {3, 5, 5, 5, 4, 5, 6, 6, 6, 6, 6, 6, 6, 5, 5, 6, 5}}, 
 {30, {3, 5, 5, 5, 4, 5, 6, 6, 6, 6, 6, 5, 6, 6, 6, 5, 5}}, 
 {30, {3, 5, 5, 5, 4, 5, 6, 6, 6, 5, 6, 6, 6, 6, 6, 5, 5}}, 
 {30, {1, 6, 6, 6, 6, 6, 5, 3, 5, 5, 5, 6, 6, 6, 6, 6, 6}}, 
 {30, {1, 6, 6, 6, 6, 5, 3, 5, 6, 5, 5, 6, 6, 6, 6, 6, 6}}, 
 {30, {1, 6, 6, 6, 5, 6, 6, 6, 5, 4, 4, 5, 6, 6, 6, 6, 6}}, 
 {30, {1, 6, 6, 6, 5, 6, 6, 5, 4, 5, 6, 5, 5, 6, 6, 6, 6}}, 
 {30, {1, 6, 6, 6, 5, 6, 6, 5, 4, 4, 5, 6, 6, 6, 6, 6, 6}}, 
 {30, {1, 6, 6, 6, 5, 3, 5, 5, 6, 6, 6, 6, 6, 5, 6, 6, 6}}, 
 {30, {1, 6, 6, 5, 6, 6, 6, 6, 5, 6, 6, 6, 1, 6, 6, 6, 6}}, 
 {30, {1, 6, 6, 5, 5, 6, 6, 5, 6, 6, 5, 4, 5, 6, 6, 6, 6}}, 
 {30, {1, 6, 6, 5, 5, 6, 5, 5, 6, 6, 5, 5, 6, 6, 5, 6, 6}}, 
 {30, {1, 6, 6, 5, 4, 5, 6, 6, 5, 5, 6, 6, 5, 6, 6, 6, 6}}, 
 {30, {1, 6, 6, 5, 4, 5, 5, 6, 6, 5, 6, 6, 6, 5, 6, 6, 6}}, 
 {29, {4, 5, 5, 5, 5, 5, 4, 5, 5, 5, 5, 5, 4, 5, 6, 6, 6, 5}}, 
 {29, {4, 5, 5, 5, 5, 4, 5, 5, 6, 5, 4, 5, 5, 5, 5, 6, 6, 5}}, 
 {29, {4, 5, 5, 5, 5, 4, 5, 5, 5, 5, 6, 5, 5, 4, 5, 6, 6, 5}}, 
 {29, {4, 5, 5, 5, 4, 5, 5, 5, 5, 6, 5, 5, 4, 5, 6, 6, 5, 5}}, 
 {29, {4, 5, 5, 5, 4, 5, 5, 5, 5, 5, 6, 6, 5, 4, 5, 5, 6, 5}}, 
 {29, {4, 5, 5, 5, 4, 5, 5, 5, 5, 4, 5, 6, 5, 5, 5, 6, 6, 5}}, 
 {29, {4, 5, 5, 4, 5, 5, 5, 5, 5, 6, 5, 5, 4, 5, 6, 6, 5, 5}}, 
 {29, {4, 5, 5, 4, 5, 5, 5, 5, 5, 5, 6, 5, 4, 5, 6, 6, 5, 5}}, 
 {29, {4, 4, 5, 5, 5, 5, 5, 5, 6, 6, 6, 5, 5, 4, 4, 5, 6, 5}}, 
 {29, {4, 4, 5, 5, 5, 5, 5, 5, 6, 5, 4, 4, 5, 5, 6, 6, 6, 5}}, 
 {29, {4, 4, 5, 5, 5, 4, 5, 5, 6, 5, 4, 5, 5, 6, 6, 6, 5, 5}}, 
 {29, {4, 4, 5, 5, 5, 4, 5, 5, 5, 4, 5, 5, 5, 6, 6, 6, 6, 5}}, 
 {29, {4, 4, 5, 5, 5, 4, 4, 5, 5, 5, 6, 6, 6, 5, 5, 5, 6, 5}}, 
 {29, {4, 4, 5, 4, 5, 5, 6, 6, 6, 6, 5, 5, 5, 5, 4, 5, 5, 5}}, 
 {29, {4, 4, 5, 4, 5, 5, 6, 6, 6, 6, 5, 5, 5, 4, 5, 5, 5, 5}}, 
 {29, {4, 4, 5, 4, 5, 5, 5, 6, 5, 5, 5, 5, 5, 5, 6, 6, 5, 5}}, 
 {29, {4, 4, 5, 4, 5, 5, 5, 5, 5, 6, 6, 5, 5, 6, 5, 5, 5, 5}}, 
 {29, {3, 5, 5, 5, 5, 6, 6, 5, 5, 5, 5, 5, 4, 5, 6, 5, 5, 5}}, 
 {29, {3, 5, 5, 5, 5, 5, 6, 5, 5, 4, 5, 5, 5, 5, 6, 6, 5, 5}}, 
 {29, {3, 5, 5, 5, 5, 5, 5, 6, 5, 4, 5, 5, 5, 5, 5, 6, 6, 5}}, 
 {29, {3, 5, 5, 5, 5, 5, 5, 5, 4, 5, 5, 5, 6, 5, 5, 6, 6, 5}}, 
 {29, {3, 5, 5, 5, 5, 5, 5, 4, 5, 6, 6, 5, 5, 5, 5, 5, 6, 5}}, 
 {29, {3, 5, 5, 5, 5, 5, 4, 5, 5, 5, 6, 5, 5, 5, 6, 6, 5, 5}}, 
 {29, {3, 5, 5, 5, 5, 4, 5, 6, 6, 5, 5, 5, 5, 5, 5, 5, 6, 5}}, 
 {29, {3, 5, 5, 5, 4, 5, 6, 5, 5, 5, 5, 5, 6, 6, 5, 5, 5, 5}}, 
 {29, {3, 5, 5, 5, 4, 5, 5, 5, 6, 6, 6, 6, 5, 5, 4, 5, 5, 5}}, 
 {29, {3, 5, 5, 5, 4, 5, 5, 5, 5, 6, 5, 5, 5, 5, 6, 6, 5, 5}}, 
 {29, {3, 5, 5, 4, 5, 5, 6, 6, 6, 6, 5, 5, 5, 5, 4, 5, 5, 5}}, 
 {29, {3, 5, 5, 4, 5, 5, 5, 5, 5, 5, 6, 5, 5, 5, 6, 6, 5, 5}}, 
 {29, {3, 5, 5, 4, 4, 5, 6, 6, 6, 6, 5, 5, 5, 5, 5, 5, 5, 5}}, 
 {29, {1, 6, 6, 6, 5, 5, 4, 5, 5, 5, 4, 4, 5, 5, 6, 6, 6, 6}}, 
 {29, {1, 6, 6, 5, 5, 5, 5, 5, 6, 5, 4, 4, 5, 5, 5, 6, 6, 6}}, 
 {29, {1, 6, 6, 5, 5, 5, 5, 5, 5, 4, 5, 5, 6, 5, 5, 5, 6, 6}}, 
 {29, {1, 6, 6, 5, 5, 5, 5, 4, 5, 5, 5, 5, 6, 5, 5, 5, 6, 6}}, 
 {29, {1, 6, 6, 5, 5, 4, 4, 5, 5, 6, 5, 5, 5, 5, 5, 6, 6, 6}}, 
 {29, {1, 6, 6, 5, 4, 5, 6, 5, 5, 5, 5, 5, 4, 5, 5, 6, 6, 6}}, 
 {29, {1, 6, 6, 5, 4, 4, 5, 5, 5, 5, 5, 6, 6, 5, 5, 5, 6, 6}}, 
 {29, {1, 6, 6, 1, 6, 6, 5, 5, 5, 5, 5, 5, 6, 6, 5, 5, 6, 6}}, 
 {29, {4, 5, 6, 5, 4, 5, 6, 6, 6, 5, 5, 5, 5, 6, 6, 6, 5}}, 
 {29, {4, 5, 5, 6, 6, 5, 4, 5, 6, 6, 5, 6, 6, 5, 5, 6, 5}}, 
 {29, {4, 5, 5, 4, 5, 5, 6, 5, 5, 6, 6, 6, 5, 6, 6, 6, 5}}, 
 {29, {4, 4, 5, 6, 6, 5, 5, 5, 6, 5, 5, 6, 6, 5, 6, 6, 5}}, 
 {29, {4, 4, 5, 6, 5, 5, 6, 6, 6, 6, 6, 5, 4, 5, 6, 6, 5}}, 
 {29, {4, 4, 5, 6, 5, 5, 6, 5, 5, 6, 5, 5, 6, 6, 6, 6, 5}}, 
 {29, {4, 4, 5, 6, 5, 4, 5, 5, 6, 6, 6, 6, 6, 5, 6, 6, 5}}, 
 {29, {4, 4, 5, 6, 5, 4, 5, 5, 6, 6, 5, 6, 6, 6, 6, 6, 5}}, 
 {29, {3, 5, 6, 6, 6, 5, 5, 5, 6, 5, 4, 5, 6, 6, 6, 6, 5}}, 
 {29, {3, 5, 6, 5, 5, 6, 6, 6, 5, 5, 6, 5, 5, 5, 6, 6, 5}}, 
 {29, {3, 5, 6, 5, 4, 5, 5, 6, 6, 6, 6, 6, 6, 5, 5, 6, 5}}, 
 {29, {3, 5, 6, 5, 4, 5, 5, 6, 6, 6, 5, 5, 6, 6, 6, 6, 5}}, 
 {29, {3, 5, 5, 6, 6, 5, 5, 6, 5, 5, 6, 5, 5, 6, 6, 6, 5}}, 
 {29, {3, 5, 5, 5, 6, 6, 5, 6, 6, 5, 5, 6, 6, 5, 5, 6, 5}}, 
 {29, {3, 5, 5, 5, 6, 5, 5, 6, 6, 5, 6, 6, 6, 5, 5, 6, 5}}, 
 {29, {3, 5, 5, 5, 5, 6, 6, 5, 6, 6, 6, 6, 5, 5, 6, 5, 5}}, 
 {29, {3, 5, 5, 5, 4, 5, 6, 6, 5, 6, 6, 6, 6, 6, 6, 5, 5}}, 
 {29, {3, 5, 5, 4, 4, 5, 5, 6, 6, 6, 6, 6, 6, 6, 6, 6, 5}}, 
 {29, {1, 6, 6, 6, 5, 5, 6, 5, 3, 5, 6, 6, 6, 6, 6, 6, 6}}, 
 {29, {1, 6, 6, 6, 5, 5, 4, 5, 6, 6, 5, 6, 6, 5, 6, 6, 6}}, 
 {29, {1, 6, 6, 5, 6, 6, 6, 6, 6, 5, 6, 6, 6, 1, 6, 6, 6}}, 
 {29, {1, 6, 6, 5, 6, 6, 6, 5, 6, 6, 5, 4, 5, 5, 6, 6, 6}}, 
 {29, {1, 6, 6, 5, 6, 6, 6, 1, 6, 6, 6, 6, 6, 5, 6, 6, 6}}, 
 {29, {1, 6, 6, 5, 4, 4, 5, 6, 6, 5, 6, 6, 6, 6, 6, 6, 6}}, 
 {29, {1, 6, 6, 1, 6, 6, 6, 5, 6, 6, 5, 6, 6, 6, 6, 6, 6}}, 
 {29, {1, 6, 1, 6, 6, 6, 6, 5, 5, 6, 6, 6, 6, 6, 6, 6, 6}}, 
 {28, {4, 4, 5, 5, 5, 4, 5, 5, 5, 5, 4, 4, 5, 5, 5, 5, 5, 5, 5}}, 
 {28, {4, 5, 5, 5, 5, 5, 5, 4, 5, 5, 6, 6, 5, 4, 5, 5, 6, 5}}, 
 {28, {4, 5, 5, 5, 5, 5, 4, 5, 5, 6, 6, 5, 5, 5, 4, 5, 6, 5}}, 
 {28, {4, 5, 5, 5, 5, 4, 5, 5, 5, 5, 6, 6, 5, 4, 5, 6, 5, 5}}, 
 {28, {4, 5, 5, 5, 4, 5, 5, 5, 6, 5, 4, 5, 5, 5, 5, 6, 6, 5}}, 
 {28, {4, 5, 5, 5, 4, 5, 5, 5, 5, 6, 6, 5, 4, 5, 5, 6, 5, 5}}, 
 {28, {4, 5, 5, 4, 5, 5, 6, 6, 5, 4, 5, 5, 5, 5, 5, 5, 6, 5}}, 
 {28, {4, 5, 5, 4, 5, 5, 5, 5, 4, 5, 5, 6, 6, 5, 5, 5, 6, 5}}, 
 {28, {4, 5, 5, 4, 5, 5, 5, 4, 5, 5, 6, 6, 5, 5, 5, 6, 5, 5}}, 
 {28, {4, 5, 5, 4, 5, 5, 5, 4, 5, 5, 6, 5, 5, 6, 6, 5, 5, 5}}, 
 {28, {4, 5, 5, 4, 5, 5, 5, 4, 5, 5, 6, 5, 5, 5, 6, 6, 5, 5}}, 
 {28, {4, 4, 5, 6, 5, 5, 5, 4, 5, 6, 5, 5, 5, 5, 5, 5, 6, 5}}, 
 {28, {4, 4, 5, 6, 5, 5, 5, 4, 5, 5, 6, 5, 5, 5, 5, 5, 6, 5}}, 
 {28, {4, 4, 5, 6, 5, 5, 4, 5, 5, 5, 5, 6, 5, 5, 5, 5, 6, 5}}, 
 {28, {4, 4, 5, 6, 5, 4, 5, 5, 6, 5, 5, 5, 5, 5, 5, 5, 6, 5}}, 
 {28, {4, 4, 5, 5, 6, 5, 5, 5, 5, 5, 5, 5, 5, 4, 5, 6, 6, 5}}, 
 {28, {4, 4, 5, 5, 5, 6, 6, 5, 4, 4, 5, 5, 5, 6, 6, 5, 5, 5}}, 
 {28, {4, 4, 5, 5, 5, 5, 6, 6, 5, 4, 4, 5, 5, 5, 6, 6, 5, 5}}, 
 {28, {4, 4, 5, 5, 5, 5, 5, 6, 6, 5, 5, 4, 4, 5, 6, 6, 5, 5}}, 
 {28, {4, 4, 5, 5, 5, 5, 5, 5, 6, 6, 6, 5, 4, 4, 5, 5, 6, 5}}, 
 {28, {4, 4, 5, 5, 4, 5, 5, 5, 5, 6, 6, 6, 6, 5, 4, 5, 5, 5}}, 
 {28, {4, 4, 5, 5, 4, 5, 5, 5, 4, 5, 6, 6, 6, 6, 5, 5, 5, 5}}, 
 {28, {4, 4, 5, 5, 4, 5, 5, 5, 4, 5, 5, 5, 5, 6, 6, 6, 6, 5}}, 
 {28, {4, 4, 5, 4, 5, 6, 5, 5, 6, 6, 5, 5, 5, 5, 5, 5, 5, 5}}, 
 {28, {3, 5, 6, 5, 5, 5, 4, 5, 5, 5, 5, 5, 5, 6, 5, 5, 6, 5}}, 
 {28, {3, 5, 6, 5, 5, 4, 4, 5, 5, 5, 5, 5, 5, 5, 6, 6, 6, 5}}, 
 {28, {3, 5, 5, 5, 5, 6, 5, 5, 5, 5, 4, 5, 5, 5, 6, 6, 5, 5}}, 
 {28, {3, 5, 5, 5, 5, 5, 6, 6, 5, 4, 5, 5, 5, 5, 5, 5, 6, 5}}, 
 {28, {3, 5, 5, 5, 5, 5, 6, 5, 4, 5, 5, 6, 6, 5, 5, 5, 5, 5}}, 
 {28, {3, 5, 5, 5, 5, 5, 5, 5, 5, 6, 5, 5, 4, 5, 6, 6, 5, 5}}, 
 {28, {3, 5, 5, 5, 5, 5, 5, 5, 5, 6, 5, 4, 5, 5, 6, 6, 5, 5}}, 
 {28, {3, 5, 5, 5, 5, 5, 4, 5, 6, 5, 5, 5, 5, 5, 6, 6, 5, 5}}, 
 {28, {3, 5, 5, 5, 5, 4, 5, 5, 5, 5, 6, 5, 5, 5, 6, 6, 5, 5}}, 
 {28, {3, 5, 5, 5, 4, 5, 6, 5, 5, 5, 6, 6, 5, 5, 5, 5, 5, 5}}, 
 {28, {3, 5, 5, 5, 4, 5, 6, 5, 5, 5, 5, 5, 5, 6, 6, 5, 5, 5}}, 
 {28, {3, 5, 5, 5, 4, 5, 5, 5, 6, 5, 5, 5, 5, 5, 5, 6, 6, 5}}, 
 {28, {3, 5, 5, 5, 4, 5, 5, 5, 5, 5, 5, 6, 5, 5, 6, 6, 5, 5}}, 
 {28, {3, 5, 5, 5, 4, 4, 5, 5, 6, 6, 6, 5, 5, 5, 5, 5, 6, 5}}, 
 {28, {3, 5, 5, 4, 5, 6, 5, 5, 5, 5, 5, 5, 5, 5, 6, 6, 5, 5}}, 
 {28, {3, 5, 5, 4, 5, 5, 5, 5, 5, 4, 5, 5, 6, 6, 6, 6, 5, 5}}, 
 {28, {1, 6, 6, 5, 5, 5, 4, 5, 5, 5, 5, 5, 5, 6, 5, 5, 6, 6}}, 
 {28, {1, 6, 6, 5, 5, 5, 4, 4, 5, 6, 5, 5, 5, 5, 5, 6, 6, 6}}, 
 {28, {1, 6, 6, 5, 5, 5, 4, 4, 5, 5, 6, 6, 5, 5, 5, 5, 6, 6}}, 
 {28, {4, 5, 6, 6, 5, 5, 5, 5, 6, 6, 5, 4, 5, 6, 6, 6, 5}}, 
 {28, {4, 5, 5, 6, 6, 6, 5, 6, 6, 5, 5, 6, 5, 4, 5, 6, 5}}, 
 {28, {4, 5, 5, 4, 5, 6, 5, 5, 6, 6, 6, 6, 5, 5, 6, 6, 5}}, 
 {28, {4, 5, 5, 4, 5, 6, 5, 5, 6, 5, 5, 6, 6, 6, 6, 6, 5}}, 
 {28, {4, 5, 5, 4, 5, 5, 5, 6, 6, 6, 5, 6, 6, 5, 6, 6, 5}}, 
 {28, {4, 4, 5, 6, 5, 5, 4, 5, 6, 6, 6, 5, 6, 6, 6, 6, 5}}, 
 {28, {4, 4, 5, 6, 5, 4, 5, 6, 6, 6, 6, 5, 5, 6, 6, 6, 5}}, 
 {28, {4, 4, 5, 6, 5, 4, 5, 6, 6, 6, 5, 5, 6, 6, 6, 6, 5}}, 
 {28, {4, 4, 5, 4, 5, 6, 5, 5, 5, 6, 6, 6, 6, 6, 6, 6, 5}}, 
 {28, {3, 5, 6, 6, 6, 5, 5, 4, 5, 6, 5, 5, 6, 6, 6, 6, 5}}, 
 {28, {3, 5, 6, 6, 5, 5, 5, 6, 5, 5, 6, 6, 5, 5, 6, 6, 5}}, 
 {28, {3, 5, 6, 5, 4, 5, 5, 6, 6, 6, 6, 5, 5, 6, 6, 6, 5}}, 
 {28, {3, 5, 6, 5, 4, 4, 5, 5, 6, 6, 6, 6, 6, 6, 6, 6, 5}}, 
 {28, {3, 5, 5, 6, 6, 5, 5, 6, 5, 5, 5, 6, 6, 5, 6, 6, 5}}, 
 {28, {3, 5, 5, 5, 6, 5, 5, 6, 6, 6, 6, 5, 6, 6, 5, 5, 5}}, 
 {28, {3, 5, 5, 4, 5, 5, 5, 6, 6, 6, 6, 6, 6, 5, 6, 6, 5}}, 
 {28, {1, 6, 6, 6, 6, 5, 5, 6, 6, 5, 3, 5, 6, 6, 6, 6, 6}}, 
 {28, {1, 6, 6, 6, 6, 5, 4, 4, 5, 6, 6, 6, 5, 6, 6, 6, 6}}, 
 {28, {1, 6, 6, 5, 6, 6, 6, 5, 6, 6, 6, 6, 6, 1, 6, 6, 6}}, 
 {28, {1, 6, 6, 5, 6, 6, 5, 5, 6, 6, 6, 5, 4, 5, 6, 6, 6}}, 
 {28, {1, 6, 6, 5, 4, 5, 5, 6, 6, 6, 5, 6, 6, 5, 6, 6, 6}}, 
 {28, {1, 6, 6, 1, 6, 6, 5, 6, 6, 6, 6, 6, 6, 5, 6, 6, 6}}, 
 {27, {4, 5, 5, 5, 5, 5, 5, 5, 4, 5, 6, 5, 4, 5, 5, 6, 6, 5}}, 
 {27, {4, 5, 5, 5, 5, 4, 5, 5, 5, 6, 6, 5, 4, 5, 5, 6, 5, 5}}, 
 {27, {4, 5, 5, 5, 5, 4, 5, 5, 5, 5, 6, 5, 4, 5, 6, 6, 5, 5}}, 
 {27, {4, 5, 5, 5, 4, 5, 5, 5, 4, 5, 5, 5, 6, 6, 5, 5, 6, 5}}, 
 {27, {4, 5, 5, 4, 5, 5, 6, 5, 5, 4, 5, 5, 5, 5, 6, 6, 5, 5}}, 
 {27, {4, 5, 5, 4, 5, 5, 5, 4, 5, 6, 5, 5, 6, 6, 5, 5, 5, 5}}, 
 {27, {4, 4, 5, 6, 5, 4, 5, 5, 5, 5, 6, 5, 5, 5, 5, 5, 6, 5}}, 
 {27, {4, 4, 5, 5, 5, 6, 5, 5, 4, 5, 6, 5, 5, 5, 6, 5, 5, 5}}, 
 {27, {4, 4, 5, 5, 5, 6, 5, 4, 5, 5, 6, 5, 5, 5, 5, 6, 5, 5}}, 
 {27, {4, 4, 5, 5, 5, 5, 6, 6, 6, 5, 5, 5, 4, 4, 5, 6, 5, 5}}, 
 {27, {4, 4, 5, 5, 5, 5, 6, 5, 4, 5, 5, 5, 6, 5, 5, 5, 6, 5}}, 
 {27, {4, 4, 5, 5, 5, 5, 6, 5, 4, 4, 5, 6, 6, 6, 5, 5, 5, 5}}, 
 {27, {4, 4, 5, 5, 5, 5, 5, 6, 5, 4, 5, 6, 5, 5, 5, 5, 6, 5}}, 
 {27, {4, 4, 5, 5, 5, 5, 4, 4, 5, 5, 6, 6, 5, 5, 5, 6, 6, 5}}, 
 {27, {4, 4, 5, 5, 5, 4, 5, 5, 6, 6, 6, 5, 5, 4, 5, 5, 6, 5}}, 
 {27, {4, 4, 5, 5, 5, 4, 4, 5, 6, 5, 5, 5, 5, 5, 6, 6, 6, 5}}, 
 {27, {4, 4, 5, 5, 4, 5, 5, 5, 6, 6, 6, 5, 5, 4, 5, 5, 6, 5}}, 
 {27, {4, 4, 5, 5, 4, 5, 5, 5, 6, 5, 5, 6, 5, 5, 5, 5, 6, 5}}, 
 {27, {4, 4, 5, 5, 4, 5, 5, 5, 5, 5, 4, 5, 6, 6, 6, 6, 5, 5}}, 
 {27, {4, 4, 5, 5, 4, 4, 5, 6, 5, 5, 5, 5, 5, 5, 6, 6, 6, 5}}, 
 {27, {4, 4, 5, 5, 4, 4, 5, 5, 6, 6, 5, 5, 5, 5, 5, 6, 6, 5}}, 
 {27, {4, 4, 5, 5, 4, 4, 5, 5, 5, 5, 6, 6, 6, 5, 5, 6, 5, 5}}, 
 {27, {4, 4, 5, 4, 5, 6, 6, 5, 5, 5, 5, 5, 5, 5, 5, 5, 6, 5}}, 
 {27, {4, 4, 5, 4, 5, 5, 6, 6, 6, 6, 5, 4, 5, 5, 5, 5, 5, 5}}, 
 {27, {4, 4, 5, 4, 5, 5, 5, 6, 6, 6, 6, 5, 5, 4, 5, 5, 5, 5}}, 
 {27, {4, 4, 5, 4, 5, 5, 5, 5, 5, 5, 6, 5, 5, 5, 6, 6, 5, 5}}, 
 {27, {4, 4, 5, 4, 5, 5, 5, 5, 4, 5, 5, 6, 6, 6, 6, 5, 5, 5}}, 
 {27, {4, 4, 5, 4, 5, 5, 4, 5, 5, 6, 6, 6, 6, 5, 5, 5, 5, 5}}, 
 {27, {4, 4, 5, 4, 4, 5, 5, 5, 6, 5, 5, 5, 5, 6, 6, 6, 5, 5}}, 
 {27, {3, 5, 6, 5, 5, 5, 4, 5, 5, 5, 5, 6, 5, 5, 5, 5, 6, 5}}, 
 {27, {3, 5, 6, 5, 5, 4, 5, 5, 5, 5, 5, 6, 5, 5, 5, 5, 6, 5}}, 
 {27, {3, 5, 5, 5, 6, 6, 5, 5, 5, 5, 5, 4, 5, 5, 5, 5, 6, 5}}, 
 {27, {3, 5, 5, 5, 5, 5, 6, 6, 5, 5, 5, 5, 4, 5, 6, 5, 5, 5}}, 
 {27, {3, 5, 5, 5, 5, 5, 6, 5, 5, 5, 6, 5, 4, 5, 5, 5, 6, 5}}, 
 {27, {3, 5, 5, 5, 5, 5, 6, 5, 5, 4, 5, 5, 5, 6, 6, 5, 5, 5}}, 
 {27, {3, 5, 5, 5, 5, 5, 5, 6, 5, 5, 5, 4, 5, 5, 5, 6, 6, 5}}, 
 {27, {3, 5, 5, 5, 5, 5, 5, 6, 5, 5, 4, 5, 5, 5, 6, 6, 5, 5}}, 
 {27, {3, 5, 5, 5, 5, 5, 5, 6, 5, 4, 5, 5, 5, 6, 6, 5, 5, 5}}, 
 {27, {3, 5, 5, 5, 5, 5, 5, 5, 6, 5, 5, 4, 5, 5, 6, 6, 5, 5}}, 
 {27, {3, 5, 5, 5, 5, 5, 5, 5, 5, 6, 5, 5, 4, 5, 5, 6, 6, 5}}, 
 {27, {3, 5, 5, 5, 5, 5, 5, 5, 5, 4, 4, 5, 5, 6, 6, 6, 6, 5}}, 
 {27, {3, 5, 5, 5, 5, 5, 4, 5, 6, 5, 5, 5, 5, 5, 5, 6, 6, 5}}, 
 {27, {3, 5, 5, 5, 5, 5, 4, 5, 5, 6, 5, 5, 5, 5, 6, 6, 5, 5}}, 
 {27, {3, 5, 5, 5, 5, 5, 4, 4, 5, 5, 6, 6, 6, 6, 5, 5, 5, 5}}, 
 {27, {3, 5, 5, 5, 5, 4, 5, 5, 6, 5, 5, 5, 5, 5, 6, 6, 5, 5}}, 
 {27, {3, 5, 5, 5, 5, 4, 5, 5, 5, 5, 5, 5, 6, 5, 5, 6, 6, 5}}, 
 {27, {3, 5, 5, 5, 5, 4, 4, 5, 5, 5, 5, 5, 5, 6, 6, 6, 6, 5}}, 
 {27, {3, 5, 5, 5, 4, 5, 5, 5, 6, 6, 5, 5, 5, 5, 6, 5, 5, 5}}, 
 {27, {3, 5, 5, 5, 4, 5, 5, 5, 5, 5, 6, 5, 5, 5, 5, 6, 6, 5}}, 
 {27, {3, 5, 5, 5, 4, 4, 5, 5, 5, 6, 6, 6, 5, 5, 5, 5, 6, 5}}, 
 {27, {3, 5, 5, 5, 4, 4, 5, 5, 5, 5, 5, 5, 6, 6, 6, 6, 5, 5}}, 
 {27, {3, 5, 5, 4, 5, 5, 6, 5, 5, 5, 5, 5, 5, 5, 6, 6, 5, 5}}, 
 {27, {3, 5, 5, 4, 5, 5, 5, 6, 5, 5, 5, 5, 5, 5, 6, 6, 5, 5}}, 
 {27, {3, 5, 5, 4, 5, 5, 5, 5, 5, 5, 5, 6, 5, 5, 5, 6, 6, 5}}, 
 {27, {1, 6, 6, 5, 5, 5, 5, 5, 4, 4, 5, 6, 5, 5, 5, 6, 6, 6}}, 
 {27, {1, 6, 6, 5, 5, 5, 5, 4, 4, 5, 5, 6, 6, 5, 5, 5, 6, 6}}, 
 {27, {1, 6, 6, 5, 5, 4, 5, 6, 5, 5, 5, 5, 4, 5, 5, 6, 6, 6}}, 
 {27, {1, 6, 6, 5, 4, 5, 5, 4, 5, 5, 5, 5, 5, 5, 6, 6, 6, 6}}, 
 {27, {1, 6, 6, 1, 6, 6, 6, 5, 5, 5, 4, 5, 5, 5, 6, 6, 6, 6}}, 
 {27, {4, 5, 6, 5, 5, 6, 5, 4, 5, 6, 6, 5, 5, 6, 6, 6, 5}}, 
 {27, {4, 5, 5, 6, 6, 5, 6, 6, 5, 4, 5, 6, 5, 5, 6, 6, 5}}, 
 {27, {4, 5, 5, 4, 5, 6, 6, 5, 5, 6, 5, 5, 6, 6, 6, 6, 5}}, 
 {27, {4, 5, 4, 5, 5, 6, 5, 5, 5, 5, 6, 6, 6, 6, 6, 6, 5}}, 
 {27, {3, 5, 6, 6, 5, 5, 6, 6, 5, 5, 4, 5, 6, 6, 6, 6, 5}}, 
 {27, {3, 5, 6, 5, 5, 5, 6, 6, 6, 5, 4, 5, 6, 6, 6, 6, 5}}, 
 {27, {3, 5, 6, 5, 5, 5, 5, 6, 5, 5, 6, 6, 5, 6, 6, 6, 5}}, 
 {27, {3, 5, 6, 5, 5, 5, 4, 5, 6, 6, 6, 5, 6, 6, 6, 6, 5}}, 
 {27, {3, 5, 5, 6, 6, 6, 6, 5, 5, 6, 5, 5, 6, 5, 5, 6, 5}}, 
 {27, {3, 5, 5, 6, 6, 6, 5, 5, 6, 5, 4, 5, 6, 6, 6, 6, 5}}, 
 {27, {3, 5, 5, 5, 6, 6, 6, 5, 5, 6, 6, 6, 5, 5, 6, 5, 5}}, 
 {27, {3, 5, 5, 5, 6, 6, 5, 6, 6, 5, 6, 6, 5, 5, 5, 6, 5}}, 
 {27, {3, 5, 5, 5, 6, 6, 5, 5, 6, 6, 6, 6, 5, 5, 6, 5, 5}}, 
 {27, {3, 5, 5, 5, 6, 6, 5, 5, 6, 6, 5, 6, 6, 5, 5, 6, 5}}, 
 {27, {3, 5, 5, 5, 4, 4, 5, 6, 6, 6, 6, 6, 6, 6, 6, 6, 5}}, 
 {27, {3, 5, 5, 4, 5, 5, 5, 6, 6, 6, 5, 6, 6, 6, 6, 6, 5}}, 
 {27, {1, 6, 6, 6, 6, 5, 5, 6, 5, 3, 5, 6, 6, 6, 6, 6, 6}}, 
 {27, {1, 6, 6, 6, 6, 5, 3, 5, 6, 6, 6, 5, 5, 6, 6, 6, 6}}, 
 {27, {1, 6, 6, 6, 5, 5, 6, 6, 6, 5, 3, 5, 6, 6, 6, 6, 6}}, 
 {27, {1, 6, 6, 5, 6, 6, 5, 5, 6, 5, 4, 5, 6, 6, 6, 6, 6}}, 
 {27, {1, 6, 6, 5, 6, 6, 1, 6, 6, 6, 5, 6, 6, 6, 6, 6, 6}}, 
 {27, {1, 6, 6, 5, 5, 6, 5, 5, 5, 6, 6, 5, 6, 6, 5, 6, 6}}, 
 {27, {1, 6, 6, 5, 4, 5, 6, 6, 6, 5, 6, 6, 5, 5, 6, 6, 6}}, 
 {27, {1, 6, 6, 5, 3, 5, 6, 6, 6, 6, 6, 6, 5, 5, 6, 6, 6}}, 
 {27, {1, 6, 6, 5, 3, 5, 5, 6, 6, 6, 5, 6, 6, 6, 6, 6, 6}}, 
 {27, {1, 6, 6, 5, 3, 5, 5, 5, 6, 6, 6, 6, 6, 6, 6, 6, 6}}, 
 {27, {1, 6, 1, 6, 6, 6, 6, 6, 5, 5, 6, 6, 6, 6, 6, 6, 6}}, 
 {27, {1, 6, 1, 6, 6, 6, 5, 5, 6, 6, 6, 6, 6, 6, 6, 6, 6}}, 
 {26, {4, 5, 5, 5, 6, 5, 4, 5, 5, 6, 5, 4, 5, 5, 6, 5, 5, 5}}, 
 {26, {4, 5, 5, 5, 5, 5, 4, 5, 6, 5, 5, 5, 4, 5, 5, 6, 6, 5}}, 
 {26, {4, 5, 5, 5, 5, 4, 5, 6, 5, 5, 4, 5, 5, 5, 5, 6, 6, 5}}, 
 {26, {4, 5, 5, 5, 4, 5, 6, 5, 5, 5, 5, 5, 5, 4, 5, 6, 6, 5}}, 
 {26, {4, 5, 5, 5, 4, 5, 5, 5, 6, 6, 5, 5, 4, 5, 5, 6, 5, 5}}, 
 {26, {4, 5, 5, 5, 4, 5, 5, 5, 5, 6, 5, 4, 5, 6, 6, 5, 5, 5}}, 
 {26, {4, 5, 5, 4, 5, 5, 6, 5, 5, 5, 5, 5, 5, 4, 5, 6, 6, 5}}, 
 {26, {4, 5, 5, 4, 5, 5, 5, 6, 5, 5, 4, 5, 5, 6, 6, 5, 5, 5}}, 
 {26, {4, 5, 5, 4, 5, 5, 5, 5, 6, 6, 5, 5, 4, 5, 5, 6, 5, 5}}, 
 {26, {4, 5, 5, 4, 5, 5, 5, 5, 5, 6, 5, 5, 4, 5, 5, 6, 6, 5}}, 
 {26, {4, 5, 5, 4, 5, 5, 5, 5, 5, 4, 5, 5, 6, 6, 5, 5, 6, 5}}, 
 {26, {4, 5, 5, 4, 5, 5, 4, 5, 5, 6, 6, 5, 5, 5, 5, 5, 6, 5}}, 
 {26, {4, 4, 5, 6, 5, 4, 5, 5, 5, 5, 5, 5, 5, 6, 5, 5, 6, 5}}, 
 {26, {4, 4, 5, 5, 5, 6, 5, 5, 5, 5, 6, 5, 5, 4, 5, 6, 5, 5}}, 
 {26, {4, 4, 5, 5, 5, 6, 5, 5, 5, 5, 6, 5, 4, 5, 5, 6, 5, 5}}, 
 {26, {4, 4, 5, 5, 5, 6, 5, 5, 5, 5, 5, 6, 5, 4, 5, 6, 5, 5}}, 
 {26, {4, 4, 5, 5, 5, 6, 5, 5, 4, 5, 6, 5, 5, 5, 5, 5, 6, 5}}, 
 {26, {4, 4, 5, 5, 5, 6, 5, 4, 5, 6, 5, 5, 5, 5, 5, 5, 6, 5}}, 
 {26, {4, 4, 5, 5, 5, 5, 6, 5, 5, 4, 4, 5, 5, 6, 6, 6, 5, 5}}, 
 {26, {4, 4, 5, 5, 5, 5, 6, 5, 4, 4, 5, 5, 5, 5, 6, 6, 6, 5}}, 
 {26, {4, 4, 5, 5, 5, 5, 5, 4, 5, 6, 6, 5, 5, 5, 6, 5, 5, 5}}, 
 {26, {4, 4, 5, 5, 5, 5, 5, 4, 5, 5, 4, 5, 5, 6, 6, 6, 6, 5}}, 
 {26, {4, 4, 5, 5, 5, 5, 4, 5, 5, 5, 5, 4, 5, 6, 6, 6, 6, 5}}, 
 {26, {4, 4, 5, 5, 5, 5, 4, 5, 5, 5, 4, 5, 6, 6, 6, 6, 5, 5}}, 
 {26, {4, 4, 5, 5, 5, 4, 4, 5, 5, 5, 6, 5, 5, 6, 6, 6, 5, 5}}, 
 {26, {4, 4, 5, 5, 4, 5, 6, 6, 5, 5, 5, 5, 5, 5, 5, 6, 5, 5}}, 
 {26, {4, 4, 5, 5, 4, 5, 5, 6, 5, 5, 5, 5, 6, 5, 5, 5, 6, 5}}, 
 {26, {4, 4, 5, 5, 4, 5, 5, 5, 5, 4, 5, 6, 6, 6, 6, 5, 5, 5}}, 
 {26, {4, 4, 5, 5, 4, 4, 5, 5, 5, 6, 6, 5, 5, 5, 6, 6, 5, 5}}, 
 {26, {4, 4, 5, 5, 4, 4, 5, 5, 5, 6, 6, 5, 5, 5, 5, 6, 6, 5}}, 
 {26, {4, 4, 5, 4, 5, 5, 5, 6, 6, 5, 5, 5, 5, 5, 6, 5, 5, 5}}, 
 {26, {4, 4, 5, 4, 5, 5, 5, 5, 5, 6, 6, 6, 6, 5, 5, 4, 5, 5}}, 
 {26, {4, 4, 5, 4, 5, 5, 5, 4, 5, 5, 6, 6, 6, 6, 5, 5, 5, 5}}, 
 {26, {4, 4, 5, 4, 4, 5, 5, 6, 6, 5, 5, 5, 5, 5, 5, 6, 6, 5}}, 
 {26, {3, 5, 6, 5, 5, 5, 5, 5, 5, 5, 5, 4, 5, 6, 5, 5, 6, 5}}, 
 {26, {3, 5, 6, 5, 5, 5, 5, 5, 5, 4, 5, 6, 5, 5, 5, 5, 6, 5}}, 
 {26, {3, 5, 6, 5, 5, 5, 5, 5, 5, 4, 5, 5, 6, 5, 5, 5, 6, 5}}, 
 {26, {3, 5, 6, 5, 5, 5, 5, 5, 4, 5, 5, 6, 5, 5, 5, 5, 6, 5}}, 
 {26, {3, 5, 6, 5, 5, 5, 5, 4, 5, 6, 5, 5, 5, 5, 5, 5, 6, 5}}, 
 {26, {3, 5, 5, 6, 6, 6, 5, 5, 5, 5, 4, 4, 5, 5, 5, 5, 6, 5}}, 
 {26, {3, 5, 5, 5, 6, 5, 5, 5, 5, 5, 4, 5, 5, 5, 6, 6, 5, 5}}, 
 {26, {3, 5, 5, 5, 5, 6, 5, 5, 5, 5, 5, 5, 4, 5, 6, 6, 5, 5}}, 
 {26, {3, 5, 5, 5, 5, 6, 5, 5, 5, 5, 5, 4, 5, 5, 6, 6, 5, 5}}, 
 {26, {3, 5, 5, 5, 5, 6, 5, 5, 5, 4, 5, 5, 5, 5, 6, 6, 5, 5}}, 
 {26, {3, 5, 5, 5, 5, 5, 6, 5, 5, 5, 4, 5, 5, 6, 6, 5, 5, 5}}, 
 {26, {3, 5, 5, 5, 5, 5, 6, 5, 5, 5, 4, 5, 5, 5, 6, 6, 5, 5}}, 
 {26, {3, 5, 5, 5, 5, 5, 5, 6, 6, 5, 4, 5, 5, 5, 6, 5, 5, 5}}, 
 {26, {3, 5, 5, 5, 5, 5, 5, 6, 5, 5, 5, 5, 4, 5, 5, 6, 6, 5}}, 
 {26, {3, 5, 5, 5, 5, 5, 5, 5, 6, 5, 5, 5, 4, 5, 6, 6, 5, 5}}, 
 {26, {3, 5, 5, 5, 5, 5, 5, 5, 5, 4, 4, 5, 6, 6, 6, 6, 5, 5}}, 
 {26, {3, 5, 5, 5, 5, 5, 5, 5, 4, 5, 6, 5, 5, 6, 6, 5, 5, 5}}, 
 {26, {3, 5, 5, 5, 5, 5, 4, 5, 5, 5, 6, 5, 5, 5, 5, 6, 6, 5}}, 
 {26, {3, 5, 5, 5, 5, 5, 4, 4, 5, 5, 5, 6, 6, 6, 6, 5, 5, 5}}, 
 {26, {3, 5, 5, 5, 5, 4, 5, 5, 5, 6, 5, 5, 5, 6, 6, 5, 5, 5}}, 
 {26, {3, 5, 5, 5, 4, 5, 5, 6, 5, 5, 5, 5, 5, 6, 6, 5, 5, 5}}, 
 {26, {3, 5, 5, 5, 4, 5, 5, 5, 6, 5, 5, 5, 5, 6, 6, 5, 5, 5}}, 
 {26, {3, 5, 5, 5, 4, 5, 5, 5, 5, 6, 6, 5, 5, 5, 6, 5, 5, 5}}, 
 {26, {3, 5, 5, 5, 4, 5, 5, 5, 5, 5, 5, 5, 6, 5, 5, 6, 6, 5}}, 
 {26, {3, 5, 5, 4, 5, 5, 6, 6, 5, 5, 5, 5, 5, 6, 5, 5, 5, 5}}, 
 {26, {3, 5, 5, 3, 5, 5, 5, 5, 5, 5, 5, 5, 6, 6, 6, 6, 5, 5}}, 
 {26, {1, 6, 6, 6, 5, 5, 4, 4, 5, 5, 5, 4, 5, 5, 6, 6, 6, 6}}, 
 {26, {1, 6, 6, 5, 5, 5, 5, 5, 5, 6, 5, 4, 4, 5, 5, 6, 6, 6}}, 
 {26, {1, 6, 6, 5, 5, 5, 5, 4, 5, 5, 6, 6, 6, 6, 1, 6, 6, 6}}, 
 {26, {1, 6, 6, 5, 5, 5, 5, 4, 5, 5, 4, 5, 5, 5, 6, 6, 6, 6}}, 
 {26, {1, 6, 6, 5, 5, 4, 5, 5, 6, 6, 5, 5, 4, 5, 5, 5, 6, 6}}, 
 {26, {1, 6, 6, 5, 5, 4, 5, 5, 5, 4, 4, 5, 5, 6, 6, 6, 6, 6}}, 
 {26, {1, 6, 6, 5, 5, 4, 4, 5, 6, 5, 5, 5, 5, 5, 5, 6, 6, 6}}, 
 {26, {1, 6, 6, 5, 4, 4, 5, 6, 5, 5, 5, 5, 5, 5, 5, 6, 6, 6}}, 
 {26, {1, 6, 6, 1, 6, 6, 6, 1, 6, 6, 6, 6, 5, 5, 5, 6, 6, 6}}, 
 {26, {1, 6, 6, 1, 6, 6, 5, 4, 5, 5, 5, 5, 5, 6, 6, 6, 6, 6}}, 
 {26, {4, 5, 6, 5, 5, 6, 5, 5, 6, 5, 5, 6, 5, 5, 6, 6, 5}}, 
 {26, {4, 4, 5, 6, 6, 5, 6, 6, 5, 4, 5, 5, 6, 6, 6, 6, 5}}, 
 {26, {4, 4, 5, 6, 6, 5, 5, 6, 6, 6, 5, 4, 5, 6, 6, 6, 5}}, 
 {26, {4, 4, 5, 6, 6, 5, 5, 4, 5, 6, 6, 6, 6, 5, 6, 6, 5}}, 
 {26, {4, 4, 5, 6, 6, 5, 4, 5, 5, 6, 6, 6, 5, 6, 6, 6, 5}}, 
 {26, {4, 4, 5, 4, 5, 6, 6, 6, 6, 6, 6, 5, 6, 6, 5, 5, 5}}, 
 {26, {4, 4, 5, 4, 5, 6, 6, 6, 6, 5, 6, 6, 6, 6, 5, 5, 5}}, 
 {26, {3, 5, 6, 6, 5, 5, 6, 6, 5, 4, 5, 5, 6, 6, 6, 6, 5}}, 
 {26, {3, 5, 6, 6, 5, 5, 6, 5, 5, 6, 5, 5, 5, 6, 6, 6, 5}}, 
 {26, {3, 5, 6, 6, 5, 5, 6, 5, 5, 5, 5, 6, 6, 5, 6, 6, 5}}, 
 {26, {3, 5, 6, 5, 5, 5, 6, 6, 6, 6, 6, 5, 4, 5, 6, 6, 5}}, 
 {26, {3, 5, 5, 6, 6, 6, 6, 6, 5, 6, 6, 5, 5, 4, 5, 6, 5}}, 
 {26, {3, 5, 5, 6, 6, 6, 5, 6, 6, 5, 4, 5, 5, 6, 6, 6, 5}}, 
 {26, {3, 5, 5, 6, 6, 6, 5, 4, 5, 5, 6, 6, 6, 5, 6, 6, 5}}, 
 {26, {3, 5, 5, 6, 6, 5, 6, 6, 5, 5, 6, 5, 5, 5, 6, 6, 5}}, 
 {26, {3, 5, 5, 6, 6, 5, 5, 6, 6, 5, 4, 5, 6, 6, 6, 6, 5}}, 
 {26, {3, 5, 5, 6, 6, 5, 5, 6, 5, 5, 6, 6, 5, 6, 6, 5, 5}}, 
 {26, {3, 5, 5, 6, 6, 5, 5, 5, 4, 5, 6, 6, 6, 6, 6, 6, 5}}, 
 {26, {3, 5, 5, 5, 6, 6, 6, 6, 6, 5, 4, 5, 6, 6, 6, 5, 5}}, 
 {26, {3, 5, 5, 5, 6, 5, 5, 5, 6, 5, 5, 6, 6, 6, 6, 6, 5}}, 
 {26, {3, 5, 5, 4, 5, 5, 6, 5, 5, 6, 6, 6, 6, 6, 6, 6, 5}}, 
 {26, {1, 6, 6, 6, 6, 5, 6, 6, 5, 3, 5, 5, 6, 6, 6, 6, 6}}, 
 {26, {1, 6, 6, 6, 5, 6, 6, 5, 5, 6, 5, 4, 5, 6, 6, 6, 6}}, 
 {26, {1, 6, 6, 6, 5, 5, 6, 6, 5, 3, 5, 6, 6, 6, 6, 6, 6}}, 
 {26, {1, 6, 6, 6, 5, 4, 5, 6, 5, 5, 6, 6, 5, 6, 6, 6, 6}}, 
 {26, {1, 6, 6, 6, 5, 3, 5, 6, 6, 6, 6, 6, 5, 5, 6, 6, 6}}, 
 {26, {1, 6, 6, 6, 5, 3, 5, 5, 5, 6, 6, 6, 6, 6, 6, 6, 6}}, 
 {26, {1, 6, 6, 5, 6, 6, 6, 6, 1, 6, 6, 6, 5, 6, 6, 6, 6}}, 
 {26, {1, 6, 6, 5, 6, 6, 6, 5, 5, 4, 5, 6, 6, 5, 6, 6, 6}}, 
 {26, {1, 6, 6, 5, 6, 6, 6, 5, 4, 5, 5, 6, 6, 6, 5, 6, 6}}, 
 {26, {1, 6, 6, 5, 6, 6, 5, 4, 5, 6, 6, 5, 5, 6, 6, 6, 6}}, 
 {26, {1, 6, 6, 5, 6, 6, 1, 6, 6, 6, 6, 5, 6, 6, 6, 6, 6}}, 
 {26, {1, 6, 6, 5, 5, 4, 5, 6, 6, 6, 5, 6, 6, 5, 6, 6, 6}}, 
 {26, {1, 6, 6, 5, 4, 5, 6, 6, 5, 6, 6, 6, 5, 5, 6, 6, 6}}, 
 {26, {1, 6, 6, 5, 4, 5, 5, 6, 6, 6, 6, 5, 6, 6, 5, 6, 6}}, 
 {25, {4, 4, 5, 5, 5, 4, 4, 5, 5, 5, 5, 4, 5, 5, 5, 5, 5, 5, 5}}, 
 {25, {4, 4, 5, 5, 4, 5, 5, 5, 5, 5, 5, 5, 5, 4, 4, 5, 5, 5, 5}}, 
 {25, {4, 5, 5, 5, 5, 5, 5, 5, 4, 5, 5, 6, 5, 4, 5, 6, 6, 5}}, 
 {25, {4, 5, 5, 5, 4, 5, 5, 5, 6, 5, 5, 4, 5, 6, 6, 5, 5, 5}}, 
 {25, {4, 5, 5, 4, 5, 5, 5, 6, 6, 5, 5, 5, 5, 4, 5, 6, 5, 5}}, 
 {25, {4, 5, 5, 4, 5, 5, 5, 5, 6, 5, 5, 5, 4, 5, 6, 6, 5, 5}}, 
 {25, {4, 5, 5, 4, 5, 5, 5, 5, 5, 6, 6, 5, 4, 5, 6, 5, 5, 5}}, 
 {25, {4, 5, 5, 4, 5, 5, 5, 5, 5, 6, 6, 5, 4, 5, 5, 6, 5, 5}}, 
 {25, {4, 5, 5, 4, 5, 5, 5, 5, 4, 5, 6, 5, 5, 6, 6, 5, 5, 5}}, 
 {25, {4, 5, 5, 4, 5, 5, 5, 4, 5, 5, 5, 6, 6, 5, 5, 6, 5, 5}}, 
 {25, {4, 5, 5, 4, 5, 5, 5, 4, 5, 5, 5, 6, 6, 5, 5, 5, 6, 5}}, 
 {25, {4, 4, 5, 6, 5, 5, 5, 5, 4, 5, 6, 5, 5, 5, 5, 5, 6, 5}}, 
 {25, {4, 4, 5, 6, 5, 5, 4, 5, 5, 5, 5, 4, 5, 5, 6, 6, 6, 5}}, 
 {25, {4, 4, 5, 5, 5, 6, 5, 4, 5, 5, 6, 5, 5, 5, 5, 5, 6, 5}}, 
 {25, {4, 4, 5, 5, 5, 6, 5, 4, 4, 5, 5, 5, 6, 6, 6, 5, 5, 5}}, 
 {25, {4, 4, 5, 5, 5, 5, 6, 6, 5, 4, 4, 5, 5, 6, 6, 5, 5, 5}}, 
 {25, {4, 4, 5, 5, 5, 5, 6, 5, 5, 5, 4, 4, 5, 5, 6, 6, 6, 5}}, 
 {25, {4, 4, 5, 5, 5, 5, 6, 5, 5, 4, 5, 6, 5, 5, 5, 5, 6, 5}}, 
 {25, {4, 4, 5, 5, 5, 5, 6, 5, 4, 5, 5, 6, 5, 5, 5, 5, 6, 5}}, 
 {25, {4, 4, 5, 5, 5, 5, 5, 6, 6, 5, 4, 4, 5, 6, 6, 5, 5, 5}}, 
 {25, {4, 4, 5, 5, 5, 5, 5, 6, 5, 4, 4, 5, 6, 6, 6, 5, 5, 5}}, 
 {25, {4, 4, 5, 5, 5, 5, 5, 5, 6, 6, 5, 4, 4, 5, 5, 6, 6, 5}}, 
 {25, {4, 4, 5, 5, 5, 5, 5, 5, 6, 5, 4, 5, 6, 5, 5, 5, 6, 5}}, 
 {25, {4, 4, 5, 5, 5, 5, 5, 5, 5, 6, 6, 6, 5, 4, 4, 5, 6, 5}}, 
 {25, {4, 4, 5, 5, 5, 5, 5, 4, 4, 5, 5, 6, 6, 5, 5, 6, 6, 5}}, 
 {25, {4, 4, 5, 5, 5, 4, 5, 6, 5, 5, 6, 5, 5, 5, 5, 5, 6, 5}}, 
 {25, {4, 4, 5, 5, 4, 5, 5, 6, 6, 6, 6, 5, 4, 5, 5, 5, 5, 5}}, 
 {25, {4, 4, 5, 5, 4, 5, 5, 6, 6, 5, 5, 4, 5, 5, 6, 6, 5, 5}}, 
 {25, {4, 4, 5, 5, 4, 5, 5, 5, 6, 5, 4, 5, 5, 6, 6, 6, 5, 5}}, 
 {25, {4, 4, 5, 5, 4, 5, 5, 5, 5, 5, 6, 6, 6, 6, 5, 4, 5, 5}}, 
 {25, {4, 4, 5, 5, 4, 5, 5, 4, 5, 5, 5, 5, 6, 6, 6, 6, 5, 5}}, 
 {25, {4, 4, 5, 4, 5, 6, 5, 5, 5, 5, 5, 5, 6, 6, 5, 5, 5, 5}}, 
 {25, {4, 4, 5, 4, 5, 6, 5, 5, 5, 5, 5, 5, 5, 5, 5, 6, 6, 5}}, 
 {25, {4, 4, 5, 4, 5, 5, 6, 6, 6, 6, 5, 5, 5, 5, 5, 4, 5, 5}}, 
 {25, {4, 4, 5, 4, 5, 5, 5, 6, 6, 6, 6, 5, 5, 5, 5, 4, 5, 5}}, 
 {25, {4, 4, 5, 4, 5, 5, 5, 6, 6, 5, 5, 5, 5, 5, 5, 6, 5, 5}}, 
 {25, {4, 4, 5, 4, 5, 5, 5, 5, 5, 5, 6, 6, 5, 5, 6, 5, 5, 5}}, 
 {25, {4, 4, 5, 4, 5, 5, 5, 5, 4, 5, 5, 5, 6, 6, 6, 6, 5, 5}}, 
 {25, {3, 5, 5, 5, 6, 5, 5, 5, 5, 5, 5, 4, 5, 5, 5, 6, 6, 5}}, 
 {25, {3, 5, 5, 5, 5, 5, 6, 5, 4, 4, 5, 5, 5, 6, 6, 6, 5, 5}}, 
 {25, {3, 5, 5, 5, 5, 5, 5, 5, 5, 6, 6, 5, 5, 4, 5, 6, 5, 5}}, 
 {25, {3, 5, 5, 5, 5, 5, 5, 5, 5, 4, 5, 5, 6, 6, 5, 5, 6, 5}}, 
 {25, {3, 5, 5, 5, 5, 5, 5, 5, 4, 5, 6, 5, 5, 5, 5, 6, 6, 5}}, 
 {25, {3, 5, 5, 5, 5, 5, 5, 5, 4, 4, 5, 5, 6, 6, 6, 6, 5, 5}}, 
 {25, {3, 5, 5, 5, 5, 5, 5, 4, 5, 5, 6, 6, 5, 5, 6, 5, 5, 5}}, 
 {25, {3, 5, 5, 5, 5, 5, 5, 4, 4, 5, 5, 5, 6, 6, 6, 6, 5, 5}}, 
 {25, {3, 5, 5, 5, 5, 5, 5, 4, 4, 5, 5, 5, 5, 6, 6, 6, 6, 5}}, 
 {25, {3, 5, 5, 5, 5, 4, 5, 5, 6, 5, 5, 5, 5, 5, 5, 6, 6, 5}}, 
 {25, {3, 5, 5, 5, 4, 5, 6, 6, 5, 5, 5, 6, 5, 5, 5, 5, 5, 5}}, 
 {25, {3, 5, 5, 5, 4, 4, 5, 5, 6, 6, 6, 6, 5, 5, 5, 5, 5, 5}}, 
 {25, {3, 5, 5, 4, 5, 5, 5, 6, 6, 5, 5, 5, 5, 5, 6, 5, 5, 5}}, 
 {25, {3, 5, 5, 4, 5, 5, 5, 6, 5, 5, 5, 5, 5, 6, 6, 5, 5, 5}}, 
 {25, {3, 5, 5, 4, 5, 5, 5, 5, 5, 5, 6, 5, 5, 5, 5, 6, 6, 5}}, 
 {25, {3, 5, 5, 4, 5, 5, 5, 5, 4, 5, 5, 5, 5, 6, 6, 6, 6, 5}}, 
 {25, {1, 6, 6, 5, 5, 6, 5, 4, 4, 5, 5, 5, 5, 5, 5, 6, 6, 6}}, 
 {25, {1, 6, 6, 5, 5, 5, 5, 5, 5, 5, 4, 5, 5, 6, 5, 5, 6, 6}}, 
 {25, {1, 6, 6, 5, 5, 4, 5, 5, 5, 5, 6, 5, 4, 5, 5, 6, 6, 6}}, 
 {25, {1, 6, 6, 1, 6, 6, 5, 5, 5, 5, 5, 5, 5, 6, 6, 5, 6, 6}}, 
 {25, {4, 5, 5, 6, 6, 5, 6, 6, 5, 5, 6, 5, 4, 5, 6, 6, 5}}, 
 {25, {4, 5, 5, 6, 6, 5, 4, 5, 6, 5, 5, 6, 6, 5, 6, 6, 5}}, 
 {25, {4, 5, 5, 6, 5, 4, 5, 6, 6, 5, 6, 6, 5, 6, 6, 5, 5}}, 
 {25, {4, 5, 5, 5, 5, 6, 6, 5, 4, 5, 6, 6, 6, 5, 6, 6, 5}}, 
 {25, {4, 5, 5, 4, 5, 6, 6, 6, 5, 4, 5, 6, 6, 6, 6, 6, 5}}, 
 {25, {4, 4, 5, 6, 6, 5, 5, 4, 5, 6, 6, 6, 5, 6, 6, 6, 5}}, 
 {25, {4, 4, 5, 6, 6, 5, 5, 4, 5, 6, 6, 5, 6, 6, 6, 6, 5}}, 
 {25, {4, 4, 5, 6, 5, 5, 6, 6, 5, 4, 5, 6, 6, 6, 6, 6, 5}}, 
 {25, {4, 4, 5, 6, 5, 4, 5, 6, 6, 6, 6, 6, 5, 5, 6, 6, 5}}, 
 {25, {4, 4, 5, 4, 5, 5, 5, 6, 6, 6, 5, 6, 6, 6, 6, 6, 5}}, 
 {25, {3, 5, 6, 6, 5, 6, 6, 6, 5, 5, 5, 4, 5, 6, 6, 6, 5}}, 
 {25, {3, 5, 6, 6, 5, 5, 5, 6, 6, 6, 5, 4, 5, 6, 6, 6, 5}}, 
 {25, {3, 5, 6, 6, 5, 5, 4, 5, 6, 6, 5, 5, 6, 6, 6, 6, 5}}, 
 {25, {3, 5, 6, 5, 5, 6, 6, 6, 6, 6, 5, 5, 4, 5, 6, 6, 5}}, 
 {25, {3, 5, 6, 5, 5, 6, 5, 5, 5, 6, 5, 5, 6, 6, 6, 6, 5}}, 
 {25, {3, 5, 5, 6, 6, 5, 5, 6, 5, 5, 6, 6, 5, 5, 6, 6, 5}}, 
 {25, {3, 5, 5, 5, 6, 6, 5, 4, 5, 6, 6, 6, 6, 6, 6, 5, 5}}, 
 {25, {3, 5, 5, 5, 5, 4, 5, 6, 6, 6, 6, 6, 5, 6, 6, 6, 5}}, 
 {25, {1, 6, 6, 6, 5, 6, 6, 6, 5, 5, 3, 5, 6, 6, 6, 6, 6}}, 
 {25, {1, 6, 6, 6, 5, 6, 6, 5, 6, 6, 6, 6, 1, 6, 6, 6, 6}}, 
 {25, {1, 6, 6, 6, 5, 4, 5, 6, 6, 5, 5, 6, 6, 5, 6, 6, 6}}, 
 {25, {1, 6, 6, 6, 5, 4, 5, 5, 6, 6, 5, 6, 6, 5, 6, 6, 6}}, 
 {25, {1, 6, 6, 5, 6, 6, 6, 6, 5, 6, 6, 6, 6, 1, 6, 6, 6}}, 
 {25, {1, 6, 6, 5, 6, 6, 6, 6, 5, 5, 3, 5, 6, 6, 6, 6, 6}}, 
 {25, {1, 6, 6, 5, 6, 6, 6, 6, 1, 6, 6, 5, 6, 6, 6, 6, 6}}, 
 {25, {1, 6, 6, 5, 6, 6, 6, 1, 6, 6, 6, 5, 6, 6, 6, 6, 6}}, 
 {25, {1, 6, 6, 5, 6, 6, 5, 6, 6, 6, 6, 6, 1, 6, 6, 6, 6}}, 
 {25, {1, 6, 6, 5, 6, 6, 5, 6, 6, 6, 6, 1, 6, 6, 6, 6, 6}}, 
 {25, {1, 6, 6, 5, 6, 6, 5, 5, 6, 6, 5, 4, 5, 6, 6, 6, 6}}, 
 {25, {1, 6, 6, 5, 6, 6, 5, 4, 5, 5, 6, 6, 6, 6, 5, 6, 6}}, 
 {25, {1, 6, 6, 5, 6, 6, 5, 4, 5, 5, 6, 6, 6, 5, 6, 6, 6}}, 
 {25, {1, 6, 6, 5, 6, 6, 5, 4, 5, 5, 6, 6, 5, 6, 6, 6, 6}}, 
 {25, {1, 6, 6, 5, 4, 5, 5, 6, 6, 5, 6, 6, 5, 6, 6, 6, 6}}, 
 {25, {1, 6, 6, 1, 6, 6, 5, 6, 6, 6, 5, 6, 6, 6, 6, 6, 6}}, 
 {24, {4, 5, 5, 5, 5, 5, 5, 4, 5, 6, 5, 5, 5, 4, 5, 6, 6, 5}}, 
 {24, {4, 5, 5, 5, 5, 4, 5, 5, 5, 6, 5, 5, 5, 4, 5, 6, 6, 5}}, 
 {24, {4, 5, 5, 5, 4, 5, 6, 5, 5, 4, 5, 6, 5, 5, 5, 5, 6, 5}}, 
 {24, {4, 5, 5, 5, 4, 5, 5, 5, 5, 6, 6, 5, 5, 5, 4, 5, 6, 5}}, 
 {24, {4, 5, 5, 5, 4, 5, 5, 5, 5, 6, 6, 5, 4, 5, 5, 5, 6, 5}}, 
 {24, {4, 5, 5, 5, 4, 5, 5, 5, 5, 4, 5, 6, 6, 5, 5, 5, 6, 5}}, 
 {24, {4, 5, 5, 5, 4, 5, 5, 5, 4, 5, 5, 5, 6, 5, 5, 6, 6, 5}}, 
 {24, {4, 5, 5, 4, 5, 6, 5, 5, 4, 5, 5, 5, 5, 5, 5, 6, 6, 5}}, 
 {24, {4, 5, 5, 4, 5, 5, 6, 6, 5, 5, 5, 5, 5, 5, 4, 5, 6, 5}}, 
 {24, {4, 5, 5, 4, 5, 5, 5, 6, 6, 5, 5, 5, 4, 5, 5, 5, 6, 5}}, 
 {24, {4, 5, 5, 4, 5, 5, 5, 6, 5, 5, 5, 4, 5, 5, 5, 6, 6, 5}}, 
 {24, {4, 5, 5, 4, 5, 5, 4, 5, 5, 5, 5, 6, 6, 5, 5, 6, 5, 5}}, 
 {24, {4, 5, 4, 5, 5, 5, 5, 5, 5, 5, 6, 5, 5, 5, 5, 6, 5, 5}}, 
 {24, {4, 4, 5, 6, 5, 5, 4, 5, 5, 5, 5, 5, 5, 6, 5, 5, 6, 5}}, 
 {24, {4, 4, 5, 5, 6, 5, 4, 5, 5, 5, 6, 5, 5, 5, 5, 6, 5, 5}}, 
 {24, {4, 4, 5, 5, 5, 6, 5, 5, 5, 5, 6, 5, 5, 4, 5, 5, 6, 5}}, 
 {24, {4, 4, 5, 5, 5, 5, 6, 5, 4, 5, 6, 5, 5, 5, 5, 5, 6, 5}}, 
 {24, {4, 4, 5, 5, 5, 5, 6, 5, 4, 4, 5, 5, 6, 6, 6, 5, 5, 5}}, 
 {24, {4, 4, 5, 5, 5, 5, 5, 5, 6, 5, 5, 5, 5, 4, 5, 6, 6, 5}}, 
 {24, {4, 4, 5, 5, 5, 5, 5, 5, 6, 5, 4, 5, 6, 5, 5, 6, 5, 5}}, 
 {24, {4, 4, 5, 5, 5, 5, 5, 4, 4, 5, 6, 5, 5, 5, 6, 6, 6, 5}}, 
 {24, {4, 4, 5, 5, 5, 5, 4, 5, 5, 5, 5, 5, 6, 5, 5, 6, 6, 5}}, 
 {24, {4, 4, 5, 5, 5, 5, 4, 4, 5, 5, 5, 6, 6, 6, 5, 5, 6, 5}}, 
 {24, {4, 4, 5, 5, 5, 5, 4, 4, 5, 5, 5, 6, 6, 5, 5, 6, 6, 5}}, 
 {24, {4, 4, 5, 5, 5, 4, 5, 5, 6, 6, 6, 5, 4, 5, 5, 5, 6, 5}}, 
 {24, {4, 4, 5, 5, 5, 4, 5, 5, 4, 5, 5, 5, 5, 6, 6, 6, 6, 5}}, 
 {24, {4, 4, 5, 5, 4, 5, 6, 6, 6, 6, 5, 5, 5, 4, 5, 5, 5, 5}}, 
 {24, {4, 4, 5, 5, 4, 5, 5, 6, 6, 6, 5, 5, 6, 5, 4, 5, 5, 5}}, 
 {24, {4, 4, 5, 5, 4, 5, 5, 6, 5, 5, 5, 5, 5, 6, 5, 5, 6, 5}}, 
 {24, {4, 4, 5, 5, 4, 5, 5, 5, 5, 6, 6, 5, 4, 5, 5, 6, 6, 5}}, 
 {24, {4, 4, 5, 5, 4, 4, 5, 5, 5, 6, 6, 6, 5, 5, 5, 6, 5, 5}}, 
 {24, {4, 4, 5, 5, 4, 4, 5, 5, 5, 5, 6, 6, 6, 5, 5, 5, 6, 5}}, 
 {24, {4, 4, 5, 4, 5, 6, 6, 5, 5, 5, 6, 5, 5, 5, 5, 5, 5, 5}}, 
 {24, {4, 4, 5, 4, 5, 6, 5, 5, 5, 5, 6, 6, 5, 5, 5, 5, 5, 5}}, 
 {24, {4, 4, 5, 4, 5, 6, 5, 5, 5, 5, 5, 5, 5, 6, 6, 5, 5, 5}}, 
 {24, {4, 4, 5, 4, 5, 5, 5, 5, 5, 5, 6, 5, 5, 5, 5, 6, 6, 5}}, 
 {24, {4, 4, 5, 4, 5, 5, 5, 5, 5, 5, 5, 6, 5, 5, 5, 6, 6, 5}}, 
 {24, {4, 4, 5, 4, 4, 5, 5, 5, 5, 6, 6, 5, 5, 5, 5, 6, 6, 5}}, 
 {24, {3, 5, 6, 5, 5, 5, 5, 5, 5, 5, 4, 4, 5, 5, 6, 6, 6, 5}}, 
 {24, {3, 5, 6, 5, 5, 4, 5, 5, 5, 6, 5, 5, 5, 5, 5, 5, 6, 5}}, 
 {24, {3, 5, 6, 5, 5, 4, 5, 5, 5, 5, 5, 5, 6, 5, 5, 5, 6, 5}}, 
 {24, {3, 5, 5, 6, 6, 6, 5, 5, 4, 4, 5, 5, 5, 5, 5, 5, 6, 5}}, 
 {24, {3, 5, 5, 6, 6, 6, 5, 4, 4, 5, 5, 5, 5, 5, 5, 5, 6, 5}}, 
 {24, {3, 5, 5, 6, 5, 4, 5, 5, 5, 5, 5, 5, 5, 5, 5, 6, 6, 5}}, 
 {24, {3, 5, 5, 5, 5, 6, 5, 5, 5, 5, 4, 5, 5, 6, 6, 5, 5, 5}}, 
 {24, {3, 5, 5, 5, 5, 6, 5, 5, 4, 5, 5, 5, 5, 5, 5, 6, 6, 5}}, 
 {24, {3, 5, 5, 5, 5, 5, 5, 6, 6, 6, 5, 4, 4, 5, 5, 5, 6, 5}}, 
 {24, {3, 5, 5, 5, 5, 5, 5, 6, 6, 5, 5, 5, 4, 5, 6, 5, 5, 5}}, 
 {24, {3, 5, 5, 5, 5, 5, 5, 6, 5, 4, 5, 5, 5, 5, 6, 6, 5, 5}}, 
 {24, {3, 5, 5, 5, 5, 5, 5, 5, 5, 4, 5, 6, 5, 5, 6, 6, 5, 5}}, 
 {24, {3, 5, 5, 5, 5, 5, 5, 4, 5, 5, 5, 6, 5, 5, 5, 6, 6, 5}}, 
 {24, {3, 5, 5, 5, 5, 5, 4, 5, 6, 5, 5, 6, 6, 5, 5, 5, 5, 5}}, 
 {24, {3, 5, 5, 5, 5, 5, 4, 4, 5, 6, 6, 6, 6, 5, 5, 5, 5, 5}}, 
 {24, {3, 5, 5, 5, 5, 4, 5, 6, 5, 5, 5, 5, 5, 6, 6, 5, 5, 5}}, 
 {24, {3, 5, 5, 5, 4, 5, 6, 6, 5, 5, 5, 5, 6, 5, 5, 5, 5, 5}}, 
 {24, {3, 5, 5, 5, 4, 5, 6, 6, 5, 5, 5, 5, 5, 5, 6, 5, 5, 5}}, 
 {24, {3, 5, 5, 5, 4, 4, 5, 5, 5, 6, 6, 6, 6, 5, 5, 5, 5, 5}}, 
 {24, {3, 5, 5, 4, 5, 6, 5, 5, 5, 5, 5, 5, 6, 6, 5, 5, 5, 5}}, 
 {24, {3, 5, 5, 4, 5, 5, 6, 5, 5, 5, 5, 5, 5, 6, 6, 5, 5, 5}}, 
 {24, {3, 5, 5, 4, 4, 5, 5, 6, 6, 6, 6, 5, 5, 5, 5, 5, 5, 5}}, 
 {24, {1, 6, 6, 6, 5, 5, 4, 5, 5, 4, 4, 5, 5, 5, 6, 6, 6, 6}}, 
 {24, {1, 6, 6, 5, 5, 6, 5, 5, 5, 5, 5, 4, 4, 5, 5, 6, 6, 6}}, 
 {24, {1, 6, 6, 5, 5, 5, 6, 5, 5, 4, 4, 5, 5, 5, 5, 6, 6, 6}}, 
 {24, {1, 6, 6, 5, 5, 5, 5, 5, 5, 6, 5, 5, 6, 6, 1, 6, 6, 6}}, 
 {24, {1, 6, 6, 5, 5, 5, 5, 5, 4, 4, 5, 5, 6, 6, 5, 5, 6, 6}}, 
 {24, {1, 6, 6, 5, 5, 5, 5, 4, 5, 5, 5, 5, 4, 5, 6, 6, 6, 6}}, 
 {24, {1, 6, 6, 5, 5, 5, 4, 4, 5, 5, 5, 5, 5, 6, 6, 5, 6, 6}}, 
 {24, {1, 6, 6, 5, 5, 4, 5, 5, 5, 6, 6, 5, 5, 4, 5, 5, 6, 6}}, 
 {24, {1, 6, 6, 5, 5, 4, 4, 5, 5, 5, 6, 6, 5, 5, 5, 5, 6, 6}}, 
 {24, {1, 6, 6, 5, 4, 5, 5, 5, 6, 6, 5, 5, 5, 4, 5, 5, 6, 6}}, 
 {24, {1, 6, 6, 1, 6, 6, 6, 5, 5, 5, 5, 4, 5, 5, 6, 6, 6, 6}}, 
 {24, {1, 6, 6, 1, 6, 6, 5, 5, 5, 5, 5, 6, 6, 5, 5, 5, 6, 6}}, 
 {24, {4, 5, 5, 4, 5, 5, 6, 5, 5, 6, 6, 6, 6, 5, 6, 6, 5}}, 
 {24, {4, 5, 4, 5, 5, 6, 6, 5, 5, 6, 6, 5, 6, 6, 6, 5, 5}}, 
 {24, {4, 5, 4, 5, 5, 5, 5, 6, 6, 5, 6, 6, 6, 6, 6, 5, 5}}, 
 {24, {4, 4, 5, 6, 6, 5, 4, 5, 6, 6, 6, 6, 5, 5, 6, 6, 5}}, 
 {24, {4, 4, 5, 6, 5, 5, 6, 6, 6, 6, 5, 4, 5, 6, 6, 6, 5}}, 
 {24, {4, 4, 5, 6, 5, 4, 5, 6, 6, 6, 6, 6, 6, 5, 5, 6, 5}}, 
 {24, {4, 4, 5, 6, 5, 4, 5, 5, 6, 6, 6, 5, 6, 6, 6, 6, 5}}, 
 {24, {3, 5, 6, 6, 6, 5, 5, 6, 5, 4, 5, 5, 6, 6, 6, 6, 5}}, 
 {24, {3, 5, 6, 6, 6, 5, 4, 5, 5, 6, 6, 5, 5, 6, 6, 6, 5}}, 
 {24, {3, 5, 6, 6, 5, 5, 6, 6, 6, 5, 5, 4, 5, 6, 6, 6, 5}}, 
 {24, {3, 5, 6, 6, 5, 5, 5, 6, 5, 5, 6, 5, 5, 6, 6, 6, 5}}, 
 {24, {3, 5, 6, 6, 5, 5, 5, 6, 5, 4, 5, 6, 6, 6, 6, 6, 5}}, 
 {24, {3, 5, 6, 6, 5, 5, 4, 5, 5, 6, 6, 5, 6, 6, 6, 6, 5}}, 
 {24, {3, 5, 6, 6, 5, 4, 5, 6, 6, 5, 5, 5, 6, 6, 6, 6, 5}}, 
 {24, {3, 5, 6, 5, 5, 4, 5, 6, 6, 6, 5, 5, 6, 6, 6, 6, 5}}, 
 {24, {3, 5, 5, 6, 6, 6, 5, 5, 6, 5, 5, 6, 5, 5, 6, 6, 5}}, 
 {24, {3, 5, 5, 6, 5, 5, 6, 6, 5, 5, 6, 6, 6, 5, 5, 6, 5}}, 
 {24, {3, 5, 5, 5, 6, 6, 5, 6, 6, 6, 6, 5, 5, 5, 6, 5, 5}}, 
 {24, {3, 5, 5, 5, 5, 6, 6, 6, 6, 6, 6, 6, 5, 4, 5, 6, 5}}, 
 {24, {3, 5, 5, 5, 5, 4, 5, 6, 6, 6, 5, 6, 6, 6, 6, 6, 5}}, 
 {24, {1, 6, 6, 6, 6, 5, 4, 4, 5, 6, 6, 5, 6, 6, 6, 6, 6}}, 
 {24, {1, 6, 6, 6, 5, 3, 5, 5, 6, 6, 6, 6, 5, 6, 6, 6, 6}}, 
 {24, {1, 6, 6, 5, 6, 6, 6, 6, 6, 6, 5, 3, 5, 5, 6, 6, 6}}, 
 {24, {1, 6, 6, 5, 5, 6, 6, 5, 4, 5, 6, 6, 6, 5, 6, 6, 6}}, 
 {24, {1, 6, 6, 5, 4, 4, 5, 6, 6, 6, 6, 6, 6, 5, 6, 6, 6}}, 
 {24, {1, 6, 6, 5, 3, 5, 6, 6, 5, 5, 6, 6, 6, 6, 6, 6, 6}}, 
 {24, {1, 6, 6, 5, 3, 5, 6, 5, 5, 6, 6, 6, 6, 6, 6, 6, 6}}, 
 {23, {4, 4, 5, 5, 5, 4, 4, 5, 5, 5, 4, 5, 5, 5, 5, 5, 5, 5, 5}}, 
 {23, {4, 5, 5, 5, 5, 6, 5, 4, 5, 6, 5, 5, 4, 5, 6, 5, 5, 5}}, 
 {23, {4, 5, 5, 5, 5, 5, 5, 5, 4, 5, 6, 5, 5, 4, 5, 6, 6, 5}}, 
 {23, {4, 5, 5, 5, 5, 5, 4, 5, 5, 5, 6, 6, 5, 4, 5, 5, 6, 5}}, 
 {23, {4, 5, 5, 5, 5, 4, 5, 5, 6, 6, 5, 5, 5, 5, 4, 5, 6, 5}}, 
 {23, {4, 5, 5, 5, 5, 4, 5, 5, 5, 6, 6, 5, 5, 5, 4, 5, 6, 5}}, 
 {23, {4, 5, 5, 5, 4, 5, 5, 6, 5, 5, 5, 5, 4, 5, 6, 6, 5, 5}}, 
 {23, {4, 5, 5, 5, 4, 5, 5, 5, 6, 6, 5, 5, 5, 5, 4, 5, 6, 5}}, 
 {23, {4, 5, 5, 5, 4, 5, 5, 5, 5, 6, 6, 5, 4, 5, 6, 5, 5, 5}}, 
 {23, {4, 5, 5, 5, 4, 5, 5, 5, 5, 5, 4, 5, 6, 6, 5, 5, 6, 5}}, 
 {23, {4, 5, 5, 4, 5, 6, 5, 5, 5, 5, 5, 4, 5, 5, 5, 6, 6, 5}}, 
 {23, {4, 5, 5, 4, 5, 5, 5, 5, 5, 5, 6, 6, 5, 5, 4, 5, 6, 5}}, 
 {23, {4, 5, 5, 4, 5, 5, 5, 5, 5, 5, 6, 6, 5, 4, 5, 5, 6, 5}}, 
 {23, {4, 5, 4, 5, 5, 5, 5, 5, 5, 6, 5, 5, 5, 5, 6, 5, 5, 5}}, 
 {23, {4, 4, 5, 6, 6, 5, 5, 4, 5, 5, 5, 4, 5, 5, 5, 6, 6, 5}}, 
 {23, {4, 4, 5, 6, 5, 5, 5, 5, 5, 5, 5, 4, 5, 6, 5, 5, 6, 5}}, 
 {23, {4, 4, 5, 6, 5, 5, 5, 4, 5, 5, 5, 6, 5, 5, 5, 5, 6, 5}}, 
 {23, {4, 4, 5, 5, 6, 5, 5, 5, 5, 5, 6, 5, 4, 5, 5, 5, 6, 5}}, 
 {23, {4, 4, 5, 5, 5, 6, 6, 5, 5, 4, 5, 5, 4, 5, 5, 6, 6, 5}}, 
 {23, {4, 4, 5, 5, 5, 5, 6, 6, 5, 5, 5, 4, 4, 5, 6, 6, 5, 5}}, 
 {23, {4, 4, 5, 5, 5, 5, 6, 6, 5, 5, 5, 4, 4, 5, 5, 6, 6, 5}}, 
 {23, {4, 4, 5, 5, 5, 5, 6, 5, 5, 6, 5, 5, 5, 4, 5, 6, 5, 5}}, 
 {23, {4, 4, 5, 5, 5, 5, 5, 6, 6, 5, 5, 5, 4, 4, 5, 6, 6, 5}}, 
 {23, {4, 4, 5, 5, 5, 5, 5, 6, 6, 5, 5, 4, 4, 5, 5, 6, 6, 5}}, 
 {23, {4, 4, 5, 5, 5, 5, 5, 5, 6, 5, 5, 4, 5, 6, 5, 5, 6, 5}}, 
 {23, {4, 4, 5, 5, 5, 5, 4, 5, 5, 4, 5, 6, 6, 6, 6, 5, 5, 5}}, 
 {23, {4, 4, 5, 5, 5, 5, 4, 4, 5, 5, 6, 5, 5, 5, 6, 6, 6, 5}}, 
 {23, {4, 4, 5, 5, 5, 4, 5, 6, 5, 5, 4, 5, 5, 6, 6, 6, 5, 5}}, 
 {23, {4, 4, 5, 5, 5, 4, 4, 5, 5, 5, 5, 6, 6, 5, 5, 6, 6, 5}}, 
 {23, {4, 4, 5, 5, 4, 5, 5, 5, 6, 6, 6, 6, 5, 4, 5, 5, 5, 5}}, 
 {23, {4, 4, 5, 5, 4, 4, 5, 5, 6, 5, 5, 5, 5, 5, 6, 6, 6, 5}}, 
 {23, {4, 4, 5, 4, 5, 5, 6, 5, 5, 5, 5, 5, 5, 5, 5, 6, 6, 5}}, 
 {23, {4, 4, 5, 4, 5, 5, 5, 5, 6, 6, 5, 5, 6, 5, 5, 5, 5, 5}}, 
 {23, {4, 4, 5, 4, 5, 5, 5, 5, 6, 6, 5, 5, 5, 5, 6, 5, 5, 5}}, 
 {23, {4, 4, 5, 4, 5, 5, 5, 5, 5, 6, 5, 5, 5, 6, 6, 5, 5, 5}}, 
 {23, {3, 5, 6, 5, 5, 5, 5, 4, 5, 5, 5, 5, 6, 5, 5, 5, 6, 5}}, 
 {23, {3, 5, 5, 6, 6, 6, 5, 5, 5, 5, 5, 5, 4, 4, 5, 5, 6, 5}}, 
 {23, {3, 5, 5, 6, 6, 6, 5, 5, 5, 5, 5, 4, 4, 5, 5, 5, 6, 5}}, 
 {23, {3, 5, 5, 5, 6, 6, 5, 5, 5, 4, 5, 5, 5, 5, 5, 5, 6, 5}}, 
 {23, {3, 5, 5, 5, 6, 5, 5, 4, 5, 5, 5, 5, 5, 6, 6, 5, 5, 5}}, 
 {23, {3, 5, 5, 5, 6, 5, 5, 4, 5, 5, 5, 5, 5, 5, 5, 6, 6, 5}}, 
 {23, {3, 5, 5, 5, 6, 5, 4, 5, 5, 5, 5, 5, 5, 5, 5, 6, 6, 5}}, 
 {23, {3, 5, 5, 5, 5, 6, 5, 5, 5, 4, 5, 5, 5, 5, 5, 6, 6, 5}}, 
 {23, {3, 5, 5, 5, 5, 6, 5, 4, 5, 5, 5, 5, 5, 5, 5, 6, 6, 5}}, 
 {23, {3, 5, 5, 5, 5, 5, 5, 6, 6, 5, 5, 4, 5, 6, 5, 5, 5, 5}}, 
 {23, {3, 5, 5, 5, 5, 5, 5, 6, 5, 5, 4, 5, 5, 6, 6, 5, 5, 5}}, 
 {23, {3, 5, 5, 5, 5, 5, 5, 5, 5, 5, 6, 5, 4, 5, 6, 6, 5, 5}}, 
 {23, {3, 5, 5, 5, 5, 5, 5, 4, 5, 5, 6, 5, 5, 5, 6, 6, 5, 5}}, 
 {23, {3, 5, 5, 5, 5, 5, 4, 5, 6, 5, 5, 5, 5, 6, 6, 5, 5, 5}}, 
 {23, {3, 5, 5, 5, 5, 4, 5, 5, 5, 6, 6, 5, 5, 5, 6, 5, 5, 5}}, 
 {23, {3, 5, 5, 5, 5, 4, 5, 5, 5, 5, 6, 5, 5, 5, 5, 6, 6, 5}}, 
 {23, {3, 5, 5, 5, 5, 4, 5, 5, 5, 4, 5, 5, 6, 6, 6, 6, 5, 5}}, 
 {23, {3, 5, 5, 5, 5, 4, 4, 5, 5, 6, 6, 6, 6, 5, 5, 5, 5, 5}}, 
 {23, {3, 5, 5, 5, 4, 5, 6, 5, 5, 6, 6, 5, 5, 5, 5, 5, 5, 5}}, 
 {23, {3, 5, 5, 5, 4, 5, 5, 6, 5, 5, 6, 6, 5, 5, 5, 5, 5, 5}}, 
 {23, {3, 5, 5, 5, 4, 5, 5, 5, 5, 6, 5, 5, 5, 6, 6, 5, 5, 5}}, 
 {23, {3, 5, 5, 5, 4, 5, 5, 5, 4, 5, 5, 5, 5, 6, 6, 6, 6, 5}}, 
 {23, {3, 5, 5, 4, 5, 5, 6, 5, 5, 5, 5, 6, 6, 5, 5, 5, 5, 5}}, 
 {23, {3, 5, 5, 4, 5, 5, 5, 5, 6, 6, 6, 6, 5, 5, 4, 5, 5, 5}}, 
 {23, {3, 5, 5, 4, 5, 5, 5, 5, 6, 6, 5, 5, 6, 5, 5, 5, 5, 5}}, 
 {23, {3, 5, 5, 4, 5, 5, 5, 5, 6, 5, 5, 5, 5, 5, 6, 6, 5, 5}}, 
 {23, {3, 5, 5, 4, 5, 5, 5, 5, 5, 5, 6, 6, 5, 5, 6, 5, 5, 5}}, 
 {23, {3, 5, 5, 4, 4, 5, 5, 5, 5, 5, 5, 6, 6, 6, 6, 5, 5, 5}}, 
 {23, {1, 6, 6, 6, 5, 5, 5, 5, 3, 5, 5, 5, 5, 5, 5, 6, 6, 6}}, 
 {23, {1, 6, 6, 5, 5, 5, 5, 5, 6, 5, 5, 4, 4, 5, 5, 6, 6, 6}}, 
 {23, {1, 6, 6, 5, 5, 5, 5, 5, 5, 5, 6, 5, 4, 4, 5, 6, 6, 6}}, 
 {23, {1, 6, 6, 5, 5, 5, 5, 5, 5, 5, 5, 6, 5, 3, 5, 6, 6, 6}}, 
 {23, {1, 6, 6, 5, 5, 5, 5, 5, 5, 4, 5, 5, 4, 5, 6, 6, 6, 6}}, 
 {23, {1, 6, 6, 5, 5, 5, 5, 5, 5, 3, 5, 5, 5, 5, 6, 6, 6, 6}}, 
 {23, {1, 6, 6, 5, 5, 5, 5, 3, 5, 5, 5, 5, 5, 5, 6, 6, 6, 6}}, 
 {23, {1, 6, 6, 5, 5, 4, 5, 5, 6, 5, 4, 5, 5, 5, 5, 6, 6, 6}}, 
 {23, {1, 6, 6, 5, 5, 4, 5, 5, 5, 6, 5, 4, 5, 5, 5, 6, 6, 6}}, 
 {23, {1, 6, 6, 5, 5, 4, 5, 5, 5, 5, 5, 6, 5, 4, 5, 6, 6, 6}}, 
 {23, {1, 6, 6, 5, 5, 4, 5, 5, 5, 5, 4, 5, 5, 6, 6, 5, 6, 6}}, 
 {23, {1, 6, 6, 5, 4, 4, 5, 5, 6, 6, 5, 5, 5, 5, 5, 5, 6, 6}}, 
 {23, {1, 6, 6, 5, 4, 4, 5, 5, 5, 5, 6, 6, 5, 5, 5, 5, 6, 6}}, 
 {23, {4, 5, 5, 6, 5, 4, 5, 6, 5, 5, 6, 6, 6, 5, 6, 6, 5}}, 
 {23, {4, 5, 5, 6, 5, 4, 5, 6, 5, 5, 6, 6, 5, 6, 6, 6, 5}}, 
 {23, {4, 5, 4, 5, 5, 6, 6, 5, 6, 6, 5, 5, 6, 6, 6, 5, 5}}, 
 {23, {4, 5, 4, 5, 5, 6, 6, 5, 5, 6, 6, 6, 5, 6, 6, 5, 5}}, 
 {23, {4, 4, 5, 5, 6, 6, 6, 5, 4, 5, 6, 6, 6, 5, 6, 6, 5}}, 
 {23, {4, 4, 5, 5, 6, 5, 5, 6, 5, 5, 6, 6, 5, 6, 6, 6, 5}}, 
 {23, {4, 4, 5, 5, 6, 5, 5, 5, 6, 6, 5, 6, 6, 5, 6, 6, 5}}, 
 {23, {4, 4, 5, 5, 5, 6, 5, 5, 6, 6, 5, 6, 6, 5, 6, 6, 5}}, 
 {23, {4, 4, 5, 4, 5, 6, 6, 6, 5, 6, 6, 6, 6, 6, 5, 5, 5}}, 
 {23, {4, 4, 5, 4, 5, 6, 6, 6, 5, 5, 6, 6, 6, 6, 6, 5, 5}}, 
 {23, {3, 5, 6, 6, 5, 6, 6, 5, 5, 5, 4, 5, 6, 6, 6, 6, 5}}, 
 {23, {3, 5, 6, 6, 5, 5, 6, 5, 4, 5, 5, 6, 6, 6, 6, 6, 5}}, 
 {23, {3, 5, 6, 6, 5, 5, 5, 6, 6, 5, 5, 6, 5, 5, 6, 6, 5}}, 
 {23, {3, 5, 6, 6, 5, 4, 5, 5, 6, 6, 6, 6, 5, 5, 6, 6, 5}}, 
 {23, {3, 5, 6, 5, 5, 5, 6, 6, 5, 4, 5, 6, 6, 6, 6, 6, 5}}, 
 {23, {3, 5, 6, 5, 4, 5, 6, 6, 6, 6, 5, 5, 5, 6, 6, 6, 5}}, 
 {23, {3, 5, 5, 6, 6, 6, 5, 6, 6, 6, 6, 5, 4, 5, 5, 6, 5}}, 
 {23, {3, 5, 5, 6, 6, 6, 5, 4, 5, 6, 6, 6, 5, 5, 6, 6, 5}}, 
 {23, {3, 5, 5, 5, 6, 6, 6, 6, 6, 6, 5, 5, 4, 5, 6, 6, 5}}, 
 {23, {3, 5, 5, 5, 5, 6, 6, 6, 5, 6, 6, 6, 5, 5, 6, 5, 5}}, 
 {23, {1, 6, 6, 6, 5, 6, 6, 6, 6, 5, 3, 5, 5, 6, 6, 6, 6}}, 
 {23, {1, 6, 6, 5, 6, 6, 6, 6, 6, 1, 6, 6, 6, 5, 6, 6, 6}}, 
 {23, {1, 6, 6, 5, 6, 6, 6, 5, 5, 6, 5, 4, 5, 6, 6, 6, 6}}, 
 {23, {1, 6, 6, 5, 6, 6, 6, 1, 6, 6, 6, 6, 5, 6, 6, 6, 6}}, 
 {23, {1, 6, 6, 5, 6, 6, 5, 6, 6, 6, 6, 6, 6, 1, 6, 6, 6}}, 
 {23, {1, 6, 6, 5, 5, 6, 5, 5, 6, 5, 5, 6, 6, 6, 5, 6, 6}}, 
 {23, {1, 6, 6, 5, 5, 4, 5, 6, 6, 5, 6, 6, 5, 6, 6, 6, 6}}, 
 {23, {1, 6, 6, 5, 4, 5, 6, 5, 5, 6, 6, 6, 5, 6, 6, 6, 6}}, 
 {22, {4, 4, 5, 5, 5, 5, 4, 4, 5, 5, 5, 5, 4, 5, 5, 5, 5, 5, 5}}, 
 {22, {4, 4, 5, 5, 5, 4, 4, 5, 5, 5, 5, 5, 4, 5, 5, 5, 5, 5, 5}}, 
 {22, {4, 4, 5, 5, 4, 5, 5, 5, 4, 4, 5, 5, 5, 5, 5, 5, 5, 5, 5}}, 
 {22, {4, 5, 5, 5, 5, 5, 5, 4, 5, 5, 5, 6, 6, 5, 4, 5, 6, 5}}, 
 {22, {4, 5, 5, 5, 5, 5, 4, 5, 5, 6, 5, 5, 4, 5, 5, 6, 6, 5}}, 
 {22, {4, 5, 5, 5, 4, 5, 5, 6, 5, 5, 5, 5, 5, 4, 5, 6, 6, 5}}, 
 {22, {4, 5, 5, 5, 4, 5, 5, 5, 5, 6, 5, 5, 5, 4, 5, 6, 6, 5}}, 
 {22, {4, 5, 5, 4, 5, 6, 5, 5, 5, 5, 5, 5, 4, 5, 5, 6, 6, 5}}, 
 {22, {4, 5, 5, 4, 5, 5, 5, 6, 6, 5, 4, 5, 5, 5, 5, 5, 6, 5}}, 
 {22, {4, 5, 5, 4, 5, 5, 5, 5, 6, 6, 5, 5, 4, 5, 5, 5, 6, 5}}, 
 {22, {4, 5, 5, 4, 5, 5, 5, 5, 6, 5, 5, 5, 4, 5, 5, 6, 6, 5}}, 
 {22, {4, 5, 5, 4, 5, 5, 5, 5, 6, 5, 5, 4, 5, 5, 5, 6, 6, 5}}, 
 {22, {4, 5, 5, 4, 5, 5, 5, 5, 5, 6, 6, 5, 5, 5, 4, 5, 6, 5}}, 
 {22, {4, 5, 5, 4, 5, 5, 5, 5, 4, 5, 6, 6, 5, 5, 5, 6, 5, 5}}, 
 {22, {4, 5, 5, 4, 5, 5, 5, 5, 4, 5, 5, 6, 6, 5, 5, 6, 5, 5}}, 
 {22, {4, 5, 5, 4, 5, 5, 5, 4, 5, 6, 6, 5, 5, 5, 6, 5, 5, 5}}, 
 {22, {4, 5, 5, 4, 5, 5, 5, 4, 5, 5, 5, 6, 5, 5, 5, 6, 6, 5}}, 
 {22, {4, 5, 5, 4, 5, 5, 4, 5, 5, 6, 5, 5, 5, 5, 6, 6, 5, 5}}, 
 {22, {4, 5, 5, 4, 5, 5, 4, 5, 5, 5, 6, 6, 5, 5, 5, 6, 5, 5}}, 
 {22, {4, 5, 5, 4, 5, 5, 4, 5, 5, 5, 5, 6, 6, 6, 5, 5, 5, 5}}, 
 {22, {4, 5, 4, 5, 5, 5, 5, 5, 5, 4, 5, 5, 6, 6, 6, 5, 5, 5}}, 
 {22, {4, 4, 5, 6, 5, 5, 5, 5, 5, 4, 5, 5, 5, 6, 5, 5, 6, 5}}, 
 {22, {4, 4, 5, 6, 5, 4, 5, 6, 5, 5, 5, 5, 5, 5, 5, 5, 6, 5}}, 
 {22, {4, 4, 5, 5, 5, 6, 5, 5, 5, 4, 5, 5, 5, 6, 5, 5, 6, 5}}, 
 {22, {4, 4, 5, 5, 5, 5, 6, 5, 5, 6, 5, 4, 5, 5, 5, 5, 6, 5}}, 
 {22, {4, 4, 5, 5, 5, 5, 6, 5, 4, 5, 6, 5, 5, 5, 5, 6, 5, 5}}, 
 {22, {4, 4, 5, 5, 5, 5, 5, 6, 6, 6, 5, 4, 4, 5, 6, 5, 5, 5}}, 
 {22, {4, 4, 5, 5, 5, 5, 5, 6, 6, 6, 5, 4, 4, 5, 5, 5, 6, 5}}, 
 {22, {4, 4, 5, 5, 5, 5, 5, 5, 5, 6, 6, 5, 4, 4, 5, 6, 6, 5}}, 
 {22, {4, 4, 5, 5, 5, 5, 5, 5, 5, 5, 4, 4, 5, 6, 6, 6, 6, 5}}, 
 {22, {4, 4, 5, 5, 5, 5, 5, 4, 5, 5, 4, 5, 6, 6, 6, 6, 5, 5}}, 
 {22, {4, 4, 5, 5, 5, 5, 4, 5, 6, 5, 5, 5, 6, 5, 5, 5, 6, 5}}, 
 {22, {4, 4, 5, 5, 5, 4, 5, 5, 6, 6, 6, 5, 5, 5, 4, 5, 6, 5}}, 
 {22, {4, 4, 5, 5, 5, 4, 5, 5, 6, 6, 5, 4, 5, 5, 6, 6, 5, 5}}, 
 {22, {4, 4, 5, 5, 4, 5, 6, 5, 5, 5, 5, 6, 5, 5, 5, 6, 5, 5}}, 
 {22, {4, 4, 5, 5, 4, 5, 5, 6, 6, 5, 5, 4, 5, 5, 5, 6, 6, 5}}, 
 {22, {4, 4, 5, 5, 4, 5, 5, 6, 5, 5, 6, 5, 5, 5, 5, 5, 6, 5}}, 
 {22, {4, 4, 5, 5, 4, 5, 5, 4, 5, 5, 5, 6, 6, 6, 6, 5, 5, 5}}, 
 {22, {4, 4, 5, 5, 4, 5, 5, 4, 5, 5, 5, 5, 5, 6, 6, 6, 6, 5}}, 
 {22, {4, 4, 5, 4, 5, 5, 5, 6, 6, 5, 5, 6, 5, 5, 5, 5, 5, 5}}, 
 {22, {4, 4, 5, 4, 5, 5, 5, 6, 6, 5, 5, 5, 6, 5, 5, 5, 5, 5}}, 
 {22, {3, 5, 6, 5, 5, 5, 5, 5, 5, 4, 5, 5, 5, 6, 5, 5, 6, 5}}, 
 {22, {3, 5, 6, 5, 5, 5, 5, 5, 4, 5, 5, 5, 5, 6, 5, 5, 6, 5}}, 
 {22, {3, 5, 6, 5, 5, 5, 5, 5, 4, 4, 5, 5, 5, 5, 6, 6, 6, 5}}, 
 {22, {3, 5, 6, 5, 5, 5, 4, 5, 5, 5, 5, 4, 5, 5, 6, 6, 6, 5}}, 
 {22, {3, 5, 6, 5, 5, 4, 5, 5, 6, 5, 5, 5, 5, 5, 5, 5, 6, 5}}, 
 {22, {3, 5, 5, 5, 6, 5, 5, 5, 5, 5, 5, 5, 4, 5, 6, 6, 5, 5}}, 
 {22, {3, 5, 5, 5, 6, 5, 5, 5, 4, 5, 5, 5, 5, 5, 6, 6, 5, 5}}, 
 {22, {3, 5, 5, 5, 5, 6, 6, 6, 5, 5, 5, 5, 5, 4, 4, 5, 6, 5}}, 
 {22, {3, 5, 5, 5, 5, 5, 6, 6, 5, 5, 4, 4, 5, 5, 6, 6, 5, 5}}, 
 {22, {3, 5, 5, 5, 5, 5, 5, 5, 6, 6, 5, 5, 4, 5, 6, 5, 5, 5}}, 
 {22, {3, 5, 5, 5, 5, 5, 5, 5, 5, 6, 6, 5, 5, 5, 4, 5, 6, 5}}, 
 {22, {3, 5, 5, 5, 5, 5, 5, 5, 5, 6, 6, 5, 4, 5, 6, 5, 5, 5}}, 
 {22, {3, 5, 5, 5, 5, 5, 5, 5, 5, 6, 6, 5, 4, 5, 5, 5, 6, 5}}, 
 {22, {3, 5, 5, 5, 5, 5, 5, 5, 5, 4, 5, 6, 6, 5, 5, 5, 6, 5}}, 
 {22, {3, 5, 5, 5, 5, 5, 5, 5, 5, 4, 5, 6, 5, 5, 5, 6, 6, 5}}, 
 {22, {3, 5, 5, 5, 5, 5, 5, 5, 4, 5, 6, 6, 5, 5, 5, 5, 6, 5}}, 
 {22, {3, 5, 5, 5, 5, 5, 5, 5, 4, 5, 5, 6, 5, 5, 6, 6, 5, 5}}, 
 {22, {3, 5, 5, 5, 5, 4, 5, 6, 5, 5, 6, 6, 5, 5, 5, 5, 5, 5}}, 
 {22, {3, 5, 5, 5, 5, 4, 5, 6, 5, 5, 5, 5, 6, 6, 5, 5, 5, 5}}, 
 {22, {3, 5, 5, 5, 5, 4, 5, 6, 5, 5, 5, 5, 5, 5, 5, 6, 6, 5}}, 
 {22, {3, 5, 5, 5, 5, 4, 5, 5, 6, 6, 5, 5, 5, 6, 5, 5, 5, 5}}, 
 {22, {3, 5, 5, 5, 5, 4, 4, 5, 6, 6, 6, 6, 5, 5, 5, 5, 5, 5}}, 
 {22, {3, 5, 5, 5, 4, 5, 5, 6, 5, 5, 5, 5, 6, 6, 5, 5, 5, 5}}, 
 {22, {3, 5, 5, 5, 4, 5, 5, 5, 5, 4, 5, 5, 6, 6, 6, 6, 5, 5}}, 
 {22, {3, 5, 5, 4, 5, 6, 5, 5, 5, 5, 5, 6, 6, 5, 5, 5, 5, 5}}, 
 {22, {3, 5, 5, 4, 5, 5, 5, 5, 5, 6, 6, 5, 5, 5, 6, 5, 5, 5}}, 
 {22, {3, 5, 5, 4, 5, 5, 5, 4, 5, 5, 5, 5, 5, 6, 6, 6, 6, 5}}, 
 {22, {3, 5, 5, 4, 4, 5, 5, 5, 5, 6, 6, 6, 5, 5, 5, 5, 6, 5}}, 
 {22, {1, 6, 6, 6, 5, 5, 5, 3, 5, 5, 5, 5, 5, 5, 5, 6, 6, 6}}, 
 {22, {1, 6, 6, 5, 5, 5, 5, 6, 5, 5, 4, 4, 5, 5, 5, 6, 6, 6}}, 
 {22, {1, 6, 6, 5, 5, 5, 4, 5, 5, 6, 5, 5, 5, 5, 5, 5, 6, 6}}, 
 {22, {1, 6, 6, 5, 5, 4, 5, 6, 5, 5, 5, 4, 5, 5, 5, 6, 6, 6}}, 
 {22, {1, 6, 6, 5, 5, 4, 5, 5, 4, 5, 5, 6, 6, 5, 5, 5, 6, 6}}, 
 {22, {1, 6, 6, 5, 4, 4, 5, 5, 5, 6, 5, 5, 5, 5, 5, 6, 6, 6}}, 
 {22, {1, 6, 6, 5, 4, 4, 5, 5, 5, 5, 6, 5, 5, 5, 5, 6, 6, 6}}, 
 {22, {1, 6, 6, 1, 6, 6, 6, 5, 5, 5, 5, 5, 4, 5, 6, 6, 6, 6}}, 
 {22, {4, 5, 5, 6, 6, 5, 6, 6, 5, 6, 6, 5, 5, 4, 5, 6, 5}}, 
 {22, {4, 5, 5, 4, 5, 6, 6, 5, 4, 5, 6, 6, 6, 6, 6, 6, 5}}, 
 {22, {4, 5, 4, 5, 5, 5, 5, 6, 6, 6, 6, 5, 5, 6, 6, 6, 5}}, 
 {22, {4, 4, 5, 6, 6, 6, 5, 5, 4, 5, 6, 6, 5, 6, 6, 6, 5}}, 
 {22, {4, 4, 5, 6, 6, 5, 6, 6, 6, 5, 4, 5, 5, 6, 6, 6, 5}}, 
 {22, {4, 4, 5, 6, 6, 5, 4, 5, 5, 6, 6, 5, 6, 6, 6, 6, 5}}, 
 {22, {4, 4, 5, 6, 5, 5, 4, 5, 6, 6, 6, 6, 6, 5, 6, 6, 5}}, 
 {22, {4, 4, 5, 5, 6, 6, 6, 6, 5, 4, 5, 6, 6, 5, 6, 6, 5}}, 
 {22, {4, 4, 5, 5, 6, 6, 6, 5, 6, 6, 6, 6, 5, 4, 5, 6, 5}}, 
 {22, {4, 4, 5, 5, 6, 6, 6, 5, 6, 6, 6, 5, 4, 5, 6, 6, 5}}, 
 {22, {4, 4, 5, 5, 6, 6, 5, 6, 6, 6, 6, 6, 5, 4, 5, 6, 5}}, 
 {22, {4, 4, 5, 4, 5, 6, 6, 5, 5, 5, 6, 6, 6, 6, 6, 6, 5}}, 
 {22, {3, 5, 6, 6, 6, 5, 5, 4, 4, 5, 6, 6, 6, 6, 6, 6, 5}}, 
 {22, {3, 5, 6, 6, 5, 5, 4, 5, 6, 6, 6, 6, 5, 5, 6, 6, 5}}, 
 {22, {3, 5, 6, 6, 5, 4, 5, 5, 6, 6, 6, 5, 5, 6, 6, 6, 5}}, 
 {22, {3, 5, 6, 6, 5, 4, 4, 5, 5, 6, 6, 6, 6, 6, 6, 6, 5}}, 
 {22, {3, 5, 6, 5, 5, 4, 5, 6, 6, 5, 5, 6, 6, 6, 6, 6, 5}}, 
 {22, {3, 5, 6, 5, 4, 5, 6, 6, 6, 6, 6, 5, 5, 5, 6, 6, 5}}, 
 {22, {3, 5, 5, 6, 6, 6, 6, 5, 4, 5, 6, 6, 6, 5, 5, 6, 5}}, 
 {22, {3, 5, 5, 6, 6, 6, 5, 6, 6, 5, 5, 4, 5, 6, 6, 6, 5}}, 
 {22, {3, 5, 5, 6, 6, 6, 5, 5, 4, 5, 6, 6, 6, 5, 6, 6, 5}}, 
 {22, {3, 5, 5, 6, 6, 5, 6, 6, 6, 5, 4, 5, 5, 6, 6, 6, 5}}, 
 {22, {3, 5, 5, 6, 5, 5, 5, 5, 6, 5, 5, 6, 6, 6, 6, 6, 5}}, 
 {22, {3, 5, 5, 5, 6, 5, 5, 6, 6, 6, 6, 6, 5, 5, 6, 5, 5}}, 
 {22, {3, 5, 5, 5, 5, 4, 5, 6, 6, 6, 6, 5, 6, 6, 6, 6, 5}}, 
 {22, {3, 5, 5, 4, 5, 5, 5, 6, 6, 5, 6, 6, 6, 6, 6, 6, 5}}, 
 {22, {3, 5, 5, 4, 4, 5, 6, 6, 6, 6, 6, 6, 6, 6, 6, 5, 5}}, 
 {22, {1, 6, 6, 6, 6, 5, 6, 6, 5, 4, 4, 5, 6, 6, 6, 6, 6}}, 
 {22, {1, 6, 6, 6, 6, 5, 5, 3, 5, 6, 6, 6, 5, 6, 6, 6, 6}}, 
 {22, {1, 6, 6, 6, 5, 6, 6, 6, 5, 3, 5, 5, 6, 6, 6, 6, 6}}, 
 {22, {1, 6, 6, 6, 5, 5, 6, 6, 5, 4, 5, 6, 6, 5, 6, 6, 6}}, 
 {22, {1, 6, 6, 6, 5, 5, 6, 5, 4, 5, 6, 6, 5, 6, 6, 6, 6}}, 
 {22, {1, 6, 6, 6, 5, 4, 5, 6, 5, 5, 6, 6, 6, 5, 6, 6, 6}}, 
 {22, {1, 6, 6, 5, 6, 6, 6, 6, 1, 6, 6, 6, 6, 6, 5, 6, 6}}, 
 {22, {1, 6, 6, 5, 6, 6, 5, 6, 6, 5, 5, 4, 5, 6, 6, 6, 6}}, 
 {22, {1, 6, 6, 5, 5, 4, 5, 6, 6, 6, 5, 6, 6, 6, 5, 6, 6}}, 
 {22, {1, 6, 6, 5, 4, 5, 6, 5, 5, 6, 6, 5, 6, 6, 6, 6, 6}}, 
 {22, {1, 6, 6, 5, 4, 4, 5, 6, 6, 6, 6, 6, 5, 6, 6, 6, 6}}, 
 {22, {1, 6, 6, 5, 3, 5, 5, 6, 6, 6, 6, 6, 6, 5, 6, 6, 6}}, 
 {21, {4, 4, 5, 5, 5, 4, 5, 5, 5, 5, 5, 4, 4, 5, 5, 5, 5, 5, 5}}, 
 {21, {4, 5, 5, 5, 6, 5, 4, 5, 5, 6, 5, 5, 5, 4, 5, 6, 5, 5}}, 
 {21, {4, 5, 5, 5, 5, 5, 5, 4, 5, 6, 5, 5, 4, 5, 5, 6, 6, 5}}, 
 {21, {4, 5, 5, 5, 5, 5, 4, 5, 6, 5, 5, 4, 5, 5, 5, 6, 6, 5}}, 
 {21, {4, 5, 5, 5, 5, 4, 5, 5, 5, 5, 4, 5, 6, 5, 5, 6, 6, 5}}, 
 {21, {4, 5, 5, 5, 4, 5, 5, 6, 5, 5, 5, 4, 5, 5, 5, 6, 6, 5}}, 
 {21, {4, 5, 5, 5, 4, 5, 5, 5, 5, 5, 4, 5, 6, 5, 5, 6, 6, 5}}, 
 {21, {4, 5, 5, 4, 5, 5, 6, 5, 5, 5, 4, 5, 5, 5, 5, 6, 6, 5}}, 
 {21, {4, 5, 5, 4, 5, 5, 5, 5, 6, 6, 5, 5, 5, 4, 5, 5, 6, 5}}, 
 {21, {4, 5, 5, 4, 5, 5, 5, 5, 6, 6, 5, 4, 5, 5, 5, 5, 6, 5}}, 
 {21, {4, 5, 5, 4, 5, 5, 5, 5, 5, 6, 5, 4, 5, 6, 6, 5, 5, 5}}, 
 {21, {4, 5, 5, 4, 5, 5, 5, 5, 5, 5, 6, 6, 5, 4, 5, 6, 5, 5}}, 
 {21, {4, 5, 5, 4, 5, 5, 5, 4, 5, 6, 5, 5, 5, 5, 5, 6, 6, 5}}, 
 {21, {4, 5, 4, 5, 5, 5, 5, 5, 5, 5, 5, 5, 5, 5, 5, 6, 6, 5}}, 
 {21, {4, 5, 4, 5, 5, 5, 5, 5, 4, 5, 5, 5, 6, 6, 6, 5, 5, 5}}, 
 {21, {4, 4, 5, 6, 5, 4, 5, 5, 5, 6, 5, 5, 5, 5, 5, 5, 6, 5}}, 
 {21, {4, 4, 5, 6, 5, 4, 5, 5, 5, 5, 5, 6, 5, 5, 5, 5, 6, 5}}, 
 {21, {4, 4, 5, 5, 6, 6, 5, 5, 5, 4, 5, 5, 4, 5, 5, 6, 6, 5}}, 
 {21, {4, 4, 5, 5, 6, 5, 5, 5, 5, 5, 5, 6, 5, 4, 5, 5, 6, 5}}, 
 {21, {4, 4, 5, 5, 6, 5, 4, 5, 6, 5, 5, 5, 5, 5, 5, 5, 6, 5}}, 
 {21, {4, 4, 5, 5, 6, 5, 4, 5, 5, 5, 5, 5, 5, 6, 5, 5, 6, 5}}, 
 {21, {4, 4, 5, 5, 5, 6, 5, 5, 5, 6, 5, 5, 5, 4, 5, 6, 5, 5}}, 
 {21, {4, 4, 5, 5, 5, 6, 5, 5, 5, 5, 5, 6, 5, 5, 4, 5, 6, 5}}, 
 {21, {4, 4, 5, 5, 5, 6, 5, 5, 5, 4, 5, 6, 5, 5, 5, 6, 5, 5}}, 
 {21, {4, 4, 5, 5, 5, 6, 5, 4, 5, 6, 5, 5, 5, 5, 5, 6, 5, 5}}, 
 {21, {4, 4, 5, 5, 5, 6, 5, 4, 5, 5, 5, 6, 5, 5, 5, 5, 6, 5}}, 
 {21, {4, 4, 5, 5, 5, 6, 5, 4, 5, 5, 5, 5, 5, 6, 5, 5, 6, 5}}, 
 {21, {4, 4, 5, 5, 5, 5, 5, 6, 5, 5, 6, 5, 5, 5, 4, 5, 6, 5}}, 
 {21, {4, 4, 5, 5, 5, 5, 5, 6, 5, 5, 5, 4, 5, 6, 5, 5, 6, 5}}, 
 {21, {4, 4, 5, 5, 5, 5, 5, 6, 5, 5, 4, 5, 6, 5, 5, 5, 6, 5}}, 
 {21, {4, 4, 5, 5, 5, 5, 5, 5, 6, 6, 5, 4, 4, 5, 6, 6, 5, 5}}, 
 {21, {4, 4, 5, 5, 5, 5, 4, 5, 6, 5, 5, 6, 5, 5, 5, 5, 6, 5}}, 
 {21, {4, 4, 5, 5, 5, 5, 4, 5, 5, 6, 6, 6, 5, 5, 4, 5, 6, 5}}, 
 {21, {4, 4, 5, 5, 5, 5, 4, 5, 5, 5, 6, 5, 5, 6, 5, 5, 6, 5}}, 
 {21, {4, 4, 5, 5, 5, 4, 5, 5, 6, 6, 6, 5, 5, 4, 5, 6, 5, 5}}, 
 {21, {4, 4, 5, 5, 5, 4, 5, 5, 6, 6, 5, 5, 4, 5, 5, 6, 6, 5}}, 
 {21, {4, 4, 5, 5, 5, 4, 4, 5, 5, 6, 5, 5, 5, 5, 6, 6, 6, 5}}, 
 {21, {4, 4, 5, 5, 5, 4, 4, 5, 5, 5, 6, 6, 5, 5, 5, 6, 6, 5}}, 
 {21, {4, 4, 5, 5, 4, 5, 6, 6, 6, 6, 5, 4, 5, 5, 5, 5, 5, 5}}, 
 {21, {4, 4, 5, 5, 4, 5, 5, 6, 6, 6, 5, 5, 4, 5, 5, 5, 6, 5}}, 
 {21, {4, 4, 5, 5, 4, 5, 5, 6, 5, 5, 5, 6, 5, 5, 5, 5, 6, 5}}, 
 {21, {4, 4, 5, 5, 4, 5, 5, 5, 6, 6, 6, 5, 5, 4, 5, 6, 5, 5}}, 
 {21, {4, 4, 5, 5, 4, 5, 5, 5, 6, 6, 5, 5, 4, 5, 5, 6, 6, 5}}, 
 {21, {4, 4, 5, 5, 4, 5, 5, 5, 5, 6, 5, 4, 5, 5, 6, 6, 6, 5}}, 
 {21, {4, 4, 5, 5, 4, 4, 5, 5, 6, 6, 5, 5, 5, 5, 6, 6, 5, 5}}, 
 {21, {4, 4, 5, 5, 4, 4, 5, 5, 6, 5, 5, 5, 5, 6, 6, 6, 5, 5}}, 
 {21, {4, 4, 5, 4, 5, 6, 6, 5, 5, 5, 5, 6, 5, 5, 5, 5, 5, 5}}, 
 {21, {4, 4, 5, 4, 5, 5, 5, 6, 5, 5, 6, 6, 5, 5, 5, 5, 5, 5}}, 
 {21, {4, 4, 5, 4, 5, 5, 5, 5, 6, 6, 6, 6, 5, 5, 5, 4, 5, 5}}, 
 {21, {4, 4, 5, 4, 5, 5, 5, 5, 6, 5, 5, 5, 5, 6, 6, 5, 5, 5}}, 
 {21, {4, 4, 5, 4, 5, 5, 5, 5, 5, 6, 6, 6, 6, 5, 4, 5, 5, 5}}, 
 {21, {4, 4, 5, 4, 5, 5, 5, 5, 5, 5, 6, 6, 5, 5, 5, 6, 5, 5}}, 
 {21, {4, 4, 5, 4, 5, 5, 5, 5, 5, 5, 5, 6, 6, 5, 5, 6, 5, 5}}, 
 {21, {4, 4, 5, 4, 4, 5, 5, 5, 5, 5, 6, 6, 6, 5, 5, 5, 6, 5}}, 
 {21, {4, 4, 5, 4, 4, 5, 5, 5, 5, 5, 6, 6, 5, 5, 6, 6, 5, 5}}, 
 {21, {3, 5, 6, 5, 5, 5, 5, 4, 4, 5, 5, 5, 5, 5, 6, 6, 6, 5}}, 
 {21, {3, 5, 6, 5, 5, 5, 4, 4, 5, 5, 5, 5, 5, 5, 6, 6, 6, 5}}, 
 {21, {3, 5, 5, 6, 6, 6, 5, 5, 5, 4, 5, 5, 5, 4, 5, 5, 6, 5}}, 
 {21, {3, 5, 5, 6, 6, 6, 5, 5, 5, 4, 4, 5, 5, 5, 5, 5, 6, 5}}, 
 {21, {3, 5, 5, 5, 6, 5, 5, 5, 5, 5, 5, 5, 5, 4, 5, 6, 6, 5}}, 
 {21, {3, 5, 5, 5, 6, 5, 5, 5, 4, 5, 5, 5, 5, 6, 6, 5, 5, 5}}, 
 {21, {3, 5, 5, 5, 6, 5, 4, 5, 5, 5, 5, 5, 5, 6, 6, 5, 5, 5}}, 
 {21, {3, 5, 5, 5, 5, 6, 6, 5, 5, 5, 4, 5, 5, 5, 6, 5, 5, 5}}, 
 {21, {3, 5, 5, 5, 5, 6, 5, 5, 5, 5, 5, 5, 5, 4, 5, 6, 6, 5}}, 
 {21, {3, 5, 5, 5, 5, 6, 5, 5, 5, 4, 5, 5, 5, 6, 6, 5, 5, 5}}, 
 {21, {3, 5, 5, 5, 5, 5, 5, 5, 5, 6, 5, 5, 5, 4, 5, 6, 6, 5}}, 
 {21, {3, 5, 5, 5, 5, 5, 5, 5, 5, 5, 6, 5, 5, 4, 5, 6, 6, 5}}, 
 {21, {3, 5, 5, 5, 5, 5, 5, 5, 5, 4, 5, 5, 6, 5, 5, 6, 6, 5}}, 
 {21, {3, 5, 5, 5, 5, 5, 5, 5, 4, 5, 6, 5, 5, 5, 6, 6, 5, 5}}, 
 {21, {3, 5, 5, 5, 5, 5, 5, 4, 5, 5, 5, 6, 5, 5, 6, 6, 5, 5}}, 
 {21, {3, 5, 5, 5, 5, 5, 4, 5, 5, 4, 5, 5, 6, 6, 6, 6, 5, 5}}, 
 {21, {3, 5, 5, 5, 5, 5, 4, 4, 5, 5, 6, 6, 6, 5, 5, 5, 6, 5}}, 
 {21, {3, 5, 5, 5, 5, 4, 5, 6, 6, 5, 5, 5, 5, 5, 6, 5, 5, 5}}, 
 {21, {3, 5, 5, 5, 5, 4, 5, 6, 5, 5, 5, 5, 5, 5, 6, 6, 5, 5}}, 
 {21, {3, 5, 5, 5, 5, 4, 5, 5, 6, 6, 5, 5, 6, 5, 5, 5, 5, 5}}, 
 {21, {3, 5, 5, 5, 5, 4, 5, 5, 5, 5, 6, 5, 5, 6, 6, 5, 5, 5}}, 
 {21, {3, 5, 5, 5, 5, 4, 5, 5, 5, 5, 5, 6, 5, 5, 5, 6, 6, 5}}, 
 {21, {3, 5, 5, 5, 5, 4, 5, 5, 5, 4, 5, 5, 5, 6, 6, 6, 6, 5}}, 
 {21, {3, 5, 5, 5, 4, 5, 5, 6, 6, 6, 5, 5, 4, 5, 5, 5, 6, 5}}, 
 {21, {3, 5, 5, 5, 4, 5, 5, 5, 6, 5, 5, 5, 6, 6, 5, 5, 5, 5}}, 
 {21, {3, 5, 5, 4, 5, 6, 6, 5, 5, 5, 5, 6, 5, 5, 5, 5, 5, 5}}, 
 {21, {3, 5, 5, 4, 5, 5, 6, 5, 5, 6, 6, 5, 5, 5, 5, 5, 5, 5}}, 
 {21, {3, 5, 5, 4, 5, 5, 6, 5, 5, 5, 6, 6, 5, 5, 5, 5, 5, 5}}, 
 {21, {3, 5, 5, 4, 5, 5, 5, 6, 6, 6, 5, 5, 4, 5, 5, 5, 6, 5}}, 
 {21, {3, 5, 5, 4, 5, 5, 5, 6, 5, 5, 5, 5, 5, 5, 5, 6, 6, 5}}, 
 {21, {3, 5, 5, 4, 5, 5, 5, 5, 5, 6, 5, 5, 5, 5, 6, 6, 5, 5}}, 
 {21, {3, 5, 5, 4, 4, 5, 5, 5, 6, 6, 6, 5, 5, 5, 5, 5, 6, 5}}, 
 {21, {3, 5, 5, 4, 4, 5, 5, 5, 5, 5, 5, 5, 5, 6, 6, 6, 6, 5}}, 
 {21, {1, 6, 6, 6, 5, 5, 4, 4, 5, 5, 4, 5, 5, 5, 6, 6, 6, 6}}, 
 {21, {1, 6, 6, 6, 1, 6, 6, 6, 5, 4, 5, 5, 5, 5, 5, 6, 6, 6}}, 
 {21, {1, 6, 6, 5, 5, 5, 5, 6, 5, 5, 5, 5, 4, 4, 5, 6, 6, 6}}, 
 {21, {1, 6, 6, 5, 5, 5, 5, 5, 5, 5, 6, 5, 3, 5, 5, 6, 6, 6}}, 
 {21, {1, 6, 6, 5, 5, 5, 5, 4, 5, 5, 5, 5, 5, 6, 5, 5, 6, 6}}, 
 {21, {1, 6, 6, 5, 5, 5, 4, 5, 5, 5, 6, 6, 6, 6, 1, 6, 6, 6}}, 
 {21, {1, 6, 6, 5, 5, 5, 4, 5, 5, 4, 5, 5, 5, 5, 6, 6, 6, 6}}, 
 {21, {1, 6, 6, 5, 5, 5, 4, 4, 5, 5, 6, 5, 5, 5, 5, 6, 6, 6}}, 
 {21, {1, 6, 6, 5, 4, 5, 5, 5, 5, 6, 5, 5, 4, 5, 5, 6, 6, 6}}, 
 {21, {1, 6, 6, 5, 4, 5, 5, 5, 5, 5, 5, 5, 4, 5, 6, 6, 6, 6}}, 
 {21, {1, 6, 6, 1, 6, 6, 6, 1, 6, 6, 6, 5, 5, 5, 6, 6, 6, 6}}, 
 {21, {1, 6, 6, 1, 6, 6, 5, 5, 5, 5, 6, 6, 5, 5, 5, 5, 6, 6}}, 
 {21, {1, 6, 6, 1, 6, 6, 5, 5, 5, 5, 4, 5, 5, 6, 6, 6, 6, 6}}, 
 {21, {1, 6, 6, 1, 6, 6, 5, 5, 4, 5, 6, 5, 5, 5, 6, 6, 6, 6}}, 
 {21, {1, 6, 6, 1, 6, 6, 5, 5, 4, 5, 5, 6, 6, 5, 5, 6, 6, 6}}, 
 {21, {4, 5, 6, 5, 4, 5, 6, 5, 5, 5, 6, 6, 6, 5, 6, 6, 5}}, 
 {21, {4, 5, 5, 6, 6, 6, 5, 6, 6, 5, 4, 5, 6, 5, 5, 6, 5}}, 
 {21, {4, 5, 5, 6, 5, 4, 5, 6, 6, 6, 5, 6, 6, 5, 5, 6, 5}}, 
 {21, {4, 5, 5, 4, 5, 5, 4, 5, 6, 6, 6, 6, 6, 6, 6, 6, 5}}, 
 {21, {4, 5, 4, 5, 5, 5, 6, 6, 6, 5, 6, 6, 5, 6, 6, 5, 5}}, 
 {21, {4, 5, 4, 5, 5, 5, 5, 6, 6, 6, 6, 6, 6, 5, 5, 6, 5}}, 
 {21, {4, 4, 5, 6, 5, 5, 6, 5, 4, 5, 6, 6, 6, 6, 6, 6, 5}}, 
 {21, {4, 4, 5, 6, 5, 5, 4, 5, 6, 6, 5, 6, 6, 6, 6, 6, 5}}, 
 {21, {4, 4, 5, 6, 5, 4, 5, 6, 6, 5, 5, 6, 6, 6, 6, 6, 5}}, 
 {21, {4, 4, 5, 5, 6, 6, 6, 6, 5, 6, 6, 6, 5, 4, 5, 6, 5}}, 
 {21, {4, 4, 5, 5, 6, 6, 5, 6, 6, 5, 4, 5, 6, 6, 6, 6, 5}}, 
 {21, {4, 4, 5, 4, 5, 6, 6, 6, 6, 6, 6, 6, 5, 5, 6, 5, 5}}, 
 {21, {3, 5, 6, 6, 5, 5, 6, 5, 5, 5, 6, 5, 5, 6, 6, 6, 5}}, 
 {21, {3, 5, 6, 6, 5, 5, 5, 6, 6, 5, 4, 5, 6, 6, 6, 6, 5}}, 
 {21, {3, 5, 6, 6, 5, 5, 4, 5, 5, 6, 6, 6, 6, 5, 6, 6, 5}}, 
 {21, {3, 5, 6, 6, 5, 4, 5, 6, 6, 6, 5, 5, 5, 6, 6, 6, 5}}, 
 {21, {3, 5, 6, 5, 4, 5, 5, 5, 6, 6, 6, 6, 5, 6, 6, 6, 5}}, 
 {21, {3, 5, 5, 6, 6, 6, 6, 5, 6, 6, 6, 5, 4, 5, 5, 6, 5}}, 
 {21, {3, 5, 5, 6, 6, 6, 5, 5, 6, 6, 6, 5, 4, 5, 6, 6, 5}}, 
 {21, {3, 5, 5, 6, 6, 6, 5, 4, 5, 6, 6, 6, 6, 5, 5, 6, 5}}, 
 {21, {3, 5, 5, 6, 6, 5, 6, 6, 6, 5, 5, 6, 5, 5, 5, 6, 5}}, 
 {21, {3, 5, 5, 6, 6, 5, 5, 4, 5, 6, 6, 6, 5, 6, 6, 6, 5}}, 
 {21, {3, 5, 5, 6, 5, 4, 5, 5, 5, 6, 6, 6, 6, 6, 6, 6, 5}}, 
 {21, {3, 5, 5, 5, 6, 6, 6, 6, 5, 5, 6, 6, 5, 5, 6, 5, 5}}, 
 {21, {3, 5, 5, 5, 5, 4, 5, 6, 6, 6, 6, 6, 6, 5, 6, 6, 5}}, 
 {21, {1, 6, 6, 6, 6, 5, 3, 5, 5, 6, 6, 5, 6, 6, 6, 6, 6}}, 
 {21, {1, 6, 6, 6, 5, 4, 4, 5, 6, 6, 6, 6, 6, 5, 6, 6, 6}}, 
 {21, {1, 6, 6, 6, 5, 4, 4, 5, 6, 6, 6, 5, 6, 6, 6, 6, 6}}, 
 {21, {1, 6, 6, 6, 5, 3, 5, 6, 6, 6, 5, 5, 6, 6, 6, 6, 6}}, 
 {21, {1, 6, 6, 5, 6, 6, 6, 6, 5, 5, 6, 5, 4, 5, 6, 6, 6}}, 
 {21, {1, 6, 6, 5, 6, 6, 6, 6, 5, 4, 5, 6, 5, 5, 6, 6, 6}}, 
 {21, {1, 6, 6, 5, 6, 6, 6, 5, 5, 6, 6, 5, 4, 5, 6, 6, 6}}, 
 {21, {1, 6, 6, 5, 6, 6, 5, 5, 4, 5, 6, 6, 6, 6, 5, 6, 6}}, 
 {21, {1, 6, 6, 5, 6, 6, 5, 4, 5, 6, 5, 5, 6, 6, 6, 6, 6}}, 
 {21, {1, 6, 6, 5, 6, 6, 1, 6, 6, 5, 6, 6, 6, 6, 6, 6, 6}}, 
 {21, {1, 6, 6, 5, 5, 6, 6, 1, 6, 6, 6, 6, 6, 6, 6, 6, 6}}, 
 {21, {1, 6, 6, 5, 4, 5, 6, 6, 5, 5, 6, 6, 6, 6, 5, 6, 6}}, 
 {21, {1, 6, 1, 6, 6, 5, 5, 6, 6, 6, 6, 6, 6, 6, 6, 6, 6}}, 
 {20, {4, 4, 5, 5, 4, 5, 5, 5, 5, 5, 4, 4, 5, 5, 5, 5, 5, 5, 5}}, 
 {20, {4, 5, 5, 5, 5, 6, 5, 4, 5, 5, 5, 6, 5, 4, 5, 5, 6, 5}}, 
 {20, {4, 5, 5, 5, 5, 5, 5, 6, 5, 4, 5, 6, 5, 5, 4, 5, 6, 5}}, 
 {20, {4, 5, 5, 5, 5, 5, 5, 4, 5, 5, 6, 5, 5, 4, 5, 6, 6, 5}}, 
 {20, {4, 5, 5, 5, 5, 4, 5, 6, 5, 5, 5, 5, 5, 4, 5, 6, 6, 5}}, 
 {20, {4, 5, 5, 5, 4, 5, 5, 5, 6, 6, 5, 4, 5, 5, 5, 5, 6, 5}}, 
 {20, {4, 5, 5, 5, 4, 5, 5, 5, 6, 5, 5, 5, 5, 4, 5, 6, 6, 5}}, 
 {20, {4, 5, 5, 5, 4, 5, 5, 5, 5, 4, 5, 6, 6, 5, 5, 6, 5, 5}}, 
 {20, {4, 5, 5, 5, 4, 5, 5, 5, 4, 5, 5, 6, 5, 5, 5, 6, 6, 5}}, 
 {20, {4, 5, 5, 4, 5, 5, 6, 5, 4, 5, 5, 5, 5, 5, 5, 6, 6, 5}}, 
 {20, {4, 5, 5, 4, 5, 5, 5, 6, 5, 4, 5, 5, 5, 5, 5, 6, 6, 5}}, 
 {20, {4, 5, 5, 4, 5, 5, 5, 5, 6, 6, 5, 4, 5, 5, 6, 5, 5, 5}}, 
 {20, {4, 5, 5, 4, 5, 5, 5, 5, 6, 6, 5, 4, 5, 5, 5, 6, 5, 5}}, 
 {20, {4, 5, 5, 4, 5, 5, 5, 5, 4, 5, 6, 5, 5, 5, 5, 6, 6, 5}}, 
 {20, {4, 5, 5, 4, 5, 5, 4, 5, 5, 6, 5, 5, 5, 5, 5, 6, 6, 5}}, 
 {20, {4, 5, 4, 5, 5, 5, 5, 5, 5, 4, 5, 5, 5, 6, 6, 6, 5, 5}}, 
 {20, {4, 4, 5, 6, 5, 5, 5, 5, 5, 4, 5, 5, 6, 5, 5, 5, 6, 5}}, 
 {20, {4, 4, 5, 6, 5, 5, 5, 5, 4, 5, 5, 5, 5, 6, 5, 5, 6, 5}}, 
 {20, {4, 4, 5, 6, 5, 5, 4, 5, 5, 5, 5, 5, 6, 5, 5, 5, 6, 5}}, 
 {20, {4, 4, 5, 5, 6, 5, 5, 4, 5, 6, 5, 5, 5, 5, 5, 5, 6, 5}}, 
 {20, {4, 4, 5, 5, 6, 5, 5, 4, 5, 5, 5, 4, 5, 6, 6, 6, 5, 5}}, 
 {20, {4, 4, 5, 5, 5, 6, 6, 6, 5, 4, 5, 5, 5, 4, 5, 5, 6, 5}}, 
 {20, {4, 4, 5, 5, 5, 6, 6, 5, 5, 4, 4, 5, 5, 6, 6, 5, 5, 5}}, 
 {20, {4, 4, 5, 5, 5, 6, 5, 5, 5, 6, 5, 4, 5, 5, 5, 5, 6, 5}}, 
 {20, {4, 4, 5, 5, 5, 6, 5, 5, 5, 4, 5, 5, 6, 5, 5, 5, 6, 5}}, 
 {20, {4, 4, 5, 5, 5, 6, 5, 5, 4, 5, 5, 5, 5, 6, 5, 5, 6, 5}}, 
 {20, {4, 4, 5, 5, 5, 6, 5, 4, 5, 6, 5, 5, 5, 5, 6, 5, 5, 5}}, 
 {20, {4, 4, 5, 5, 5, 5, 6, 6, 5, 5, 4, 4, 5, 6, 6, 5, 5, 5}}, 
 {20, {4, 4, 5, 5, 5, 5, 6, 5, 5, 6, 5, 5, 5, 4, 5, 5, 6, 5}}, 
 {20, {4, 4, 5, 5, 5, 5, 6, 5, 5, 6, 5, 5, 4, 5, 5, 5, 6, 5}}, 
 {20, {4, 4, 5, 5, 5, 5, 6, 5, 5, 5, 5, 6, 5, 4, 5, 6, 5, 5}}, 
 {20, {4, 4, 5, 5, 5, 5, 6, 5, 5, 5, 4, 5, 6, 5, 5, 5, 6, 5}}, 
 {20, {4, 4, 5, 5, 5, 5, 6, 5, 4, 5, 5, 6, 5, 5, 5, 6, 5, 5}}, 
 {20, {4, 4, 5, 5, 5, 5, 6, 5, 4, 5, 5, 5, 6, 5, 5, 6, 5, 5}}, 
 {20, {4, 4, 5, 5, 5, 5, 5, 6, 5, 5, 4, 4, 5, 6, 6, 6, 5, 5}}, 
 {20, {4, 4, 5, 5, 5, 5, 5, 5, 6, 6, 5, 5, 4, 4, 5, 6, 6, 5}}, 
 {20, {4, 4, 5, 5, 5, 5, 5, 5, 6, 5, 4, 4, 5, 6, 6, 6, 5, 5}}, 
 {20, {4, 4, 5, 5, 5, 5, 5, 5, 5, 6, 5, 4, 4, 5, 6, 6, 6, 5}}, 
 {20, {4, 4, 5, 5, 5, 5, 4, 4, 5, 5, 5, 6, 6, 5, 6, 6, 5, 5}}, 
 {20, {4, 4, 5, 5, 5, 4, 5, 6, 6, 6, 5, 5, 4, 5, 5, 5, 6, 5}}, 
 {20, {4, 4, 5, 5, 5, 4, 5, 5, 5, 6, 6, 6, 6, 5, 4, 5, 5, 5}}, 
 {20, {4, 4, 5, 5, 5, 4, 4, 5, 5, 5, 5, 6, 6, 6, 5, 5, 6, 5}}, 
 {20, {4, 4, 5, 5, 4, 5, 5, 6, 6, 5, 5, 5, 4, 5, 6, 6, 5, 5}}, 
 {20, {4, 4, 5, 5, 4, 5, 5, 6, 6, 5, 5, 5, 4, 5, 5, 6, 6, 5}}, 
 {20, {4, 4, 5, 5, 4, 5, 5, 6, 5, 4, 5, 5, 5, 6, 6, 6, 5, 5}}, 
 {20, {4, 4, 5, 5, 4, 5, 5, 5, 5, 6, 6, 6, 5, 5, 4, 5, 6, 5}}, 
 {20, {4, 4, 5, 5, 4, 4, 5, 5, 5, 6, 6, 5, 5, 6, 6, 5, 5, 5}}, 
 {20, {4, 4, 5, 5, 4, 4, 5, 5, 5, 6, 5, 5, 5, 6, 6, 6, 5, 5}}, 
 {20, {4, 4, 5, 5, 4, 4, 5, 5, 5, 5, 6, 6, 5, 6, 6, 5, 5, 5}}, 
 {20, {4, 4, 5, 4, 5, 5, 6, 6, 6, 6, 5, 5, 4, 5, 5, 5, 5, 5}}, 
 {20, {4, 4, 5, 4, 5, 5, 6, 6, 6, 5, 5, 5, 5, 4, 5, 5, 6, 5}}, 
 {20, {4, 4, 5, 4, 5, 5, 6, 6, 5, 5, 4, 5, 5, 6, 6, 5, 5, 5}}, 
 {20, {4, 4, 5, 4, 5, 5, 5, 5, 6, 6, 6, 6, 5, 5, 4, 5, 5, 5}}, 
 {20, {4, 4, 5, 4, 5, 5, 5, 5, 6, 6, 5, 5, 5, 6, 5, 5, 5, 5}}, 
 {20, {4, 4, 5, 4, 5, 5, 5, 5, 6, 5, 5, 5, 6, 6, 5, 5, 5, 5}}, 
 {20, {4, 4, 5, 4, 4, 5, 5, 6, 6, 6, 5, 5, 5, 5, 5, 5, 6, 5}}, 
 {20, {4, 4, 5, 4, 4, 5, 5, 5, 6, 5, 5, 5, 5, 5, 6, 6, 6, 5}}, 
 {20, {4, 4, 5, 4, 4, 5, 5, 5, 4, 5, 5, 6, 6, 6, 6, 6, 5, 5}}, 
 {20, {3, 5, 6, 5, 5, 5, 5, 5, 5, 5, 4, 5, 5, 6, 5, 5, 6, 5}}, 
 {20, {3, 5, 6, 5, 5, 5, 5, 5, 5, 4, 4, 5, 5, 5, 6, 6, 6, 5}}, 
 {20, {3, 5, 6, 5, 5, 5, 5, 4, 5, 5, 5, 5, 5, 6, 5, 5, 6, 5}}, 
 {20, {3, 5, 6, 5, 4, 4, 5, 5, 5, 5, 5, 5, 5, 5, 6, 6, 6, 5}}, 
 {20, {3, 5, 5, 6, 6, 6, 5, 5, 4, 5, 5, 5, 4, 5, 5, 5, 6, 5}}, 
 {20, {3, 5, 5, 6, 5, 5, 5, 5, 5, 5, 5, 4, 5, 5, 5, 6, 6, 5}}, 
 {20, {3, 5, 5, 6, 5, 5, 5, 4, 5, 5, 5, 5, 5, 5, 5, 6, 6, 5}}, 
 {20, {3, 5, 5, 5, 6, 6, 6, 5, 5, 5, 5, 5, 4, 4, 5, 5, 6, 5}}, 
 {20, {3, 5, 5, 5, 6, 6, 6, 5, 5, 5, 5, 4, 4, 5, 5, 5, 6, 5}}, 
 {20, {3, 5, 5, 5, 6, 6, 5, 4, 5, 5, 5, 5, 5, 5, 5, 5, 6, 5}}, 
 {20, {3, 5, 5, 5, 6, 5, 5, 5, 5, 5, 5, 5, 4, 5, 5, 6, 6, 5}}, 
 {20, {3, 5, 5, 5, 6, 5, 5, 5, 5, 4, 5, 5, 5, 5, 6, 6, 5, 5}}, 
 {20, {3, 5, 5, 5, 6, 5, 5, 4, 5, 5, 5, 5, 5, 5, 6, 6, 5, 5}}, 
 {20, {3, 5, 5, 5, 5, 6, 6, 6, 5, 5, 4, 4, 5, 5, 5, 5, 6, 5}}, 
 {20, {3, 5, 5, 5, 5, 6, 6, 5, 5, 5, 5, 5, 5, 5, 4, 5, 6, 5}}, 
 {20, {3, 5, 5, 5, 5, 6, 6, 5, 5, 5, 5, 4, 5, 5, 6, 5, 5, 5}}, 
 {20, {3, 5, 5, 5, 5, 5, 6, 6, 5, 5, 4, 5, 5, 6, 5, 5, 5, 5}}, 
 {20, {3, 5, 5, 5, 5, 5, 6, 6, 5, 5, 4, 5, 5, 5, 6, 5, 5, 5}}, 
 {20, {3, 5, 5, 5, 5, 5, 6, 5, 5, 5, 4, 5, 5, 5, 5, 6, 6, 5}}, 
 {20, {3, 5, 5, 5, 5, 5, 6, 5, 5, 4, 5, 5, 6, 6, 5, 5, 5, 5}}, 
 {20, {3, 5, 5, 5, 5, 5, 6, 5, 5, 4, 5, 5, 5, 5, 5, 6, 6, 5}}, 
 {20, {3, 5, 5, 5, 5, 5, 6, 5, 4, 5, 5, 5, 5, 6, 6, 5, 5, 5}}, 
 {20, {3, 5, 5, 5, 5, 5, 6, 5, 4, 5, 5, 5, 5, 5, 6, 6, 5, 5}}, 
 {20, {3, 5, 5, 5, 5, 5, 5, 6, 6, 5, 4, 5, 6, 5, 5, 5, 5, 5}}, 
 {20, {3, 5, 5, 5, 5, 5, 5, 6, 5, 5, 4, 5, 5, 5, 5, 6, 6, 5}}, 
 {20, {3, 5, 5, 5, 5, 5, 5, 5, 6, 6, 5, 4, 5, 6, 5, 5, 5, 5}}, 
 {20, {3, 5, 5, 5, 5, 5, 5, 5, 6, 5, 5, 4, 5, 5, 5, 6, 6, 5}}, 
 {20, {3, 5, 5, 5, 5, 5, 5, 5, 5, 5, 5, 6, 6, 5, 4, 5, 6, 5}}, 
 {20, {3, 5, 5, 5, 5, 5, 5, 5, 4, 4, 5, 6, 6, 6, 6, 5, 5, 5}}, 
 {20, {3, 5, 5, 5, 5, 5, 5, 4, 5, 5, 6, 5, 5, 5, 5, 6, 6, 5}}, 
 {20, {3, 5, 5, 5, 5, 5, 5, 4, 5, 5, 5, 5, 6, 6, 5, 5, 6, 5}}, 
 {20, {3, 5, 5, 5, 5, 5, 5, 4, 5, 5, 5, 5, 6, 5, 5, 6, 6, 5}}, 
 {20, {3, 5, 5, 5, 5, 5, 4, 5, 6, 6, 5, 5, 5, 5, 5, 5, 6, 5}}, 
 {20, {3, 5, 5, 5, 5, 5, 4, 5, 5, 5, 4, 5, 5, 6, 6, 6, 6, 5}}, 
 {20, {3, 5, 5, 5, 5, 5, 4, 4, 5, 5, 5, 5, 6, 6, 6, 6, 5, 5}}, 
 {20, {3, 5, 5, 5, 5, 5, 4, 4, 5, 5, 5, 5, 5, 6, 6, 6, 6, 5}}, 
 {20, {3, 5, 5, 5, 5, 4, 5, 5, 6, 5, 5, 5, 6, 6, 5, 5, 5, 5}}, 
 {20, {3, 5, 5, 5, 5, 4, 5, 5, 5, 6, 6, 5, 5, 6, 5, 5, 5, 5}}, 
 {20, {3, 5, 5, 5, 5, 4, 5, 5, 5, 5, 4, 5, 5, 6, 6, 6, 6, 5}}, 
 {20, {3, 5, 5, 5, 4, 5, 5, 6, 6, 5, 5, 6, 5, 5, 5, 5, 5, 5}}, 
 {20, {3, 5, 5, 5, 4, 5, 5, 5, 5, 6, 5, 5, 5, 5, 5, 6, 6, 5}}, 
 {20, {3, 5, 5, 5, 4, 5, 5, 5, 5, 5, 6, 6, 5, 5, 6, 5, 5, 5}}, 
 {20, {3, 5, 5, 5, 4, 5, 5, 5, 5, 5, 5, 6, 5, 5, 5, 6, 6, 5}}, 
 {20, {3, 5, 5, 5, 4, 5, 5, 5, 5, 5, 4, 5, 5, 6, 6, 6, 6, 5}}, 
 {20, {3, 5, 5, 5, 4, 5, 5, 4, 5, 5, 6, 6, 6, 6, 5, 5, 5, 5}}, 
 {20, {3, 5, 5, 5, 4, 4, 5, 5, 5, 5, 5, 6, 6, 6, 6, 5, 5, 5}}, 
 {20, {3, 5, 5, 5, 4, 4, 5, 5, 5, 5, 5, 6, 6, 6, 5, 5, 6, 5}}, 
 {20, {3, 5, 5, 4, 5, 6, 5, 5, 5, 6, 6, 5, 5, 5, 5, 5, 5, 5}}, 
 {20, {3, 5, 5, 4, 5, 5, 6, 6, 6, 5, 5, 4, 5, 5, 5, 5, 6, 5}}, 
 {20, {3, 5, 5, 4, 5, 5, 5, 6, 6, 5, 5, 5, 5, 6, 5, 5, 5, 5}}, 
 {20, {3, 5, 5, 4, 5, 5, 5, 6, 5, 5, 5, 5, 6, 5, 5, 5, 6, 5}}, 
 {20, {3, 5, 5, 4, 5, 5, 5, 5, 5, 5, 5, 6, 5, 5, 6, 6, 5, 5}}, 
 {20, {3, 5, 5, 3, 5, 6, 5, 5, 5, 5, 5, 5, 5, 5, 6, 6, 6, 5}}, 
 {20, {3, 5, 5, 3, 5, 5, 5, 5, 5, 5, 5, 5, 5, 6, 6, 6, 6, 5}}, 
 {20, {1, 6, 6, 6, 5, 5, 4, 5, 5, 5, 5, 4, 4, 5, 6, 6, 6, 6}}, 
 {20, {1, 6, 6, 5, 5, 5, 6, 5, 5, 5, 5, 4, 4, 5, 5, 6, 6, 6}}, 
 {20, {1, 6, 6, 5, 5, 5, 6, 5, 3, 5, 5, 5, 5, 5, 5, 6, 6, 6}}, 
 {20, {1, 6, 6, 5, 5, 5, 5, 6, 5, 4, 5, 5, 4, 5, 5, 6, 6, 6}}, 
 {20, {1, 6, 6, 5, 5, 5, 5, 5, 6, 5, 3, 5, 5, 5, 5, 6, 6, 6}}, 
 {20, {1, 6, 6, 5, 5, 5, 5, 5, 3, 5, 6, 5, 5, 5, 5, 6, 6, 6}}, 
 {20, {1, 6, 6, 5, 5, 5, 4, 5, 5, 5, 5, 5, 4, 5, 6, 6, 6, 6}}, 
 {20, {1, 6, 6, 5, 5, 5, 3, 5, 6, 5, 5, 5, 5, 5, 5, 6, 6, 6}}, 
 {20, {1, 6, 6, 5, 5, 4, 5, 5, 5, 6, 5, 5, 5, 4, 5, 6, 6, 6}}, 
 {20, {1, 6, 6, 5, 5, 4, 5, 5, 5, 4, 5, 6, 5, 5, 5, 6, 6, 6}}, 
 {20, {1, 6, 6, 5, 4, 4, 5, 5, 6, 5, 5, 5, 5, 5, 5, 6, 6, 6}}, 
 {20, {1, 6, 6, 5, 4, 4, 5, 5, 5, 6, 6, 5, 5, 5, 5, 5, 6, 6}}, 
 {20, {1, 6, 1, 6, 6, 5, 5, 5, 5, 5, 5, 5, 5, 6, 6, 6, 6, 6}}, 
 {20, {4, 5, 6, 5, 5, 5, 6, 6, 5, 6, 6, 5, 4, 5, 6, 6, 5}}, 
 {20, {4, 5, 6, 5, 5, 5, 6, 5, 4, 5, 6, 6, 5, 6, 6, 6, 5}}, 
 {20, {4, 5, 5, 4, 5, 6, 5, 5, 6, 6, 6, 6, 6, 5, 5, 6, 5}}, 
 {20, {4, 4, 5, 6, 5, 5, 4, 5, 6, 6, 6, 6, 5, 6, 6, 6, 5}}, 
 {20, {4, 4, 5, 5, 6, 6, 6, 6, 6, 5, 6, 6, 5, 4, 5, 6, 5}}, 
 {20, {4, 4, 5, 5, 6, 5, 4, 5, 6, 6, 6, 5, 6, 6, 6, 6, 5}}, 
 {20, {4, 4, 5, 5, 4, 5, 6, 6, 5, 6, 6, 6, 6, 5, 6, 6, 5}}, 
 {20, {4, 4, 5, 4, 5, 6, 6, 5, 5, 6, 6, 6, 6, 6, 6, 5, 5}}, 
 {20, {3, 5, 6, 6, 5, 5, 4, 5, 6, 5, 5, 6, 6, 6, 6, 6, 5}}, 
 {20, {3, 5, 6, 5, 5, 5, 4, 5, 6, 6, 6, 6, 5, 6, 6, 6, 5}}, 
 {20, {3, 5, 6, 5, 5, 5, 4, 5, 6, 6, 5, 6, 6, 6, 6, 6, 5}}, 
 {20, {3, 5, 5, 6, 6, 6, 6, 5, 6, 6, 6, 5, 5, 4, 5, 6, 5}}, 
 {20, {3, 5, 5, 6, 6, 6, 5, 5, 6, 6, 6, 6, 5, 4, 5, 6, 5}}, 
 {20, {3, 5, 5, 6, 6, 5, 6, 6, 5, 5, 4, 5, 6, 6, 6, 6, 5}}, 
 {20, {3, 5, 5, 6, 6, 5, 5, 5, 6, 5, 5, 6, 6, 5, 6, 6, 5}}, 
 {20, {3, 5, 5, 6, 6, 5, 4, 5, 5, 6, 6, 6, 5, 6, 6, 6, 5}}, 
 {20, {3, 5, 5, 6, 5, 5, 6, 6, 5, 6, 6, 6, 5, 5, 5, 6, 5}}, 
 {20, {3, 5, 5, 5, 6, 6, 6, 5, 6, 6, 6, 5, 5, 5, 6, 5, 5}}, 
 {20, {3, 5, 5, 5, 6, 5, 5, 6, 6, 6, 5, 6, 6, 5, 5, 6, 5}}, 
 {20, {3, 5, 5, 4, 5, 6, 6, 6, 5, 5, 6, 6, 6, 6, 6, 5, 5}}, 
 {20, {1, 6, 6, 6, 6, 5, 3, 5, 6, 6, 5, 5, 6, 6, 6, 6, 6}}, 
 {20, {1, 6, 6, 6, 5, 6, 6, 6, 6, 5, 4, 4, 5, 6, 6, 6, 6}}, 
 {20, {1, 6, 6, 6, 5, 6, 6, 5, 6, 6, 6, 1, 6, 6, 6, 6, 6}}, 
 {20, {1, 6, 6, 6, 5, 5, 6, 5, 4, 5, 6, 6, 6, 5, 6, 6, 6}}, 
 {20, {1, 6, 6, 6, 5, 5, 3, 5, 6, 6, 6, 6, 6, 5, 6, 6, 6}}, 
 {20, {1, 6, 6, 6, 5, 5, 3, 5, 6, 6, 6, 6, 5, 6, 6, 6, 6}}, 
 {20, {1, 6, 6, 6, 5, 5, 3, 5, 6, 6, 5, 6, 6, 6, 6, 6, 6}}, 
 {20, {1, 6, 6, 6, 1, 6, 6, 6, 6, 5, 6, 6, 5, 6, 6, 6, 6}}, 
 {20, {1, 6, 6, 6, 1, 6, 6, 6, 5, 6, 6, 5, 6, 6, 6, 6, 6}}, 
 {20, {1, 6, 6, 5, 6, 6, 6, 6, 6, 5, 5, 3, 5, 6, 6, 6, 6}}, 
 {20, {1, 6, 6, 5, 6, 6, 6, 6, 5, 4, 4, 5, 6, 6, 6, 6, 6}}, 
 {20, {1, 6, 6, 5, 6, 6, 6, 5, 6, 6, 6, 6, 1, 6, 6, 6, 6}}, 
 {20, {1, 6, 6, 5, 6, 6, 6, 5, 4, 4, 5, 6, 6, 6, 6, 6, 6}}, 
 {20, {1, 6, 6, 5, 6, 6, 6, 1, 6, 6, 5, 6, 6, 6, 6, 6, 6}}, 
 {20, {1, 6, 6, 5, 5, 6, 6, 6, 6, 6, 5, 3, 5, 6, 6, 6, 6}}, 
 {20, {1, 6, 6, 5, 5, 6, 6, 6, 5, 4, 5, 6, 6, 5, 6, 6, 6}}, 
 {20, {1, 6, 6, 5, 5, 6, 5, 3, 5, 6, 6, 6, 6, 6, 6, 6, 6}}, 
 {20, {1, 6, 6, 5, 3, 5, 5, 6, 6, 6, 6, 6, 6, 6, 5, 6, 6}}, 
 {20, {1, 6, 6, 5, 3, 5, 5, 6, 6, 6, 6, 6, 5, 6, 6, 6, 6}}, 
 {19, {4, 4, 5, 5, 5, 4, 5, 5, 5, 5, 5, 5, 5, 4, 4, 5, 5, 5, 5}}, 
 {19, {4, 4, 5, 5, 4, 5, 5, 5, 5, 4, 4, 5, 5, 5, 5, 5, 5, 5, 5}}, 
 {19, {4, 5, 5, 5, 6, 5, 4, 5, 5, 5, 6, 5, 5, 4, 5, 5, 6, 5}}, 
 {19, {4, 5, 5, 5, 5, 5, 5, 5, 4, 5, 5, 6, 6, 5, 4, 5, 6, 5}}, 
 {19, {4, 5, 5, 5, 5, 5, 4, 5, 5, 6, 5, 5, 5, 4, 5, 6, 6, 5}}, 
 {19, {4, 5, 5, 5, 5, 4, 5, 5, 6, 5, 5, 4, 5, 5, 5, 6, 6, 5}}, 
 {19, {4, 5, 5, 4, 5, 6, 5, 5, 5, 4, 5, 5, 5, 5, 5, 6, 6, 5}}, 
 {19, {4, 5, 5, 4, 5, 5, 6, 5, 4, 5, 6, 5, 5, 5, 5, 6, 5, 5}}, 
 {19, {4, 5, 5, 4, 5, 5, 5, 5, 6, 5, 5, 4, 5, 6, 6, 5, 5, 5}}, 
 {19, {4, 5, 5, 4, 5, 5, 5, 5, 6, 5, 4, 5, 6, 6, 5, 5, 5, 5}}, 
 {19, {4, 5, 5, 4, 5, 5, 5, 4, 5, 5, 5, 5, 6, 6, 5, 5, 6, 5}}, 
 {19, {4, 5, 5, 4, 5, 5, 5, 4, 5, 5, 4, 5, 5, 6, 6, 6, 6, 5}}, 
 {19, {4, 5, 5, 4, 5, 5, 4, 5, 5, 5, 6, 5, 5, 6, 6, 5, 5, 5}}, 
 {19, {4, 5, 5, 4, 5, 5, 4, 5, 5, 5, 4, 5, 5, 6, 6, 6, 6, 5}}, 
 {19, {4, 5, 4, 5, 5, 5, 6, 5, 5, 5, 5, 5, 5, 5, 5, 6, 5, 5}}, 
 {19, {4, 5, 4, 5, 5, 5, 5, 5, 4, 5, 5, 6, 6, 6, 5, 5, 5, 5}}, 
 {19, {4, 5, 4, 5, 5, 5, 5, 4, 5, 5, 6, 6, 6, 5, 5, 5, 5, 5}}, 
 {19, {4, 5, 4, 5, 5, 5, 5, 4, 5, 5, 5, 5, 6, 6, 6, 5, 5, 5}}, 
 {19, {4, 4, 5, 5, 6, 6, 6, 5, 5, 5, 4, 5, 5, 4, 5, 5, 6, 5}}, 
 {19, {4, 4, 5, 5, 6, 5, 5, 5, 5, 6, 5, 4, 5, 5, 5, 5, 6, 5}}, 
 {19, {4, 4, 5, 5, 6, 5, 5, 5, 5, 5, 6, 5, 5, 4, 5, 5, 6, 5}}, 
 {19, {4, 4, 5, 5, 6, 5, 5, 5, 5, 5, 5, 6, 5, 5, 4, 5, 6, 5}}, 
 {19, {4, 4, 5, 5, 6, 5, 5, 4, 5, 5, 4, 5, 5, 6, 6, 6, 5, 5}}, 
 {19, {4, 4, 5, 5, 6, 5, 4, 5, 5, 6, 5, 5, 5, 5, 5, 5, 6, 5}}, 
 {19, {4, 4, 5, 5, 6, 5, 4, 5, 5, 5, 6, 5, 5, 5, 5, 5, 6, 5}}, 
 {19, {4, 4, 5, 5, 5, 6, 5, 5, 5, 6, 5, 5, 4, 5, 5, 5, 6, 5}}, 
 {19, {4, 4, 5, 5, 5, 5, 6, 5, 5, 5, 5, 6, 5, 5, 4, 5, 6, 5}}, 
 {19, {4, 4, 5, 5, 5, 5, 6, 5, 5, 4, 4, 5, 5, 5, 6, 6, 6, 5}}, 
 {19, {4, 4, 5, 5, 5, 5, 5, 4, 5, 5, 5, 4, 5, 6, 6, 6, 6, 5}}, 
 {19, {4, 4, 5, 5, 5, 4, 5, 6, 5, 5, 5, 6, 5, 5, 5, 5, 6, 5}}, 
 {19, {4, 4, 5, 5, 5, 4, 5, 6, 5, 5, 5, 5, 5, 6, 5, 5, 6, 5}}, 
 {19, {4, 4, 5, 5, 5, 4, 5, 5, 6, 5, 5, 6, 5, 5, 5, 6, 5, 5}}, 
 {19, {4, 4, 5, 5, 5, 4, 5, 5, 5, 6, 6, 5, 5, 4, 5, 6, 6, 5}}, 
 {19, {4, 4, 5, 5, 4, 5, 6, 6, 5, 5, 4, 5, 6, 6, 5, 5, 5, 5}}, 
 {19, {4, 4, 5, 5, 4, 5, 6, 5, 5, 5, 4, 5, 6, 6, 6, 5, 5, 5}}, 
 {19, {4, 4, 5, 5, 4, 5, 6, 5, 5, 5, 4, 5, 5, 6, 6, 6, 5, 5}}, 
 {19, {4, 4, 5, 5, 4, 5, 6, 5, 5, 4, 5, 5, 6, 6, 6, 5, 5, 5}}, 
 {19, {4, 4, 5, 5, 4, 5, 5, 5, 4, 5, 5, 6, 6, 6, 5, 5, 6, 5}}, 
 {19, {4, 4, 5, 5, 4, 5, 5, 4, 5, 5, 6, 6, 5, 5, 5, 6, 6, 5}}, 
 {19, {4, 4, 5, 5, 4, 4, 5, 5, 5, 5, 6, 5, 5, 6, 6, 6, 5, 5}}, 
 {19, {4, 4, 5, 5, 4, 4, 5, 5, 5, 5, 5, 6, 6, 6, 5, 5, 6, 5}}, 
 {19, {4, 4, 5, 5, 4, 4, 5, 5, 5, 5, 5, 6, 6, 5, 6, 6, 5, 5}}, 
 {19, {4, 4, 5, 5, 4, 4, 5, 5, 4, 5, 5, 6, 6, 6, 6, 6, 5, 5}}, 
 {19, {4, 4, 5, 4, 5, 5, 6, 5, 5, 5, 6, 6, 5, 5, 5, 5, 5, 5}}, 
 {19, {4, 4, 5, 4, 5, 5, 5, 5, 6, 5, 5, 6, 6, 5, 5, 5, 5, 5}}, 
 {19, {4, 4, 5, 4, 5, 5, 5, 5, 6, 5, 5, 5, 5, 5, 5, 6, 6, 5}}, 
 {19, {4, 4, 5, 4, 5, 5, 5, 5, 4, 5, 6, 6, 6, 6, 5, 5, 5, 5}}, 
 {19, {4, 4, 5, 4, 5, 5, 4, 5, 5, 5, 5, 5, 6, 6, 6, 6, 5, 5}}, 
 {19, {4, 4, 5, 4, 4, 5, 5, 5, 6, 6, 6, 5, 5, 5, 5, 6, 5, 5}}, 
 {19, {4, 4, 5, 4, 4, 5, 5, 5, 6, 6, 5, 5, 5, 5, 5, 6, 6, 5}}, 
 {19, {4, 4, 5, 4, 4, 5, 5, 5, 5, 6, 6, 6, 5, 5, 5, 5, 6, 5}}, 
 {19, {4, 4, 5, 4, 4, 5, 5, 5, 5, 6, 5, 5, 5, 5, 6, 6, 6, 5}}, 
 {19, {4, 4, 5, 4, 4, 5, 5, 5, 5, 5, 6, 6, 6, 5, 5, 6, 5, 5}}, 
 {19, {3, 5, 6, 6, 5, 5, 4, 5, 5, 5, 5, 4, 5, 5, 5, 6, 6, 5}}, 
 {19, {3, 5, 6, 5, 5, 5, 5, 5, 5, 5, 4, 5, 6, 5, 5, 5, 6, 5}}, 
 {19, {3, 5, 6, 5, 5, 5, 5, 4, 5, 5, 5, 6, 5, 5, 5, 5, 6, 5}}, 
 {19, {3, 5, 5, 6, 6, 5, 5, 5, 5, 5, 5, 5, 4, 4, 5, 6, 6, 5}}, 
 {19, {3, 5, 5, 6, 6, 5, 5, 5, 4, 4, 5, 5, 5, 5, 6, 6, 5, 5}}, 
 {19, {3, 5, 5, 5, 6, 5, 5, 5, 5, 5, 6, 5, 5, 4, 5, 5, 6, 5}}, 
 {19, {3, 5, 5, 5, 5, 6, 6, 6, 5, 5, 5, 5, 4, 4, 5, 5, 6, 5}}, 
 {19, {3, 5, 5, 5, 5, 6, 6, 6, 5, 5, 5, 4, 4, 5, 5, 5, 6, 5}}, 
 {19, {3, 5, 5, 5, 5, 6, 6, 5, 5, 4, 5, 5, 5, 5, 6, 5, 5, 5}}, 
 {19, {3, 5, 5, 5, 5, 6, 6, 5, 4, 5, 5, 5, 5, 5, 5, 5, 6, 5}}, 
 {19, {3, 5, 5, 5, 5, 5, 6, 5, 4, 5, 5, 5, 6, 6, 5, 5, 5, 5}}, 
 {19, {3, 5, 5, 5, 5, 5, 5, 5, 6, 6, 5, 5, 5, 5, 4, 5, 6, 5}}, 
 {19, {3, 5, 5, 5, 5, 5, 5, 5, 5, 5, 4, 5, 6, 5, 5, 6, 6, 5}}, 
 {19, {3, 5, 5, 5, 5, 5, 5, 5, 4, 5, 5, 6, 5, 5, 5, 6, 6, 5}}, 
 {19, {3, 5, 5, 5, 5, 5, 5, 4, 5, 5, 6, 5, 5, 6, 6, 5, 5, 5}}, 
 {19, {3, 5, 5, 5, 5, 5, 5, 4, 4, 5, 5, 6, 6, 6, 6, 5, 5, 5}}, 
 {19, {3, 5, 5, 5, 5, 5, 4, 5, 5, 6, 6, 5, 5, 6, 5, 5, 5, 5}}, 
 {19, {3, 5, 5, 5, 5, 4, 5, 6, 6, 5, 5, 5, 5, 6, 5, 5, 5, 5}}, 
 {19, {3, 5, 5, 5, 5, 4, 5, 5, 6, 5, 5, 6, 6, 5, 5, 5, 5, 5}}, 
 {19, {3, 5, 5, 5, 5, 4, 5, 5, 6, 5, 5, 5, 5, 6, 6, 5, 5, 5}}, 
 {19, {3, 5, 5, 5, 5, 4, 4, 5, 6, 5, 5, 5, 5, 5, 6, 6, 6, 5}}, 
 {19, {3, 5, 5, 5, 4, 5, 5, 5, 6, 5, 5, 6, 6, 5, 5, 5, 5, 5}}, 
 {19, {3, 5, 5, 5, 4, 4, 5, 6, 6, 6, 5, 5, 5, 5, 5, 5, 6, 5}}, 
 {19, {3, 5, 5, 5, 4, 4, 5, 5, 5, 5, 6, 6, 6, 5, 5, 5, 6, 5}}, 
 {19, {3, 5, 5, 4, 5, 6, 5, 5, 5, 5, 6, 6, 5, 5, 5, 5, 5, 5}}, 
 {19, {3, 5, 5, 4, 5, 5, 5, 6, 5, 5, 6, 6, 5, 5, 5, 5, 5, 5}}, 
 {19, {3, 5, 5, 4, 5, 5, 5, 5, 6, 6, 5, 5, 5, 5, 6, 5, 5, 5}}, 
 {19, {3, 5, 5, 4, 5, 5, 5, 5, 5, 6, 6, 5, 5, 6, 5, 5, 5, 5}}, 
 {19, {3, 5, 5, 4, 4, 5, 5, 5, 5, 5, 6, 6, 6, 5, 5, 5, 6, 5}}, 
 {19, {1, 6, 6, 6, 5, 5, 5, 4, 5, 5, 4, 4, 5, 5, 6, 6, 6, 6}}, 
 {19, {1, 6, 6, 5, 5, 5, 6, 5, 5, 4, 5, 5, 4, 5, 5, 6, 6, 6}}, 
 {19, {1, 6, 6, 5, 5, 5, 6, 5, 4, 4, 5, 5, 5, 5, 5, 6, 6, 6}}, 
 {19, {1, 6, 6, 5, 5, 5, 5, 5, 3, 5, 5, 5, 5, 5, 6, 6, 6, 6}}, 
 {19, {1, 6, 6, 5, 5, 5, 5, 4, 5, 6, 5, 5, 4, 5, 5, 6, 6, 6}}, 
 {19, {1, 6, 6, 5, 5, 5, 5, 4, 4, 5, 6, 5, 5, 5, 5, 6, 6, 6}}, 
 {19, {1, 6, 6, 5, 5, 5, 5, 4, 4, 5, 5, 5, 6, 6, 5, 5, 6, 6}}, 
 {19, {1, 6, 6, 5, 5, 5, 5, 4, 4, 5, 5, 5, 6, 5, 5, 6, 6, 6}}, 
 {19, {1, 6, 6, 5, 5, 5, 4, 5, 6, 5, 5, 5, 4, 5, 5, 6, 6, 6}}, 
 {19, {1, 6, 6, 5, 5, 5, 4, 5, 5, 5, 6, 6, 6, 1, 6, 6, 6, 6}}, 
 {19, {1, 6, 6, 5, 5, 4, 4, 5, 6, 6, 5, 5, 5, 5, 5, 5, 6, 6}}, 
 {19, {1, 6, 6, 5, 5, 4, 4, 5, 5, 5, 5, 6, 5, 5, 5, 6, 6, 6}}, 
 {19, {1, 6, 6, 5, 5, 4, 4, 5, 5, 5, 4, 5, 5, 6, 6, 6, 6, 6}}, 
 {19, {1, 6, 6, 5, 4, 5, 6, 5, 5, 4, 5, 5, 5, 5, 5, 6, 6, 6}}, 
 {19, {1, 6, 6, 5, 4, 5, 5, 6, 6, 5, 5, 5, 4, 5, 5, 5, 6, 6}}, 
 {19, {1, 6, 6, 5, 4, 5, 5, 6, 5, 5, 5, 4, 5, 5, 5, 6, 6, 6}}, 
 {19, {1, 6, 6, 5, 4, 5, 5, 5, 6, 5, 4, 5, 5, 5, 5, 6, 6, 6}}, 
 {19, {1, 6, 6, 5, 4, 5, 5, 5, 5, 5, 6, 6, 5, 5, 4, 5, 6, 6}}, 
 {19, {1, 6, 6, 5, 3, 5, 5, 5, 5, 5, 6, 5, 5, 5, 5, 6, 6, 6}}, 
 {19, {4, 5, 6, 5, 5, 6, 6, 5, 4, 5, 6, 5, 5, 6, 6, 6, 5}}, 
 {19, {4, 5, 5, 6, 5, 5, 6, 6, 6, 5, 6, 6, 5, 4, 5, 6, 5}}, 
 {19, {4, 5, 5, 6, 5, 5, 6, 6, 5, 6, 6, 5, 4, 5, 6, 6, 5}}, 
 {19, {4, 5, 5, 4, 5, 6, 6, 5, 5, 6, 6, 6, 5, 5, 6, 6, 5}}, 
 {19, {4, 5, 4, 5, 5, 6, 6, 6, 6, 5, 5, 6, 6, 5, 5, 6, 5}}, 
 {19, {4, 4, 5, 6, 6, 5, 5, 6, 6, 5, 4, 5, 6, 6, 6, 6, 5}}, 
 {19, {4, 4, 5, 5, 6, 6, 6, 5, 6, 6, 5, 4, 5, 6, 6, 6, 5}}, 
 {19, {4, 4, 5, 5, 4, 4, 5, 6, 6, 6, 6, 6, 6, 6, 6, 6, 5}}, 
 {19, {3, 5, 6, 6, 6, 6, 5, 5, 4, 4, 5, 6, 6, 6, 6, 6, 5}}, 
 {19, {3, 5, 6, 6, 6, 5, 4, 4, 5, 5, 6, 6, 6, 6, 6, 6, 5}}, 
 {19, {3, 5, 6, 5, 4, 5, 6, 6, 6, 6, 6, 6, 5, 5, 5, 6, 5}}, 
 {19, {3, 5, 6, 5, 4, 5, 6, 6, 5, 5, 5, 6, 6, 6, 6, 6, 5}}, 
 {19, {3, 5, 6, 5, 4, 5, 5, 5, 6, 6, 6, 5, 6, 6, 6, 6, 5}}, 
 {19, {3, 5, 5, 6, 6, 6, 6, 6, 5, 6, 6, 5, 4, 5, 5, 6, 5}}, 
 {19, {3, 5, 5, 6, 6, 5, 5, 4, 5, 6, 6, 5, 6, 6, 6, 6, 5}}, 
 {19, {3, 5, 5, 6, 5, 5, 6, 6, 5, 5, 6, 6, 5, 6, 6, 5, 5}}, 
 {19, {3, 5, 5, 5, 5, 6, 6, 6, 6, 5, 6, 6, 5, 5, 6, 5, 5}}, 
 {19, {3, 5, 5, 5, 5, 6, 5, 4, 5, 6, 6, 6, 6, 6, 6, 6, 5}}, 
 {19, {3, 5, 5, 5, 5, 4, 5, 6, 6, 5, 6, 6, 6, 6, 6, 6, 5}}, 
 {19, {3, 5, 5, 4, 5, 6, 6, 6, 5, 6, 6, 6, 6, 6, 5, 5, 5}}, 
 {19, {3, 5, 5, 4, 5, 5, 6, 6, 6, 6, 5, 6, 6, 5, 6, 6, 5}}, 
 {19, {3, 5, 5, 4, 5, 5, 6, 6, 5, 6, 6, 6, 6, 5, 6, 6, 5}}, 
 {19, {1, 6, 6, 6, 6, 6, 5, 3, 5, 6, 5, 5, 6, 6, 6, 6, 6}}, 
 {19, {1, 6, 6, 6, 6, 5, 5, 5, 3, 5, 6, 6, 6, 6, 6, 6, 6}}, 
 {19, {1, 6, 6, 6, 6, 5, 3, 5, 5, 6, 6, 6, 5, 6, 6, 6, 6}}, 
 {19, {1, 6, 6, 6, 5, 6, 6, 6, 1, 6, 6, 6, 5, 6, 6, 6, 6}}, 
 {19, {1, 6, 6, 6, 5, 3, 5, 6, 6, 6, 6, 5, 5, 6, 6, 6, 6}}, 
 {19, {1, 6, 6, 5, 6, 6, 5, 5, 3, 5, 6, 6, 6, 6, 6, 6, 6}}, 
 {19, {1, 6, 6, 5, 5, 4, 5, 6, 6, 5, 6, 6, 6, 5, 6, 6, 6}}, 
 {19, {1, 6, 6, 5, 4, 5, 6, 6, 5, 6, 6, 5, 5, 6, 6, 6, 6}}, 
 {19, {1, 6, 6, 5, 4, 4, 5, 6, 6, 6, 5, 6, 6, 6, 6, 6, 6}}, 
 {19, {1, 6, 6, 1, 6, 6, 6, 6, 5, 6, 6, 6, 5, 6, 6, 6, 6}}, 
 {18, {4, 4, 5, 5, 4, 4, 5, 5, 5, 4, 5, 5, 5, 5, 5, 5, 5, 5, 5}}, 
 {18, {4, 5, 5, 5, 6, 5, 4, 5, 6, 5, 5, 4, 5, 5, 6, 5, 5, 5}}, 
 {18, {4, 5, 5, 5, 6, 5, 4, 5, 5, 5, 6, 5, 5, 4, 5, 6, 5, 5}}, 
 {18, {4, 5, 5, 5, 5, 5, 6, 5, 4, 5, 5, 6, 5, 4, 5, 5, 6, 5}}, 
 {18, {4, 5, 5, 5, 5, 5, 4, 5, 5, 6, 5, 4, 5, 5, 5, 6, 6, 5}}, 
 {18, {4, 5, 5, 5, 5, 5, 4, 5, 5, 5, 6, 6, 5, 5, 4, 5, 6, 5}}, 
 {18, {4, 5, 5, 5, 5, 4, 5, 6, 5, 5, 4, 5, 6, 5, 5, 5, 6, 5}}, 
 {18, {4, 5, 5, 5, 4, 5, 5, 6, 5, 4, 5, 5, 6, 5, 5, 5, 6, 5}}, 
 {18, {4, 5, 5, 5, 4, 5, 5, 5, 5, 5, 5, 6, 6, 5, 4, 5, 6, 5}}, 
 {18, {4, 5, 5, 4, 5, 5, 5, 6, 5, 5, 5, 4, 5, 6, 6, 5, 5, 5}}, 
 {18, {4, 5, 5, 4, 5, 5, 5, 5, 5, 6, 5, 4, 5, 5, 5, 6, 6, 5}}, 
 {18, {4, 5, 5, 4, 5, 5, 5, 5, 5, 5, 6, 5, 5, 4, 5, 6, 6, 5}}, 
 {18, {4, 5, 5, 4, 5, 5, 4, 5, 5, 5, 6, 6, 5, 5, 5, 5, 6, 5}}, 
 {18, {4, 5, 4, 5, 5, 5, 5, 5, 4, 5, 5, 5, 5, 6, 6, 6, 5, 5}}, 
 {18, {4, 5, 4, 5, 5, 5, 5, 4, 5, 5, 5, 6, 6, 6, 5, 5, 5, 5}}, 
 {18, {4, 5, 4, 5, 5, 5, 4, 5, 5, 5, 5, 5, 5, 6, 6, 6, 5, 5}}, 
 {18, {4, 4, 5, 6, 5, 5, 5, 5, 5, 5, 4, 5, 5, 6, 5, 5, 6, 5}}, 
 {18, {4, 4, 5, 5, 6, 6, 6, 5, 5, 4, 5, 5, 5, 5, 4, 5, 6, 5}}, 
 {18, {4, 4, 5, 5, 6, 6, 6, 5, 4, 5, 5, 5, 4, 5, 5, 5, 6, 5}}, 
 {18, {4, 4, 5, 5, 6, 6, 5, 5, 4, 5, 5, 5, 4, 5, 5, 6, 6, 5}}, 
 {18, {4, 4, 5, 5, 6, 5, 5, 5, 5, 5, 5, 5, 6, 5, 4, 5, 6, 5}}, 
 {18, {4, 4, 5, 5, 6, 5, 5, 5, 5, 4, 5, 5, 5, 6, 5, 5, 6, 5}}, 
 {18, {4, 4, 5, 5, 5, 6, 5, 5, 5, 6, 5, 4, 5, 5, 5, 6, 5, 5}}, 
 {18, {4, 4, 5, 5, 5, 6, 5, 5, 5, 5, 5, 4, 5, 6, 5, 5, 6, 5}}, 
 {18, {4, 4, 5, 5, 5, 6, 5, 5, 4, 5, 5, 6, 5, 5, 5, 5, 6, 5}}, 
 {18, {4, 4, 5, 5, 5, 6, 5, 4, 5, 5, 4, 5, 5, 6, 6, 6, 5, 5}}, 
 {18, {4, 4, 5, 5, 5, 5, 6, 5, 5, 5, 6, 5, 4, 5, 5, 5, 6, 5}}, 
 {18, {4, 4, 5, 5, 5, 5, 5, 6, 6, 5, 4, 4, 5, 5, 6, 6, 5, 5}}, 
 {18, {4, 4, 5, 5, 5, 5, 5, 6, 5, 5, 6, 5, 4, 5, 5, 5, 6, 5}}, 
 {18, {4, 4, 5, 5, 5, 5, 5, 6, 5, 5, 5, 6, 5, 5, 4, 5, 6, 5}}, 
 {18, {4, 4, 5, 5, 5, 5, 5, 6, 5, 4, 5, 5, 5, 6, 5, 5, 6, 5}}, 
 {18, {4, 4, 5, 5, 5, 5, 5, 6, 5, 4, 4, 5, 5, 5, 6, 6, 6, 5}}, 
 {18, {4, 4, 5, 5, 5, 5, 5, 4, 5, 6, 6, 5, 5, 4, 5, 6, 6, 5}}, 
 {18, {4, 4, 5, 5, 5, 5, 5, 4, 4, 5, 5, 6, 6, 6, 5, 5, 6, 5}}, 
 {18, {4, 4, 5, 5, 5, 5, 4, 5, 5, 6, 6, 6, 5, 4, 5, 5, 6, 5}}, 
 {18, {4, 4, 5, 5, 5, 5, 4, 5, 5, 6, 6, 5, 4, 5, 6, 6, 5, 5}}, 
 {18, {4, 4, 5, 5, 5, 5, 4, 5, 5, 6, 6, 5, 4, 5, 5, 6, 6, 5}}, 
 {18, {4, 4, 5, 5, 5, 5, 4, 5, 5, 4, 5, 5, 5, 6, 6, 6, 6, 5}}, 
 {18, {4, 4, 5, 5, 5, 4, 5, 5, 6, 5, 5, 5, 6, 5, 5, 6, 5, 5}}, 
 {18, {4, 4, 5, 5, 5, 4, 4, 5, 5, 5, 5, 5, 6, 6, 5, 6, 6, 5}}, 
 {18, {4, 4, 5, 5, 4, 5, 6, 5, 5, 6, 6, 6, 5, 5, 4, 5, 5, 5}}, 
 {18, {4, 4, 5, 5, 4, 5, 6, 5, 5, 5, 5, 5, 6, 5, 5, 6, 5, 5}}, 
 {18, {4, 4, 5, 5, 4, 5, 6, 5, 5, 5, 5, 5, 6, 5, 5, 5, 6, 5}}, 
 {18, {4, 4, 5, 5, 4, 5, 6, 5, 5, 5, 5, 5, 5, 6, 5, 5, 6, 5}}, 
 {18, {4, 4, 5, 5, 4, 5, 5, 6, 6, 6, 5, 5, 5, 4, 5, 6, 5, 5}}, 
 {18, {4, 4, 5, 5, 4, 5, 5, 6, 6, 6, 5, 4, 5, 6, 5, 5, 5, 5}}, 
 {18, {4, 4, 5, 5, 4, 5, 5, 5, 6, 6, 6, 6, 5, 5, 5, 4, 5, 5}}, 
 {18, {4, 4, 5, 5, 4, 5, 5, 5, 5, 6, 6, 5, 5, 4, 5, 6, 6, 5}}, 
 {18, {4, 4, 5, 5, 4, 5, 5, 5, 5, 6, 5, 5, 6, 5, 5, 5, 6, 5}}, 
 {18, {4, 4, 5, 5, 4, 5, 5, 5, 5, 6, 5, 5, 5, 6, 5, 5, 6, 5}}, 
 {18, {4, 4, 5, 5, 4, 5, 5, 4, 5, 5, 6, 6, 6, 5, 5, 6, 5, 5}}, 
 {18, {4, 4, 5, 5, 4, 4, 5, 5, 5, 6, 5, 5, 6, 6, 6, 5, 5, 5}}, 
 {18, {4, 4, 5, 5, 4, 4, 5, 5, 5, 6, 5, 5, 5, 5, 6, 6, 6, 5}}, 
 {18, {4, 4, 5, 4, 5, 6, 6, 5, 5, 5, 5, 5, 5, 6, 5, 5, 5, 5}}, 
 {18, {4, 4, 5, 4, 5, 6, 6, 5, 5, 5, 5, 5, 5, 5, 6, 5, 5, 5}}, 
 {18, {4, 4, 5, 4, 5, 5, 6, 6, 6, 5, 5, 5, 4, 5, 5, 5, 6, 5}}, 
 {18, {4, 4, 5, 4, 5, 5, 6, 6, 6, 5, 5, 4, 5, 6, 5, 5, 5, 5}}, 
 {18, {4, 4, 5, 4, 5, 5, 6, 6, 5, 5, 4, 5, 5, 5, 6, 6, 5, 5}}, 
 {18, {4, 4, 5, 4, 5, 5, 5, 5, 5, 6, 6, 5, 5, 5, 6, 5, 5, 5}}, 
 {18, {4, 4, 5, 4, 5, 5, 5, 5, 5, 6, 5, 4, 5, 5, 6, 6, 6, 5}}, 
 {18, {4, 4, 5, 4, 4, 5, 5, 6, 5, 5, 5, 5, 5, 6, 6, 6, 5, 5}}, 
 {18, {3, 5, 6, 5, 5, 5, 5, 4, 4, 5, 5, 6, 6, 5, 5, 5, 6, 5}}, 
 {18, {3, 5, 6, 5, 5, 4, 5, 5, 5, 5, 4, 5, 5, 5, 6, 6, 6, 5}}, 
 {18, {3, 5, 6, 5, 4, 5, 5, 5, 6, 5, 5, 5, 5, 5, 5, 5, 6, 5}}, 
 {18, {3, 5, 5, 6, 6, 6, 5, 5, 5, 5, 5, 5, 5, 4, 4, 5, 6, 5}}, 
 {18, {3, 5, 5, 6, 6, 6, 5, 5, 4, 5, 5, 5, 5, 4, 5, 5, 6, 5}}, 
 {18, {3, 5, 5, 5, 6, 5, 5, 5, 5, 5, 4, 5, 5, 6, 6, 5, 5, 5}}, 
 {18, {3, 5, 5, 5, 6, 5, 5, 5, 4, 5, 5, 5, 5, 5, 5, 6, 6, 5}}, 
 {18, {3, 5, 5, 5, 5, 6, 6, 5, 5, 5, 4, 4, 5, 5, 5, 6, 6, 5}}, 
 {18, {3, 5, 5, 5, 5, 6, 6, 5, 5, 4, 4, 5, 5, 6, 6, 5, 5, 5}}, 
 {18, {3, 5, 5, 5, 5, 6, 5, 5, 5, 4, 5, 5, 6, 6, 5, 5, 5, 5}}, 
 {18, {3, 5, 5, 5, 5, 5, 6, 6, 6, 5, 5, 5, 5, 4, 4, 5, 6, 5}}, 
 {18, {3, 5, 5, 5, 5, 5, 6, 6, 5, 5, 5, 5, 5, 5, 4, 5, 6, 5}}, 
 {18, {3, 5, 5, 5, 5, 5, 6, 6, 5, 5, 5, 4, 5, 6, 5, 5, 5, 5}}, 
 {18, {3, 5, 5, 5, 5, 5, 5, 6, 6, 5, 5, 4, 5, 5, 5, 6, 5, 5}}, 
 {18, {3, 5, 5, 5, 5, 5, 5, 5, 6, 5, 4, 5, 5, 5, 5, 6, 6, 5}}, 
 {18, {3, 5, 5, 5, 5, 5, 5, 4, 5, 5, 4, 5, 5, 6, 6, 6, 6, 5}}, 
 {18, {3, 5, 5, 5, 5, 5, 5, 4, 4, 5, 6, 5, 5, 5, 6, 6, 6, 5}}, 
 {18, {3, 5, 5, 5, 5, 5, 4, 5, 5, 5, 6, 6, 5, 5, 6, 5, 5, 5}}, 
 {18, {3, 5, 5, 5, 5, 4, 5, 6, 5, 5, 5, 6, 6, 5, 5, 5, 5, 5}}, 
 {18, {3, 5, 5, 5, 4, 5, 6, 6, 6, 6, 5, 5, 4, 5, 5, 5, 5, 5}}, 
 {18, {3, 5, 5, 5, 4, 5, 6, 6, 5, 5, 5, 5, 5, 6, 5, 5, 5, 5}}, 
 {18, {3, 5, 5, 5, 4, 4, 5, 6, 5, 5, 5, 5, 5, 5, 6, 6, 6, 5}}, 
 {18, {3, 5, 5, 4, 5, 5, 6, 6, 6, 6, 5, 5, 4, 5, 5, 5, 5, 5}}, 
 {18, {3, 5, 5, 4, 5, 5, 6, 6, 5, 5, 5, 6, 5, 5, 5, 5, 5, 5}}, 
 {18, {3, 5, 5, 4, 5, 5, 5, 6, 6, 6, 6, 5, 5, 4, 5, 5, 5, 5}}, 
 {18, {3, 5, 5, 4, 5, 5, 5, 6, 5, 5, 5, 6, 6, 5, 5, 5, 5, 5}}, 
 {18, {3, 5, 5, 4, 5, 5, 5, 6, 5, 5, 5, 5, 6, 6, 5, 5, 5, 5}}, 
 {18, {3, 5, 5, 4, 5, 5, 5, 5, 6, 5, 5, 6, 6, 5, 5, 5, 5, 5}}, 
 {18, {3, 5, 5, 4, 5, 5, 5, 5, 5, 5, 5, 5, 6, 5, 5, 6, 6, 5}}, 
 {18, {3, 5, 5, 4, 5, 5, 5, 5, 5, 4, 5, 5, 5, 6, 6, 6, 6, 5}}, 
 {18, {3, 5, 5, 4, 5, 5, 5, 4, 5, 5, 6, 6, 6, 6, 5, 5, 5, 5}}, 
 {18, {3, 5, 5, 4, 4, 5, 6, 5, 5, 5, 5, 5, 5, 5, 6, 6, 6, 5}}, 
 {18, {3, 5, 5, 4, 4, 5, 5, 6, 6, 6, 5, 5, 5, 5, 5, 5, 6, 5}}, 
 {18, {1, 6, 6, 6, 5, 5, 4, 5, 4, 4, 5, 5, 5, 5, 6, 6, 6, 6}}, 
 {18, {1, 6, 6, 6, 5, 5, 4, 4, 5, 5, 5, 5, 4, 5, 6, 6, 6, 6}}, 
 {18, {1, 6, 6, 6, 5, 4, 4, 5, 5, 5, 5, 4, 5, 5, 6, 6, 6, 6}}, 
 {18, {1, 6, 6, 5, 5, 5, 5, 5, 6, 5, 5, 5, 6, 6, 1, 6, 6, 6}}, 
 {18, {1, 6, 6, 5, 5, 5, 5, 5, 5, 5, 4, 4, 5, 6, 6, 5, 6, 6}}, 
 {18, {1, 6, 6, 5, 5, 5, 5, 4, 5, 5, 6, 5, 4, 5, 5, 6, 6, 6}}, 
 {18, {1, 6, 6, 5, 5, 5, 4, 5, 6, 5, 5, 4, 5, 5, 5, 6, 6, 6}}, 
 {18, {1, 6, 6, 5, 5, 5, 4, 4, 5, 5, 5, 5, 6, 6, 5, 5, 6, 6}}, 
 {18, {1, 6, 6, 5, 5, 4, 5, 5, 5, 6, 5, 5, 4, 5, 5, 6, 6, 6}}, 
 {18, {1, 6, 6, 5, 5, 4, 5, 5, 5, 5, 6, 6, 5, 4, 5, 5, 6, 6}}, 
 {18, {1, 6, 6, 5, 5, 4, 4, 5, 5, 5, 6, 5, 5, 5, 5, 6, 6, 6}}, 
 {18, {1, 6, 6, 5, 4, 5, 6, 6, 5, 5, 5, 5, 4, 5, 5, 5, 6, 6}}, 
 {18, {1, 6, 6, 5, 4, 5, 5, 6, 5, 5, 5, 5, 4, 5, 5, 6, 6, 6}}, 
 {18, {1, 6, 6, 5, 4, 5, 5, 5, 5, 5, 5, 5, 5, 6, 5, 5, 6, 6}}, 
 {18, {1, 6, 6, 5, 4, 5, 5, 5, 4, 5, 5, 6, 5, 5, 5, 6, 6, 6}}, 
 {18, {1, 6, 6, 5, 4, 4, 5, 5, 5, 5, 5, 5, 6, 6, 5, 5, 6, 6}}, 
 {18, {1, 6, 6, 5, 3, 5, 5, 5, 5, 5, 5, 6, 6, 5, 5, 5, 6, 6}}, 
 {18, {1, 6, 6, 1, 6, 6, 6, 5, 4, 5, 5, 5, 5, 5, 6, 6, 6, 6}}, 
 {18, {1, 6, 6, 1, 6, 6, 5, 5, 5, 5, 5, 4, 5, 6, 6, 6, 6, 6}}, 
 {18, {1, 6, 6, 1, 6, 6, 1, 6, 6, 6, 5, 5, 5, 6, 6, 6, 6, 6}}, 
 {18, {4, 5, 6, 5, 5, 6, 6, 5, 5, 6, 6, 5, 4, 5, 6, 6, 5}}, 
 {18, {4, 5, 6, 5, 5, 6, 5, 4, 5, 6, 5, 5, 6, 6, 6, 6, 5}}, 
 {18, {4, 5, 6, 5, 5, 5, 6, 6, 5, 4, 5, 6, 6, 5, 6, 6, 5}}, 
 {18, {4, 5, 5, 6, 5, 5, 6, 5, 5, 4, 5, 6, 6, 6, 6, 6, 5}}, 
 {18, {4, 5, 5, 6, 5, 4, 5, 6, 6, 5, 6, 6, 5, 5, 6, 6, 5}}, 
 {18, {4, 5, 4, 5, 5, 5, 6, 5, 5, 5, 6, 6, 6, 6, 6, 6, 5}}, 
 {18, {4, 4, 5, 6, 6, 6, 5, 4, 5, 6, 6, 5, 5, 6, 6, 6, 5}}, 
 {18, {4, 4, 5, 6, 6, 5, 4, 5, 6, 6, 6, 5, 5, 6, 6, 6, 5}}, 
 {18, {4, 4, 5, 5, 6, 6, 5, 6, 6, 6, 6, 5, 4, 5, 6, 6, 5}}, 
 {18, {4, 4, 5, 5, 6, 6, 5, 6, 6, 6, 5, 4, 5, 6, 6, 6, 5}}, 
 {18, {4, 4, 5, 5, 6, 5, 4, 5, 6, 6, 6, 6, 6, 5, 6, 6, 5}}, 
 {18, {4, 4, 5, 5, 4, 5, 6, 6, 5, 6, 6, 6, 5, 6, 6, 6, 5}}, 
 {18, {4, 4, 5, 4, 5, 6, 6, 6, 6, 5, 6, 6, 5, 5, 6, 6, 5}}, 
 {18, {3, 5, 6, 6, 6, 5, 5, 4, 5, 6, 6, 5, 5, 6, 6, 6, 5}}, 
 {18, {3, 5, 6, 6, 6, 5, 4, 5, 6, 6, 5, 5, 5, 6, 6, 6, 5}}, 
 {18, {3, 5, 6, 6, 6, 5, 4, 5, 5, 5, 6, 6, 5, 6, 6, 6, 5}}, 
 {18, {3, 5, 6, 5, 4, 5, 5, 5, 6, 6, 5, 6, 6, 6, 6, 6, 5}}, 
 {18, {3, 5, 5, 6, 6, 6, 5, 6, 6, 6, 5, 5, 4, 5, 6, 6, 5}}, 
 {18, {3, 5, 5, 6, 6, 6, 5, 4, 5, 5, 6, 6, 5, 6, 6, 6, 5}}, 
 {18, {3, 5, 5, 6, 6, 5, 6, 6, 6, 6, 6, 5, 5, 4, 5, 6, 5}}, 
 {18, {3, 5, 5, 6, 6, 5, 6, 6, 6, 6, 5, 5, 4, 5, 6, 6, 5}}, 
 {18, {3, 5, 5, 6, 6, 5, 5, 6, 5, 4, 5, 6, 6, 6, 6, 6, 5}}, 
 {18, {3, 5, 5, 6, 5, 5, 6, 6, 6, 5, 5, 6, 6, 5, 5, 6, 5}}, 
 {18, {3, 5, 5, 5, 6, 6, 6, 6, 5, 6, 6, 5, 5, 5, 6, 5, 5}}, 
 {18, {3, 5, 5, 4, 5, 6, 6, 6, 6, 6, 6, 5, 6, 6, 5, 5, 5}}, 
 {18, {3, 5, 5, 4, 5, 6, 6, 6, 6, 5, 6, 6, 6, 6, 5, 5, 5}}, 
 {18, {3, 5, 5, 4, 5, 5, 6, 6, 5, 6, 6, 6, 5, 6, 6, 6, 5}}, 
 {18, {1, 6, 6, 6, 5, 3, 5, 5, 6, 6, 6, 5, 6, 6, 6, 6, 6}}, 
 {18, {1, 6, 6, 5, 6, 6, 6, 6, 6, 6, 5, 4, 4, 5, 6, 6, 6}}, 
 {18, {1, 6, 6, 5, 5, 6, 6, 6, 5, 6, 6, 5, 4, 5, 6, 6, 6}}, 
 {18, {1, 6, 6, 5, 5, 6, 6, 6, 5, 4, 5, 6, 6, 6, 5, 6, 6}}, 
 {18, {1, 6, 6, 5, 5, 6, 6, 6, 5, 3, 5, 6, 6, 6, 6, 6, 6}}, 
 {18, {1, 6, 6, 5, 5, 4, 5, 6, 6, 6, 6, 5, 6, 6, 5, 6, 6}}, 
 {18, {1, 6, 6, 5, 5, 4, 5, 6, 6, 5, 6, 6, 6, 6, 5, 6, 6}}, 
 {18, {1, 6, 6, 5, 4, 5, 5, 6, 6, 5, 6, 6, 6, 6, 5, 6, 6}}, 
 {18, {1, 6, 6, 5, 3, 5, 6, 6, 6, 6, 6, 5, 5, 6, 6, 6, 6}}, 
 {18, {1, 6, 6, 5, 3, 5, 5, 6, 6, 5, 6, 6, 6, 6, 6, 6, 6}}, 
 {17, {4, 4, 5, 5, 5, 5, 4, 5, 5, 5, 5, 5, 4, 4, 5, 5, 5, 5, 5}}, 
 {17, {4, 4, 5, 5, 4, 5, 5, 5, 5, 4, 5, 5, 5, 5, 5, 5, 4, 5, 5}}, 
 {17, {4, 4, 5, 5, 4, 4, 5, 5, 5, 5, 4, 5, 5, 5, 5, 5, 5, 5, 5}}, 
 {17, {4, 4, 5, 5, 4, 4, 5, 5, 4, 5, 5, 5, 5, 5, 5, 5, 5, 5, 5}}, 
 {17, {4, 5, 5, 5, 6, 5, 4, 5, 5, 5, 6, 5, 4, 5, 6, 5, 5, 5}}, 
 {17, {4, 5, 5, 5, 5, 5, 6, 5, 4, 5, 5, 6, 5, 4, 5, 6, 5, 5}}, 
 {17, {4, 5, 5, 5, 5, 4, 5, 5, 5, 5, 5, 6, 5, 4, 5, 6, 6, 5}}, 
 {17, {4, 5, 5, 5, 4, 5, 6, 5, 5, 5, 4, 5, 6, 5, 5, 5, 6, 5}}, 
 {17, {4, 5, 5, 5, 4, 5, 6, 5, 5, 5, 4, 5, 5, 6, 5, 5, 6, 5}}, 
 {17, {4, 5, 5, 5, 4, 5, 5, 6, 5, 5, 5, 6, 5, 4, 5, 5, 6, 5}}, 
 {17, {4, 5, 5, 5, 4, 5, 5, 5, 5, 5, 6, 5, 5, 4, 5, 6, 6, 5}}, 
 {17, {4, 5, 5, 5, 4, 5, 5, 5, 5, 5, 5, 6, 5, 4, 5, 6, 6, 5}}, 
 {17, {4, 5, 5, 4, 5, 5, 6, 5, 5, 4, 5, 5, 5, 5, 5, 6, 6, 5}}, 
 {17, {4, 5, 5, 4, 5, 5, 5, 6, 6, 5, 4, 5, 5, 5, 5, 6, 5, 5}}, 
 {17, {4, 5, 5, 4, 5, 5, 5, 6, 5, 5, 4, 5, 5, 5, 5, 6, 6, 5}}, 
 {17, {4, 5, 5, 4, 5, 5, 5, 5, 5, 5, 4, 5, 6, 5, 5, 6, 6, 5}}, 
 {17, {4, 5, 5, 4, 5, 5, 5, 5, 4, 5, 6, 6, 5, 5, 6, 5, 5, 5}}, 
 {17, {4, 5, 5, 4, 5, 5, 5, 4, 5, 5, 5, 5, 6, 5, 5, 6, 6, 5}}, 
 {17, {4, 5, 5, 4, 5, 5, 4, 5, 5, 5, 5, 6, 6, 5, 5, 5, 6, 5}}, 
 {17, {4, 5, 5, 4, 5, 5, 4, 5, 5, 5, 5, 6, 5, 5, 6, 6, 5, 5}}, 
 {17, {4, 5, 4, 5, 5, 5, 5, 6, 5, 5, 5, 5, 5, 5, 6, 5, 5, 5}}, 
 {17, {4, 5, 4, 5, 5, 5, 5, 5, 5, 6, 5, 5, 5, 5, 5, 6, 5, 5}}, 
 {17, {4, 5, 4, 5, 5, 5, 5, 5, 5, 5, 6, 5, 5, 5, 6, 5, 5, 5}}, 
 {17, {4, 5, 4, 5, 5, 5, 5, 5, 5, 5, 4, 5, 5, 6, 6, 6, 5, 5}}, 
 {17, {4, 5, 4, 5, 5, 4, 5, 5, 6, 6, 6, 5, 5, 5, 5, 5, 5, 5}}, 
 {17, {4, 4, 5, 5, 6, 6, 6, 5, 5, 5, 4, 5, 5, 5, 4, 5, 6, 5}}, 
 {17, {4, 4, 5, 5, 6, 6, 6, 5, 5, 4, 5, 5, 5, 4, 5, 5, 6, 5}}, 
 {17, {4, 4, 5, 5, 6, 6, 5, 4, 5, 5, 5, 4, 5, 5, 6, 6, 5, 5}}, 
 {17, {4, 4, 5, 5, 6, 5, 5, 5, 5, 6, 5, 5, 4, 5, 5, 5, 6, 5}}, 
 {17, {4, 4, 5, 5, 6, 5, 5, 5, 5, 5, 6, 5, 5, 5, 4, 5, 6, 5}}, 
 {17, {4, 4, 5, 5, 6, 5, 5, 5, 4, 5, 6, 5, 5, 5, 5, 6, 5, 5}}, 
 {17, {4, 4, 5, 5, 6, 5, 4, 5, 5, 5, 5, 5, 6, 5, 5, 6, 5, 5}}, 
 {17, {4, 4, 5, 5, 5, 6, 5, 5, 5, 6, 5, 5, 5, 5, 4, 5, 6, 5}}, 
 {17, {4, 4, 5, 5, 5, 6, 5, 5, 5, 6, 5, 5, 4, 5, 5, 6, 5, 5}}, 
 {17, {4, 4, 5, 5, 5, 6, 5, 5, 5, 5, 5, 5, 6, 5, 4, 5, 6, 5}}, 
 {17, {4, 4, 5, 5, 5, 6, 5, 5, 5, 4, 5, 6, 5, 5, 6, 5, 5, 5}}, 
 {17, {4, 4, 5, 5, 5, 6, 5, 5, 5, 4, 5, 5, 6, 5, 5, 6, 5, 5}}, 
 {17, {4, 4, 5, 5, 5, 6, 5, 5, 5, 4, 4, 5, 5, 5, 6, 6, 6, 5}}, 
 {17, {4, 4, 5, 5, 5, 6, 5, 5, 4, 4, 5, 5, 6, 6, 6, 5, 5, 5}}, 
 {17, {4, 4, 5, 5, 5, 6, 5, 4, 4, 5, 6, 5, 5, 5, 6, 6, 5, 5}}, 
 {17, {4, 4, 5, 5, 5, 5, 6, 5, 5, 6, 5, 4, 5, 5, 6, 5, 5, 5}}, 
 {17, {4, 4, 5, 5, 5, 5, 6, 5, 5, 6, 5, 4, 5, 5, 5, 6, 5, 5}}, 
 {17, {4, 4, 5, 5, 5, 5, 6, 5, 5, 5, 6, 5, 4, 5, 6, 5, 5, 5}}, 
 {17, {4, 4, 5, 5, 5, 5, 6, 5, 5, 4, 5, 5, 5, 6, 5, 5, 6, 5}}, 
 {17, {4, 4, 5, 5, 5, 5, 5, 6, 5, 5, 6, 5, 4, 5, 6, 5, 5, 5}}, 
 {17, {4, 4, 5, 5, 5, 5, 5, 6, 5, 5, 4, 4, 5, 5, 6, 6, 6, 5}}, 
 {17, {4, 4, 5, 5, 5, 5, 5, 6, 5, 4, 5, 5, 6, 5, 5, 5, 6, 5}}, 
 {17, {4, 4, 5, 5, 5, 5, 5, 5, 6, 5, 5, 6, 5, 4, 5, 5, 6, 5}}, 
 {17, {4, 4, 5, 5, 5, 5, 4, 5, 6, 6, 6, 5, 4, 5, 6, 5, 5, 5}}, 
 {17, {4, 4, 5, 5, 5, 4, 5, 6, 5, 5, 5, 6, 5, 5, 5, 6, 5, 5}}, 
 {17, {4, 4, 5, 5, 5, 4, 5, 6, 5, 5, 5, 4, 5, 5, 6, 6, 6, 5}}, 
 {17, {4, 4, 5, 5, 5, 4, 5, 5, 6, 6, 5, 5, 4, 5, 6, 6, 5, 5}}, 
 {17, {4, 4, 5, 5, 5, 4, 5, 5, 6, 5, 5, 5, 6, 5, 5, 5, 6, 5}}, 
 {17, {4, 4, 5, 5, 5, 4, 5, 5, 5, 6, 6, 6, 5, 4, 5, 5, 6, 5}}, 
 {17, {4, 4, 5, 5, 5, 4, 4, 5, 5, 5, 6, 5, 5, 5, 6, 6, 6, 5}}, 
 {17, {4, 4, 5, 5, 5, 4, 4, 5, 5, 5, 5, 6, 6, 5, 6, 6, 5, 5}}, 
 {17, {4, 4, 5, 5, 5, 4, 4, 5, 5, 5, 5, 6, 5, 5, 6, 6, 6, 5}}, 
 {17, {4, 4, 5, 5, 4, 5, 6, 6, 6, 5, 5, 4, 5, 6, 5, 5, 5, 5}}, 
 {17, {4, 4, 5, 5, 4, 5, 5, 6, 6, 5, 4, 5, 5, 6, 6, 5, 5, 5}}, 
 {17, {4, 4, 5, 5, 4, 5, 5, 6, 5, 4, 5, 5, 6, 6, 6, 5, 5, 5}}, 
 {17, {4, 4, 5, 5, 4, 5, 5, 5, 6, 6, 5, 5, 4, 5, 6, 6, 5, 5}}, 
 {17, {4, 4, 5, 5, 4, 5, 5, 5, 6, 5, 5, 5, 5, 6, 5, 5, 6, 5}}, 
 {17, {4, 4, 5, 5, 4, 5, 5, 5, 6, 5, 5, 4, 5, 5, 6, 6, 6, 5}}, 
 {17, {4, 4, 5, 5, 4, 5, 5, 4, 5, 6, 6, 6, 6, 5, 5, 5, 5, 5}}, 
 {17, {4, 4, 5, 5, 4, 4, 5, 5, 5, 5, 6, 6, 5, 5, 5, 6, 6, 5}}, 
 {17, {4, 4, 5, 5, 4, 4, 5, 5, 5, 5, 5, 6, 6, 5, 5, 6, 6, 5}}, 
 {17, {4, 4, 5, 5, 4, 4, 5, 5, 5, 5, 5, 5, 6, 6, 5, 6, 6, 5}}, 
 {17, {4, 4, 5, 4, 5, 5, 6, 6, 5, 5, 4, 5, 6, 6, 5, 5, 5, 5}}, 
 {17, {4, 4, 5, 4, 5, 5, 5, 5, 5, 6, 5, 5, 6, 6, 5, 5, 5, 5}}, 
 {17, {4, 4, 5, 4, 5, 5, 5, 4, 5, 5, 5, 5, 5, 6, 6, 6, 6, 5}}, 
 {17, {4, 4, 5, 4, 5, 5, 4, 5, 5, 6, 6, 5, 5, 5, 6, 6, 5, 5}}, 
 {17, {4, 4, 5, 4, 4, 5, 6, 5, 5, 5, 5, 5, 5, 5, 6, 6, 6, 5}}, 
 {17, {4, 4, 5, 4, 4, 5, 5, 5, 5, 6, 6, 5, 5, 5, 6, 6, 5, 5}}, 
 {17, {4, 4, 5, 4, 4, 5, 5, 5, 5, 5, 6, 6, 5, 5, 5, 6, 6, 5}}, 
 {17, {4, 4, 4, 4, 5, 5, 5, 5, 5, 5, 5, 6, 6, 6, 6, 5, 5, 5}}, 
 {17, {3, 5, 6, 5, 5, 5, 4, 5, 5, 5, 4, 5, 5, 5, 6, 6, 6, 5}}, 
 {17, {3, 5, 6, 5, 5, 4, 5, 5, 5, 4, 5, 5, 5, 5, 6, 6, 6, 5}}, 
 {17, {3, 5, 6, 5, 4, 5, 5, 5, 5, 5, 6, 5, 5, 5, 5, 5, 6, 5}}, 
 {17, {3, 5, 5, 6, 6, 5, 5, 4, 4, 5, 5, 5, 5, 5, 5, 6, 6, 5}}, 
 {17, {3, 5, 5, 6, 5, 5, 5, 5, 5, 5, 5, 4, 5, 5, 6, 6, 5, 5}}, 
 {17, {3, 5, 5, 6, 5, 5, 5, 5, 5, 5, 4, 5, 5, 5, 6, 6, 5, 5}}, 
 {17, {3, 5, 5, 5, 6, 6, 6, 5, 5, 5, 5, 5, 5, 4, 4, 5, 6, 5}}, 
 {17, {3, 5, 5, 5, 6, 6, 6, 5, 5, 5, 4, 4, 5, 5, 5, 5, 6, 5}}, 
 {17, {3, 5, 5, 5, 6, 6, 6, 5, 5, 4, 4, 5, 5, 5, 5, 5, 6, 5}}, 
 {17, {3, 5, 5, 5, 6, 6, 5, 5, 5, 5, 5, 4, 5, 5, 5, 6, 5, 5}}, 
 {17, {3, 5, 5, 5, 5, 6, 6, 5, 4, 5, 5, 5, 5, 5, 6, 5, 5, 5}}, 
 {17, {3, 5, 5, 5, 5, 6, 5, 5, 4, 5, 5, 5, 5, 5, 6, 6, 5, 5}}, 
 {17, {3, 5, 5, 5, 5, 5, 6, 5, 5, 5, 5, 5, 4, 5, 6, 6, 5, 5}}, 
 {17, {3, 5, 5, 5, 5, 5, 6, 5, 5, 4, 5, 6, 5, 5, 5, 5, 6, 5}}, 
 {17, {3, 5, 5, 5, 5, 5, 5, 6, 5, 5, 5, 5, 5, 4, 5, 6, 6, 5}}, 
 {17, {3, 5, 5, 5, 5, 5, 4, 5, 5, 6, 6, 6, 5, 5, 4, 5, 6, 5}}, 
 {17, {3, 5, 5, 5, 5, 5, 4, 5, 5, 6, 5, 5, 6, 6, 5, 5, 5, 5}}, 
 {17, {3, 5, 5, 5, 5, 4, 5, 6, 6, 5, 5, 5, 6, 5, 5, 5, 5, 5}}, 
 {17, {3, 5, 5, 5, 5, 4, 5, 6, 5, 5, 5, 5, 5, 6, 5, 5, 6, 5}}, 
 {17, {3, 5, 5, 5, 4, 5, 6, 5, 5, 6, 5, 5, 5, 5, 5, 5, 6, 5}}, 
 {17, {3, 5, 5, 5, 4, 5, 5, 6, 6, 5, 5, 5, 5, 5, 5, 6, 5, 5}}, 
 {17, {3, 5, 5, 5, 4, 5, 5, 5, 5, 6, 6, 5, 5, 6, 5, 5, 5, 5}}, 
 {17, {3, 5, 5, 5, 4, 5, 5, 5, 5, 4, 5, 5, 5, 6, 6, 6, 6, 5}}, 
 {17, {3, 5, 5, 5, 4, 5, 5, 5, 4, 5, 5, 6, 6, 6, 6, 5, 5, 5}}, 
 {17, {3, 5, 5, 5, 4, 5, 5, 4, 5, 5, 5, 5, 5, 6, 6, 6, 6, 5}}, 
 {17, {3, 5, 5, 5, 4, 4, 5, 6, 5, 5, 5, 5, 5, 6, 6, 6, 5, 5}}, 
 {17, {3, 5, 5, 4, 5, 6, 5, 5, 5, 5, 5, 6, 5, 5, 5, 5, 6, 5}}, 
 {17, {3, 5, 5, 4, 5, 6, 5, 5, 5, 5, 5, 5, 5, 6, 6, 5, 5, 5}}, 
 {17, {3, 5, 5, 4, 5, 5, 6, 6, 6, 6, 5, 5, 5, 4, 5, 5, 5, 5}}, 
 {17, {3, 5, 5, 4, 5, 5, 6, 6, 5, 5, 5, 5, 5, 5, 5, 6, 5, 5}}, 
 {17, {3, 5, 5, 4, 5, 5, 6, 5, 5, 5, 5, 5, 6, 6, 5, 5, 5, 5}}, 
 {17, {3, 5, 5, 4, 5, 5, 5, 6, 6, 6, 6, 5, 5, 5, 4, 5, 5, 5}}, 
 {17, {3, 5, 5, 4, 5, 5, 5, 6, 5, 5, 5, 5, 5, 6, 5, 5, 6, 5}}, 
 {17, {3, 5, 5, 4, 5, 5, 5, 5, 5, 6, 6, 6, 6, 5, 4, 5, 5, 5}}, 
 {17, {3, 5, 5, 4, 5, 5, 5, 4, 5, 5, 6, 6, 6, 5, 5, 5, 6, 5}}, 
 {17, {3, 5, 5, 4, 5, 5, 5, 4, 5, 5, 5, 5, 6, 6, 6, 6, 5, 5}}, 
 {17, {3, 5, 5, 4, 4, 5, 5, 5, 5, 5, 5, 6, 6, 6, 5, 5, 6, 5}}, 
 {17, {1, 6, 6, 6, 5, 5, 5, 5, 5, 3, 5, 5, 5, 5, 5, 6, 6, 6}}, 
 {17, {1, 6, 6, 6, 5, 5, 5, 4, 4, 5, 5, 4, 5, 5, 6, 6, 6, 6}}, 
 {17, {1, 6, 6, 6, 5, 5, 4, 4, 5, 4, 5, 5, 5, 5, 6, 6, 6, 6}}, 
 {17, {1, 6, 6, 6, 5, 4, 5, 5, 5, 5, 4, 4, 5, 5, 6, 6, 6, 6}}, 
 {17, {1, 6, 6, 5, 5, 5, 6, 5, 5, 5, 4, 4, 5, 5, 5, 6, 6, 6}}, 
 {17, {1, 6, 6, 5, 5, 5, 6, 5, 4, 5, 5, 5, 4, 5, 5, 6, 6, 6}}, 
 {17, {1, 6, 6, 5, 5, 5, 5, 6, 5, 3, 5, 5, 5, 5, 5, 6, 6, 6}}, 
 {17, {1, 6, 6, 5, 5, 5, 5, 5, 6, 5, 5, 5, 4, 4, 5, 6, 6, 6}}, 
 {17, {1, 6, 6, 5, 5, 5, 5, 5, 5, 4, 4, 5, 6, 5, 5, 6, 6, 6}}, 
 {17, {1, 6, 6, 5, 5, 5, 5, 5, 4, 5, 5, 6, 5, 4, 5, 6, 6, 6}}, 
 {17, {1, 6, 6, 5, 5, 5, 5, 5, 4, 4, 5, 6, 6, 5, 5, 5, 6, 6}}, 
 {17, {1, 6, 6, 5, 5, 5, 5, 5, 4, 4, 5, 5, 5, 6, 6, 5, 6, 6}}, 
 {17, {1, 6, 6, 5, 5, 5, 5, 4, 4, 5, 6, 6, 5, 5, 5, 5, 6, 6}}, 
 {17, {1, 6, 6, 5, 5, 5, 4, 5, 5, 5, 6, 5, 4, 5, 5, 6, 6, 6}}, 
 {17, {1, 6, 6, 5, 5, 4, 5, 6, 5, 5, 4, 5, 5, 5, 5, 6, 6, 6}}, 
 {17, {1, 6, 6, 5, 5, 4, 5, 5, 5, 5, 4, 5, 6, 5, 5, 6, 6, 6}}, 
 {17, {1, 6, 6, 5, 5, 4, 5, 5, 5, 4, 5, 5, 5, 6, 6, 5, 6, 6}}, 
 {17, {1, 6, 6, 5, 5, 4, 5, 5, 4, 5, 6, 5, 5, 5, 5, 6, 6, 6}}, 
 {17, {1, 6, 6, 5, 5, 4, 5, 5, 4, 4, 5, 5, 5, 6, 6, 6, 6, 6}}, 
 {17, {1, 6, 6, 5, 5, 4, 4, 5, 5, 5, 5, 5, 6, 6, 5, 5, 6, 6}}, 
 {17, {1, 6, 6, 5, 5, 3, 5, 5, 5, 5, 5, 5, 5, 5, 6, 6, 6, 6}}, 
 {17, {1, 6, 6, 5, 4, 5, 5, 6, 6, 5, 5, 4, 5, 5, 5, 5, 6, 6}}, 
 {17, {1, 6, 6, 5, 4, 5, 5, 5, 6, 5, 5, 5, 4, 5, 5, 6, 6, 6}}, 
 {17, {1, 6, 6, 5, 4, 5, 5, 5, 5, 6, 5, 4, 5, 5, 5, 6, 6, 6}}, 
 {17, {1, 6, 6, 5, 4, 5, 5, 5, 4, 5, 5, 5, 6, 5, 5, 6, 6, 6}}, 
 {17, {1, 6, 6, 5, 4, 4, 5, 5, 5, 5, 5, 5, 6, 5, 5, 6, 6, 6}}, 
 {17, {1, 6, 6, 5, 3, 5, 5, 5, 5, 6, 5, 5, 5, 5, 5, 6, 6, 6}}, 
 {17, {1, 6, 6, 1, 6, 6, 6, 5, 5, 4, 5, 6, 5, 5, 5, 6, 6, 6}}, 
 {17, {1, 6, 6, 1, 6, 6, 6, 5, 5, 4, 4, 5, 5, 6, 6, 6, 6, 6}}, 
 {17, {1, 6, 6, 1, 6, 6, 5, 4, 5, 6, 6, 5, 5, 5, 5, 6, 6, 6}}, 
 {17, {4, 5, 6, 5, 4, 5, 6, 5, 5, 5, 6, 6, 5, 6, 6, 6, 5}}, 
 {17, {4, 5, 5, 6, 6, 5, 6, 6, 6, 5, 5, 6, 5, 4, 5, 6, 5}}, 
 {17, {4, 5, 5, 6, 5, 5, 4, 5, 6, 6, 6, 5, 5, 6, 6, 6, 5}}, 
 {17, {4, 5, 5, 6, 5, 4, 5, 6, 6, 5, 6, 6, 6, 5, 5, 6, 5}}, 
 {17, {4, 5, 4, 5, 5, 5, 6, 6, 6, 6, 5, 5, 5, 6, 6, 6, 5}}, 
 {17, {4, 4, 5, 6, 6, 6, 5, 4, 5, 6, 5, 5, 6, 6, 6, 6, 5}}, 
 {17, {4, 4, 5, 6, 6, 5, 5, 6, 5, 4, 5, 6, 6, 6, 6, 6, 5}}, 
 {17, {4, 4, 5, 6, 6, 5, 4, 5, 6, 6, 5, 5, 6, 6, 6, 6, 5}}, 
 {17, {4, 4, 5, 6, 5, 4, 5, 6, 5, 5, 6, 6, 6, 6, 6, 6, 5}}, 
 {17, {4, 4, 5, 5, 4, 5, 6, 6, 5, 6, 6, 5, 6, 6, 6, 6, 5}}, 
 {17, {4, 4, 5, 4, 5, 6, 6, 6, 6, 6, 5, 6, 6, 6, 5, 5, 5}}, 
 {17, {4, 4, 5, 4, 5, 5, 6, 6, 6, 5, 6, 6, 6, 5, 6, 6, 5}}, 
 {17, {4, 4, 5, 4, 5, 5, 6, 6, 5, 6, 6, 6, 6, 5, 6, 6, 5}}, 
 {17, {3, 5, 6, 6, 5, 5, 6, 6, 6, 5, 4, 5, 5, 6, 6, 6, 5}}, 
 {17, {3, 5, 6, 6, 5, 5, 4, 4, 5, 6, 6, 6, 6, 6, 6, 6, 5}}, 
 {17, {3, 5, 6, 5, 5, 6, 5, 5, 6, 6, 5, 6, 6, 5, 5, 6, 5}}, 
 {17, {3, 5, 6, 5, 5, 6, 5, 5, 4, 5, 6, 6, 6, 6, 6, 6, 5}}, 
 {17, {3, 5, 5, 6, 6, 6, 6, 6, 5, 4, 5, 6, 6, 5, 5, 6, 5}}, 
 {17, {3, 5, 5, 6, 6, 6, 6, 5, 6, 6, 5, 5, 4, 5, 6, 6, 5}}, 
 {17, {3, 5, 5, 6, 6, 6, 6, 5, 4, 5, 6, 5, 5, 6, 6, 6, 5}}, 
 {17, {3, 5, 5, 6, 6, 6, 5, 5, 4, 5, 6, 6, 5, 6, 6, 6, 5}}, 
 {17, {3, 5, 5, 6, 6, 5, 4, 5, 6, 6, 6, 5, 5, 6, 6, 6, 5}}, 
 {17, {3, 5, 5, 6, 5, 5, 6, 5, 5, 6, 6, 5, 5, 6, 6, 6, 5}}, 
 {17, {3, 5, 5, 6, 5, 4, 5, 5, 6, 6, 6, 6, 5, 6, 6, 6, 5}}, 
 {17, {3, 5, 5, 5, 6, 6, 6, 6, 6, 6, 5, 4, 5, 6, 6, 5, 5}}, 
 {17, {3, 5, 5, 5, 6, 5, 5, 6, 6, 5, 5, 6, 6, 5, 6, 6, 5}}, 
 {17, {1, 6, 6, 6, 5, 3, 5, 5, 6, 6, 5, 6, 6, 6, 6, 6, 6}}, 
 {17, {1, 6, 6, 5, 6, 6, 6, 5, 5, 3, 5, 6, 6, 6, 6, 6, 6}}, 
 {17, {1, 6, 6, 5, 6, 6, 6, 1, 6, 6, 6, 6, 6, 6, 5, 6, 6}}, 
 {17, {1, 6, 6, 5, 6, 6, 5, 5, 4, 5, 6, 6, 6, 5, 6, 6, 6}}, 
 {17, {1, 6, 6, 5, 5, 6, 5, 4, 5, 6, 6, 6, 6, 5, 6, 6, 6}}, 
 {17, {1, 6, 6, 5, 4, 5, 6, 5, 5, 6, 6, 6, 6, 5, 6, 6, 6}}, 
 {16, {4, 4, 5, 5, 5, 4, 5, 5, 5, 5, 5, 5, 4, 4, 5, 5, 5, 5, 5}}, 
 {16, {4, 4, 5, 5, 5, 4, 5, 5, 5, 4, 5, 5, 5, 5, 5, 4, 5, 5, 5}}, 
 {16, {4, 4, 5, 5, 4, 5, 5, 5, 5, 5, 5, 4, 4, 5, 5, 5, 5, 5, 5}}, 
 {16, {4, 5, 5, 5, 6, 5, 4, 5, 5, 6, 5, 5, 5, 4, 5, 5, 6, 5}}, 
 {16, {4, 5, 5, 5, 5, 5, 6, 5, 5, 4, 5, 6, 5, 4, 5, 6, 5, 5}}, 
 {16, {4, 5, 5, 5, 4, 5, 6, 5, 5, 4, 5, 5, 5, 6, 5, 5, 6, 5}}, 
 {16, {4, 5, 5, 5, 4, 5, 5, 6, 5, 5, 4, 5, 5, 5, 5, 6, 6, 5}}, 
 {16, {4, 5, 5, 4, 5, 6, 5, 5, 5, 5, 5, 5, 5, 4, 5, 6, 6, 5}}, 
 {16, {4, 5, 5, 4, 5, 6, 5, 5, 5, 5, 4, 5, 5, 5, 5, 6, 6, 5}}, 
 {16, {4, 5, 5, 4, 5, 5, 5, 5, 5, 6, 6, 5, 4, 5, 5, 5, 6, 5}}, 
 {16, {4, 5, 5, 4, 5, 5, 5, 5, 4, 5, 5, 5, 6, 6, 5, 5, 6, 5}}, 
 {16, {4, 5, 5, 4, 5, 5, 4, 5, 6, 5, 5, 5, 5, 5, 5, 6, 6, 5}}, 
 {16, {4, 5, 4, 5, 5, 5, 6, 5, 5, 5, 5, 5, 5, 5, 6, 5, 5, 5}}, 
 {16, {4, 5, 4, 5, 5, 5, 5, 5, 5, 5, 6, 5, 5, 6, 5, 5, 5, 5}}, 
 {16, {4, 5, 4, 5, 5, 5, 5, 4, 5, 5, 5, 5, 5, 6, 6, 6, 5, 5}}, 
 {16, {4, 5, 4, 5, 5, 5, 4, 5, 5, 6, 6, 6, 5, 5, 5, 5, 5, 5}}, 
 {16, {4, 4, 5, 6, 5, 5, 5, 5, 4, 5, 5, 5, 6, 5, 5, 5, 6, 5}}, 
 {16, {4, 4, 5, 6, 5, 5, 5, 4, 5, 5, 5, 5, 6, 5, 5, 5, 6, 5}}, 
 {16, {4, 4, 5, 6, 5, 5, 4, 5, 5, 5, 4, 5, 5, 5, 6, 6, 6, 5}}, 
 {16, {4, 4, 5, 5, 6, 6, 5, 4, 4, 5, 6, 5, 5, 5, 5, 5, 6, 5}}, 
 {16, {4, 4, 5, 5, 6, 5, 5, 6, 5, 5, 5, 5, 5, 5, 4, 5, 6, 5}}, 
 {16, {4, 4, 5, 5, 6, 5, 5, 5, 6, 5, 5, 5, 4, 5, 5, 5, 6, 5}}, 
 {16, {4, 4, 5, 5, 6, 5, 5, 5, 5, 5, 4, 5, 6, 5, 5, 5, 6, 5}}, 
 {16, {4, 4, 5, 5, 6, 5, 5, 5, 5, 4, 5, 6, 5, 5, 5, 6, 5, 5}}, 
 {16, {4, 4, 5, 5, 6, 5, 5, 5, 5, 4, 5, 6, 5, 5, 5, 5, 6, 5}}, 
 {16, {4, 4, 5, 5, 6, 5, 5, 5, 4, 5, 6, 5, 5, 5, 5, 5, 6, 5}}, 
 {16, {4, 4, 5, 5, 6, 5, 5, 5, 4, 5, 5, 6, 5, 5, 5, 5, 6, 5}}, 
 {16, {4, 4, 5, 5, 6, 5, 5, 4, 5, 5, 6, 5, 5, 5, 5, 6, 5, 5}}, 
 {16, {4, 4, 5, 5, 6, 5, 5, 4, 5, 5, 6, 5, 5, 5, 5, 5, 6, 5}}, 
 {16, {4, 4, 5, 5, 6, 5, 5, 4, 5, 5, 5, 5, 6, 5, 5, 5, 6, 5}}, 
 {16, {4, 4, 5, 5, 6, 5, 4, 5, 5, 6, 5, 5, 5, 5, 5, 6, 5, 5}}, 
 {16, {4, 4, 5, 5, 6, 5, 4, 5, 5, 5, 4, 5, 5, 6, 6, 6, 5, 5}}, 
 {16, {4, 4, 5, 5, 5, 6, 6, 6, 5, 5, 4, 5, 5, 4, 5, 5, 6, 5}}, 
 {16, {4, 4, 5, 5, 5, 6, 5, 5, 4, 5, 6, 5, 5, 5, 5, 6, 5, 5}}, 
 {16, {4, 4, 5, 5, 5, 6, 5, 4, 5, 5, 5, 5, 6, 5, 5, 6, 5, 5}}, 
 {16, {4, 4, 5, 5, 5, 5, 6, 6, 6, 5, 4, 4, 5, 5, 5, 6, 5, 5}}, 
 {16, {4, 4, 5, 5, 5, 5, 6, 6, 5, 4, 4, 5, 6, 6, 5, 5, 5, 5}}, 
 {16, {4, 4, 5, 5, 5, 5, 6, 5, 5, 6, 5, 5, 5, 5, 4, 5, 6, 5}}, 
 {16, {4, 4, 5, 5, 5, 5, 6, 5, 4, 5, 6, 5, 5, 6, 5, 5, 5, 5}}, 
 {16, {4, 4, 5, 5, 5, 5, 5, 5, 6, 5, 4, 5, 5, 6, 5, 5, 6, 5}}, 
 {16, {4, 4, 5, 5, 5, 5, 5, 5, 5, 6, 5, 4, 5, 6, 5, 5, 6, 5}}, 
 {16, {4, 4, 5, 5, 5, 5, 5, 5, 4, 5, 6, 5, 5, 6, 5, 5, 6, 5}}, 
 {16, {4, 4, 5, 5, 5, 5, 5, 4, 5, 6, 5, 5, 6, 5, 5, 5, 6, 5}}, 
 {16, {4, 4, 5, 5, 5, 5, 4, 5, 6, 6, 6, 5, 5, 4, 5, 5, 6, 5}}, 
 {16, {4, 4, 5, 5, 5, 5, 4, 5, 6, 5, 5, 4, 5, 5, 6, 6, 6, 5}}, 
 {16, {4, 4, 5, 5, 5, 5, 4, 5, 5, 6, 6, 6, 5, 4, 5, 6, 5, 5}}, 
 {16, {4, 4, 5, 5, 5, 5, 4, 5, 5, 6, 5, 4, 5, 6, 6, 6, 5, 5}}, 
 {16, {4, 4, 5, 5, 5, 5, 4, 5, 5, 6, 5, 4, 5, 5, 6, 6, 6, 5}}, 
 {16, {4, 4, 5, 5, 5, 4, 5, 6, 6, 5, 5, 4, 5, 5, 6, 6, 5, 5}}, 
 {16, {4, 4, 5, 5, 5, 4, 5, 6, 5, 5, 5, 6, 5, 5, 6, 5, 5, 5}}, 
 {16, {4, 4, 5, 5, 5, 4, 5, 5, 6, 6, 5, 4, 5, 6, 6, 5, 5, 5}}, 
 {16, {4, 4, 5, 5, 5, 4, 5, 5, 6, 6, 5, 4, 5, 5, 5, 6, 6, 5}}, 
 {16, {4, 4, 5, 5, 5, 4, 5, 5, 5, 6, 5, 5, 5, 6, 5, 5, 6, 5}}, 
 {16, {4, 4, 5, 5, 5, 4, 5, 5, 4, 5, 5, 6, 6, 5, 5, 6, 6, 5}}, 
 {16, {4, 4, 5, 5, 4, 5, 6, 6, 5, 5, 4, 5, 5, 6, 6, 5, 5, 5}}, 
 {16, {4, 4, 5, 5, 4, 5, 6, 5, 5, 6, 5, 5, 6, 5, 5, 5, 5, 5}}, 
 {16, {4, 4, 5, 5, 4, 5, 6, 5, 5, 5, 6, 5, 5, 5, 5, 5, 6, 5}}, 
 {16, {4, 4, 5, 5, 4, 5, 6, 5, 5, 5, 4, 5, 5, 5, 6, 6, 6, 5}}, 
 {16, {4, 4, 5, 5, 4, 5, 5, 6, 6, 5, 5, 4, 5, 6, 6, 5, 5, 5}}, 
 {16, {4, 4, 5, 5, 4, 5, 5, 6, 5, 5, 5, 6, 5, 5, 5, 6, 5, 5}}, 
 {16, {4, 4, 5, 5, 4, 5, 5, 5, 6, 6, 6, 5, 5, 5, 4, 5, 6, 5}}, 
 {16, {4, 4, 5, 5, 4, 5, 5, 5, 6, 6, 6, 5, 4, 5, 5, 5, 6, 5}}, 
 {16, {4, 4, 5, 5, 4, 5, 5, 5, 6, 5, 5, 4, 5, 6, 6, 6, 5, 5}}, 
 {16, {4, 4, 5, 5, 4, 5, 5, 5, 5, 6, 5, 5, 6, 5, 5, 6, 5, 5}}, 
 {16, {4, 4, 5, 5, 4, 5, 5, 5, 5, 5, 6, 5, 5, 6, 5, 5, 6, 5}}, 
 {16, {4, 4, 5, 5, 4, 5, 5, 5, 4, 5, 6, 5, 5, 6, 6, 6, 5, 5}}, 
 {16, {4, 4, 5, 5, 4, 5, 5, 4, 5, 5, 6, 6, 6, 5, 5, 5, 6, 5}}, 
 {16, {4, 4, 5, 5, 4, 4, 5, 5, 5, 5, 6, 5, 5, 5, 6, 6, 6, 5}}, 
 {16, {4, 4, 5, 4, 5, 5, 5, 5, 5, 4, 5, 5, 5, 6, 6, 6, 6, 5}}, 
 {16, {4, 4, 5, 4, 5, 5, 5, 4, 5, 5, 5, 5, 6, 6, 6, 6, 5, 5}}, 
 {16, {4, 4, 5, 4, 4, 5, 5, 5, 5, 6, 5, 5, 5, 6, 6, 6, 5, 5}}, 
 {16, {3, 5, 6, 6, 5, 5, 5, 5, 4, 4, 5, 5, 5, 5, 5, 6, 6, 5}}, 
 {16, {3, 5, 6, 6, 5, 4, 5, 5, 5, 5, 5, 5, 4, 5, 5, 6, 6, 5}}, 
 {16, {3, 5, 6, 5, 4, 5, 5, 6, 5, 5, 5, 5, 5, 5, 5, 5, 6, 5}}, 
 {16, {3, 5, 5, 6, 5, 5, 5, 4, 5, 5, 5, 5, 5, 5, 6, 6, 5, 5}}, 
 {16, {3, 5, 5, 6, 5, 4, 5, 5, 5, 5, 5, 6, 5, 5, 5, 5, 6, 5}}, 
 {16, {3, 5, 5, 5, 6, 6, 5, 5, 5, 5, 5, 5, 5, 5, 4, 5, 6, 5}}, 
 {16, {3, 5, 5, 5, 6, 6, 5, 5, 5, 4, 5, 5, 5, 5, 5, 6, 5, 5}}, 
 {16, {3, 5, 5, 5, 6, 5, 5, 5, 5, 5, 5, 4, 5, 6, 6, 5, 5, 5}}, 
 {16, {3, 5, 5, 5, 5, 6, 5, 5, 5, 5, 5, 6, 5, 4, 5, 5, 6, 5}}, 
 {16, {3, 5, 5, 5, 5, 6, 5, 4, 5, 5, 5, 6, 5, 5, 5, 5, 6, 5}}, 
 {16, {3, 5, 5, 5, 5, 6, 5, 4, 5, 5, 5, 5, 5, 6, 6, 5, 5, 5}}, 
 {16, {3, 5, 5, 5, 5, 5, 6, 6, 6, 5, 4, 4, 5, 5, 5, 5, 6, 5}}, 
 {16, {3, 5, 5, 5, 5, 5, 6, 6, 5, 5, 5, 4, 5, 5, 6, 5, 5, 5}}, 
 {16, {3, 5, 5, 5, 5, 5, 6, 5, 5, 6, 5, 5, 5, 4, 5, 5, 6, 5}}, 
 {16, {3, 5, 5, 5, 5, 5, 5, 6, 6, 5, 4, 5, 5, 6, 5, 5, 5, 5}}, 
 {16, {3, 5, 5, 5, 5, 5, 5, 5, 6, 6, 5, 4, 5, 5, 6, 5, 5, 5}}, 
 {16, {3, 5, 5, 5, 5, 5, 5, 5, 5, 6, 5, 4, 5, 5, 5, 6, 6, 5}}, 
 {16, {3, 5, 5, 5, 5, 5, 5, 4, 5, 6, 5, 5, 5, 6, 6, 5, 5, 5}}, 
 {16, {3, 5, 5, 5, 5, 5, 4, 5, 6, 5, 5, 5, 6, 6, 5, 5, 5, 5}}, 
 {16, {3, 5, 5, 5, 5, 5, 4, 5, 5, 5, 6, 6, 5, 5, 5, 6, 5, 5}}, 
 {16, {3, 5, 5, 5, 5, 4, 5, 5, 5, 6, 5, 5, 6, 6, 5, 5, 5, 5}}, 
 {16, {3, 5, 5, 5, 5, 4, 5, 5, 5, 6, 5, 5, 6, 5, 5, 5, 6, 5}}, 
 {16, {3, 5, 5, 5, 5, 4, 5, 5, 5, 5, 5, 6, 5, 5, 6, 6, 5, 5}}, 
 {16, {3, 5, 5, 5, 5, 4, 5, 5, 4, 5, 5, 5, 6, 6, 6, 6, 5, 5}}, 
 {16, {3, 5, 5, 5, 5, 4, 5, 5, 4, 5, 5, 5, 5, 6, 6, 6, 6, 5}}, 
 {16, {3, 5, 5, 5, 4, 5, 6, 6, 5, 5, 6, 5, 5, 5, 5, 5, 5, 5}}, 
 {16, {3, 5, 5, 5, 4, 5, 5, 6, 6, 6, 5, 5, 5, 4, 5, 5, 6, 5}}, 
 {16, {3, 5, 5, 5, 4, 5, 5, 5, 6, 5, 4, 5, 5, 6, 6, 6, 5, 5}}, 
 {16, {3, 5, 5, 5, 4, 4, 5, 6, 5, 5, 5, 5, 6, 6, 6, 5, 5, 5}}, 
 {16, {3, 5, 5, 4, 5, 6, 5, 5, 6, 6, 5, 5, 5, 5, 5, 5, 5, 5}}, 
 {16, {3, 5, 5, 4, 5, 5, 5, 6, 6, 5, 5, 5, 6, 5, 5, 5, 5, 5}}, 
 {16, {3, 5, 5, 4, 4, 5, 5, 6, 6, 6, 5, 5, 5, 6, 5, 5, 5, 5}}, 
 {16, {1, 6, 6, 6, 5, 5, 5, 5, 4, 4, 5, 5, 4, 5, 6, 6, 6, 6}}, 
 {16, {1, 6, 6, 6, 5, 5, 5, 4, 4, 5, 5, 5, 4, 5, 6, 6, 6, 6}}, 
 {16, {1, 6, 6, 6, 5, 4, 5, 5, 5, 5, 5, 5, 5, 4, 5, 6, 6, 6}}, 
 {16, {1, 6, 6, 5, 5, 5, 6, 6, 6, 6, 1, 6, 6, 6, 1, 6, 6, 6}}, 
 {16, {1, 6, 6, 5, 5, 5, 5, 5, 5, 4, 4, 5, 6, 6, 5, 5, 6, 6}}, 
 {16, {1, 6, 6, 5, 5, 5, 4, 4, 5, 5, 4, 5, 5, 6, 6, 6, 6, 6}}, 
 {16, {1, 6, 6, 5, 5, 4, 5, 5, 6, 5, 5, 5, 4, 5, 5, 6, 6, 6}}, 
 {16, {1, 6, 6, 5, 5, 4, 4, 5, 5, 5, 5, 5, 5, 6, 6, 5, 6, 6}}, 
 {16, {1, 6, 6, 5, 5, 3, 5, 6, 5, 5, 5, 5, 5, 5, 5, 6, 6, 6}}, 
 {16, {1, 6, 6, 5, 4, 5, 5, 5, 5, 5, 6, 6, 6, 6, 1, 6, 6, 6}}, 
 {16, {1, 6, 6, 5, 4, 5, 5, 5, 5, 5, 6, 5, 4, 5, 5, 6, 6, 6}}, 
 {16, {1, 6, 6, 5, 4, 5, 5, 5, 4, 5, 6, 5, 5, 5, 5, 6, 6, 6}}, 
 {16, {1, 6, 6, 5, 4, 4, 5, 6, 6, 5, 5, 5, 5, 5, 5, 5, 6, 6}}, 
 {16, {1, 6, 6, 5, 4, 4, 5, 5, 5, 5, 5, 5, 5, 6, 6, 5, 6, 6}}, 
 {16, {1, 6, 6, 5, 4, 4, 5, 5, 5, 5, 4, 5, 5, 6, 6, 6, 6, 6}}, 
 {16, {1, 6, 6, 5, 3, 5, 5, 6, 6, 5, 5, 5, 5, 5, 5, 5, 6, 6}}, 
 {16, {1, 6, 6, 5, 3, 5, 5, 5, 5, 6, 6, 5, 5, 5, 5, 5, 6, 6}}, 
 {16, {1, 6, 6, 1, 6, 6, 5, 5, 5, 4, 5, 5, 5, 6, 6, 6, 6, 6}}, 
 {16, {4, 5, 6, 5, 5, 6, 6, 5, 4, 5, 6, 6, 6, 5, 5, 6, 5}}, 
 {16, {4, 5, 5, 6, 6, 5, 6, 6, 6, 5, 4, 5, 6, 5, 5, 6, 5}}, 
 {16, {4, 5, 5, 6, 6, 5, 5, 6, 5, 4, 5, 6, 6, 5, 6, 6, 5}}, 
 {16, {4, 5, 5, 6, 5, 5, 6, 5, 4, 5, 6, 6, 6, 5, 6, 6, 5}}, 
 {16, {4, 5, 5, 4, 5, 6, 6, 6, 5, 5, 6, 5, 5, 6, 6, 6, 5}}, 
 {16, {4, 5, 5, 4, 5, 6, 5, 4, 5, 6, 6, 6, 6, 6, 6, 6, 5}}, 
 {16, {4, 5, 4, 5, 5, 6, 6, 6, 5, 6, 6, 6, 5, 5, 5, 6, 5}}, 
 {16, {4, 5, 4, 5, 5, 6, 6, 6, 5, 5, 5, 6, 6, 5, 6, 6, 5}}, 
 {16, {4, 5, 4, 5, 5, 5, 6, 6, 6, 6, 5, 6, 6, 5, 5, 6, 5}}, 
 {16, {4, 4, 5, 5, 6, 6, 5, 4, 5, 6, 6, 6, 6, 5, 6, 6, 5}}, 
 {16, {4, 4, 5, 5, 6, 6, 5, 4, 5, 6, 6, 5, 6, 6, 6, 6, 5}}, 
 {16, {4, 4, 5, 5, 6, 5, 4, 5, 6, 6, 6, 6, 5, 6, 6, 6, 5}}, 
 {16, {3, 5, 6, 6, 6, 5, 5, 6, 5, 5, 4, 5, 6, 6, 6, 6, 5}}, 
 {16, {3, 5, 6, 6, 5, 4, 5, 6, 6, 6, 6, 5, 5, 5, 6, 6, 5}}, 
 {16, {3, 5, 6, 6, 5, 4, 5, 5, 6, 6, 5, 5, 6, 6, 6, 6, 5}}, 
 {16, {3, 5, 6, 6, 5, 4, 5, 5, 6, 5, 5, 6, 6, 6, 6, 6, 5}}, 
 {16, {3, 5, 5, 6, 6, 6, 6, 6, 6, 5, 5, 6, 5, 4, 5, 6, 5}}, 
 {16, {3, 5, 5, 6, 6, 6, 6, 5, 6, 6, 5, 4, 5, 5, 6, 6, 5}}, 
 {16, {3, 5, 5, 6, 6, 6, 6, 5, 5, 6, 5, 4, 5, 6, 6, 6, 5}}, 
 {16, {3, 5, 5, 6, 6, 6, 5, 6, 6, 6, 5, 4, 5, 5, 6, 6, 5}}, 
 {16, {3, 5, 5, 6, 6, 6, 5, 4, 5, 6, 6, 5, 5, 6, 6, 6, 5}}, 
 {16, {3, 5, 5, 6, 6, 5, 6, 6, 6, 6, 5, 4, 5, 5, 6, 6, 5}}, 
 {16, {3, 5, 5, 6, 6, 5, 5, 6, 6, 6, 6, 6, 5, 4, 5, 6, 5}}, 
 {16, {3, 5, 5, 6, 6, 5, 4, 5, 6, 6, 6, 6, 6, 5, 5, 6, 5}}, 
 {16, {3, 5, 5, 6, 6, 5, 4, 5, 6, 6, 6, 6, 5, 5, 6, 6, 5}}, 
 {16, {3, 5, 5, 5, 6, 6, 6, 6, 6, 6, 6, 5, 4, 5, 6, 5, 5}}, 
 {16, {3, 5, 5, 5, 6, 6, 6, 5, 6, 6, 5, 5, 6, 5, 5, 6, 5}}, 
 {16, {3, 5, 5, 5, 6, 5, 5, 6, 6, 5, 6, 6, 5, 5, 6, 6, 5}}, 
 {16, {3, 5, 5, 5, 6, 5, 5, 5, 6, 6, 5, 6, 6, 5, 6, 6, 5}}, 
 {16, {3, 5, 5, 5, 5, 6, 6, 5, 4, 5, 6, 6, 6, 6, 6, 6, 5}}, 
 {16, {3, 5, 5, 5, 5, 6, 5, 5, 6, 6, 5, 6, 6, 5, 6, 6, 5}}, 
 {16, {3, 5, 5, 4, 5, 6, 6, 6, 6, 6, 6, 6, 5, 5, 6, 5, 5}}, 
 {16, {1, 6, 6, 6, 5, 6, 6, 5, 5, 3, 5, 6, 6, 6, 6, 6, 6}}, 
 {16, {1, 6, 6, 6, 5, 4, 4, 5, 6, 6, 6, 6, 5, 6, 6, 6, 6}}, 
 {16, {1, 6, 6, 6, 5, 3, 5, 6, 5, 5, 6, 6, 6, 6, 6, 6, 6}}, 
 {16, {1, 6, 6, 5, 6, 6, 6, 6, 6, 5, 4, 4, 5, 6, 6, 6, 6}}, 
 {16, {1, 6, 6, 5, 6, 6, 6, 5, 4, 5, 6, 5, 5, 6, 6, 6, 6}}, 
 {16, {1, 6, 6, 5, 6, 6, 5, 4, 5, 6, 6, 6, 5, 5, 6, 6, 6}}, 
 {16, {1, 6, 6, 5, 6, 6, 5, 4, 4, 5, 6, 6, 6, 6, 6, 6, 6}}, 
 {16, {1, 6, 6, 5, 5, 6, 5, 4, 5, 6, 6, 6, 6, 6, 5, 6, 6}}, 
 {16, {1, 6, 6, 5, 4, 4, 5, 6, 6, 6, 6, 6, 6, 6, 5, 6, 6}}, 
 {16, {1, 6, 1, 6, 6, 6, 6, 6, 6, 5, 5, 6, 6, 6, 6, 6, 6}}, 
 {15, {4, 4, 5, 5, 5, 4, 5, 5, 5, 4, 5, 5, 5, 4, 5, 5, 5, 5, 5}}, 
 {15, {4, 4, 5, 5, 4, 5, 5, 4, 5, 5, 5, 4, 5, 5, 5, 5, 5, 5, 5}}, 
 {15, {4, 4, 5, 4, 5, 5, 5, 5, 5, 5, 5, 5, 4, 4, 5, 5, 5, 5, 5}}, 
 {15, {4, 4, 5, 4, 4, 5, 5, 5, 5, 5, 4, 5, 5, 5, 5, 5, 5, 5, 5}}, 
 {15, {3, 5, 5, 5, 5, 4, 4, 5, 5, 5, 5, 5, 5, 5, 4, 5, 5, 5, 5}}, 
 {15, {4, 5, 5, 5, 5, 6, 5, 5, 4, 5, 6, 5, 5, 5, 4, 5, 6, 5}}, 
 {15, {4, 5, 5, 5, 5, 6, 5, 4, 5, 6, 5, 5, 5, 4, 5, 5, 6, 5}}, 
 {15, {4, 5, 5, 5, 5, 6, 5, 4, 5, 6, 5, 5, 4, 5, 5, 5, 6, 5}}, 
 {15, {4, 5, 5, 5, 5, 6, 5, 4, 5, 5, 6, 5, 4, 5, 5, 6, 5, 5}}, 
 {15, {4, 5, 5, 5, 5, 5, 6, 5, 4, 5, 6, 5, 5, 4, 5, 6, 5, 5}}, 
 {15, {4, 5, 5, 5, 5, 5, 4, 5, 5, 5, 6, 5, 5, 4, 5, 6, 6, 5}}, 
 {15, {4, 5, 5, 5, 5, 4, 5, 6, 5, 4, 5, 5, 5, 5, 5, 6, 6, 5}}, 
 {15, {4, 5, 5, 5, 4, 5, 5, 6, 5, 5, 6, 5, 4, 5, 5, 5, 6, 5}}, 
 {15, {4, 5, 5, 5, 4, 5, 5, 6, 5, 5, 4, 5, 6, 5, 5, 5, 6, 5}}, 
 {15, {4, 5, 5, 5, 4, 5, 5, 5, 5, 4, 5, 5, 6, 5, 5, 6, 6, 5}}, 
 {15, {4, 5, 5, 4, 5, 6, 5, 5, 4, 5, 5, 5, 6, 5, 5, 5, 6, 5}}, 
 {15, {4, 5, 5, 4, 5, 5, 6, 5, 5, 5, 6, 5, 5, 4, 5, 5, 6, 5}}, 
 {15, {4, 5, 5, 4, 5, 5, 5, 6, 5, 5, 5, 5, 4, 5, 5, 6, 6, 5}}, 
 {15, {4, 5, 5, 4, 5, 5, 5, 5, 6, 6, 5, 5, 5, 5, 4, 5, 6, 5}}, 
 {15, {4, 5, 5, 4, 5, 5, 5, 5, 5, 5, 5, 6, 5, 4, 5, 6, 6, 5}}, 
 {15, {4, 5, 5, 4, 5, 5, 5, 5, 5, 5, 4, 5, 6, 6, 5, 5, 6, 5}}, 
 {15, {4, 5, 5, 4, 5, 5, 5, 5, 5, 4, 5, 6, 6, 5, 5, 5, 6, 5}}, 
 {15, {4, 5, 5, 4, 5, 5, 4, 5, 5, 5, 5, 6, 5, 5, 5, 6, 6, 5}}, 
 {15, {4, 5, 5, 4, 5, 5, 4, 5, 5, 5, 4, 5, 6, 6, 6, 6, 5, 5}}, 
 {15, {4, 5, 4, 5, 5, 5, 5, 5, 6, 5, 5, 5, 5, 5, 5, 6, 5, 5}}, 
 {15, {4, 5, 4, 5, 5, 5, 4, 5, 5, 5, 6, 6, 6, 5, 5, 5, 5, 5}}, 
 {15, {4, 4, 5, 6, 5, 4, 5, 5, 5, 4, 5, 5, 5, 5, 6, 6, 6, 5}}, 
 {15, {4, 4, 5, 5, 6, 6, 6, 5, 5, 5, 5, 4, 5, 5, 4, 5, 6, 5}}, 
 {15, {4, 4, 5, 5, 6, 6, 6, 5, 5, 4, 5, 5, 4, 5, 5, 5, 6, 5}}, 
 {15, {4, 4, 5, 5, 6, 6, 5, 5, 4, 5, 5, 4, 5, 5, 6, 6, 5, 5}}, 
 {15, {4, 4, 5, 5, 6, 5, 5, 5, 5, 6, 6, 5, 5, 4, 4, 5, 6, 5}}, 
 {15, {4, 4, 5, 5, 6, 5, 4, 5, 5, 5, 5, 6, 5, 5, 5, 5, 6, 5}}, 
 {15, {4, 4, 5, 5, 6, 5, 4, 5, 5, 5, 5, 4, 5, 5, 6, 6, 6, 5}}, 
 {15, {4, 4, 5, 5, 5, 6, 6, 5, 5, 4, 5, 5, 4, 5, 6, 6, 5, 5}}, 
 {15, {4, 4, 5, 5, 5, 6, 5, 5, 6, 5, 5, 4, 5, 5, 5, 6, 5, 5}}, 
 {15, {4, 4, 5, 5, 5, 6, 5, 5, 6, 5, 5, 4, 5, 5, 5, 5, 6, 5}}, 
 {15, {4, 4, 5, 5, 5, 6, 5, 5, 4, 5, 5, 5, 6, 5, 5, 5, 6, 5}}, 
 {15, {4, 4, 5, 5, 5, 6, 5, 4, 5, 5, 6, 5, 5, 5, 6, 5, 5, 5}}, 
 {15, {4, 4, 5, 5, 5, 6, 5, 4, 5, 5, 5, 5, 6, 5, 5, 5, 6, 5}}, 
 {15, {4, 4, 5, 5, 5, 5, 6, 6, 5, 4, 5, 5, 4, 5, 5, 6, 6, 5}}, 
 {15, {4, 4, 5, 5, 5, 5, 6, 5, 5, 5, 5, 4, 5, 6, 5, 5, 6, 5}}, 
 {15, {4, 4, 5, 5, 5, 5, 6, 5, 5, 5, 4, 5, 6, 5, 5, 6, 5, 5}}, 
 {15, {4, 4, 5, 5, 5, 5, 6, 5, 5, 4, 5, 6, 5, 5, 5, 6, 5, 5}}, 
 {15, {4, 4, 5, 5, 5, 5, 6, 5, 4, 5, 5, 6, 5, 5, 6, 5, 5, 5}}, 
 {15, {4, 4, 5, 5, 5, 5, 5, 6, 6, 6, 5, 4, 4, 5, 5, 6, 5, 5}}, 
 {15, {4, 4, 5, 5, 5, 5, 5, 6, 5, 5, 6, 5, 5, 4, 5, 6, 5, 5}}, 
 {15, {4, 4, 5, 5, 5, 5, 5, 6, 5, 5, 4, 5, 6, 5, 5, 6, 5, 5}}, 
 {15, {4, 4, 5, 5, 5, 5, 5, 5, 4, 4, 5, 6, 6, 5, 5, 6, 6, 5}}, 
 {15, {4, 4, 5, 5, 5, 5, 5, 5, 4, 4, 5, 6, 5, 5, 6, 6, 6, 5}}, 
 {15, {4, 4, 5, 5, 5, 5, 5, 5, 4, 4, 5, 5, 6, 6, 5, 6, 6, 5}}, 
 {15, {4, 4, 5, 5, 5, 5, 4, 5, 6, 6, 5, 4, 5, 5, 6, 6, 5, 5}}, 
 {15, {4, 4, 5, 5, 5, 5, 4, 5, 6, 5, 5, 6, 5, 5, 6, 5, 5, 5}}, 
 {15, {4, 4, 5, 5, 5, 5, 4, 5, 6, 5, 5, 4, 5, 6, 6, 6, 5, 5}}, 
 {15, {4, 4, 5, 5, 5, 5, 4, 4, 5, 5, 5, 6, 5, 5, 6, 6, 6, 5}}, 
 {15, {4, 4, 5, 5, 5, 4, 5, 6, 6, 6, 5, 5, 5, 4, 5, 5, 6, 5}}, 
 {15, {4, 4, 5, 5, 5, 4, 5, 6, 6, 5, 5, 4, 5, 5, 5, 6, 6, 5}}, 
 {15, {4, 4, 5, 5, 5, 4, 5, 6, 5, 5, 5, 5, 6, 5, 5, 5, 6, 5}}, 
 {15, {4, 4, 5, 5, 5, 4, 5, 6, 5, 4, 5, 5, 5, 6, 6, 6, 5, 5}}, 
 {15, {4, 4, 5, 5, 5, 4, 5, 5, 6, 6, 6, 5, 4, 5, 6, 5, 5, 5}}, 
 {15, {4, 4, 5, 5, 5, 4, 5, 5, 6, 5, 5, 4, 5, 6, 6, 6, 5, 5}}, 
 {15, {4, 4, 5, 5, 5, 4, 5, 5, 5, 6, 6, 6, 5, 5, 4, 5, 6, 5}}, 
 {15, {4, 4, 5, 5, 5, 4, 5, 5, 5, 6, 5, 4, 5, 5, 6, 6, 6, 5}}, 
 {15, {4, 4, 5, 5, 5, 4, 4, 5, 5, 5, 6, 6, 6, 5, 5, 6, 5, 5}}, 
 {15, {4, 4, 5, 5, 4, 5, 6, 5, 5, 6, 5, 5, 5, 5, 5, 5, 6, 5}}, 
 {15, {4, 4, 5, 5, 4, 5, 6, 5, 5, 5, 6, 5, 5, 5, 5, 6, 5, 5}}, 
 {15, {4, 4, 5, 5, 4, 5, 6, 5, 5, 5, 5, 6, 6, 6, 5, 4, 5, 5}}, 
 {15, {4, 4, 5, 5, 4, 5, 6, 5, 5, 4, 5, 6, 6, 6, 5, 5, 5, 5}}, 
 {15, {4, 4, 5, 5, 4, 5, 5, 6, 6, 6, 5, 5, 5, 6, 5, 4, 5, 5}}, 
 {15, {4, 4, 5, 5, 4, 5, 5, 6, 6, 6, 5, 5, 4, 5, 5, 6, 5, 5}}, 
 {15, {4, 4, 5, 5, 4, 5, 5, 6, 6, 5, 6, 6, 5, 5, 4, 5, 5, 5}}, 
 {15, {4, 4, 5, 5, 4, 5, 5, 5, 6, 6, 5, 6, 6, 5, 5, 4, 5, 5}}, 
 {15, {4, 4, 5, 5, 4, 5, 5, 5, 6, 5, 5, 5, 6, 5, 5, 6, 5, 5}}, 
 {15, {4, 4, 5, 5, 4, 5, 5, 5, 5, 5, 5, 4, 5, 6, 6, 6, 6, 5}}, 
 {15, {4, 4, 5, 4, 5, 5, 6, 6, 5, 5, 4, 5, 5, 5, 5, 6, 6, 5}}, 
 {15, {4, 4, 5, 4, 5, 5, 6, 5, 5, 6, 6, 5, 5, 5, 5, 5, 5, 5}}, 
 {15, {4, 4, 5, 4, 5, 5, 5, 4, 5, 5, 6, 6, 6, 5, 5, 5, 6, 5}}, 
 {15, {4, 4, 5, 4, 5, 5, 5, 4, 5, 5, 5, 6, 6, 6, 6, 5, 5, 5}}, 
 {15, {4, 4, 5, 4, 4, 5, 5, 5, 6, 5, 5, 5, 6, 6, 6, 5, 5, 5}}, 
 {15, {4, 4, 5, 4, 4, 5, 5, 5, 5, 5, 6, 5, 5, 6, 6, 6, 5, 5}}, 
 {15, {4, 4, 5, 4, 4, 5, 5, 5, 5, 5, 5, 6, 6, 6, 5, 5, 6, 5}}, 
 {15, {4, 4, 5, 4, 4, 5, 5, 5, 5, 5, 5, 6, 6, 5, 5, 6, 6, 5}}, 
 {15, {4, 4, 4, 4, 5, 5, 5, 5, 5, 5, 6, 6, 6, 6, 5, 5, 5, 5}}, 
 {15, {3, 5, 6, 6, 5, 5, 4, 4, 5, 5, 5, 5, 5, 5, 5, 6, 6, 5}}, 
 {15, {3, 5, 6, 5, 5, 5, 5, 4, 5, 5, 5, 4, 5, 5, 6, 6, 6, 5}}, 
 {15, {3, 5, 6, 5, 4, 5, 5, 5, 5, 5, 5, 6, 5, 5, 5, 5, 6, 5}}, 
 {15, {3, 5, 5, 6, 6, 6, 5, 5, 5, 5, 4, 5, 5, 4, 5, 5, 6, 5}}, 
 {15, {3, 5, 5, 6, 6, 5, 5, 4, 4, 5, 5, 5, 5, 5, 6, 6, 5, 5}}, 
 {15, {3, 5, 5, 6, 5, 5, 5, 5, 5, 5, 5, 5, 5, 5, 5, 6, 5, 5}}, 
 {15, {3, 5, 5, 5, 6, 6, 5, 5, 4, 4, 5, 5, 5, 5, 6, 6, 5, 5}}, 
 {15, {3, 5, 5, 5, 6, 5, 5, 6, 5, 4, 5, 5, 5, 5, 5, 5, 6, 5}}, 
 {15, {3, 5, 5, 5, 6, 5, 5, 5, 5, 6, 5, 5, 4, 5, 5, 5, 6, 5}}, 
 {15, {3, 5, 5, 5, 6, 5, 5, 5, 5, 6, 5, 4, 5, 5, 5, 5, 6, 5}}, 
 {15, {3, 5, 5, 5, 6, 5, 5, 5, 5, 4, 4, 5, 5, 6, 6, 6, 5, 5}}, 
 {15, {3, 5, 5, 5, 5, 6, 6, 5, 5, 5, 5, 5, 5, 4, 5, 6, 5, 5}}, 
 {15, {3, 5, 5, 5, 5, 6, 5, 4, 5, 5, 5, 5, 6, 6, 5, 5, 5, 5}}, 
 {15, {3, 5, 5, 5, 5, 6, 5, 4, 4, 5, 5, 5, 5, 6, 6, 6, 5, 5}}, 
 {15, {3, 5, 5, 5, 5, 5, 6, 6, 5, 5, 5, 4, 4, 5, 6, 6, 5, 5}}, 
 {15, {3, 5, 5, 5, 5, 5, 6, 6, 5, 5, 5, 4, 4, 5, 5, 6, 6, 5}}, 
 {15, {3, 5, 5, 5, 5, 5, 6, 6, 5, 4, 4, 5, 5, 5, 6, 6, 5, 5}}, 
 {15, {3, 5, 5, 5, 5, 5, 6, 5, 5, 5, 4, 5, 6, 6, 5, 5, 5, 5}}, 
 {15, {3, 5, 5, 5, 5, 5, 6, 5, 5, 4, 5, 6, 6, 5, 5, 5, 5, 5}}, 
 {15, {3, 5, 5, 5, 5, 5, 6, 5, 4, 4, 5, 5, 5, 5, 6, 6, 6, 5}}, 
 {15, {3, 5, 5, 5, 5, 5, 5, 6, 6, 6, 5, 5, 5, 4, 4, 5, 6, 5}}, 
 {15, {3, 5, 5, 5, 5, 5, 5, 6, 5, 5, 5, 4, 5, 6, 6, 5, 5, 5}}, 
 {15, {3, 5, 5, 5, 5, 5, 5, 6, 5, 4, 5, 5, 6, 5, 5, 5, 6, 5}}, 
 {15, {3, 5, 5, 5, 5, 5, 5, 6, 5, 4, 5, 5, 5, 6, 5, 5, 6, 5}}, 
 {15, {3, 5, 5, 5, 5, 5, 5, 6, 5, 4, 4, 5, 5, 6, 6, 6, 5, 5}}, 
 {15, {3, 5, 5, 5, 5, 5, 5, 5, 6, 6, 6, 5, 5, 4, 4, 5, 6, 5}}, 
 {15, {3, 5, 5, 5, 5, 5, 5, 5, 6, 6, 5, 5, 5, 4, 5, 6, 5, 5}}, 
 {15, {3, 5, 5, 5, 5, 5, 5, 5, 5, 5, 5, 6, 5, 4, 5, 6, 6, 5}}, 
 {15, {3, 5, 5, 5, 5, 5, 5, 4, 5, 6, 6, 5, 5, 6, 5, 5, 5, 5}}, 
 {15, {3, 5, 5, 5, 5, 5, 5, 4, 4, 5, 6, 6, 6, 6, 5, 5, 5, 5}}, 
 {15, {3, 5, 5, 5, 5, 5, 5, 3, 5, 5, 5, 5, 5, 6, 6, 6, 6, 5}}, 
 {15, {3, 5, 5, 5, 5, 5, 4, 5, 5, 6, 6, 5, 5, 5, 6, 5, 5, 5}}, 
 {15, {3, 5, 5, 5, 5, 5, 4, 5, 5, 6, 5, 5, 5, 5, 5, 6, 6, 5}}, 
 {15, {3, 5, 5, 5, 5, 5, 4, 5, 5, 5, 5, 5, 6, 5, 5, 6, 6, 5}}, 
 {15, {3, 5, 5, 5, 5, 5, 4, 5, 5, 5, 5, 4, 5, 6, 6, 6, 6, 5}}, 
 {15, {3, 5, 5, 5, 5, 4, 5, 5, 6, 6, 6, 5, 5, 4, 5, 5, 6, 5}}, 
 {15, {3, 5, 5, 5, 5, 4, 5, 5, 6, 6, 5, 5, 5, 5, 5, 6, 5, 5}}, 
 {15, {3, 5, 5, 5, 5, 4, 5, 5, 6, 6, 5, 5, 4, 5, 6, 6, 5, 5}}, 
 {15, {3, 5, 5, 5, 5, 4, 5, 5, 6, 5, 4, 5, 5, 6, 6, 6, 5, 5}}, 
 {15, {3, 5, 5, 5, 5, 4, 5, 5, 5, 5, 6, 6, 5, 5, 6, 5, 5, 5}}, 
 {15, {3, 5, 5, 5, 5, 4, 5, 5, 5, 5, 5, 4, 5, 6, 6, 6, 6, 5}}, 
 {15, {3, 5, 5, 5, 5, 4, 4, 5, 5, 5, 5, 6, 6, 6, 6, 5, 5, 5}}, 
 {15, {3, 5, 5, 5, 4, 5, 6, 6, 5, 5, 5, 4, 5, 5, 6, 6, 5, 5}}, 
 {15, {3, 5, 5, 5, 4, 5, 5, 6, 5, 5, 6, 5, 5, 5, 5, 5, 6, 5}}, 
 {15, {3, 5, 5, 5, 4, 5, 5, 5, 6, 6, 6, 5, 5, 4, 5, 5, 6, 5}}, 
 {15, {3, 5, 5, 5, 4, 5, 5, 5, 6, 6, 6, 5, 4, 5, 5, 5, 6, 5}}, 
 {15, {3, 5, 5, 5, 4, 5, 5, 5, 6, 6, 5, 5, 5, 6, 5, 5, 5, 5}}, 
 {15, {3, 5, 5, 5, 4, 5, 5, 5, 4, 5, 6, 6, 6, 6, 5, 5, 5, 5}}, 
 {15, {3, 5, 5, 5, 4, 4, 5, 6, 6, 6, 6, 5, 5, 5, 5, 5, 5, 5}}, 
 {15, {3, 5, 5, 5, 4, 4, 5, 5, 6, 6, 5, 5, 5, 5, 6, 6, 5, 5}}, 
 {15, {3, 5, 5, 5, 4, 4, 5, 5, 5, 6, 5, 5, 5, 6, 6, 6, 5, 5}}, 
 {15, {3, 5, 5, 5, 3, 5, 5, 5, 5, 6, 6, 6, 5, 5, 5, 5, 6, 5}}, 
 {15, {3, 5, 5, 4, 5, 5, 6, 6, 6, 5, 5, 5, 6, 5, 4, 5, 5, 5}}, 
 {15, {3, 5, 5, 4, 5, 5, 6, 6, 6, 5, 5, 5, 4, 5, 5, 5, 6, 5}}, 
 {15, {3, 5, 5, 4, 5, 5, 5, 6, 6, 5, 5, 5, 5, 5, 5, 6, 5, 5}}, 
 {15, {3, 5, 5, 4, 5, 5, 5, 5, 6, 6, 6, 6, 5, 5, 5, 4, 5, 5}}, 
 {15, {3, 5, 5, 4, 5, 5, 5, 5, 6, 5, 5, 5, 6, 6, 5, 5, 5, 5}}, 
 {15, {3, 5, 5, 4, 5, 5, 5, 5, 6, 5, 5, 5, 5, 6, 6, 5, 5, 5}}, 
 {15, {3, 5, 5, 4, 5, 5, 5, 5, 5, 6, 6, 6, 6, 5, 5, 4, 5, 5}}, 
 {15, {3, 5, 5, 4, 4, 5, 5, 5, 6, 6, 5, 5, 5, 5, 6, 6, 5, 5}}, 
 {15, {3, 5, 5, 3, 5, 5, 5, 5, 5, 5, 6, 6, 6, 6, 5, 5, 5, 5}}, 
 {15, {1, 6, 6, 6, 5, 4, 4, 5, 5, 5, 4, 5, 5, 5, 6, 6, 6, 6}}, 
 {15, {1, 6, 6, 6, 5, 4, 4, 5, 5, 4, 5, 5, 5, 5, 6, 6, 6, 6}}, 
 {15, {1, 6, 6, 5, 5, 6, 5, 5, 5, 5, 4, 4, 5, 5, 5, 6, 6, 6}}, 
 {15, {1, 6, 6, 5, 5, 5, 5, 5, 5, 5, 5, 5, 3, 5, 6, 6, 6, 6}}, 
 {15, {1, 6, 6, 5, 5, 5, 5, 5, 5, 5, 3, 5, 5, 5, 6, 6, 6, 6}}, 
 {15, {1, 6, 6, 5, 5, 5, 5, 5, 5, 4, 4, 5, 5, 6, 6, 5, 6, 6}}, 
 {15, {1, 6, 6, 5, 5, 5, 4, 5, 5, 6, 5, 5, 4, 5, 5, 6, 6, 6}}, 
 {15, {1, 6, 6, 5, 5, 5, 4, 4, 5, 6, 6, 5, 5, 5, 5, 5, 6, 6}}, 
 {15, {1, 6, 6, 5, 5, 5, 4, 4, 5, 5, 5, 6, 6, 5, 5, 5, 6, 6}}, 
 {15, {1, 6, 6, 5, 5, 5, 3, 5, 5, 5, 5, 5, 5, 5, 6, 6, 6, 6}}, 
 {15, {1, 6, 6, 5, 5, 4, 5, 6, 5, 4, 5, 5, 5, 5, 5, 6, 6, 6}}, 
 {15, {1, 6, 6, 5, 5, 4, 4, 5, 5, 5, 5, 5, 6, 5, 5, 6, 6, 6}}, 
 {15, {1, 6, 6, 5, 5, 3, 5, 6, 6, 5, 5, 5, 5, 5, 5, 5, 6, 6}}, 
 {15, {1, 6, 6, 5, 4, 5, 5, 6, 6, 5, 5, 5, 5, 4, 5, 5, 6, 6}}, 
 {15, {1, 6, 6, 5, 4, 5, 5, 5, 6, 6, 5, 5, 4, 5, 5, 5, 6, 6}}, 
 {15, {1, 6, 6, 5, 4, 5, 5, 5, 5, 6, 6, 5, 5, 4, 5, 5, 6, 6}}, 
 {15, {1, 6, 6, 5, 4, 5, 5, 5, 4, 4, 5, 5, 5, 6, 6, 6, 6, 6}}, 
 {15, {1, 6, 6, 5, 4, 4, 5, 5, 5, 5, 5, 6, 5, 5, 5, 6, 6, 6}}, 
 {15, {1, 6, 6, 5, 3, 5, 5, 5, 5, 5, 5, 5, 6, 5, 5, 6, 6, 6}}, 
 {15, {1, 6, 6, 1, 6, 6, 5, 5, 6, 6, 5, 5, 4, 5, 5, 6, 6, 6}}, 
 {15, {4, 5, 4, 5, 5, 5, 6, 6, 5, 6, 6, 6, 5, 6, 6, 5, 5}}, 
 {15, {4, 5, 4, 5, 5, 5, 5, 6, 6, 6, 5, 5, 6, 6, 6, 6, 5}}, 
 {15, {4, 4, 5, 5, 6, 6, 6, 6, 5, 6, 6, 5, 4, 5, 6, 6, 5}}, 
 {15, {4, 4, 5, 5, 6, 5, 4, 5, 6, 6, 5, 6, 6, 6, 6, 6, 5}}, 
 {15, {4, 4, 5, 5, 4, 5, 6, 6, 6, 5, 6, 6, 5, 6, 6, 6, 5}}, 
 {15, {4, 4, 5, 4, 5, 6, 6, 5, 6, 6, 6, 6, 6, 6, 5, 5, 5}}, 
 {15, {3, 5, 6, 6, 6, 6, 5, 4, 4, 5, 5, 6, 6, 6, 6, 6, 5}}, 
 {15, {3, 5, 6, 6, 6, 5, 5, 5, 4, 5, 6, 6, 5, 6, 6, 6, 5}}, 
 {15, {3, 5, 6, 6, 5, 5, 4, 5, 6, 6, 6, 5, 5, 6, 6, 6, 5}}, 
 {15, {3, 5, 6, 6, 5, 4, 5, 6, 5, 5, 5, 6, 6, 6, 6, 6, 5}}, 
 {15, {3, 5, 6, 5, 5, 5, 4, 5, 6, 6, 6, 6, 6, 5, 6, 6, 5}}, 
 {15, {3, 5, 6, 5, 5, 4, 4, 5, 6, 6, 6, 6, 6, 6, 6, 6, 5}}, 
 {15, {3, 5, 6, 5, 4, 5, 6, 5, 5, 5, 6, 6, 6, 6, 6, 6, 5}}, 
 {15, {3, 5, 5, 6, 6, 6, 6, 6, 6, 5, 4, 5, 6, 5, 5, 6, 5}}, 
 {15, {3, 5, 5, 6, 6, 6, 6, 6, 6, 5, 4, 4, 5, 6, 6, 6, 5}}, 
 {15, {3, 5, 5, 6, 6, 6, 6, 6, 5, 5, 6, 6, 5, 4, 5, 6, 5}}, 
 {15, {3, 5, 5, 6, 6, 6, 6, 5, 5, 6, 6, 6, 5, 4, 5, 6, 5}}, 
 {15, {3, 5, 5, 6, 6, 6, 6, 5, 5, 6, 6, 5, 4, 5, 6, 6, 5}}, 
 {15, {3, 5, 5, 6, 6, 6, 5, 3, 5, 6, 6, 6, 6, 6, 6, 5, 5}}, 
 {15, {3, 5, 5, 6, 6, 5, 6, 6, 5, 5, 5, 6, 5, 5, 6, 6, 5}}, 
 {15, {3, 5, 5, 6, 6, 5, 6, 6, 5, 4, 5, 5, 6, 6, 6, 6, 5}}, 
 {15, {3, 5, 5, 6, 6, 5, 4, 5, 5, 6, 6, 6, 6, 5, 6, 6, 5}}, 
 {15, {3, 5, 5, 6, 5, 5, 5, 6, 6, 6, 5, 6, 6, 5, 5, 6, 5}}, 
 {15, {3, 5, 5, 6, 5, 5, 5, 6, 6, 5, 6, 6, 6, 5, 5, 6, 5}}, 
 {15, {3, 5, 5, 6, 5, 4, 5, 5, 6, 6, 6, 5, 6, 6, 6, 6, 5}}, 
 {15, {3, 5, 5, 6, 5, 3, 5, 6, 6, 6, 6, 6, 6, 6, 6, 5, 5}}, 
 {15, {3, 5, 5, 5, 6, 5, 5, 6, 5, 5, 6, 6, 5, 6, 6, 6, 5}}, 
 {15, {1, 6, 6, 6, 6, 5, 5, 3, 5, 6, 6, 5, 6, 6, 6, 6, 6}}, 
 {15, {1, 6, 6, 6, 5, 6, 6, 5, 3, 5, 5, 6, 6, 6, 6, 6, 6}}, 
 {15, {1, 6, 6, 5, 6, 6, 6, 5, 3, 5, 5, 6, 6, 6, 6, 6, 6}}, 
 {15, {1, 6, 6, 5, 6, 6, 5, 3, 5, 5, 6, 6, 6, 6, 6, 6, 6}}, 
 {15, {1, 6, 6, 5, 5, 6, 5, 4, 5, 6, 6, 5, 6, 6, 6, 6, 6}}, 
 {15, {1, 6, 6, 5, 5, 3, 5, 6, 6, 6, 6, 5, 6, 6, 6, 6, 6}}, 
 {15, {1, 6, 6, 5, 4, 5, 6, 6, 5, 5, 6, 6, 6, 5, 6, 6, 6}}, 
 {15, {1, 6, 6, 5, 4, 4, 5, 6, 6, 6, 6, 5, 6, 6, 6, 6, 6}}, 
 {15, {1, 6, 6, 5, 3, 5, 5, 6, 6, 6, 6, 5, 6, 6, 6, 6, 6}}, 
 {14, {4, 4, 5, 5, 5, 5, 4, 4, 5, 5, 5, 5, 5, 4, 5, 5, 5, 5, 5}}, 
 {14, {4, 5, 5, 5, 5, 6, 5, 5, 4, 5, 6, 5, 4, 5, 6, 5, 5, 5}}, 
 {14, {4, 5, 5, 5, 5, 6, 5, 4, 5, 6, 5, 5, 5, 4, 5, 6, 5, 5}}, 
 {14, {4, 5, 5, 5, 5, 5, 6, 5, 5, 4, 5, 6, 5, 4, 5, 5, 6, 5}}, 
 {14, {4, 5, 5, 5, 5, 5, 5, 4, 5, 5, 5, 6, 5, 4, 5, 6, 6, 5}}, 
 {14, {4, 5, 5, 5, 5, 5, 4, 5, 6, 5, 5, 5, 5, 4, 5, 6, 6, 5}}, 
 {14, {4, 5, 5, 5, 5, 5, 4, 5, 5, 5, 5, 6, 6, 5, 4, 5, 6, 5}}, 
 {14, {4, 5, 5, 5, 4, 5, 5, 5, 6, 5, 4, 5, 5, 6, 5, 5, 6, 5}}, 
 {14, {4, 5, 5, 4, 5, 5, 6, 5, 4, 5, 5, 5, 6, 5, 5, 6, 5, 5}}, 
 {14, {4, 5, 5, 4, 5, 5, 5, 4, 5, 6, 6, 5, 5, 6, 5, 5, 5, 5}}, 
 {14, {4, 5, 5, 4, 5, 5, 4, 5, 5, 5, 6, 5, 5, 5, 5, 6, 6, 5}}, 
 {14, {4, 5, 5, 4, 5, 5, 4, 5, 5, 5, 5, 4, 5, 6, 6, 6, 6, 5}}, 
 {14, {4, 5, 4, 5, 5, 5, 5, 6, 5, 5, 5, 5, 5, 5, 5, 6, 5, 5}}, 
 {14, {4, 5, 4, 5, 5, 5, 5, 5, 6, 5, 5, 5, 5, 5, 6, 5, 5, 5}}, 
 {14, {4, 5, 4, 5, 5, 5, 5, 5, 5, 6, 5, 4, 5, 5, 6, 6, 5, 5}}, 
 {14, {4, 5, 4, 5, 5, 5, 5, 4, 5, 6, 6, 6, 5, 5, 5, 5, 5, 5}}, 
 {14, {4, 5, 4, 5, 5, 5, 4, 5, 5, 5, 5, 5, 6, 6, 6, 5, 5, 5}}, 
 {14, {4, 4, 5, 6, 5, 4, 5, 5, 5, 5, 5, 5, 6, 5, 5, 5, 6, 5}}, 
 {14, {4, 4, 5, 6, 5, 4, 5, 5, 5, 5, 5, 4, 5, 5, 6, 6, 6, 5}}, 
 {14, {4, 4, 5, 6, 5, 4, 4, 5, 6, 5, 5, 5, 5, 5, 5, 6, 6, 5}}, 
 {14, {4, 4, 5, 5, 6, 5, 5, 5, 6, 5, 4, 5, 5, 5, 5, 5, 6, 5}}, 
 {14, {4, 4, 5, 5, 6, 5, 5, 5, 5, 6, 5, 5, 5, 4, 5, 5, 6, 5}}, 
 {14, {4, 4, 5, 5, 6, 5, 5, 5, 5, 5, 5, 4, 5, 6, 5, 5, 6, 5}}, 
 {14, {4, 4, 5, 5, 6, 5, 5, 5, 4, 5, 5, 5, 6, 5, 5, 6, 5, 5}}, 
 {14, {4, 4, 5, 5, 6, 5, 5, 4, 5, 6, 5, 5, 5, 5, 5, 6, 5, 5}}, 
 {14, {4, 4, 5, 5, 6, 5, 4, 5, 6, 5, 5, 5, 5, 5, 5, 6, 5, 5}}, 
 {14, {4, 4, 5, 5, 6, 5, 4, 5, 5, 5, 4, 5, 5, 5, 6, 6, 6, 5}}, 
 {14, {4, 4, 5, 5, 6, 5, 4, 4, 5, 5, 6, 6, 5, 5, 5, 5, 6, 5}}, 
 {14, {4, 4, 5, 5, 5, 6, 6, 6, 5, 4, 5, 5, 5, 5, 4, 5, 6, 5}}, 
 {14, {4, 4, 5, 5, 5, 6, 6, 5, 5, 4, 4, 5, 5, 5, 6, 6, 5, 5}}, 
 {14, {4, 4, 5, 5, 5, 6, 6, 5, 4, 5, 5, 4, 5, 5, 6, 6, 5, 5}}, 
 {14, {4, 4, 5, 5, 5, 6, 5, 5, 5, 6, 5, 5, 5, 4, 5, 5, 6, 5}}, 
 {14, {4, 4, 5, 5, 5, 6, 5, 5, 5, 5, 6, 5, 5, 5, 4, 5, 6, 5}}, 
 {14, {4, 4, 5, 5, 5, 6, 5, 5, 5, 5, 4, 5, 6, 5, 5, 6, 5, 5}}, 
 {14, {4, 4, 5, 5, 5, 6, 5, 5, 5, 4, 5, 6, 5, 5, 5, 5, 6, 5}}, 
 {14, {4, 4, 5, 5, 5, 6, 5, 5, 4, 5, 5, 6, 5, 5, 6, 5, 5, 5}}, 
 {14, {4, 4, 5, 5, 5, 6, 5, 5, 4, 5, 5, 4, 5, 6, 6, 6, 5, 5}}, 
 {14, {4, 4, 5, 5, 5, 6, 5, 4, 5, 5, 5, 6, 5, 5, 5, 6, 5, 5}}, 
 {14, {4, 4, 5, 5, 5, 6, 5, 4, 5, 5, 5, 4, 5, 5, 6, 6, 6, 5}}, 
 {14, {4, 4, 5, 5, 5, 5, 6, 6, 6, 5, 4, 5, 5, 4, 5, 5, 6, 5}}, 
 {14, {4, 4, 5, 5, 5, 5, 6, 6, 5, 5, 4, 5, 5, 4, 5, 6, 6, 5}}, 
 {14, {4, 4, 5, 5, 5, 5, 6, 5, 5, 6, 5, 5, 4, 5, 6, 5, 5, 5}}, 
 {14, {4, 4, 5, 5, 5, 5, 6, 5, 5, 5, 6, 5, 4, 5, 5, 6, 5, 5}}, 
 {14, {4, 4, 5, 5, 5, 5, 6, 5, 5, 5, 5, 5, 6, 5, 4, 5, 6, 5}}, 
 {14, {4, 4, 5, 5, 5, 5, 6, 5, 4, 4, 5, 6, 5, 5, 6, 6, 5, 5}}, 
 {14, {4, 4, 5, 5, 5, 5, 5, 6, 5, 5, 6, 5, 5, 4, 5, 5, 6, 5}}, 
 {14, {4, 4, 5, 5, 5, 5, 5, 6, 5, 5, 5, 5, 6, 5, 4, 5, 6, 5}}, 
 {14, {4, 4, 5, 5, 5, 5, 5, 5, 6, 5, 5, 6, 5, 4, 5, 6, 5, 5}}, 
 {14, {4, 4, 5, 5, 5, 5, 5, 4, 5, 6, 6, 5, 4, 5, 5, 6, 6, 5}}, 
 {14, {4, 4, 5, 5, 5, 5, 4, 5, 6, 6, 6, 5, 5, 4, 5, 6, 5, 5}}, 
 {14, {4, 4, 5, 5, 5, 5, 4, 5, 6, 6, 5, 5, 4, 5, 6, 6, 5, 5}}, 
 {14, {4, 4, 5, 5, 5, 5, 4, 4, 5, 5, 6, 5, 5, 6, 6, 6, 5, 5}}, 
 {14, {4, 4, 5, 5, 5, 5, 4, 4, 5, 5, 5, 5, 6, 6, 5, 6, 6, 5}}, 
 {14, {4, 4, 5, 5, 5, 4, 5, 6, 6, 6, 5, 5, 4, 5, 5, 6, 5, 5}}, 
 {14, {4, 4, 5, 5, 5, 4, 5, 6, 6, 6, 5, 4, 5, 5, 5, 5, 6, 5}}, 
 {14, {4, 4, 5, 5, 5, 4, 5, 6, 6, 5, 5, 5, 4, 5, 5, 6, 6, 5}}, 
 {14, {4, 4, 5, 5, 5, 4, 5, 6, 5, 5, 6, 6, 6, 5, 4, 5, 5, 5}}, 
 {14, {4, 4, 5, 5, 5, 4, 5, 6, 5, 5, 6, 5, 5, 5, 6, 5, 5, 5}}, 
 {14, {4, 4, 5, 5, 5, 4, 5, 6, 5, 5, 5, 4, 5, 6, 6, 6, 5, 5}}, 
 {14, {4, 4, 5, 5, 5, 4, 5, 6, 5, 4, 5, 5, 6, 6, 6, 5, 5, 5}}, 
 {14, {4, 4, 5, 5, 5, 4, 5, 5, 6, 5, 5, 6, 5, 5, 5, 5, 6, 5}}, 
 {14, {4, 4, 5, 5, 5, 4, 5, 5, 6, 5, 5, 4, 5, 5, 6, 6, 6, 5}}, 
 {14, {4, 4, 5, 5, 5, 4, 5, 5, 5, 5, 5, 4, 5, 6, 6, 6, 6, 5}}, 
 {14, {4, 4, 5, 5, 5, 4, 5, 5, 4, 5, 5, 6, 6, 6, 5, 5, 6, 5}}, 
 {14, {4, 4, 5, 5, 5, 4, 4, 5, 5, 6, 6, 5, 5, 5, 6, 6, 5, 5}}, 
 {14, {4, 4, 5, 5, 4, 5, 6, 6, 6, 5, 4, 5, 5, 6, 5, 5, 5, 5}}, 
 {14, {4, 4, 5, 5, 4, 5, 6, 6, 5, 5, 4, 5, 5, 5, 5, 6, 6, 5}}, 
 {14, {4, 4, 5, 5, 4, 5, 5, 6, 6, 6, 5, 5, 5, 5, 4, 5, 6, 5}}, 
 {14, {4, 4, 5, 5, 4, 5, 5, 6, 6, 6, 5, 4, 5, 5, 6, 5, 5, 5}}, 
 {14, {4, 4, 5, 5, 4, 5, 5, 6, 5, 5, 5, 5, 6, 5, 5, 6, 5, 5}}, 
 {14, {4, 4, 5, 5, 4, 5, 5, 6, 5, 5, 5, 4, 5, 5, 6, 6, 6, 5}}, 
 {14, {4, 4, 5, 5, 4, 5, 5, 5, 6, 6, 5, 5, 5, 4, 5, 6, 6, 5}}, 
 {14, {4, 4, 5, 5, 4, 5, 5, 5, 6, 6, 5, 4, 5, 5, 6, 6, 5, 5}}, 
 {14, {4, 4, 5, 5, 4, 5, 5, 5, 6, 5, 4, 5, 5, 5, 6, 6, 6, 5}}, 
 {14, {4, 4, 5, 5, 4, 5, 5, 5, 5, 4, 5, 5, 6, 6, 5, 6, 6, 5}}, 
 {14, {4, 4, 5, 5, 4, 4, 5, 5, 5, 5, 5, 6, 5, 5, 6, 6, 6, 5}}, 
 {14, {4, 4, 5, 4, 5, 6, 6, 6, 5, 5, 4, 5, 5, 5, 5, 5, 6, 5}}, 
 {14, {4, 4, 5, 4, 5, 6, 6, 5, 5, 5, 5, 5, 6, 5, 5, 5, 5, 5}}, 
 {14, {4, 4, 5, 4, 5, 6, 5, 5, 5, 6, 5, 5, 5, 5, 5, 5, 6, 5}}, 
 {14, {4, 4, 5, 4, 5, 5, 6, 6, 5, 5, 5, 5, 4, 5, 5, 6, 6, 5}}, 
 {14, {4, 4, 5, 4, 5, 5, 6, 6, 5, 4, 5, 5, 6, 6, 5, 5, 5, 5}}, 
 {14, {4, 4, 5, 4, 5, 5, 5, 6, 6, 6, 5, 5, 4, 5, 5, 5, 6, 5}}, 
 {14, {4, 4, 5, 4, 5, 5, 5, 6, 5, 5, 4, 5, 5, 6, 6, 6, 5, 5}}, 
 {14, {4, 4, 5, 4, 5, 5, 5, 5, 5, 6, 6, 6, 5, 5, 4, 5, 6, 5}}, 
 {14, {4, 4, 5, 4, 5, 5, 5, 5, 5, 5, 6, 6, 6, 6, 5, 4, 5, 5}}, 
 {14, {4, 4, 5, 4, 5, 5, 5, 5, 5, 5, 4, 5, 6, 6, 6, 6, 5, 5}}, 
 {14, {4, 4, 5, 4, 5, 5, 5, 5, 4, 5, 5, 5, 5, 6, 6, 6, 6, 5}}, 
 {14, {4, 4, 5, 4, 4, 5, 6, 5, 5, 5, 5, 6, 6, 5, 5, 5, 6, 5}}, 
 {14, {4, 4, 4, 4, 5, 5, 5, 5, 5, 5, 5, 5, 6, 6, 6, 6, 5, 5}}, 
 {14, {3, 5, 6, 5, 5, 5, 5, 5, 5, 5, 5, 4, 4, 5, 6, 6, 6, 5}}, 
 {14, {3, 5, 6, 5, 5, 5, 4, 5, 5, 5, 5, 5, 4, 5, 6, 6, 6, 5}}, 
 {14, {3, 5, 6, 5, 5, 4, 5, 5, 5, 5, 5, 5, 5, 6, 5, 5, 6, 5}}, 
 {14, {3, 5, 6, 5, 5, 4, 4, 5, 5, 5, 5, 6, 6, 5, 5, 5, 6, 5}}, 
 {14, {3, 5, 6, 5, 4, 4, 5, 6, 6, 5, 5, 5, 5, 5, 5, 5, 6, 5}}, 
 {14, {3, 5, 5, 6, 6, 5, 5, 5, 5, 5, 6, 5, 4, 4, 5, 5, 6, 5}}, 
 {14, {3, 5, 5, 6, 6, 5, 5, 5, 5, 5, 5, 4, 4, 5, 5, 6, 6, 5}}, 
 {14, {3, 5, 5, 6, 6, 5, 5, 5, 5, 4, 4, 5, 5, 5, 5, 6, 6, 5}}, 
 {14, {3, 5, 5, 6, 6, 5, 4, 4, 5, 5, 5, 5, 5, 5, 6, 6, 5, 5}}, 
 {14, {3, 5, 5, 6, 5, 5, 5, 5, 5, 5, 6, 5, 4, 5, 5, 5, 6, 5}}, 
 {14, {3, 5, 5, 6, 5, 5, 5, 5, 5, 5, 5, 5, 4, 5, 5, 6, 6, 5}}, 
 {14, {3, 5, 5, 6, 5, 5, 5, 5, 4, 5, 5, 5, 5, 5, 6, 6, 5, 5}}, 
 {14, {3, 5, 5, 6, 5, 5, 5, 4, 5, 6, 5, 5, 5, 5, 5, 5, 6, 5}}, 
 {14, {3, 5, 5, 5, 6, 6, 5, 5, 5, 5, 5, 5, 5, 4, 5, 6, 5, 5}}, 
 {14, {3, 5, 5, 5, 6, 6, 5, 5, 4, 4, 5, 5, 5, 6, 6, 5, 5, 5}}, 
 {14, {3, 5, 5, 5, 6, 5, 5, 5, 5, 4, 5, 6, 5, 5, 5, 5, 6, 5}}, 
 {14, {3, 5, 5, 5, 6, 5, 5, 5, 4, 5, 5, 5, 5, 6, 5, 5, 6, 5}}, 
 {14, {3, 5, 5, 5, 6, 5, 4, 5, 5, 5, 5, 5, 5, 6, 5, 5, 6, 5}}, 
 {14, {3, 5, 5, 5, 6, 5, 4, 4, 5, 5, 6, 6, 5, 5, 5, 5, 6, 5}}, 
 {14, {3, 5, 5, 5, 5, 6, 5, 5, 5, 4, 5, 6, 5, 5, 5, 5, 6, 5}}, 
 {14, {3, 5, 5, 5, 5, 6, 5, 4, 4, 5, 5, 6, 6, 6, 5, 5, 5, 5}}, 
 {14, {3, 5, 5, 5, 5, 6, 5, 4, 4, 5, 5, 5, 5, 5, 6, 6, 6, 5}}, 
 {14, {3, 5, 5, 5, 5, 5, 6, 6, 6, 5, 5, 5, 5, 3, 5, 6, 5, 5}}, 
 {14, {3, 5, 5, 5, 5, 5, 6, 6, 6, 5, 5, 4, 4, 5, 5, 5, 6, 5}}, 
 {14, {3, 5, 5, 5, 5, 5, 6, 6, 5, 4, 5, 5, 5, 6, 5, 5, 5, 5}}, 
 {14, {3, 5, 5, 5, 5, 5, 6, 6, 5, 4, 4, 5, 5, 5, 5, 6, 6, 5}}, 
 {14, {3, 5, 5, 5, 5, 5, 5, 6, 6, 6, 5, 5, 4, 4, 5, 5, 6, 5}}, 
 {14, {3, 5, 5, 5, 5, 5, 5, 6, 5, 5, 5, 5, 4, 5, 6, 6, 5, 5}}, 
 {14, {3, 5, 5, 5, 5, 5, 5, 6, 5, 5, 4, 5, 6, 6, 5, 5, 5, 5}}, 
 {14, {3, 5, 5, 5, 5, 5, 5, 5, 6, 5, 4, 5, 6, 6, 5, 5, 5, 5}}, 
 {14, {3, 5, 5, 5, 5, 5, 5, 5, 6, 5, 4, 5, 5, 6, 5, 5, 6, 5}}, 
 {14, {3, 5, 5, 5, 5, 5, 5, 5, 5, 6, 5, 4, 5, 6, 6, 5, 5, 5}}, 
 {14, {3, 5, 5, 5, 5, 5, 5, 4, 5, 6, 6, 5, 5, 5, 5, 6, 5, 5}}, 
 {14, {3, 5, 5, 5, 5, 5, 5, 4, 4, 5, 5, 6, 6, 6, 5, 5, 6, 5}}, 
 {14, {3, 5, 5, 5, 5, 5, 5, 3, 5, 5, 6, 6, 6, 5, 5, 5, 6, 5}}, 
 {14, {3, 5, 5, 5, 5, 5, 4, 4, 5, 5, 6, 6, 5, 5, 5, 6, 6, 5}}, 
 {14, {3, 5, 5, 5, 5, 5, 4, 4, 5, 5, 5, 6, 6, 6, 5, 5, 6, 5}}, 
 {14, {3, 5, 5, 5, 5, 4, 4, 5, 5, 6, 6, 5, 5, 5, 6, 6, 5, 5}}, 
 {14, {3, 5, 5, 5, 5, 4, 4, 5, 5, 5, 6, 6, 6, 5, 5, 5, 6, 5}}, 
 {14, {3, 5, 5, 5, 4, 5, 5, 6, 6, 6, 6, 5, 4, 5, 5, 5, 5, 5}}, 
 {14, {3, 5, 5, 5, 4, 5, 5, 6, 5, 4, 5, 5, 5, 5, 6, 6, 6, 5}}, 
 {14, {3, 5, 5, 5, 4, 5, 5, 5, 6, 5, 5, 5, 6, 5, 5, 5, 6, 5}}, 
 {14, {3, 5, 5, 5, 4, 5, 5, 5, 5, 6, 5, 5, 6, 6, 5, 5, 5, 5}}, 
 {14, {3, 5, 5, 5, 4, 5, 5, 5, 5, 5, 6, 5, 5, 6, 6, 5, 5, 5}}, 
 {14, {3, 5, 5, 5, 4, 5, 5, 5, 5, 5, 5, 6, 6, 5, 5, 6, 5, 5}}, 
 {14, {3, 5, 5, 5, 4, 5, 5, 4, 5, 5, 5, 6, 6, 6, 6, 5, 5, 5}}, 
 {14, {3, 5, 5, 5, 4, 5, 5, 4, 5, 5, 5, 5, 6, 6, 6, 6, 5, 5}}, 
 {14, {3, 5, 5, 5, 4, 4, 5, 5, 6, 6, 5, 5, 6, 6, 5, 5, 5, 5}}, 
 {14, {3, 5, 5, 5, 4, 4, 5, 5, 5, 5, 5, 6, 6, 5, 6, 6, 5, 5}}, 
 {14, {3, 5, 5, 4, 5, 6, 6, 5, 5, 5, 5, 5, 5, 6, 5, 5, 5, 5}}, 
 {14, {3, 5, 5, 4, 5, 5, 6, 6, 6, 5, 5, 5, 5, 4, 5, 5, 6, 5}}, 
 {14, {3, 5, 5, 4, 5, 5, 6, 6, 5, 5, 4, 5, 5, 5, 6, 6, 5, 5}}, 
 {14, {3, 5, 5, 4, 5, 5, 6, 5, 5, 5, 5, 5, 6, 5, 5, 5, 6, 5}}, 
 {14, {3, 5, 5, 4, 5, 5, 5, 5, 6, 6, 5, 5, 5, 6, 5, 5, 5, 5}}, 
 {14, {3, 5, 5, 4, 5, 5, 5, 5, 5, 6, 5, 5, 5, 6, 6, 5, 5, 5}}, 
 {14, {3, 5, 5, 4, 4, 5, 6, 5, 5, 5, 5, 5, 5, 6, 6, 6, 5, 5}}, 
 {14, {3, 5, 5, 4, 4, 5, 5, 5, 5, 6, 6, 5, 5, 5, 5, 6, 6, 5}}, 
 {14, {1, 6, 6, 6, 5, 4, 5, 5, 5, 5, 5, 4, 4, 5, 6, 6, 6, 6}}, 
 {14, {1, 6, 6, 6, 1, 6, 6, 6, 5, 5, 5, 4, 5, 5, 5, 6, 6, 6}}, 
 {14, {1, 6, 6, 6, 1, 6, 6, 6, 5, 4, 4, 5, 5, 5, 6, 6, 6, 6}}, 
 {14, {1, 6, 6, 5, 5, 6, 5, 5, 5, 4, 4, 5, 5, 5, 5, 6, 6, 6}}, 
 {14, {1, 6, 6, 5, 5, 5, 6, 5, 4, 5, 5, 4, 5, 5, 5, 6, 6, 6}}, 
 {14, {1, 6, 6, 5, 5, 5, 5, 6, 5, 5, 5, 5, 6, 6, 1, 6, 6, 6}}, 
 {14, {1, 6, 6, 5, 5, 5, 5, 6, 5, 4, 4, 5, 6, 5, 5, 5, 6, 6}}, 
 {14, {1, 6, 6, 5, 5, 5, 5, 5, 4, 5, 6, 6, 6, 6, 1, 6, 6, 6}}, 
 {14, {1, 6, 6, 5, 5, 5, 5, 4, 5, 6, 5, 5, 5, 4, 5, 6, 6, 6}}, 
 {14, {1, 6, 6, 5, 5, 4, 5, 5, 5, 5, 6, 6, 1, 6, 6, 6, 6, 6}}, 
 {14, {1, 6, 6, 5, 5, 4, 5, 5, 4, 5, 5, 5, 6, 5, 5, 6, 6, 6}}, 
 {14, {1, 6, 6, 5, 5, 3, 5, 5, 5, 5, 6, 5, 5, 5, 5, 6, 6, 6}}, 
 {14, {1, 6, 6, 5, 4, 5, 5, 6, 6, 5, 4, 5, 5, 5, 5, 5, 6, 6}}, 
 {14, {1, 6, 6, 5, 4, 5, 5, 5, 5, 5, 6, 6, 6, 1, 6, 6, 6, 6}}, 
 {14, {1, 6, 6, 5, 4, 5, 5, 5, 5, 4, 5, 6, 5, 5, 5, 6, 6, 6}}, 
 {14, {1, 6, 6, 5, 4, 5, 5, 5, 5, 4, 5, 5, 6, 6, 5, 5, 6, 6}}, 
 {14, {1, 6, 6, 5, 4, 5, 5, 4, 5, 5, 6, 6, 5, 5, 5, 5, 6, 6}}, 
 {14, {1, 6, 6, 5, 4, 4, 5, 5, 5, 6, 5, 4, 5, 5, 6, 6, 6, 6}}, 
 {14, {1, 6, 6, 5, 3, 5, 6, 6, 5, 5, 5, 5, 5, 5, 5, 5, 6, 6}}, 
 {14, {1, 6, 6, 5, 3, 5, 5, 6, 5, 5, 5, 5, 5, 5, 5, 6, 6, 6}}, 
 {14, {1, 6, 6, 5, 3, 5, 5, 5, 5, 5, 5, 6, 5, 5, 5, 6, 6, 6}}, 
 {14, {1, 6, 6, 1, 6, 6, 5, 5, 5, 6, 6, 6, 6, 6, 1, 6, 6, 6}}, 
 {14, {1, 6, 6, 1, 6, 6, 5, 4, 5, 5, 6, 6, 5, 5, 5, 6, 6, 6}}, 
 {14, {1, 6, 1, 6, 6, 6, 5, 5, 5, 5, 4, 5, 5, 6, 6, 6, 6, 6}}, 
 {14, {4, 5, 6, 5, 5, 6, 6, 5, 5, 6, 5, 4, 5, 6, 6, 6, 5}}, 
 {14, {4, 5, 6, 5, 4, 5, 6, 5, 5, 6, 6, 5, 5, 6, 6, 6, 5}}, 
 {14, {4, 5, 6, 5, 4, 5, 6, 5, 5, 6, 5, 5, 6, 6, 6, 6, 5}}, 
 {14, {4, 5, 5, 6, 6, 5, 6, 6, 5, 5, 6, 6, 5, 4, 5, 6, 5}}, 
 {14, {4, 5, 5, 6, 5, 5, 6, 5, 4, 5, 6, 6, 5, 6, 6, 6, 5}}, 
 {14, {4, 5, 5, 4, 5, 6, 6, 6, 6, 5, 4, 5, 6, 6, 6, 6, 5}}, 
 {14, {4, 5, 4, 5, 6, 5, 5, 5, 5, 5, 6, 6, 6, 6, 6, 6, 5}}, 
 {14, {4, 5, 4, 5, 5, 6, 5, 5, 6, 6, 5, 6, 6, 6, 6, 5, 5}}, 
 {14, {4, 5, 4, 5, 5, 5, 6, 6, 6, 6, 6, 5, 5, 5, 6, 6, 5}}, 
 {14, {4, 5, 4, 5, 5, 5, 5, 5, 6, 6, 6, 6, 6, 5, 6, 6, 5}}, 
 {14, {4, 4, 5, 6, 6, 6, 5, 5, 6, 5, 4, 5, 6, 6, 6, 6, 5}}, 
 {14, {4, 4, 5, 5, 6, 6, 6, 5, 4, 5, 6, 6, 5, 6, 6, 6, 5}}, 
 {14, {4, 4, 5, 5, 4, 5, 6, 6, 6, 6, 5, 6, 6, 5, 6, 6, 5}}, 
 {14, {4, 4, 5, 4, 5, 6, 6, 6, 5, 5, 6, 6, 6, 5, 6, 6, 5}}, 
 {14, {4, 4, 5, 4, 5, 6, 5, 5, 6, 6, 6, 6, 6, 5, 6, 6, 5}}, 
 {14, {4, 4, 5, 4, 5, 5, 6, 6, 5, 6, 6, 5, 6, 6, 6, 6, 5}}, 
 {14, {4, 4, 5, 4, 4, 5, 6, 6, 6, 5, 6, 6, 6, 6, 6, 6, 5}}, 
 {14, {3, 5, 6, 6, 5, 6, 6, 5, 4, 5, 5, 5, 6, 6, 6, 6, 5}}, 
 {14, {3, 5, 6, 6, 5, 5, 6, 5, 5, 4, 5, 6, 6, 6, 6, 6, 5}}, 
 {14, {3, 5, 5, 6, 5, 5, 6, 6, 5, 5, 5, 6, 6, 5, 6, 6, 5}}, 
 {14, {3, 5, 5, 6, 5, 5, 5, 6, 6, 6, 6, 6, 5, 5, 6, 5, 5}}, 
 {14, {3, 5, 5, 5, 6, 6, 6, 6, 6, 5, 5, 6, 5, 5, 6, 5, 5}}, 
 {14, {3, 5, 5, 5, 4, 5, 6, 6, 5, 6, 6, 5, 6, 6, 6, 6, 5}}, 
 {14, {3, 5, 5, 4, 5, 6, 6, 6, 6, 6, 5, 6, 6, 6, 5, 5, 5}}, 
 {14, {3, 5, 5, 4, 5, 6, 6, 6, 5, 6, 6, 6, 6, 5, 5, 6, 5}}, 
 {14, {3, 5, 5, 4, 5, 6, 6, 5, 6, 6, 6, 6, 6, 5, 5, 6, 5}}, 
 {14, {3, 5, 5, 4, 5, 6, 6, 5, 5, 6, 6, 6, 5, 6, 6, 6, 5}}, 
 {14, {1, 6, 6, 5, 6, 6, 6, 6, 6, 6, 5, 5, 3, 5, 6, 6, 6}}, 
 {14, {1, 6, 6, 5, 6, 6, 6, 6, 5, 3, 5, 5, 6, 6, 6, 6, 6}}, 
 {14, {1, 6, 6, 5, 6, 6, 6, 5, 4, 5, 6, 6, 5, 5, 6, 6, 6}}, 
 {14, {1, 6, 6, 5, 5, 6, 6, 6, 6, 5, 3, 5, 6, 6, 6, 6, 6}}, 
 {14, {1, 6, 6, 5, 5, 6, 6, 5, 6, 6, 6, 5, 4, 5, 6, 6, 6}}, 
 {14, {1, 6, 6, 5, 5, 6, 6, 5, 4, 5, 6, 6, 5, 6, 6, 6, 6}}, 
 {14, {1, 6, 6, 5, 5, 5, 3, 5, 6, 6, 6, 6, 6, 6, 6, 6, 6}}, 
 {14, {1, 6, 6, 5, 4, 5, 6, 6, 6, 6, 5, 6, 6, 5, 5, 6, 6}}, 
 {14, {1, 6, 6, 5, 4, 5, 6, 5, 5, 6, 6, 6, 6, 6, 5, 6, 6}}, 
 {14, {1, 6, 1, 6, 6, 6, 6, 5, 6, 6, 5, 6, 6, 6, 6, 6, 6}}, 
 {13, {4, 4, 5, 5, 5, 4, 5, 5, 4, 5, 5, 5, 5, 4, 5, 5, 5, 5, 5}}, 
 {13, {4, 4, 5, 5, 4, 5, 5, 5, 5, 5, 5, 4, 5, 5, 5, 4, 5, 5, 5}}, 
 {13, {4, 4, 5, 5, 4, 5, 5, 5, 5, 5, 4, 5, 5, 4, 5, 5, 5, 5, 5}}, 
 {13, {4, 4, 5, 5, 4, 5, 5, 5, 5, 4, 5, 5, 5, 5, 4, 5, 5, 5, 5}}, 
 {13, {4, 4, 5, 5, 4, 5, 5, 5, 5, 4, 5, 5, 4, 5, 5, 5, 5, 5, 5}}, 
 {13, {4, 4, 5, 4, 5, 5, 5, 5, 5, 5, 5, 5, 5, 5, 4, 4, 5, 5, 5}}, 
 {13, {4, 4, 5, 4, 5, 5, 5, 5, 5, 5, 4, 4, 5, 5, 5, 5, 5, 5, 5}}, 
 {13, {4, 4, 5, 4, 4, 5, 5, 5, 5, 4, 5, 5, 5, 5, 5, 5, 5, 5, 5}}, 
 {13, {4, 4, 5, 4, 4, 5, 5, 5, 4, 5, 5, 5, 5, 5, 5, 5, 5, 5, 5}}, 
 {13, {4, 5, 5, 5, 5, 6, 5, 4, 5, 5, 6, 5, 4, 5, 5, 5, 6, 5}}, 
 {13, {4, 5, 5, 5, 5, 5, 6, 5, 4, 5, 6, 5, 4, 5, 5, 5, 6, 5}}, 
 {13, {4, 5, 5, 5, 5, 5, 6, 5, 4, 5, 5, 5, 6, 5, 4, 5, 6, 5}}, 
 {13, {4, 5, 5, 5, 5, 5, 5, 5, 5, 4, 5, 6, 5, 4, 5, 6, 6, 5}}, 
 {13, {4, 5, 5, 5, 5, 5, 4, 5, 5, 5, 5, 6, 5, 4, 5, 6, 6, 5}}, 
 {13, {4, 5, 5, 5, 5, 4, 5, 6, 5, 5, 5, 4, 5, 6, 5, 5, 6, 5}}, 
 {13, {4, 5, 5, 5, 4, 5, 6, 5, 4, 5, 5, 5, 5, 5, 5, 6, 6, 5}}, 
 {13, {4, 5, 5, 5, 4, 5, 5, 6, 5, 4, 5, 5, 6, 5, 5, 6, 5, 5}}, 
 {13, {4, 5, 5, 4, 5, 6, 5, 4, 5, 5, 5, 5, 5, 5, 5, 6, 6, 5}}, 
 {13, {4, 5, 5, 4, 5, 5, 5, 5, 6, 5, 5, 5, 5, 4, 5, 6, 6, 5}}, 
 {13, {4, 5, 5, 4, 5, 5, 5, 5, 4, 5, 5, 5, 6, 5, 5, 6, 6, 5}}, 
 {13, {4, 5, 5, 4, 5, 5, 4, 5, 5, 4, 5, 5, 6, 6, 6, 6, 5, 5}}, 
 {13, {4, 5, 4, 5, 5, 5, 6, 5, 5, 4, 5, 5, 5, 6, 6, 5, 5, 5}}, 
 {13, {4, 5, 4, 5, 5, 5, 5, 5, 6, 5, 5, 5, 5, 6, 5, 5, 5, 5}}, 
 {13, {4, 5, 4, 5, 5, 5, 5, 5, 5, 5, 6, 5, 5, 5, 5, 5, 6, 5}}, 
 {13, {4, 5, 4, 5, 5, 5, 5, 5, 5, 5, 5, 6, 5, 5, 5, 6, 5, 5}}, 
 {13, {4, 5, 4, 5, 5, 5, 5, 5, 5, 5, 5, 5, 6, 5, 5, 6, 5, 5}}, 
 {13, {4, 5, 4, 5, 5, 5, 5, 5, 5, 5, 4, 5, 6, 6, 6, 5, 5, 5}}, 
 {13, {4, 4, 5, 6, 6, 5, 4, 5, 5, 5, 5, 4, 5, 5, 5, 6, 6, 5}}, 
 {13, {4, 4, 5, 6, 6, 5, 4, 5, 5, 5, 4, 5, 5, 5, 5, 6, 6, 5}}, 
 {13, {4, 4, 5, 6, 5, 5, 5, 4, 5, 5, 5, 5, 5, 6, 5, 5, 6, 5}}, 
 {13, {4, 4, 5, 6, 5, 5, 5, 4, 5, 5, 5, 5, 4, 5, 6, 6, 6, 5}}, 
 {13, {4, 4, 5, 5, 6, 6, 5, 5, 5, 5, 5, 6, 5, 4, 4, 5, 6, 5}}, 
 {13, {4, 4, 5, 5, 6, 5, 5, 6, 5, 5, 5, 5, 5, 4, 5, 5, 6, 5}}, 
 {13, {4, 4, 5, 5, 6, 5, 5, 6, 5, 5, 4, 5, 5, 5, 5, 5, 6, 5}}, 
 {13, {4, 4, 5, 5, 6, 5, 5, 5, 6, 5, 5, 4, 5, 5, 5, 5, 6, 5}}, 
 {13, {4, 4, 5, 5, 6, 5, 5, 4, 5, 5, 5, 6, 5, 5, 5, 6, 5, 5}}, 
 {13, {4, 4, 5, 5, 6, 5, 5, 4, 5, 5, 5, 6, 5, 5, 5, 5, 6, 5}}, 
 {13, {4, 4, 5, 5, 6, 5, 5, 4, 5, 5, 5, 4, 5, 5, 6, 6, 6, 5}}, 
 {13, {4, 4, 5, 5, 6, 5, 4, 5, 5, 4, 5, 5, 5, 6, 6, 6, 5, 5}}, 
 {13, {4, 4, 5, 5, 6, 5, 4, 4, 5, 6, 5, 5, 5, 5, 6, 6, 5, 5}}, 
 {13, {4, 4, 5, 5, 5, 6, 6, 5, 4, 4, 5, 6, 5, 5, 5, 5, 6, 5}}, 
 {13, {4, 4, 5, 5, 5, 6, 5, 5, 6, 5, 5, 5, 5, 4, 5, 6, 5, 5}}, 
 {13, {4, 4, 5, 5, 5, 6, 5, 5, 5, 5, 4, 5, 6, 5, 5, 5, 6, 5}}, 
 {13, {4, 4, 5, 5, 5, 6, 5, 5, 4, 5, 5, 6, 5, 5, 5, 6, 5, 5}}, 
 {13, {4, 4, 5, 5, 5, 6, 5, 4, 5, 5, 5, 6, 5, 5, 6, 5, 5, 5}}, 
 {13, {4, 4, 5, 5, 5, 5, 6, 6, 6, 5, 5, 4, 5, 5, 4, 5, 6, 5}}, 
 {13, {4, 4, 5, 5, 5, 5, 6, 6, 5, 4, 4, 5, 5, 5, 5, 6, 6, 5}}, 
 {13, {4, 4, 5, 5, 5, 5, 6, 5, 5, 5, 4, 4, 5, 6, 6, 6, 5, 5}}, 
 {13, {4, 4, 5, 5, 5, 5, 6, 5, 5, 4, 4, 5, 6, 6, 6, 5, 5, 5}}, 
 {13, {4, 4, 5, 5, 5, 5, 5, 6, 5, 5, 5, 4, 4, 5, 6, 6, 6, 5}}, 
 {13, {4, 4, 5, 5, 5, 5, 5, 4, 4, 5, 5, 5, 6, 6, 5, 6, 6, 5}}, 
 {13, {4, 4, 5, 5, 5, 5, 4, 5, 6, 6, 6, 6, 5, 4, 5, 5, 5, 5}}, 
 {13, {4, 4, 5, 5, 5, 5, 4, 5, 6, 6, 5, 5, 4, 5, 5, 6, 6, 5}}, 
 {13, {4, 4, 5, 5, 5, 5, 4, 5, 6, 5, 5, 6, 5, 5, 5, 6, 5, 5}}, 
 {13, {4, 4, 5, 5, 5, 5, 4, 5, 6, 5, 5, 5, 4, 5, 6, 6, 6, 5}}, 
 {13, {4, 4, 5, 5, 5, 5, 4, 5, 6, 5, 4, 5, 5, 6, 6, 6, 5, 5}}, 
 {13, {4, 4, 5, 5, 5, 4, 5, 6, 6, 5, 4, 5, 5, 6, 6, 5, 5, 5}}, 
 {13, {4, 4, 5, 5, 5, 4, 5, 6, 5, 5, 6, 5, 5, 5, 5, 6, 5, 5}}, 
 {13, {4, 4, 5, 5, 5, 4, 5, 6, 5, 5, 4, 5, 6, 6, 6, 5, 5, 5}}, 
 {13, {4, 4, 5, 5, 5, 4, 5, 5, 6, 6, 5, 5, 5, 4, 5, 6, 6, 5}}, 
 {13, {4, 4, 5, 5, 5, 4, 5, 5, 5, 6, 6, 5, 4, 5, 5, 6, 6, 5}}, 
 {13, {4, 4, 5, 5, 5, 4, 5, 5, 5, 6, 5, 4, 5, 6, 6, 6, 5, 5}}, 
 {13, {4, 4, 5, 5, 5, 4, 5, 5, 5, 5, 6, 5, 5, 6, 5, 5, 6, 5}}, 
 {13, {4, 4, 5, 5, 5, 4, 5, 5, 5, 4, 5, 6, 6, 5, 5, 6, 6, 5}}, 
 {13, {4, 4, 5, 5, 4, 5, 6, 6, 6, 5, 5, 4, 5, 5, 5, 5, 6, 5}}, 
 {13, {4, 4, 5, 5, 4, 5, 6, 5, 5, 6, 5, 5, 5, 6, 5, 5, 5, 5}}, 
 {13, {4, 4, 5, 5, 4, 5, 6, 5, 5, 5, 5, 4, 5, 5, 6, 6, 6, 5}}, 
 {13, {4, 4, 5, 5, 4, 5, 6, 5, 5, 4, 5, 5, 5, 6, 6, 6, 5, 5}}, 
 {13, {4, 4, 5, 5, 4, 5, 5, 6, 6, 6, 6, 6, 5, 4, 4, 5, 5, 5}}, 
 {13, {4, 4, 5, 5, 4, 5, 5, 6, 5, 4, 5, 5, 5, 5, 6, 6, 6, 5}}, 
 {13, {4, 4, 5, 5, 4, 5, 5, 5, 6, 6, 5, 4, 5, 5, 5, 6, 6, 5}}, 
 {13, {4, 4, 5, 5, 4, 5, 5, 5, 6, 5, 5, 6, 5, 5, 5, 6, 5, 5}}, 
 {13, {4, 4, 5, 5, 4, 5, 5, 5, 5, 6, 6, 6, 5, 4, 5, 5, 6, 5}}, 
 {13, {4, 4, 5, 5, 4, 5, 5, 5, 5, 4, 5, 6, 6, 5, 6, 6, 5, 5}}, 
 {13, {4, 4, 5, 5, 4, 5, 5, 5, 4, 5, 6, 6, 6, 5, 5, 5, 6, 5}}, 
 {13, {4, 4, 5, 5, 4, 5, 5, 5, 4, 5, 5, 6, 6, 5, 6, 6, 5, 5}}, 
 {13, {4, 4, 5, 5, 4, 5, 5, 5, 4, 5, 5, 6, 6, 5, 5, 6, 6, 5}}, 
 {13, {4, 4, 5, 4, 5, 6, 6, 6, 6, 5, 5, 5, 5, 5, 5, 4, 5, 5}}, 
 {13, {4, 4, 5, 4, 5, 6, 6, 6, 6, 5, 5, 4, 5, 5, 5, 5, 5, 5}}, 
 {13, {4, 4, 5, 4, 5, 6, 6, 5, 5, 6, 5, 5, 5, 5, 5, 5, 5, 5}}, 
 {13, {4, 4, 5, 4, 5, 6, 5, 5, 5, 5, 6, 5, 5, 5, 5, 5, 6, 5}}, 
 {13, {4, 4, 5, 4, 5, 5, 6, 6, 6, 5, 5, 5, 6, 5, 4, 5, 5, 5}}, 
 {13, {4, 4, 5, 4, 5, 5, 6, 6, 5, 4, 5, 5, 5, 5, 6, 6, 5, 5}}, 
 {13, {4, 4, 5, 4, 5, 5, 6, 5, 5, 5, 6, 6, 6, 5, 5, 4, 5, 5}}, 
 {13, {4, 4, 5, 4, 5, 5, 6, 5, 5, 5, 5, 6, 5, 5, 5, 5, 6, 5}}, 
 {13, {4, 4, 5, 4, 5, 5, 6, 5, 4, 5, 6, 6, 6, 5, 5, 5, 5, 5}}, 
 {13, {4, 4, 5, 4, 5, 5, 5, 6, 6, 6, 5, 5, 5, 4, 5, 6, 5, 5}}, 
 {13, {4, 4, 5, 4, 5, 5, 5, 6, 5, 4, 5, 5, 5, 6, 6, 6, 5, 5}}, 
 {13, {4, 4, 5, 4, 5, 5, 5, 5, 5, 5, 4, 5, 5, 6, 6, 6, 6, 5}}, 
 {13, {4, 4, 5, 4, 5, 5, 5, 5, 4, 4, 5, 6, 6, 6, 6, 6, 5, 5}}, 
 {13, {4, 4, 5, 4, 5, 5, 4, 5, 6, 6, 6, 6, 5, 5, 5, 5, 5, 5}}, 
 {13, {4, 4, 5, 4, 5, 5, 4, 5, 5, 5, 5, 6, 6, 6, 6, 5, 5, 5}}, 
 {13, {4, 4, 5, 4, 5, 5, 4, 5, 5, 5, 5, 5, 5, 6, 6, 6, 6, 5}}, 
 {13, {4, 4, 5, 4, 4, 5, 5, 6, 5, 5, 5, 5, 5, 5, 6, 6, 6, 5}}, 
 {13, {4, 4, 5, 4, 4, 5, 5, 5, 6, 6, 5, 5, 5, 5, 6, 6, 5, 5}}, 
 {13, {4, 4, 5, 4, 4, 5, 5, 5, 5, 6, 6, 5, 5, 6, 6, 5, 5, 5}}, 
 {13, {4, 4, 5, 4, 4, 5, 5, 5, 5, 6, 5, 5, 6, 6, 6, 5, 5, 5}}, 
 {13, {4, 4, 5, 4, 4, 5, 5, 5, 5, 5, 6, 6, 5, 6, 6, 5, 5, 5}}, 
 {13, {4, 4, 5, 4, 4, 5, 5, 5, 5, 5, 5, 6, 5, 5, 6, 6, 6, 5}}, 
 {13, {3, 5, 6, 5, 5, 5, 5, 5, 4, 5, 5, 4, 5, 5, 6, 6, 6, 5}}, 
 {13, {3, 5, 6, 5, 5, 5, 5, 5, 4, 4, 5, 5, 6, 6, 5, 5, 6, 5}}, 
 {13, {3, 5, 6, 5, 4, 5, 5, 5, 5, 5, 5, 5, 5, 6, 5, 5, 6, 5}}, 
 {13, {3, 5, 5, 6, 6, 6, 5, 5, 5, 4, 5, 5, 4, 5, 5, 5, 6, 5}}, 
 {13, {3, 5, 5, 6, 6, 5, 5, 5, 5, 6, 5, 4, 4, 5, 5, 5, 6, 5}}, 
 {13, {3, 5, 5, 6, 5, 5, 5, 5, 5, 6, 5, 5, 4, 5, 5, 5, 6, 5}}, 
 {13, {3, 5, 5, 6, 5, 5, 5, 5, 5, 5, 4, 5, 5, 5, 5, 6, 6, 5}}, 
 {13, {3, 5, 5, 5, 6, 6, 5, 5, 4, 4, 5, 5, 5, 5, 5, 6, 6, 5}}, 
 {13, {3, 5, 5, 5, 6, 5, 5, 5, 5, 5, 5, 6, 5, 4, 5, 5, 6, 5}}, 
 {13, {3, 5, 5, 5, 6, 5, 5, 5, 5, 4, 5, 5, 5, 6, 6, 5, 5, 5}}, 
 {13, {3, 5, 5, 5, 6, 5, 5, 5, 4, 5, 5, 6, 5, 5, 5, 5, 6, 5}}, 
 {13, {3, 5, 5, 5, 6, 5, 5, 5, 4, 4, 5, 5, 5, 6, 6, 6, 5, 5}}, 
 {13, {3, 5, 5, 5, 6, 5, 5, 5, 4, 4, 5, 5, 5, 5, 6, 6, 6, 5}}, 
 {13, {3, 5, 5, 5, 6, 5, 4, 5, 5, 6, 5, 5, 5, 5, 6, 5, 5, 5}}, 
 {13, {3, 5, 5, 5, 6, 5, 4, 4, 5, 5, 5, 5, 5, 5, 6, 6, 6, 5}}, 
 {13, {3, 5, 5, 5, 5, 6, 5, 5, 4, 5, 5, 5, 6, 5, 5, 5, 6, 5}}, 
 {13, {3, 5, 5, 5, 5, 6, 5, 5, 4, 5, 5, 5, 5, 6, 6, 5, 5, 5}}, 
 {13, {3, 5, 5, 5, 5, 5, 6, 6, 5, 5, 4, 5, 5, 5, 5, 6, 5, 5}}, 
 {13, {3, 5, 5, 5, 5, 5, 6, 5, 5, 5, 5, 5, 5, 4, 5, 6, 6, 5}}, 
 {13, {3, 5, 5, 5, 5, 5, 6, 5, 5, 5, 4, 4, 5, 6, 6, 6, 5, 5}}, 
 {13, {3, 5, 5, 5, 5, 5, 6, 5, 5, 4, 4, 5, 5, 5, 6, 6, 6, 5}}, 
 {13, {3, 5, 5, 5, 5, 5, 6, 5, 4, 5, 5, 6, 5, 5, 5, 5, 6, 5}}, 
 {13, {3, 5, 5, 5, 5, 5, 6, 5, 3, 5, 5, 5, 5, 6, 6, 6, 5, 5}}, 
 {13, {3, 5, 5, 5, 5, 5, 5, 6, 6, 5, 5, 5, 5, 4, 5, 6, 5, 5}}, 
 {13, {3, 5, 5, 5, 5, 5, 5, 6, 5, 4, 5, 6, 6, 5, 5, 5, 5, 5}}, 
 {13, {3, 5, 5, 5, 5, 5, 5, 6, 5, 4, 5, 6, 5, 5, 5, 5, 6, 5}}, 
 {13, {3, 5, 5, 5, 5, 5, 5, 5, 6, 5, 5, 4, 5, 6, 6, 5, 5, 5}}, 
 {13, {3, 5, 5, 5, 5, 5, 5, 5, 6, 5, 3, 5, 6, 6, 6, 5, 5, 5}}, 
 {13, {3, 5, 5, 5, 5, 5, 5, 5, 3, 5, 5, 5, 6, 6, 6, 6, 5, 5}}, 
 {13, {3, 5, 5, 5, 5, 5, 5, 4, 5, 5, 4, 5, 6, 6, 6, 6, 5, 5}}, 
 {13, {3, 5, 5, 5, 5, 5, 4, 5, 5, 6, 6, 5, 5, 5, 5, 6, 5, 5}}, 
 {13, {3, 5, 5, 5, 5, 5, 4, 5, 5, 6, 5, 5, 5, 6, 6, 5, 5, 5}}, 
 {13, {3, 5, 5, 5, 5, 5, 4, 5, 5, 5, 5, 6, 5, 5, 6, 6, 5, 5}}, 
 {13, {3, 5, 5, 5, 5, 5, 4, 4, 5, 6, 6, 6, 5, 5, 5, 5, 6, 5}}, 
 {13, {3, 5, 5, 5, 5, 5, 4, 4, 5, 6, 5, 5, 5, 6, 6, 6, 5, 5}}, 
 {13, {3, 5, 5, 5, 5, 4, 5, 5, 6, 6, 6, 6, 5, 4, 5, 5, 5, 5}}, 
 {13, {3, 5, 5, 5, 5, 4, 4, 5, 5, 5, 5, 6, 6, 6, 5, 5, 6, 5}}, 
 {13, {3, 5, 5, 5, 4, 5, 6, 5, 5, 5, 5, 5, 6, 5, 5, 5, 6, 5}}, 
 {13, {3, 5, 5, 5, 4, 5, 5, 5, 6, 5, 5, 6, 5, 5, 5, 5, 6, 5}}, 
 {13, {3, 5, 5, 5, 4, 4, 5, 5, 5, 6, 6, 5, 5, 5, 6, 6, 5, 5}}, 
 {13, {3, 5, 5, 5, 4, 4, 5, 5, 5, 5, 6, 5, 5, 6, 6, 6, 5, 5}}, 
 {13, {3, 5, 5, 4, 5, 5, 6, 5, 5, 6, 5, 5, 5, 5, 5, 5, 6, 5}}, 
 {13, {3, 5, 5, 4, 5, 5, 5, 5, 4, 5, 6, 5, 5, 6, 6, 6, 5, 5}}, 
 {13, {3, 5, 5, 4, 5, 5, 5, 4, 5, 6, 6, 6, 6, 5, 5, 5, 5, 5}}, 
 {13, {3, 5, 5, 4, 5, 5, 5, 4, 5, 5, 5, 6, 6, 6, 6, 5, 5, 5}}, 
 {13, {3, 5, 5, 4, 4, 5, 6, 6, 6, 5, 5, 5, 5, 5, 5, 5, 6, 5}}, 
 {13, {3, 5, 5, 4, 4, 5, 6, 6, 5, 5, 5, 5, 5, 5, 6, 6, 5, 5}}, 
 {13, {3, 5, 5, 4, 4, 5, 6, 5, 5, 5, 6, 6, 6, 5, 5, 5, 5, 5}}, 
 {13, {3, 5, 5, 4, 4, 5, 5, 5, 6, 6, 5, 5, 5, 5, 5, 6, 6, 5}}, 
 {13, {3, 5, 5, 4, 4, 5, 5, 5, 5, 5, 6, 6, 5, 5, 6, 6, 5, 5}}, 
 {13, {3, 5, 5, 3, 5, 5, 6, 6, 6, 5, 5, 5, 5, 5, 5, 5, 6, 5}}, 
 {13, {1, 6, 6, 6, 5, 5, 5, 4, 5, 5, 5, 5, 3, 5, 6, 6, 6, 6}}, 
 {13, {1, 6, 6, 6, 5, 3, 5, 5, 4, 5, 5, 5, 5, 5, 6, 6, 6, 6}}, 
 {13, {1, 6, 6, 6, 1, 6, 6, 6, 5, 5, 4, 4, 5, 5, 6, 6, 6, 6}}, 
 {13, {1, 6, 6, 5, 5, 6, 5, 4, 5, 5, 5, 5, 4, 5, 5, 6, 6, 6}}, 
 {13, {1, 6, 6, 5, 5, 5, 6, 6, 6, 1, 6, 6, 6, 1, 6, 6, 6, 6}}, 
 {13, {1, 6, 6, 5, 5, 5, 6, 5, 5, 5, 5, 5, 4, 4, 5, 6, 6, 6}}, 
 {13, {1, 6, 6, 5, 5, 5, 5, 5, 5, 5, 5, 3, 5, 6, 6, 5, 6, 6}}, 
 {13, {1, 6, 6, 5, 5, 5, 5, 5, 5, 5, 3, 5, 6, 5, 5, 6, 6, 6}}, 
 {13, {1, 6, 6, 5, 5, 5, 5, 5, 5, 5, 3, 5, 5, 6, 6, 5, 6, 6}}, 
 {13, {1, 6, 6, 5, 5, 5, 5, 5, 4, 5, 6, 6, 6, 1, 6, 6, 6, 6}}, 
 {13, {1, 6, 6, 5, 5, 5, 5, 5, 4, 5, 6, 5, 5, 4, 5, 6, 6, 6}}, 
 {13, {1, 6, 6, 5, 5, 5, 5, 4, 4, 5, 5, 6, 5, 5, 5, 6, 6, 6}}, 
 {13, {1, 6, 6, 5, 5, 5, 4, 5, 5, 4, 5, 5, 6, 6, 5, 5, 6, 6}}, 
 {13, {1, 6, 6, 5, 5, 5, 4, 4, 5, 5, 5, 6, 5, 5, 5, 6, 6, 6}}, 
 {13, {1, 6, 6, 5, 5, 5, 4, 4, 5, 5, 5, 5, 6, 5, 5, 6, 6, 6}}, 
 {13, {1, 6, 6, 5, 5, 4, 5, 6, 5, 5, 5, 4, 4, 5, 6, 6, 6, 6}}, 
 {13, {1, 6, 6, 5, 4, 5, 6, 5, 5, 5, 4, 5, 5, 5, 5, 6, 6, 6}}, 
 {13, {1, 6, 6, 5, 4, 5, 5, 5, 5, 5, 6, 5, 5, 4, 5, 6, 6, 6}}, 
 {13, {1, 6, 6, 5, 4, 5, 5, 5, 5, 5, 5, 6, 6, 5, 4, 5, 6, 6}}, 
 {13, {1, 6, 6, 5, 4, 5, 5, 5, 5, 4, 4, 5, 5, 6, 6, 6, 6, 6}}, 
 {13, {1, 6, 6, 5, 4, 5, 5, 4, 5, 5, 6, 5, 5, 5, 5, 6, 6, 6}}, 
 {13, {1, 6, 6, 5, 3, 5, 5, 5, 5, 5, 6, 6, 5, 5, 5, 5, 6, 6}}, 
 {13, {1, 6, 6, 5, 3, 5, 5, 5, 5, 5, 5, 5, 6, 6, 5, 5, 6, 6}}, 
 {13, {1, 6, 6, 5, 3, 5, 5, 5, 5, 5, 5, 5, 5, 6, 6, 5, 6, 6}}, 
 {13, {1, 6, 6, 1, 6, 6, 6, 5, 4, 4, 5, 5, 5, 6, 6, 6, 6, 6}}, 
 {13, {1, 6, 6, 1, 6, 6, 5, 6, 6, 5, 5, 4, 5, 5, 5, 6, 6, 6}}, 
 {13, {1, 6, 6, 1, 6, 6, 5, 5, 5, 6, 5, 4, 5, 5, 6, 6, 6, 6}}, 
 {13, {1, 6, 6, 1, 6, 6, 5, 5, 4, 4, 5, 5, 6, 6, 6, 6, 6, 6}}, 
 {13, {4, 5, 6, 5, 5, 6, 5, 5, 6, 6, 5, 5, 6, 5, 5, 6, 5}}, 
 {13, {4, 5, 6, 5, 4, 5, 6, 6, 5, 5, 6, 5, 5, 6, 6, 6, 5}}, 
 {13, {4, 5, 6, 5, 4, 5, 6, 6, 5, 5, 5, 6, 6, 5, 6, 6, 5}}, 
 {13, {4, 5, 6, 5, 4, 5, 6, 5, 5, 6, 6, 6, 5, 5, 6, 6, 5}}, 
 {13, {4, 5, 5, 5, 6, 6, 5, 6, 6, 5, 6, 6, 5, 4, 5, 6, 5}}, 
 {13, {4, 5, 4, 5, 5, 6, 5, 5, 6, 6, 6, 6, 5, 6, 6, 5, 5}}, 
 {13, {4, 5, 4, 5, 5, 5, 6, 6, 5, 5, 6, 6, 5, 6, 6, 6, 5}}, 
 {13, {4, 5, 4, 5, 5, 5, 5, 6, 6, 6, 6, 6, 5, 5, 6, 6, 5}}, 
 {13, {4, 4, 5, 6, 6, 5, 4, 5, 6, 5, 5, 6, 6, 6, 6, 6, 5}}, 
 {13, {4, 4, 5, 4, 5, 6, 6, 6, 6, 6, 5, 6, 6, 5, 5, 6, 5}}, 
 {13, {4, 4, 5, 4, 5, 5, 6, 6, 6, 6, 5, 6, 6, 5, 6, 6, 5}}, 
 {13, {3, 5, 6, 6, 5, 5, 5, 4, 5, 6, 6, 6, 6, 5, 6, 6, 5}}, 
 {13, {3, 5, 6, 6, 5, 4, 5, 5, 5, 6, 6, 6, 6, 5, 6, 6, 5}}, 
 {13, {3, 5, 6, 5, 5, 6, 5, 5, 6, 5, 5, 6, 6, 5, 6, 6, 5}}, 
 {13, {3, 5, 6, 5, 4, 5, 5, 5, 6, 6, 6, 6, 6, 5, 6, 6, 5}}, 
 {13, {3, 5, 5, 6, 6, 6, 6, 6, 5, 5, 6, 5, 4, 5, 6, 6, 5}}, 
 {13, {3, 5, 5, 6, 6, 6, 6, 6, 5, 4, 5, 6, 5, 5, 6, 6, 5}}, 
 {13, {3, 5, 5, 6, 6, 6, 6, 6, 5, 4, 4, 5, 6, 6, 6, 6, 5}}, 
 {13, {3, 5, 5, 6, 6, 6, 6, 5, 4, 4, 5, 6, 6, 6, 6, 6, 5}}, 
 {13, {3, 5, 5, 6, 6, 6, 6, 5, 3, 5, 5, 6, 6, 6, 6, 6, 5}}, 
 {13, {3, 5, 5, 6, 6, 6, 5, 4, 5, 6, 6, 5, 6, 6, 6, 5, 5}}, 
 {13, {3, 5, 5, 6, 6, 5, 5, 6, 6, 6, 6, 5, 4, 5, 6, 6, 5}}, 
 {13, {3, 5, 5, 6, 5, 5, 6, 6, 5, 6, 6, 5, 5, 6, 6, 5, 5}}, 
 {13, {3, 5, 5, 6, 5, 5, 6, 6, 5, 5, 6, 5, 5, 6, 6, 6, 5}}, 
 {13, {3, 5, 5, 6, 5, 5, 6, 5, 5, 5, 6, 6, 6, 5, 6, 6, 5}}, 
 {13, {3, 5, 5, 6, 5, 5, 5, 6, 6, 5, 6, 6, 5, 5, 6, 6, 5}}, 
 {13, {3, 5, 5, 6, 5, 4, 5, 6, 6, 6, 6, 6, 6, 5, 5, 6, 5}}, 
 {13, {3, 5, 5, 6, 5, 4, 5, 5, 6, 6, 5, 6, 6, 6, 6, 6, 5}}, 
 {13, {3, 5, 5, 5, 6, 6, 5, 5, 6, 5, 5, 6, 6, 5, 6, 6, 5}}, 
 {13, {3, 5, 5, 5, 6, 5, 5, 6, 5, 5, 6, 6, 6, 5, 6, 6, 5}}, 
 {13, {3, 5, 5, 5, 5, 6, 6, 5, 6, 6, 5, 6, 6, 5, 5, 6, 5}}, 
 {13, {3, 5, 5, 4, 5, 5, 6, 6, 6, 5, 6, 6, 5, 6, 6, 6, 5}}, 
 {13, {3, 5, 5, 4, 5, 5, 6, 6, 5, 6, 6, 5, 6, 6, 6, 6, 5}}, 
 {13, {3, 5, 5, 4, 4, 5, 6, 6, 6, 5, 6, 6, 6, 6, 6, 6, 5}}, 
 {13, {3, 5, 5, 3, 5, 5, 6, 6, 6, 6, 6, 6, 6, 6, 6, 5, 5}}, 
 {13, {3, 5, 4, 4, 5, 5, 5, 6, 6, 6, 6, 6, 6, 6, 6, 6, 5}}, 
 {13, {1, 6, 6, 6, 6, 5, 6, 6, 5, 5, 3, 5, 6, 6, 6, 6, 6}}, 
 {13, {1, 6, 6, 6, 1, 6, 6, 6, 5, 6, 6, 6, 6, 5, 6, 6, 6}}, 
 {13, {1, 6, 6, 5, 6, 6, 6, 6, 6, 5, 3, 5, 5, 6, 6, 6, 6}}, 
 {13, {1, 6, 6, 5, 6, 6, 6, 5, 6, 6, 1, 6, 6, 6, 6, 6, 6}}, 
 {13, {1, 6, 6, 5, 5, 6, 6, 5, 4, 5, 6, 6, 6, 6, 5, 6, 6}}, 
 {12, {4, 4, 5, 5, 5, 4, 5, 5, 5, 4, 4, 5, 5, 5, 5, 5, 5, 5, 5}}, 
 {12, {4, 4, 5, 5, 4, 5, 5, 5, 5, 5, 5, 5, 5, 5, 4, 4, 5, 5, 5}}, 
 {12, {4, 4, 5, 5, 4, 4, 5, 5, 5, 5, 5, 5, 4, 5, 5, 5, 5, 5, 5}}, 
 {12, {4, 4, 5, 4, 4, 5, 5, 4, 5, 5, 5, 5, 5, 5, 5, 5, 5, 5, 5}}, 
 {12, {3, 5, 5, 5, 5, 5, 5, 4, 4, 5, 5, 5, 4, 5, 5, 5, 5, 5, 5}}, 
 {12, {3, 5, 5, 5, 5, 5, 4, 4, 5, 5, 5, 4, 5, 5, 5, 5, 5, 5, 5}}, 
 {12, {3, 5, 5, 4, 4, 5, 5, 5, 5, 5, 5, 5, 4, 5, 5, 5, 5, 5, 5}}, 
 {12, {4, 5, 5, 5, 6, 5, 5, 4, 5, 5, 6, 5, 5, 4, 5, 6, 5, 5}}, 
 {12, {4, 5, 5, 5, 6, 5, 5, 4, 5, 5, 6, 5, 5, 4, 5, 5, 6, 5}}, 
 {12, {4, 5, 5, 5, 6, 5, 4, 5, 5, 5, 6, 5, 4, 5, 5, 6, 5, 5}}, 
 {12, {4, 5, 5, 5, 6, 5, 4, 5, 5, 5, 6, 5, 4, 5, 5, 5, 6, 5}}, 
 {12, {4, 5, 5, 5, 5, 5, 6, 5, 4, 5, 6, 5, 5, 4, 5, 5, 6, 5}}, 
 {12, {4, 5, 5, 5, 5, 4, 5, 6, 5, 4, 5, 5, 6, 5, 5, 5, 6, 5}}, 
 {12, {4, 5, 5, 5, 5, 4, 5, 5, 6, 5, 5, 5, 5, 4, 5, 6, 6, 5}}, 
 {12, {4, 5, 5, 4, 5, 6, 5, 5, 5, 4, 5, 6, 5, 5, 5, 5, 6, 5}}, 
 {12, {4, 5, 5, 4, 5, 5, 6, 5, 5, 5, 5, 4, 5, 5, 5, 6, 6, 5}}, 
 {12, {4, 5, 5, 4, 5, 5, 5, 6, 6, 5, 5, 5, 5, 5, 4, 5, 6, 5}}, 
 {12, {4, 5, 5, 4, 5, 5, 5, 6, 5, 4, 5, 6, 5, 5, 5, 5, 6, 5}}, 
 {12, {4, 5, 5, 4, 5, 5, 5, 5, 6, 5, 4, 5, 5, 5, 5, 6, 6, 5}}, 
 {12, {4, 5, 5, 4, 5, 5, 5, 5, 5, 6, 5, 5, 5, 4, 5, 6, 6, 5}}, 
 {12, {4, 5, 5, 4, 5, 5, 5, 5, 5, 4, 5, 6, 5, 5, 5, 6, 6, 5}}, 
 {12, {4, 5, 5, 4, 5, 5, 5, 5, 5, 4, 5, 5, 6, 5, 5, 6, 6, 5}}, 
 {12, {4, 5, 5, 4, 5, 5, 5, 5, 4, 5, 5, 6, 5, 5, 5, 6, 6, 5}}, 
 {12, {4, 5, 5, 4, 5, 5, 5, 4, 5, 5, 6, 5, 5, 5, 5, 6, 6, 5}}, 
 {12, {4, 5, 4, 5, 5, 6, 5, 5, 5, 5, 5, 5, 5, 5, 5, 5, 6, 5}}, 
 {12, {4, 5, 4, 5, 5, 5, 5, 5, 5, 6, 5, 5, 6, 5, 5, 5, 5, 5}}, 
 {12, {4, 5, 4, 5, 5, 5, 5, 5, 5, 5, 5, 6, 5, 5, 5, 5, 6, 5}}, 
 {12, {4, 5, 4, 5, 5, 5, 4, 5, 5, 6, 5, 5, 5, 5, 6, 6, 5, 5}}, 
 {12, {4, 5, 4, 5, 5, 5, 4, 5, 5, 5, 5, 6, 6, 6, 5, 5, 5, 5}}, 
 {12, {4, 5, 4, 5, 5, 4, 5, 5, 5, 5, 5, 5, 5, 6, 6, 6, 5, 5}}, 
 {12, {4, 4, 5, 6, 5, 4, 5, 5, 6, 6, 5, 5, 4, 5, 5, 5, 6, 5}}, 
 {12, {4, 4, 5, 5, 6, 5, 5, 6, 5, 5, 5, 5, 4, 5, 5, 5, 6, 5}}, 
 {12, {4, 4, 5, 5, 6, 5, 4, 5, 5, 5, 5, 6, 5, 5, 5, 6, 5, 5}}, 
 {12, {4, 4, 5, 5, 6, 5, 4, 5, 5, 5, 5, 5, 6, 5, 5, 5, 6, 5}}, 
 {12, {4, 4, 5, 5, 5, 6, 6, 6, 5, 5, 5, 4, 4, 5, 5, 6, 5, 5}}, 
 {12, {4, 4, 5, 5, 5, 6, 6, 6, 5, 5, 4, 5, 5, 4, 5, 6, 5, 5}}, 
 {12, {4, 4, 5, 5, 5, 6, 6, 5, 5, 5, 6, 5, 4, 4, 5, 6, 5, 5}}, 
 {12, {4, 4, 5, 5, 5, 6, 5, 5, 6, 5, 5, 5, 4, 5, 5, 5, 6, 5}}, 
 {12, {4, 4, 5, 5, 5, 6, 5, 5, 5, 4, 4, 5, 5, 6, 6, 6, 5, 5}}, 
 {12, {4, 4, 5, 5, 5, 6, 5, 5, 4, 5, 5, 5, 6, 5, 5, 6, 5, 5}}, 
 {12, {4, 4, 5, 5, 5, 6, 5, 4, 5, 5, 5, 4, 5, 6, 6, 6, 5, 5}}, 
 {12, {4, 4, 5, 5, 5, 6, 5, 4, 4, 5, 6, 5, 5, 5, 5, 6, 6, 5}}, 
 {12, {4, 4, 5, 5, 5, 5, 6, 6, 6, 5, 5, 5, 5, 4, 4, 5, 6, 5}}, 
 {12, {4, 4, 5, 5, 5, 5, 6, 5, 5, 5, 6, 5, 5, 4, 5, 6, 5, 5}}, 
 {12, {4, 4, 5, 5, 5, 5, 6, 5, 5, 4, 5, 5, 6, 5, 5, 5, 6, 5}}, 
 {12, {4, 4, 5, 5, 5, 5, 6, 5, 4, 5, 6, 5, 5, 5, 6, 5, 5, 5}}, 
 {12, {4, 4, 5, 5, 5, 5, 6, 5, 4, 5, 5, 5, 5, 6, 5, 5, 6, 5}}, 
 {12, {4, 4, 5, 5, 5, 5, 6, 5, 4, 5, 5, 4, 5, 5, 6, 6, 6, 5}}, 
 {12, {4, 4, 5, 5, 5, 5, 5, 6, 6, 5, 5, 6, 5, 4, 4, 5, 6, 5}}, 
 {12, {4, 4, 5, 5, 5, 5, 5, 6, 6, 5, 4, 4, 5, 5, 5, 6, 6, 5}}, 
 {12, {4, 4, 5, 5, 5, 5, 5, 6, 5, 4, 5, 6, 5, 5, 6, 5, 5, 5}}, 
 {12, {4, 4, 5, 5, 5, 5, 5, 4, 5, 6, 6, 6, 5, 5, 4, 5, 6, 5}}, 
 {12, {4, 4, 5, 5, 5, 5, 5, 4, 5, 6, 5, 4, 5, 5, 6, 6, 6, 5}}, 
 {12, {4, 4, 5, 5, 5, 5, 4, 5, 6, 5, 5, 5, 6, 5, 5, 6, 5, 5}}, 
 {12, {4, 4, 5, 5, 5, 5, 4, 5, 5, 6, 6, 5, 5, 4, 5, 6, 6, 5}}, 
 {12, {4, 4, 5, 5, 5, 4, 5, 6, 6, 6, 5, 4, 5, 6, 5, 5, 5, 5}}, 
 {12, {4, 4, 5, 5, 5, 4, 5, 5, 6, 5, 4, 5, 6, 6, 6, 5, 5, 5}}, 
 {12, {4, 4, 5, 5, 5, 4, 4, 5, 5, 6, 6, 5, 5, 5, 5, 6, 6, 5}}, 
 {12, {4, 4, 5, 5, 4, 5, 6, 6, 6, 5, 5, 5, 4, 5, 6, 5, 5, 5}}, 
 {12, {4, 4, 5, 5, 4, 5, 6, 6, 5, 5, 5, 4, 5, 5, 6, 6, 5, 5}}, 
 {12, {4, 4, 5, 5, 4, 5, 6, 5, 5, 5, 6, 5, 5, 6, 5, 5, 5, 5}}, 
 {12, {4, 4, 5, 5, 4, 5, 5, 6, 6, 5, 5, 5, 6, 6, 5, 4, 5, 5}}, 
 {12, {4, 4, 5, 5, 4, 5, 5, 6, 6, 5, 4, 5, 5, 5, 6, 6, 5, 5}}, 
 {12, {4, 4, 5, 5, 4, 5, 5, 6, 5, 5, 4, 5, 5, 6, 6, 6, 5, 5}}, 
 {12, {4, 4, 5, 5, 4, 5, 5, 5, 6, 6, 6, 5, 4, 5, 5, 6, 5, 5}}, 
 {12, {4, 4, 5, 5, 4, 5, 5, 5, 6, 6, 5, 6, 6, 5, 4, 5, 5, 5}}, 
 {12, {4, 4, 5, 5, 4, 5, 5, 5, 6, 5, 5, 5, 6, 5, 5, 5, 6, 5}}, 
 {12, {4, 4, 5, 5, 4, 5, 5, 5, 4, 5, 6, 5, 5, 5, 6, 6, 6, 5}}, 
 {12, {4, 4, 5, 5, 4, 5, 5, 4, 5, 5, 6, 6, 5, 6, 6, 5, 5, 5}}, 
 {12, {4, 4, 5, 5, 4, 5, 5, 4, 5, 5, 6, 6, 5, 5, 6, 6, 5, 5}}, 
 {12, {4, 4, 5, 5, 4, 5, 5, 4, 5, 5, 5, 6, 6, 6, 5, 5, 6, 5}}, 
 {12, {4, 4, 5, 5, 4, 5, 5, 4, 5, 5, 5, 6, 6, 5, 5, 6, 6, 5}}, 
 {12, {4, 4, 5, 5, 4, 4, 5, 5, 4, 5, 6, 6, 6, 6, 6, 5, 5, 5}}, 
 {12, {4, 4, 5, 4, 5, 6, 6, 6, 6, 5, 5, 5, 5, 5, 4, 5, 5, 5}}, 
 {12, {4, 4, 5, 4, 5, 6, 6, 6, 6, 5, 5, 5, 5, 4, 5, 5, 5, 5}}, 
 {12, {4, 4, 5, 4, 5, 6, 6, 5, 5, 5, 5, 5, 5, 5, 5, 6, 5, 5}}, 
 {12, {4, 4, 5, 4, 5, 6, 5, 5, 5, 5, 5, 6, 5, 5, 5, 5, 6, 5}}, 
 {12, {4, 4, 5, 4, 5, 5, 6, 6, 6, 5, 5, 6, 5, 5, 5, 4, 5, 5}}, 
 {12, {4, 4, 5, 4, 5, 5, 6, 6, 5, 5, 5, 4, 5, 5, 6, 6, 5, 5}}, 
 {12, {4, 4, 5, 4, 5, 5, 6, 5, 5, 5, 6, 5, 5, 5, 6, 5, 5, 5}}, 
 {12, {4, 4, 5, 4, 5, 5, 6, 5, 5, 5, 6, 5, 5, 5, 5, 6, 5, 5}}, 
 {12, {4, 4, 5, 4, 5, 5, 6, 5, 5, 5, 6, 5, 5, 5, 5, 5, 6, 5}}, 
 {12, {4, 4, 5, 4, 5, 5, 6, 5, 5, 5, 5, 5, 6, 5, 5, 5, 6, 5}}, 
 {12, {4, 4, 5, 4, 5, 5, 5, 6, 6, 6, 5, 5, 5, 6, 5, 4, 5, 5}}, 
 {12, {4, 4, 5, 4, 5, 5, 5, 6, 6, 6, 5, 5, 5, 4, 5, 5, 6, 5}}, 
 {12, {4, 4, 5, 4, 5, 5, 5, 6, 6, 5, 5, 4, 5, 5, 5, 6, 6, 5}}, 
 {12, {4, 4, 5, 4, 5, 5, 5, 6, 5, 5, 6, 5, 5, 5, 5, 5, 6, 5}}, 
 {12, {4, 4, 5, 4, 5, 5, 5, 5, 6, 6, 6, 5, 5, 4, 5, 5, 6, 5}}, 
 {12, {4, 4, 5, 4, 5, 5, 5, 5, 5, 5, 6, 6, 6, 6, 5, 5, 4, 5}}, 
 {12, {4, 4, 5, 4, 5, 5, 5, 5, 5, 5, 6, 5, 5, 6, 5, 5, 6, 5}}, 
 {12, {4, 4, 5, 4, 5, 5, 5, 5, 5, 4, 5, 6, 6, 6, 6, 5, 5, 5}}, 
 {12, {4, 4, 5, 4, 5, 5, 5, 4, 5, 6, 6, 6, 6, 5, 5, 5, 5, 5}}, 
 {12, {4, 4, 5, 4, 4, 5, 5, 5, 5, 6, 6, 6, 5, 5, 5, 6, 5, 5}}, 
 {12, {4, 4, 5, 4, 4, 5, 5, 5, 4, 5, 6, 6, 6, 6, 6, 5, 5, 5}}, 
 {12, {4, 4, 5, 4, 4, 5, 5, 4, 5, 5, 6, 6, 5, 6, 6, 6, 5, 5}}, 
 {12, {3, 5, 6, 6, 5, 4, 4, 5, 5, 5, 5, 5, 5, 5, 5, 6, 6, 5}}, 
 {12, {3, 5, 6, 5, 5, 5, 4, 5, 5, 4, 5, 5, 5, 5, 6, 6, 6, 5}}, 
 {12, {3, 5, 6, 5, 5, 4, 5, 5, 6, 6, 5, 5, 4, 5, 5, 5, 6, 5}}, 
 {12, {3, 5, 6, 5, 5, 4, 4, 5, 6, 5, 5, 5, 5, 5, 5, 6, 6, 5}}, 
 {12, {3, 5, 6, 5, 5, 4, 4, 5, 5, 5, 5, 5, 6, 6, 5, 5, 6, 5}}, 
 {12, {3, 5, 6, 5, 4, 5, 5, 5, 5, 5, 5, 5, 6, 5, 5, 5, 6, 5}}, 
 {12, {3, 5, 5, 6, 6, 6, 5, 4, 5, 5, 5, 5, 4, 5, 5, 5, 6, 5}}, 
 {12, {3, 5, 5, 6, 6, 5, 5, 5, 5, 5, 5, 6, 5, 4, 4, 5, 6, 5}}, 
 {12, {3, 5, 5, 6, 6, 5, 5, 4, 5, 5, 5, 5, 4, 5, 5, 6, 6, 5}}, 
 {12, {3, 5, 5, 6, 6, 5, 4, 5, 5, 5, 4, 5, 5, 5, 6, 6, 5, 5}}, 
 {12, {3, 5, 5, 6, 5, 5, 5, 5, 6, 5, 5, 5, 4, 5, 5, 5, 6, 5}}, 
 {12, {3, 5, 5, 6, 5, 5, 5, 5, 5, 6, 5, 5, 5, 4, 5, 5, 6, 5}}, 
 {12, {3, 5, 5, 6, 5, 4, 5, 5, 5, 5, 5, 5, 5, 5, 6, 6, 5, 5}}, 
 {12, {3, 5, 5, 5, 6, 6, 5, 5, 5, 5, 4, 5, 5, 5, 5, 6, 5, 5}}, 
 {12, {3, 5, 5, 5, 6, 6, 5, 5, 5, 4, 4, 5, 5, 5, 6, 6, 5, 5}}, 
 {12, {3, 5, 5, 5, 6, 5, 5, 5, 4, 5, 5, 5, 6, 5, 5, 5, 6, 5}}, 
 {12, {3, 5, 5, 5, 6, 5, 5, 5, 4, 5, 5, 5, 4, 5, 6, 6, 6, 5}}, 
 {12, {3, 5, 5, 5, 5, 6, 6, 6, 5, 5, 5, 5, 3, 5, 5, 5, 6, 5}}, 
 {12, {3, 5, 5, 5, 5, 6, 6, 5, 5, 5, 5, 4, 5, 5, 5, 6, 5, 5}}, 
 {12, {3, 5, 5, 5, 5, 6, 6, 5, 5, 4, 5, 5, 4, 5, 5, 6, 6, 5}}, 
 {12, {3, 5, 5, 5, 5, 6, 5, 5, 6, 5, 5, 5, 5, 4, 5, 5, 6, 5}}, 
 {12, {3, 5, 5, 5, 5, 6, 5, 5, 5, 6, 5, 5, 5, 4, 5, 5, 6, 5}}, 
 {12, {3, 5, 5, 5, 5, 6, 5, 5, 5, 5, 6, 5, 4, 5, 5, 5, 6, 5}}, 
 {12, {3, 5, 5, 5, 5, 6, 5, 5, 5, 5, 5, 4, 4, 5, 6, 6, 6, 5}}, 
 {12, {3, 5, 5, 5, 5, 6, 5, 5, 4, 5, 5, 5, 6, 6, 5, 5, 5, 5}}, 
 {12, {3, 5, 5, 5, 5, 6, 5, 4, 5, 5, 6, 5, 5, 5, 5, 5, 6, 5}}, 
 {12, {3, 5, 5, 5, 5, 5, 6, 6, 6, 5, 5, 5, 4, 4, 5, 5, 6, 5}}, 
 {12, {3, 5, 5, 5, 5, 5, 6, 6, 5, 5, 5, 4, 5, 5, 5, 6, 5, 5}}, 
 {12, {3, 5, 5, 5, 5, 5, 6, 6, 5, 5, 4, 4, 5, 5, 5, 6, 6, 5}}, 
 {12, {3, 5, 5, 5, 5, 5, 6, 6, 5, 4, 4, 5, 6, 5, 5, 5, 6, 5}}, 
 {12, {3, 5, 5, 5, 5, 5, 6, 5, 5, 5, 5, 6, 5, 5, 4, 5, 6, 5}}, 
 {12, {3, 5, 5, 5, 5, 5, 6, 5, 4, 5, 6, 5, 5, 5, 6, 5, 5, 5}}, 
 {12, {3, 5, 5, 5, 5, 5, 5, 6, 6, 5, 5, 4, 4, 5, 6, 6, 5, 5}}, 
 {12, {3, 5, 5, 5, 5, 5, 5, 6, 5, 5, 6, 5, 5, 4, 5, 5, 6, 5}}, 
 {12, {3, 5, 5, 5, 5, 5, 5, 6, 5, 4, 4, 5, 5, 5, 6, 6, 6, 5}}, 
 {12, {3, 5, 5, 5, 5, 5, 5, 6, 5, 3, 5, 6, 6, 6, 5, 5, 5, 5}}, 
 {12, {3, 5, 5, 5, 5, 5, 5, 5, 6, 6, 6, 5, 4, 4, 5, 5, 6, 5}}, 
 {12, {3, 5, 5, 5, 5, 5, 5, 5, 6, 6, 5, 4, 5, 5, 5, 6, 5, 5}}, 
 {12, {3, 5, 5, 5, 5, 5, 5, 5, 6, 5, 5, 4, 5, 6, 5, 5, 6, 5}}, 
 {12, {3, 5, 5, 5, 5, 5, 5, 5, 6, 5, 4, 5, 6, 5, 5, 5, 6, 5}}, 
 {12, {3, 5, 5, 5, 5, 5, 5, 5, 6, 5, 4, 4, 5, 6, 6, 6, 5, 5}}, 
 {12, {3, 5, 5, 5, 5, 5, 5, 5, 5, 5, 6, 6, 5, 4, 5, 6, 5, 5}}, 
 {12, {3, 5, 5, 5, 5, 5, 5, 5, 5, 5, 3, 5, 5, 6, 6, 6, 6, 5}}, 
 {12, {3, 5, 5, 5, 5, 5, 5, 5, 4, 5, 6, 6, 5, 5, 6, 5, 5, 5}}, 
 {12, {3, 5, 5, 5, 5, 5, 5, 5, 4, 5, 6, 6, 5, 5, 5, 6, 5, 5}}, 
 {12, {3, 5, 5, 5, 5, 5, 5, 5, 3, 5, 5, 5, 5, 6, 6, 6, 6, 5}}, 
 {12, {3, 5, 5, 5, 5, 5, 5, 4, 5, 5, 5, 4, 5, 6, 6, 6, 6, 5}}, 
 {12, {3, 5, 5, 5, 5, 5, 5, 4, 4, 5, 6, 6, 5, 5, 5, 6, 6, 5}}, 
 {12, {3, 5, 5, 5, 5, 5, 4, 5, 6, 6, 5, 5, 5, 5, 6, 5, 5, 5}}, 
 {12, {3, 5, 5, 5, 5, 5, 4, 5, 5, 6, 6, 6, 5, 4, 5, 5, 6, 5}}, 
 {12, {3, 5, 5, 5, 5, 5, 4, 5, 5, 5, 5, 6, 5, 5, 5, 6, 6, 5}}, 
 {12, {3, 5, 5, 5, 5, 5, 4, 4, 5, 6, 6, 5, 5, 5, 5, 6, 6, 5}}, 
 {12, {3, 5, 5, 5, 5, 4, 5, 5, 6, 6, 6, 5, 5, 5, 4, 5, 6, 5}}, 
 {12, {3, 5, 5, 5, 5, 4, 5, 5, 6, 6, 5, 4, 5, 5, 6, 6, 5, 5}}, 
 {12, {3, 5, 5, 5, 5, 4, 5, 5, 5, 6, 5, 5, 5, 6, 5, 5, 6, 5}}, 
 {12, {3, 5, 5, 5, 5, 4, 5, 5, 5, 4, 5, 5, 6, 6, 5, 6, 6, 5}}, 
 {12, {3, 5, 5, 5, 5, 4, 4, 5, 6, 6, 5, 5, 5, 5, 6, 6, 5, 5}}, 
 {12, {3, 5, 5, 5, 5, 4, 4, 5, 5, 6, 6, 5, 5, 5, 5, 6, 6, 5}}, 
 {12, {3, 5, 5, 5, 4, 5, 6, 6, 6, 5, 5, 4, 5, 5, 5, 5, 6, 5}}, 
 {12, {3, 5, 5, 5, 4, 5, 6, 6, 5, 5, 4, 5, 5, 5, 6, 6, 5, 5}}, 
 {12, {3, 5, 5, 5, 4, 5, 6, 5, 5, 5, 5, 6, 5, 5, 5, 5, 6, 5}}, 
 {12, {3, 5, 5, 5, 4, 5, 6, 5, 5, 4, 5, 5, 6, 6, 6, 5, 5, 5}}, 
 {12, {3, 5, 5, 5, 4, 5, 6, 5, 4, 5, 5, 6, 6, 6, 5, 5, 5, 5}}, 
 {12, {3, 5, 5, 5, 4, 5, 5, 6, 6, 5, 4, 5, 5, 6, 6, 5, 5, 5}}, 
 {12, {3, 5, 5, 5, 4, 5, 5, 6, 5, 5, 5, 6, 5, 5, 5, 5, 6, 5}}, 
 {12, {3, 5, 5, 5, 4, 5, 5, 5, 6, 6, 5, 5, 5, 5, 5, 6, 5, 5}}, 
 {12, {3, 5, 5, 5, 4, 5, 5, 5, 6, 6, 5, 5, 4, 5, 5, 6, 6, 5}}, 
 {12, {3, 5, 5, 5, 4, 5, 5, 5, 6, 5, 4, 5, 5, 5, 6, 6, 6, 5}}, 
 {12, {3, 5, 5, 5, 4, 5, 5, 5, 5, 6, 5, 4, 5, 5, 6, 6, 6, 5}}, 
 {12, {3, 5, 5, 5, 4, 5, 5, 5, 5, 4, 5, 6, 6, 6, 6, 5, 5, 5}}, 
 {12, {3, 5, 5, 5, 4, 5, 5, 5, 4, 5, 5, 6, 6, 6, 5, 5, 6, 5}}, 
 {12, {3, 5, 5, 5, 4, 4, 5, 6, 6, 5, 5, 5, 5, 5, 5, 6, 6, 5}}, 
 {12, {3, 5, 5, 5, 4, 4, 5, 6, 5, 5, 5, 6, 6, 6, 5, 5, 5, 5}}, 
 {12, {3, 5, 5, 4, 5, 6, 6, 6, 6, 5, 5, 5, 5, 4, 5, 5, 5, 5}}, 
 {12, {3, 5, 5, 4, 5, 6, 6, 6, 5, 5, 5, 5, 5, 4, 5, 5, 6, 5}}, 
 {12, {3, 5, 5, 4, 5, 6, 6, 5, 5, 5, 5, 5, 6, 5, 5, 5, 5, 5}}, 
 {12, {3, 5, 5, 4, 5, 6, 5, 5, 5, 5, 6, 5, 5, 5, 5, 5, 6, 5}}, 
 {12, {3, 5, 5, 4, 5, 5, 6, 6, 6, 5, 5, 5, 5, 5, 4, 5, 6, 5}}, 
 {12, {3, 5, 5, 4, 5, 5, 6, 5, 5, 5, 6, 5, 5, 5, 5, 5, 6, 5}}, 
 {12, {3, 5, 5, 4, 5, 5, 6, 5, 5, 5, 5, 6, 5, 5, 5, 5, 6, 5}}, 
 {12, {3, 5, 5, 4, 5, 5, 5, 6, 6, 5, 5, 5, 4, 5, 5, 6, 6, 5}}, 
 {12, {3, 5, 5, 4, 5, 5, 5, 5, 5, 6, 5, 5, 6, 6, 5, 5, 5, 5}}, 
 {12, {3, 5, 5, 4, 5, 5, 5, 5, 5, 5, 6, 6, 6, 6, 5, 4, 5, 5}}, 
 {12, {3, 5, 5, 4, 5, 5, 5, 5, 5, 5, 4, 5, 6, 6, 6, 6, 5, 5}}, 
 {12, {3, 5, 5, 4, 5, 5, 5, 5, 5, 5, 4, 5, 5, 6, 6, 6, 6, 5}}, 
 {12, {3, 5, 5, 4, 5, 5, 4, 5, 5, 6, 6, 6, 6, 5, 5, 5, 5, 5}}, 
 {12, {3, 5, 5, 4, 4, 5, 6, 6, 5, 5, 5, 5, 5, 5, 5, 6, 6, 5}}, 
 {12, {3, 5, 5, 4, 4, 5, 5, 6, 6, 5, 5, 5, 5, 5, 5, 6, 6, 5}}, 
 {12, {3, 5, 5, 4, 4, 5, 5, 6, 5, 5, 5, 5, 5, 6, 6, 6, 5, 5}}, 
 {12, {3, 5, 5, 4, 4, 5, 5, 5, 5, 5, 6, 5, 5, 5, 6, 6, 6, 5}}, 
 {12, {1, 6, 6, 6, 5, 5, 5, 5, 4, 5, 5, 5, 3, 5, 6, 6, 6, 6}}, 
 {12, {1, 6, 6, 6, 5, 5, 5, 4, 5, 5, 5, 4, 4, 5, 6, 6, 6, 6}}, 
 {12, {1, 6, 6, 6, 5, 5, 5, 4, 5, 4, 4, 5, 5, 5, 6, 6, 6, 6}}, 
 {12, {1, 6, 6, 6, 5, 5, 4, 5, 5, 5, 5, 5, 3, 5, 6, 6, 6, 6}}, 
 {12, {1, 6, 6, 6, 5, 5, 4, 5, 5, 5, 5, 3, 5, 5, 6, 6, 6, 6}}, 
 {12, {1, 6, 6, 6, 5, 5, 3, 5, 5, 5, 4, 5, 5, 5, 6, 6, 6, 6}}, 
 {12, {1, 6, 6, 6, 5, 4, 5, 5, 5, 4, 4, 5, 5, 5, 6, 6, 6, 6}}, 
 {12, {1, 6, 6, 6, 5, 4, 5, 5, 4, 4, 5, 5, 5, 5, 6, 6, 6, 6}}, 
 {12, {1, 6, 6, 6, 1, 6, 6, 6, 5, 5, 5, 4, 4, 5, 6, 6, 6, 6}}, 
 {12, {1, 6, 6, 5, 5, 6, 5, 5, 5, 4, 5, 5, 5, 4, 5, 6, 6, 6}}, 
 {12, {1, 6, 6, 5, 5, 5, 5, 6, 5, 5, 5, 3, 5, 5, 5, 6, 6, 6}}, 
 {12, {1, 6, 6, 5, 5, 5, 5, 5, 5, 6, 5, 5, 5, 3, 5, 6, 6, 6}}, 
 {12, {1, 6, 6, 5, 5, 5, 5, 4, 4, 5, 5, 5, 5, 6, 6, 5, 6, 6}}, 
 {12, {1, 6, 6, 5, 5, 5, 5, 3, 5, 6, 6, 5, 5, 5, 5, 5, 6, 6}}, 
 {12, {1, 6, 6, 5, 5, 5, 4, 5, 5, 6, 5, 4, 5, 5, 5, 6, 6, 6}}, 
 {12, {1, 6, 6, 5, 5, 5, 4, 5, 5, 5, 6, 5, 5, 4, 5, 6, 6, 6}}, 
 {12, {1, 6, 6, 5, 5, 5, 4, 5, 5, 4, 4, 5, 5, 6, 6, 6, 6, 6}}, 
 {12, {1, 6, 6, 5, 5, 4, 5, 6, 6, 5, 5, 4, 5, 5, 5, 5, 6, 6}}, 
 {12, {1, 6, 6, 5, 5, 4, 5, 6, 6, 5, 4, 5, 5, 5, 5, 5, 6, 6}}, 
 {12, {1, 6, 6, 5, 5, 4, 5, 5, 5, 5, 5, 4, 5, 6, 6, 5, 6, 6}}, 
 {12, {1, 6, 6, 5, 5, 4, 5, 5, 4, 5, 5, 6, 5, 5, 5, 6, 6, 6}}, 
 {12, {1, 6, 6, 5, 5, 3, 5, 5, 6, 6, 5, 5, 5, 5, 5, 5, 6, 6}}, 
 {12, {1, 6, 6, 5, 4, 5, 5, 6, 6, 5, 5, 4, 4, 5, 5, 6, 6, 6}}, 
 {12, {1, 6, 6, 5, 4, 5, 5, 6, 5, 4, 4, 5, 5, 5, 6, 6, 6, 6}}, 
 {12, {1, 6, 6, 5, 4, 5, 5, 5, 4, 5, 5, 6, 6, 5, 5, 5, 6, 6}}, 
 {12, {1, 6, 6, 5, 4, 4, 5, 6, 6, 5, 5, 4, 5, 5, 5, 6, 6, 6}}, 
 {12, {1, 6, 6, 5, 4, 4, 5, 5, 4, 5, 5, 5, 5, 6, 6, 6, 6, 6}}, 
 {12, {1, 6, 6, 1, 6, 6, 6, 6, 5, 5, 4, 4, 5, 5, 6, 6, 6, 6}}, 
 {12, {1, 6, 6, 1, 6, 6, 6, 5, 4, 5, 6, 5, 5, 5, 5, 6, 6, 6}}, 
 {12, {4, 5, 6, 5, 5, 6, 5, 5, 6, 6, 5, 4, 5, 6, 6, 6, 5}}, 
 {12, {4, 5, 5, 6, 5, 5, 6, 6, 5, 6, 6, 6, 5, 4, 5, 6, 5}}, 
 {12, {4, 5, 5, 6, 5, 5, 6, 6, 5, 4, 5, 6, 6, 5, 6, 6, 5}}, 
 {12, {4, 5, 5, 5, 6, 5, 4, 5, 6, 6, 5, 6, 6, 5, 6, 6, 5}}, 
 {12, {4, 5, 5, 5, 5, 4, 5, 6, 6, 5, 6, 6, 6, 5, 6, 6, 5}}, 
 {12, {4, 5, 4, 5, 5, 5, 6, 6, 5, 6, 6, 5, 6, 6, 6, 5, 5}}, 
 {12, {4, 4, 5, 5, 6, 6, 5, 4, 5, 6, 6, 6, 5, 6, 6, 6, 5}}, 
 {12, {4, 4, 5, 5, 4, 5, 6, 6, 6, 5, 6, 6, 6, 5, 6, 6, 5}}, 
 {12, {4, 4, 5, 4, 5, 6, 6, 6, 5, 6, 6, 5, 5, 6, 6, 6, 5}}, 
 {12, {4, 4, 5, 4, 5, 5, 6, 6, 6, 5, 6, 6, 5, 6, 6, 6, 5}}, 
 {12, {4, 4, 4, 4, 5, 5, 5, 6, 6, 6, 6, 6, 6, 6, 6, 6, 5}}, 
 {12, {3, 5, 6, 6, 5, 4, 5, 5, 5, 6, 6, 6, 5, 6, 6, 6, 5}}, 
 {12, {3, 5, 6, 6, 5, 4, 5, 5, 5, 6, 6, 5, 6, 6, 6, 6, 5}}, 
 {12, {3, 5, 5, 6, 6, 6, 6, 5, 5, 4, 5, 6, 6, 5, 6, 6, 5}}, 
 {12, {3, 5, 5, 6, 6, 6, 6, 5, 4, 5, 5, 6, 6, 5, 6, 6, 5}}, 
 {12, {3, 5, 5, 6, 6, 6, 5, 3, 5, 5, 6, 6, 6, 6, 6, 6, 5}}, 
 {12, {3, 5, 5, 6, 6, 5, 6, 6, 6, 6, 6, 5, 4, 5, 5, 6, 5}}, 
 {12, {3, 5, 5, 6, 6, 5, 6, 6, 6, 5, 5, 4, 5, 6, 6, 6, 5}}, 
 {12, {3, 5, 5, 6, 6, 5, 5, 6, 6, 6, 5, 4, 5, 6, 6, 6, 5}}, 
 {12, {3, 5, 5, 6, 6, 5, 5, 4, 5, 6, 6, 6, 6, 5, 6, 6, 5}}, 
 {12, {3, 5, 5, 6, 6, 5, 4, 4, 5, 6, 6, 6, 6, 6, 6, 6, 5}}, 
 {12, {3, 5, 5, 6, 5, 5, 6, 6, 6, 6, 5, 5, 6, 5, 5, 6, 5}}, 
 {12, {3, 5, 5, 6, 5, 4, 5, 6, 6, 6, 6, 5, 6, 6, 6, 5, 5}}, 
 {12, {3, 5, 5, 6, 5, 4, 5, 5, 6, 6, 6, 6, 6, 5, 6, 6, 5}}, 
 {12, {3, 5, 5, 5, 6, 6, 5, 6, 6, 6, 6, 6, 5, 4, 5, 6, 5}}, 
 {12, {3, 5, 5, 5, 6, 6, 5, 6, 6, 6, 5, 5, 6, 5, 5, 6, 5}}, 
 {12, {3, 5, 5, 5, 6, 6, 5, 6, 6, 5, 5, 6, 5, 5, 6, 6, 5}}, 
 {12, {3, 5, 5, 5, 6, 5, 4, 5, 6, 6, 5, 6, 6, 6, 6, 6, 5}}, 
 {12, {3, 5, 5, 5, 4, 5, 6, 6, 6, 5, 6, 6, 5, 6, 6, 6, 5}}, 
 {12, {3, 5, 5, 4, 5, 6, 6, 6, 6, 5, 6, 6, 6, 5, 5, 6, 5}}, 
 {12, {3, 5, 5, 4, 5, 6, 6, 6, 5, 5, 6, 6, 6, 5, 6, 6, 5}}, 
 {12, {3, 5, 5, 4, 5, 6, 6, 5, 5, 6, 6, 6, 6, 5, 6, 6, 5}}, 
 {12, {3, 5, 5, 4, 4, 5, 6, 6, 6, 6, 6, 6, 6, 5, 6, 6, 5}}, 
 {12, {1, 6, 6, 6, 5, 6, 6, 6, 6, 5, 5, 3, 5, 6, 6, 6, 6}}, 
 {12, {1, 6, 6, 5, 5, 6, 6, 6, 6, 6, 6, 5, 3, 5, 6, 6, 6}}, 
 {12, {1, 6, 6, 5, 5, 3, 5, 6, 6, 6, 6, 6, 5, 6, 6, 6, 6}}, 
 {12, {1, 6, 6, 5, 4, 5, 6, 6, 6, 5, 6, 6, 6, 5, 5, 6, 6}}, 
 {11, {4, 4, 5, 5, 5, 4, 5, 5, 5, 5, 4, 5, 5, 4, 5, 5, 5, 5, 5}}, 
 {11, {4, 4, 5, 5, 5, 4, 5, 5, 5, 4, 5, 5, 5, 5, 4, 5, 5, 5, 5}}, 
 {11, {4, 4, 5, 5, 4, 5, 5, 5, 5, 5, 5, 5, 4, 4, 5, 5, 5, 5, 5}}, 
 {11, {4, 4, 5, 5, 4, 5, 5, 5, 5, 5, 4, 5, 5, 5, 5, 4, 5, 5, 5}}, 
 {11, {4, 4, 5, 5, 4, 5, 5, 4, 5, 5, 5, 5, 4, 5, 5, 5, 5, 5, 5}}, 
 {11, {4, 4, 5, 5, 4, 4, 5, 5, 5, 5, 5, 4, 5, 5, 5, 5, 5, 5, 5}}, 
 {11, {4, 4, 5, 5, 4, 4, 5, 5, 5, 4, 4, 5, 6, 5, 5, 5, 5, 5, 5}}, 
 {11, {4, 4, 5, 4, 5, 5, 5, 5, 4, 4, 5, 5, 5, 5, 5, 5, 5, 5, 5}}, 
 {11, {3, 5, 5, 5, 5, 5, 4, 5, 5, 4, 4, 5, 5, 5, 5, 5, 5, 5, 5}}, 
 {11, {3, 5, 5, 5, 5, 4, 5, 5, 4, 4, 5, 5, 5, 5, 5, 5, 5, 5, 5}}, 
 {11, {3, 5, 5, 5, 4, 4, 5, 5, 5, 5, 5, 5, 5, 5, 5, 4, 5, 5, 5}}, 
 {11, {3, 5, 5, 4, 5, 5, 5, 4, 4, 5, 5, 5, 5, 5, 5, 5, 5, 5, 5}}, 
 {11, {4, 5, 5, 5, 6, 5, 5, 4, 5, 5, 5, 6, 5, 5, 4, 5, 6, 5}}, 
 {11, {4, 5, 5, 5, 6, 5, 4, 5, 6, 5, 5, 4, 5, 5, 5, 6, 5, 5}}, 
 {11, {4, 5, 5, 5, 6, 5, 4, 5, 5, 5, 6, 5, 5, 5, 4, 5, 6, 5}}, 
 {11, {4, 5, 5, 5, 5, 6, 5, 4, 5, 5, 6, 5, 5, 4, 5, 5, 6, 5}}, 
 {11, {4, 5, 5, 5, 5, 6, 5, 4, 5, 5, 5, 6, 5, 5, 4, 5, 6, 5}}, 
 {11, {4, 5, 5, 5, 5, 5, 6, 5, 4, 5, 6, 5, 4, 5, 6, 5, 5, 5}}, 
 {11, {4, 5, 5, 5, 5, 4, 5, 5, 6, 5, 5, 6, 5, 4, 5, 5, 6, 5}}, 
 {11, {4, 5, 5, 5, 4, 5, 6, 5, 5, 5, 5, 4, 5, 6, 5, 5, 6, 5}}, 
 {11, {4, 5, 5, 5, 4, 5, 5, 5, 6, 5, 5, 6, 5, 4, 5, 5, 6, 5}}, 
 {11, {4, 5, 5, 5, 4, 5, 5, 5, 6, 5, 5, 4, 5, 6, 5, 5, 6, 5}}, 
 {11, {4, 5, 5, 5, 4, 5, 5, 5, 5, 6, 5, 4, 5, 6, 5, 5, 6, 5}}, 
 {11, {4, 5, 5, 4, 5, 6, 5, 5, 5, 5, 4, 5, 6, 5, 5, 5, 6, 5}}, 
 {11, {4, 5, 5, 4, 5, 6, 5, 5, 5, 4, 5, 5, 6, 5, 5, 5, 6, 5}}, 
 {11, {4, 5, 5, 4, 5, 5, 6, 5, 5, 4, 5, 5, 5, 6, 5, 5, 6, 5}}, 
 {11, {4, 5, 5, 4, 5, 5, 6, 5, 4, 5, 6, 5, 5, 5, 5, 5, 6, 5}}, 
 {11, {4, 5, 5, 4, 5, 5, 5, 5, 5, 4, 5, 6, 6, 5, 5, 6, 5, 5}}, 
 {11, {4, 5, 4, 5, 5, 5, 6, 5, 5, 5, 5, 5, 4, 5, 6, 6, 5, 5}}, 
 {11, {4, 5, 4, 5, 5, 5, 6, 5, 5, 5, 4, 5, 5, 6, 6, 5, 5, 5}}, 
 {11, {4, 5, 4, 5, 5, 5, 6, 5, 4, 5, 5, 5, 5, 5, 6, 6, 5, 5}}, 
 {11, {4, 5, 4, 5, 5, 5, 5, 5, 6, 5, 5, 5, 5, 5, 5, 5, 6, 5}}, 
 {11, {4, 5, 4, 5, 5, 5, 5, 5, 5, 6, 5, 5, 5, 6, 5, 5, 5, 5}}, 
 {11, {4, 5, 4, 5, 5, 4, 5, 5, 5, 5, 5, 6, 6, 6, 5, 5, 5, 5}}, 
 {11, {4, 4, 5, 6, 5, 5, 4, 5, 5, 5, 6, 5, 4, 5, 5, 6, 6, 5}}, 
 {11, {4, 4, 5, 6, 5, 4, 5, 5, 5, 6, 6, 5, 5, 4, 5, 5, 6, 5}}, 
 {11, {4, 4, 5, 6, 5, 4, 5, 5, 5, 5, 5, 5, 4, 5, 6, 6, 6, 5}}, 
 {11, {4, 4, 5, 5, 6, 6, 5, 5, 5, 4, 5, 6, 5, 4, 5, 5, 6, 5}}, 
 {11, {4, 4, 5, 5, 6, 6, 5, 4, 5, 5, 5, 5, 4, 5, 5, 6, 6, 5}}, 
 {11, {4, 4, 5, 5, 6, 5, 5, 6, 5, 5, 5, 4, 5, 5, 5, 5, 6, 5}}, 
 {11, {4, 4, 5, 5, 6, 5, 5, 6, 5, 4, 5, 5, 5, 5, 5, 5, 6, 5}}, 
 {11, {4, 4, 5, 5, 6, 5, 5, 5, 6, 5, 5, 5, 5, 4, 5, 5, 6, 5}}, 
 {11, {4, 4, 5, 5, 6, 5, 5, 5, 5, 6, 5, 5, 5, 5, 4, 5, 6, 5}}, 
 {11, {4, 4, 5, 5, 6, 5, 5, 5, 5, 5, 4, 5, 5, 6, 5, 5, 6, 5}}, 
 {11, {4, 4, 5, 5, 6, 5, 4, 5, 6, 5, 5, 4, 5, 5, 6, 6, 5, 5}}, 
 {11, {4, 4, 5, 5, 6, 5, 4, 5, 5, 5, 5, 4, 5, 6, 6, 6, 5, 5}}, 
 {11, {4, 4, 5, 5, 6, 5, 4, 4, 5, 6, 6, 5, 5, 5, 5, 6, 5, 5}}, 
 {11, {4, 4, 5, 5, 5, 6, 6, 6, 5, 5, 5, 4, 5, 5, 4, 5, 6, 5}}, 
 {11, {4, 4, 5, 5, 5, 6, 6, 6, 5, 4, 5, 5, 5, 4, 5, 6, 5, 5}}, 
 {11, {4, 4, 5, 5, 5, 6, 6, 5, 5, 6, 5, 4, 4, 5, 5, 6, 5, 5}}, 
 {11, {4, 4, 5, 5, 5, 6, 5, 5, 6, 6, 5, 5, 4, 4, 5, 6, 5, 5}}, 
 {11, {4, 4, 5, 5, 5, 5, 6, 6, 6, 5, 4, 4, 5, 5, 6, 5, 5, 5}}, 
 {11, {4, 4, 5, 5, 5, 5, 6, 6, 5, 5, 6, 5, 4, 4, 5, 5, 6, 5}}, 
 {11, {4, 4, 5, 5, 5, 5, 6, 5, 5, 5, 6, 5, 5, 5, 4, 5, 6, 5}}, 
 {11, {4, 4, 5, 5, 5, 5, 6, 5, 4, 4, 5, 6, 5, 5, 5, 6, 6, 5}}, 
 {11, {4, 4, 5, 5, 5, 5, 5, 6, 5, 5, 6, 5, 4, 5, 5, 6, 5, 5}}, 
 {11, {4, 4, 5, 5, 5, 5, 5, 6, 5, 4, 5, 5, 6, 5, 5, 6, 5, 5}}, 
 {11, {4, 4, 5, 5, 5, 5, 5, 5, 6, 5, 5, 6, 5, 5, 4, 5, 6, 5}}, 
 {11, {4, 4, 5, 5, 5, 5, 5, 5, 6, 5, 5, 4, 4, 5, 6, 6, 6, 5}}, 
 {11, {4, 4, 5, 5, 5, 5, 5, 4, 5, 6, 5, 5, 5, 6, 5, 5, 6, 5}}, 
 {11, {4, 4, 5, 5, 5, 5, 4, 5, 6, 6, 5, 5, 5, 4, 5, 6, 6, 5}}, 
 {11, {4, 4, 5, 5, 5, 5, 4, 5, 6, 5, 5, 5, 5, 6, 5, 5, 6, 5}}, 
 {11, {4, 4, 5, 5, 5, 5, 4, 5, 6, 5, 4, 5, 5, 5, 6, 6, 6, 5}}, 
 {11, {4, 4, 5, 5, 5, 5, 4, 5, 5, 4, 5, 6, 6, 5, 5, 6, 6, 5}}, 
 {11, {4, 4, 5, 5, 5, 4, 5, 6, 6, 5, 5, 5, 4, 5, 6, 6, 5, 5}}, 
 {11, {4, 4, 5, 5, 5, 4, 5, 6, 6, 5, 5, 4, 5, 6, 6, 5, 5, 5}}, 
 {11, {4, 4, 5, 5, 5, 4, 5, 6, 5, 5, 5, 5, 6, 5, 5, 6, 5, 5}}, 
 {11, {4, 4, 5, 5, 5, 4, 5, 6, 5, 4, 5, 6, 6, 6, 5, 5, 5, 5}}, 
 {11, {4, 4, 5, 5, 5, 4, 5, 5, 6, 6, 6, 5, 4, 5, 5, 6, 5, 5}}, 
 {11, {4, 4, 5, 5, 5, 4, 5, 5, 6, 5, 5, 6, 5, 5, 6, 5, 5, 5}}, 
 {11, {4, 4, 5, 5, 5, 4, 5, 5, 5, 6, 5, 5, 6, 5, 5, 6, 5, 5}}, 
 {11, {4, 4, 5, 5, 5, 4, 5, 5, 5, 4, 5, 6, 6, 5, 6, 6, 5, 5}}, 
 {11, {4, 4, 5, 5, 5, 4, 5, 5, 4, 5, 6, 6, 6, 5, 5, 5, 6, 5}}, 
 {11, {4, 4, 5, 5, 5, 4, 4, 5, 5, 6, 6, 5, 5, 6, 5, 5, 6, 5}}, 
 {11, {4, 4, 5, 5, 5, 4, 4, 5, 5, 6, 5, 5, 6, 6, 5, 5, 6, 5}}, 
 {11, {4, 4, 5, 5, 4, 5, 6, 6, 6, 5, 5, 5, 5, 5, 4, 5, 6, 5}}, 
 {11, {4, 4, 5, 5, 4, 5, 6, 6, 6, 5, 5, 5, 5, 4, 5, 5, 6, 5}}, 
 {11, {4, 4, 5, 5, 4, 5, 6, 6, 6, 5, 5, 5, 4, 5, 5, 5, 6, 5}}, 
 {11, {4, 4, 5, 5, 4, 5, 6, 6, 6, 5, 5, 4, 5, 5, 6, 5, 5, 5}}, 
 {11, {4, 4, 5, 5, 4, 5, 6, 6, 6, 5, 5, 4, 5, 5, 5, 6, 5, 5}}, 
 {11, {4, 4, 5, 5, 4, 5, 6, 6, 5, 5, 4, 5, 5, 5, 6, 6, 5, 5}}, 
 {11, {4, 4, 5, 5, 4, 5, 6, 6, 5, 4, 5, 5, 6, 6, 5, 5, 5, 5}}, 
 {11, {4, 4, 5, 5, 4, 5, 6, 6, 5, 4, 5, 5, 5, 6, 6, 5, 5, 5}}, 
 {11, {4, 4, 5, 5, 4, 5, 6, 5, 5, 6, 6, 5, 5, 5, 4, 5, 6, 5}}, 
 {11, {4, 4, 5, 5, 4, 5, 6, 5, 5, 6, 5, 5, 5, 5, 6, 5, 5, 5}}, 
 {11, {4, 4, 5, 5, 4, 5, 6, 5, 5, 5, 6, 6, 6, 5, 4, 5, 5, 5}}, 
 {11, {4, 4, 5, 5, 4, 5, 6, 5, 5, 5, 5, 6, 5, 5, 6, 5, 5, 5}}, 
 {11, {4, 4, 5, 5, 4, 5, 6, 5, 5, 5, 5, 4, 5, 6, 6, 6, 5, 5}}, 
 {11, {4, 4, 5, 5, 4, 5, 5, 6, 6, 6, 5, 4, 5, 5, 5, 6, 5, 5}}, 
 {11, {4, 4, 5, 5, 4, 5, 5, 6, 6, 5, 5, 6, 6, 5, 4, 5, 5, 5}}, 
 {11, {4, 4, 5, 5, 4, 5, 5, 6, 5, 5, 6, 5, 5, 6, 5, 5, 5, 5}}, 
 {11, {4, 4, 5, 5, 4, 5, 5, 6, 5, 5, 6, 5, 5, 5, 6, 5, 5, 5}}, 
 {11, {4, 4, 5, 5, 4, 5, 5, 6, 5, 5, 5, 6, 5, 5, 6, 5, 5, 5}}, 
 {11, {4, 4, 5, 5, 4, 5, 5, 5, 6, 6, 6, 5, 5, 6, 5, 4, 5, 5}}, 
 {11, {4, 4, 5, 5, 4, 5, 5, 5, 6, 6, 5, 5, 6, 6, 5, 4, 5, 5}}, 
 {11, {4, 4, 5, 5, 4, 5, 5, 5, 5, 4, 5, 6, 5, 5, 6, 6, 6, 5}}, 
 {11, {4, 4, 5, 5, 4, 5, 5, 5, 4, 5, 6, 6, 5, 5, 6, 6, 5, 5}}, 
 {11, {4, 4, 5, 5, 4, 5, 5, 5, 4, 5, 5, 5, 6, 6, 5, 6, 6, 5}}, 
 {11, {4, 4, 5, 5, 4, 5, 5, 4, 5, 6, 6, 6, 5, 5, 5, 6, 5, 5}}, 
 {11, {4, 4, 5, 5, 4, 5, 5, 4, 5, 6, 5, 5, 5, 6, 6, 6, 5, 5}}, 
 {11, {4, 4, 5, 5, 4, 4, 5, 5, 5, 4, 5, 5, 6, 6, 6, 6, 6, 5}}, 
 {11, {4, 4, 5, 4, 5, 6, 6, 6, 6, 5, 5, 5, 4, 5, 5, 5, 5, 5}}, 
 {11, {4, 4, 5, 4, 5, 6, 6, 6, 6, 5, 4, 5, 5, 5, 5, 5, 5, 5}}, 
 {11, {4, 4, 5, 4, 5, 6, 5, 5, 5, 4, 5, 5, 5, 6, 6, 6, 5, 5}}, 
 {11, {4, 4, 5, 4, 5, 6, 5, 5, 5, 4, 5, 5, 5, 5, 6, 6, 6, 5}}, 
 {11, {4, 4, 5, 4, 5, 5, 6, 6, 6, 5, 5, 5, 5, 4, 5, 6, 5, 5}}, 
 {11, {4, 4, 5, 4, 5, 5, 6, 6, 5, 5, 6, 6, 5, 5, 4, 5, 5, 5}}, 
 {11, {4, 4, 5, 4, 5, 5, 6, 6, 5, 5, 5, 4, 5, 6, 6, 5, 5, 5}}, 
 {11, {4, 4, 5, 4, 5, 5, 6, 6, 5, 4, 5, 5, 5, 5, 5, 6, 6, 5}}, 
 {11, {4, 4, 5, 4, 5, 5, 6, 5, 5, 5, 5, 5, 5, 6, 5, 5, 6, 5}}, 
 {11, {4, 4, 5, 4, 5, 5, 5, 6, 6, 6, 6, 5, 4, 5, 5, 5, 5, 5}}, 
 {11, {4, 4, 5, 4, 5, 5, 5, 6, 6, 5, 5, 5, 4, 5, 6, 6, 5, 5}}, 
 {11, {4, 4, 5, 4, 5, 5, 5, 6, 5, 5, 5, 6, 5, 5, 5, 6, 5, 5}}, 
 {11, {4, 4, 5, 4, 5, 5, 5, 6, 5, 5, 5, 5, 6, 5, 5, 5, 6, 5}}, 
 {11, {4, 4, 5, 4, 5, 5, 5, 5, 6, 5, 5, 5, 6, 5, 5, 5, 6, 5}}, 
 {11, {4, 4, 5, 4, 5, 5, 5, 5, 5, 6, 5, 5, 6, 5, 5, 5, 6, 5}}, 
 {11, {4, 4, 5, 4, 5, 5, 4, 5, 5, 6, 6, 6, 5, 5, 5, 5, 6, 5}}, 
 {11, {4, 4, 5, 4, 4, 5, 5, 5, 5, 4, 5, 5, 6, 6, 6, 6, 6, 5}}, 
 {11, {4, 4, 5, 4, 4, 5, 5, 4, 5, 5, 5, 5, 6, 6, 6, 6, 6, 5}}, 
 {11, {3, 5, 6, 6, 5, 5, 5, 4, 4, 5, 5, 5, 5, 5, 5, 6, 6, 5}}, 
 {11, {3, 5, 6, 6, 5, 5, 4, 5, 5, 5, 4, 5, 5, 5, 5, 6, 6, 5}}, 
 {11, {3, 5, 6, 5, 5, 5, 5, 6, 5, 5, 5, 4, 4, 5, 5, 6, 6, 5}}, 
 {11, {3, 5, 6, 5, 5, 5, 5, 5, 5, 6, 5, 4, 4, 5, 5, 6, 6, 5}}, 
 {11, {3, 5, 6, 5, 5, 5, 5, 4, 4, 5, 6, 5, 5, 5, 5, 6, 6, 5}}, 
 {11, {3, 5, 6, 5, 5, 5, 4, 4, 5, 6, 6, 5, 5, 5, 5, 5, 6, 5}}, 
 {11, {3, 5, 6, 5, 5, 4, 5, 5, 5, 5, 6, 5, 4, 5, 5, 6, 6, 5}}, 
 {11, {3, 5, 6, 5, 5, 4, 5, 5, 5, 5, 5, 5, 4, 5, 6, 6, 6, 5}}, 
 {11, {3, 5, 6, 5, 4, 5, 5, 5, 5, 6, 5, 5, 5, 5, 5, 5, 6, 5}}, 
 {11, {3, 5, 5, 6, 6, 6, 5, 5, 4, 5, 5, 4, 5, 5, 5, 5, 6, 5}}, 
 {11, {3, 5, 5, 6, 6, 5, 5, 5, 6, 5, 4, 4, 5, 5, 5, 5, 6, 5}}, 
 {11, {3, 5, 5, 6, 6, 5, 4, 4, 5, 5, 5, 5, 5, 6, 5, 5, 6, 5}}, 
 {11, {3, 5, 5, 6, 5, 5, 6, 5, 5, 5, 5, 5, 5, 4, 5, 5, 6, 5}}, 
 {11, {3, 5, 5, 6, 5, 5, 5, 5, 4, 5, 5, 5, 5, 5, 5, 6, 6, 5}}, 
 {11, {3, 5, 5, 6, 5, 4, 5, 6, 5, 5, 5, 5, 5, 5, 5, 5, 6, 5}}, 
 {11, {3, 5, 5, 5, 6, 6, 6, 5, 5, 5, 5, 5, 4, 4, 5, 6, 5, 5}}, 
 {11, {3, 5, 5, 5, 6, 6, 5, 5, 4, 5, 5, 5, 5, 5, 5, 6, 5, 5}}, 
 {11, {3, 5, 5, 5, 6, 5, 5, 6, 5, 5, 5, 4, 5, 5, 5, 5, 6, 5}}, 
 {11, {3, 5, 5, 5, 6, 5, 5, 5, 5, 5, 6, 5, 4, 5, 5, 5, 6, 5}}, 
 {11, {3, 5, 5, 5, 6, 5, 5, 5, 5, 5, 5, 6, 5, 5, 4, 5, 6, 5}}, 
 {11, {3, 5, 5, 5, 6, 5, 5, 5, 5, 4, 5, 5, 5, 5, 5, 6, 6, 5}}, 
 {11, {3, 5, 5, 5, 6, 5, 5, 5, 4, 5, 6, 5, 5, 5, 5, 5, 6, 5}}, 
 {11, {3, 5, 5, 5, 6, 5, 4, 5, 6, 5, 5, 5, 5, 5, 5, 5, 6, 5}}, 
 {11, {3, 5, 5, 5, 6, 5, 4, 5, 5, 6, 5, 5, 5, 5, 5, 5, 6, 5}}, 
 {11, {3, 5, 5, 5, 6, 5, 4, 5, 5, 5, 6, 5, 5, 5, 5, 5, 6, 5}}, 
 {11, {3, 5, 5, 5, 5, 6, 6, 6, 5, 5, 4, 5, 5, 5, 4, 5, 6, 5}}, 
 {11, {3, 5, 5, 5, 5, 6, 6, 6, 5, 5, 4, 5, 5, 4, 5, 5, 6, 5}}, 
 {11, {3, 5, 5, 5, 5, 6, 6, 6, 5, 4, 4, 5, 5, 5, 6, 5, 5, 5}}, 
 {11, {3, 5, 5, 5, 5, 6, 6, 5, 5, 5, 4, 5, 5, 5, 5, 6, 5, 5}}, 
 {11, {3, 5, 5, 5, 5, 6, 6, 5, 5, 4, 4, 5, 5, 5, 5, 6, 6, 5}}, 
 {11, {3, 5, 5, 5, 5, 6, 5, 5, 5, 5, 5, 6, 5, 5, 4, 5, 6, 5}}, 
 {11, {3, 5, 5, 5, 5, 6, 5, 5, 5, 5, 5, 4, 5, 6, 6, 5, 5, 5}}, 
 {11, {3, 5, 5, 5, 5, 6, 5, 5, 5, 5, 4, 5, 5, 5, 5, 6, 6, 5}}, 
 {11, {3, 5, 5, 5, 5, 6, 5, 5, 5, 4, 5, 5, 6, 5, 5, 5, 6, 5}}, 
 {11, {3, 5, 5, 5, 5, 6, 5, 5, 5, 4, 4, 5, 5, 6, 6, 6, 5, 5}}, 
 {11, {3, 5, 5, 5, 5, 6, 5, 4, 4, 5, 5, 5, 6, 6, 6, 5, 5, 5}}, 
 {11, {3, 5, 5, 5, 5, 5, 6, 6, 6, 5, 5, 5, 5, 3, 5, 5, 6, 5}}, 
 {11, {3, 5, 5, 5, 5, 5, 6, 6, 5, 5, 5, 5, 3, 5, 5, 6, 6, 5}}, 
 {11, {3, 5, 5, 5, 5, 5, 6, 5, 5, 6, 5, 4, 5, 5, 5, 5, 6, 5}}, 
 {11, {3, 5, 5, 5, 5, 5, 6, 5, 5, 5, 4, 4, 5, 5, 6, 6, 6, 5}}, 
 {11, {3, 5, 5, 5, 5, 5, 6, 5, 5, 4, 5, 5, 5, 6, 5, 5, 6, 5}}, 
 {11, {3, 5, 5, 5, 5, 5, 6, 5, 4, 5, 6, 5, 5, 5, 5, 5, 6, 5}}, 
 {11, {3, 5, 5, 5, 5, 5, 6, 5, 4, 5, 5, 5, 6, 5, 5, 5, 6, 5}}, 
 {11, {3, 5, 5, 5, 5, 5, 5, 6, 6, 6, 5, 5, 3, 5, 5, 5, 6, 5}}, 
 {11, {3, 5, 5, 5, 5, 5, 5, 6, 5, 5, 4, 5, 6, 5, 5, 5, 6, 5}}, 
 {11, {3, 5, 5, 5, 5, 5, 5, 5, 6, 6, 5, 5, 4, 5, 5, 6, 5, 5}}, 
 {11, {3, 5, 5, 5, 5, 5, 5, 5, 6, 5, 5, 5, 5, 4, 5, 6, 6, 5}}, 
 {11, {3, 5, 5, 5, 5, 5, 5, 5, 5, 6, 6, 5, 3, 5, 6, 6, 5, 5}}, 
 {11, {3, 5, 5, 5, 5, 5, 5, 5, 5, 6, 5, 4, 5, 6, 5, 5, 6, 5}}, 
 {11, {3, 5, 5, 5, 5, 5, 5, 5, 5, 6, 5, 4, 4, 5, 6, 6, 6, 5}}, 
 {11, {3, 5, 5, 5, 5, 5, 5, 5, 5, 6, 5, 3, 5, 5, 6, 6, 6, 5}}, 
 {11, {3, 5, 5, 5, 5, 5, 5, 5, 4, 5, 6, 5, 5, 6, 5, 5, 6, 5}}, 
 {11, {3, 5, 5, 5, 5, 5, 5, 4, 5, 5, 6, 6, 5, 5, 5, 6, 5, 5}}, 
 {11, {3, 5, 5, 5, 5, 5, 5, 4, 5, 5, 5, 6, 6, 5, 5, 6, 5, 5}}, 
 {11, {3, 5, 5, 5, 5, 5, 5, 3, 5, 5, 5, 5, 6, 6, 6, 6, 5, 5}}, 
 {11, {3, 5, 5, 5, 5, 5, 4, 5, 5, 5, 6, 5, 5, 6, 6, 5, 5, 5}}, 
 {11, {3, 5, 5, 5, 5, 5, 4, 4, 5, 6, 6, 5, 5, 5, 6, 6, 5, 5}}, 
 {11, {3, 5, 5, 5, 5, 5, 4, 4, 5, 5, 6, 5, 5, 6, 6, 6, 5, 5}}, 
 {11, {3, 5, 5, 5, 5, 4, 5, 6, 6, 5, 5, 6, 5, 5, 5, 5, 5, 5}}, 
 {11, {3, 5, 5, 5, 5, 4, 5, 6, 5, 5, 4, 5, 5, 6, 6, 6, 5, 5}}, 
 {11, {3, 5, 5, 5, 5, 4, 5, 5, 6, 6, 6, 5, 4, 5, 5, 5, 6, 5}}, 
 {11, {3, 5, 5, 5, 5, 4, 5, 5, 6, 6, 5, 5, 4, 5, 5, 6, 6, 5}}, 
 {11, {3, 5, 5, 5, 5, 4, 5, 5, 6, 6, 5, 4, 5, 5, 5, 6, 6, 5}}, 
 {11, {3, 5, 5, 5, 5, 4, 5, 5, 6, 5, 5, 6, 5, 5, 5, 5, 6, 5}}, 
 {11, {3, 5, 5, 5, 5, 4, 5, 5, 5, 5, 6, 5, 5, 6, 5, 5, 6, 5}}, 
 {11, {3, 5, 5, 5, 5, 4, 5, 5, 5, 5, 4, 5, 6, 6, 6, 6, 5, 5}}, 
 {11, {3, 5, 5, 5, 5, 4, 5, 5, 4, 5, 5, 6, 6, 6, 6, 5, 5, 5}}, 
 {11, {3, 5, 5, 5, 5, 4, 4, 5, 6, 6, 5, 6, 6, 5, 5, 5, 5, 5}}, 
 {11, {3, 5, 5, 5, 5, 4, 4, 5, 5, 6, 6, 6, 5, 5, 5, 5, 6, 5}}, 
 {11, {3, 5, 5, 5, 5, 4, 4, 5, 5, 6, 6, 5, 5, 6, 6, 5, 5, 5}}, 
 {11, {3, 5, 5, 5, 4, 5, 6, 6, 6, 5, 5, 5, 5, 4, 5, 5, 6, 5}}, 
 {11, {3, 5, 5, 5, 4, 5, 6, 5, 5, 5, 4, 5, 5, 6, 6, 6, 5, 5}}, 
 {11, {3, 5, 5, 5, 4, 5, 5, 6, 6, 6, 5, 5, 5, 5, 4, 5, 6, 5}}, 
 {11, {3, 5, 5, 5, 4, 5, 5, 6, 6, 5, 5, 4, 5, 6, 5, 5, 6, 5}}, 
 {11, {3, 5, 5, 5, 4, 5, 5, 6, 6, 5, 5, 4, 5, 5, 6, 6, 5, 5}}, 
 {11, {3, 5, 5, 5, 4, 5, 5, 6, 6, 5, 4, 5, 5, 5, 6, 6, 5, 5}}, 
 {11, {3, 5, 5, 5, 4, 5, 5, 5, 6, 6, 5, 5, 6, 5, 5, 5, 5, 5}}, 
 {11, {3, 5, 5, 5, 4, 5, 5, 5, 6, 6, 5, 4, 5, 5, 6, 6, 5, 5}}, 
 {11, {3, 5, 5, 5, 4, 5, 5, 5, 6, 5, 5, 4, 5, 5, 6, 6, 6, 5}}, 
 {11, {3, 5, 5, 5, 4, 5, 5, 5, 5, 6, 6, 6, 5, 4, 5, 5, 6, 5}}, 
 {11, {3, 5, 5, 5, 4, 5, 5, 4, 5, 6, 5, 5, 5, 6, 6, 6, 5, 5}}, 
 {11, {3, 5, 5, 5, 4, 4, 5, 6, 6, 5, 5, 6, 6, 5, 5, 5, 5, 5}}, 
 {11, {3, 5, 5, 5, 4, 4, 5, 5, 6, 6, 6, 5, 5, 5, 6, 5, 5, 5}}, 
 {11, {3, 5, 5, 5, 4, 4, 5, 5, 5, 6, 5, 5, 5, 5, 6, 6, 6, 5}}, 
 {11, {3, 5, 5, 4, 5, 6, 6, 6, 5, 5, 4, 5, 5, 5, 5, 5, 6, 5}}, 
 {11, {3, 5, 5, 4, 5, 6, 6, 5, 5, 5, 5, 5, 5, 5, 6, 5, 5, 5}}, 
 {11, {3, 5, 5, 4, 5, 6, 5, 5, 5, 5, 5, 5, 5, 6, 5, 5, 6, 5}}, 
 {11, {3, 5, 5, 4, 5, 6, 5, 5, 4, 5, 6, 6, 6, 5, 5, 5, 5, 5}}, 
 {11, {3, 5, 5, 4, 5, 5, 6, 6, 6, 5, 5, 5, 5, 6, 5, 4, 5, 5}}, 
 {11, {3, 5, 5, 4, 5, 5, 6, 6, 6, 5, 4, 5, 6, 5, 5, 5, 5, 5}}, 
 {11, {3, 5, 5, 4, 5, 5, 6, 6, 5, 5, 4, 5, 5, 6, 6, 5, 5, 5}}, 
 {11, {3, 5, 5, 4, 5, 5, 5, 6, 6, 6, 5, 4, 5, 6, 5, 5, 5, 5}}, 
 {11, {3, 5, 5, 4, 5, 5, 5, 6, 6, 5, 5, 5, 4, 5, 6, 6, 5, 5}}, 
 {11, {3, 5, 5, 4, 5, 5, 5, 6, 5, 5, 6, 5, 5, 5, 5, 5, 6, 5}}, 
 {11, {3, 5, 5, 4, 5, 5, 5, 6, 5, 5, 4, 5, 5, 6, 6, 6, 5, 5}}, 
 {11, {3, 5, 5, 4, 5, 5, 5, 5, 5, 5, 6, 6, 5, 5, 5, 6, 5, 5}}, 
 {11, {3, 5, 5, 4, 5, 5, 5, 5, 5, 5, 6, 5, 5, 6, 6, 5, 5, 5}}, 
 {11, {3, 5, 5, 4, 5, 5, 5, 4, 5, 6, 5, 5, 5, 5, 6, 6, 6, 5}}, 
 {11, {3, 5, 5, 4, 5, 5, 4, 5, 5, 5, 6, 6, 6, 6, 5, 5, 5, 5}}, 
 {11, {3, 5, 5, 4, 5, 5, 4, 5, 5, 5, 5, 6, 6, 6, 6, 5, 5, 5}}, 
 {11, {3, 5, 5, 4, 5, 5, 4, 5, 5, 5, 5, 5, 6, 6, 6, 6, 5, 5}}, 
 {11, {3, 5, 5, 4, 4, 5, 6, 5, 5, 5, 5, 6, 6, 6, 5, 5, 5, 5}}, 
 {11, {3, 5, 5, 4, 4, 5, 5, 6, 6, 5, 5, 5, 5, 5, 6, 6, 5, 5}}, 
 {11, {3, 5, 5, 4, 4, 5, 5, 6, 5, 5, 5, 5, 5, 5, 6, 6, 6, 5}}, 
 {11, {3, 5, 5, 4, 4, 5, 5, 5, 5, 5, 6, 6, 5, 5, 5, 6, 6, 5}}, 
 {11, {3, 5, 4, 4, 5, 5, 5, 5, 6, 6, 6, 6, 5, 5, 5, 5, 5, 5}}, 
 {11, {1, 6, 6, 6, 5, 5, 4, 5, 4, 4, 5, 6, 5, 5, 5, 6, 6, 6}}, 
 {11, {1, 6, 6, 6, 5, 4, 4, 5, 5, 5, 5, 5, 4, 5, 6, 6, 6, 6}}, 
 {11, {1, 6, 6, 6, 5, 3, 5, 6, 5, 5, 5, 5, 5, 4, 5, 6, 6, 6}}, 
 {11, {1, 6, 6, 5, 5, 6, 5, 5, 5, 5, 5, 5, 4, 4, 5, 6, 6, 6}}, 
 {11, {1, 6, 6, 5, 5, 6, 5, 5, 4, 5, 5, 5, 4, 5, 5, 6, 6, 6}}, 
 {11, {1, 6, 6, 5, 5, 6, 5, 5, 4, 4, 5, 5, 5, 5, 5, 6, 6, 6}}, 
 {11, {1, 6, 6, 5, 5, 6, 5, 4, 5, 5, 4, 5, 5, 5, 5, 6, 6, 6}}, 
 {11, {1, 6, 6, 5, 5, 5, 6, 5, 5, 5, 5, 5, 5, 3, 5, 6, 6, 6}}, 
 {11, {1, 6, 6, 5, 5, 5, 5, 6, 5, 5, 5, 5, 3, 5, 5, 6, 6, 6}}, 
 {11, {1, 6, 6, 5, 5, 5, 5, 5, 6, 5, 5, 5, 3, 5, 5, 6, 6, 6}}, 
 {11, {1, 6, 6, 5, 5, 5, 5, 5, 5, 6, 5, 3, 5, 5, 5, 6, 6, 6}}, 
 {11, {1, 6, 6, 5, 5, 5, 5, 5, 5, 5, 3, 5, 6, 6, 5, 5, 6, 6}}, 
 {11, {1, 6, 6, 5, 5, 5, 5, 5, 5, 4, 5, 6, 5, 4, 5, 6, 6, 6}}, 
 {11, {1, 6, 6, 5, 5, 5, 5, 5, 3, 5, 5, 5, 6, 5, 5, 6, 6, 6}}, 
 {11, {1, 6, 6, 5, 5, 5, 5, 4, 5, 5, 5, 6, 5, 4, 5, 6, 6, 6}}, 
 {11, {1, 6, 6, 5, 5, 5, 5, 3, 5, 6, 5, 5, 5, 5, 5, 6, 6, 6}}, 
 {11, {1, 6, 6, 5, 5, 5, 4, 5, 6, 5, 5, 5, 5, 4, 5, 6, 6, 6}}, 
 {11, {1, 6, 6, 5, 5, 5, 4, 5, 5, 5, 6, 6, 1, 6, 6, 6, 6, 6}}, 
 {11, {1, 6, 6, 5, 5, 5, 4, 5, 5, 5, 5, 6, 5, 4, 5, 6, 6, 6}}, 
 {11, {1, 6, 6, 5, 5, 5, 4, 5, 5, 5, 4, 4, 5, 6, 6, 6, 6, 6}}, 
 {11, {1, 6, 6, 5, 5, 5, 4, 5, 5, 4, 5, 5, 6, 5, 5, 6, 6, 6}}, 
 {11, {1, 6, 6, 5, 5, 4, 5, 6, 5, 5, 5, 5, 5, 4, 5, 6, 6, 6}}, 
 {11, {1, 6, 6, 5, 5, 4, 5, 5, 6, 6, 5, 4, 4, 5, 5, 6, 6, 6}}, 
 {11, {1, 6, 6, 5, 5, 4, 5, 5, 6, 5, 5, 6, 6, 6, 1, 6, 6, 6}}, 
 {11, {1, 6, 6, 5, 5, 4, 5, 5, 6, 5, 5, 5, 5, 4, 5, 6, 6, 6}}, 
 {11, {1, 6, 6, 5, 5, 4, 5, 5, 5, 5, 4, 4, 5, 6, 6, 6, 6, 6}}, 
 {11, {1, 6, 6, 5, 5, 4, 5, 5, 5, 4, 5, 6, 6, 5, 5, 5, 6, 6}}, 
 {11, {1, 6, 6, 5, 5, 4, 4, 5, 5, 5, 6, 5, 5, 6, 5, 5, 6, 6}}, 
 {11, {1, 6, 6, 5, 5, 4, 4, 5, 5, 5, 5, 4, 5, 6, 6, 6, 6, 6}}, 
 {11, {1, 6, 6, 5, 5, 4, 4, 5, 5, 4, 5, 5, 5, 6, 6, 6, 6, 6}}, 
 {11, {1, 6, 6, 5, 5, 3, 5, 5, 5, 6, 6, 5, 5, 5, 5, 5, 6, 6}}, 
 {11, {1, 6, 6, 5, 5, 3, 5, 5, 5, 5, 6, 6, 5, 5, 5, 5, 6, 6}}, 
 {11, {1, 6, 6, 5, 4, 5, 6, 6, 5, 5, 5, 5, 5, 4, 5, 5, 6, 6}}, 
 {11, {1, 6, 6, 5, 4, 5, 6, 5, 5, 5, 5, 6, 6, 6, 1, 6, 6, 6}}, 
 {11, {1, 6, 6, 5, 4, 5, 6, 5, 5, 4, 4, 5, 5, 5, 6, 6, 6, 6}}, 
 {11, {1, 6, 6, 5, 4, 5, 5, 5, 5, 5, 6, 6, 5, 4, 5, 5, 6, 6}}, 
 {11, {1, 6, 6, 5, 4, 5, 5, 4, 5, 6, 5, 5, 5, 5, 5, 6, 6, 6}}, 
 {11, {1, 6, 6, 5, 3, 5, 6, 5, 5, 5, 5, 6, 5, 5, 5, 5, 6, 6}}, 
 {11, {1, 6, 6, 5, 3, 5, 5, 5, 6, 5, 5, 5, 5, 5, 5, 6, 6, 6}}, 
 {11, {1, 6, 6, 1, 6, 6, 5, 6, 6, 5, 5, 5, 4, 5, 5, 6, 6, 6}}, 
 {11, {1, 6, 6, 1, 6, 6, 5, 5, 5, 6, 6, 6, 1, 6, 6, 6, 6, 6}}, 
 {11, {1, 6, 6, 1, 6, 6, 5, 5, 5, 6, 6, 5, 4, 5, 5, 6, 6, 6}}, 
 {11, {1, 6, 6, 1, 6, 6, 5, 5, 5, 5, 6, 5, 4, 5, 6, 6, 6, 6}}, 
 {11, {1, 6, 6, 1, 6, 6, 5, 4, 5, 5, 6, 6, 6, 5, 5, 5, 6, 6}}, 
 {11, {1, 6, 6, 1, 6, 6, 1, 6, 6, 6, 6, 5, 5, 5, 6, 6, 6, 6}}, 
 {11, {1, 6, 1, 6, 6, 5, 5, 5, 5, 5, 6, 6, 5, 5, 5, 6, 6, 6}}, 
 {11, {4, 5, 6, 5, 5, 6, 6, 5, 4, 5, 6, 6, 5, 5, 6, 6, 5}}, 
 {11, {4, 5, 6, 5, 5, 6, 5, 5, 6, 6, 6, 5, 4, 5, 6, 6, 5}}, 
 {11, {4, 5, 4, 5, 5, 6, 6, 5, 6, 6, 6, 6, 5, 5, 5, 6, 5}}, 
 {11, {4, 5, 4, 5, 5, 6, 6, 5, 6, 6, 6, 5, 5, 5, 6, 6, 5}}, 
 {11, {4, 5, 4, 5, 5, 6, 6, 5, 5, 5, 6, 6, 6, 5, 6, 6, 5}}, 
 {11, {4, 5, 4, 5, 5, 5, 5, 6, 6, 6, 5, 6, 6, 5, 6, 6, 5}}, 
 {11, {4, 5, 4, 5, 5, 5, 5, 6, 6, 5, 5, 6, 6, 6, 6, 6, 5}}, 
 {11, {4, 5, 4, 5, 5, 4, 5, 6, 6, 6, 6, 6, 6, 6, 6, 5, 5}}, 
 {11, {4, 4, 5, 6, 5, 5, 6, 6, 6, 5, 4, 5, 6, 6, 6, 6, 5}}, 
 {11, {4, 4, 5, 4, 5, 6, 6, 6, 5, 5, 6, 6, 5, 6, 6, 6, 5}}, 
 {11, {4, 4, 5, 4, 5, 6, 6, 5, 5, 6, 6, 6, 6, 5, 6, 6, 5}}, 
 {11, {3, 5, 6, 6, 5, 6, 6, 6, 5, 4, 5, 5, 5, 6, 6, 6, 5}}, 
 {11, {3, 5, 6, 6, 5, 5, 5, 5, 6, 5, 5, 6, 6, 5, 6, 6, 5}}, 
 {11, {3, 5, 6, 6, 5, 5, 5, 4, 5, 6, 6, 5, 6, 6, 6, 6, 5}}, 
 {11, {3, 5, 5, 6, 6, 6, 6, 6, 6, 6, 6, 5, 4, 4, 5, 6, 5}}, 
 {11, {3, 5, 5, 6, 6, 6, 6, 5, 3, 5, 6, 6, 6, 6, 6, 5, 5}}, 
 {11, {3, 5, 5, 6, 6, 6, 5, 6, 6, 6, 6, 5, 5, 4, 5, 6, 5}}, 
 {11, {3, 5, 5, 6, 6, 5, 6, 6, 6, 6, 6, 6, 5, 3, 5, 6, 5}}, 
 {11, {3, 5, 5, 6, 6, 5, 6, 6, 6, 5, 4, 5, 6, 6, 6, 5, 5}}, 
 {11, {3, 5, 5, 6, 6, 5, 4, 5, 6, 6, 5, 5, 6, 6, 6, 6, 5}}, 
 {11, {3, 5, 5, 6, 6, 5, 4, 5, 5, 6, 6, 5, 6, 6, 6, 6, 5}}, 
 {11, {3, 5, 5, 6, 5, 5, 6, 6, 6, 5, 5, 6, 5, 5, 6, 6, 5}}, 
 {11, {3, 5, 5, 6, 5, 5, 5, 6, 6, 5, 5, 6, 6, 5, 6, 6, 5}}, 
 {11, {3, 5, 5, 6, 5, 5, 5, 4, 5, 6, 6, 6, 6, 6, 6, 6, 5}}, 
 {11, {3, 5, 5, 6, 5, 5, 4, 5, 6, 6, 6, 6, 6, 6, 6, 5, 5}}, 
 {11, {3, 5, 5, 6, 5, 4, 5, 6, 6, 6, 5, 5, 6, 6, 6, 6, 5}}, 
 {11, {3, 5, 5, 6, 5, 3, 5, 5, 6, 6, 6, 6, 6, 6, 6, 6, 5}}, 
 {11, {3, 5, 5, 5, 6, 6, 6, 6, 6, 6, 6, 5, 3, 5, 6, 6, 5}}, 
 {11, {3, 5, 5, 5, 4, 5, 6, 6, 5, 6, 6, 6, 6, 5, 6, 6, 5}}, 
 {11, {3, 5, 5, 4, 5, 6, 6, 6, 5, 5, 6, 6, 5, 6, 6, 6, 5}}, 
 {11, {3, 5, 5, 4, 5, 6, 5, 5, 6, 6, 6, 6, 6, 5, 6, 6, 5}}, 
 {11, {3, 5, 5, 4, 5, 6, 5, 5, 6, 6, 5, 6, 6, 6, 6, 6, 5}}, 
 {11, {3, 5, 5, 4, 4, 5, 6, 6, 5, 6, 6, 6, 6, 6, 6, 6, 5}}, 
 {11, {1, 6, 6, 5, 6, 6, 6, 6, 6, 6, 5, 6, 6, 1, 6, 6, 6}}, 
 {11, {1, 6, 6, 5, 5, 6, 5, 4, 5, 6, 6, 6, 5, 6, 6, 6, 6}}, 
 {10, {4, 4, 5, 5, 5, 5, 4, 5, 5, 4, 5, 5, 5, 4, 5, 5, 5, 5, 5}}, 
 {10, {4, 4, 5, 5, 5, 4, 5, 5, 5, 5, 4, 5, 5, 5, 4, 5, 5, 5, 5}}, 
 {10, {4, 4, 5, 5, 4, 5, 5, 5, 5, 4, 5, 5, 5, 4, 5, 5, 5, 5, 5}}, 
 {10, {4, 4, 5, 5, 4, 5, 5, 5, 4, 5, 5, 5, 5, 5, 5, 5, 4, 5, 5}}, 
 {10, {4, 4, 5, 5, 4, 5, 5, 5, 4, 5, 5, 5, 5, 5, 4, 5, 5, 5, 5}}, 
 {10, {4, 4, 5, 4, 5, 5, 5, 4, 5, 5, 5, 5, 5, 5, 5, 5, 4, 5, 5}}, 
 {10, {4, 4, 5, 4, 5, 5, 4, 4, 5, 5, 5, 5, 5, 5, 5, 5, 5, 5, 5}}, 
 {10, {3, 5, 5, 4, 4, 5, 5, 5, 5, 5, 5, 4, 5, 5, 5, 5, 5, 5, 5}}, 
 {10, {4, 5, 5, 5, 6, 5, 4, 5, 5, 6, 5, 4, 5, 5, 5, 6, 5, 5}}, 
 {10, {4, 5, 5, 5, 5, 6, 5, 5, 4, 5, 6, 5, 5, 4, 5, 5, 6, 5}}, 
 {10, {4, 5, 5, 5, 5, 6, 5, 5, 4, 5, 6, 5, 4, 5, 5, 5, 6, 5}}, 
 {10, {4, 5, 5, 5, 5, 6, 5, 5, 4, 5, 5, 6, 5, 4, 5, 5, 6, 5}}, 
 {10, {4, 5, 5, 5, 5, 6, 5, 4, 5, 5, 6, 5, 5, 4, 5, 6, 5, 5}}, 
 {10, {4, 5, 5, 5, 5, 6, 5, 4, 5, 5, 6, 5, 4, 5, 6, 5, 5, 5}}, 
 {10, {4, 5, 5, 5, 5, 5, 5, 6, 5, 4, 5, 6, 5, 4, 5, 5, 6, 5}}, 
 {10, {4, 5, 5, 5, 5, 5, 5, 4, 5, 6, 5, 4, 5, 5, 5, 6, 6, 5}}, 
 {10, {4, 5, 5, 5, 5, 5, 4, 5, 6, 5, 5, 4, 5, 6, 5, 5, 6, 5}}, 
 {10, {4, 5, 5, 5, 5, 4, 5, 6, 5, 4, 5, 6, 5, 5, 5, 5, 6, 5}}, 
 {10, {4, 5, 5, 5, 5, 4, 5, 5, 6, 5, 4, 5, 6, 5, 5, 5, 6, 5}}, 
 {10, {4, 5, 5, 5, 5, 4, 5, 5, 5, 5, 5, 6, 6, 5, 4, 5, 6, 5}}, 
 {10, {4, 5, 5, 5, 4, 5, 5, 6, 5, 5, 5, 5, 6, 5, 4, 5, 6, 5}}, 
 {10, {4, 5, 5, 5, 4, 5, 5, 6, 5, 4, 5, 6, 5, 5, 5, 6, 5, 5}}, 
 {10, {4, 5, 5, 5, 4, 5, 5, 6, 5, 4, 5, 6, 5, 5, 5, 5, 6, 5}}, 
 {10, {4, 5, 5, 5, 4, 5, 5, 6, 5, 4, 5, 5, 5, 6, 5, 5, 6, 5}}, 
 {10, {4, 5, 5, 5, 4, 5, 5, 5, 6, 5, 5, 6, 5, 4, 5, 6, 5, 5}}, 
 {10, {4, 5, 5, 4, 5, 5, 6, 5, 5, 5, 6, 5, 4, 5, 5, 5, 6, 5}}, 
 {10, {4, 5, 5, 4, 5, 5, 6, 5, 5, 5, 5, 5, 6, 5, 4, 5, 6, 5}}, 
 {10, {4, 5, 5, 4, 5, 5, 6, 5, 4, 5, 5, 5, 5, 6, 5, 5, 6, 5}}, 
 {10, {4, 5, 5, 4, 5, 5, 5, 6, 5, 5, 6, 5, 5, 4, 5, 6, 5, 5}}, 
 {10, {4, 5, 5, 4, 5, 5, 5, 6, 5, 5, 5, 6, 5, 4, 5, 6, 5, 5}}, 
 {10, {4, 5, 5, 4, 5, 5, 5, 6, 5, 5, 5, 5, 6, 5, 4, 5, 6, 5}}, 
 {10, {4, 5, 5, 4, 5, 5, 5, 5, 6, 5, 5, 6, 5, 4, 5, 6, 5, 5}}, 
 {10, {4, 5, 5, 4, 5, 5, 5, 4, 5, 5, 5, 4, 5, 6, 6, 6, 6, 5}}, 
 {10, {4, 5, 5, 4, 5, 5, 4, 5, 5, 5, 5, 5, 6, 5, 5, 6, 6, 5}}, 
 {10, {4, 5, 4, 5, 5, 6, 5, 5, 5, 5, 5, 5, 5, 5, 5, 6, 5, 5}}, 
 {10, {4, 5, 4, 5, 5, 5, 6, 6, 5, 5, 4, 5, 5, 5, 5, 6, 5, 5}}, 
 {10, {4, 5, 4, 5, 5, 5, 6, 5, 5, 5, 5, 5, 5, 5, 5, 5, 6, 5}}, 
 {10, {4, 5, 4, 5, 5, 5, 5, 5, 5, 6, 6, 5, 5, 4, 5, 6, 5, 5}}, 
 {10, {4, 5, 4, 5, 5, 5, 5, 5, 5, 5, 5, 6, 5, 5, 6, 5, 5, 5}}, 
 {10, {4, 5, 4, 5, 5, 5, 5, 5, 5, 4, 5, 6, 5, 5, 6, 6, 5, 5}}, 
 {10, {4, 5, 4, 5, 5, 5, 5, 5, 4, 5, 6, 6, 6, 5, 5, 5, 5, 5}}, 
 {10, {4, 5, 4, 5, 5, 5, 5, 4, 5, 5, 5, 5, 5, 5, 6, 6, 6, 5}}, 
 {10, {4, 5, 4, 5, 5, 5, 4, 5, 6, 6, 6, 5, 5, 5, 5, 5, 5, 5}}, 
 {10, {4, 4, 5, 6, 5, 5, 5, 5, 4, 5, 5, 5, 4, 5, 6, 6, 6, 5}}, 
 {10, {4, 4, 5, 6, 5, 5, 4, 5, 5, 5, 6, 5, 5, 4, 5, 6, 6, 5}}, 
 {10, {4, 4, 5, 6, 5, 5, 4, 5, 5, 4, 5, 5, 5, 5, 6, 6, 6, 5}}, 
 {10, {4, 4, 5, 5, 6, 6, 6, 5, 4, 5, 5, 4, 5, 5, 5, 5, 6, 5}}, 
 {10, {4, 4, 5, 5, 6, 5, 5, 5, 6, 5, 4, 4, 5, 5, 6, 6, 5, 5}}, 
 {10, {4, 4, 5, 5, 6, 5, 5, 5, 5, 5, 4, 5, 6, 5, 5, 6, 5, 5}}, 
 {10, {4, 4, 5, 5, 6, 5, 5, 5, 5, 4, 5, 5, 6, 5, 5, 5, 6, 5}}, 
 {10, {4, 4, 5, 5, 6, 5, 5, 5, 4, 5, 5, 6, 5, 5, 5, 6, 5, 5}}, 
 {10, {4, 4, 5, 5, 6, 5, 4, 4, 5, 6, 6, 5, 5, 5, 5, 5, 6, 5}}, 
 {10, {4, 4, 5, 5, 5, 6, 6, 6, 5, 5, 4, 5, 5, 5, 4, 5, 6, 5}}, 
 {10, {4, 4, 5, 5, 5, 6, 6, 5, 5, 5, 5, 6, 5, 4, 4, 5, 6, 5}}, 
 {10, {4, 4, 5, 5, 5, 6, 6, 5, 5, 5, 4, 5, 5, 4, 5, 6, 6, 5}}, 
 {10, {4, 4, 5, 5, 5, 6, 6, 5, 4, 5, 5, 5, 4, 5, 6, 6, 5, 5}}, 
 {10, {4, 4, 5, 5, 5, 6, 5, 5, 6, 6, 5, 5, 5, 4, 4, 5, 6, 5}}, 
 {10, {4, 4, 5, 5, 5, 6, 5, 5, 6, 5, 5, 5, 5, 5, 4, 5, 6, 5}}, 
 {10, {4, 4, 5, 5, 5, 6, 5, 5, 6, 5, 5, 5, 5, 4, 5, 5, 6, 5}}, 
 {10, {4, 4, 5, 5, 5, 6, 5, 5, 6, 5, 4, 5, 5, 5, 5, 6, 5, 5}}, 
 {10, {4, 4, 5, 5, 5, 6, 5, 4, 5, 5, 6, 6, 5, 4, 5, 5, 6, 5}}, 
 {10, {4, 4, 5, 5, 5, 6, 5, 4, 5, 5, 4, 5, 6, 6, 6, 5, 5, 5}}, 
 {10, {4, 4, 5, 5, 5, 5, 6, 6, 5, 5, 6, 5, 5, 4, 4, 5, 6, 5}}, 
 {10, {4, 4, 5, 5, 5, 5, 6, 6, 5, 4, 4, 5, 5, 6, 5, 5, 6, 5}}, 
 {10, {4, 4, 5, 5, 5, 5, 6, 5, 5, 5, 6, 5, 5, 4, 5, 5, 6, 5}}, 
 {10, {4, 4, 5, 5, 5, 5, 6, 5, 5, 4, 5, 6, 5, 5, 6, 5, 5, 5}}, 
 {10, {4, 4, 5, 5, 5, 5, 5, 6, 5, 5, 5, 6, 5, 4, 5, 6, 5, 5}}, 
 {10, {4, 4, 5, 5, 5, 5, 5, 6, 5, 5, 4, 5, 5, 6, 5, 5, 6, 5}}, 
 {10, {4, 4, 5, 5, 5, 5, 5, 4, 5, 6, 5, 4, 5, 6, 6, 6, 5, 5}}, 
 {10, {4, 4, 5, 5, 5, 5, 5, 4, 4, 5, 5, 6, 6, 5, 6, 6, 5, 5}}, 
 {10, {4, 4, 5, 5, 5, 5, 4, 5, 6, 6, 6, 5, 4, 5, 5, 5, 6, 5}}, 
 {10, {4, 4, 5, 5, 5, 5, 4, 5, 5, 5, 6, 5, 4, 5, 6, 6, 6, 5}}, 
 {10, {4, 4, 5, 5, 5, 4, 5, 6, 6, 5, 4, 5, 6, 6, 5, 5, 5, 5}}, 
 {10, {4, 4, 5, 5, 5, 4, 5, 6, 6, 5, 4, 5, 5, 5, 6, 6, 5, 5}}, 
 {10, {4, 4, 5, 5, 5, 4, 5, 6, 5, 5, 4, 5, 5, 5, 6, 6, 6, 5}}, 
 {10, {4, 4, 5, 5, 5, 4, 5, 5, 6, 6, 5, 6, 6, 5, 4, 5, 5, 5}}, 
 {10, {4, 4, 5, 5, 5, 4, 5, 5, 5, 6, 6, 5, 4, 5, 6, 6, 5, 5}}, 
 {10, {4, 4, 5, 5, 5, 4, 5, 5, 5, 4, 5, 5, 6, 6, 5, 6, 6, 5}}, 
 {10, {4, 4, 5, 5, 5, 4, 5, 5, 4, 5, 6, 6, 5, 5, 6, 6, 5, 5}}, 
 {10, {4, 4, 5, 5, 5, 4, 5, 5, 4, 5, 6, 5, 5, 5, 6, 6, 6, 5}}, 
 {10, {4, 4, 5, 5, 5, 4, 4, 5, 5, 6, 6, 6, 6, 5, 4, 5, 6, 5}}, 
 {10, {4, 4, 5, 5, 4, 5, 6, 6, 5, 6, 6, 5, 5, 4, 5, 5, 5, 5}}, 
 {10, {4, 4, 5, 5, 4, 5, 6, 6, 5, 5, 6, 6, 5, 5, 4, 5, 5, 5}}, 
 {10, {4, 4, 5, 5, 4, 5, 6, 6, 5, 5, 5, 4, 5, 5, 5, 6, 6, 5}}, 
 {10, {4, 4, 5, 5, 4, 5, 6, 5, 5, 6, 5, 4, 5, 5, 6, 6, 5, 5}}, 
 {10, {4, 4, 5, 5, 4, 5, 6, 5, 5, 4, 5, 5, 5, 5, 6, 6, 6, 5}}, 
 {10, {4, 4, 5, 5, 4, 5, 6, 5, 4, 5, 5, 6, 6, 6, 5, 5, 5, 5}}, 
 {10, {4, 4, 5, 5, 4, 5, 5, 6, 6, 5, 4, 5, 5, 5, 5, 6, 6, 5}}, 
 {10, {4, 4, 5, 5, 4, 5, 5, 6, 5, 5, 5, 4, 5, 6, 6, 6, 5, 5}}, 
 {10, {4, 4, 5, 5, 4, 5, 5, 6, 5, 4, 5, 6, 6, 6, 5, 5, 5, 5}}, 
 {10, {4, 4, 5, 5, 4, 5, 5, 5, 4, 5, 6, 6, 5, 5, 5, 6, 6, 5}}, 
 {10, {4, 4, 5, 5, 4, 5, 5, 5, 4, 4, 5, 6, 6, 6, 6, 6, 5, 5}}, 
 {10, {4, 4, 5, 5, 4, 5, 5, 4, 5, 6, 6, 6, 5, 5, 5, 5, 6, 5}}, 
 {10, {4, 4, 5, 5, 4, 5, 5, 4, 5, 6, 5, 5, 6, 6, 6, 5, 5, 5}}, 
 {10, {4, 4, 5, 5, 4, 5, 5, 4, 5, 6, 5, 5, 5, 5, 6, 6, 6, 5}}, 
 {10, {4, 4, 5, 5, 4, 5, 5, 4, 5, 5, 6, 5, 5, 6, 6, 6, 5, 5}}, 
 {10, {4, 4, 5, 5, 4, 5, 5, 4, 4, 5, 6, 5, 5, 6, 6, 6, 6, 5}}, 
 {10, {4, 4, 5, 5, 4, 4, 5, 5, 5, 6, 6, 5, 5, 6, 5, 5, 6, 5}}, 
 {10, {4, 4, 5, 5, 4, 4, 5, 5, 4, 5, 5, 5, 6, 6, 6, 6, 6, 5}}, 
 {10, {4, 4, 5, 4, 5, 6, 6, 5, 5, 5, 4, 5, 5, 5, 6, 6, 5, 5}}, 
 {10, {4, 4, 5, 4, 5, 6, 5, 5, 6, 5, 5, 5, 5, 5, 5, 5, 6, 5}}, 
 {10, {4, 4, 5, 4, 5, 6, 5, 5, 5, 5, 6, 6, 6, 5, 5, 4, 5, 5}}, 
 {10, {4, 4, 5, 4, 5, 6, 5, 5, 5, 5, 6, 5, 5, 5, 5, 6, 5, 5}}, 
 {10, {4, 4, 5, 4, 5, 6, 5, 5, 5, 5, 5, 5, 6, 5, 5, 5, 6, 5}}, 
 {10, {4, 4, 5, 4, 5, 6, 5, 5, 5, 5, 5, 4, 5, 5, 6, 6, 6, 5}}, 
 {10, {4, 4, 5, 4, 5, 5, 6, 6, 6, 6, 6, 5, 5, 4, 4, 5, 5, 5}}, 
 {10, {4, 4, 5, 4, 5, 5, 6, 6, 6, 5, 5, 5, 6, 5, 5, 4, 5, 5}}, 
 {10, {4, 4, 5, 4, 5, 5, 6, 6, 6, 5, 5, 5, 4, 5, 5, 6, 5, 5}}, 
 {10, {4, 4, 5, 4, 5, 5, 6, 6, 5, 5, 5, 6, 6, 5, 4, 5, 5, 5}}, 
 {10, {4, 4, 5, 4, 5, 5, 6, 6, 5, 5, 5, 5, 6, 6, 5, 4, 5, 5}}, 
 {10, {4, 4, 5, 4, 5, 5, 6, 5, 5, 5, 4, 5, 5, 6, 6, 6, 5, 5}}, 
 {10, {4, 4, 5, 4, 5, 5, 5, 6, 6, 5, 5, 6, 6, 5, 5, 4, 5, 5}}, 
 {10, {4, 4, 5, 4, 5, 5, 5, 6, 6, 5, 5, 4, 5, 6, 6, 5, 5, 5}}, 
 {10, {4, 4, 5, 4, 5, 5, 5, 6, 5, 5, 6, 5, 5, 5, 6, 5, 5, 5}}, 
 {10, {4, 4, 5, 4, 5, 5, 5, 6, 5, 5, 6, 5, 5, 5, 5, 6, 5, 5}}, 
 {10, {4, 4, 5, 4, 5, 5, 5, 6, 5, 5, 5, 6, 5, 5, 5, 5, 6, 5}}, 
 {10, {4, 4, 5, 4, 5, 5, 5, 6, 5, 4, 5, 5, 5, 5, 6, 6, 6, 5}}, 
 {10, {4, 4, 5, 4, 5, 5, 5, 5, 6, 5, 5, 6, 5, 5, 6, 5, 5, 5}}, 
 {10, {4, 4, 5, 4, 5, 5, 5, 5, 6, 5, 5, 6, 5, 5, 5, 6, 5, 5}}, 
 {10, {4, 4, 5, 4, 5, 5, 5, 5, 5, 6, 6, 6, 5, 4, 5, 6, 5, 5}}, 
 {10, {4, 4, 5, 4, 5, 5, 5, 5, 5, 6, 6, 5, 5, 4, 5, 6, 6, 5}}, 
 {10, {4, 4, 5, 4, 5, 5, 5, 5, 4, 5, 6, 6, 5, 5, 6, 6, 5, 5}}, 
 {10, {4, 4, 5, 4, 5, 5, 5, 5, 4, 5, 6, 5, 5, 5, 6, 6, 6, 5}}, 
 {10, {4, 4, 5, 4, 5, 5, 5, 5, 4, 5, 5, 6, 6, 6, 5, 5, 6, 5}}, 
 {10, {4, 4, 5, 4, 5, 5, 5, 4, 5, 6, 5, 5, 5, 6, 6, 6, 5, 5}}, 
 {10, {4, 4, 5, 4, 5, 5, 5, 4, 5, 5, 5, 5, 6, 6, 5, 6, 6, 5}}, 
 {10, {4, 4, 5, 4, 4, 5, 5, 6, 6, 5, 5, 5, 5, 5, 6, 6, 5, 5}}, 
 {10, {4, 4, 5, 4, 4, 5, 5, 5, 6, 6, 6, 6, 5, 5, 4, 5, 6, 5}}, 
 {10, {4, 4, 5, 4, 4, 5, 5, 5, 5, 5, 6, 5, 5, 5, 6, 6, 6, 5}}, 
 {10, {4, 4, 5, 4, 4, 5, 5, 5, 5, 5, 5, 6, 6, 5, 6, 6, 5, 5}},
```
