## Supplementary material for "On the structure of multiple stable equilibria in competitive ecological systems": Mathematica+Julia code and examples: basins.m

```
(* Code to generate and analyze periodic niche axis models *)

(* this part originally from  unstable.m *)

(* this is the interaction we use *)

f[3, x_] := Exp[- x^4]

total[x_, i_, a_, n_] := Sum[f [ i, x - j a], {j, - n, n}]

(* ok, let's look at a system of n interacting things on a circle
 of fixed size *)

potential[d_, size_, k_:10, i_:3] := N[total[d, i, size, k]]


(* matrix of species with niche overlap interaction *)
(* size = S/\Delta *)
(* example matrix[30, 7.5]  gives S= 30, \Delta = 4 system from paper *)

(* ------------------generate interaction matrix--------- *)

matrix[n_, size_] :=
 Table[potential[Min[Abs[i-j], n -Abs[i-j]]*(size/n), size],
        {i,0, n -1},{j, 0, n -1}]

(* assume spacing 0.25 (\Delta = 4)*)

mx[n_] := mx[n] = matrix[n, n/4.0]


(* analyze eigenvectors and eigenvalues *)

vector[m_, n_] :=Table[E^(2 Pi I k m/n),{k, 0, n -1}]

eigenvalues[n_, size_] :=
 With[{mm = matrix[n, size]},
  Table[Re [(mm.vector[m,n])[[1]]],{ m, 0, n-1}]]

(* given some general positions of spikes, give potential *)

general[list_, weights_, length_, x_] :=
 Sum[f[3, (x-list[[i]])/length]*weights[[i]],{i, 1,Length[list]}]

(* code originally from multiple.m *)


(* first let's identify which subsets have good equilibria *)

subset[m_, set_] :=
 Table[m[[set[[i]],  set[[j]]]],{ i, 1,Length[set]},{j, 1,Length[set]}]

(* is there a good equilibrium? *)

positive[list_] :=Apply[And, Map[#> 0 &, list]]

(* finds stable equilibria *)

(* need to check against invasion *)

replace[size_, set_, numbers_, position_:1]:=
 If[size < position,{},
  If[And[Length[set] > 0, set[[1]] == position], 
        Join[{numbers[[1]]}, replace[size,Drop[set, 1], Drop[numbers, 1],
          position + 1]],
        Join[{0}, replace[size, set, numbers, position + 1]]]]

(* checks if stable to invasion *)
(* returns True if stable, also vector with info on directions *)

invasion[m_, set_] :=
 Block[{ss= subset[m, set], l =Length[set], lm =Length[m], pp, pa, r0, u},
  pp = Inverse[ss].Table[1,{l}];
  pa = replace[lm, set, pp];
(*   Print["test", m, pa]; *)
  r0 = Table[1,{lm}] -m.pa;
(*   Print[r0]; *)
  u =Table[Or[MemberQ[set,  i], r0[[i]] < 0], {i, 1, lm}];
(*   Print[u]; *)
  {Apply[And, u], u}]

(* key function stable: determines if system described by interaction
matrix m has a stable subset equilibrium given by set *)

stable[m_, set_] :=
 With[{ss= subset[m, set], l =Length[set]},
  And[positive[Inverse[ss].Table[1,{l}]],
      positive[Eigensystem[ss][[1]]],
      invasion[m, set][[1]]]]

(* unstable equilibria, but stable to invasion *)

unstable[m_, set_] :=
 With[{ss= subset[m, set], l =Length[set]},
  And[positive[Inverse[ss].Table[1,{l}]],
      Not[positive[Eigensystem[ss][[1]]]],
      invasion[m, set][[1]]]]

(* ----------------------code to find stable equilibria----------- *)

(* stable[m] finds all stable subset equilibria *)

stable[m_] :=
 Select[Drop[Subsets[Table[i,{i, 1, Length[m]}]], 1], stable[m, #]&]

(*  finds stable equilibria only checking all subsets for first k, then including specific positions *)

stable[m_, k_, extras_] :=
  Select[Map[Join[#, extras]&,Drop[Subsets[Table[i,{i, 1, k}]], 1]], 
        stable[m, #]&]

unstable[m_] :=
 Select[Drop[Subsets[Table[i,{i, 1, Length[m]}]], 1], unstable[m, #]&]


(* analyzing equilibria *)

Lyapunov[m_, v_] := -2 Apply[Plus, v] +v.m.v

(* -----------returns Lyapunov function, n*'s for given set----------- *)

energy[m_, set_] :=
 With[{ss= subset[m, set], l =Length[set]},
  With[{v= Inverse[ss].Table[1,{l}]},
   {Lyapunov[ss, v], v}]]

(* some useful tools for analyzing solutions *)
(* from analysis.m *)

loop[list_, n_, offset_: 200] :=
 Join[list, offset +Take[list, n]]

differences[list_] :=
 Table[list[[i + 1]]-list[[i]],{ i, 1,Length[list] -1}]

(* getting complete set, from complete.m *)

(* positions to gaps *)

gaps[all_, n_] := Map[differences[loop[#, 1, n]] &, all]

(* goes from gaps to positions *)

configuration[gaps_] :=
 Table[1 +Sum[gaps[[i]],{i, 1, k}],{ k, 0,Length[gaps] -1}]

(* gets canonical order for a given gap configuration including reversal *)

canonical[list_] :=
 First[Sort[Table[RotateLeft[list, k],{k, 0,Length[list] -1}]]]

canonical[list_, 1] :=
 First[Sort[{canonical[Reverse[list]], canonical[list]}]]

(* extracts minimal canonical representatives from list *)

extract[list_] := Union[Map[canonical, list]]

sequences[n_] := extract[gaps[stable[mx[n]], n]]

(* only loops over first k, with fixed additional stuff *)

sequences[n_, k_, extras_] := extract[gaps[stable[mx[n], k, extras], n]]

systematic[n_:30] :=
 Block[{s},
     For[j = 1, j <= n, j += 1,
    s = sequences[j];
    Print[j, ":", Length[s]];
    {j, s} >>> "systematic-sequences.m"]]

(* from analysis.m: check for allowed results on simulation data *)


(* need to use a matrix m and types and hits from a julia run set
  (e.g. m = matrix[60, 15.0]; << r10-output.m) *)


allowed[mx_, t_, h_] :=
 Map[{t[[#]], h[[#]]}&, Select[Table[i,{i, 1,Length[t]}],
                stable[mx, configuration[t[[#]]]]&]]

sortedresults[r_] := Reverse[Sort[Map[Reverse, r]]]
```
