## Supplementary material for "On the structure of multiple stable equilibria in competitive ecological systems": Mathematica+Julia code and examples: combine.jl

```
# extracted from analysis.jl, 11/9/24

# code to take two lists, 1 of solutions and 1 of instances, add new result

#function augment(list::Vector{Any}, numbers::Vector{Any}, new::Vector{Any}, number)
function augment(list::Vector{Vector{Int64}}, numbers::Vector{Int64}, new::Vector{Int64}, number::Int64)
#  print(new,number)
  for i in 1:length(list)
    if list[i] == new
      numbers[i] += number
      return i
    end
  end
  push!(list, new)
  push!(numbers, number)
end

#assumes information is in canonical order of gaps
#modified to only use valid indices 11/28/23
function extraction(i1, i2, size) #extract information from indices i1--i2
    o = open(string("combined"), "a")
#    t = Vector{Any}(undef, 0)
#    h = Vector{Any}(undef, 0)
    t = Vector{Vector{Int64}}(undef, 0)
    h = Vector{Int64}(undef, 0)
    for k in i1:i2
    	if isassigned(types, k)
            tt = types[k]
	    hh = hits[k]
	    for l in 1:length(tt)
#	        print("k =", k, "l =", l, "\n")
#avoids bad answers
	        if length(tt[l]) > 0
	            augment(t, h, tt[l], hh[l])
	        end
	    end
	end
    end
    write(o, "types =" * string(t, "\n"))
    write(o, "hits =" * string(h, "\n"))
    close(o)
end
```
