## Supplementary material for "On the structure of multiple stable equilibria in competitive ecological systems": Mathematica+Julia code and examples: combined.m

```
(* this has the results of 200 runs at size 30, in mathematica format
  for processing by model.m *)


types ={{4, 5, 5, 6, 5, 5}, {6, 6, 6, 6, 6}, {4, 5, 5, 5, 6, 5}, {5, 5, 5, 5, 5, 5}, {3, 5, 5, 6, 6, 5}, {4, 4, 5, 6, 6, 5}, {1, 6, 6, 5, 6, 6}}
hits ={19, 57, 37, 50, 14, 11, 12}
```
