## Supplementary material for "On the structure of multiple stable equilibria in competitive ecological systems": Mathematica+Julia code and examples: examples-2.txt

# This demonstrates use of julia code to produce files with simulation
# results.  Combines 2*100 simulations at size S= 30, \Delta = 4

# use Julia code to produce file "output" with two sets of simulations

% julia niches.jl 30 7.5 100 1
% julia niches.jl 30 7.5 100 2

# if using a cluster to produce multiple files, combine with cat

# format of output file:
# matrix[i] = interaction matrix
# parameters[i] = parameters given to program
# types[i] = canonical form of solutions reached
# hits[i] = number of times each solution is reached

# now add header to file for processing/combining multiple simulation sets

% cat header-10.jl output > results-200.jl

# now use combine.jl to combine results

% julia
               _
   _       _ _(_)_     |  Documentation: https://docs.julialang.org
  (_)     | (_) (_)    |
   _ _   _| |_  __ _   |  Type "?" for help, "]?" for Pkg help.
  | | | | | | |/ _` |  |
  | | |_| | | | (_| |  |  Version 1.10.4 (2024-06-04)
 _/ |\__'_|_|_|\__'_|  |  Official https://julialang.org/ release
|__/                   |

┌ Warning: Terminal not fully functional
└ @ Base client.jl:422
julia> include("results-200.jl")
7-element Vector{Int64}:
 11
 30
 18
 22
  8
  7
  4

julia> include("combine.jl")
extraction (generic function with 1 method)

# this combines data from runs 1--2 for size 30.

julia> extraction(1, 2, 30)

julia> exit()

# at the end we have a file combined, containing all the simulation
# results organized by type and number of hits.
