## Supplementary material for "On the structure of multiple stable equilibria in competitive ecological systems": Mathematica+Julia code and examples: header-10.jl

```
# header file for up to 10 sets of simulation results
matrix = Vector{Any}(undef, 10)
parameters = Vector{Any}(undef, 10)
types = Vector{Vector{Vector{Int64}}}(undef, 10)
hits = Vector{Vector{Int64}}(undef, 10)
```
