## Supplementary material for "On the structure of multiple stable equilibria in competitive ecological systems": Mathematica+Julia code and examples: m4-90.m

```
(* m4, w4 from 10^6 simulations at size S=90 *)
(* m4 =model[90, 4] *)

m4 =
{{{{5}, 0.5470107926600054}, {{6}, 0.38462241705021505}, 
  {{4}, 0.05649095803214271}, {{1}, 0.005149219698169079}, 
  {{3}, 0.006726612559467657}}, {{{5, 5}, 0.3596491772376277}, 
  {{5, 6}, 0.13365378779645629}, {{6, 6}, 0.2458194095555897}, 
  {{6, 5}, 0.13365378779645629}, {{4, 5}, 0.04698121506645385}, 
  {{5, 4}, 0.04698121506645385}, {{1, 6}, 0.005149219698169079}, 
  {{6, 1}, 0.005149219698169079}, {{4, 4}, 0.009509742965688869}, 
  {{3, 5}, 0.006726612559467657}, {{5, 3}, 0.006726612559467657}}, 
 {{{5, 5, 5}, 0.22123023523849084}, {{5, 5, 6}, 0.10205023250470567}, 
  {{5, 6, 6}, 0.09781363374003854}, {{6, 6, 6}, 0.14291497260334768}, 
  {{6, 6, 5}, 0.09781363374003854}, {{6, 5, 5}, 0.10205023250470567}, 
  {{6, 5, 6}, 0.015503386708331774}, {{5, 6, 5}, 0.03583982902791717}, 
  {{4, 5, 5}, 0.032398840936547754}, {{5, 5, 4}, 0.032398840936547754}, 
  {{5, 4, 5}, 0.03755964346855523}, {{6, 5, 4}, 0.013375306922935683}, 
  {{4, 5, 6}, 0.013375306922935683}, {{1, 6, 6}, 0.0050908032122034845}, 
  {{6, 6, 1}, 0.0050908032122034845}, {{6, 1, 6}, 0.005149219698169079}, 
  {{4, 4, 5}, 0.009421571597898613}, {{5, 4, 4}, 0.009421571597898613}, 
  {{3, 5, 5}, 0.003969868557883409}, {{6, 5, 3}, 0.002724861660483147}, 
  {{5, 3, 5}, 0.006726612559467657}, {{5, 5, 3}, 0.003969868557883409}, 
  {{3, 5, 6}, 0.002724861660483147}, {{4, 5, 4}, 0.0011751848658693067}, 
  {{1, 6, 1}, 0.00005809145746502708}, {{4, 4, 4}, 0.00008817136779025473}, 
  {{3, 5, 4}, 0.00003188234110110082}, {{4, 5, 3}, 0.00003188234110110082}, 
  {{1, 6, 5}, 3.250285005672929*^-7}, {{5, 6, 1}, 3.250285005672929*^-7}}, 
 {{{5, 5, 5, 5}, 0.13472017675877204}, {{5, 5, 5, 6}, 0.06524022295064064}, 
  {{5, 5, 6, 6}, 0.07226532986581109}, {{5, 6, 6, 6}, 0.059025884856112536}, 
  {{6, 6, 6, 6}, 0.08064991280667251}, {{6, 6, 6, 5}, 0.059025884856112536}, 
  {{6, 6, 5, 5}, 0.07226532986581109}, {{6, 5, 5, 5}, 0.06524022295064064}, 
  {{5, 6, 6, 5}, 0.03709709836420247}, {{6, 5, 5, 6}, 0.02482668150687704}, 
  {{6, 6, 5, 6}, 0.015503386708331774}, {{6, 5, 6, 6}, 0.015503386708331774}, 
  {{5, 5, 6, 5}, 0.029784902638894575}, {{5, 6, 5, 5}, 0.029784902638894575}, 
  {{4, 5, 5, 5}, 0.018949486611573745}, {{6, 5, 5, 4}, 0.010744733075662557}, 
  {{5, 5, 4, 5}, 0.02624135328157343}, {{5, 4, 5, 5}, 0.02624135328157343}, 
  {{5, 5, 5, 4}, 0.018949486611573745}, {{4, 5, 5, 6}, 0.010744733075662557}, 
  {{6, 6, 5, 4}, 0.008440221910549437}, {{6, 5, 4, 5}, 0.01062491575113525}, 
  {{5, 4, 5, 6}, 0.01062491575113525}, {{4, 5, 6, 6}, 0.008440221910549437}, 
  {{5, 6, 5, 4}, 0.004935085012386245}, {{4, 5, 6, 5}, 0.004935085012386245}, 
  {{1, 6, 6, 6}, 0.0032391749405626287}, 
  {{6, 6, 6, 1}, 0.0032391749405626287}, 
  {{6, 6, 1, 6}, 0.0050908032122034845}, 
  {{6, 1, 6, 6}, 0.0050908032122034845}, 
  {{4, 4, 5, 5}, 0.006157487654974327}, {{5, 5, 4, 4}, 0.006157487654974327}, 
  {{5, 4, 4, 5}, 0.009377899586640571}, {{1, 6, 6, 5}, 
   0.0016906505197235272}, {{5, 6, 6, 1}, 0.0016906505197235272}, 
  {{6, 5, 4, 4}, 0.0027503911718004324}, 
  {{4, 4, 5, 6}, 0.0027503911718004324}, 
  {{4, 5, 5, 4}, 0.0023559247642937634}, 
  {{3, 5, 5, 6}, 0.001238594971525435}, {{6, 6, 5, 3}, 
   0.0016046952553462313}, {{6, 5, 3, 5}, 0.002724861660483147}, 
  {{5, 3, 5, 5}, 0.003969868557883409}, {{6, 5, 5, 3}, 0.001238594971525435}, 
  {{5, 5, 3, 5}, 0.003969868557883409}, {{5, 3, 5, 6}, 0.002724861660483147}, 
  {{3, 5, 6, 6}, 0.0016046952553462313}, 
  {{3, 5, 5, 5}, 0.002320348917504398}, {{5, 5, 5, 3}, 0.002320348917504398}, 
  {{5, 6, 5, 3}, 0.0011198413766363485}, 
  {{3, 5, 6, 5}, 0.0011198413766363485}, 
  {{1, 6, 6, 1}, 0.00016097775191732834}, 
  {{4, 4, 5, 4}, 0.00048181043002275254}, 
  {{4, 5, 4, 5}, 0.0006933744358465541}, 
  {{5, 4, 5, 4}, 0.0006933744358465541}, 
  {{4, 5, 4, 4}, 0.00048181043002275254}, 
  {{3, 5, 5, 4}, 0.00034869648501769306}, 
  {{4, 5, 5, 3}, 0.00034869648501769306}, 
  {{1, 6, 1, 6}, 0.00005809145746502708}, 
  {{6, 1, 6, 1}, 0.00005809145746502708}, 
  {{3, 5, 5, 3}, 0.00006222818383588353}, 
  {{4, 4, 4, 4}, 0.00004449935653221301}, 
  {{4, 4, 4, 5}, 0.00004367201125804172}, 
  {{5, 4, 4, 4}, 0.00004367201125804172}, 
  {{3, 5, 4, 4}, 0.00003188234110110082}, 
  {{5, 3, 5, 4}, 0.00003188234110110082}, 
  {{4, 4, 5, 3}, 0.00003188234110110082}, 
  {{4, 5, 3, 5}, 0.00003188234110110082}, 
  {{1, 6, 5, 3}, 3.250285005672929*^-7}, 
  {{6, 1, 6, 5}, 3.250285005672929*^-7}, 
  {{3, 5, 6, 1}, 3.250285005672929*^-7}, 
  {{5, 6, 1, 6}, 3.250285005672929*^-7}}}

w4 =
{{{{5}, 0.5470107926600054}, {{6}, 0.38462241705021505}, 
  {{4}, 0.05649095803214271}, {{1}, 0.005149219698169079}, 
  {{3}, 0.006726612559467657}}, {{{5, 5}, 1.2019524327171383}, 
  {{5, 6}, 0.635258936839385}, {{6, 6}, 1.6616784426844013}, 
  {{6, 5}, 0.635258936839385}, {{4, 5}, 1.5203704130692115}, 
  {{5, 4}, 1.5203704130692115}, {{1, 6}, 2.5999524616097536}, 
  {{6, 1}, 2.5999524616097536}, {{4, 4}, 2.9799631848057304}, 
  {{3, 5}, 1.8281174949715298}, {{5, 3}, 1.8281174949715298}}, 
 {{{5, 5, 5}, 0.9355828108566409}, {{5, 5, 6}, 1.161313704825896}, 
  {{5, 6, 6}, 1.1450818824951712}, {{6, 6, 6}, 0.9096618487260919}, 
  {{6, 6, 5}, 1.1450818824951712}, {{6, 5, 5}, 1.161313704825896}, 
  {{6, 5, 6}, 0.4747444895298451}, {{5, 6, 5}, 0.7716813054950212}, 
  {{4, 5, 5}, 1.0488709400631813}, {{5, 5, 4}, 1.0488709400631813}, 
  {{5, 4, 5}, 0.9612844382597505}, {{6, 5, 4}, 1.1651829894460028}, 
  {{4, 5, 6}, 1.1651829894460028}, {{1, 6, 6}, 1.546903809790235}, 
  {{6, 6, 1}, 1.546903809790235}, {{6, 1, 6}, 1.0000000000000004}, 
  {{4, 4, 5}, 1.1912674337360984}, {{5, 4, 4}, 1.1912674337360984}, 
  {{3, 5, 5}, 0.8976283028625377}, {{6, 5, 3}, 1.6579165554583384}, 
  {{5, 3, 5}, 1.}, {{5, 5, 3}, 0.8976283028625377}, 
  {{3, 5, 6}, 1.657916555458338}, {{4, 5, 4}, 0.29124172572985324}, 
  {{1, 6, 1}, 0.8426825902080946}, {{4, 4, 4}, 0.055076833097830645}, 
  {{3, 5, 4}, 0.055185558295115594}, {{4, 5, 3}, 0.055185558295115594}, 
  {{1, 6, 5}, 0.00018164913904733916}, {{5, 6, 1}, 0.00018164913904733916}}, 
 {{{5, 5, 5, 5}, 0.9899716386320482}, {{5, 5, 5, 6}, 1.0392882756454294}, 
  {{5, 5, 6, 6}, 0.9676044770356874}, {{5, 6, 6, 6}, 1.0379622385719358}, 
  {{6, 6, 6, 6}, 0.9706543665951987}, {{6, 6, 6, 5}, 1.0379622385719356}, 
  {{6, 6, 5, 5}, 0.9676044770356874}, {{6, 5, 5, 5}, 1.0392882756454294}, 
  {{5, 6, 6, 5}, 0.9531413844929948}, {{6, 5, 5, 6}, 0.8573728897137426}, 
  {{6, 6, 5, 6}, 1.3664126634091824}, {{6, 5, 6, 6}, 1.3664126634091822}, 
  {{5, 5, 6, 5}, 1.0884225436477488}, {{5, 6, 5, 5}, 1.0884225436477488}, 
  {{4, 5, 5, 5}, 0.9508293186726413}, {{6, 5, 5, 4}, 1.168775851734088}, 
  {{5, 5, 4, 5}, 1.0131168487088058}, {{5, 4, 5, 5}, 1.0131168487088058}, 
  {{5, 5, 5, 4}, 0.9508293186726413}, {{4, 5, 5, 6}, 1.168775851734088}, 
  {{6, 6, 5, 4}, 0.8622475855699548}, {{6, 5, 4, 5}, 0.9936296034129525}, 
  {{5, 4, 5, 6}, 0.9936296034129525}, {{4, 5, 6, 6}, 0.8622475855699547}, 
  {{5, 6, 5, 4}, 1.3759613147113865}, {{4, 5, 6, 5}, 1.3759613147113865}, 
  {{1, 6, 6, 6}, 1.0944263359651087}, {{6, 6, 6, 1}, 1.0944263359651087}, 
  {{6, 6, 1, 6}, 0.9999999999999996}, {{6, 1, 6, 6}, 0.9999999999999996}, 
  {{4, 4, 5, 5}, 0.9477089192801317}, {{5, 5, 4, 4}, 0.9477089192801317}, 
  {{5, 4, 4, 5}, 1.0046797557559515}, {{1, 6, 6, 5}, 0.8346113971398059}, 
  {{5, 6, 6, 1}, 0.8346113971398059}, {{6, 5, 4, 4}, 1.025395956722993}, 
  {{4, 4, 5, 6}, 1.025395956722993}, {{4, 5, 5, 4}, 0.8072007208349868}, 
  {{3, 5, 5, 6}, 1.0995582909126513}, {{6, 6, 5, 3}, 0.8046925646232388}, 
  {{6, 5, 3, 5}, 1.}, {{5, 3, 5, 5}, 1.}, {{6, 5, 5, 3}, 1.0995582909126513}, 
  {{5, 5, 3, 5}, 1.0000000000000002}, {{5, 3, 5, 6}, 1.0000000000000002}, 
  {{3, 5, 6, 6}, 0.804692564623239}, {{3, 5, 5, 5}, 0.9501928504970153}, 
  {{5, 5, 5, 3}, 0.9501928504970152}, {{5, 6, 5, 3}, 1.5325949629123656}, 
  {{3, 5, 6, 5}, 1.5325949629123663}, {{1, 6, 6, 1}, 1.5268958346348338}, 
  {{4, 4, 5, 4}, 2.0444236632963757}, {{4, 5, 4, 5}, 0.7380136919956536}, 
  {{5, 4, 5, 4}, 0.7380136919956536}, {{4, 5, 4, 4}, 2.0444236632963757}, 
  {{3, 5, 5, 4}, 0.9750368723896611}, {{4, 5, 5, 3}, 0.9750368723896611}, 
  {{1, 6, 1, 6}, 0.9999999999999994}, {{6, 1, 6, 1}, 0.9999999999999994}, 
  {{3, 5, 5, 3}, 1.4200836985172587}, {{4, 4, 4, 4}, 54.43363712813288}, 
  {{4, 4, 4, 5}, 0.49994363226551003}, {{5, 4, 4, 4}, 0.4999436322655102}, 
  {{3, 5, 4, 4}, 4.986558195549088}, {{5, 3, 5, 4}, 1.0000000000000002}, 
  {{4, 4, 5, 3}, 4.9865581955490885}, {{4, 5, 3, 5}, 1.}, 
  {{1, 6, 5, 3}, 49.049751675377905}, {{6, 1, 6, 5}, 0.9999999999999997}, 
  {{3, 5, 6, 1}, 49.049751675377905}, {{5, 6, 1, 6}, 0.9999999999999997}}};
```
