## Supplementary material for "On the structure of multiple stable equilibria in competitive ecological systems": Mathematica+Julia code and examples: model-30.m

```
(* k = 3 model based on 200 simulations *)
m4 = {{{{6}, 0.38407699037620296}, {{5}, 0.5249343832020997}, 
  {{4}, 0.06824146981627296}, {{3}, 0.012248468941382328}, 
  {{1}, 0.010498687664041995}}, {{{6, 6}, 0.2922134733158355}, 
  {{5, 5}, 0.37270341207349084}, {{4, 5}, 0.05861767279090114}, 
  {{5, 6}, 0.08136482939632546}, {{6, 5}, 0.08136482939632546}, 
  {{5, 4}, 0.05861767279090114}, {{3, 5}, 0.012248468941382328}, 
  {{5, 3}, 0.012248468941382328}, {{1, 6}, 0.010498687664041995}, 
  {{6, 1}, 0.010498687664041995}, {{4, 4}, 0.00962379702537183}}, 
 {{{6, 6, 6}, 0.24934383202099739}, {{5, 5, 5}, 0.29483814523184604}, 
  {{4, 5, 5}, 0.03280839895013123}, {{5, 5, 6}, 0.0389326334208224}, 
  {{5, 6, 5}, 0.04899387576552931}, {{6, 5, 4}, 0.025809273840769906}, 
  {{5, 4, 5}, 0.04899387576552931}, {{6, 5, 5}, 0.0389326334208224}, 
  {{5, 5, 4}, 0.03280839895013123}, {{4, 5, 6}, 0.025809273840769906}, 
  {{3, 5, 5}, 0.006124234470691164}, {{5, 6, 6}, 0.03237095363079615}, 
  {{6, 6, 5}, 0.03237095363079615}, {{6, 5, 3}, 0.006124234470691164}, 
  {{5, 3, 5}, 0.012248468941382328}, {{5, 5, 3}, 0.006124234470691164}, 
  {{3, 5, 6}, 0.006124234470691164}, {{1, 6, 6}, 0.010498687664041995}, 
  {{6, 5, 6}, 0.010498687664041995}, {{6, 6, 1}, 0.010498687664041995}, 
  {{6, 1, 6}, 0.010498687664041995}, {{4, 4, 5}, 0.00962379702537183}, 
  {{5, 4, 4}, 0.00962379702537183}}}
w4 = {{{{6}, 0.38407699037620296}, {{5}, 0.5249343832020997}, 
  {{4}, 0.06824146981627296}, {{3}, 0.012248468941382328}, 
  {{1}, 0.010498687664041995}}, {{{6, 6}, 1.9809050388904166}, 
  {{5, 5}, 1.3525500000000001}, {{4, 5}, 1.636346153846154}, 
  {{5, 6}, 0.4035649202733485}, {{6, 5}, 0.4035649202733485}, 
  {{5, 4}, 1.636346153846154}, {{3, 5}, 1.905}, {{5, 3}, 1.905}, 
  {{1, 6}, 2.603644646924829}, {{6, 1}, 2.603644646924829}, 
  {{4, 4}, 2.0665680473372783}}, {{{6, 6, 6}, 1.1215443364767474}, 
  {{5, 5, 5}, 1.1141969186008067}, {{4, 5, 5}, 0.7883119613201597}, 
  {{5, 5, 6}, 0.673936089656217}, {{5, 6, 5}, 2.8424095271129612}, 
  {{6, 5, 4}, 2.840635532017333}, {{5, 4, 5}, 0.9730452216529294}, 
  {{6, 5, 5}, 0.6739360896562169}, {{5, 5, 4}, 0.7883119613201598}, 
  {{4, 5, 6}, 2.840635532017333}, {{3, 5, 5}, 0.704225352112676}, 
  {{5, 6, 6}, 0.5229218981392055}, {{6, 6, 5}, 0.5229218981392055}, 
  {{6, 5, 3}, 3.225806451612904}, {{5, 3, 5}, 1.}, 
  {{5, 5, 3}, 0.704225352112676}, {{3, 5, 6}, 3.225806451612904}, 
  {{1, 6, 6}, 1.3143712574850304}, {{6, 5, 6}, 0.8324661810613944}, 
  {{6, 6, 1}, 1.3143712574850304}, {{6, 1, 6}, 1.0000000000000002}, 
  {{4, 4, 5}, 1.164179104477612}, {{5, 4, 4}, 1.164179104477612}}}
```
