## Supplementary material for "On the structure of multiple stable equilibria in competitive ecological systems": Mathematica+Julia code and examples: model.m

```
(* modeling statistics with local model *)

(* this code builds and analyzes statistical mechanical models from
frequency data of equilibria for periodic niche models *)

(* some key functions, see also examples log file: *)

(* model[n, k]: computes the probabilities of all sequences 
 p(g_1,...,g_j), j <= k from equilibrium distribution data given in
 distribution[n] *)

(* weightsmodel[n, k]: computes the connected probability weight,
 i.e. e^(-\beta(g_1, ..., g_j) = p_c(g_1, ..., g_j)) *)

(* pp[sequence_, t_]: extracts the probability/weight for a given
 sequence from a model t (could be model or weightsmodel) *)

(* build[m, w, l, k]: produces a random (non-periodic) sequence of
  length l using k-gap model m + weight model w *)

(* periodic[m, w, l, k]:  adds probability with p = 0 if non-periodic *)

(* transfer[k, w, pt = True/False]: computes Markov matrix M or M^
  if pt = True/False, using k-gap weight model w. *)

(* -------------------------------- *)

(* various useful tools, not all used directly for producing results
of paper *)

(* given a set of gaps and occurrences, compute  probability *)

likelihood[x_, data_] :=
 With[{total = Apply[Plus, Map[First, data]]*1.0},
  Apply[Plus, Map[#[[1]]*Count[#[[2]], x] &, data]]/total]

likelihoods[data_] :=
 Table[likelihood[i, data],{i, 1, 6}]

(* now computes probability per lattice site *)

probabilities[data_] :=
 With[{l = likelihoods[data], size = Apply[Plus, data[[1, 2]]]},
  Table[l[[i]]*i/size,{i, 1, 6}]]

(* these properly add to 1 *)

(* computes probability per gap *)

fraction[x_, data_] :=
 With[{total = Apply[Plus, Map[First, data]]*1.0},
  Apply[Plus, 
   Map[N[#[[1]]*Count[#[[2]], x]/Length[#[[2]]]] &, data]]/total]

fractions[data_] :=
 Table[fraction[i, data],{i, 1, 6}]

(* now, given fractions with different g, predict probabilities in data *)

relative[rule_, {p_, sequence_}] :=
 Product[rule[[sequence[[i]]]],{i, 1, Length[sequence]}]*Length[sequence]

predictions[data_] :=
 With[{f = fractions[data]},
  Map[{relative[f, #], #[[1]], #[[2]]} &, data]]

normalize[list_] :=
 With[{x = Apply[Plus, Map[First, list]], y = Apply[Plus, Map[#[[2]] &, list]]},
   Map[{N[#[[1]]/x],  N[#[[2]]/y], #[[3]]} &, list]]

comparison[data_] :=
 Map[Map[Log, Drop[#, -1]]&, normalize[predictions[data]]]

information[ps_] :=
 Apply[Plus, Map[-# * Log[#]/Log[2.0] &, ps]]

(* given a sequence, compute occurrence of pairs at spacing k *)

pairs[s_, k_] :=
 Table[{s[[i]], s[[Mod[i + k -1,Length[s]]+ 1]]},{i, 1,Length[s]}]

(* ----------------------------------------- *)
(* stuff from here is used in function model[n, k] *)

(* given a sequence, occurrence of subsequences *)

subs1[s_, k_] :=
 Table[Take[Join[s, s],{i, i + k -1}],{i, 1,Length[s]}]

subs[s_, k_] := Join[subs1[s, k], subs1[Reverse[s], k]]

(* given a sequence with multiplicity, gives tally with multiplicity *)
(* apply to f = pairs or subs *)

tally[{n_, list_}, f_, k_] :=
 Map[{#[[1]], n #[[2]]}&, Tally[f@@{list, k}]]

(* combine a bunch of things with the same first element *)

combinesingle[list_] :={list[[1, 1]], Apply[Plus, Map[#[[2]]&, list]]}

(* combine tallies *)

combine[ts_] := Map[combinesingle, GatherBy[ts, First]]

(* all pairs/sequences from data *)

collection[data_, f_, k_] :=
  combine[Apply[Join, Map[tally[#, f, k] &, data]]]

(* converts statistics to probabilities *)

convert[s_] :=
 With[{total = N[Apply[Plus, Map[#[[2]] &, s]]]},
  Map[{#[[1]], #[[2]]/total}&, s]]

model[n_, k_] :=
 Table[convert[collection[distribution[n], subs, i]],{i, 1, k}]

(* look at pairs *)
pairing[n_, k_] :=
 {convert[ collection[distribution[n], subs, 1]],
  convert[collection[distribution[n], pairs, k]]}

(* from earlier model.m *)

(* compute in terms of probability weights (exponential of terms in L) *)

(* t = list of probabilities *)

p[n_, t_] := With[{item = Select[t[[1]], #[[1]] =={n} &]},
  If[item =={}, 0, item[[1, 2]]]]

(* extract probability associated with a sequence from rules *)

pp[sequence_, t_]:=
 With[{item = Select[t[[Length[sequence]]], #[[1]] ==sequence &]},
  If[item =={}, 0, 
     item[[1, 2]]]]

weight1[{n_, m_}, t_] := pp[{n, m}, t]/(pp[{n}, t]pp[{m}, t])

(* gets probabilities from weights at given level *)
partial[s_, k_, t_] := 
  Product[weight[Take[s,{i, i + k -1}], t],{i, 1,Length[s]-k+ 1}]

weight[s_, t_] := pp[s, t]/Product[partial[s, k, t],{k, 1,Length[s]-1}]

(* gets weights at a certain level *)

weights[k_, t_] := 
 Map[{#, weight[#, t]} &, Map[First, t[[k]]]]

weightsmodel[n_, k_] := (* weightsmodel[n, k] = *)
 With[{m = model[n, k]}, 
  Table[weights[i, m],{i, 1, k}]]

(* =================================================== *)
(* the remainder of this code does other various useful things but not
carefully documented, use at your own risk!*)

(* predicted probability of a sequence given model *)
(* note:pp generally gets probability in a model. *)

(* note, predict does not include distinct realizations. *)

predict[sequence_, k_, wm_] :=
  With[{s= Join[sequence, sequence]},
    Product[Product[pp[Take[s, {j, j + i}], wm],{j, 1, Length[sequence]}],
        {i, 0, k-1}]]

(* note: includes factor from distinct, probably don't use this function unless you are careful *)

logpredict[sequence_, k_, wm_] :=
  With[{s= Join[sequence, sequence]},
     Sum[Sum[Log[pp[Take[s, {j, j + i}], wm]],{j, 1, Length[sequence]}],
        {i, 0, k-1}]]+Log[distinct[sequence]]

(* probability of a sequence of gaps without periodicity *)


logpredictsingle[sequence_, k_, wm_] :=
  With[{s= sequence},
     Sum[Sum[Log[pp[Take[s, {j, j + i}], wm]],{j, 1, Length[sequence]-i}],
        {i, 0, k-1}]]

  
logpredictpair[sequence_, k_, wm_] :=
  {distinct[sequence],
  With[{s= Join[sequence, sequence]},
     Sum[Sum[Log[pp[Take[s, {j, j + i}], wm]],{j, 1, Length[sequence]}],
        {i, 0, k-1}]]}

(* how many times to count a sequence *)

rotations[s_] :=
 Table[RotateRight[s, i],{i, 1,Length[s]}]

distinct[s_] :=
 Length[Union[Join[rotations[s], rotations[Reverse[s]]]]]*
  Apply[Plus, s]/(Length[s])

(* comparing model to actual numbers *)

compare[data_, k_, wm_] :=
 With[{total = N[Apply[Plus, Map[First, data]]]},
  Table[{Log[data[[i,1]]/total],  logpredict[ data[[i, 2]], k, wm],
         data[[i, 2]]},
    {i, 1,Length[data]}]]

comparenumbers[n_, k_, wm_] :=
  Map[Drop[#, -1] &, compare[distribution[n], k, wm]]

(* wm = weightsmodel[90, k] *)

(* looking at two point correlators *)

correlate[{i_, j_}, k_, n_] := correlate[{i, j}, k, n] =
 Table[weight[{i, j}, pairing[n, x]],{x, 1, k}]


(* compute entropy given weights *)

conditional[initial_, final_, t_] :=
 conditional[initial, final, t] =
  Product[pp[Append[Take[initial, -i], final], t],{i, 0,Length[initial]}]

(* only takes beginning from some i, used in periodic construction *)

conditional[initial_, final_, ii_, t_] :=
  Product[pp[Append[Take[initial, -i], final], t],{i, ii,Length[initial]}]

(* Compute total entropy*)

entropy[list_] :=
 Sum[If[list[[i]] > 0,-list[[i]]Log[ list[[i]]], 0],{ i, 1,Length[list]}]

(*  Variance for log probability *)

entropy2[list_] :=
 Sum[If[list[[i]] > 0,-list[[i]]Log[ list[[i]]]^2, 0],{ i, 1,Length[list]}]


conditionals[initial_, t_] :=
 Table[conditional[initial, i, t],{i, 1, 6}]

(*  Looking at all conditionals for k = 4 *)

(* Map[{#[[1]],#[[2]]*entropy[conditionals[ #[[1]], w4]]} &, m4[[3]]] *)


(* computes entropy based on model m, w with k initial states (Use k = 3!) *)

entropy[m_, w_, k_] :=
 Apply[Plus,
        Map[#[[2]]*entropy[conditionals[ #[[1]], w]] &, m[[k]]]]

(*  variance *)
entropy2[m_, w_, k_] :=
 Apply[Plus,
        Map[#[[2]]*entropy2[conditionals[ #[[1]], w]] &, m[[k]]]]

(* average gap size  (gapsize[m4[[1]]]) *)

gapsize[m_] :=Sum[m[[i, 1, 1]]*m[[i, 2]],{i, 1, 5}]

(* computes average of ln p, without weighting by p *)

entropy0[list_] :=
 Sum[If[list[[i]] > 0, {1, Log[ list[[i]]]}, 0],{ i, 1,Length[list]}]

entropy0[m_, w_, k_] :=
 With[{x = Apply[Plus,
        Map[entropy0[conditionals[ #[[1]], w]] &, m[[k]]]]},
    x[[2]]/x[[1]]]

(* builds a sequence of length at least l, 
  returns probability so far, sequence *)
(* needs probability model, weights *)

(* Given a set of options, probabilities, picks one: *)

(* gets seeds, weighted by length of first gap *)

seeds[m_, k_] :=
  Map[{#[[2]]*#[[1, 1]], #[[1]]}&, m[[k]]]

(* ----------------key function--------------------- *)
(* builds a random sequence of length l using k-gap model m, w *)

build[m_, w_, l_, k_] :=
  With[{s= seeds[m, k]},
    initial = RandomChoice[Map[First, s] -> s];
    t = Apply[Plus, Map[First, s]];
    current = {Log[initial[[1]]/t], initial[[2]]};
    While[Apply[Plus, current[[2]]] < l,
(*       Print[current];  *) 
       c = conditionals[Take[current[[2]], 1-k], w];
       next = RandomChoice[c -> {1, 2, 3, 4, 5, 6}];
       current ={current[[1]]+ Log[c[[next]]], Append[current[[2]], next]}];
    current]

(* makes a  sequence, *)
(* checks if  is periodic and if so additional probability factor 
   returns (probability factor 0 or p_extra, total probability, sequence)
*)

periodic[m_, w_, l_, k_] :=
  With[{b= build[m, w, l, k]},
    p1 = E^b[[1]]; s= b[[2]];
    double = Join[s, s];
    ls=Length[s];
    If[Apply[Plus, s]!= l,{0, p1, s},
       With[{extra = 
          Product[conditional[Take[double,{ls-k + i+1,ls+i-1}],
                              double[[ls+ i]], i, w],{i, 1, k -1}]},
            {extra, p1*extra, s}]]]

(* computes n of these built sequences and gets statistics on
 log probabilities *)

statistics[m_, w_, k_, n_, l_] :=
  Block[{l1 = 0, l2 = 0, l3 = 0, s},
     For[j = 1, j <= n, j += 1,
        s = build[m, w, l, k];
        l1 = l1 + s[[1]]; l2 = l2 + s[[1]]^2; l3 = l3 + s[[1]]^3
        ];
     {l, k, n, (l1/n)/l,  (l2/n -(l1/n)^2)/l, 
        (l3/n-3 (l2/n)(l1/n)+ 2 (l1/n)^3)/l^(3/2.0)}]
(*  *)

(* gets statistics for a range of l's *)
statistics[m_, w_, k_, n_, la_, lb_, step_] :=
 Table[statistics[m, w, k, n, l],{l, la, lb, step}]

(* looking at some random uncorrelated stuff central limit theorem *)

central[l_, n_] :=
  Block[{l1 = 0.0, l2 = 0.0, l3 = 0.0, s},
     For[j = 1, j <= n, j += 1,
        s = Sum[RandomChoice[{1, 2,  4, 8, 16}],{k, 1, l}];
        l1 = l1 + s; l2 = l2 + s^2; l3 = l3 + s^3
        ];
     {l, k, n, l1/n, l2/n -(l1/n)^2, l3/n-3 (l2/n)(l1/n)+ 2 (l1/n)^3}]
(*  *)

(* theoretical prediction for distribution *)

distribute[x_, n_] := E^(-(x + 0.143 n)^2/(2 * 0.049 *n))

(* gets probability that a sequence is acceptable/periodic *)

acceptable[m_, w_, l_, k_, n_] :=
 Mean[Map[First,Table[periodic[m, w, l, k],{i, 1, n}]]]

(* -------------------------------------------- *)
(* key function: computes matrix M or M^, "transfer matrix" *)

(* computing transfer matrix for k-1 point functions, with and without
   p's *)

transfer[k_, w_, pt_] :=
  With[{indices = Map[First, w[[k -1]]]},
    l =Length[indices];
    Table[If[Drop[indices[[i]], 1] ==
             Drop[indices[[j]], -1],
                If[pt, conditional[indices[[i]], Last[indices[[j]]], w],
                   If[conditional[indices[[i]], Last[indices[[j]]], w]>0,
                        1, 0]], 0],{i, 1, l},{j, 1, l}]]

(* weighted entropy: computes expected ln p per gap with weights 1 *)
(* 1/9/24: corrected to include proper weighting factor depending on previous state on lattice *)

lastg[k_, w_] := Map[Last[First[#]]&, w[[k -1]]]
  
(*  trying to get entropy per lattice site *)

latticeentropy[k_, w_] :=
 With[{tf = transfer[k, w, False], tt = transfer[k, w, True]},
   ll = lastg[k, w];
   ww =Eigensystem[N[Transpose[tf]]];
   ee = ww[[1, 1]];
   w1 = Abs[ww[[2, 1]]];
   w2 = w1/Apply[Plus, w1];
   Print[w2];
   Sum[w2[[i]]*(entropy0[tt[[i]]][[2]]/(ee*ll[[i]])),{i, 1,Length[w1]}]
]

(*  proper distribution *)

normalize1[v_] := Abs[v]/Apply[Plus, Abs[v]]

wentropy[k_, w_] :=
 With[{tf = transfer[k, w, False], tt = transfer[k, w, True]},
   wwl =Eigensystem[N[Transpose[tf]]];
   eel = wwl[[1, 1]];
   w1l = normalize1[wwl[[2, 1]]];
   wwr = Eigensystem[N[tf]];
   eer = wwr[[1, 1]];
   w1r = normalize1[wwr[[2, 1]]];
   w1 = Table[w1l[[i]]*w1r[[i]], {i, 1,Length[w1r]}];
   w1m = Apply[Plus, w1];
   Print[eel, eer, w1l, w1r, w1/w1m, w1m];
   Sum[If[tf[[i, j]]== 1,
          w1l[[i]]*w1r[[j]]*Log[tt[[i, j]]]/(eer*w1m), 0],
       {i, 1,Length[w1r]}, {j, 1, Length[w1r]}]   
]

frequencies3[k_, w_] :=
  With[{tf = transfer[k, w, False], tt = transfer[k, w, True]},
   wwl =Eigensystem[N[Transpose[tf]]];
   eel = wwl[[1, 1]];
   w1l = normalize1[wwl[[2, 1]]];
   wwr = Eigensystem[N[tf]];
   eer = wwr[[1, 1]];
   w1r = normalize1[wwr[[2, 1]]];
   w1 = Table[w1l[[i]]*w1r[[i]], {i, 1,Length[w1r]}];
   w1m = Apply[Plus, w1];
   w1/w1m]

(*  Average Size at position i out of k -1 with unit weighting *)

averagegap1[i_, k_, w_] :=
  With[{ f = frequencies3[k, w]},
    Sum[ f[[j]]*w[[k -1, j, 1, i]], { j, 1,Length[ f]}]]
```
