## Supplementary material for "On the structure of multiple stable equilibria in competitive ecological systems": Mathematica+Julia code and examples: niches.jl

```
# solves dynamics of LV with multiple stable equilibria
# based on version 7 
# but this starts with a circulant matrix with standard structure
# parameters are n = lattice size, l = system size
# 9/10/23: put in gaps, canonical form

#uncomment when running directly to ensure package
#import Pkg
#Pkg.add("DifferentialEquations")

#needed for running on cluster
#using Pkg
#project_dir="v1.6"
#Pkg.activate(project_dir)
#Pkg.add("DifferentialEquations")

using DifferentialEquations
using Random
using LinearAlgebra

# initialize a matrix  with parameters n, l

function interaction(distance)
 return exp(-distance^4)
end

function potential(x, l, k = 10)
 return sum(interaction(x-j*l) for j = -k:k)
end

function fixmatrix(n, l)
 y = Matrix{Float64}(undef, n, n)
 for i in 1:n
  for j in 1:n
   y[i, j] = potential(min(abs(i-j),n-abs(i-j))*l/n, l)
  end
 end
 return y
end

# get gaps, canonical form

function gaps(list, size)
 push!([list[i + 1]-list[i] for i in 1:(length(list)-1)], list[1] -last(list)+ size)
end

function rotate(list, k)
 x = vcat(list, list)
 [x[i] for i in k +1:k + length(list)]
end

#gets the smallest thing in a list
function least(lists)
 if length(lists) == 1
  return lists[1]
 elseif length(lists) == 2
  if lists[1] < lists[2]
   return lists[1]
  else
   return lists[2]
  end
 else
  return least([least([lists[1], lists[2]]), least([lists[i] for i in 3:length(lists)])])
 end
end

function canonical(list)
 least([least([rotate(list, i) for i in 1:length(list)]),
        least([reverse(rotate(list, i)) for i in 1:length(list)])])
end

# start with initial values, run for time t, give output

function run(m::Matrix{Float64}, initial, tt)
 f(u, p, t) = u - u.*(m*u)
 problem = ODEProblem(f, initial, (0.0, tt))
 solution = solve(problem)
 solution(tt)
end
 
# find what species are present

function remaining(list, tolerance)
 [z for z in 1:length(list) if list[z] > tolerance]
end

# find what species have diverged

function diverged(list, tolerance)
 [z for z in 1:length(list) if list[z] > tolerance]
end


# run system with random initial values many times and get species
#  remaining
# each time around do multiple passes and get rid of negative populations
#for fixed matrices, don't worry about declaring symmetric (?)

function randomrun(m::Matrix{Float64}, tt, blocks::Int, scale = 1.0, tolerance = 0.000001)
 i = map(x -> abs(scale*x), randn(size(m)[1]))
 for j in 1:blocks
  r = run(m, i, tt)
  i = map(x -> (x < 0.0) ? 0.0 : x, r)
  x = remaining(r, 1000000.0)
  if length(x) > 0
   return -x
  end
  #print(j, r, i, x)
 end
# (i, r, remaining(r, tolerance))
 remaining(i, tolerance)
end

# code to take two lists, 1 of solutions and 1 of instances, add new result

function augment(list::Vector{Vector{Int64}}, numbers::Vector{Int64}, new::Vector{Int64})
#function augment(list, numbers, new)
  for i in 1:length(list)
    if list[i] == new
      numbers[i] += 1
      return i
    end
  end
  push!(list, new)
  push!(numbers, 1)
end

#runs and keeps list of results
#modified for niche matrices.

function dynamics(n, l, count, index) #n, count as integers
    m = fixmatrix(n, l)
    o = open(string("output"), "a")
    write(o, "matrix[" * string(index) * "] =" * string(m, "\n"))
    write(o, "parameters[" * string(index) * "] =" * string([n, l], "\n"))
    types = Vector{Vector{Int64}}(undef, 0)
    hits = Vector{Int64}(undef, 0)
    for k in 1:count
    	r = randomrun(m, 100, 200)
	augment(types, hits, canonical(gaps(r, n)))
    end
    write(o, "types[" * string(index) * "] =" * string(types, "\n"))
    write(o, "hits[" * string(index) * "] =" * string(hits, "\n"))
    close(o)
end

# dynamics(ARGS[1],ARGS[2])


# runs a single time with a niche matrix with given parameters
#  mathematica initialization

dynamics(parse(Int, ARGS[1]),parse(Float64, ARGS[2]),parse(Int, ARGS[3]),parse(Int, ARGS[4]))
```
