## Supplementary material for "On the structure of multiple stable equilibria in competitive ecological systems": Mathematica+Julia code and examples: README

Files to accompany bioarxiv version of paper "On the structure of
multiple stable equilibria in competitive ecological systems" 
by Taylor and O'Dwyer

This contains 3 sets of files:

(1) Mathematica code to analyze periodic niche interaction matrices
and associated equilibria

(2) Julia code to run simulations and collect resulting stable
equilibrium states

(3) Mathematica code to produce and analyze a statistical model given
data from many simulations.

More details are in files examples-i.txt, i = 1, 2, 3, which give
examples of use with commentary.

----------------------------------------

(1) Mathematica code to analyze matrices and equilibria

basins.m: mathematica code to compute periodic niche interaction
  matrices and analyze associated stable equilibria

examples-1.txt: examples of usage of basins.m

----------------------------------------

(2) Julia code to run simulations

niches.jl: code to run simulations
combine.jl: code to combine output from multiple simulations

examples-2.txt: example of usage (2 * 100 runs at S= 30)

output: results of simulations (output of niches.jl)
header-10.jl: auxiliary header file to combine simulation results
results-200.jl: header + output
combined: combined results of 2*100 simulations (output of combine.jl)

----------------------------------------

(3) Mathematica code to build statistical model from data

model.m: code to take simulation results and build statistical model
combined.m: data from combined simulations, in mathematica format

examples-3.txt: examples of usage of model.m code

model-30.m: statistical mechanical model computed from code and simple
 data set above in file combined.m
 (w4 contains e^(-alpha), e^(-beta)'s, m4 is probabilities p(g_1), etc.)

sorted-200.m: tabulation of equilibria and frequencies for sample 200 runs

tree.m: code to generate all possible stable subset equilibria
 compatible with a given local model

90-all.m: results of 10^6 simulations at size S= 90 (99.5% gave stable
 solutions).
m4-90.m: statistical mechanical model built from data from 90-all at k = 4.
