## Supplementary material for "On the structure of multiple stable equilibria in competitive ecological systems": Mathematica+Julia code and examples: results-200.jl

```
# header file for up to 10 sets of simulation results
matrix = Vector{Any}(undef, 10)
parameters = Vector{Any}(undef, 10)
types = Vector{Vector{Vector{Int64}}}(undef, 10)
hits = Vector{Vector{Int64}}(undef, 10)
matrix[1] =[1.0 0.9961013694701175 0.9394130628134758 0.7287633299194912 0.36787944117144233 0.08703836765622351 0.006329715427485747 8.448756028504651e-5 1.1253517471925912e-7 7.404699497933558e-12 1.0848552640429378e-17 1.4524053946493149e-25 6.639677199580735e-36 3.526349876712582e-49 6.740378884131016e-66 2.6157710955041962e-86 6.740378884131016e-66 3.526349876712582e-49 6.639677199580735e-36 1.4524053946493149e-25 1.0848552640429378e-17 7.404699497933558e-12 1.1253517471925912e-7 8.448756028504651e-5 0.006329715427485747 0.08703836765622351 0.36787944117144233 0.7287633299194912 0.9394130628134758 0.9961013694701175; 0.9961013694701175 1.0 0.9961013694701175 0.9394130628134758 0.7287633299194912 0.36787944117144233 0.08703836765622351 0.006329715427485747 8.448756028504651e-5 1.1253517471925912e-7 7.404699497933558e-12 1.0848552640429378e-17 1.4524053946493149e-25 6.639677199580735e-36 3.526349876712582e-49 6.740378884131016e-66 2.6157710955041962e-86 6.740378884131016e-66 3.526349876712582e-49 6.639677199580735e-36 1.4524053946493149e-25 1.0848552640429378e-17 7.404699497933558e-12 1.1253517471925912e-7 8.448756028504651e-5 0.006329715427485747 0.08703836765622351 0.36787944117144233 0.7287633299194912 0.9394130628134758; 0.9394130628134758 0.9961013694701175 1.0 0.9961013694701175 0.9394130628134758 0.7287633299194912 0.36787944117144233 0.08703836765622351 0.006329715427485747 8.448756028504651e-5 1.1253517471925912e-7 7.404699497933558e-12 1.0848552640429378e-17 1.4524053946493149e-25 6.639677199580735e-36 3.526349876712582e-49 6.740378884131016e-66 2.6157710955041962e-86 6.740378884131016e-66 3.526349876712582e-49 6.639677199580735e-36 1.4524053946493149e-25 1.0848552640429378e-17 7.404699497933558e-12 1.1253517471925912e-7 8.448756028504651e-5 0.006329715427485747 0.08703836765622351 0.36787944117144233 0.7287633299194912; 0.7287633299194912 0.9394130628134758 0.9961013694701175 1.0 0.9961013694701175 0.9394130628134758 0.7287633299194912 0.36787944117144233 0.08703836765622351 0.006329715427485747 8.448756028504651e-5 1.1253517471925912e-7 7.404699497933558e-12 1.0848552640429378e-17 1.4524053946493149e-25 6.639677199580735e-36 3.526349876712582e-49 6.740378884131016e-66 2.6157710955041962e-86 6.740378884131016e-66 3.526349876712582e-49 6.639677199580735e-36 1.4524053946493149e-25 1.0848552640429378e-17 7.404699497933558e-12 1.1253517471925912e-7 8.448756028504651e-5 0.006329715427485747 0.08703836765622351 0.36787944117144233; 0.36787944117144233 0.7287633299194912 0.9394130628134758 0.9961013694701175 1.0 0.9961013694701175 0.9394130628134758 0.7287633299194912 0.36787944117144233 0.08703836765622351 0.006329715427485747 8.448756028504651e-5 1.1253517471925912e-7 7.404699497933558e-12 1.0848552640429378e-17 1.4524053946493149e-25 6.639677199580735e-36 3.526349876712582e-49 6.740378884131016e-66 2.6157710955041962e-86 6.740378884131016e-66 3.526349876712582e-49 6.639677199580735e-36 1.4524053946493149e-25 1.0848552640429378e-17 7.404699497933558e-12 1.1253517471925912e-7 8.448756028504651e-5 0.006329715427485747 0.08703836765622351; 0.08703836765622351 0.36787944117144233 0.7287633299194912 0.9394130628134758 0.9961013694701175 1.0 0.9961013694701175 0.9394130628134758 0.7287633299194912 0.36787944117144233 0.08703836765622351 0.006329715427485747 8.448756028504651e-5 1.1253517471925912e-7 7.404699497933558e-12 1.0848552640429378e-17 1.4524053946493149e-25 6.639677199580735e-36 3.526349876712582e-49 6.740378884131016e-66 2.6157710955041962e-86 6.740378884131016e-66 3.526349876712582e-49 6.639677199580735e-36 1.4524053946493149e-25 1.0848552640429378e-17 7.404699497933558e-12 1.1253517471925912e-7 8.448756028504651e-5 0.006329715427485747; 0.006329715427485747 0.08703836765622351 0.36787944117144233 0.7287633299194912 0.9394130628134758 0.9961013694701175 1.0 0.9961013694701175 0.9394130628134758 0.7287633299194912 0.36787944117144233 0.08703836765622351 0.006329715427485747 8.448756028504651e-5 1.1253517471925912e-7 7.404699497933558e-12 1.0848552640429378e-17 1.4524053946493149e-25 6.639677199580735e-36 3.526349876712582e-49 6.740378884131016e-66 2.6157710955041962e-86 6.740378884131016e-66 3.526349876712582e-49 6.639677199580735e-36 1.4524053946493149e-25 1.0848552640429378e-17 7.404699497933558e-12 1.1253517471925912e-7 8.448756028504651e-5; 8.448756028504651e-5 0.006329715427485747 0.08703836765622351 0.36787944117144233 0.7287633299194912 0.9394130628134758 0.9961013694701175 1.0 0.9961013694701175 0.9394130628134758 0.7287633299194912 0.36787944117144233 0.08703836765622351 0.006329715427485747 8.448756028504651e-5 1.1253517471925912e-7 7.404699497933558e-12 1.0848552640429378e-17 1.4524053946493149e-25 6.639677199580735e-36 3.526349876712582e-49 6.740378884131016e-66 2.6157710955041962e-86 6.740378884131016e-66 3.526349876712582e-49 6.639677199580735e-36 1.4524053946493149e-25 1.0848552640429378e-17 7.404699497933558e-12 1.1253517471925912e-7; 1.1253517471925912e-7 8.448756028504651e-5 0.006329715427485747 0.08703836765622351 0.36787944117144233 0.7287633299194912 0.9394130628134758 0.9961013694701175 1.0 0.9961013694701175 0.9394130628134758 0.7287633299194912 0.36787944117144233 0.08703836765622351 0.006329715427485747 8.448756028504651e-5 1.1253517471925912e-7 7.404699497933558e-12 1.0848552640429378e-17 1.4524053946493149e-25 6.639677199580735e-36 3.526349876712582e-49 6.740378884131016e-66 2.6157710955041962e-86 6.740378884131016e-66 3.526349876712582e-49 6.639677199580735e-36 1.4524053946493149e-25 1.0848552640429378e-17 7.404699497933558e-12; 7.404699497933558e-12 1.1253517471925912e-7 8.448756028504651e-5 0.006329715427485747 0.08703836765622351 0.36787944117144233 0.7287633299194912 0.9394130628134758 0.9961013694701175 1.0 0.9961013694701175 0.9394130628134758 0.7287633299194912 0.36787944117144233 0.08703836765622351 0.006329715427485747 8.448756028504651e-5 1.1253517471925912e-7 7.404699497933558e-12 1.0848552640429378e-17 1.4524053946493149e-25 6.639677199580735e-36 3.526349876712582e-49 6.740378884131016e-66 2.6157710955041962e-86 6.740378884131016e-66 3.526349876712582e-49 6.639677199580735e-36 1.4524053946493149e-25 1.0848552640429378e-17; 1.0848552640429378e-17 7.404699497933558e-12 1.1253517471925912e-7 8.448756028504651e-5 0.006329715427485747 0.08703836765622351 0.36787944117144233 0.7287633299194912 0.9394130628134758 0.9961013694701175 1.0 0.9961013694701175 0.9394130628134758 0.7287633299194912 0.36787944117144233 0.08703836765622351 0.006329715427485747 8.448756028504651e-5 1.1253517471925912e-7 7.404699497933558e-12 1.0848552640429378e-17 1.4524053946493149e-25 6.639677199580735e-36 3.526349876712582e-49 6.740378884131016e-66 2.6157710955041962e-86 6.740378884131016e-66 3.526349876712582e-49 6.639677199580735e-36 1.4524053946493149e-25; 1.4524053946493149e-25 1.0848552640429378e-17 7.404699497933558e-12 1.1253517471925912e-7 8.448756028504651e-5 0.006329715427485747 0.08703836765622351 0.36787944117144233 0.7287633299194912 0.9394130628134758 0.9961013694701175 1.0 0.9961013694701175 0.9394130628134758 0.7287633299194912 0.36787944117144233 0.08703836765622351 0.006329715427485747 8.448756028504651e-5 1.1253517471925912e-7 7.404699497933558e-12 1.0848552640429378e-17 1.4524053946493149e-25 6.639677199580735e-36 3.526349876712582e-49 6.740378884131016e-66 2.6157710955041962e-86 6.740378884131016e-66 3.526349876712582e-49 6.639677199580735e-36; 6.639677199580735e-36 1.4524053946493149e-25 1.0848552640429378e-17 7.404699497933558e-12 1.1253517471925912e-7 8.448756028504651e-5 0.006329715427485747 0.08703836765622351 0.36787944117144233 0.7287633299194912 0.9394130628134758 0.9961013694701175 1.0 0.9961013694701175 0.9394130628134758 0.7287633299194912 0.36787944117144233 0.08703836765622351 0.006329715427485747 8.448756028504651e-5 1.1253517471925912e-7 7.404699497933558e-12 1.0848552640429378e-17 1.4524053946493149e-25 6.639677199580735e-36 3.526349876712582e-49 6.740378884131016e-66 2.6157710955041962e-86 6.740378884131016e-66 3.526349876712582e-49; 3.526349876712582e-49 6.639677199580735e-36 1.4524053946493149e-25 1.0848552640429378e-17 7.404699497933558e-12 1.1253517471925912e-7 8.448756028504651e-5 0.006329715427485747 0.08703836765622351 0.36787944117144233 0.7287633299194912 0.9394130628134758 0.9961013694701175 1.0 0.9961013694701175 0.9394130628134758 0.7287633299194912 0.36787944117144233 0.08703836765622351 0.006329715427485747 8.448756028504651e-5 1.1253517471925912e-7 7.404699497933558e-12 1.0848552640429378e-17 1.4524053946493149e-25 6.639677199580735e-36 3.526349876712582e-49 6.740378884131016e-66 2.6157710955041962e-86 6.740378884131016e-66; 6.740378884131016e-66 3.526349876712582e-49 6.639677199580735e-36 1.4524053946493149e-25 1.0848552640429378e-17 7.404699497933558e-12 1.1253517471925912e-7 8.448756028504651e-5 0.006329715427485747 0.08703836765622351 0.36787944117144233 0.7287633299194912 0.9394130628134758 0.9961013694701175 1.0 0.9961013694701175 0.9394130628134758 0.7287633299194912 0.36787944117144233 0.08703836765622351 0.006329715427485747 8.448756028504651e-5 1.1253517471925912e-7 7.404699497933558e-12 1.0848552640429378e-17 1.4524053946493149e-25 6.639677199580735e-36 3.526349876712582e-49 6.740378884131016e-66 2.6157710955041962e-86; 2.6157710955041962e-86 6.740378884131016e-66 3.526349876712582e-49 6.639677199580735e-36 1.4524053946493149e-25 1.0848552640429378e-17 7.404699497933558e-12 1.1253517471925912e-7 8.448756028504651e-5 0.006329715427485747 0.08703836765622351 0.36787944117144233 0.7287633299194912 0.9394130628134758 0.9961013694701175 1.0 0.9961013694701175 0.9394130628134758 0.7287633299194912 0.36787944117144233 0.08703836765622351 0.006329715427485747 8.448756028504651e-5 1.1253517471925912e-7 7.404699497933558e-12 1.0848552640429378e-17 1.4524053946493149e-25 6.639677199580735e-36 3.526349876712582e-49 6.740378884131016e-66; 6.740378884131016e-66 2.6157710955041962e-86 6.740378884131016e-66 3.526349876712582e-49 6.639677199580735e-36 1.4524053946493149e-25 1.0848552640429378e-17 7.404699497933558e-12 1.1253517471925912e-7 8.448756028504651e-5 0.006329715427485747 0.08703836765622351 0.36787944117144233 0.7287633299194912 0.9394130628134758 0.9961013694701175 1.0 0.9961013694701175 0.9394130628134758 0.7287633299194912 0.36787944117144233 0.08703836765622351 0.006329715427485747 8.448756028504651e-5 1.1253517471925912e-7 7.404699497933558e-12 1.0848552640429378e-17 1.4524053946493149e-25 6.639677199580735e-36 3.526349876712582e-49; 3.526349876712582e-49 6.740378884131016e-66 2.6157710955041962e-86 6.740378884131016e-66 3.526349876712582e-49 6.639677199580735e-36 1.4524053946493149e-25 1.0848552640429378e-17 7.404699497933558e-12 1.1253517471925912e-7 8.448756028504651e-5 0.006329715427485747 0.08703836765622351 0.36787944117144233 0.7287633299194912 0.9394130628134758 0.9961013694701175 1.0 0.9961013694701175 0.9394130628134758 0.7287633299194912 0.36787944117144233 0.08703836765622351 0.006329715427485747 8.448756028504651e-5 1.1253517471925912e-7 7.404699497933558e-12 1.0848552640429378e-17 1.4524053946493149e-25 6.639677199580735e-36; 6.639677199580735e-36 3.526349876712582e-49 6.740378884131016e-66 2.6157710955041962e-86 6.740378884131016e-66 3.526349876712582e-49 6.639677199580735e-36 1.4524053946493149e-25 1.0848552640429378e-17 7.404699497933558e-12 1.1253517471925912e-7 8.448756028504651e-5 0.006329715427485747 0.08703836765622351 0.36787944117144233 0.7287633299194912 0.9394130628134758 0.9961013694701175 1.0 0.9961013694701175 0.9394130628134758 0.7287633299194912 0.36787944117144233 0.08703836765622351 0.006329715427485747 8.448756028504651e-5 1.1253517471925912e-7 7.404699497933558e-12 1.0848552640429378e-17 1.4524053946493149e-25; 1.4524053946493149e-25 6.639677199580735e-36 3.526349876712582e-49 6.740378884131016e-66 2.6157710955041962e-86 6.740378884131016e-66 3.526349876712582e-49 6.639677199580735e-36 1.4524053946493149e-25 1.0848552640429378e-17 7.404699497933558e-12 1.1253517471925912e-7 8.448756028504651e-5 0.006329715427485747 0.08703836765622351 0.36787944117144233 0.7287633299194912 0.9394130628134758 0.9961013694701175 1.0 0.9961013694701175 0.9394130628134758 0.7287633299194912 0.36787944117144233 0.08703836765622351 0.006329715427485747 8.448756028504651e-5 1.1253517471925912e-7 7.404699497933558e-12 1.0848552640429378e-17; 1.0848552640429378e-17 1.4524053946493149e-25 6.639677199580735e-36 3.526349876712582e-49 6.740378884131016e-66 2.6157710955041962e-86 6.740378884131016e-66 3.526349876712582e-49 6.639677199580735e-36 1.4524053946493149e-25 1.0848552640429378e-17 7.404699497933558e-12 1.1253517471925912e-7 8.448756028504651e-5 0.006329715427485747 0.08703836765622351 0.36787944117144233 0.7287633299194912 0.9394130628134758 0.9961013694701175 1.0 0.9961013694701175 0.9394130628134758 0.7287633299194912 0.36787944117144233 0.08703836765622351 0.006329715427485747 8.448756028504651e-5 1.1253517471925912e-7 7.404699497933558e-12; 7.404699497933558e-12 1.0848552640429378e-17 1.4524053946493149e-25 6.639677199580735e-36 3.526349876712582e-49 6.740378884131016e-66 2.6157710955041962e-86 6.740378884131016e-66 3.526349876712582e-49 6.639677199580735e-36 1.4524053946493149e-25 1.0848552640429378e-17 7.404699497933558e-12 1.1253517471925912e-7 8.448756028504651e-5 0.006329715427485747 0.08703836765622351 0.36787944117144233 0.7287633299194912 0.9394130628134758 0.9961013694701175 1.0 0.9961013694701175 0.9394130628134758 0.7287633299194912 0.36787944117144233 0.08703836765622351 0.006329715427485747 8.448756028504651e-5 1.1253517471925912e-7; 1.1253517471925912e-7 7.404699497933558e-12 1.0848552640429378e-17 1.4524053946493149e-25 6.639677199580735e-36 3.526349876712582e-49 6.740378884131016e-66 2.6157710955041962e-86 6.740378884131016e-66 3.526349876712582e-49 6.639677199580735e-36 1.4524053946493149e-25 1.0848552640429378e-17 7.404699497933558e-12 1.1253517471925912e-7 8.448756028504651e-5 0.006329715427485747 0.08703836765622351 0.36787944117144233 0.7287633299194912 0.9394130628134758 0.9961013694701175 1.0 0.9961013694701175 0.9394130628134758 0.7287633299194912 0.36787944117144233 0.08703836765622351 0.006329715427485747 8.448756028504651e-5; 8.448756028504651e-5 1.1253517471925912e-7 7.404699497933558e-12 1.0848552640429378e-17 1.4524053946493149e-25 6.639677199580735e-36 3.526349876712582e-49 6.740378884131016e-66 2.6157710955041962e-86 6.740378884131016e-66 3.526349876712582e-49 6.639677199580735e-36 1.4524053946493149e-25 1.0848552640429378e-17 7.404699497933558e-12 1.1253517471925912e-7 8.448756028504651e-5 0.006329715427485747 0.08703836765622351 0.36787944117144233 0.7287633299194912 0.9394130628134758 0.9961013694701175 1.0 0.9961013694701175 0.9394130628134758 0.7287633299194912 0.36787944117144233 0.08703836765622351 0.006329715427485747; 0.006329715427485747 8.448756028504651e-5 1.1253517471925912e-7 7.404699497933558e-12 1.0848552640429378e-17 1.4524053946493149e-25 6.639677199580735e-36 3.526349876712582e-49 6.740378884131016e-66 2.6157710955041962e-86 6.740378884131016e-66 3.526349876712582e-49 6.639677199580735e-36 1.4524053946493149e-25 1.0848552640429378e-17 7.404699497933558e-12 1.1253517471925912e-7 8.448756028504651e-5 0.006329715427485747 0.08703836765622351 0.36787944117144233 0.7287633299194912 0.9394130628134758 0.9961013694701175 1.0 0.9961013694701175 0.9394130628134758 0.7287633299194912 0.36787944117144233 0.08703836765622351; 0.08703836765622351 0.006329715427485747 8.448756028504651e-5 1.1253517471925912e-7 7.404699497933558e-12 1.0848552640429378e-17 1.4524053946493149e-25 6.639677199580735e-36 3.526349876712582e-49 6.740378884131016e-66 2.6157710955041962e-86 6.740378884131016e-66 3.526349876712582e-49 6.639677199580735e-36 1.4524053946493149e-25 1.0848552640429378e-17 7.404699497933558e-12 1.1253517471925912e-7 8.448756028504651e-5 0.006329715427485747 0.08703836765622351 0.36787944117144233 0.7287633299194912 0.9394130628134758 0.9961013694701175 1.0 0.9961013694701175 0.9394130628134758 0.7287633299194912 0.36787944117144233; 0.36787944117144233 0.08703836765622351 0.006329715427485747 8.448756028504651e-5 1.1253517471925912e-7 7.404699497933558e-12 1.0848552640429378e-17 1.4524053946493149e-25 6.639677199580735e-36 3.526349876712582e-49 6.740378884131016e-66 2.6157710955041962e-86 6.740378884131016e-66 3.526349876712582e-49 6.639677199580735e-36 1.4524053946493149e-25 1.0848552640429378e-17 7.404699497933558e-12 1.1253517471925912e-7 8.448756028504651e-5 0.006329715427485747 0.08703836765622351 0.36787944117144233 0.7287633299194912 0.9394130628134758 0.9961013694701175 1.0 0.9961013694701175 0.9394130628134758 0.7287633299194912; 0.7287633299194912 0.36787944117144233 0.08703836765622351 0.006329715427485747 8.448756028504651e-5 1.1253517471925912e-7 7.404699497933558e-12 1.0848552640429378e-17 1.4524053946493149e-25 6.639677199580735e-36 3.526349876712582e-49 6.740378884131016e-66 2.6157710955041962e-86 6.740378884131016e-66 3.526349876712582e-49 6.639677199580735e-36 1.4524053946493149e-25 1.0848552640429378e-17 7.404699497933558e-12 1.1253517471925912e-7 8.448756028504651e-5 0.006329715427485747 0.08703836765622351 0.36787944117144233 0.7287633299194912 0.9394130628134758 0.9961013694701175 1.0 0.9961013694701175 0.9394130628134758; 0.9394130628134758 0.7287633299194912 0.36787944117144233 0.08703836765622351 0.006329715427485747 8.448756028504651e-5 1.1253517471925912e-7 7.404699497933558e-12 1.0848552640429378e-17 1.4524053946493149e-25 6.639677199580735e-36 3.526349876712582e-49 6.740378884131016e-66 2.6157710955041962e-86 6.740378884131016e-66 3.526349876712582e-49 6.639677199580735e-36 1.4524053946493149e-25 1.0848552640429378e-17 7.404699497933558e-12 1.1253517471925912e-7 8.448756028504651e-5 0.006329715427485747 0.08703836765622351 0.36787944117144233 0.7287633299194912 0.9394130628134758 0.9961013694701175 1.0 0.9961013694701175; 0.9961013694701175 0.9394130628134758 0.7287633299194912 0.36787944117144233 0.08703836765622351 0.006329715427485747 8.448756028504651e-5 1.1253517471925912e-7 7.404699497933558e-12 1.0848552640429378e-17 1.4524053946493149e-25 6.639677199580735e-36 3.526349876712582e-49 6.740378884131016e-66 2.6157710955041962e-86 6.740378884131016e-66 3.526349876712582e-49 6.639677199580735e-36 1.4524053946493149e-25 1.0848552640429378e-17 7.404699497933558e-12 1.1253517471925912e-7 8.448756028504651e-5 0.006329715427485747 0.08703836765622351 0.36787944117144233 0.7287633299194912 0.9394130628134758 0.9961013694701175 1.0]
parameters[1] =[30.0, 7.5]
types[1] =[[4, 5, 5, 6, 5, 5], [6, 6, 6, 6, 6], [4, 5, 5, 5, 6, 5], [5, 5, 5, 5, 5, 5], [3, 5, 5, 6, 6, 5], [4, 4, 5, 6, 6, 5], [1, 6, 6, 5, 6, 6]]
hits[1] =[8, 27, 19, 28, 10, 4, 4]
matrix[2] =[1.0 0.9961013694701175 0.9394130628134758 0.7287633299194912 0.36787944117144233 0.08703836765622351 0.006329715427485747 8.448756028504651e-5 1.1253517471925912e-7 7.404699497933558e-12 1.0848552640429378e-17 1.4524053946493149e-25 6.639677199580735e-36 3.526349876712582e-49 6.740378884131016e-66 2.6157710955041962e-86 6.740378884131016e-66 3.526349876712582e-49 6.639677199580735e-36 1.4524053946493149e-25 1.0848552640429378e-17 7.404699497933558e-12 1.1253517471925912e-7 8.448756028504651e-5 0.006329715427485747 0.08703836765622351 0.36787944117144233 0.7287633299194912 0.9394130628134758 0.9961013694701175; 0.9961013694701175 1.0 0.9961013694701175 0.9394130628134758 0.7287633299194912 0.36787944117144233 0.08703836765622351 0.006329715427485747 8.448756028504651e-5 1.1253517471925912e-7 7.404699497933558e-12 1.0848552640429378e-17 1.4524053946493149e-25 6.639677199580735e-36 3.526349876712582e-49 6.740378884131016e-66 2.6157710955041962e-86 6.740378884131016e-66 3.526349876712582e-49 6.639677199580735e-36 1.4524053946493149e-25 1.0848552640429378e-17 7.404699497933558e-12 1.1253517471925912e-7 8.448756028504651e-5 0.006329715427485747 0.08703836765622351 0.36787944117144233 0.7287633299194912 0.9394130628134758; 0.9394130628134758 0.9961013694701175 1.0 0.9961013694701175 0.9394130628134758 0.7287633299194912 0.36787944117144233 0.08703836765622351 0.006329715427485747 8.448756028504651e-5 1.1253517471925912e-7 7.404699497933558e-12 1.0848552640429378e-17 1.4524053946493149e-25 6.639677199580735e-36 3.526349876712582e-49 6.740378884131016e-66 2.6157710955041962e-86 6.740378884131016e-66 3.526349876712582e-49 6.639677199580735e-36 1.4524053946493149e-25 1.0848552640429378e-17 7.404699497933558e-12 1.1253517471925912e-7 8.448756028504651e-5 0.006329715427485747 0.08703836765622351 0.36787944117144233 0.7287633299194912; 0.7287633299194912 0.9394130628134758 0.9961013694701175 1.0 0.9961013694701175 0.9394130628134758 0.7287633299194912 0.36787944117144233 0.08703836765622351 0.006329715427485747 8.448756028504651e-5 1.1253517471925912e-7 7.404699497933558e-12 1.0848552640429378e-17 1.4524053946493149e-25 6.639677199580735e-36 3.526349876712582e-49 6.740378884131016e-66 2.6157710955041962e-86 6.740378884131016e-66 3.526349876712582e-49 6.639677199580735e-36 1.4524053946493149e-25 1.0848552640429378e-17 7.404699497933558e-12 1.1253517471925912e-7 8.448756028504651e-5 0.006329715427485747 0.08703836765622351 0.36787944117144233; 0.36787944117144233 0.7287633299194912 0.9394130628134758 0.9961013694701175 1.0 0.9961013694701175 0.9394130628134758 0.7287633299194912 0.36787944117144233 0.08703836765622351 0.006329715427485747 8.448756028504651e-5 1.1253517471925912e-7 7.404699497933558e-12 1.0848552640429378e-17 1.4524053946493149e-25 6.639677199580735e-36 3.526349876712582e-49 6.740378884131016e-66 2.6157710955041962e-86 6.740378884131016e-66 3.526349876712582e-49 6.639677199580735e-36 1.4524053946493149e-25 1.0848552640429378e-17 7.404699497933558e-12 1.1253517471925912e-7 8.448756028504651e-5 0.006329715427485747 0.08703836765622351; 0.08703836765622351 0.36787944117144233 0.7287633299194912 0.9394130628134758 0.9961013694701175 1.0 0.9961013694701175 0.9394130628134758 0.7287633299194912 0.36787944117144233 0.08703836765622351 0.006329715427485747 8.448756028504651e-5 1.1253517471925912e-7 7.404699497933558e-12 1.0848552640429378e-17 1.4524053946493149e-25 6.639677199580735e-36 3.526349876712582e-49 6.740378884131016e-66 2.6157710955041962e-86 6.740378884131016e-66 3.526349876712582e-49 6.639677199580735e-36 1.4524053946493149e-25 1.0848552640429378e-17 7.404699497933558e-12 1.1253517471925912e-7 8.448756028504651e-5 0.006329715427485747; 0.006329715427485747 0.08703836765622351 0.36787944117144233 0.7287633299194912 0.9394130628134758 0.9961013694701175 1.0 0.9961013694701175 0.9394130628134758 0.7287633299194912 0.36787944117144233 0.08703836765622351 0.006329715427485747 8.448756028504651e-5 1.1253517471925912e-7 7.404699497933558e-12 1.0848552640429378e-17 1.4524053946493149e-25 6.639677199580735e-36 3.526349876712582e-49 6.740378884131016e-66 2.6157710955041962e-86 6.740378884131016e-66 3.526349876712582e-49 6.639677199580735e-36 1.4524053946493149e-25 1.0848552640429378e-17 7.404699497933558e-12 1.1253517471925912e-7 8.448756028504651e-5; 8.448756028504651e-5 0.006329715427485747 0.08703836765622351 0.36787944117144233 0.7287633299194912 0.9394130628134758 0.9961013694701175 1.0 0.9961013694701175 0.9394130628134758 0.7287633299194912 0.36787944117144233 0.08703836765622351 0.006329715427485747 8.448756028504651e-5 1.1253517471925912e-7 7.404699497933558e-12 1.0848552640429378e-17 1.4524053946493149e-25 6.639677199580735e-36 3.526349876712582e-49 6.740378884131016e-66 2.6157710955041962e-86 6.740378884131016e-66 3.526349876712582e-49 6.639677199580735e-36 1.4524053946493149e-25 1.0848552640429378e-17 7.404699497933558e-12 1.1253517471925912e-7; 1.1253517471925912e-7 8.448756028504651e-5 0.006329715427485747 0.08703836765622351 0.36787944117144233 0.7287633299194912 0.9394130628134758 0.9961013694701175 1.0 0.9961013694701175 0.9394130628134758 0.7287633299194912 0.36787944117144233 0.08703836765622351 0.006329715427485747 8.448756028504651e-5 1.1253517471925912e-7 7.404699497933558e-12 1.0848552640429378e-17 1.4524053946493149e-25 6.639677199580735e-36 3.526349876712582e-49 6.740378884131016e-66 2.6157710955041962e-86 6.740378884131016e-66 3.526349876712582e-49 6.639677199580735e-36 1.4524053946493149e-25 1.0848552640429378e-17 7.404699497933558e-12; 7.404699497933558e-12 1.1253517471925912e-7 8.448756028504651e-5 0.006329715427485747 0.08703836765622351 0.36787944117144233 0.7287633299194912 0.9394130628134758 0.9961013694701175 1.0 0.9961013694701175 0.9394130628134758 0.7287633299194912 0.36787944117144233 0.08703836765622351 0.006329715427485747 8.448756028504651e-5 1.1253517471925912e-7 7.404699497933558e-12 1.0848552640429378e-17 1.4524053946493149e-25 6.639677199580735e-36 3.526349876712582e-49 6.740378884131016e-66 2.6157710955041962e-86 6.740378884131016e-66 3.526349876712582e-49 6.639677199580735e-36 1.4524053946493149e-25 1.0848552640429378e-17; 1.0848552640429378e-17 7.404699497933558e-12 1.1253517471925912e-7 8.448756028504651e-5 0.006329715427485747 0.08703836765622351 0.36787944117144233 0.7287633299194912 0.9394130628134758 0.9961013694701175 1.0 0.9961013694701175 0.9394130628134758 0.7287633299194912 0.36787944117144233 0.08703836765622351 0.006329715427485747 8.448756028504651e-5 1.1253517471925912e-7 7.404699497933558e-12 1.0848552640429378e-17 1.4524053946493149e-25 6.639677199580735e-36 3.526349876712582e-49 6.740378884131016e-66 2.6157710955041962e-86 6.740378884131016e-66 3.526349876712582e-49 6.639677199580735e-36 1.4524053946493149e-25; 1.4524053946493149e-25 1.0848552640429378e-17 7.404699497933558e-12 1.1253517471925912e-7 8.448756028504651e-5 0.006329715427485747 0.08703836765622351 0.36787944117144233 0.7287633299194912 0.9394130628134758 0.9961013694701175 1.0 0.9961013694701175 0.9394130628134758 0.7287633299194912 0.36787944117144233 0.08703836765622351 0.006329715427485747 8.448756028504651e-5 1.1253517471925912e-7 7.404699497933558e-12 1.0848552640429378e-17 1.4524053946493149e-25 6.639677199580735e-36 3.526349876712582e-49 6.740378884131016e-66 2.6157710955041962e-86 6.740378884131016e-66 3.526349876712582e-49 6.639677199580735e-36; 6.639677199580735e-36 1.4524053946493149e-25 1.0848552640429378e-17 7.404699497933558e-12 1.1253517471925912e-7 8.448756028504651e-5 0.006329715427485747 0.08703836765622351 0.36787944117144233 0.7287633299194912 0.9394130628134758 0.9961013694701175 1.0 0.9961013694701175 0.9394130628134758 0.7287633299194912 0.36787944117144233 0.08703836765622351 0.006329715427485747 8.448756028504651e-5 1.1253517471925912e-7 7.404699497933558e-12 1.0848552640429378e-17 1.4524053946493149e-25 6.639677199580735e-36 3.526349876712582e-49 6.740378884131016e-66 2.6157710955041962e-86 6.740378884131016e-66 3.526349876712582e-49; 3.526349876712582e-49 6.639677199580735e-36 1.4524053946493149e-25 1.0848552640429378e-17 7.404699497933558e-12 1.1253517471925912e-7 8.448756028504651e-5 0.006329715427485747 0.08703836765622351 0.36787944117144233 0.7287633299194912 0.9394130628134758 0.9961013694701175 1.0 0.9961013694701175 0.9394130628134758 0.7287633299194912 0.36787944117144233 0.08703836765622351 0.006329715427485747 8.448756028504651e-5 1.1253517471925912e-7 7.404699497933558e-12 1.0848552640429378e-17 1.4524053946493149e-25 6.639677199580735e-36 3.526349876712582e-49 6.740378884131016e-66 2.6157710955041962e-86 6.740378884131016e-66; 6.740378884131016e-66 3.526349876712582e-49 6.639677199580735e-36 1.4524053946493149e-25 1.0848552640429378e-17 7.404699497933558e-12 1.1253517471925912e-7 8.448756028504651e-5 0.006329715427485747 0.08703836765622351 0.36787944117144233 0.7287633299194912 0.9394130628134758 0.9961013694701175 1.0 0.9961013694701175 0.9394130628134758 0.7287633299194912 0.36787944117144233 0.08703836765622351 0.006329715427485747 8.448756028504651e-5 1.1253517471925912e-7 7.404699497933558e-12 1.0848552640429378e-17 1.4524053946493149e-25 6.639677199580735e-36 3.526349876712582e-49 6.740378884131016e-66 2.6157710955041962e-86; 2.6157710955041962e-86 6.740378884131016e-66 3.526349876712582e-49 6.639677199580735e-36 1.4524053946493149e-25 1.0848552640429378e-17 7.404699497933558e-12 1.1253517471925912e-7 8.448756028504651e-5 0.006329715427485747 0.08703836765622351 0.36787944117144233 0.7287633299194912 0.9394130628134758 0.9961013694701175 1.0 0.9961013694701175 0.9394130628134758 0.7287633299194912 0.36787944117144233 0.08703836765622351 0.006329715427485747 8.448756028504651e-5 1.1253517471925912e-7 7.404699497933558e-12 1.0848552640429378e-17 1.4524053946493149e-25 6.639677199580735e-36 3.526349876712582e-49 6.740378884131016e-66; 6.740378884131016e-66 2.6157710955041962e-86 6.740378884131016e-66 3.526349876712582e-49 6.639677199580735e-36 1.4524053946493149e-25 1.0848552640429378e-17 7.404699497933558e-12 1.1253517471925912e-7 8.448756028504651e-5 0.006329715427485747 0.08703836765622351 0.36787944117144233 0.7287633299194912 0.9394130628134758 0.9961013694701175 1.0 0.9961013694701175 0.9394130628134758 0.7287633299194912 0.36787944117144233 0.08703836765622351 0.006329715427485747 8.448756028504651e-5 1.1253517471925912e-7 7.404699497933558e-12 1.0848552640429378e-17 1.4524053946493149e-25 6.639677199580735e-36 3.526349876712582e-49; 3.526349876712582e-49 6.740378884131016e-66 2.6157710955041962e-86 6.740378884131016e-66 3.526349876712582e-49 6.639677199580735e-36 1.4524053946493149e-25 1.0848552640429378e-17 7.404699497933558e-12 1.1253517471925912e-7 8.448756028504651e-5 0.006329715427485747 0.08703836765622351 0.36787944117144233 0.7287633299194912 0.9394130628134758 0.9961013694701175 1.0 0.9961013694701175 0.9394130628134758 0.7287633299194912 0.36787944117144233 0.08703836765622351 0.006329715427485747 8.448756028504651e-5 1.1253517471925912e-7 7.404699497933558e-12 1.0848552640429378e-17 1.4524053946493149e-25 6.639677199580735e-36; 6.639677199580735e-36 3.526349876712582e-49 6.740378884131016e-66 2.6157710955041962e-86 6.740378884131016e-66 3.526349876712582e-49 6.639677199580735e-36 1.4524053946493149e-25 1.0848552640429378e-17 7.404699497933558e-12 1.1253517471925912e-7 8.448756028504651e-5 0.006329715427485747 0.08703836765622351 0.36787944117144233 0.7287633299194912 0.9394130628134758 0.9961013694701175 1.0 0.9961013694701175 0.9394130628134758 0.7287633299194912 0.36787944117144233 0.08703836765622351 0.006329715427485747 8.448756028504651e-5 1.1253517471925912e-7 7.404699497933558e-12 1.0848552640429378e-17 1.4524053946493149e-25; 1.4524053946493149e-25 6.639677199580735e-36 3.526349876712582e-49 6.740378884131016e-66 2.6157710955041962e-86 6.740378884131016e-66 3.526349876712582e-49 6.639677199580735e-36 1.4524053946493149e-25 1.0848552640429378e-17 7.404699497933558e-12 1.1253517471925912e-7 8.448756028504651e-5 0.006329715427485747 0.08703836765622351 0.36787944117144233 0.7287633299194912 0.9394130628134758 0.9961013694701175 1.0 0.9961013694701175 0.9394130628134758 0.7287633299194912 0.36787944117144233 0.08703836765622351 0.006329715427485747 8.448756028504651e-5 1.1253517471925912e-7 7.404699497933558e-12 1.0848552640429378e-17; 1.0848552640429378e-17 1.4524053946493149e-25 6.639677199580735e-36 3.526349876712582e-49 6.740378884131016e-66 2.6157710955041962e-86 6.740378884131016e-66 3.526349876712582e-49 6.639677199580735e-36 1.4524053946493149e-25 1.0848552640429378e-17 7.404699497933558e-12 1.1253517471925912e-7 8.448756028504651e-5 0.006329715427485747 0.08703836765622351 0.36787944117144233 0.7287633299194912 0.9394130628134758 0.9961013694701175 1.0 0.9961013694701175 0.9394130628134758 0.7287633299194912 0.36787944117144233 0.08703836765622351 0.006329715427485747 8.448756028504651e-5 1.1253517471925912e-7 7.404699497933558e-12; 7.404699497933558e-12 1.0848552640429378e-17 1.4524053946493149e-25 6.639677199580735e-36 3.526349876712582e-49 6.740378884131016e-66 2.6157710955041962e-86 6.740378884131016e-66 3.526349876712582e-49 6.639677199580735e-36 1.4524053946493149e-25 1.0848552640429378e-17 7.404699497933558e-12 1.1253517471925912e-7 8.448756028504651e-5 0.006329715427485747 0.08703836765622351 0.36787944117144233 0.7287633299194912 0.9394130628134758 0.9961013694701175 1.0 0.9961013694701175 0.9394130628134758 0.7287633299194912 0.36787944117144233 0.08703836765622351 0.006329715427485747 8.448756028504651e-5 1.1253517471925912e-7; 1.1253517471925912e-7 7.404699497933558e-12 1.0848552640429378e-17 1.4524053946493149e-25 6.639677199580735e-36 3.526349876712582e-49 6.740378884131016e-66 2.6157710955041962e-86 6.740378884131016e-66 3.526349876712582e-49 6.639677199580735e-36 1.4524053946493149e-25 1.0848552640429378e-17 7.404699497933558e-12 1.1253517471925912e-7 8.448756028504651e-5 0.006329715427485747 0.08703836765622351 0.36787944117144233 0.7287633299194912 0.9394130628134758 0.9961013694701175 1.0 0.9961013694701175 0.9394130628134758 0.7287633299194912 0.36787944117144233 0.08703836765622351 0.006329715427485747 8.448756028504651e-5; 8.448756028504651e-5 1.1253517471925912e-7 7.404699497933558e-12 1.0848552640429378e-17 1.4524053946493149e-25 6.639677199580735e-36 3.526349876712582e-49 6.740378884131016e-66 2.6157710955041962e-86 6.740378884131016e-66 3.526349876712582e-49 6.639677199580735e-36 1.4524053946493149e-25 1.0848552640429378e-17 7.404699497933558e-12 1.1253517471925912e-7 8.448756028504651e-5 0.006329715427485747 0.08703836765622351 0.36787944117144233 0.7287633299194912 0.9394130628134758 0.9961013694701175 1.0 0.9961013694701175 0.9394130628134758 0.7287633299194912 0.36787944117144233 0.08703836765622351 0.006329715427485747; 0.006329715427485747 8.448756028504651e-5 1.1253517471925912e-7 7.404699497933558e-12 1.0848552640429378e-17 1.4524053946493149e-25 6.639677199580735e-36 3.526349876712582e-49 6.740378884131016e-66 2.6157710955041962e-86 6.740378884131016e-66 3.526349876712582e-49 6.639677199580735e-36 1.4524053946493149e-25 1.0848552640429378e-17 7.404699497933558e-12 1.1253517471925912e-7 8.448756028504651e-5 0.006329715427485747 0.08703836765622351 0.36787944117144233 0.7287633299194912 0.9394130628134758 0.9961013694701175 1.0 0.9961013694701175 0.9394130628134758 0.7287633299194912 0.36787944117144233 0.08703836765622351; 0.08703836765622351 0.006329715427485747 8.448756028504651e-5 1.1253517471925912e-7 7.404699497933558e-12 1.0848552640429378e-17 1.4524053946493149e-25 6.639677199580735e-36 3.526349876712582e-49 6.740378884131016e-66 2.6157710955041962e-86 6.740378884131016e-66 3.526349876712582e-49 6.639677199580735e-36 1.4524053946493149e-25 1.0848552640429378e-17 7.404699497933558e-12 1.1253517471925912e-7 8.448756028504651e-5 0.006329715427485747 0.08703836765622351 0.36787944117144233 0.7287633299194912 0.9394130628134758 0.9961013694701175 1.0 0.9961013694701175 0.9394130628134758 0.7287633299194912 0.36787944117144233; 0.36787944117144233 0.08703836765622351 0.006329715427485747 8.448756028504651e-5 1.1253517471925912e-7 7.404699497933558e-12 1.0848552640429378e-17 1.4524053946493149e-25 6.639677199580735e-36 3.526349876712582e-49 6.740378884131016e-66 2.6157710955041962e-86 6.740378884131016e-66 3.526349876712582e-49 6.639677199580735e-36 1.4524053946493149e-25 1.0848552640429378e-17 7.404699497933558e-12 1.1253517471925912e-7 8.448756028504651e-5 0.006329715427485747 0.08703836765622351 0.36787944117144233 0.7287633299194912 0.9394130628134758 0.9961013694701175 1.0 0.9961013694701175 0.9394130628134758 0.7287633299194912; 0.7287633299194912 0.36787944117144233 0.08703836765622351 0.006329715427485747 8.448756028504651e-5 1.1253517471925912e-7 7.404699497933558e-12 1.0848552640429378e-17 1.4524053946493149e-25 6.639677199580735e-36 3.526349876712582e-49 6.740378884131016e-66 2.6157710955041962e-86 6.740378884131016e-66 3.526349876712582e-49 6.639677199580735e-36 1.4524053946493149e-25 1.0848552640429378e-17 7.404699497933558e-12 1.1253517471925912e-7 8.448756028504651e-5 0.006329715427485747 0.08703836765622351 0.36787944117144233 0.7287633299194912 0.9394130628134758 0.9961013694701175 1.0 0.9961013694701175 0.9394130628134758; 0.9394130628134758 0.7287633299194912 0.36787944117144233 0.08703836765622351 0.006329715427485747 8.448756028504651e-5 1.1253517471925912e-7 7.404699497933558e-12 1.0848552640429378e-17 1.4524053946493149e-25 6.639677199580735e-36 3.526349876712582e-49 6.740378884131016e-66 2.6157710955041962e-86 6.740378884131016e-66 3.526349876712582e-49 6.639677199580735e-36 1.4524053946493149e-25 1.0848552640429378e-17 7.404699497933558e-12 1.1253517471925912e-7 8.448756028504651e-5 0.006329715427485747 0.08703836765622351 0.36787944117144233 0.7287633299194912 0.9394130628134758 0.9961013694701175 1.0 0.9961013694701175; 0.9961013694701175 0.9394130628134758 0.7287633299194912 0.36787944117144233 0.08703836765622351 0.006329715427485747 8.448756028504651e-5 1.1253517471925912e-7 7.404699497933558e-12 1.0848552640429378e-17 1.4524053946493149e-25 6.639677199580735e-36 3.526349876712582e-49 6.740378884131016e-66 2.6157710955041962e-86 6.740378884131016e-66 3.526349876712582e-49 6.639677199580735e-36 1.4524053946493149e-25 1.0848552640429378e-17 7.404699497933558e-12 1.1253517471925912e-7 8.448756028504651e-5 0.006329715427485747 0.08703836765622351 0.36787944117144233 0.7287633299194912 0.9394130628134758 0.9961013694701175 1.0]
parameters[2] =[30.0, 7.5]
types[2] =[[4, 5, 5, 6, 5, 5], [6, 6, 6, 6, 6], [4, 5, 5, 5, 6, 5], [5, 5, 5, 5, 5, 5], [1, 6, 6, 5, 6, 6], [4, 4, 5, 6, 6, 5], [3, 5, 5, 6, 6, 5]]
hits[2] =[11, 30, 18, 22, 8, 7, 4]
```
