## Supplementary material for "On the structure of multiple stable equilibria in competitive ecological systems": Mathematica+Julia code and examples: sorted-200.m

```
distribution[30] =
 {{57, {6, 6, 6, 6, 6}}, {50, {5, 5, 5, 5, 5, 5}}, {37, {4, 5, 5, 5, 6, 5}}, 
 {19, {4, 5, 5, 6, 5, 5}}, {14, {3, 5, 5, 6, 6, 5}}, 
 {12, {1, 6, 6, 5, 6, 6}}, {11, {4, 4, 5, 6, 6, 5}}}
```
