## Supplementary material for "On the structure of multiple stable equilibria in competitive ecological systems": Mathematica+Julia code and examples: tree.m

```
(* builds a tree based on statistical model to find all possible solutions *)
(* note, these must be checked for stability *)

(* assumes have a model (e.g. m4), use set of possibilities at length k *)
(* also assumes have loaded stability code (e.g. basins.m) *)

(* key functions: *)

(* canonicalstable[possible[rules, k]]: finds all possible periodic
 sequences given local k-gap possible subspace from rules, puts in
 canonical order and only returns stable solutions. *)

(* solutions[rules, k] number of solutions for rules, given k *)

(* ------------------------------------------------------ *)

(* we will use rules from m4.m *)

<< m4.m

(* rules is just the list of allowed 4-gap sequences *)

r4 = Map[First, m4[[4]]]

(* ------------------------------------------------------- *)

(* expands a set of partials by one, dropping anything over the limit *)

expand1[p_, rules_] :=
 Map[Append[p, Last[#]]&, 
        Select[rules, Drop[#, -1] == Take[p, 1-Length[rules[[1]]]] &]]

expands[ps_, rules_, limit_] :=
 Select[Apply[Join, Map[expand1[#, rules] &, ps]], Apply[Plus, #]<= limit &]

expandsg[ps_, rules_] :=
 Apply[Join, Map[expand1[#, rules] &, ps]]

(* collects everything of correct size satisfying periodicity *)

periodicq[s_,rules_] :=
 With[{double = Join[s, s], l =Length[rules[[1]]]},
  Apply[And,Table[MemberQ[rules, Take[double,{i, i + l-1}]],
                  {i,Length[s]-l+2,Length[s]}]]]

size1[p_] := Apply[Plus, p]

possible[rules_, length_] :=
 Reap[part = rules;
      While[part != {},
        Map[Sow, 
            Select[part, And[size1[#]== length, periodicq[#, rules]] &]];
        part = expands[part, rules, length]]][[2,1]]

possibleg[rules_, length_] :=
  Block[{},
	part = rules;
	i = Length[part[[1]]];
	list = {{i, part}};
      While[i <= length,
	    i = i +1;
	    part = expandsg[part, rules];
	    list = Append[list, {i, part}];
	    Print [i,": ",Length[ part]]];
      list]

(* finds canonical ones that are stable *)

canonicalstable[list_] :=
 With[{mx1 = matrix[Apply[Plus, list[[1]]], Apply[Plus, list[[1]]]/4.0]},
     Select[Union[Map[canonical[#, 1] &, list]],
        stable[mx1, configuration[#]]&]]


(* how many times to count a sequence *)

rotations[s_] :=
 Table[RotateRight[s, i],{i, 1,Length[s]}]

distinct[s_] :=
 Length[Union[Join[rotations[s], rotations[Reverse[s]]]]]*
  Apply[Plus, s]/(Length[s])

(* counting distinct solutions at size l *)

solutions[rules_, l_] :=
 With[{a = canonicalstable[possible[rules, l]]},
  {Apply[Plus, Map[distinct, a]], a}]

(* output to file *)

producesolutions[rules_, max_]  :=
 Block[{r},
  For[j = 20, j <= max, j += 1,
    r = solutions[rules, j];
    Print[j, ":", r[[1]]];
    {j, r} >>> "model-solutions.m"]]
```
